# Accessible and reproducible deployment reveals the practical boundaries of single-cell foundation models

**DOI:** 10.64898/2026.01.06.698060

**Authors:** Siyu Hou, Penghui Yang, Wenjing Ma, Jinxi Xiang, Jade Xiaoqing Wang, Hui Wan, Ying Ma, Xiang Zhou

## Abstract

Single-cell foundation models (scFMs) have been widely promoted as a unifying paradigm for transcriptomic analysis, yet whether large-scale pretraining translates into reproducible biological advantages remains unclear. Their adoption is further hindered by heterogeneous implementations, preprocessing requirements, and computational environments. Here we develop a unified, automated, and reproducible framework for standardized deployment and controlled evaluation of scFMs across datasets, computational environments, training regimes, and downstream analyses, substantially lowering the technical barriers to their use. Leveraging this framework, we systematically investigate thirteen scFMs alongside established methods across nearly one hundred datasets spanning diverse biological contexts. Our analyses reveal clear practical boundaries to scFM utility. First, increased model scale, architectural complexity, pretraining corpus size, or input encoding does not consistently translate into superior downstream performance. Instead, measurable properties of embedding geometry provide a model-agnostic, representation-level explanation for differences in zero-shot performance across diverse model families. Second, the benefits of pretrained representations depend strongly on the biological and supervision regime: scFMs provide their clearest advantages under extremely limited supervision, particularly for rare-cell annotation and open-set detection of source-absent cell states, whereas established methods remain competitive or preferable in most other settings. Task-matched analyses further show that scFM representations transfer inconsistently to spatial-domain recovery, while their gene embeddings capture broad functional relatedness without reliably recovering context-specific regulatory relationships. Together, these results establish that scFM utility is neither universal nor determined simply by model scale alone, but varies with learned representation geometry, biological context, and supervision. By combining reproducible deployment with large-scale empirical and mechanistic investigation, our framework provides a principled foundation for determining when foundation-model pretraining offers genuine practical value and when simpler approaches remain sufficient.

## 1 Introduction

Inspired by recent advances in large language models, single-cell foundation models (scFMs; also referred to as scLLMs) have emerged as a promising paradigm for modeling gene expression at scale [1–3]. By leveraging transformer-based architectures and self-supervised learning on massive transcriptomic corpora, scFMs aim to learn transferable representations applicable to diverse downstream tasks, including cell type annotation, batch effect correction, perturbation prediction and multi-omics integration [1, 4]. These proposed capabilities position scFMs as a promising and unifying paradigm for diverse analytical tasks in single-cell biology.

Despite growing enthusiasm, scFMs remain difficult to deploy in practice. These models are large, complex systems that often require substantial computational expertise, placing them beyond the reach of many non-specialists. Existing scFMs differ widely in architecture, training objectives, preprocessing requirements, and software implementations, and are frequently applied under heterogeneous preprocessing pipelines, training regimes, and execution environments. Consequently, choices in data processing, embedding extraction, and supervised adaptation can materially affect downstream performance. At present, standardized, accessible deployment remains lacking, with little consensus on best practices for using scFMs across biological applications. These challenges are compounded by incomplete documentation, tightly coupled software dependencies, and incompatible computing environments, making installation and execution time-consuming and error-prone. Together, these technical barriers hinder reproducibility, accessibility, and broad adoption of scFMs, particularly for experimental biologists and smaller laboratories.

Even if these practical barriers were eliminated, a central scientific question would remain unresolved: under what conditions do scFMs provide meaningful advantages over established single-cell analysis approaches? The appeal of scFMs rests partly on the premise that increasing model scale and expanding self-supervised pretraining will produce representations that transfer more effectively across biological contexts. However, systematic evidence supporting this assumption remains limited. Early scFMs were primarily developed from a computational perspective, with downstream evaluations that were limited in scope and often performed on small or simplified datasets, leaving generalization to diverse, realistic biological scenarios unclear. Existing comparative studies likewise typically examined only a few well-known methods such as scGPT[4], Geneformer[5] and scFoundation[6], across a restricted set of tasks and datasets [7–10]. Differences in configuration, embedding extraction and training procedure further complicate comparisons among studies. Consequently, it remains unclear how well scFMs generalize across biological contexts, when they consistently outperform simpler established methods, and where foundation-model pretraining provides genuine practical value.

Beyond establishing whether scFMs work, an even more fundamental question is why they work. Current studies overwhelmingly emphasize performance rankings while providing limited insight into the representations learned during large-scale pretraining. As a result, we still lack a mechanistic understanding of what properties of these representations enable improved performance and generalization, which downstream tasks truly benefit from foundation-model pretraining, and when simpler classical methods should instead be expected to perform equally well or even better. Recent reports, for example, suggest that scFMs fail to outperform simple linear baselines for gene perturbation effect prediction [11], underscoring that increased model scale alone does not guarantee superior biological performance. Addressing these questions requires standardized, reproducible evaluation frameworks that not only compare predictive accuracy across diverse biological settings, but also enable systematic investigation of the mechanisms underlying scFM representations and their downstream utility.

To address these practical, empirical, and conceptual challenges, we developed a unified, standardized, and extensible framework that harmonizes scFM implementations and enables reproducible deployment and controlled evaluation across datasets, training regimes, and downstream tasks. This framework substantially lowers the technical barrier to using scFMs while providing a controlled foundation for rigorous and reproducible analyses across computational platforms and diverse biological settings. Leveraging this framework, we conducted a large-scale and systematic investigation of thirteen scFMs alongside established baselines across 94 datasets spanning zero-shot, frozen-embedding transfer, and supervised adaptation. Crucially, we move beyond ranking models to ask when, and why, scFMs provide meaningful advantages. We show that measurable properties of embedding geometry provide a representation-level explanation for performance differences across discrete cell-type organization and continuous trajectory recovery, whereas model architecture, scale, pretraining corpus size, and input encoding do not reliably predict representation quality. Across the evaluated downstream settings, established and task-native methods generally match or outperform scFMs, while selected pretrained representations provide their clearest advantages under severe label limitation, particularly for rare-cell annotation, and in open-set detection of source-absent cell types. Frozen scFM representations also transfer inconsistently to spatial-domain recovery, while their gene embeddings capture broad functional relatedness without reliably recovering context-specific regulatory relationships. Together, this work establishes an accessible and reproducible foundation for the informed deployment, interpretation, and future development of single-cell foundation models, providing practical guidance for their application and new insights into the principles governing their success.

## 2 Results

### 2.1 A unified, containerized framework for reproducible deployment and evaluation of scFMs

To lower the technical barrier and make single-cell foundation models (scFMs) easily accessible, we developed a unified, workflow-based infrastructure that standardizes model execution, embedding extraction and downstream evaluation across heterogeneous implementations. Implemented in Nextflow [12], the framework supports single-command execution of end-to-end inference and downstream analysis, with model and task modules that can be extended independently (Fig. 1).

**Fig. 1.**
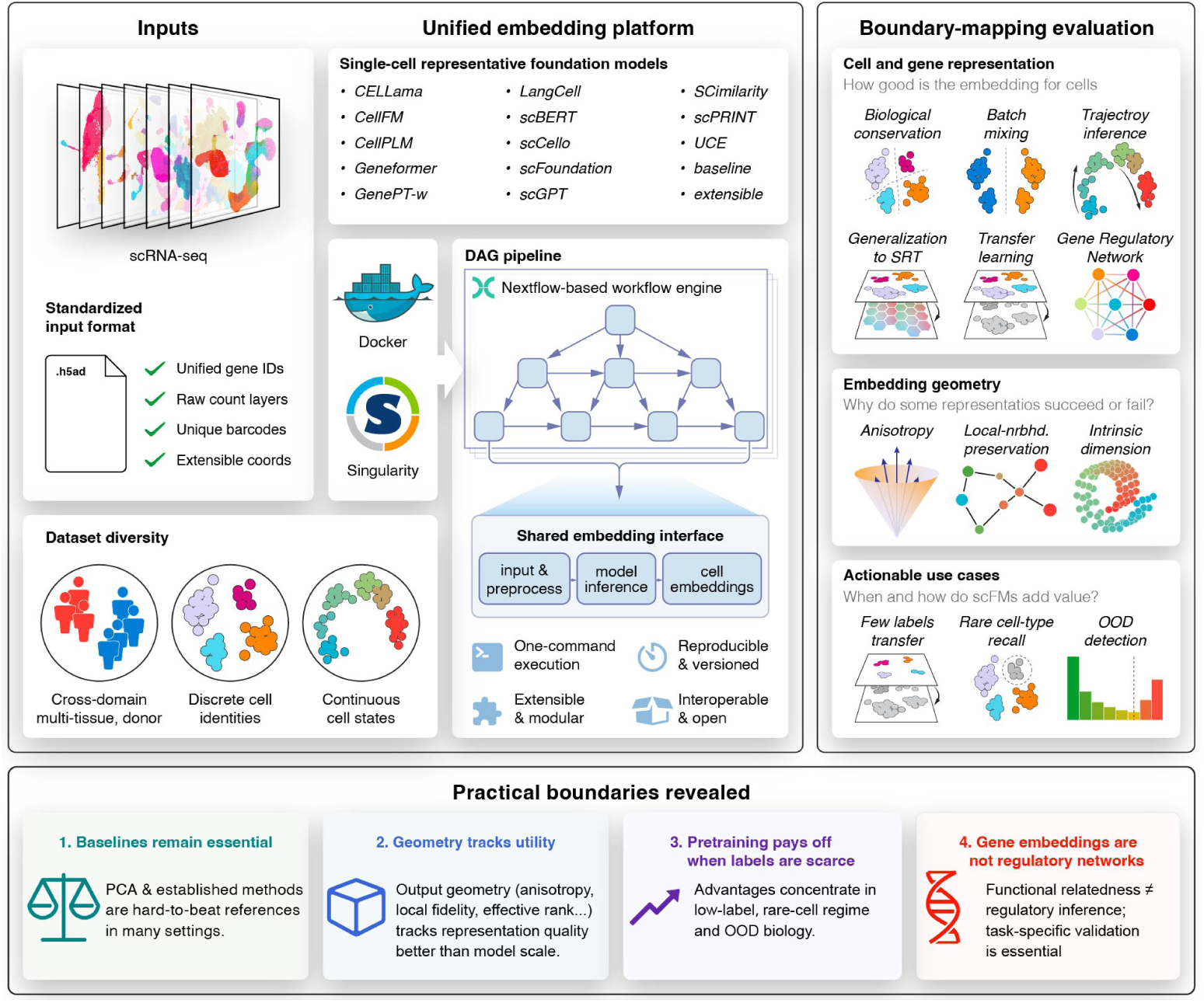
Overview of the deployment and evaluation framework and study design. **Left**, standardized transcriptomic inputs and evaluation datasets spanning tissues, donors and studies, with discrete cell-identity and continuous cell-state annotations used for evaluation and, where specified, frozen-embedding transfer or supervised adaptation; spatial coordinates are retained for SRT analyses. **Center**, a containerized Nextflow platform for 13 core pretrained scFM representations, with a shared AnnData embedding interface and task-dependent reference methods. **Right**, downstream evaluation of biological conservation, batch mixing, trajectory inference, SRT deployment, transfer learning and gene-level regulatory readouts. **Bottom**, the principal practical boundaries and associations examined in the study, including the competitiveness of reference methods, associations between embedding geometry and downstream performance, benefits in selected label-scarce settings and the limits of gene-embedding similarity for regulatory inference.

A central design goal was to provide a model-agnostic embedding interface. Each model receives a standardized AnnData [13] object with harmonized gene identifiers and metadata and returns embeddings with aligned cell metadata in the same format. This common interface decouples downstream tasks from model-specific preprocessing and implementation details, allowing the same analysis and evaluation code to be applied uniformly across models. When official implementations required specific preprocessing steps (e.g., gene identifier conventions, normalization or tokenization), these were encapsulated within the corresponding wrapper to match the authors’ intended usage, rather than being imposed globally.

To support reproducibility and isolate software dependencies, each method is executed in a dedicated Docker or Singularity container with fixed library versions and runtime requirements. The workflow records software versions and key parameters, allowing analyses to be re-executed across computing platforms. We used released code and pretrained weights wherever available, applying only documented fixes required for execution without altering the underlying model methodology.

The framework not only enables the deployment and investigation of scFMs at scale but also improves their accessibility. Users can now run individual models with a single command, and can extend the model zoo by implementing a lightweight wrapper that conforms to the standardized input–output contract. Collectively, this infrastructure lowers the technical barrier for deploying scFMs, enables reproducible comparisons under matched conditions, and provides a practical foundation for community-wide evaluation as new models and datasets emerge.

### 2.2 A controlled tissue atlas reveals selective zero-shot advantages of scFMs

We first asked whether frozen single-cell foundation-model embeddings provided a consistent advantage over simple expression-based representations. To address this question, we constructed a controlled panel of 26 tissue datasets from Tabula Sapiens v2 (TSv2) [14], comprising 548,977 cells and 14–32 annotated cell types per tissue. These data were generated and processed within a harmonized atlas, and our audit of the available documentation found that TSv2 was not documented in the pretraining corpora of any evaluated scFMs (Supplementary Materials). We evaluated 20 representations: 13 frozen pretrained scFMs (CELLama[15], CellFM[16], CellPLM[17], Geneformer[5], GenePT-w[18], LangCell[19], scBERT[20], scCello[21], scFoundation[6], scGPT[4], SCimilarity[22], scPRINT[23] and UCE[24]), a large-scale pretrained scVI [25, 26] as a non-foundation counterpart to scFMs, a classic expression-based PCA reference, three established target-data integration methods (Harmony[27], Seurat CCA and Seurat RPCA [28]), and the standard target-data workflows of scVI and scGPT, denoted scVI (de novo) and scGPT (integrated), respectively. Unlike the frozen pretrained representations, the last six methods derive representations directly from the target dataset and therefore served as target-data reference comparators.

Because batch effect correction is not a native objective of most frozen scFM workflows, but rather an expected byproduct of learning biologically relevant embeddings, we evaluated preservation of biological structure and removal of batch variation as two separate properties. The primary zero-shot comparison thus focused on conservation of biological variance from cell identity, assessed using nine biological-conservation metrics grouped into three categories: cluster concordance, continuous cell-type separation, and local cell-type-neighbourhood structure; batch mixing was analysed separately in the 19 multi-batch tissues (Methods).

Across the full 20-representation panel, the overall performance hierarchy among the methods was broadly consistent across the 26 tissues (Kendall’s *W* = 0.723; Friedman *χ*^2^ = 357.212, d.f. = 19, *P* = 3.30×10^−64^; Fig. 2a). SCimilarity achieved the best mean weighted biological-conservation family rank (3.98), whereas scFoundation (6.15) and PCA (6.60) were closely matched and ranked second and third, respectively (Fig. 2a). SCimilarity was the only frozen scFM with a significantly better aggregate family rank than PCA after Holm correction (paired tissue-level tests; adjusted *P* = 0.0145). The modest advantage of scFoundation over PCA was not statistically significant (adjusted *P* = 0.816), and neither CellPLM nor UCE differed significantly from PCA. Several representations, including Geneformer, scPRINT, CellFM, CELLama, GenePT-w and scBERT, ranked substantially below PCA. Metric-level paired comparisons further showed that a better mean rank did not imply statistically significant improvement across individual biological metrics (Supplementary Results).

**Fig. 2.**
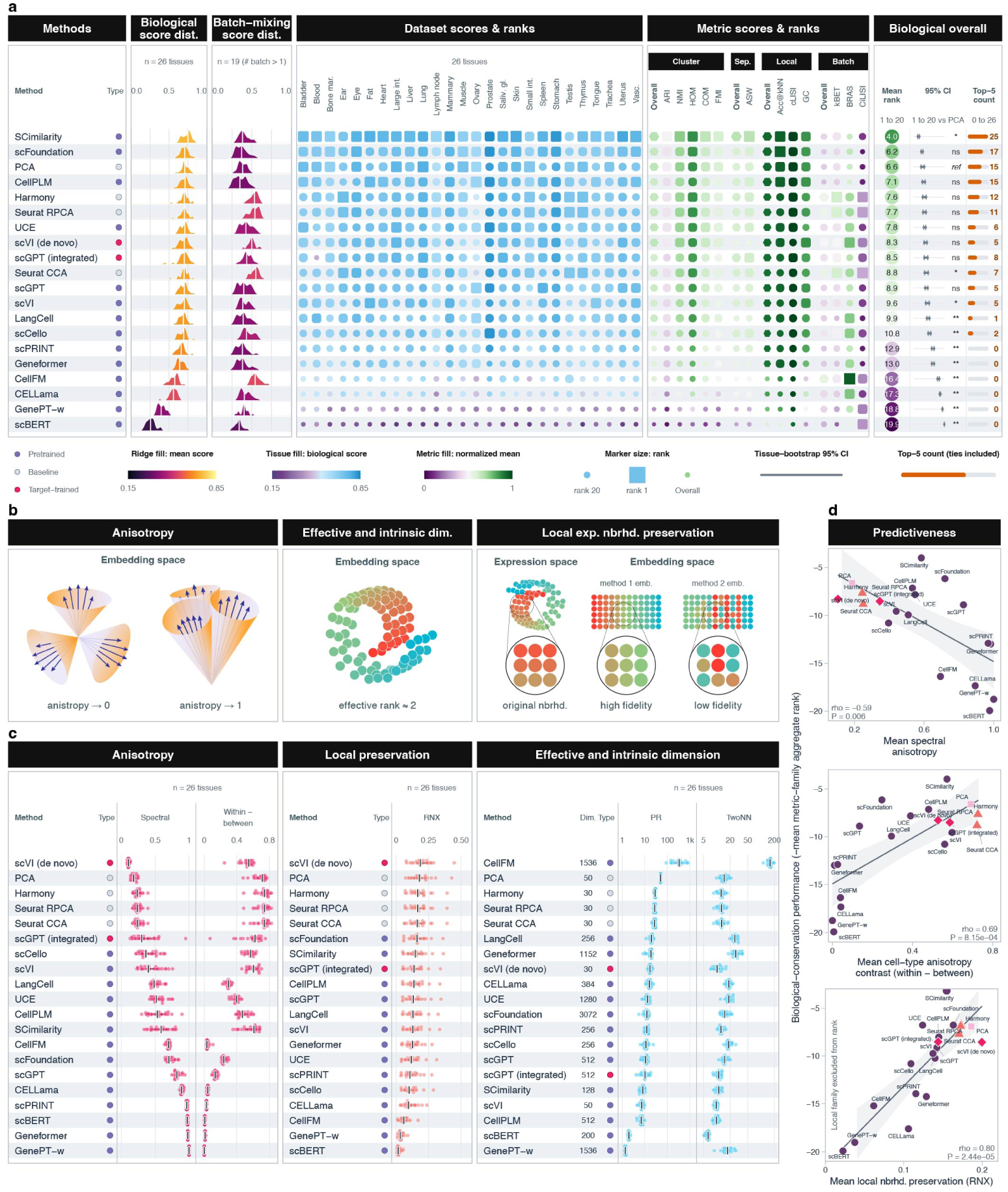
Frozen representation benchmarking and geometric diagnostics. **a,** Biological-conservation ranks for 20 representations across 26 TSv2 tissues. Biological scores combine cluster concordance (32.5%), continuous cell-type separation (10%), and local cell-type-neighbourhood structure (32.5%), with weights renormalized over the biological components. Batch mixing is summarized separately for 19 eligible multi-batch tissues. Intervals are 95% percentile intervals from 10,000 complete-tissue bootstrap replicates. Symbols indicate two-sided paired Wilcoxon signed-rank tests versus PCA across 26 tissues, with Holm correction over 19 comparisons. **b,** Schematics illustrating global anisotropy, effective and intrinsic dimensionality, and local expression-neighbourhood preservation. **c,** Tissue-level distributions of spectral anisotropy, cell-type anisotropy contrast, local expression-neighbourhood preservation (*R_NX_*; *k* = 15, 30, 50), participation ratio (PR) and TwoNN intrinsic dimension. Points denote tissues; ticks and bars indicate medians and interquartile ranges. **d,** Mean geometry properties versus biological-conservation performance (negative mean metric-family aggregate rank) across 20 representations. The *R*_7_*NX* analysis excludes Acc@kNN, cLISI and graph connectivity to avoid part–whole overlap. Spearman correlations are −0.591, 0.687 and 0.798, respectively; *P* values are nominal asymptotic two-sided values. Lines and bands show ordinary-least-squares fits and pointwise 95% confidence intervals.

Established target-data integration methods provided a complementary comparison. Harmony, Seurat RPCA and de novo scVI moved into the upper performance tier when batch mixing was considered together with biological conservation, although they did not uniformly improve biological conservation alone (Supplementary Results). In contrast, the frozen scFMs exhibited more polarized behaviors: some preserved biological structure while retaining batch effects, whereas others mixed batches at the expense of biological conservation. Notably, the latter suggests that effective batch mixing does not necessarily reflect successful integration, but may instead arise from diminished biological resolution. This pattern reinforces the trade-off between biological conservation and batch mixing [29], and none of the evaluated frozen scFMs shifted the balance toward simultaneous improvements in both properties.

### 2.3 Embedding geometry, but not model characteristics, explains zero-shot representation quality across scFMs

The wide variation in biological-conservation performance across scFMs motivated us to ask: what properties distinguish a biologically useful representation from one that fails? In particular, we investigated whether the observed performance differences among methods can be explained by measurable properties of the embedding geometry. We first investigated global (spectral) anisotropy, a geometric property in which cell vectors share a dominant global direction and occupy a narrow cone in the latent space. Across the 20 representations, greater spectral anisotropy was associated with poorer biological-conservation performance (Spearman *ρ* = −0.591, nominal *P* = 0.00607; Fig. 2c,d). PCA and the established integration methods exhibited relatively low global anisotropy, whereas GenePT-w, Geneformer, scBERT and scPRINT showed pronounced directional concentration.

However, low global anisotropy alone was not sufficient to achieve good biological performance: a representation may be globally diffuse yet still fail to organize cells according to biological identity. To capture whether directional structure is biologically meaningful, we decomposed cosine alignment into within- and between-cell-type components. The difference between these quantities, defined as within-cell-type minus between-cell-type anisotropy, measures whether directional coherence is specific to cells sharing a biological identity rather than being imposed on all cells. This cell-type-specific anisotropy emerged as a substantially stronger predictor of downstream performance (*ρ* = 0.687, nominal *P* = 8.15×10^−4^). Among frozen scFMs, SCimilarity showed the strongest median cell-type anisotropy contrast, with scCello and CellPLM also displaying clear cell-type-specific organization (Fig. 2c). By contrast, CellFM did not exhibit strong global cone collapse, but its within-cell-type alignment was nearly indistinguishable from its between-cell-type alignment. Its embedding was therefore dispersed without effectively organizing variation around annotated biological identities.

The second geometric diagnostic was local expression-neighbourhood preservation. We quantified this by chance-corrected *R_NX_*between within-batch neighbour sets in each representation and a reference constructed directly in analytic-Pearson-residual expression space. This analysis examines whether an embedding preserves fine-grained local expression relationships, rather than merely separating broad cell classes. Across all 20 representations, mean *R_NX_*was strongly associated with downstream performance (*ρ* = 0.798, nominal *P* = 2.44×10^−5^; Fig. 2c,d); this association was even stronger among the 13 frozen scFMs (*ρ* = 0.901, nominal *P* = 2.61×10^−5^; Supplementary Results). Established methods, including de novo scVI, classic HVGs+PCA, Harmony, and Seurat maintained relatively high local fidelity. Among frozen scFMs, scFoundation and SCimilarity were among the strongest, whereas scBERT, GenePT-w, and CellFM substantially distorted the reference neighbourhood structure.

Together, these results highlight two key geometric determinants of representation quality: avoidance of non-specific directional collapse and preservation of local expression neighbourhoods.

To determine whether these geometric patterns also generalized from discrete cell-type organization to continuous developmental structure, we next examined the trajectory inference setting across nine developmental systems spanning linear, bifurcating and tree-structured trajectories. Trajectory inference provides a complementary setting to cell typing as the continuous ordering and branching are reconstructed from intercellular distances, neighbourhood graphs and manifold structure rather than from discrete cell-type assignments. We therefore examined whether the same geometric properties also tracked the performance of continuous trajectory recovery relative to PCA. Trajectory recovery was similar for the leading scFMs and PCA (mean aggregate ranks 6.52 for scFoundation, 7.21 for CellPLM and 7.27 for PCA), and no method outperformed PCA after multiple-testing correction (Extended Fig. 2). Across the full trajectory panel, greater spectral anisotropy was associated with poorer trajectory recovery (*ρ* = −0.446, *P* = 0.0759), while expression-reference *R_NX_*was positively associated with PCA-relative recovery across nine developmental systems and four trajectory-inference methods (*ρ* = 0.775, *P* = 4.50×10^−4^; Extended Fig. 1g,h). The recurrence of these associations across discrete cell-type organization and continuous developmental structure supports their use as representation-level diagnostics.

Additional geometric probes revealed qualitatively distinct failure modes across scFMs, rather than a single monotonic “better-to-worse” performance axis (Fig. 2c). On one hand, GenePT-w and scBERT produced very few effective dimensions (mean participation ratios of approximately 1.36 and 1.99, respectively), consistent with severe variance concentration. At the opposite extreme, CellFM distributed variation across hundreds of effective dimensions (mean participation ratio approximately 357; mean TwoNN intrinsic dimension approximately 159), yet this high-dimensional variation did not translate into strong biological organization. These observations indicate that both dimensional collapse and high-dimensional but biologically unstructured variation can compromise representation quality. Notably, many well-behaved embeddings had effective dimensionalities on the order of only 10–30, suggesting substantial redundancy in the hundreds- to thousands-dimensional representations commonly used by current scFMs. This suggests that high nominal dimensionality is not necessary for capturing the biologically relevant dominant structure and that considerably more compact representations may suffice. Importantly, this observation provides an empirical reference for selecting representation bottlenecks or projection dimensions and motivates systematic investigation of whether explicitly targeting this dimensional regime can improve representation quality.

Partial *η*^2^ analysis provided a complementary separation of cell-type- and batch-associated variation (Extend Fig. 1a,b). SCimilarity showed strong cell-type-associated structure (mean cell-type partial *η*^2^ = 0.535), whereas CellFM and scBERT showed substantially weaker values (0.074 and 0.166, respectively). CellFM also had little batch-associated variation (mean batch partial *η*^2^ = 0.0065), demonstrating that weak batch association is not necessarily beneficial when biological structure is simultaneously lost. Many frozen scFM embeddings retained measurable batch-associated variation, further indicating that scFMs do not inherently disentangle biological structure from technical variation.

Finally, we asked whether these geometric and performance differences are predictable from broad model characteristics. In general, they were not. Neither reported pretraining corpus size (Spearman *ρ* = −0.237, permutation *P* = 0.480, *n* = 11 models) nor total parameter count (*ρ* = 0.392, permutation *P* = 0.211, *n* = 12) was significantly associated with PCA-relative biological performance. Model family accounted for a moderate descriptive fraction of between-model variation (*R*^2^ = 0.374), but the association was not significant under exact group-preserving permutation (*P* = 0.095). Whether the input preserved expression magnitude explained little variation (*R*^2^ = 0.018, exact *P* = 0.653), and none of the four model-characteristic tests remained significant after Holm correction (all adjusted *P* ≥ 0.379; Extended Data Fig. 1c-f). Notably, models with superficially similar designs could nevertheless fall at opposite ends of both the geometric and performance spectra, as illustrated by the ESM2-based UCE and scPRINT embeddings and by the continuous-expression scFoundation and CellFM embeddings. Collectively, these analyses show that broad model descriptors such as scale, architectural family, and input encoding are poor indicators of zero-shot representation quality; by contrast, the geometry of the output embedding provides a more direct diagnostic of whether biological structure has been retained.

### 2.4 Pretrained representations show selective benefits in label-limited supervised transfer and open-set detection

Single-cell foundation models are designed to support transfer across biological domains with limited supervision. We evaluated this premise for supervised cell-type transfer under domain shift, distinguishing frozen-embedding transfer from model-specific supervised adaptation. We then asked how performance depended on the number of labelled reference cells and whether the query contained cell types absent from the reference.

In data-rich, closed-set transfer with abundant labeled reference cells and no semantically out-of-distribution (OOD) cell types at test time, established annotation methods performed best. CellTypist reached a median macro-F1 of 0.749, compared with 0.720 for scFoundation, the highest-scoring scFM. Pretrained scVI (0.707) and PCA (0.704) fell within the overall scFM range. Thus, under standard closed-set transfer with ample labels, we did not observe a consistent advantage for pretrained representations.

The ranking of methods changed when labelled reference cells were scarce. With only five labelled cells per class, scFoundation, scCello and UCE reached 90% of their own macro-F1 at the maximum label budget (*N* = 500); PCA required *N* = 50. With one labelled cell per class, rare-type recall (*<* 1%, LR-based) was 0.68 for UCE and 0.65 for scFoundation, compared with 0.61 for PCA and 0.48 for CellTypist; these gaps narrowed as labels accumulated (Extended Fig. 3). The main benefit of scFMs therefore appeared to be label efficiency rather than a higher data-rich performance ceiling. In this sense, pretrained representations function less as a universal performance boost and more as a prior that is most valuable when data are limited. Additionally, model-specific supervised adaptation had heterogeneous effects. Across the 13 scFMs, mean macro-F1 changed by a median of +0.03 relative to the corresponding frozen readout (Supplementary Results). The sole large gain was scBERT (+0.29), whose near-random frozen embedding is effectively rescued by adaptation, indicating that fine-tuning chiefly compensates for a weak base representation rather than extracting substantial new task signal from an already strong one.

The clearest benefit of pretrained embeddings emerged in open-set detection of source-absent cell types. Across 37 eligible transfer directions, feature-space distance scores were more informative than logit- or confidence-based scores. In particular, Mahalanobis distance computed directly in the frozen embedding space flagged novel cells with a mean AUROC of 0.82–0.84 for the leading foundation models (scFoundation, LangCell, UCE). This is exactly the axis on which the annotation baselines fall short. CellTypist and SingleR do not produce feature embeddings that are directly comparable in this setting, so we instead used their native scores as model-derived features, which showed weaker performance (AUROC 0.72–0.76). CellTypist, the single strongest closed-set classifier, was among the *weakest* novelty detectors (0.73). Among the methods that do produce an embedding, the foundation models led over the classical PCA embedding (0.73), with only the pretrained VAE baseline scVI (0.82) keeping pace. Moreover, even when a novel cell went undetected and was forced into a reference label, the error degraded gracefully: the assigned label shared the true type’s broad class in ~80% of cases and its coarse compartment in over 90%, and in up to ~40% was an ontological ancestor of the true cell type (UCE, scFoundation). Thus, novel-type detection is a property of the embedding’s distance geometry, and its residual errors respect cell-lineage structure—both are features largely invisible to conventional accuracy benchmarks (Fig. 3c,d).

**Fig. 3.**
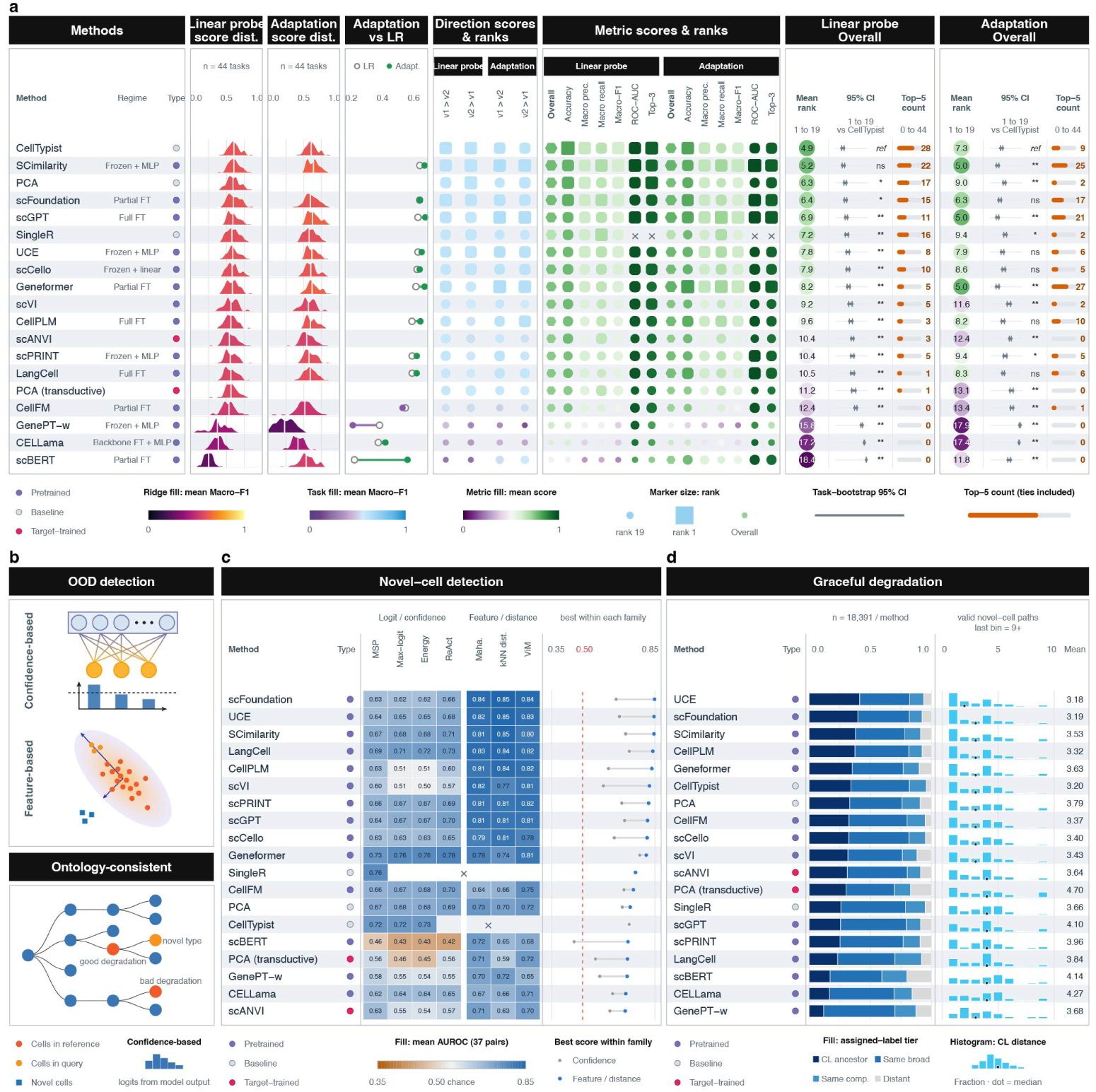
Supervised transfer and open-set semantic novel-cell detection. **a,** Frozen-embedding transfer and native established-method performance are compared with model-specific supervised adaptation across 44 directed cross-dataset transfers, with the complete query population retained, including source-absent types. Ridges show direction-level macro-F1, connectors show changes after adaptation, and matrices show direction- and metric-level scores and ranks. The right columns show mean rank with 95% intervals from 2,000 tissue-cluster bootstrap replicates. Symbols indicate two-sided paired Wilcoxon tests against CellTypist with Holm correction across methods (∗∗, adjusted *P <* 0.01; ∗, adjusted *P <* 0.05; ns, not significant; ref, CellTypist). CellTypist, SingleR, and scANVI are established methods with their own methodological frameworks and are included alongside these approaches solely for comparison; although they perform supervised transfer, they do not fall under our definitions of either linear probing or model-specific adaptation. **b,** Schematics of confidence- and feature-based novelty detection and ontology-based assessment of forced reference-label assignments. **c,** Mean novel-type-detection AUROC across 37 eligible transfer directions; crosses denote unavailable endpoints and dots compare the best score in each family. **d,** Ontological proximity of forced assignments for 18,391 source-absent cells per method. Bars show Cell Ontology ancestor, same broad class, same compartment and distant assignments; histograms show path distance and dots indicate medians.

Taken together, these results refine the value of single-cell pretraining. Pretraining front-loads a representation whose benefit is realized cheaply, with a few labels and a linear probe, and is concentrated in the regimes that matter operationally: scarce labels, rare cell types and unseen states. For closed-set annotation with abundant labels, the advantage is modest: a well-trained VAE and even PCA largely keep pace; no foundation model surpassed CellTypist, and fine-tuning adds little. The distinctive payoff of pretraining lies instead in open-set generalization: only a learned embedding supports the distance-based detection of novel cell types, an axis on which the foundation models lead and classifier baselines structurally cannot compete, and even the resulting errors respect cell-lineage structure. For routine annotation, frozen embeddings with a linear probe are therefore the pragmatic default; the case for a foundation model is strongest when labels are few or the biology is new.

### 2.5 scFMs transfer unreliably to spatial transcriptomics

To test whether scFMs generalize beyond scRNA-seq, we evaluated their direct transfer to spatially resolved transcriptomics (SRT). We assessed spatial-domain clustering across three platforms and, separately, cell-type clustering in MERFISH. This setting presents a substantial modality shift from scRNA-seq: spatial transcriptomics platforms such as Visium and ST do not achieve single-cell resolution, whereas imaging-based methods such as MERFISH measure restricted gene panels. Performance on these datasets therefore provides a stringent test of representation robustness beyond the training modality. We evaluated 12 dorsolateral prefrontal cortex sections (SpatialLIBD, Visium platform) [30], 8 breast cancer sections (HER2ST, ST platform) [31] and 5 hypothalamic sections (MERFISH) [32]. The spatial-domain comparison comprised all 13 frozen scFMs, pretrained scVI, de novo scVI, PCA, and NOVAE [33], an SRT-native foundation model that additionally uses the target spatial coordinates. Under the multi-metric aggregate, NOVAE ranked first in each dataset and achieved the best mean rank across the 25 sections (2.87), followed by scFoundation (5.98), CellPLM (6.01), scVI trained de novo (6.32) and scGPT (6.39) (Fig. 4). NOVAE significantly outperformed all comparators (all Holm-adjusted *P* = 1.96×10^−4^), whereas PCA ranked seventh overall (mean rank 7.61), ahead of 10 of the 13 frozen scFMs. When we excluded NOVAE, the top-performing specialized method, the best remaining approach differed by dataset: scFoundation on SpatialLIBD, scGPT on HER2ST and PCA on MERFISH. Overall, spatially native modeling produced the strongest domain recovery, whereas frozen scRNA-seq representations transferred inconsistently and showed no consistent advantage over PCA.

**Fig. 4.**
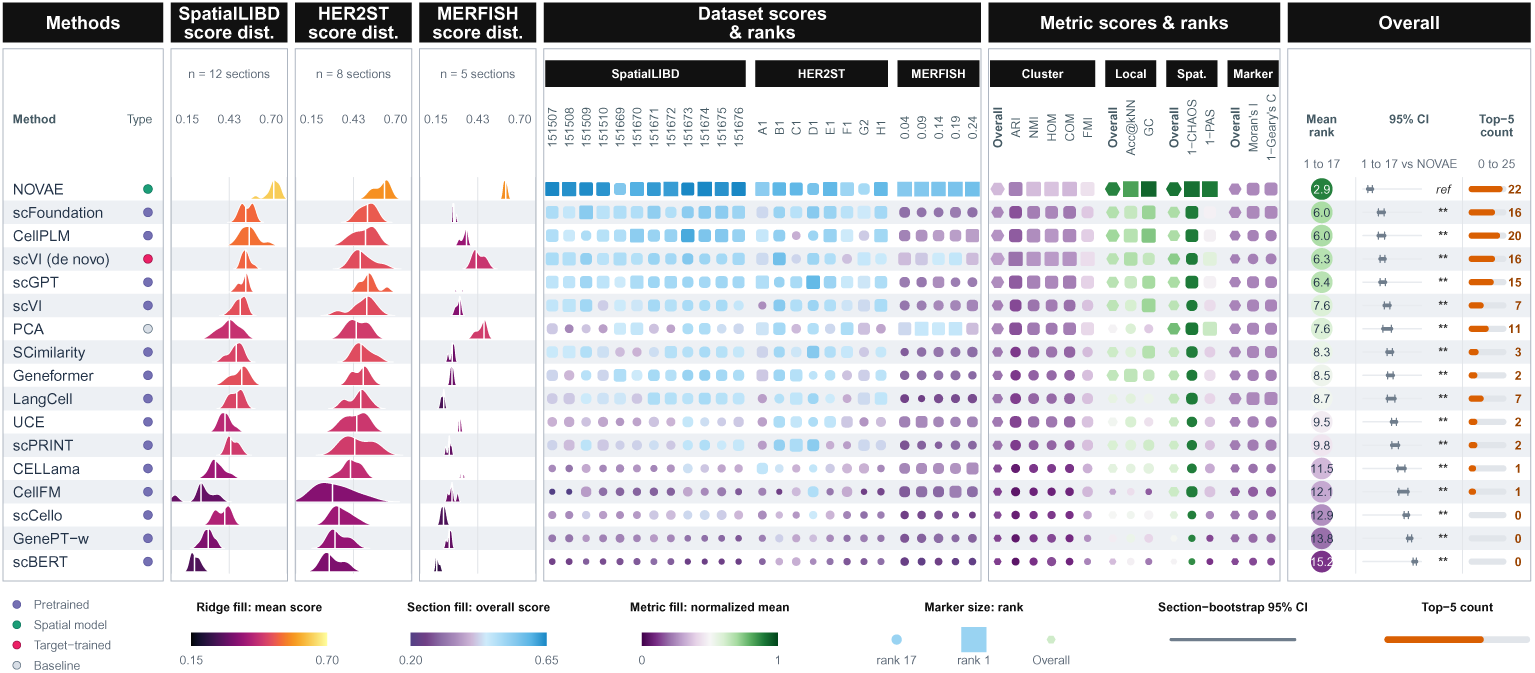
Spatial-domain recovery across spatial-transcriptomics datasets. Seventeen methods were evaluated across 25 sections from SpatialLIBD (*n* = 12), HER2ST (*n* = 8) and MERFISH (*n* = 5); task-native NOVAE additionally uses spatial coordinates. Ridges show section-level aggregate-score distributions, and the matrix shows scores and ranks for individual sections. Aggregate ranks combine cluster concordance, local structure, spatial coherence and spatial-marker autocorrelation. The right columns show mean rank with 95% section-bootstrap intervals and the number of sections ranked in the top five. Symbols indicate two-sided paired Wilcoxon tests against NOVAE with Holm correction across 16 comparisons (∗∗, adjusted *P <* 0.01; ∗, adjusted *P <* 0.05; ns, not significant; ref, NOVAE). Full metric definitions and rank construction are provided in Methods and Supplementary Methods.

The pattern changed in the separate MERFISH cell-type analysis. In this analysis, CELLama had the highest mean ARI (0.358), followed by scVI trained de novo (0.350) and PCA (0.317), whereas NOVAE ranked 16th of 17 (0.041; Supplementary Results). This result is consistent with cell-type identity being an intrinsic rather than a spatial property. More generally, the reversal indicates that relative method performance depended on which biological target the labels captured; although it does not, by itself, explain the underlying mechanism.

Notably, applying the spatial diffusion strategy [34] improved spatial-domain recovery of different methods mainly at single-cell resolution, with less benefit at spot resolution. However, the same diffusion-based adaptation benefited the PCA baseline at least as much as the scFM representations, while all adapted representations still underperformed the SRT-native NOVAE. Together, these results suggest that the observed gains arise primarily from the injected spatial context rather than from scFM pretraining, highlighting the need for modality-aware modeling or adaptation. Together these results show that scFMs pretrained on scRNA-seq do not transfer reliably to SRT: they neither match a purpose-built spatial model on spatial-domain tasks nor consistently outperform a simple PCA baseline, and their relative ranking shifts with resolution, gene-panel breadth and evaluation metric.

### 2.6 Gene embeddings capture broad functional relatedness but not reliably context-specific regulatory structure

The preceding analyses evaluated whether cell embeddings preserve biological structure. We next asked whether the gene-level representations exposed by scFMs provide a useful substrate for functional and regulatory inference. This distinction is important because different model outputs support different biological interpretations: a model may provide a static gene-embedding table, contextual gene representations derived from the target cells, or a model-specific regulatory-network readout, each of which reflects a distinct type of biological information. We therefore evaluated four complementary properties: broad, context-independent functional relatedness; regulatory-edge prioritization in target cells; recovery of recorded transcription-factor regulons; and retrieval of genes with large observed responses to transcription-factor perturbation.

We first evaluated functional coherence on fixed common-gene universes using high-confidence physical protein–protein interactions and nearest-neighbour retrieval of shared Gene Ontology Biological Process, Reactome and KEGG annotations (Fig. 5b). The text-derived GenePT-w representation was the clear exception to the generally modest performance of transcriptome-derived gene embeddings. GenePT-w achieved a STRING physical-interaction AUROC of 0.827 and an AUPRC of 0.641, and obtained mean average precisions of 0.156, 0.196 and 0.247 for GO-BP, Reactome and KEGG retrieval, respectively. By comparison, the corresponding co-expression values were 0.585 and 0.286 for PPI and 0.070, 0.072 and 0.075 for the three retrieval tasks. Other scFM representations contained heterogeneous functional signal: several exceeded co-expression for physical-interaction ranking, but most showed only modest pathway-level coherence, and the static scFoundation and CellFM embeddings were close to random for physical interactions (AUROC 0.508 and 0.498). Recomputing contextual gene representations did not yield a uniform improvement. For example, contextualization increased the scFoundation physical-interaction AUROC from 0.508 to 0.639, whereas Geneformer changed from 0.675 to 0.629. Thus, transcriptome pretraining does not automatically organize the accessible gene space according to curated molecular function. Conversely, GenePT-w shows that gene-description language embeddings can supply a strong functional-semantic prior, although this may partly reflect biological knowledge already represented in the source text rather than de novo inference from expression data.

**Fig. 5.**
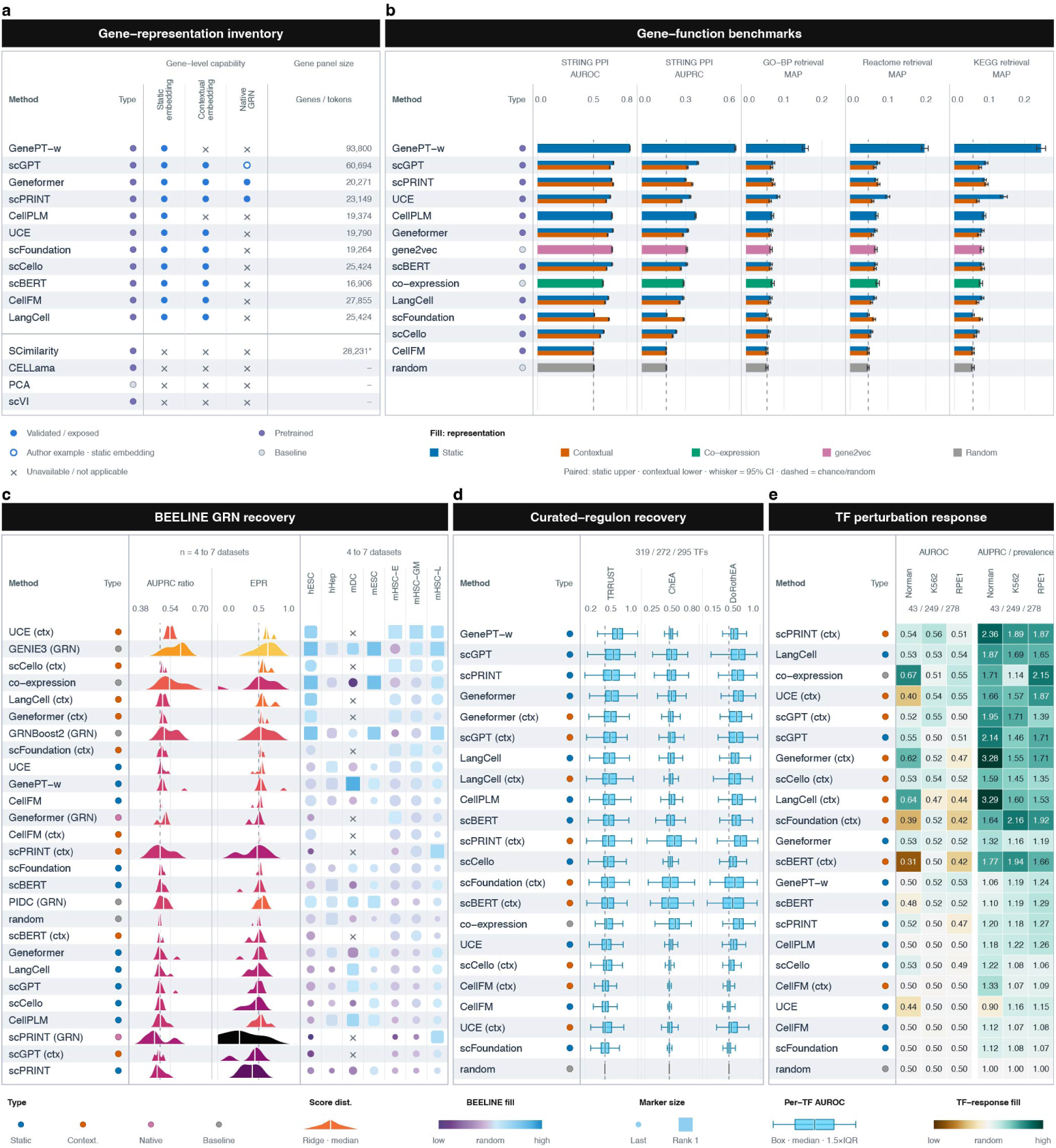
Functional and regulatory information in gene-level model outputs. **a,** Availability of static, contextual and model-native outputs across 15 methods; the hollow scGPT symbol denotes its static-embedding GRN route. **b,** Functional coherence of static and contextual representations on a common universe of 16,508 genes, compared with established co-expression and gene2vec references and a random reference. **c,** BEELINE regulatory-edge recovery for static representations and established GRN methods across seven datasets, and for contextual and model-native outputs across four count-compatible datasets. AUPRC and early-precision ratios relative to random are displayed as *r/*(1 +*r*), for which 0.5 denotes random expectation. **d,** Recovery of TRRUST, ChEA and DoRothEA ABC regulons (*n* = 319, 272 and 295 transcription factors). **e,** Retrieval of the 100 largest observed response genes after single-transcription-factor perturbation in Norman (*n* = 43), Replogle K562 (*n* = 249) and Replogle RPE1 (*n* = 278) experiments; contextual representations use control cells only.

Broad functional relatedness, however, did not translate into recovery of directed, context-specific regulation. We thus evaluated static representations alongside the task-specific algorithms GENIE3, GRNBoost2 and PIDC on seven BEELINE datasets from five independent studies [35]. To test whether target-cell context improved regulatory information, we additionally computed contextual states from nine eligible models and completed native outputs from Geneformer and scPRINT on the four datasets with available count matrices. The results shows that regulatory-edge recovery was weak and inconsistent across scFM output types (Fig. 5c and Supplementary results). Among methods designed specifically for GRN inference, GENIE3 was strongest overall and reached an aggregate AUPRC ratio of 1.27; no scFM-derived static, contextual or native output exceeded this value. Contextualization nevertheless added detectable signal: on the common four-dataset panel, the combined AUPRC and early-precision score improved over the corresponding static representation for six of the nine eligible models. UCE ranked first within the contextual track, but its AUPRC ratio remained 1.15, and the advantage was not established statistically. GenePT-w led the static scFM track but reached an AUPRC ratio of only 1.12, showing that its strong functional-semantic organization did not transfer to context-specific regulatory-edge recovery. Neither of the completed model-native routes produced a consistent advantage. Target-cell contextualization can therefore expose additional regulatory signal, but current scFM outputs remain substantially less reliable than required for direct GRN inference.

The two target-recovery analyses further separated retrieval of recorded knowledge from recovery of experimentally observed responses (Fig. 5d,e). GenePT-w had the highest median AUROC for TRRUST (0.689), a resource assembled from published regulatory relationships and therefore closely aligned with a text-derived biological prior. Its advantage did not persist for ChEA or DoRothEA ABC: ChEA recovery was weak across methods, whereas contextual scPRINT led DoRothEA ABC with a median AUROC of 0.685. The distinction was clearest across the 570 single-transcription-factor CRISPR perturbations. Contextual Geneformer, LangCell and scPRINT yielded mean AUPRC lifts of 2.454, 2.429 and 2.118, respectively, whereas GenePT-w was close to chance (AUROC 0.512; AUPRC lift 1.134). In addition, cell-embedding-based perturbation prediction struggled to outperform a simple additive model (Supplementary results). Thus, text-derived embeddings can recover biological relationships represented in the literature without necessarily identifying the regulatory responses expressed in target cells. These endpoints recover recorded regulons or genes with large observed responses; they do not identify direct targets or predict complete post-perturbation transcriptomes.

Together, these analyses show that current scFM gene-level outputs are not sufficiently reliable for direct use in GRN inference or perturbation-response modelling. Broad functional knowledge was evident, most clearly in the text-derived GenePT-w representation, but did not translate consistently into context-specific regulatory edges, experimentally observed response genes, or strong cell-level representations. The official expression-weighted aggregation of GenePT-w, for example, produced highly anisotropic cell embeddings, with a mean participation ratio of approximately 1.36, and poor downstream performance (Fig. 2). This contrast points to a gene-to-cell aggregation bottleneck, whereby a shared directional component can dominate the weighted sum of individually informative gene vectors. Gene representation, contextual regulatory inference, perturbation response, and gene-to-cell aggregation should therefore be treated as distinct model capabilities that require task-specific validation.

### 2.7 Computational resource profiling

To characterize computational demands, we implemented a unified resource-profiling protocol covering CPU memory usage, runtime and GPU memory footprint. All experiments were conducted on an HPC cluster equipped with NVIDIA H200 GPUs (140 GB) and submitted via Slurm with a fixed allocation of 256 GB system memory, four CPU cores and one H200 GPU per run.

Because GPU memory is strongly influenced by batch size and runtime depends on data scale, we standardized inputs to 10,000 cells and 30,000 genes. For GPU-accelerated methods, inference was performed with batch sizes of 8, 16, 32, 64 and 128 whenever supported. CELLama, implemented within LangChain, does not expose an explicit batch-size parameter, and scFoundation performs inference in a cell-by-cell manner by design. SCimilarity was evaluated in both its default CPU configuration and an optional GPU-enabled configuration, whereas GenePT-w runs exclusively on CPUs; therefore, GPU memory is not reported for GenePT-w.

As shown in Fig. 6a, peak resident set size (RSS) was largely insensitive to batch size across methods. SCimilarity showed the highest peak RSS due to CPU-based execution, while scPRINT, scCello and Geneformer also exhibited higher CPU memory usage. As batch size increased, both peak allocated and reserved GPU memory increased accordingly (Fig. 6b,c). When batch size was set to 32 or lower, all GPU-enabled methods could be executed on 40 GB GPUs. scFoundation, scBERT and Geneformer exhibited higher GPU memory usage, with scBERT and Geneformer showing more pronounced increases at larger batch sizes. CellPLM displayed distinct scaling: batch size primarily affects effective sequence length rather than the number of simultaneously processed cells, resulting in minimal changes in GPU memory while improving throughput.

**Fig. 6.**
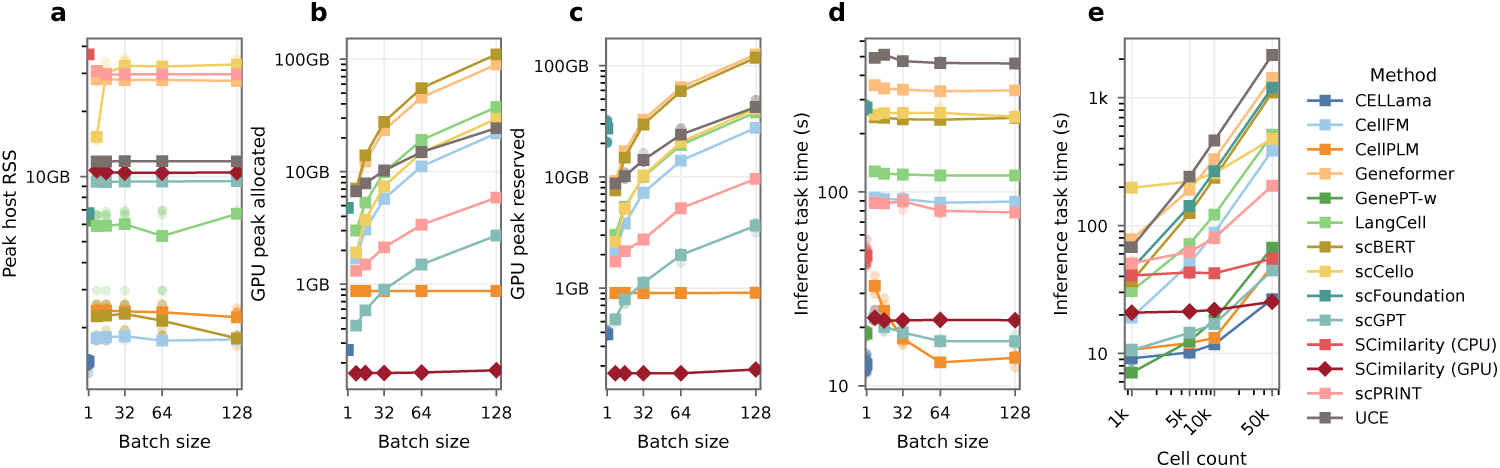
Computational resource profiling of single-cell foundation models. **a**, peak resident set size (RSS) as a function of batch size. **b**, peak GPU memory allocated as a function of batch size. **c**, peak GPU memory reserved as a function of batch size. **d**, wall-clock runtime as a function of batch size. **e**, runtime as a function of the number of processed cells.

For most methods, end-to-end runtime was dominated by data preprocessing, model loading and I/O, and batch size had limited impact on total inference time (Fig. 6d). Runtime scaled approximately linearly with cell number across methods (Fig. 6e); CELLama, CellPLM, scGPT and GenePT-w were faster, and SCimilarity became increasingly advantageous as the number of cells increased.

## 3 Discussion

In this study, we developed a unified and reproducible framework for standardized deployment and controlled evaluation of scFMs across diverse datasets, computational environments, training regimes, and downstream analyses, substantially lowering the technical barrier to their use. Leveraging this framework, we conducted a systematic and reproducible evaluation of single-cell foundation models (scFMs) across diverse analytical scenarios. Across the evaluated settings, scFMs did not show a uniform advantage over established methods. Instead, their utility was more closely associated with the biological structure retained in their learned representation than with model scale or architecture alone. Established methods remained competitive in standard settings, whereas selected pretrained representations provided advantages primarily when supervision or reference information was severely limited. These findings suggest that the practical value of scFMs lies not in replacing conventional methods broadly, but in providing useful representations under specific biological and statistical regimes.

Across zero-shot analyses, a consistent pattern emerged. Most scFMs preserved recognizable biological structure, but this rarely translated into consistent improvement over PCA. Importantly, our geometric analyses identified measurable properties of the embedding space that were more informative than conventional measures of model complexity in explaining model performance. Representations that better preserved local expression neighbourhoods and organized directional structure according to cell identity tended to show stronger biological conservation and hence better model performance, whereas non-specific directional concentration was associated with poorer performance. Local expression fidelity also tracked a model’s ability to recover continuous developmental trajectories. In contrast, standard model characteristics including parameter count, pretraining-corpus size, architectural family, and nominal embedding dimension were weak predictors of representation quality across the evaluated methods. These results suggest that representation geometry, rather than model scale itself, may provide a more useful basis for understanding and improving scFM representations.

The limitations of general-purpose pretraining became particularly apparent when downstream inference required information specific to the target modality or biological context. Under modality shift, frozen scRNA-seq representations did not consistently improve spatial-domain recovery, whereas a spatially native representation was more effective for recovering tissue domains. At the gene level, text-derived embeddings captured broad curated functional relationships, but scFM outputs did not reliably recover context-specific regulatory edges or retrieve genes with large responses to transcription-factor perturbation. Although contextualization introduced additional regulatory signal for several models, it did not produce a consistent advantage. Moreover, in the text-derived model, strong gene-level organization did not translate into an informative cell representation. Thus, broad biological structure in a representation should not be equated with reliable task-specific biological utility: a representation can encode meaningful relationships while still lacking the information required for a particular inference task.

Limited supervision substantially changed this picture, but again in a selective manner. Established annotators remained strongest for conventional closed-set transfer, while selected pretrained representations showed greater label efficiency compared to existing methods at the smallest label budgets; those differences diminished as more supervision became available. A related benefit emerged in open-set settings in which the correct cell type was absent from the reference: in several pretrained spaces, absolute feature distance provided useful information beyond classifier confidence, whereas forced known-label assignments often remained biologically close to the true type. These results suggest that the principal advantage of pretraining was not a higher performance ceiling, but greater utility when supervision or reference coverage was most limited.

Taken together, our results position single-cell pretraining as a selective source of transferrable biological structure rather than a general replacement for established analysis methods. Pretrained representations could preserve useful local or semantic relationships, but these properties do not guarantee improved performance in closed-set annotation, spatial-domain recovery or regulatory analysis. The benefit of pretraining therefore depends on whether the information encoded by the representation matches the information required by the downstream task.

These findings suggest several testable directions for future model development. First, representation learning objectives could be designed to explicitly preserve biologically meaningful local expression neighbourhoods while avoiding non-specific directional concentration in the embedded representation. Second, spatial and regulatory applications may benefit from models that incorporate modality-specific information and mechanisms for representing cellular context rather than relying solely on general-purpose transcriptomic pretraining. Third, model development should be accompanied by transparent and reproducible evaluation of representation properties, not only downstream prediction accuracy. Such evaluation could help determine whether improvements reflect genuinely more informative biological representations or task-specific adaptation. By lowering technical barriers and enabling the same scientific questions to be revisited as models evolve, the framework presented here provides a foundation for more reproducible development, evaluation, and deployment of scFMs.

## 4 Methods

### 4.1 Data collection and curation

#### 4.1.1 Single-cell RNA-seq datasets

We assembled a diverse panel of single-cell RNA sequencing (scRNA-seq) datasets from public repositories, including CellxGene, the Dynverse trajectory benchmark datasets [36], and scPerturb [37]. Datasets were selected to satisfy three criteria: (i) availability of high-quality cell type annotations, (ii) presence of multiple donors or experimental samples, and (iii) coverage of diverse biological contexts, including immune systems, brain tissues, developmental processes and tumors. Raw count matrices and accompanying metadata were retrieved using repository-provided application programming interfaces or downloaded directly from supplementary materials when necessary. Additional task-specific datasets, eligibility criteria and sample counts are described in the Supplementary Methods.

#### 4.1.2 Quality control, normalization, and feature preparation

A uniform quality-control (QC) pipeline was applied to all scRNA-seq datasets before model execution and benchmark construction. To ensure fair comparison across models, we standardized the input format by using raw count matrices as the canonical input for all datasets. Although different single-cell foundation models require distinct input representations (e.g., raw counts or log-normalized values), all model-specific preprocessing steps were applied downstream of this standardized input to match each model’s expected format.

We disabled all model-specific internal QC procedures and required that QC be performed exclusively at the data preprocessing stage. This design ensures that all models encode the same set of input cells and minimizes confounding effects introduced by model-specific filtering strategies. Several auxiliary features required by scFMs were constructed during preprocessing, including gene symbols and Ensembl gene identifiers, as different models rely on distinct tokenization schemes. CellxGene datasets typically provide complete annotations; when gene identifiers were missing, we used the mygene package [38] to map and recover missing information. Each cell was additionally assigned a unique barcode or identifier to ensure consistent tracking across models and tasks.

No normalization or feature selection was performed at this stage. Instead, after constructing standardized inputs, each model applied its own internal normalization or feature-selection procedures according to its original design.

#### 4.1.3 Train, validation, and test splitting

Analysis-specific partitions were fixed and shared across methods. The zero-shot cell-representation, trajectory and spatial analyses did not use cell labels for model fitting. For supervised transfer, entire source domains were used for training and held-out target domains for evaluation. The primary frozen-transfer and label-efficiency comparisons were closed-set; source-absent target types were retained in the full-query supervised-adaptation and open-set detection analyses. All preprocessing and classifier fitting in the inductive comparisons used source data alone.

For model-specific supervised adaptation, 20% of the labeled source cells were reserved for validation and stratified by cell type. Label-efficiency analyses used identical source-cell samples and held-out targets across methods at each label budget.

### 4.2 Model implementation and wrapping

To enable fair, reproducible and extensible benchmarking, we implemented a unified model-wrapping framework that standardizes the execution of diverse single-cell foundation models while preserving their original architectures and training procedures. For each evaluated method, we treated the released codebase and pretrained weights as the authoritative implementation and did not modify model internals, loss functions or optimization strategies. In cases where released implementations contained software bugs or execution errors that prevented correct model operation, we applied minimal fixes at the implementation level to restore intended functionality, without altering the methodological objectives or core computational logic of the original models.

Each model was wrapped with a standardized interface that defines (i) input specifications, including required gene identifiers, expression formats and metadata; (ii) model execution modes, including embedding extraction, frozen-embedding transfer and model-specific supervised adaptation where available; and (iii) standardized output formats for downstream evaluation. This abstraction layer enables all models to consume harmonized inputs and produce comparable outputs while respecting model-specific preprocessing and inference logic.

To ensure reproducibility and minimize dependency conflicts, each wrapped model was encapsulated in a dedicated container environment that includes the exact software dependencies, library versions and runtime configurations specified by the original authors. When such information was incomplete, inconsistent or led to unresolved dependency conflicts, we resolved these issues conservatively to enable successful execution while preserving the original software environment as closely as possible. These containers were orchestrated through a workflow manager (Nextflow), allowing models to be executed as modular components within a directed acyclic graph (DAG) pipeline. This design ensures deterministic execution across computing platforms and enables one-command execution of all benchmark tasks.

Where models provided official supervised-training tutorials, we followed the recommended protocols and hyperparameter settings. For models lacking documented supervised-adaptation pipelines, we exposed the fixed pretrained embeddings to a standardized downstream classifier without introducing model-specific optimization. All hyperparameter choices and execution settings are explicitly recorded in configuration files to ensure full transparency and reproducibility.

Importantly, the wrapping framework is model-agnostic and extensible. New models can be integrated by implementing the same standardized interface without modifying existing workflows. This design lowers the technical barrier for non–computer science users to run and evaluate single-cell foundation models, while enabling the benchmark to evolve as new methods become available.

### 4.3 Model zoo

We evaluated thirteen single-cell foundation models spanning the major architectural and training paradigms in the field, together with classical baselines. To make the comparison interpretable, we catalogued each model along three documented design axes: its *training paradigm*, its *input encoding* (how a transcriptome is presented to the model) and its *backbone* (Supplementary Table S3). We then grouped the models into three families by training paradigm.

1. **Self-supervised masked or generative encoders**, which represent a cell as a set or sequence of gene tokens and are trained to reconstruct masked genes or expression values. This dominant paradigm comprises Geneformer[5], scGPT[4], scBERT[20], scFoundation[6], CellFM[16], CellPLM[17], scPRINT[23], and UCE[24]. Members differ chiefly in how expression is encoded: as a *rank* ordering of gene tokens with magnitude discarded (Geneformer); as *quantile-binned* expression values (scGPT; scBERT, which pairs binned values with gene2vec gene embeddings); as *continuous* expression values (scFoundation, via a learnable auto-discretization with read-depth tokens; CellFM; and CellPLM, which forms an expression-weighted sum of learnable gene embeddings under an inter-cell Gaussian-mixture latent); or via *protein-language-model (ESM2) gene embeddings* for gene identity combined with expression (scPRINT, trained by expression denoising, and UCE, which uses expression only as a gene-sampling weight).
2. **Contrastive or label-supervised encoders**, which shape the embedding space using biological or textual supervision rather than pure reconstruction: SCimilarity[22], a multilayer-perceptron autoencoder trained by supervised metric (triplet) learning on curated cell-type labels; scCello[21], a Transformer trained with cell-type-ontology–guided contrastive objectives on top of a masked gene-prediction (Geneformer-style) base; and LangCell[19], whose Geneformer-based cell encoder is aligned to text by cell–text contrastive learning.
3. **Language-model–derived encoders**, which textualize the transcriptome and reuse general-purpose language-model embeddings: GenePT[18] (GenePT-w), which represents a cell as an expression-weighted average of frozen GPT-derived gene-description embeddings, and CELLama[15], which renders a cell as a rank-ordered sentence of its top-expressed gene symbols embedded by a sentence transformer.

Apart from SCimilarity (a multilayer perceptron) and the language-model–derived encoders, the backbones are all Transformer variants—including efficient- or linear-attention formulations (scBERT’s Performer, CellPLM’s Flowformer), a retention network (CellFM’s ERetNet) and an asymmetric encoder–decoder (scFoundation’s xTrimoGene)—so backbone architecture does not by itself distinguish the models. Because these axes co-vary—each model differs simultaneously in encoding, objective and backbone, and several (e.g. scCello, LangCell) combine multiple objectives—we treat this taxonomy as a descriptive scaffold for interpreting performance rather than as a set of independently manipulable factors. Full per-model implementation details, including pretrained checkpoints and the minimal fixes applied to enable faithful execution, are given below (M1–M13).

#### M1: scGPT (v0.2.4)

We adopt the “scGPT human” pretrained model as the base model. For zero-shot analysis and frozen-embedding transfer, we directly employ the embedding pipeline provided in scGPT to encode cells and obtain the latent representations. For supervised adaptation, although scGPT does not provide an integrated training pipeline, it includes an example training script that we follow with the released hyperparame-ters. Specifically, we set ‘mask_ratio’ = 0, ‘n_bins’ = 51, disable ‘MVC’, ‘ADV’, ‘CCE’, and set ‘ecs_thres’ = 0 and ‘dab_weight’ = 0. For cell annotation, we enable the ‘CLS’ option (‘CLS’ = True). After training or frozen embedding extraction, the cell representations are obtained directly from the scGPT embedding pipeline following the official usage instructions (https://github.com/bowang-lab/scGPT).

#### M2: Geneformer (v0.1.0)

We employ the Geneformer-V2-316M pretrained model as the base model. Geneformer provides an embedding pipeline, in which we set model type = “Pretrained”, emb_mode = “cell”, and model version = “V2” to obtain the cell-level embeddings. For supervised adaptation, 20% of the labelled source data were selected as a validation set to determine the optimal checkpoint. We used the released partial-updating route with freeze_layers=2, a learning rate of 5×10^−5^, weight decay of 0.01 and cosine decay with a warm-up ratio of 0.1. A minimum of 1,000 optimization steps was used so that the schedule remained meaningful for small reference datasets. We did not perform seed search. After training, cell embeddings were obtained following the official Geneformer pipeline (https://huggingface.co/ctheodoris/Geneformer).

#### M3: scFoundation (github version 397631c)

We utilize the only pretrained checkpoint released by the authors. Following the official preprocessing script provided in the repository, we perform identical data preprocessing steps. Regardless of how many gene panels are present in the input dataset, all inputs are aligned to a unified set of 19,264 genes, consistent with the model’s expected input dimension. For the zero-shot embedding task, we employ the official inference pipeline with the following configurations: input_type = “singlecell”, output_type = “cell”, pool_type = “all”, tgthighres = “t4”, disabling pre normalized, and setting version = “ce”. These settings follow the authors’ recommended usage for producing cell-level embeddings. For supervised adaptation, scFoundation does not provide a ready-to-use training pipeline. The repository includes the intended classifier architecture and trainable scope but lacks the essential training procedure, hyperparameter specifications or end-to-end scripts. We therefore retained the indicated partial-updating route: token and position embeddings and all encoder blocks except transformer_encoder[−2] were fixed, while that block and a max-pooling classification head were trained. We completed the training and inference pipeline using warm-up and cosine learning-rate decay. Additionally, we observe that scFoundation exhibits abnormally increasing GPU memory usage during inference when executed in full precision. To mitigate this issue and ensure reliable evaluation, we enable automatic mixed precision (AMP), which reduces memory footprint while preserving inference stability. The full implementation follows the official code structure as closely as possible, with necessary extensions to support end-to-end supervised adaptation and evaluation (https://github.com/biomap-research/scFoundation).

#### M4: CELLama (v0.1.0)

We employ the all-MiniLM-L6-v2 pretrained language model as the encoder within CELLama. Following the authors’ recommendation, for each cell we extract the top 30 highly expressed genes, convert them into a text-based “sentence,” and feed this sentence into the pretrained encoder to obtain the corresponding cell representation. Supervised adaptation used the released sentence-encoder procedure with four top-*k* configurations (*k* = 16, 20, 24, 28), two HVG set sizes (500 and 1,200), 1,000 validation examples and a learning rate of 1×10^−5^. Source and query cells were then embedded with the adapted encoder at *k* = 30, and the shared two-layer classifier was trained on the labelled source embeddings. All other settings followed the CELLama training script (https://github.com/portrai-io/CELLama).

#### M5: SCimilarity (v0.4.1)

We adopt model v1.1 as the pretrained model. Although SCimilarity provides a well-designed API and detailed usage tutorials, it offers limited flexibility for user-defined model customization. Following the official preprocessing workflow, we use the authors’ functions to align gene panels to the model’s expected feature space and perform all required data preprocessing steps. SCimilarity does not provide a model-specific supervised-adaptation interface. We therefore obtain cell embeddings using the official SCimilarity pipeline, keep the representation fixed and train the shared two-layer classifier on the labelled source embeddings. All steps follow the SCimilarity documentation (https://github.com/Genentech/scimilarity) as closely as possible while extending the downstream pipeline.

#### M6: scBERT (v1.0.0)

We use the panglao pretrain checkpoint, which is the only pretrained model released by the authors. The repository provides a minimal input data example; following this example, we align the gene panel accordingly and apply identical preprocessing steps to construct the model-compatible input sequences. Because scBERT does not provide a complete, ready-to-run embedding pipeline, we implement the embedding procedure by closely examining the authors’ code fragments and the model construction script. During this process, we identify that, in the original training procedure, the authors append a CLS token at the end of each cell’s gene sequence to encode cell-level features. Consequently, when generating embeddings, we extract the final token vector from the output sequence and use this vector as the cell representation, following the model’s intended design. For supervised adaptation, we extended the released implementation while retaining its optimizer, schedule and epoch cap. All pretrained parameters were fixed except the final normalization module and performer.net.layers[−2], which were trained together with a new classification head. All steps adhere as closely as possible to the official code structure, with necessary additions to support embedding extraction and supervised adaptation (https://github.com/TencentAILabHealthcare/scBERT).

#### M7: CellFM (github version 5054a2a)

We employ CellFM_80M weight as the pretrained model. CellFM is the only method in our benchmark that is natively trained and inferred using MindSpore. Although the authors provide a PyTorch implementation, it is currently limited to the cell type annotation task.

During inference, we identify several issues in the released codebase. First, the original data preparation logic does not allow setting mask_ratio = 0, i.e., disabling masking. We correct this behavior to enable unmasked inference. Second, in the released implementation, the inference path reuses the training-time data augmentation logic, where gene expression values are sampled to enhance training diversity. For embedding extraction, we modify the code to ensure that the model receives the original expression values rather than the sampled expressions used during training. At the time of manuscript preparation, the upstream repository still applies the training logic during inference, and our benchmark uses the corrected inference procedure.

Furthermore, we observe substantial issues in the decoder parameters of the released pretrained model: some decoder layers contain parameters that remain entirely zero (indicating initialization without effective training), while others have parameters with absolute values consistently below 1e-15, leading to nearly all-zero outputs. Due to these issues, we restrict the use of CellFM to the encoder component only in all experiments.

For supervised adaptation, we reference the official MindSpore training procedure provided by the authors and reimplement the annotation logic in PyTorch, ensuring compatibility with our benchmarking framework. The MindSpore-style annotation head and optimization strategy were retained, with parameters whose names contain encoder or head. updated and the remaining extractor fixed (https://github.com/biomed-AI/CellFM-torch).

#### M8: LangCell (github version 69e41ef)

LangCell consists of two modality-specific encoders, namely a cell encoder and a text encoder, trained via cross-modal contrastive learning. We use the only checkpoint released by the authors as the pretrained model. Since all tasks in our benchmark involve single-modality input (cellular expression data only), to ensure a fair comparison across methods, we use only the cell encoder as the embedding model for LangCell.

Although the original LangCell training framework is designed for contrastive learning between cell expression and text modalities, we follow the authors’ model usage and architectural design, and adapt the inference and training procedures to fit the specific tasks in our benchmark. Specifically, we extract cell-level embeddings directly from the pretrained cell encoder, and modify the downstream usage accordingly to support our evaluation tasks. All adaptations preserve the original model structure while removing text-modality dependencies, enabling LangCell to be evaluated consistently alongside other single-cell foundation models (https://github.com/PharMolix/LangCell).

#### M9: scCello (github version 767585b)

We use scCello-zeroshot as the pretrained model. Although scCello provides an official codebase, we identify multiple implementation issues that prevent direct and reproducible usage, and therefore apply a series of necessary corrections to enable stable benchmarking.

First, the cell type annotation script is hard-coded to read a specific input file and does not expose a user-facing interface for arbitrary datasets; we modify the script to accept external user data. Second, the codebase depends on an outdated version of the transformers library, with implicit coupling to the training logic, yet the required version is not documented. We resolve this by updating and fixing the dependency to a compatible, explicitly specified version. Third, the original implementation enforces distributed training and does not provide an option to disable Weights & Biases (wandb) logging; we adjust the training configuration to allow single-process execution and disable wandb when not required. Finally, we correct an inconsistency in the configuration files, where the documentation indicates a labels field, but the training code expects label instead.

All aforementioned fixes are applied without altering the core model architecture or training objectives. After these corrections, we use the official scCello workflow to generate embeddings; supervised adaptation keeps the pretrained encoder fixed and trains the released linear classification head (https://github.com/DeepGraphLearning/scCello).

#### M10: scPRINT (v2.3.5)

We employ the v2-medium checkpoint as the pretrained model. scPRINT is released as a relatively complete and well-packaged library, and we use its official embedding interface with the following parameters: how = “random expr”, max_len = 4000, and pred_embedding = [”cell_type_ontology_term_id”].

scPRINT includes its own data preprocessing pipeline. To ensure that all input cells are encoded in our benchmark, we disable all built-in filtering steps or set the corresponding thresholds to minimal values, thereby preventing the exclusion of cells during preprocessing. Cell-level embeddings are then generated using the official scPRINT embedder.

scPRINT does not provide an executable end-to-end supervised annotation script for arbitrary h5ad inputs. We therefore kept the released representation fixed and trained the shared two-layer classifier on labelled source embeddings. In addition, scPRINT automatically regenerates obs names during preprocessing, which complicates alignment with downstream task labels and evaluation.

We also identify a data-related bug in the released label decoders: the field “self_reported_ethnicity ontology_term_id” may contain multiple labels, causing mapping failures during decoding. To ensure stable execution, we retain only the first label for this field. All fixes and adjustments are applied without modifying the core scPRINT model or embedding mechanism, enabling consistent evaluation within our benchmark (https://github.com/cantinilab/scPRINT).

#### M11: CellPLM (v0.1.0)

. We use the 20231027_85M.best checkpoint as the pretrained model. CellPLM provides a relatively complete training and inference pipeline, and in our benchmark we adopt the default hyperparameter settings released by the authors, with only minor modifications to address implementation details.

Specifically, although CellPLM employs early stopping during training, the original pipeline does not save intermediate checkpoints nor return the best-performing model at the end of training. Moreover, the released workflow tightly couples training and prediction, making it difficult to independently evaluate on a held-out test dataset. To enable standardized benchmarking, we decouple the training and prediction stages, explicitly track the best validation performance, and ensure that the corresponding model state is used for downstream evaluation.

In addition, we modify the output interface to return logits directly, which is required for consistent evaluation across tasks in our benchmark. All adjustments preserve the original model architecture and optimization strategy while enabling reliable testing and fair comparison with other single-cell foundation models (https://github.com/OmicsML/CellPLM).

#### M12: GenePT (github version 3602699)

GenePT includes two variants, namely the “w” model and the “s” model. In our benchmark, we directly use GenePT-w as the evaluated model. We adopt the precomputed gene embeddings released by the authors, which are generated using the ChatGPT ada-002 embedding model. Following the official preprocessing protocol, we compute cell-level representations by applying the authors’ weighted aggregation scheme over gene embeddings.

GenePT is designed as an encoding-only model and does not provide a model-specific supervised-adaptation procedure. We therefore kept the GenePT-w representation fixed and trained the shared two-layer classifier on its labelled source embeddings (https://github.com/yiqunchen/GenePT).

#### M13: UCE (github version 8227a65)

We employ the 33l_8ep_1024t_1280 checkpoint as the pretrained model. UCE provides a complete and well-documented pipeline, and for embedding extraction we follow the authors’ official workflow and default parameter settings almost entirely. The only modification we make is to wrap the output embeddings into a unified format, ensuring consistency with the output structure used by all other methods in our benchmark.

UCE does not provide a model-specific supervised-adaptation procedure. For label-efficiency analysis and supervised adaptation, we therefore kept the UCE representation fixed and trained the corresponding source-labelled readout on its embeddings (https://github.com/snap-stanford/UCE).

### 4.4 Training and inference protocols

#### 4.4.1 Cross-domain supervised transfer and label-efficiency analysis

Cell-type labels were harmonized through Cell Ontology identifiers. The data-rich, closed-set analysis comprised 179 fine-label-eligible reference-to-query directions spanning five operational shift strata: cross-dataset, cross-donor, cross-chemistry, cross-platform and cross-tissue. Models were fitted using up to 200 labelled reference cells per class and evaluated on the complete held-out query; query labels were used only for evaluation. The primary frozen-embedding readout was unweighted multi-nomial logistic regression after source-fitted feature scaling. CellTypist and SingleR used their native prediction procedures, whereas scANVI was treated as transductive because it used unlabelled query expression but not query labels.

Label efficiency and model-specific supervised adaptation used the same panel of 44 directed TSv1–TSv2 transfers. Shared labels were required to contain at least 10 cells in both domains. For label efficiency, at each *N* ∈ {1, 5, 10, 50, 100, 200, 500}, up to *N* labelled reference cells per class were sampled ten times at each budget, and predictions were evaluated on the complete shared-label query at its natural class frequencies. Reference-scarce types were defined as shared types representing less than 1% of the source cells before label-budget sampling; if no type met this threshold, the least abundant shared type was used.

Thirty-seven of the 44 directions contained both source-represented and source-absent query types and were eligible for open-set detection. Detection performance was summarized by direction-level AUROC; detector definitions, sensitivity readouts and ontology-based error categories are described in the Supplementary Methods.

#### 4.4.2 Model-specific supervised adaptation

For supervised adaptation, we followed the procedure provided by the original authors whenever a complete training script or tutorial was available. The executed procedures included full-backbone updating, partial-backbone updating and supervised heads trained on fixed embeddings. When a released procedure was incomplete, we retained the intended trainable scope and completed the training and inference steps. When no supervised-adaptation procedure was provided, the pretrained representation was kept fixed and a shared two-layer multilayer-perceptron classifier was trained with cross-entropy loss. The shared classifier consisted of layer normalization, dropout, a hidden linear layer with GELU activation and an output layer matching the number of classes.

For each transfer direction, validation cells were drawn only from the labelled reference and stratified by cell type; query labels were not used for training, checkpoint selection or hyperparameter choice. Model-specific optimization and trainable-parameter settings are reported in the Supplementary Information. Adapted predictions and the corresponding frozen-embedding logistic-regression predictions were evaluated on the same complete query, including source-absent types.

For the HVGs+PCA reference, 2,000 highly variable genes were selected using the source, followed by library-size normalization, log-transformation, scaling and PCA to 50 dimensions. The query was projected using the source-fitted transformation and evaluated with the same source-trained readouts.

#### 4.4.3 Hardware and implementation

All experiments were implemented in Python using PyTorch and the HuggingFace Transformers ecosystem, together with Scanpy and other widely used single-cell analysis libraries. Model training, embedding extraction and downstream evaluations were executed on a high-performance computing cluster equipped primarily with NVIDIA H200 GPUs (140 GB memory). To assess hardware portability, all scFMs embedding pipelines were additionally validated on NVIDIA H100 GPUs and RTX 5000 Ada GPUs with a minimum of 32 GB GPU memory.

All jobs were submitted through the Slurm workload manager with standardized resource allocation, including 256 GB system memory and four CPU cores per task, unless otherwise specified. This configuration ensured consistent execution across models and datasets while accommodating the heterogeneous computational requirements of different methods.

To facilitate reproducibility and scalability, we automated all embedding, training and evaluation workflows using Nextflow. Software versions and execution parameters were explicitly recorded, and all model environments were encapsulated in Docker or Singularity images. This design enables deterministic execution across computing platforms and allows the full benchmark to be reproduced with a single command.

#### 4.4.4 Runtime and memory measurements

Inference resources were profiled on a standardized input containing 10,000 cells and 30,000 genes using an NVIDIA H200 GPU, 256 GB host memory and four CPU cores. Peak resident set size and end-to-end wall time were recorded for all methods; peak allocated and reserved device memory were additionally recorded for GPU-enabled methods over supported batch sizes. Batch-size and cell-count settings each used five replicate input matrices. Unsupported configurations and device-memory measurements for CPU-only methods were treated as not applicable. Timer boundaries, precision settings and complete profiles are provided in the Supplementary Information.

### 4.5 Embedding-geometry analysis

Geometry was evaluated independently for each dataset–representation pair using the cell embedding output by each method. To bound computation, we retained all cells in datasets containing at most 20,000 cells and otherwise sampled 20,000 cells without replacement. The sampled cells were fixed within each dataset and shared across methods.

Let *Z*R*^n^*^×^*^d^* denote the uncentred embedding of one dataset. We defined spectral anisotropy as the fraction of total embedding energy captured by its leading singular direction,

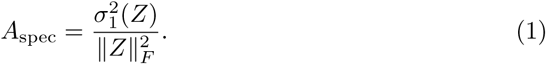

The leading singular value was estimated by randomized singular-value decomposition. Values approaching one indicate that cell vectors share a dominant global direction, consistent with cone-like collapse.

We next distinguished non-specific directional concentration from cell-type-specific organization. For cells *i* and *j*, cosine similarity was computed from their output vectors. Within-cell-type and between-cell-type anisotropy were

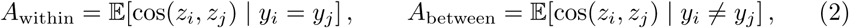

and the cell-type anisotropy contrast was Δ*A* = *A*_within_ *A*_between_. For each dataset, 100,000 pairs were sampled with replacement for each condition. Cell types, rather than cells, were sampled uniformly before drawing cells within types, preventing abundant populations from dominating the estimate. A larger Δ*A* indicates that directional coherence is specific to shared biological identity rather than imposed globally on all cells.

Local expression-neighbourhood preservation was quantified relative to an analytic-Pearson-residual gene-expression reference constructed without dimensionality reduction or cell-type labels. Starting from raw UMI counts, mitochondrial, ERCC and ribosomal-protein genes were excluded, as were genes detected in fewer than 10 cells or having fewer than 10 total counts. Within each batch, expected counts were *µ_ig_* = *s_i_p_g_*, where *s_i_* is the library size of cell *i* and *p_g_* is the batch-specific proportion of counts assigned to gene *g*. Analytic Pearson residuals were

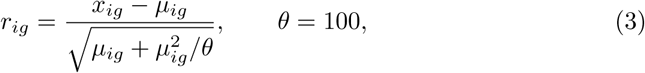

and were clipped to ± √*nb* for a batch of *n_b_* cells. Residual variance was converted to a percentile within each eligible batch, and the 2,000 genes with the highest median percentile across batches were retained. No cell-type labels entered reference construction.

Reference neighbours were computed by Euclidean distance directly in the unscaled residual gene-expression space, without dimensionality reduction. Candidate neighbours were computed by the same distance from the unstandardized coordinates of each representation. Both graphs were constructed separately within each batch, preventing between-batch structure from determining preservation; batches with fewer than 100 cells were excluded. Exact nearest neighbours were used for at most 5,000 cells and approximation (by PyNNDescent) otherwise. For neighbourhood size *k*, mean overlap and its chance-corrected form were

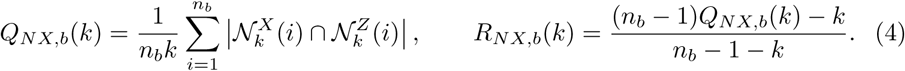

Thus, *R_NX_* = 1 denotes identical neighbourhoods and *R_NX_*= 0 the overlap expected at random. Batch estimates were weighted by their eligible cell counts, and *R_NX_* was then averaged equally over *k* = 15, 30 and 50 to give one value per tissue–representation pair.

Secondary diagnostics were used to identify distinct geometric failure modes. After centring and scaling each coordinate to unit population variance, the participation ratio was

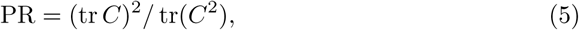

where *C* is the embedding covariance matrix, and normalized PR was PR*/d*. TwoNN intrinsic dimension was estimated in both unstandardized and coordinate-standardized spaces as

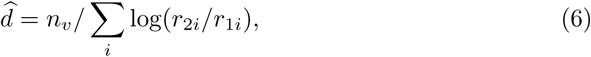

using cells with finite 0 *< r*_1*i*_ *< r*_2*i*_. These quantities were treated as descriptive indicators of dimensional collapse or high-dimensional but weakly organized structure, rather than as monotonic measures of representation quality.

To quantify label-associated structure, each standardized embedding coordinate was fitted with the additive model *z* ∼ 1 + cell type + batch. For cell type or batch *E*, the effect sum of squares was SS*_E_* = SSE_−_*_E_* − SSE*_F_*, where *F* denotes the full model and −*E* the reduced model omitting that effect. Partial eta-squared was

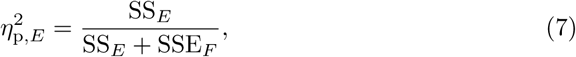

and was averaged equally across estimable embedding coordinates. Cell-type partial *η*^2^ therefore measures cell-type-associated structure conditional on batch, and batch partial *η*^2^ measures batch-associated structure conditional on cell type. Batch effects were unavailable for single-batch tissues. The two effect sizes have different denominators and were reported separately rather than interpreted as additive variance fractions.

For performance associations, each geometric statistic and performance outcome was first averaged equally across the 26 tissues for each representation. The anisotropy analyses used the complete biological-conservation family-rank aggregate. To avoid part–whole overlap, only the primary local expression-neighbourhood preservation analysis excluded Acc@kNN, cLISI and graph connectivity from its performance outcome. Performance was the negative of this aggregate rank, so that larger values indicate better performance. Spearman correlations were calculated across all 20 representations; the neighbourhood-preservation association was also repeated after restriction to the 13 frozen scFMs. The complete biological-conservation aggregate was retained as a sensitivity outcome for that association. Reported *P* values were two-sided, nominal and not multiplicity-adjusted. These model-level correlations were interpreted as diagnostic associations between observed geometry and performance, not as evidence that altering one statistic would causally improve an embedding.

### 4.6 Evaluation metrics

#### 4.6.1 Zero-shot clustering and embedding evaluation

To evaluate zero-shot representation quality, we assessed whether model-derived embeddings preserve biologically meaningful structure without task-specific training. For each dataset, embeddings were computed using pretrained models and clustered using a unified Leiden-based clustering procedure. A shared nearest-neighbour graph was constructed from each embedding, and Leiden clustering was evaluated over a common resolution grid. The selected resolution minimized the absolute difference between the number of Leiden clusters and the number of reference cell types, with normalized mutual information (NMI) used to break ties. The same procedure was applied to all embeddings. Clustering performance and embedding quality were quantified using complementary metrics capturing cluster concordance, continuous cell-type separation, local neighbourhood structure and batch mixing.

We evaluated agreement between predicted cluster labels and reference cell-type annotations using multiple label-based metrics, including the adjusted Rand index (ARI), normalized mutual information (NMI), homogeneity (HOM), completeness (COM), and the Fowlkes–Mallows index (FMI). These metrics quantify different aspects of clustering quality, including overall label agreement (ARI), mutual dependence between predicted and true labels (NMI), cluster purity (HOM), cluster completeness (COM), and pairwise sample agreement (FMI). Higher values indicate better correspondence between clustering results and biological annotations.

To assess continuous cell-type separation in the embedding space independently of clustering assignments, we computed the average silhouette width (ASW) using reference cell-type labels. Higher ASW values indicate greater within-cell-type compactness and between-cell-type separation. However, ASW favors compact, approximately spherical and well-separated groups and may therefore provide an unreliable assessment of biologically valid embeddings with different geometries [39]. We consequently treated ASW as a complementary rather than primary evaluation metric and assigned it a lower weight in the composite performance score.

To quantify local label consistency in the embedding space, we computed k-nearest-neighbor classifier accuracy (Acc@kNN). Specifically, a k-nearest-neighbor (kNN) classifier was trained to predict ground-truth cell-type labels based solely on the learned embeddings. Accuracy was estimated using stratified 5-fold cross-validation to ensure robustness across label distributions. Acc@kNN reflects how well local neighborhoods in the embedding space preserve biological identity, with higher values indicating stronger local discriminative structure.

We further evaluated local biological neighborhood purity using the cell-type local inverse Simpson’s index (cLISI), which quantifies the diversity of cell-type labels within local neighborhoods in the embedding space. Higher cLISI values indicate greater enrichment of the same cell type within local neighbourhoods. Graph connectivity (GC) was calculated on a kNN graph and measures the extent to which cells sharing a reference biological label remain connected. Higher GC indicates stronger preservation of biological population connectivity.

Batch mixing was evaluated only for datasets containing at least two batches. We used k-nearest-neighbour batch effect test (kBET), batch removal adapted silhouette (BRAS), and cell-type-conditioned integration LISI (CiLISI) as the three batch-mixing metrics included in the aggregate benchmark. kBET evaluates whether local batch compositions are consistent with the corresponding global batch distribution, with higher acceptance rates indicating better mixing. BRAS was calculated using the revised formulation designed to address limitations of the conventional batch silhouette. It was evaluated within reference cell types using cosine distance and the mean distance to cells in all other batches. Higher BRAS indicates better mixing of batches within biological cell types. CiLISI was computed by evaluating batch-label diversity separately within each reference cell type and then aggregating the normalized cell-level values. CiLISI was scaled to [0, 1], with higher values indicating stronger batch diversity within biologically matched neighbourhoods.

For spatially resolved transcriptomics (SRT) datasets, we additionally evaluated spatial coherence and domain continuity using Moran’s I, Geary’s C, percentage of abnormal spots (PAS), and CHAOS. Moran’s I and Geary’s C measure global and local spatial autocorrelation of domain-specific marker gene expression, respectively. PAS quantifies local spatial label inconsistency based on majority voting among spatial neighbours, whereas CHAOS measures within-domain spatial compactness. Except for Moran’s I, which is directly maximized, the remaining spatial metrics were transformed using 1 metric so that higher values consistently indicate better spatial domain identification performance.

All biological-conservation metrics were calculated on the full set of embedded cells. Batch-aware metrics were computed using the complete data for each multi-batch dataset and were recorded as not applicable for single-batch datasets. Rather than averaging raw metric values with different interpretations and correlation structures, methods were ranked separately within each dataset and metric. For the primary cell-representation analysis, metric ranks were first averaged within three biological families: cluster concordance (ARI, NMI, HOM, COM and FMI), continuous cell-type separation (ASW), and local cell-type neighbourhood structure (Acc@kNN, cLISI and GC). The corresponding family weights were 0.325, 0.10 and 0.325 and were renormalized over the biological total. Batch mixing was summarized separately by equally averaging ranks for kBET, BRAS and CiLISI in eligible multi-batch datasets. Lower aggregate ranks indicate better performance.

Across-dataset statistical inference treated datasets, rather than individual cells, as the independent experimental units. Overall method differences were assessed using Friedman tests, and pairwise comparisons with PCA were performed using two-sided paired Wilcoxon signed-rank tests with multiplicity correction. Uncertainty in mean aggregate ranks and paired rank differences was quantified using 10,000 dataset-level bootstrap resamples. In complementary metric-level analyses, each method was compared with the corresponding PCA score from the same dataset and metric; these paired differences were visualized as boxplots and tested against zero using two-sided paired Wilcoxon signed-rank tests with Benjamini–Hochberg correction across the method-by-metric comparisons.

##### Adjusted Rand Index (ARI)

Given two partitions of *n* cells, the reference labels *U* = {*U*_1_*, …, U_r_*} and predicted clusters *V* = {*V*_1_*, …, V_s_*}, the adjusted Rand index is defined as

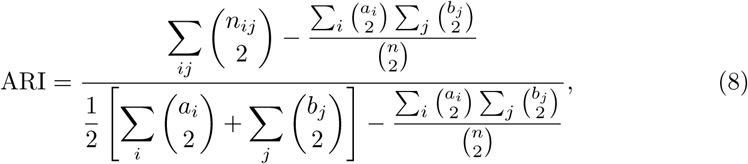

where *n_ij_* = |*U_i_* ∩ *V_j_*|, *a_i_* = |*U_i_*|, and *b_j_* = |*V_j_*|. ARI corrects for chance agreement and ranges from −1 to 1, with higher values indicating better agreement.

##### Normalized Mutual Information (NMI)

The normalized mutual information between partitions *U* and *V* is defined as

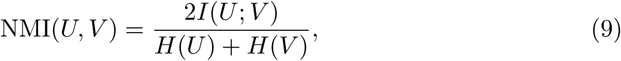

where *I*(*U*; *V*) is the mutual information and *H*(*U*), *H*(*V*) are the entropies of the two partitions. NMI ranges from 0 to 1, with higher values indicating stronger agreement.

##### Homogeneity (HOM) and Completeness (COM)

Homogeneity measures whether each predicted cluster contains cells from a single class,

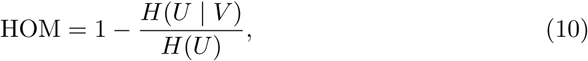

while completeness measures whether all cells of a given class are assigned to the same cluster,

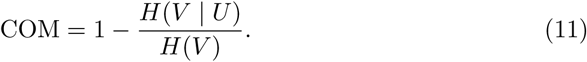

Both metrics range from 0 to 1, with higher values indicating better clustering quality.

##### Fowlkes–Mallows Index (FMI)

The Fowlkes–Mallows index is defined as

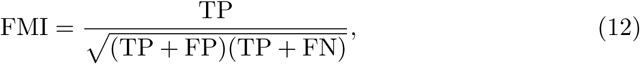

where TP, FP and FN denote the numbers of true-positive, false-positive and false-negative cell pairs, respectively. FMI ranges from 0 to 1, with higher values indicating better agreement.

##### Average Silhouette Width (ASW)

For each cell *i*, the silhouette score is defined as

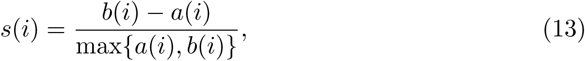

where *a*(*i*) is the average distance between *i* and all other cells of the same cell type, and *b*(*i*) is the minimum average distance between *i* and cells of a different cell type. ASW is the mean of *s*(*i*) across all cells and ranges from 1 to 1, with higher values indicating better separation of cell types.

##### kNN Classifier Accuracy (Acc@kNN)

The kNN classifier accuracy evaluates the discriminative power of learned embeddings by measuring how well a simple k-nearest-neighbor classifier can recover the ground-truth labels. Given an embedding matrix *X* ∈ R*^n^*^×^*^d^* for *n* cells and corresponding true labels 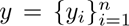, a kNN classifier assigns each sample to the majority label among its *k* nearest neighbors in the embedding space under a chosen distance metric (Euclidean distance by default).

To obtain a robust estimate, we perform stratified *K*-fold cross-validation. Specifically, the dataset is split into *K* = 5 folds with preserved label proportions. For each fold, a kNN classifier is trained on the remaining folds and evaluated on the held-out fold. The final accuracy is computed as

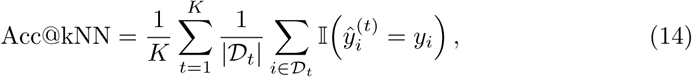

where D*_t_* denotes the test set of the *t*-th fold, *y_i_* is the ground-truth label, and 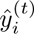 is the predicted label from the kNN classifier trained without *_t_*.

Acc@kNN ranges from 0 to 1, with higher values indicating that the learned embeddings better preserve class structure and are more discriminative with respect to the true labels.

##### Cell-type Local Inverse Simpson’s Index (cLISI)

For cell *i*, the unscaled local inverse Simpson’s index is

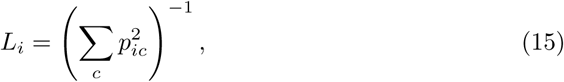

where *p_ic_* denotes the neighbourhood probability assigned to reference cell type *c*. Let *C* be the number of reference cell types and let 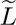 denote the median cell-level LISI value. We report the scaled cLISI score

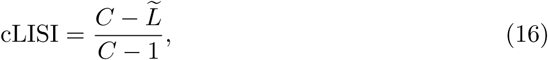

which lies in [0, 1] and is oriented so that higher values indicate stronger local biological purity.

##### Graph Connectivity (GC)

Graph connectivity measures the fraction of cells from each reference biological label contained in its largest connected component of the embedding kNN graph,

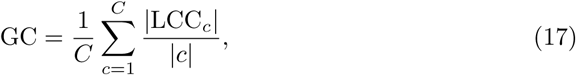

where LCC*_c_* is the largest connected component among cells carrying label *c*. GC ranges from 0 to 1, with higher values indicating stronger preservation of biological connectivity.

##### Batch Removal Adapted Silhouette (BRAS)

BRAS assesses residual batch separation within each reference cell type using a modified silhouette construction. For cell *i*, let *a_i_* denote its mean cosine distance to cells from the same batch and cell type, and let *b_i_^*^* denote its mean cosine distance to cells from all other batches within the same cell type. The modified silhouette value is

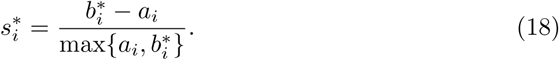

For reference cell type *c*, BRAS is calculated as

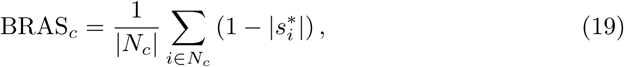

where *N_c_* is the set of cells annotated as cell type *c*. The final score is the unweighted mean across the *C* eligible cell types,

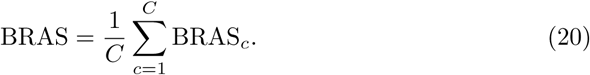

Unlike conventional batch ASW, this definition uses the mean distance to all alternative batches rather than only the nearest one and is therefore less susceptible to the nearest-cluster problem. BRAS ranges from 0 to 1, with higher values indicating better batch mixing while conditioning on biological identity.

##### Cell-type-conditioned Integration LISI (CiLISI)

Within each eligible reference cell type, batch-label LISI was calculated as

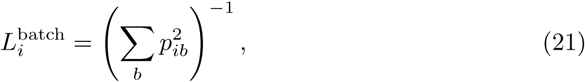

where *p_ib_* is the local probability assigned to batch *b*. For a cell type represented across *B* batches, the value was normalized as

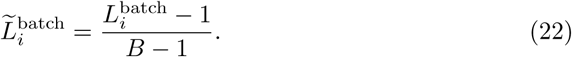

Normalized values were restricted to [0, 1] and averaged over eligible cells. Higher CiLISI indicates stronger mixing of batches within the same biological population.

##### k-nearest Neighbor Batch Effect Test (kBET)

kBET evaluates whether local batch label distributions deviate from the global batch distribution using a chi-square test. The kBET score is defined as the fraction of cells for which the null hypothesis of batch mixing is not rejected,

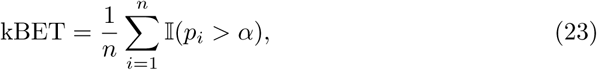

where *p_i_* is the *p*-value of the test for cell *i*, and *α* is the significance threshold. kBET ranges from 0 to 1, with higher values indicating better batch mixing.

##### Moran’s I

Moran’s I measures spatial autocorrelation of a variable over a spatial neighbor graph. Let *x_i_* denote the expression value of a gene at spot/cell *i*, *x* be the mean of 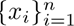, and *w_ij_* be the spatial weight between *i* and *j* derived from the spatial neighbor graph. Moran’s I is defined as

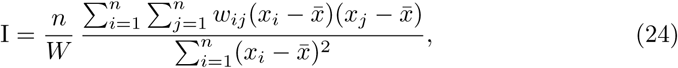

where *W* = ∑_*i*_∑_*j*_ *w_ij_*. Moran’s I typically ranges from 1 to 1: positive values indicate that similar values cluster together in space, while negative values indicate spatial dispersion. In our implementation, we normalize and log-transform the raw counts, select marker genes by ranking genes for each predicted domain, and compute Moran’s I for the selected genes on a spatial neighbor graph. The final score is the median Moran’s I across the selected genes, where a higher value indicates better spatial domain clustering.

##### Geary’s C

Geary’s C is a spatial autocorrelation statistic that emphasizes local differences. Using the same notation as above, Geary’s C is defined as

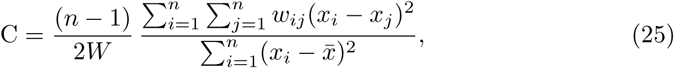

where *W* = ∑_*i*_∑_*j*_ *w_ij_*. Geary’s C typically lies in [0, 2], with smaller values indicating stronger positive spatial autocorrelation. In our experiments, Geary’s C is computed for selected marker genes and summarized by the median across genes. Since a lower Geary’s C corresponds to better spatial clustering, we report the transformed score 1 − C in the final evaluation.

##### CHAOS Score (CHAOS)

CHAOS evaluates the spatial continuity of identified domains by measuring within-domain compactness. Let **s***_i_* ∈ R*^d^* denote the standardized spatial coordinate of spot *i*, and let C(*i*) be its predicted cluster label. For each cluster *k*, consider the set *S_k_* = *i*:C(*i*) = *k*. For a spot *i*∈*S_k_*, define the distance to its nearest neighbor within the same cluster as

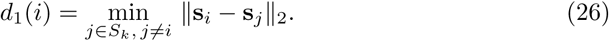

The CHAOS score is computed as

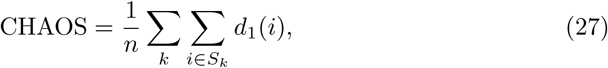

where *n* is the total number of spots. A lower CHAOS indicates better spatial continuity; therefore, we use the transformed score 1 CHAOS in the final evaluation so that larger values indicate better performance.

##### Percentage of Abnormal Spots (PAS)

PAS measures local inconsistency of predicted labels based on spatial neighborhoods. Let N*_k_*(*i*) denote the set of *k* nearest spatial neighbors of spot *i* (excluding itself), and let *y_i_*be the predicted label. A spot is defined as abnormal if more than half of its neighbors have different labels:

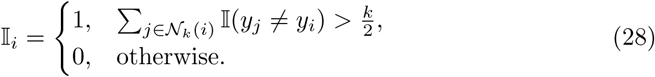

The PAS metric is defined as

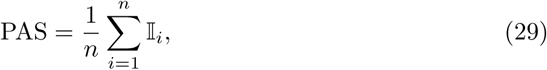

which ranges from 0 to 1. A lower PAS indicates more spatially consistent domain assignments; thus, we report 1 PAS as the final score, where larger values indicate better performance.

#### 4.6.2 Zero-shot trajectory inference evaluation

To evaluate whether zero-shot embeddings preserve biologically meaningful continuous structure, we assessed 17 methods across nine developmental systems spanning linear, bifurcating and tree-structured trajectories. Each representation was analysed independently with Slingshot [40], PAGA [41], diffusion pseudotime [42] and Monocle 3 [43] while retaining the same cells, reference trajectory and root information. Performance was evaluated against the known trajectories using the Dynverse framework.

Twelve direction-consistent measures were organized into five families describing cell position, position recoverability, topology, feature relevance and branch or milestone assignment. Methods were ranked within dataset, inference method and metric; ranks were then averaged within metric family, across applicable families and across inference methods. Outcomes unavailable by construction, including topology and branch endpoints for diffusion pseudotime, were treated as not applicable. Paired dataset-level rank differences from PCA were evaluated with exact sign-flip tests and Holm correction across 16 comparisons.

Backend-specific settings, unavailable endpoints and the trajectory geometry analysis are described in the Supplementary Methods.

#### 4.6.3 Supervised transfer, label-efficiency and open-set evaluation

Cell-type transfer was evaluated using complementary classification metrics that capture overall and per-class performance under both natural class imbalance and limited reference labels. The same definitions were used for frozen-embedding transfer, native reference methods and model-specific supervised adaptation.

We quantified overall prediction performance using accuracy, defined as the fraction of correctly predicted labels among all query samples. Accuracy provides a global measure of classification performance but can be biased toward majority classes.

To ensure equal weighting of all classes regardless of class frequency, we computed macro-averaged precision, recall and F1 score. Per-class precision, recall and F1 were computed independently and then averaged across classes. Macro recall is equivalent to balanced accuracy. Recall for reference-scarce types was averaged over the source-defined scarce types eligible in each transfer direction.

We further evaluated probabilistic predictions using the area under the receiver operating characteristic curve (ROC–AUC). For multiclass classification, ROC–AUC was computed using class probability outputs and aggregated across classes following the implementation in scikit-learn. Higher ROC–AUC values indicate better separation between true and false class assignments.

To assess whether correct labels were ranked among the most likely predictions, we computed top-*k* accuracy, defined as the proportion of samples for which the true label appeared within the top *k* predicted classes ranked by model confidence. Unless otherwise specified, we report top-3 accuracy. This metric is particularly relevant for evaluating embedding quality in ambiguous or closely related cell populations.

For label-efficiency analyses, metrics were computed on the complete shared-lable query and summarized across ten source-cell samples at each label budget. Model-specific supervised adaptation and the corresponding frozen readout were evaluated on the same complete held-out query. For open-set evaluation, source-absent query types were treated as the positive class and detection performance was summarized by direction-level AUROC.

##### Accuracy

Let *y_i_*{1*, …, C*} denote the ground-truth label of sample *i*, and *y*^*_i_* the predicted label. Classification accuracy is defined as

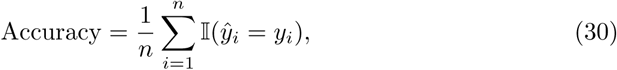

where I(·) is the indicator function and *n* is the number of samples.

##### Macro-averaged precision, recall and F1 score

For each class *c*, precision and recall are defined as

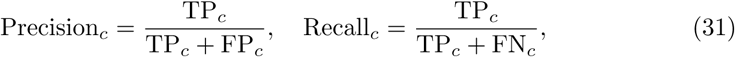

where TP*_c_*, FP*_c_* and FN*_c_* denote the numbers of true positives, false positives and false negatives for class *c*, respectively. Macro-averaged precision and recall are computed as

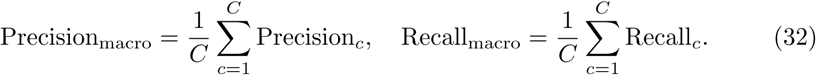

The macro-averaged F1 score is the arithmetic mean of the class-specific F1 scores:

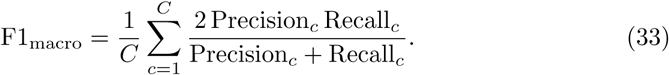

##### Receiver operating characteristic area under the curve (ROC–AUC)

Let *s_ic_* denote the predicted probability score for sample *i* belonging to class *c*. ROC–AUC measures the probability that a randomly chosen positive sample is assigned a higher score than a randomly chosen negative sample. For multiclass classification, ROC–AUC was computed using class probability outputs and aggregated across classes following the implementation in scikit-learn.

##### Top-k accuracy

Let R*_k_*(*i*) denote the set of *k* classes with the highest predicted scores for sample *i*. Top-*k* accuracy is defined as

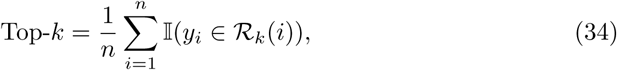

which measures the fraction of samples for which the true label is ranked among the top *k* predicted classes.

#### 4.6.4 Gene-level functional and regulatory evaluation

Static gene embeddings, contextual gene representations and model-native regulatory outputs were evaluated separately because they represent distinct forms of gene-level information. Broad functional organization was assessed using AUROC and AUPRC for recovery of STRING physical interactions and mean average precision for retrieval of shared Gene Ontology Biological Process, Reactome and KEGG annotations. Context-specific regulatory-edge recovery was evaluated on the BEELINE datasets using the AUPRC ratio, defined as observed AUPRC divided by positive-edge prevalence, and the early-precision ratio relative to random ranking. For display, ratios were mapped as *r/*(1 + *r*), whereas rankings used the untransformed values.

Recorded regulon recovery was summarized by per-transcription-factor AUROC for TRRUST, ChEA and DoRothEA ABC. Perturbation-response retrieval used the Norman CRISPRa and Replogle K562 and RPE1 CRISPRi datasets. For each eligible transcription-factor perturbation, the 100 genes with the largest absolute mean-expression response relative to control were treated as positives, and retrieval was evaluated using AUROC and AUPRC divided by positive prevalence. Contextual representations were computed from control cells only. Gene-identity mapping, candidate-universe construction, resource versions, resampling procedures, study-level aggregation and the separate cell-level double-perturbation analysis are described in the Supplementary Methods.

## 5 Data availability

All 94 transcriptomic datasets analysed in this study are publicly available. These comprised 24 Tabula Sapiens v1 tissue datasets, 26 Tabula Sapiens v2 tissue datasets, nine developmental trajectory datasets, 25 spatial transcriptomic sections, seven BEELINE datasets and three perturbation datasets.

The 50 Tabula Sapiens tissue datasets were derived from the publicly available collection in the CZ CELLxGENE Discover portal (https://cellxgene.cziscience.com/collections/e5f58829-1a66-40b5-a624-9046778e74f5). We partitioned the collection into v1 and v2 according to the ‘published 2022’ field in the observation metadata and processed the data following the official Tabula Sapiens Consortium preprocessing pipeline (https://colab.research.google.com/drive/1Z6Fsv7kEsn6ah1fCFqoXEQfFrJuPIyyQ). To ensure donor independence between releases, all donors present in v1 were excluded from the v2 datasets.

The nine developmental trajectory datasets from the Dynverse collection are available at https://zenodo.org/records/1443566. The spatial data comprised 12 Spa-tialLIBD sections (https://research.libd.org/spatialLIBD/), eight HER2ST sections (https://github.com/almaan/her2st) and five MERFISH hypothalamic preoptic-region sections (https://datadryad.org/dataset/doi:10.5061/dryad.8t8s248).

The seven experimental BEELINE datasets (hESC, hHep, mDC, mESC, mHSC-E, mHSC-GM and mHSC-L) are available from the BEELINE input-data archive (https://doi.org/10.5281/zenodo.3378975). The Norman CRISPRa and Replogle K562 and RPE1 CRISPRi datasets were obtained from the harmonized scPerturb release (https://zenodo.org/records/13350497). Versions and access details for the gene-level annotation resources, together with dataset-specific preparation and eligibility criteria, are provided in the Supplementary Information.

## 6 Code availability

The code used to implement the framework, perform the analyses and generate the results in this study is publicly available and has been deposited on Github at https://github.com/Svvord/scFoundry,under the MIT license.).

---

**Extended Figure 1.**
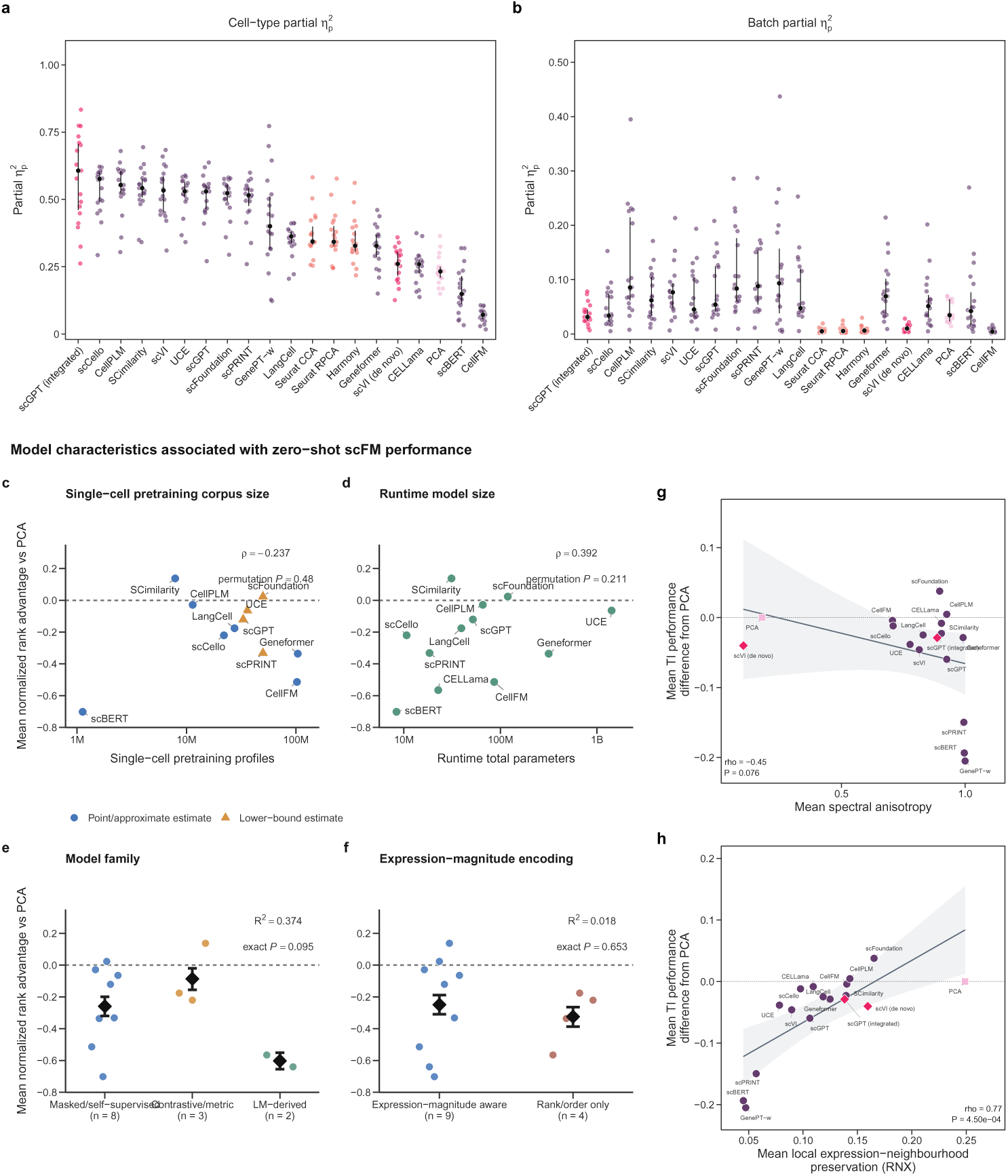
Complementary embedding-geometry diagnostics and model characteristics analysis. **a,b,** Tissue-level distributions of cell-type and batch partial eta-squared, respectively; points denote tissues and black markers and vertical lines denote the median and interquartile range. **c–f**, Model characteristics associated with TSv2 zero-shot biological-conservation performance: single-cell pretraining corpus size, runtime model size, model family and expression-magnitude encoding. **g,h,** Associations of trajectory-inference performance relative to PCA with mean spectral anisotropy and local expression-neighbourhood preservation (*R_NX_*), respectively.

**Extended Figure 2.**
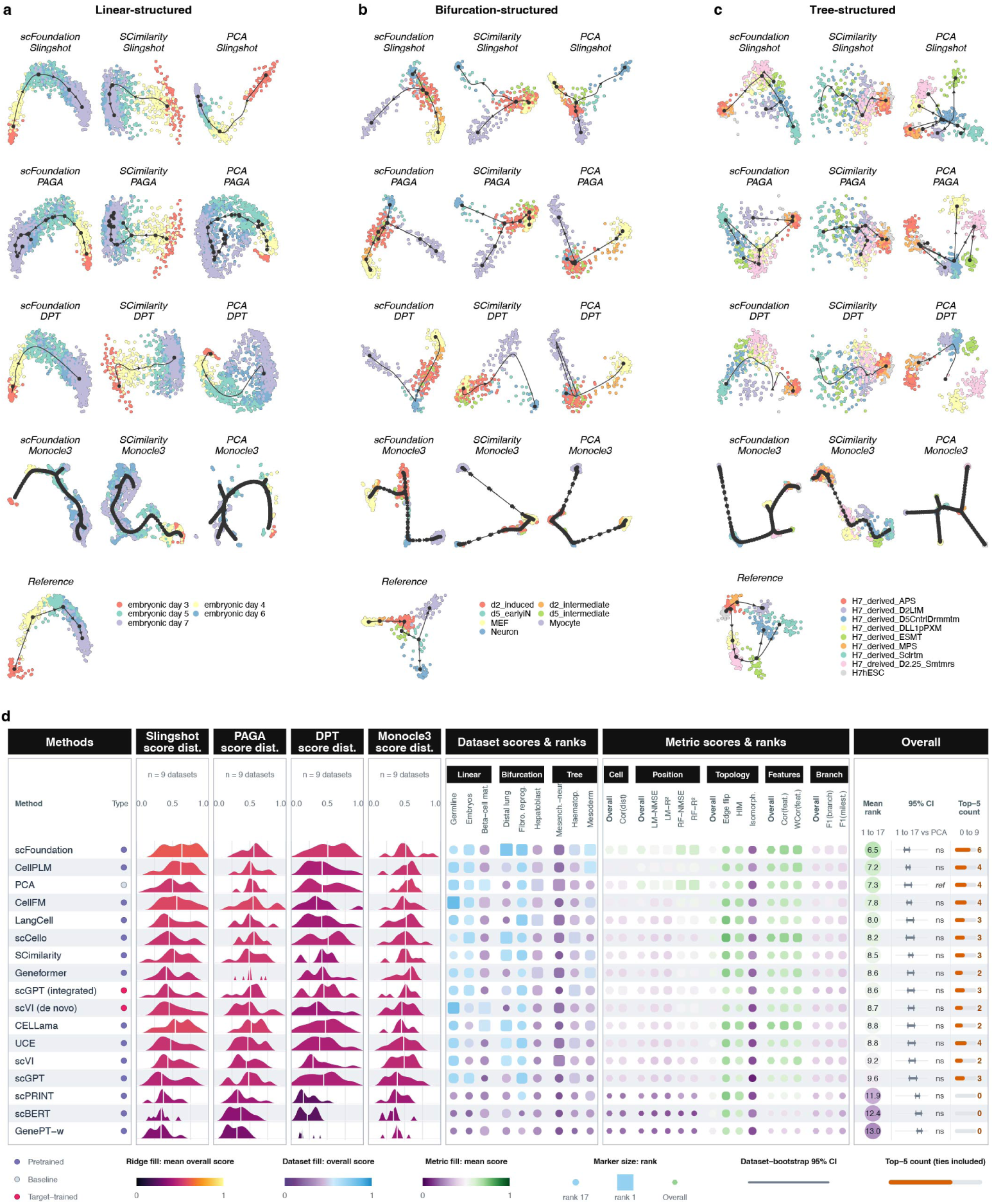
Trajectory inference across developmental structures and inference methods. **a–c,** Selected examples from a linear trajectory (human-embryos petropoulos), a bifurcating trajectory (fibroblast-reprogramming treutlein) and a tree-structured trajectory (mesoderm-development loh). Within each dataset, columns show scFoundation, SCimilarity and PCA, and rows show Slingshot, PAGA, DPT and Monocle 3; cells are coloured by the reference annotation, and black curves and nodes show the inferred trajectory. Layouts were reconstructed independently and are not comparable in orientation or scale. **d,** Aggregate trajectory-inference performance for 17 methods across nine datasets and four inference methods. Ridges show method-level score distributions for each inference method; matrices summarize dataset- and metric-level scores and ranks. The right columns show mean aggregate rank, 95% trajectory-class-stratified bootstrap intervals and the number of datasets ranked in the top five. Symbols indicate paired two-sided exact sign-flip tests against PCA with Holm correction across 16 comparisons.

**Extended Figure 3.**
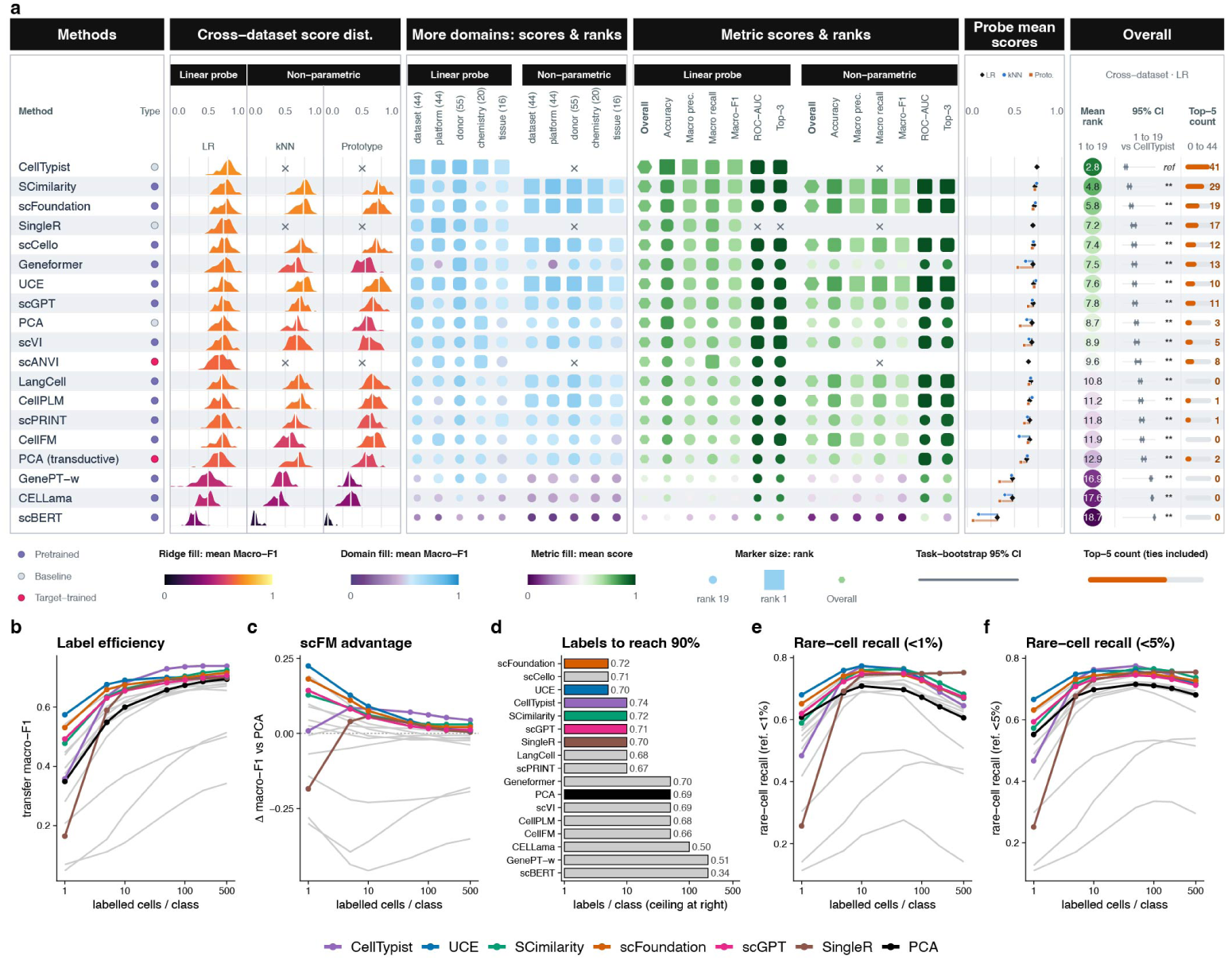
Supervised cell-type transfer across domains in a closed-set setting. **a,** Closed-set transfer benchmark. Classifiers were fitted on a labelled reference domain and evaluated on the full query domain (cell types with ≥ 10 cells in both domains; source capped at 200 cells per class). Frozen scFM, PCA and scVI embeddings were read out by a linear probe, k-NN (*k* = 15) and a nearest-prototype classifier; CellTypist, SingleR and scANVI were run natively. Methods are ordered by overall mean rank on the 44 cross-dataset tasks (linear probe; accuracy and macro precision/recall/F1; error bars, bootstrap 95% CI across tasks). Panels show per-task macro-F1 distributions, mean macro-F1 (fill) and mean rank (text) across the four other domain shifts (*n* = 44, 55, 20 and 16 pairs), and six annotation metrics for the linear-probe and prototype read-outs (SingleR provides no class probabilities). Rank 1 = best. **b–f,** Label efficiency on the same 44 pairs. For each *N* ∈ 1, 5, 10, 50, 100, 200, 500, the linear probe was fitted on up to *N* labelled source cells per class (10 random draws) and the full query predicted; lines, mean over pairs and draws. **b,** Transfer macro-F1. **c,** Macro-F1 advantage over PCA. **d,** Labels per class needed to reach 90% of each method’s own macro-F1 at the maximum budget, *N* = 500. **e,f,** Rare-cell recall: mean recall over reference-scarce types, defined as *<* 1% (**e**) or *<* 5% (**f**) of reference cells. CellTypist and SingleR were trained natively on the same subsampled reference. Non-highlighted methods are shown in grey.

## Acknowledgements

This study was supported by the National Institutes of Health (NIH) grants R01GM126553, R01HG011883, R01HG009124, and R01GM144960 (all to X.Z.).

## Author contributions statement

X.Z. and S.H. designed the project. S.H. led the project, conceived the algorithm, designed the framework, developed the software, performed data analysis. S.H and P.Y. contributed to result visualizations. P.Y., W.M., J.W., and H.W. contributed to formatting the results. J.X. provided input on model interpretation. Y.M. provided guidance on the application and validation of the DRIFT algorithm. X.Z. supervised the project. X.Z. and S.H. wrote and edited the manuscript with input from all authors. All authors read and approved the final manuscript.

