## Supplementary Information for "Accessible and reproducible deployment reveals the practical boundaries of single-cell foundation models"

Siyu Hou et al.

August 31, 2026

**3 Supplementary Tables** **291**

### Supplementary Methods

#### 1.1 Dataset collection and preprocessing for the zero-shot cell-representation benchmark

Single-cell RNA sequencing (scRNA-seq) data for zero-shot benchmarking were obtained from the publicly available Tabula Sapiens collection through the CZ CELLxGENE Discover portal (https://cellxgene.cziscience.com/collections/e5f58829-1a66-40b5-a624-9046778e74f5). Samples were partitioned into two releases, referred to as **v1** and **v2**, based on the `published_2022` field in the observation metadata. The v2 release corresponds to the curated Tabula Sapiens v2 dataset provided by the CZ Virtual Cell Models project, whereas the v1 release was processed using the same workflow to ensure consistency across releases.

Data preprocessing followed the official Tabula Sapiens Consortium pipeline and the processing workflow described by the CZ Virtual Cell Models project. Briefly, gene count matrices were merged with the corresponding cell metadata using `Scanpy`. Cells annotated with ambiguous or uninformative cell-type labels (for example, unassigned, unknown or unclassified) were removed. Genes with zero counts across all cells were excluded, and cells expressing fewer than 10 detected genes within each donor were discarded. Ensembl gene identifiers were standardized by removing version suffixes, normalized expression matrices were replaced with the corresponding raw count matrices, and raw counts were validated before downstream analyses.

To construct an independent benchmark, all donors represented in the v1 release were excluded from the v2 dataset, ensuring that no donor was shared between releases. Following preprocessing and donor filtering, the Tabula Sapiens collection contributed 50 tissue-specific datasets: 24 from v1 and 26 from v2. These datasets span 28 tissues, with 22 represented in both releases, kidney and pancreas unique to v1, and ear, ovary, stomach and testis unique to v2. The complete study analysed 94 transcriptomic datasets: these 50 Tabula Sapiens tissues, nine developmental trajectory datasets, 25 spatial transcriptomic sections, seven BEELINE datasets and three perturbation datasets (Table S12).

We manually audited the publications, supplementary information and released checkpoint documentation to determine whether the Tabula Sapiens releases were included in the pretraining data (Table S1). TSv1 was present in the pretraining corpus of most evaluated scFMs, whereas TSv2 was absent from all evaluated pretraining corpora. TSv2 was therefore used for the primary representation benchmark, and the paired TSv1–TSv2 analysis assessed whether the principal performance hierarchy was robust across atlas releases.

**Table S1: Audit of Tabula Sapiens exposure in the released pretrained models.**

| Model | TSv1 in pretraining | TSv2 in pretraining | Rationale |
|---|---|---|---|
| CELLama | No | No | The default CELLama encoder is a frozen sentence transformer and was not pretrained on any single-cell atlas. |
| CellFM | Yes | No | CellFM metadata include GEO accession GSE201333, corresponding to TSv1; TSv2 samples are not listed in the public pretraining metadata. |
| CellPLM | Yes | No | CellPLM lists Tabula Sapiens in its pretraining data, corresponding to TSv1; TSv2 was released after the reported corpus was assembled. |
| Geneformer | Yes | No | Genecorpus-104M includes the TSv1 DOI/source entry; we found no evidence that TSv2 samples were included. |
| GenePT-w | No | No | GenePT derives gene embeddings from natural-language gene descriptions and has no single-cell-atlas pretraining stage. |
| LangCell | Likely no | No | LangCell uses scLibrary, a CELLxGENE-derived cell–text corpus, but describes TSv1 as a downstream benchmark rather than a pretraining source; TSv2 was released later. |
| scBERT | No | No | scBERT was pretrained on PanglaoDB-derived profiles that predate TSv1. |
| scCello | Yes | No | scCello was trained on CELLxGENE Census release 2023-07-25, which included TSv1; TSv2 corresponds to a later release. |
| scFoundation | Yes | No | scFoundation pretraining metadata include GEO accession GSE201333 for TSv1; TSv2 is not listed among the reported sources. |
| scGPT | Yes | No | The scGPT whole-human checkpoint was trained on CELLxGENE Census release 2023-05-15, which included TSv1; TSv2 was released later. |
| SCimilarity | Yes | No | SCimilarity reports Tabula Sapiens as part of its training data, corresponding to TSv1; TSv2 was not available at the time of training. |
| scPRINT | Yes | No | scPRINT v1 was trained on CELLxGENE Census LTS 2023-12-15, which included TSv1; TSv2 was released later. |
| UCE | Yes | No | The UCE paper states that TSv1 was used for training and TSv2 was held out for evaluation. |

*Note.* TSv1 includes GEO accession GSE201333. TSv2 denotes the evaluation set after removing all donors represented in TSv1. Exposure refers to the publicly released pretrained checkpoints or default model configurations; downstream supervised adaptation and task-specific classifiers are not counted as pretraining.

#### 1.2 Zero-shot cell-representation evaluation in Tabula Sapiens v2

##### *1.2.1 Study design and analysis scope*

We evaluated whether pretrained single-cell foundation models (scFMs) produce cell representations that preserve biological structure without optimization on the evaluation data. The complete benchmark compared 20 representations within every tissue. The principal zero-shot scientific contrast focused on 13 frozen scFM representations, with PCA as the primary expression-based reference and a pretrained scVI representation as a deep-learning non-foundation comparator. For practical context, the same evaluation also included Harmony, Seurat CCA, Seurat RPCA, scVI trained de novo and scGPT fine-tuned for integration. These five target-data methods were applied independently in each tissue and are therefore explicitly treated as practical comparators rather than zero-shot models. PCA was likewise estimated independently in each target tissue, but served as the primary expression-based reference rather than as an integration method.

Throughout this study, *zero-shot* denotes that model parameters were not updated on the target Tabula Sapiens v2 (TSv2) dataset and that neither cell-type annotations nor batch identities were supplied during embedding extraction. Pretraining exposure to TSv1 and TSv2 was determined by the manual source audit reported in Table S1.

Biological-conservation ranks were calculated in this fixed 20-representation universe. Statements specifically concerning zero-shot scFMs were subsequently restricted to the 13 frozen scFMs. Preservation of biological structure was the primary outcome for this analysis. Batch mixing was evaluated as a separate, fully reported supplementary objective because most frozen scFM methods do not implement target-data batch correction and because good apparent mixing can also result from loss of biological structure. We therefore report both the individual batch-mixing metrics and a biological-plus-batch aggregate, while keeping the biological-only aggregate as the basis of the principal zero-shot comparison and the embedding-geometry analyses.

##### *1.2.2 Tabula Sapiens v2 tissue panel*

We analysed 26 tissue-resolved human single-cell RNA-sequencing datasets from the processed TSv2 benchmark collection [1, 2]. The processed release followed CELLxGENE schema version 5.2.0. The tissues were bladder, blood, bone marrow, ear, eye, fat, heart, large intestine, liver, lung, lymph node, mammary gland, muscle, ovary, prostate, salivary gland, skin, small intestine, spleen, stomach, testis, thymus, tongue, trachea, uterus and vasculature. Following upstream release-level curation, the panel contained 548,977 cells. Individual tissues contained 2,044–72,290 cells, 14–32 annotated cell types and 21,808–37,066 retained gene features.

As described in Section 1.1, upstream data curation preceded the local benchmark-preparation step. Here, *data preparation* refers solely to conversion of the processed data into the input format required by our framework, including identifier and metadata harmonization. This step did not modify the expression matrix. Specifically, no additional cell or gene filtering, quality control, library-size normalization, log transformation or highly variable gene selection was performed. The original observation index was retained in `obs["barcode"]`. A usable `feature_name` was used as the gene symbol; otherwise, the original feature index or Ensembl identifier was used as a fallback. Ensembl identifiers were retained from `ensembl_id`, or from `feature_id` when the former was unavailable. A file was rejected if the derived feature identifiers were not unique. The auxiliary AnnData `raw` slot was removed so that downstream analyses used `X`; this did not modify the values stored in `X`.

The curated `cell_type` field was used as the biological reference annotation. We defined a batch as a unique assay–donor combination,

$$\texttt{batch\_id} = \texttt{assay} \mid \texttt{donor\_id}, \tag{S1}$$

so that profiles obtained using different assays from the same donor were not treated as one technical batch. Batch count ranged from one to seven per tissue. Nineteen tissues contained multiple batches; eye, lung, mammary gland, prostate, skin, testis and uterus contained one batch and were therefore not informative for batch-mixing metrics. Nineteen tissues consisted exclusively of annotated 10x Genomics 3′ v3 profiles. The remaining seven were dominated by that assay but also included 10x Genomics 5′ v2 or Smart-seq2/3 profiles.

###### *1.2.3 Frozen representations and reference methods*

Each output was stored as the matrix of a new AnnData object, with the original cell metadata restored and rows retained or realigned by barcode. We used a prespecified deployment-matched representation for each model: the official embedding interface where available and an explicitly documented benchmark-defined extraction rule otherwise. The benchmark used each method's native representation directly; no common post hoc centring, feature-wise scaling or dimensionality reduction was applied.

**Frozen scFM embedding extraction.** We extracted one embedding per cell from CELLama, CellFM, CellPLM, Geneformer, GenePT-w, LangCell, SCimilarity, UCE, scBERT, scCello, scFoundation, scGPT and scPRINT. Model-specific preprocessing followed the input encoding expected by each released implementation. No reference cell-type label or TSv2 batch identity was supplied during frozen inference.

CELLama represented the 30 most highly expressed genes in each cell using its all-MiniLM-L6-v2 sentence encoder, without adding observation metadata to the input sentence. CellFM used the CellFM 80M checkpoint. Per-cell total counts were capped at $10^5$ and rounded; cells with more than 2,048 non-zero genes were expression-weighted sampled to the supported sequence length. Masking was disabled, and the 1,536-dimensional cell representation was calculated as the mean of the encoded non-zero gene-token states after excluding the first three special tokens. A constant placeholder cell type and batch were provided only to satisfy the released data-loader interface; the original labels were removed before inference. CellPLM indexed genes by Ensembl identifier and used the released `CellEmbeddingPipeline.predict` interface with checkpoint `20231027_85M.best.ckpt`.

Geneformer used the Geneformer-V2-316M checkpoint and the released rank-based tokenizer. Per-cell count totals were supplied to the tokenizer and the V2 CLS state was retained. Barcode completeness and output order were verified before the embedding was joined back to the original metadata. GenePT-w first normalized each cell to $10^4$ counts and applied $\log(1 + x)$. It then formed an expression-weighted combination of fixed text-embedding-ada-002-derived gene vectors. Genes absent from the released lookup received a zero vector; the weighted sum was divided by the number of input genes. LangCell applied its rank-based cell tokenizer, extracted the CLS state of the released cell BERT and applied the learned 256-dimensional projection followed by L2 normalization.

For scBERT, genes were aligned and zero-padded to the 16,906-gene Panglao reference. Counts were

**Table S2: Cell representations included in the Tabula Sapiens v2 analysis.** Target-data fitting indicates whether the representation was estimated or updated separately within each evaluation tissue.

| Representation | Analysis role | Checkpoint or implementation | Dimensions | Target-data estimation | Batch used |
|---|---|---|---|---|---|
| CELLama | frozen scFM | all-MiniLM-L6-v2 | 384 | no | no |
| CellFM | frozen scFM | CellFM 80M | 1,536 | no | no |
| CellPLM | frozen scFM | 20231027_85M.best.ckpt | 512 | no | no |
| Geneformer | frozen scFM | Geneformer-V2-316M | 1,152 | no | no |
| GenePT-w | frozen scFM (LM-derived) | text-embedding-ada-002 | 1,536 | no | no |
| LangCell | frozen scFM | released cell encoder | 256 | no | no |
| SCimilarity | frozen scFM | model v1.1 | 128 | no | no |
| UCE | frozen scFM | 33l_8ep_1024t_1280 | 1,280 | no | no |
| scBERT | frozen scFM | Panglao checkpoint | 200 | no | no |
| scCello | frozen scFM | scCello-zeroshot | 256 | no | no |
| scFoundation | frozen scFM | 01B-resolution | 3,072 | no | no |
| scGPT | frozen scFM | scGPT_human | 512 | no | no |
| scPRINT | frozen scFM | v2-medium | 256 | no | no |
| pretrained scVI | pretrained non-foundation reference | Census 2024-07-01 | 50 | no | no |
| PCA | expression reference | per-tissue HVG PCA | 50 | yes | no |
| Harmony | integration reference | Seurat HarmonyIntegration | 30 | yes | if > 1 batch |
| Seurat CCA | integration reference | Seurat CCAIntegration | 30 | yes | if > 1 batch |
| Seurat RPCA | integration reference | Seurat RPCAIntegration | 30 | yes | if > 1 batch |
| scVI (de novo) | target-trained reference | random initialization | 30 | yes | if > 1 batch |
| scGPT (integrated) | target-fine-tuned scFM | scGPT integration | 512 | yes | if > 1 batch |

**Table S3: Design taxonomy of the evaluated single-cell foundation models.**

| Model | Family | Input encoding | Training objective | Backbone |
|---|---|---|---|---|
| Geneformer | Masked/self-supervised | Gene rank | Masked gene modelling | BERT encoder |
| scGPT | Masked/self-supervised | Quantile-binned value (51) | Masked value modelling | Transformer |
| scBERT | Masked/self-supervised | Value-binned (7) + gene2vec | Masked value modelling | Performer |
| scFoundation | Masked/self-supervised | Continuous value + read depth | Masked value regression | xTrimoGene encoder–decoder |
| CellFM | Masked/self-supervised | Continuous value | Masked value regression | ERetNet |
| CellPLM | Masked/self-supervised | Continuous value (gene-embedding sum) | Masked reconstruction (GM-VAE) | Flowformer |
| scPRINT | Masked/self-supervised | Continuous value + ESM2 identity | Expression denoising | Transformer |
| UCE | Masked/self-supervised | ESM2 gene tokens; expression-weighted sampling | Masked binary expression | Transformer |
| scCello | Contrastive/metric (+ masked) | Gene rank | Masked gene + ontology contrastive | BERT encoder |
| LangCell | Contrastive/metric (+ masked) | Gene rank | Cell–text contrastive | Dual BERT |
| SCimilarity | Contrastive/metric | Continuous value vector | Triplet metric learning + reconstruction | MLP autoencoder |
| GenePT-w | Language-model-derived | Gene-description text | Frozen language-model reuse | GPT-3.5 embedding |
| CELLama | Language-model-derived | Gene rank as text | Sentence-encoder fine-tuning | MiniLM |

normalized to $10^4$, transformed as $\log_2(1 + x)$, and encoded using the implementation's `bin_num=5` setting with a seven-state input vocabulary (`CLASS=bin_num+2`) and the Panglao pretrained checkpoint. The final sequence-token state was used as the 200-dimensional cell representation. scCello used its released zero-shot transformation and the 256-dimensional representation at the terminal sequence position. scFoundation aligned and zero-padded counts to its 19,264-gene input space and used the released cell-embedding routine with `pool_type=all`, `tgthighres=t4`, `pre_normalized=F`, `version=ce` and the 01B-resolution checkpoint.

Frozen scGPT normalized counts to $10^4$, applied $\log(1 + x)$, and used the official `embed_data` interface with the `scGPT_human` checkpoint. SCimilarity aligned counts to the model gene order, zero-padded absent reference genes when necessary, applied the released log-normalization procedure and extracted the 128-dimensional model embedding. scPRINT used raw counts indexed by Ensembl identifier, the v2-medium checkpoint and its `random expr` embedding mode with a maximum sequence length of 4,000; internal cell and gene filtering was disabled so that the curated cell set was retained. UCE used the released 33-layer checkpoint and official single-AnnData inference procedure, including expression-weighted sampling of 1,024 genes, to obtain the 1,280-dimensional `X_uce` representation.

**PCA and pretrained scVI.** PCA was constructed independently in each tissue. Counts were normalized to a library size of $10^4$, transformed with $\log(1 + x)$, and the 2,000 most variable genes were selected using the Scanpy Seurat-flavour procedure. Values were standardized and capped at an absolute value of 10, after which 50 principal components were computed using the ARPACK solver with random seed 0.

For the pretrained scVI comparator, genes were indexed by Ensembl identifier and projected into the CZI CELLxGENE Census scVI model dated 1 July 2024. Query cells were assigned to the model's `unassigned` batch category. `prepare_query_anndata` and `load_query_data` were used to align the query to the reference model, and the loaded model was used directly to obtain the 50-dimensional latent representation without target-data training.

**Target-data integration references.** Harmony, Seurat CCA, Seurat RPCA, scVI (de novo) and scGPT (integrated) were applied independently to each tissue. Only `batch_id` was used as a covariate; cell-type annotations were withheld throughout model fitting.

Harmony, Seurat CCA and Seurat RPCA were run through Seurat v5.5 `IntegrateLayers`. The count matrix was split into batch-specific layers, followed by `NormalizeData`, selection of 2,000 variable genes, `ScaleData` and a 30-component PCA. Integration then used `HarmonyIntegration`, `CCAIntegration` or `RPCAIntegration`, respectively. The random seed was 0. For CCA and RPCA, insufficient-anchor failures were handled by trying `k.weight` values of 100, 50, 30, 20, 10 and 5 in that order. For a single-batch tissue, the uncorrected 30-dimensional Seurat PCA representation was returned.

scVI (de novo) received raw counts and attempted batch-aware Poisson selection of 2,000 genes; all genes were retained if this selection failed. A two-layer scVI model with a 30-dimensional latent space was initialized and trained separately in each tissue, with `batch_id` registered as the only covariate, random seed 0 and the epoch count selected automatically by scvi-tools.

scGPT (integrated) was adapted from the released batch-integration recipe. The `scGPT_human` checkpoint was fine-tuned separately in each tissue for 15 epochs using 1,200 HVGs, 51 expression bins, 40% random masking, batch size 64 and learning rate $10^{-4}$. Zero-valued genes were retained (`include_zero_gene=True`). The enabled objectives were masked expression modelling, masked value prediction from the cell embedding

(MVC/GEPC), elastic cell similarity (threshold 0.8) and adversarial batch prediction with domain-specific batch normalization. The cell-type classification objective was disabled. An 80:20 label-free training–validation split was used and the checkpoint with the lowest validation masked-MSE was retained. The final 512-dimensional cell states were L2 normalized. For single-batch tissues, adversarial batch prediction, domain-specific batch normalization and batch-label input were disabled, but unsupervised target-data fine-tuning was otherwise retained.

For gene selection in this implementation, batch-aware Seurat-v3 HVG selection was attempted first. If that implementation was unavailable or failed, the method used `cell_ranger`, followed when necessary by a final non-batch-aware `cell_ranger` retry.

##### *1.2.4 Leiden clustering and biological-conservation metrics*

All metrics were calculated directly from the complete saved embedding matrix. The benchmark did not centre, scale or project the representation before evaluation. Cells without a reference `cell_type` value would be removed; the curated panel yielded a complete 20-method by 26-tissue biological metric matrix. UMAP coordinates were generated only for visualization and were never used as input to a benchmark metric.

**Leiden clustering.** For every tissue–representation pair, a separate 15-nearest-neighbour graph was constructed from Euclidean distances using Scanpy. Leiden clustering used the igraph backend, an undirected graph, two iterations and random seed 0. Resolutions from 0.1 to 3.0 were scanned in increments of 0.1. If none of these resolutions produced at least as many clusters as there were annotated cell types, the scan was extended in increments of 0.5 to a maximum resolution of 6.0. The selected resolution minimized the absolute difference between the number of Leiden clusters and the number of reference cell types; ties were resolved in favour of the solution with the larger NMI. We refer to this as reference-informed resolution calibration: reference labels were not supplied to the embedding or integration method, but both the number of reference classes and NMI entered this post hoc evaluation rule. The same rule and resolution grid were applied to every representation. K-means clustering was also evaluated as a sensitivity analysis but was not used for the primary manuscript result.

**Cluster-concordance family.** Agreement between the selected Leiden partition and the reference cell types was quantified using adjusted Rand index (ARI), normalized mutual information (NMI; arithmetic normalization), homogeneity (HOM), completeness (COM) and Fowlkes–Mallows index (FMI). These metrics capture chance-adjusted pairwise agreement, shared information, cluster purity, recovery of complete reference classes and pairwise precision–recall agreement, respectively. All five were oriented so that larger values indicated greater concordance.

More explicitly, let $n_{uv}$ denote the number of cells shared by reference class $u$ and Leiden cluster $v$, let $a_u = \sum_v n_{uv}$, $b_v = \sum_u n_{uv}$, and let $n$ be the total number of cells. ARI was calculated as

$$\mathrm{ARI} = \frac{\displaystyle\sum_{uv} \binom{n_{uv}}{2} - \frac{\sum_u \binom{a_u}{2} \sum_v \binom{b_v}{2}}{\binom{n}{2}}}{\displaystyle\frac{1}{2}\left[\sum_u \binom{a_u}{2} + \sum_v \binom{b_v}{2}\right] - \frac{\sum_u \binom{a_u}{2} \sum_v \binom{b_v}{2}}{\binom{n}{2}}}. \tag{S2}$$

For reference labels $Y$ and Leiden assignments $Z$, arithmetic NMI was

$$\mathrm{NMI}(Y, Z) = \frac{2I(Y; Z)}{H(Y) + H(Z)}. \tag{S3}$$

Homogeneity and completeness were

$$\mathrm{HOM} = 1 - \frac{H(Y \mid Z)}{H(Y)}, \qquad \mathrm{COM} = 1 - \frac{H(Z \mid Y)}{H(Z)}, \tag{S4}$$

with the standard zero-entropy conventions used by scikit-learn. Finally, if TP, FP and FN denote pairs of cells assigned to the same cluster and/or sharing a reference label, FMI was

$$\mathrm{FMI} = \frac{\mathrm{TP}}{\sqrt{(\mathrm{TP} + \mathrm{FP})(\mathrm{TP} + \mathrm{FN})}}. \tag{S5}$$

**Continuous cell-type separation.** Average silhouette width (ASW) was calculated from Euclidean distances in the native embedding and the reference cell-type labels:

$$s_i = \frac{b_i - a_i}{\max(a_i, b_i)}, \tag{S6}$$

where $a_i$ is the mean distance from cell $i$ to cells of the same type and $b_i$ is the minimum mean distance to a different cell type. The reported ASW was the mean over cells and was retained on its native $[-1, 1]$ scale. Because ASW is sensitive to distance geometry, anisotropy, unequal group sizes and departures from compact convex clusters, it was treated as a complementary family and given less weight than the cluster and local-neighbourhood families.

**Local cell-type-neighbourhood family.** Five-nearest-neighbour classification accuracy (Acc@kNN) was estimated using a Euclidean kNN classifier and stratified fivefold cross-validation. Folds were shuffled with random seed 42 and the mean fold accuracy was reported. Entire cell types represented by fewer than five cells were excluded because they could not be distributed across all folds. Each retained cell was predicted only when it belonged to the held-out fold; Acc@kNN was the mean of the five fold-specific proportions of correctly predicted labels.

$$\mathrm{Acc@kNN} = \frac{1}{5} \sum_{q=1}^{5} \frac{1}{|\mathcal{D}_q|} \sum_{i \in \mathcal{D}_q} \mathbf{1}(\widehat{y}_i^{(q)} = y_i), \tag{S7}$$

where $\mathcal{D}_q$ denotes held-out fold $q$, and $\widehat{y}_i^{(q)}$ is predicted using only the other four folds.

Cell-type local inverse Simpson's index (cLISI) was calculated with `scib-metrics` from an approximate Euclidean nearest-neighbour graph using perplexity 30 and 90 requested neighbours. If $\widetilde{L}$ is the median of

$$L_i^{\mathrm{celltype}} = \left( \sum_{c=1}^{C} p_{ic}^2 \right)^{-1}, \tag{S8}$$

where $p_{ic}$ is the LISI kernel estimate of the local probability of cell type $c$, and $C$ is the number of reference cell types, the scaled score was

$$\text{cLISI} = \frac{C - \widetilde{L}}{C - 1}. \quad \text{(S9)}$$

It ranges from 0 to 1 and is oriented so that larger values indicate greater local cell-type purity.

Graph connectivity (GC) was calculated on a 15-nearest-neighbour graph:

$$\text{GC} = \frac{1}{C} \sum_{c=1}^{C} \frac{|\text{LCC}_c|}{|c|}, \quad \text{(S10)}$$

where $\text{LCC}_c$ is the largest connected component of the subgraph induced by cells of type $c$. Larger values indicate that cells sharing a biological identity remain connected. Approximate neighbour graphs used PyNNDescent with Euclidean distance, low-memory mode and random seed 0; requested neighbourhood sizes were capped at $n - 1$. No cells were subsampled for biological-conservation evaluation.

##### *1.2.5 Supplementary batch-mixing evaluation*

Batch mixing was evaluated as a separate objective in the 19 multi-batch tissues. All 470,978 cells from these tissues were used; no batch-metric subsampling was applied. The seven single-batch tissues, containing 77,999 cells in total, were recorded as not applicable rather than assigned a batch score of zero or one. A cell was eligible only when both `cell_type` and `batch_id` were non-missing. The curated panel contained no missing values that affected the completed analysis. Although the implementation supports deterministic label-by-batch-stratified subsampling, the production setting was `batch_max_cells=0`; thus every eligible cell was retained.

For each method–tissue pair, one approximate Euclidean neighbour graph was constructed with PyNNDescent using low-memory mode and random seed 0. The requested graph size was the larger of the kBET neighbourhood (50) and three times the LISI perplexity (90), and was capped at $n - 1$. The first 50 neighbours were supplied to kBET and the first 90 to the auxiliary global iLISI calculation. CiLISI used separate within-cell-type graphs, as described below. Batch metrics were computed from the native embedding without centring, scaling or additional dimensionality reduction.

Three higher-is-better metrics entered the batch family. kBET was calculated using `scib_metrics.kbet_per_label` on the first 50 neighbours of an approximate Euclidean graph, with $\alpha = 0.05$ and 100 diffusion components. Within each eligible reference cell type, the test compares the observed batch composition of local neighbourhoods with the batch proportions expected under local mixing. The function returned a kBET acceptance score for each testable cell type, and the reported tissue-level scalar was the unweighted mean of the available cell-type-level scores; it was therefore not weighted by cell-type abundance. Larger values indicate that a greater proportion of tested neighbourhoods failed to reject the local-mixing null. The per-label implementation used diffusion-map distances to equalize cell-type-specific neighbourhood graphs. Cell types with fewer than ten cells, represented by only one batch or otherwise unable to support the test were omitted by the library implementation.

Conceptually, if $O_{ib}$ is the observed number of neighbours from batch $b$ around tested cell $i$, $k_i$ is its neighbourhood size and $\pi_{cb}$ is the frequency of batch $b$ within cell type $c$, the local Pearson statistic is

$$X_i^2 = \sum_b \frac{(O_{ib} - k_i \pi_{cb})^2}{k_i \pi_{cb}}. \quad \text{(S11)}$$

The neighbourhood is accepted when the associated $P$ value exceeds $\alpha$. Within each eligible cell type, cell-level acceptance indicators are averaged to obtain a label-level score; the final kBET value is the unweighted mean of the available label-level scores.

Batch-removal-adapted silhouette (BRAS) was calculated using `scib-metrics` v0.5.6 with cosine distance, a pairwise-distance chunk size of 256, with `between_cluster_distances` set to `mean_other`. Within each eligible cell type, $a_i$ is the mean cosine distance between cell $i$ and cells from the same batch, whereas $b_i^*$ is the mean cosine distance to cells from all other batches of that cell type. Defining

$$s_i^* = \frac{b_i^* - a_i}{\max(a_i, b_i^*)}, \tag{S12}$$

the cell-level mixing score is $1 - |s_i^*|$. Values were averaged within cell type and then averaged equally across eligible cell types. This `mean_other` construction avoids defining the between-batch distance by only the nearest alternative batch. Embeddings with non-finite or zero-norm cell vectors were rejected for cosine BRAS, and completed values were required to lie in $[0, 1]$.

For cell-type-conditioned integration LISI (CiLISI), batch-label LISI was calculated separately within each cell type containing at least ten cells and at least two batches. The perplexity was the smaller of 30 and $n_c - 1$, and the requested number of neighbours was $\max(3\,\text{perplexity}, \text{perplexity} + 1)$. For a cell type represented across $B_c$ batches, the local inverse Simpson index was

$$L_i^{\text{batch}} = \left( \sum_{b=1}^{B_c} p_{ib}^2 \right)^{-1}, \tag{S13}$$

where $p_{ib}$ is the LISI kernel estimate of the local probability of batch $b$. Each cell's value was scaled as

$$\widetilde{L}_i^{\text{batch}} = \frac{L_i^{\text{batch}} - 1}{B_c - 1}, \tag{S14}$$

clipped to $[0, 1]$, and then pooled over all eligible cells. This final average was therefore cell-weighted rather than an unweighted mean of cell-type summaries. Global iLISI was saved as an auxiliary diagnostic but excluded from aggregation to avoid assigning duplicate weight to related LISI-based statistics. Specifically, global iLISI used the 90-neighbour graph, perplexity 30 and `scale=True`, without conditioning on cell type. It can therefore increase when biologically distinct populations are mixed and was retained only to aid diagnosis.

A batch metric was written as missing, together with an explicit reason, when the tissue contained fewer than two batches, when no eligible label remained, or when the metric's own eligibility rule could not be satisfied. Missing batch values were never replaced by a best-case or worst-case score. BRAS outputs were additionally required to be finite and within $[0, 1]$ for every multi-batch method–tissue pair.

##### *1.2.6 Descriptive embedding visualization*

UMAP was used only to visualize cell-type and batch organization and did not enter any metric or statistical test. Coordinates were generated separately for every method–tissue embedding using Scanpy. A 15-nearest-neighbour graph was constructed from the saved representation with cosine distance, followed by UMAP with random seed 42. The same saved two-dimensional coordinates were then reused for the cell-type-coloured and batch-coloured panels; the batch panels therefore did not involve a second UMAP fit.

##### *1.2.7 Metric-family aggregation and statistical inference*

All raw metrics were oriented so that larger values indicated better performance. For tissue $t$, method $m$ and metric $j$, the 20 representations were ranked in descending score order,

$$r_{tmj} = \mathrm{rank}_m(-s_{tmj}), \tag{S15}$$

with average ranks assigned to ties. Metrics were grouped into cluster concordance (ARI, NMI, HOM, COM and FMI), continuous cell-type separation (ASW), local cell-type-neighbourhood structure (Acc@kNN, cLISI and GC), and, in the supplementary analysis, batch mixing (kBET, BRAS and CiLISI). The family rank was the unweighted arithmetic mean of its constituent metric ranks,

$$\bar{r}_{tmf} = \frac{1}{|f|} \sum_{j \in f} r_{tmj}. \tag{S16}$$

For any analysis with available family set $\mathcal{A}_{tm}$, family ranks were combined as

$$R_{tm} = \frac{\sum_{f \in \mathcal{A}_{tm}} w_f \bar{r}_{tmf}}{\sum_{f \in \mathcal{A}_{tm}} w_f}. \tag{S17}$$

This available-weight denominator is important for single-batch tissues: unavailable batch information changes the denominator but is never encoded as a numerical batch score. No second ranking was applied after family aggregation.

Following the scIB principle that preservation of biological structure should receive greater weight in an integration summary [3], the biological families received 0.75 of the combined weight. Cluster concordance and local neighbourhood structure each received 0.325. ASW was retained with weight 0.10 as an auxiliary measure of global separation because silhouette-based summaries can be poorly aligned with integration quality [4]. The primary biological-conservation aggregate was

$$R_{tm}^{\mathrm{bio}} = \frac{0.325\, \bar{r}_{tm,\mathrm{cluster}} + 0.10\, \bar{r}_{tm,\mathrm{ASW}} + 0.325\, \bar{r}_{tm,\mathrm{local}}}{0.325 + 0.10 + 0.325}. \tag{S18}$$

The effective biological-only family weights were therefore 0.4333, 0.1333 and 0.4333. This lower contribution from ASW retained information about continuous separation while limiting the influence of a single global distance-based statistic. Family ranks and tissue-level aggregate values were not re-ranked. A method's overall summary was the equal-weight arithmetic mean of $R_{tm}^{\mathrm{bio}}$ over the 26 tissues; tissue contributions did not depend on cell count.

For the supplementary biological-plus-batch analysis, the batch-family rank was

$$\bar{r}_{tm,\mathrm{batch}} = \frac{r_{tm,\mathrm{kBET}} + r_{tm,\mathrm{BRAS}} + r_{tm,\mathrm{CiLISI}}}{3}, \tag{S19}$$

and the aggregate in a multi-batch tissue was

$$R_{tm}^{\mathrm{bio+batch}} = 0.325\, \bar{r}_{tm,\mathrm{cluster}} + 0.10\, \bar{r}_{tm,\mathrm{ASW}} + 0.325\, \bar{r}_{tm,\mathrm{local}} + 0.25\, \bar{r}_{tm,\mathrm{batch}}. \tag{S20}$$

The four weights sum to one. In a single-batch tissue, the unavailable batch family was omitted and the three biological weights were renormalized, making the resulting value identical to $R^{\text{bio}}_{tm}$. Thus, tissue-level biological-plus-batch values remained available for all 26 tissues, whereas inference on an individual batch metric or on the batch-family rank used only the 19 multi-batch tissues. The stored simple mean of all individual metric ranks was secondary and was not the family-balanced manuscript outcome.

The four-family aggregate included batch metrics in 19 tissues and the three biological families in the seven single-batch tissues. Batch-mixing results were based on the individual batch metrics and batch-family rank in the 19 informative tissues.

Per-tissue tables additionally displayed arithmetic means of raw metric values within each family. Cross-tissue statistical inference used the rank-first family aggregate defined above.

**Statistical analysis.** Tissues, rather than cells, donors, batches, cross-validation folds or embedding dimensions, were treated as the paired experimental units. Each tissue contributed one value per method, and tissues were weighted equally. The primary biological matrix contained 20 representations, 26 tissues and nine complete metrics.

Global differences among methods were tested using the Friedman repeated-measures test applied to the tissue-level metric-family aggregate. Kendall's coefficient of concordance was derived from the Friedman statistic,

$$W = \frac{\chi^2}{N(k-1)}, \tag{S21}$$

where $N = 26$ tissues and $k = 20$ methods. This statistic quantifies the consistency of relative method ranks across tissues.

For method $m$, the PCA-relative aggregate-rank effect in tissue $t$ was

$$\Delta_{tm} = R^{\text{bio}}_{t,\text{PCA}} - R^{\text{bio}}_{tm}, \tag{S22}$$

so that a positive value favoured method $m$. Tissue-matched differences were tested using two-sided paired Wilcoxon signed-rank tests (`zero_method="wilcox"`). Holm correction was applied across the 19 non-PCA comparisons within the aggregate analysis. The same procedure was repeated with pretrained scVI as a secondary reference.

Uncertainty in mean raw scores, mean family ranks, mean aggregate ranks and paired differences from a reference method was quantified using 10,000 paired tissue-level bootstrap resamples with random seed 20260602. Tissues were sampled with replacement and the complete vector of methods for a selected tissue was retained together. We report percentile intervals defined by the 2.5th and 97.5th percentiles. Biological-only and biological-plus-batch aggregate analyses resampled all 26 tissues; raw batch and batch-family analyses resampled the 19 complete multi-batch tissues. Influence was examined by omitting each tissue in turn, recomputing the equal-weight mean aggregate rank and recording changes in the final method order.

The supplementary biological-plus-batch aggregate was subjected to the same Friedman, Kendall-$W$, paired Wilcoxon, bootstrap and leave-one-tissue-out procedures using all 26 tissues. For its seven single-batch tissues this aggregate reduces to the renormalized biological-only value, as defined above. In contrast, analyses intended to make a statement specifically about batch mixing—an individual batch metric or the batch-family rank—were restricted to the 19 multi-batch tissues. These two estimands were reported separately: the former summarizes joint biological preservation and mixing over the full panel, whereas the latter estimates batch mixing only where that property is observable.

Metric-level PCA comparisons were performed separately from the primary aggregate-rank tests. For every tissue and metric, we calculated

$$d_{tmj} = s_{tmj} - s_{t,\mathrm{PCA},j}. \tag{S23}$$

The paired differences were visualized as boxplots and tested against zero using a two-sided Wilcoxon signed-rank test with a normal approximation and continuity correction; at least two non-zero tissue pairs were required. The displayed significance symbols used a single Benjamini–Hochberg adjustment over all finite non-PCA method–metric comparisons across the nine biological and three batch metrics (228 tests in the completed analysis). This global false-discovery-rate analysis is complementary to, and deliberately distinct from, the Holm-corrected primary comparisons of family aggregate ranks. Biological metric comparisons contained 26 tissue pairs; batch metric comparisons contained 19. A missing single-batch result did not enter either the test statistic or the multiplicity count as an artificial observation.

To describe empirical redundancy among metrics, we correlated method ranks between every pair of metrics within each tissue. Pearson correlation of precomputed average-tie ranks is equivalent to Spearman rank correlation. Tissue-specific correlations based on at least five common methods were Fisher-$z$ transformed, averaged with weights equal to the number of common methods minus three and back-transformed; at least three valid tissues were required. This analysis was descriptive, was used to assess convergence within the defined metric families, and was not assigned inferential $P$ values.

###### *1.2.8 Reproducibility and software*

Embedding extraction, target-data fitting and evaluation were orchestrated with Nextflow, and method environments were isolated in containers. The canonical biological benchmark used Python 3.10.19, NumPy 2.2.6, pandas 2.3.3, scikit-learn 1.7.2, Scanpy 1.11.5, AnnData 0.11.4, PyNNDescent 0.5.13 and scib-metrics 0.5.6. The completed result contained 26 embedding files for each of the 20 representations. Duplicate method–tissue–metric records were rejected, and a complete expected method–tissue–metric grid was audited before aggregation.

Explicit inference or fitting seeds were fixed for PCA (0), CellFM (0), CellPLM (42), pretrained scVI (0), scVI (de novo) (0), scGPT (integrated) (0) and the Harmony/Seurat methods (0); biological Leiden and approximate neighbour graphs used seed 0 and Acc@kNN folds used seed 42. The current UCE official inference path performs NumPy-based expression-weighted sampling, and the scPRINT `random expr` path also contains a stochastic gene-selection step; these methods used their released sampling paths.

##### 1.3 Tabula Sapiens v1–v2 cross-release robustness sensitivity analysis

###### *1.3.1 Study design*

We compared the Tabula Sapiens v1 (TSv1) and Tabula Sapiens v2 (TSv2) evaluations to determine whether the zero-shot cell-representation hierarchy was robust across atlas releases. The manual pretraining-corpus audit is reported in Table S1. The releases were matched by tissue; their cells, donors, batch composition and cell-type composition differed.

The primary comparison used biological-conservation performance. A supplementary analysis repeated the comparison using the joint biological-plus-batch aggregate. Both analyses retained the same 20-representation panel used in the TSv2 benchmark.

##### *1.3.2 Atlas preparation and tissue-level matching*

TSv1 comprised 24 tissue datasets and 444,116 cells. Individual tissues contained 4,665–51,173 cells, 13–38 annotated cell types and 2–11 batches. All 24 TSv1 tissues were multi-batch. TSv2 comprised 26 tissue datasets and 548,977 cells, with 2,044–72,290 cells and 14–32 annotated cell types per tissue; 19 tissues were multi-batch and seven were single-batch.

TSv1 and TSv2 were processed as described in Section 1.1. As in TSv2, batch identity was defined as the assay–donor combination,

$$\texttt{batch\_id} = \texttt{assay} \mid \texttt{donor\_id}. \tag{S24}$$

The same metadata harmonization, embedding extraction, target-data reference methods and benchmark implementations were used for the two atlas versions. Each version's curated `cell_type` annotation was used as its biological reference.

Tissue names were standardized and were then matched exactly. Twenty-two tissues were shared by name: bladder, blood, bone marrow, eye, fat, heart, large intestine, liver, lung, lymph node, mammary, muscle, prostate, salivary gland, skin, small intestine, spleen, thymus, tongue, trachea, uterus and vasculature. Kidney and pancreas occurred only in TSv1, whereas ear, ovary, stomach and testis occurred only in TSv2. The 22 shared tissues contained 422,894 TSv1 cells and 456,403 TSv2 cells. Matching was performed only at the tissue-name level: cells, donors, batches, genes and annotation labels were not paired across versions. Accordingly, tissue, rather than cell or donor, was the experimental unit in paired cross-version analyses.

##### *1.3.3 Aligned representation panel and benchmark outcomes*

The two releases were evaluated with the same 20-representation roster: CELLama, CellFM, CellPLM, Geneformer, GenePT-w, Harmony, LangCell, PCA, SCimilarity, Seurat CCA, Seurat RPCA, UCE, scBERT, scCello, scFoundation, scGPT, scGPT (integrated), scPRINT, pretrained scVI and scVI (de novo).

For each version, biological conservation was evaluated using nine higher-is-better metrics: adjusted Rand index (ARI), normalized mutual information (NMI), homogeneity (HOM), completeness (COM), Fowlkes–Mallows index (FMI), cell-type average silhouette width (ASW), five-nearest-neighbour classification accuracy (Acc@kNN), cell-type local inverse Simpson's index (cLISI) and graph connectivity (GC). The supplementary batch analysis used three additional higher-is-better metrics: kBET acceptance, batch-removal-adapted silhouette (BRAS; cosine distance and mean distance to all other batches within cell type) and cell-type-conditioned integration LISI (CiLISI). Global iLISI was retained as a diagnostic but did not enter the aggregate. Metric definitions and model-specific embedding procedures were otherwise identical to those described for the TSv2 zero-shot analysis (Methods 1.2).

For version $v$, tissue $t$, method $m$ and metric $j$, methods were ranked within each tissue and metric in descending order of the raw score $s_{vtmj}$:

$$r_{vtmj} = \text{rank}_m(-s_{vtmj}), \tag{S25}$$

with average ranks assigned to ties. Metrics were grouped into cluster concordance (ARI, NMI, HOM, COM and FMI), continuous separation (ASW), local cell-type-neighbourhood structure (Acc@kNN, cLISI and GC), and batch mixing (kBET, BRAS and CiLISI). The rank for family $f$ was the unweighted mean of its constituent metric ranks,

$$\bar{r}_{vtmf} = \frac{1}{|f|} \sum_{j \in f} r_{vtmj}. \tag{S26}$$

Family ranks were combined using prespecified weights of 0.325 for cluster concordance, 0.10 for continuous separation, 0.325 for local-neighbourhood structure and 0.25 for batch mixing:

$$R_{vtm} = \frac{\sum_{f \in \mathcal{A}_{vtm}} w_f \bar{r}_{vtmf}}{\sum_{f \in \mathcal{A}_{vtm}} w_f}, \tag{S27}$$

where $\mathcal{A}_{vtm}$ is the set of applicable families. No second ranking was performed after family aggregation. In the primary biological-only analysis, the first three weights were renormalized by their sum of 0.75, giving effective weights of 0.4333, 0.1333 and 0.4333. In a multi-batch tissue, the supplementary aggregate used all four weights, which sum to one. If batch metrics were not applicable, the batch family was omitted rather than assigned a favourable or unfavourable value, and the three biological weights were renormalized.

##### *1.3.4 Atlas-wide rank concordance*

We first compared the broad method hierarchy across the complete version-specific tissue panels. Let $\mathcal{T}_v$ denote all tissues in version $v$, with $N_1 = 24$ for TSv1 and $N_2 = 26$ for TSv2. The equal-tissue mean aggregate rank for method $m$ was

$$\bar{R}_{vm} = \frac{1}{N_v} \sum_{t \in \mathcal{T}_v} R_{vtm}, \tag{S28}$$

and the atlas-wide rank position was

$$Q_{vm} = \text{rank}_m(\bar{R}_{vm}), \tag{S29}$$

where smaller values indicate better performance. Thus, this descriptive analysis used all available tissues in each atlas, rather than restricting both versions to the 22 shared tissue names.

Concordance between the two 20-method rank vectors was summarized using Spearman's rank correlation, Kendall's $\tau$, pairwise-ordering agreement and top-$k$ overlap for $k = 3$ and 5. Pairwise-ordering agreement was the proportion of method pairs with the same non-tied ordering in TSv1 and TSv2:

$$\text{agreement} = \frac{1}{|\mathcal{P}|} \sum_{(m,m') \in \mathcal{P}} \mathbf{1}[(Q_{1m} - Q_{1m'})(Q_{2m} - Q_{2m'}) > 0], \tag{S30}$$

where $\mathcal{P}$ contains method pairs that are non-tied in both versions. Top-$k$ overlap was $|\text{Top}_{k,1} \cap \text{Top}_{k,2}|/k$. Two-sided nominal $P$ values were reported for Spearman's correlation and Kendall's $\tau$. These statistics summarize concordance within the fixed 20-method panel.

##### *1.3.5 Tissue-matched PCA-relative analysis*

The primary cross-version inference used the 22 tissue names shared between TSv1 and TSv2 and expressed each method's performance relative to PCA within the same atlas version and tissue. For the aggregate rank, the PCA-relative advantage was

$$D_{vtm} = R_{vt,\text{PCA}} - R_{vtm}. \tag{S31}$$

Positive values indicate that method $m$ ranked better than PCA. The cross-version change was

$$C_{tm} = D_{2tm} - D_{1tm}, \tag{S32}$$

so that a positive value indicates a larger PCA-relative advantage in TSv2 and a negative value indicates attenuation of that advantage in TSv2. For each of the 19 non-PCA methods, $C_{tm}$ was tested against zero across tissues using a two-sided paired Wilcoxon signed-rank test with `zero_method="wilcox"`. Holm adjustment was applied across the 19 methods within the aggregate-rank analysis. A paired $t$-test was stored as a parametric sensitivity analysis but was not used for the primary inference or figure annotations.

Raw metrics were analysed in parallel. For metric $j$, we calculated the within-version difference from PCA,

$$d_{vtmj} = s_{vtmj} - s_{vt,\mathrm{PCA},j}, \tag{S33}$$

and, as a secondary scale-free display, the PCA-normalized ratio

$$q_{vtmj} = \frac{s_{vtmj}}{s_{vt,\mathrm{PCA},j}}. \tag{S34}$$

The ratio was defined only when the corresponding PCA score was strictly positive and both version-specific ratios were finite. Aggregate-rank advantage and raw-score difference were the primary endpoints; ratios were used to display relative changes on a common scale.

For every method–metric combination, TSv1 and TSv2 were restricted to the same finite tissue pairs before plotting or testing. The version difference in $d_{vtmj}$, and separately in $q_{vtmj}$, was tested using the same two-sided paired Wilcoxon procedure. Holm correction was applied across the 19 non-PCA methods separately within each metric and separately for the difference and ratio measures. Thus, the multiplicity correction was not pooled across unrelated metrics or across the two measures. Lines joining points in the ratio plots connect the same tissue-name pair used in the corresponding paired test.

###### *1.3.6 Supplementary biological-plus-batch comparison*

The complete analysis was repeated with the biological-plus-batch aggregate. All 24 TSv1 tissues were multi-batch and therefore contributed all four metric families. In TSv2, 19 tissues contributed all four families, whereas eye, lung, mammary, prostate, skin, testis and uterus were single-batch and their aggregate reduced to the renormalized biological-only value. Direct four-family comparisons therefore used tissues with batch metrics in both releases.

For direct paired inference on the four-family aggregate, we restricted the analysis to the 16 shared tissues that were multi-batch in both versions: bladder, blood, bone marrow, fat, heart, large intestine, liver, lymph node, muscle, salivary gland, small intestine, spleen, thymus, tongue, trachea and vasculature. PCA-relative aggregate-rank changes were tested and Holm-adjusted as above. In raw-metric analyses, biological metrics could use up to all 22 shared tissues, whereas kBET, BRAS and CiLISI used only the 16 jointly multi-batch tissues with finite values. Results specific to batch mixing were interpreted from these 16 tissues rather than from the heterogeneous full-panel aggregate.

##### *1.3.7 Uncertainty and implementation*

The version-specific analyses quantified uncertainty by resampling complete tissues with replacement 10,000 times using random seed 20260602. The complete vector of methods was retained whenever a tissue was sampled, and 95% percentile intervals were taken from the 2.5th and 97.5th percentiles. Bootstrap intervals describe the version-specific estimates. Cross-version inference used the direct tissue-level paired tests described above, whereas Friedman statistics and Kendall's $W$ summarize method ranks within each atlas.

The comparison was implemented in Python 3.10.19 with NumPy 2.2.6, pandas 2.3.3, SciPy 1.15.3, Matplotlib 3.10.8 and seaborn 0.13.2; final redraws used R and ggplot2.

#### 1.4 Local expression-neighbourhood preservation in Tabula Sapiens v2

Local fidelity was evaluated against a reference constructed directly from raw UMI counts, without dimensionality reduction or cell-type labels. For each tissue we retained all cells when $n \leq 20{,}000$ and otherwise sampled 20,000 cells without replacement; the sampled cells were fixed across representations. Mitochondrial, ERCC and ribosomal-protein genes and genes detected in fewer than ten cells or with fewer than ten total counts were excluded. Analytic Pearson residuals were calculated separately within each assay–donor batch using $\theta = 100$ and clipped to $\pm\sqrt{n_b}$. Residual variance was converted to a within-batch percentile, and the 2,000 genes with the highest median percentile across eligible batches were retained.

Reference neighbours were computed by Euclidean distance directly in the unscaled residual gene-expression space; candidate neighbours were computed by Euclidean distance in each native, unstandardized representation. Both graphs were constructed within batch. Batches with fewer than 101 cells were excluded; exact nearest neighbours were used for at most 5,000 cells and PyNNDescent otherwise. For batch $b$ and neighbourhood size $k$,

$$Q_{NX,b}(k) = \frac{1}{n_b k} \sum_{i=1}^{n_b} \left| \mathcal{N}_k^X(i) \cap \mathcal{N}_k^Z(i) \right|, \qquad R_{NX,b}(k) = \frac{(n_b - 1) Q_{NX,b}(k) - k}{n_b - 1 - k}. \tag{S35}$$

Thus, $R_{NX} = 1$ denotes identical neighbourhoods and $R_{NX} = 0$ the overlap expected at random. Batch estimates were weighted by eligible cell count and averaged equally over $k = 15$, 30 and 50.

For association with biological-conservation performance, geometry and performance were first averaged equally across the 26 tissues for each of the 20 representations. To avoid part–whole overlap, the primary performance outcome omitted Acc@kNN, cLISI and graph connectivity, retained the original within-tissue cluster-concordance and ASW family ranks, and combined them as $(0.325 R_{\text{cluster}} + 0.10 R_{\text{ASW}})/0.425$ without re-ranking methods. Performance was the negative aggregate rank. Spearman correlations were calculated across all 20 representations and, as a prespecified scope sensitivity, across the 13 frozen scFMs. Reported $P$ values were two-sided, nominal and unadjusted; method was the association unit.

#### 1.5 Additional representation-geometry and model-characteristic analyses

For an uncentred cell embedding $Z \in \mathbb{R}^{n \times d}$, spectral anisotropy was defined as the fraction of embedding energy in its leading singular direction,

$$A_{\text{spec}} = \frac{\sigma_1^2(Z)}{\|Z\|_F^2}. \tag{S36}$$

The leading singular value was estimated by randomized singular-value decomposition. We separately calculated mean cosine similarity among cells of the same type, $A_{\text{within}}$, and among cells of different types, $A_{\text{between}}$, using 100,000 cell pairs per condition and uniform sampling over cell types. Their difference, $\Delta A = A_{\text{within}} - A_{\text{between}}$, measures the extent to which directional coherence follows biological identity rather than a common global direction.

Effective dimensionality was characterized after centring and scaling each coordinate to unit variance. For covariance matrix $C$, the participation ratio was

$$\mathrm{PR} = \frac{(\operatorname{tr} C)^2}{\operatorname{tr}(C^2)}. \tag{S37}$$

TwoNN intrinsic dimension was estimated from the ratio of second- to first-nearest-neighbour distances in raw and coordinate-standardized spaces. To distinguish biological from technical organization, each standardized coordinate was fitted with $z \sim 1 + \text{cell type} + \text{batch}$; cell-type and batch effects were summarized separately by partial $\eta^2$ averaged over estimable coordinates.

Geometry was calculated for each tissue–representation pair and then averaged equally over the 26 tissues. Spectral anisotropy and the cell-type anisotropy contrast were related to biological-conservation performance by Spearman correlation across the 20 representations. Associations with reported pretraining corpus size and runtime parameter count were evaluated by Spearman correlation; model family and retention of expression magnitude were evaluated by explained variance with exact permutation tests. Holm correction was applied across these four model-characteristic tests. The corresponding model taxonomy is reported in Table S3.

##### 1.6 Dataset-level moderators and shared errors in rare cell populations

**Dataset-level moderators.** This analysis was restricted to the 13 frozen scFMs, with PCA as the paired reference. For each tissue, method and biological-score family, the outcome was the raw family score of the scFM minus that of PCA. The three families were cluster–label concordance (the mean of ARI, NMI, homogeneity, completeness and FMI), continuous cell-type separation (ASW), and local cell-type-neighbourhood structure (the mean of Acc@kNN, cLISI and graph connectivity). Six standardized dataset properties entered each family-specific model simultaneously with sum-to-zero method effects: $\log_{10}$ cell count, log cell-type count, normalized cell-type entropy, $\log_{10}$ median detected genes per cell, rare-cell burden (the fraction of cells belonging to types comprising less than 1% of the tissue), and the effective rank of the first 200 reference-expression principal components. The maximum variance-inflation factor was 2.16. Confidence intervals used 5,000 complete-tissue bootstrap resamples; conditional $P$ values used 10,000 Freedman–Lane permutations of complete tissue blocks. Holm correction was applied jointly across the $3 \times 6 = 18$ family–moderator tests.

**Rare-cell and consensus-error readout.** For every tissue, one cell-type-stratified fivefold assignment was shared across the complete 20-representation panel. Each fixed representation was evaluated with a five-nearest-neighbour Euclidean label readout, excluding held-out labels and same-fold neighbours. Cell types represented by fewer than five cells were excluded. Rare types comprised less than 1% of cells within a tissue, intermediate types 1–10%, and common types more than 10%. Accuracy was first averaged across eligible cell types within each abundance class and tissue. Rare-type accuracy for each of the 19 non-PCA representations was compared with PCA across the 26 tissues using two-sided paired Wilcoxon signed-rank tests with Holm correction across the 19 comparisons.

A consensus-hard cell in the complete 20-readout panel was defined as one misclassified by at least 16 methods. For the complementary pooled-composition and adjusted-odds analysis, the panel was restricted to PCA and the 13 frozen scFMs and the corresponding threshold was at least 12 of 14 incorrect readouts. Consensus-hard event counts were modelled at the tissue-by-cell-type level using binomial logistic regression with random intercepts for tissue and for cell type nested within tissue; Holm correction covered the rare-versus-common and intermediate-versus-common contrasts.

#### 1.7 Transfer to continuous developmental trajectories

##### *1.7.1 Study design and developmental trajectory data*

We evaluated transfer to continuous biological dynamics using nine reference single-cell RNA-sequencing datasets distributed with the Dynverse trajectory collection [5, 6]. The panel comprised three linear trajectories (human female germline development, human embryo development and pancreatic beta-cell maturation), three bifurcating trajectories (distal lung epithelium, fibroblast reprogramming and hepatoblast differentiation), and three tree-structured trajectories (embryonic mesenchyme-to-neuron differentiation, haematopoiesis and mesoderm development). The datasets contained 59–1,289 cells each (4,751 cells in total). Reference milestone networks, cell progressions and start-state annotations were retained from the distributed Dynverse objects.

Raw count matrices were exported from the Dynverse R objects to AnnData format. Cell identifiers and the curated reference metadata were preserved; the prepared matrices used here were integer-valued raw counts and retained a raw-count layer. Gene identifiers followed the same pre-existing cleaned input objects used by the representation analysis. No trajectory labels, milestone labels or pseudotime values were supplied to an embedding model. The identical cells and cell order were used for every representation, inference framework and reference comparison.

###### *Cell representations*

We analysed 17 representations: the 13 original frozen scFMs (CELLama, CellFM, CellPLM, Geneformer, GenePT-w, LangCell, SCimilarity, UCE, scBERT, scCello, scFoundation, scGPT and scPRINT), pretrained scVI, an expression-based PCA reference, and two target-fitted comparators, scGPT (integrated) and scVI (de novo). Model embeddings were produced through the same containerized Nextflow entry point used elsewhere in the study. The weights of the 13 frozen scFMs and pretrained scVI were not updated on the trajectory datasets. scGPT (integrated) was fitted without trajectory labels on each target expression matrix, and scVI (de novo) was initialized and trained separately within each dataset; neither method received cell-state, milestone, trajectory or pseudotime labels. The embedding matrices were saved as AnnData objects and aligned to the reference cell identifiers before trajectory analysis. All 153 dataset–representation combinations were required to contain the complete cell set, finite values and a unique dataset–method key.

The PCA reference was calculated from raw counts by library-size normalization to $10^4$ counts per cell, $\log(1 + x)$ transformation, selection of 2,000 highly variable genes using the Seurat dispersion procedure, scaling with values clipped at 10, and 50-component PCA using the ARPACK solver and random state 0. This PCA representation served as a conventional expression baseline. The local-fidelity reference described below was constructed directly in analytic-Pearson-residual gene space and did not use PCA.

*Trajectory reconstruction*

Each representation was analysed independently with four complementary trajectory-inference frameworks: Slingshot, partition-based graph abstraction (PAGA), diffusion pseudotime (DPT) and Monocle 3 [7–10]. The input cells, representation, reference trajectory and root prior were held fixed within each comparison. Stochastic operations used fixed seeds derived from a common base seed of 0.

**Slingshot.** The complete representation was substituted for the expression matrix in the Dynverse data wrapper and passed to the official Dynverse Slingshot container through `dyno::infer_trajectory`, with the known `start_id` prior and `ndim=50`. For representations with more than 50 coordinates, the wrapper's standard dimensionality-reduction step operated within this 50-dimensional ceiling before clustering and principal-curve fitting; representations with no more than 50 coordinates were supplied without this truncation. The inferred milestone network and cell progression were retained in Dynverse format.

**PAGA.** A 15-nearest-neighbour graph was constructed directly from the complete native embedding using Euclidean distance in Scanpy. Leiden clustering used the igraph implementation, an undirected graph, two iterations, resolution 1.0 and random state 0. The fallback resolution sequence 2, 5 and 10 was defined for a single-cluster result; none of the 153 runs required it. PAGA was fitted to the Leiden groups, and the official `adata.uns["paga"]["connectivities_tree"]` output was used without custom edge thresholding or graph repair. The tree was rooted in the group containing the known start cell and its edges were oriented away from this root. PAGA connectivity weights were retained in the exported graph, whereas unit edge lengths and hard one-hot cell-to-group assignments were used in the exported trajectory representation.

**DPT.** DPT used the same 15-nearest-neighbour Euclidean graph constructed from the complete native embedding. A diffusion map was computed and DPT was run from the known start cell using ten diffusion components and the documented default `n_branchings=0`. Pseudotime values were linearly rescaled to $[0, 1]$ and exported as a directed linear trajectory. No non-finite DPT pseudotime values occurred. Because this output contains ordering but no inferred branching graph, topology metrics and branch-level F1 were treated as structurally non-applicable rather than as observed zero performance.

**Monocle 3.** Monocle 3 learns its principal graph in UMAP space and therefore required an adapter for an already learned cell representation. Rather than selecting the first 50 latent coordinates, which are not generally ordered by importance, we calculated a centred but unscaled PCA from the complete embedding and retained at most 50 components. This preserved the relative variance of the learned coordinates while avoiding coordinate-order assumptions. A two-dimensional UMAP was then constructed with cosine distance, 15 neighbours, minimum distance 0.1, spectral initialization, one thread and random seed 0. The supplied UMAP was clustered with Leiden ($k = 20$, resolution $10^{-3}$, two iterations), followed by `learn_graph` and `order_cells` with `use_partition=FALSE` and `close_loop=FALSE`. The graph was rooted at the known start cell. A one-gene placeholder count matrix was used only to instantiate the Monocle 3 object; graph learning operated on the supplied embedding-derived UMAP. Pseudotime was rescaled to $[0, 1]$, and the principal graph was oriented away from the root. Principal-graph edge weights were retained as edge lengths, and each cell was assigned to its closest principal-graph vertex. This adapter follows the graph-learning portion of the Monocle 3 procedure for supplied cell representations.

###### *1.7.2 Reference-based trajectory metrics*

Inferred and reference trajectories were compared using `dyneval` v0.9.9 and `dynwrap` v1.2.5. Before metric calculation, cell waypoints were added with the default `dynwrap::add_cell_waypoints()` setting of 100 waypoints, using a deterministic dataset–method–framework seed. The metric engine returned 16 fields, of which we retained 12 direction-consistent effect-size metrics organized a priori into five biological families:

1. **Cell-position distance:** correlation of geodesic cell–cell distances between the inferred and reference trajectories.
2. **Reference-position recoverability:** normalized mean-squared error and $R^2$ from linear-model and random-forest recovery of reference position from the inferred trajectory.
3. **Topology:** edge-flip similarity, Hamming–Ipsen–Mikhailov similarity and graph isomorphism.
4. **Feature relevance:** correlation and weighted correlation of trajectory-associated feature importance.
5. **Branch assignment:** branch F1 and milestone F1.

All retained metrics were oriented so that larger values indicated greater agreement with the reference. Raw linear- and random-forest mean-squared error were excluded because they duplicate their normalized summaries and would overweight position recoverability. The feature-importance Kolmogorov–Smirnov and Wilcoxon outputs were excluded because the installed implementation returns tail probabilities rather than effect-size scores; the stored Kolmogorov–Smirnov direction also conflicts with the package metadata. For DPT, edge flip, Hamming–Ipsen–Mikhailov similarity, isomorphism and branch F1 were designated structural missing values.

*Canonical rank summary and paired comparison with PCA*

For metric $r$, dataset $t$, inference framework $b$ and representation $m$, methods were ranked within the fixed $t \times b \times r$ block, with rank 1 denoting the best value and ties assigned their average rank. Ranks were averaged first within metric family and then across the families applicable to that framework. Family summaries and aggregate values were not re-ranked. The primary specification therefore used five families for Slingshot, PAGA and Monocle 3. DPT used cell-position distance, position recoverability, feature relevance and milestone assignment, with equal weights renormalized over these four applicable families.

A common-denominator sensitivity analysis used the same four families for every framework: cell-position distance, position recoverability, feature relevance and milestone assignment. This analysis retained eight metrics: geodesic-distance correlation; linear and random-forest normalized mean-squared error and $R^2$; feature-importance correlation and weighted correlation; and milestone F1.

Framework-specific aggregate ranks were first averaged within each dataset–representation pair. The nine datasets, rather than cells, milestones, waypoints or the 36 dataset–framework blocks, were treated as the inferential units. We obtained 95% percentile stability intervals from 10,000 paired dataset-bootstrap replicates stratified by trajectory class, resampling three linear, three bifurcating and three tree-structured datasets with replacement within their respective strata.

PCA-relative rank differences were defined as

$$\Delta_{tm}^{\text{rank}} = R_{t,\text{PCA}} - R_{tm}, \tag{S38}$$

where $R_{tm}$ is the four-framework mean aggregate rank. Positive values favour the evaluated representation. Each non-PCA representation was compared with PCA by a two-sided exact paired sign-flip test over all $2^9$ sign assignments, followed by Holm correction across 16 comparisons.

An omnibus Friedman statistic was calculated from the within-dataset method ranks. Its $P$ value was obtained from 100,000 permutations of method labels within complete dataset blocks, and Kendall's $W$ was reported as a measure of concordance among dataset-level orderings. Pairwise framework agreement was summarized by Spearman correlation of method-level mean ranks. Mean ranks were also summarized within each of the three trajectory classes (three datasets per class).

*Spectral anisotropy*

Geometry was computed directly from each native cell embedding, using every cell in each dataset. Let $Z_{tm} \in \mathbb{R}^{n_t \times d_m}$ denote the uncentred embedding for dataset $t$ and representation $m$, and let $\sigma_1(Z_{tm})$ be its leading singular value. Spectral anisotropy was defined as

$$A_{tm} = \frac{\sigma_1^2(Z_{tm})}{\|Z_{tm}\|_F^2}. \tag{S39}$$

This quantity lies in $[0, 1]$. Values approaching 1 indicate that most raw embedding energy is concentrated along one global direction, consistent with a cone-like collapse; smaller values indicate a more diffuse spectrum. The embedding was deliberately not centred, standardized or dimension-normalized, matching the definition used in the tissue analysis. The leading singular value was estimated with randomized SVD using five power iterations and a deterministic dataset–method seed. Because the measure is sensitive to origin, coordinate-wise scaling and embedding dimensionality, it was interpreted as an empirical property of the representation emitted by each model, not as a coordinate-invariant latent-space quantity.

*Local expression-neighbourhood fidelity*

The expression reference was constructed separately within each complete dataset because no uniform technical-batch field was available and a stage field could be confounded with biological progression. Raw non-negative integer counts were used. Mitochondrial, ERCC and ribosomal-protein genes and genes detected in fewer than three cells or with fewer than three total counts were removed. Analytic Pearson residuals were calculated with overdispersion parameter $\theta = 100$, and up to 2,000 genes were selected by residual variance. Exact Euclidean neighbour graphs were computed directly in this residual gene space and in each native representation, without dimensionality reduction or biological labels. Self-neighbours were excluded and no cells were subsampled.

For dataset $t$, representation $m$ and neighbourhood size $k$, let $Q_{NX,tm}(k)$ be the mean fraction of the $k$ expression-reference neighbours recovered in the representation. Chance-corrected preservation was

$$R_{NX,tm}(k) = \frac{(n_t - 1)Q_{NX,tm}(k) - k}{n_t - 1 - k}. \tag{S40}$$

The dataset-level fidelity outcome was the equal-weight mean over $k \in \{5, 10, 15\}$. Because the reference is a direct gene-space construction rather than an expression PCA, PCA was included in the primary 17-representation association.

*Continuous trajectory-performance outcome for geometry analysis*

The geometry analysis used continuous `dyneval` values rather than the canonical rank composite. Let $s_{tbmr}$ be the score for dataset $t$, inference framework $b$, representation $m$ and retained metric $r$, and let $q_r = 1$ for a higher-is-better metric and $-1$ for a lower-is-better metric. The paired, direction-aligned difference from PCA was

$$\delta_{tbmr} = q_r \left(s_{tbmr} - s_{tb,\text{PCA},r}\right). \tag{S41}$$

All 12 retained fields were stored in a higher-is-better orientation in the canonical analysis; excluded raw MSE and tail-probability fields did not enter this outcome. For each metric family $f$, differences were averaged across the applicable metrics,

$$D_{tbmf} = \frac{1}{|f_{tb}|} \sum_{r \in f_{tb}} \delta_{tbmr}, \tag{S42}$$

and family values were then averaged with equal weight,

$$D_{tbm} = \frac{1}{|F_{tb}|} \sum_{f \in F_{tb}} D_{tbmf}. \tag{S43}$$

DPT structural missing values were removed before these averages, so its primary continuous score used four families whereas the other frameworks used five. Frameworks were averaged within dataset and datasets within representation:

$$D_{tm} = \frac{1}{4} \sum_{b} D_{tbm}, \qquad \overline{D}_m = \frac{1}{9} \sum_{t} D_{tm}. \tag{S44}$$

Thus, $\overline{D}_m > 0$ denotes better mean direction-aligned trajectory performance than PCA and $\overline{D}_{\text{PCA}} = 0$ by construction. The calculation gives equal weight to metric families, inference frameworks and datasets and prevents a family containing more reported metrics from dominating the outcome.

*Association analysis and sensitivity analyses*

The primary geometry association related mean $R_{NX}$ to the continuous PCA-relative trajectory outcome across all 17 representations. The analysis was repeated within the 13 frozen scFMs. Spearman correlation was calculated after first averaging geometry and performance across the same nine datasets. The representation, rather than a cell or a dataset–backend row, was the association unit.

Uncertainty due to the finite developmental-system panel was summarized by 10,000 trajectory-class-stratified dataset-bootstrap replicates. Within each replicate, three datasets were sampled with replacement from each of the linear, bifurcating and tree-structured strata, after which method-level geometry and performance means and their Spearman correlation were recalculated. The 2.5th and 97.5th percentiles are reported as stability intervals across developmental systems.

Two-sided $P$ values were calculated from 100,000 whole-method-label permutations, with a plus-one Monte Carlo correction. The $R_{NX}$ analysis used seed 20260811, and the spectral-anisotropy analysis used seed 20260725. Robustness analyses repeated the association at each neighbourhood size, within the frozen-scFM scope, with the common-denominator trajectory outcome, with the canonical rank outcome, after removing the complete reference-position-recoverability family, and after omitting one method at a time.

The same method-level association analysis related mean spectral anisotropy to the continuous trajectory outcome across all 17 representations. The $R_{NX}$ and anisotropy analyses therefore used the same developmental-system panel and performance outcome while probing local and global properties of the delivered representations, respectively.

*Trajectory visualizations*

The cross-backend maps were assembled separately for each dataset. Rows show the four inference backends, and each row contains the reference trajectory followed by the same 17 representations. Cells are coloured by the reference milestone or state; nodes, edges and arrows show the corresponding inferred milestone graph and orientation. Layouts were learned separately for every representation and backend; absolute orientation and scale therefore vary between panels.

Canonical heatmaps use the 12 retained fields and the within-block average-tie rank procedure described above; family and overall values were not re-ranked, and DPT structural missing values are shown as blanks. The dataset-specific metric panels show the complete `dyneval` outputs and their within-family summaries for each representation and backend. Pseudotime ribbons use the complete reference progression for linear datasets and the longest reference lineage for branching or tree-structured datasets. Cells are ordered by reference progression, and pseudotime is min–max normalized within each dataset–backend–representation panel. Monocle 3 and DPT use their exported cell pseudotime, whereas Slingshot and PAGA pseudotime is derived from their inferred Dynwrap progressions.

*Reproducibility and software*

Trajectory metric calculation used `dyneval` v0.9.9 and `dynwrap` v1.2.5. The Slingshot wrapper used `tislingshot` v2.0.1 and `slingshot` v1.8.0. PAGA, DPT and geometry calculations used Scanpy v1.11.5 and AnnData v0.11.4; exact nearest neighbours used scikit-learn v1.7.2. Monocle 3 analyses used Monocle 3 v1.4.26. Geometry and association calculations used Python v3.10.19, NumPy v2.2.6, pandas v2.3.3 and SciPy v1.15.3. All stochastic stages used recorded deterministic seeds; the canonical rank bootstrap used seed 20260723, the Friedman method-label permutation used seed 20260724 and the $R_{NX}$ association analysis used seed 20260811. The complete 9-dataset × 4-framework × 17-representation grid contained 612 blocks (153 per backend), with finite values for every applicable metric.

#### 1.8 Gene-level evaluation

##### *1.8.1 Eligibility and output roles*

We distinguished static gene representations, contextual gene representations and model-native regulatory outputs because they encode different information and support different biological questions. Eleven methods contributed static gene or token representations: GenePT-w, scGPT, Geneformer, scPRINT, CellPLM, UCE, scFoundation, scCello, scBERT, CellFM and LangCell. Nine contributed contextual gene representations computed from expression data. Geneformer and scPRINT contributed the two completed model-native regulatory outputs. Static and contextual vectors were evaluated through gene–gene similarity, whereas the native outputs retained their method-specific directed, unsigned scores. The three output classes were analysed separately rather than combined into a single gene-level ranking. The static scBERT representation is the gene2vec matrix and therefore coincides with the gene2vec reference.

##### 1.8.2 Contextual reference, gene identities and similarity

For gene-function and curated-regulon analyses, contextual states were calculated from a fixed 112-cell Tabula Sapiens v2 reference. The reference contained 16 cells from each of seven compartments (immune, stromal, epithelial, endothelial, muscle, neural and germline), spanning all 26 analysed tissues and 55 native cell types. Cells were balanced over tissue and cell type within each compartment. Perturbation-response retrieval instead used 112 control cells selected independently within each perturbation dataset.

The evaluation universe was fixed before contextual extraction as the ordered identity intersection of the 11 static vocabularies (16,508 genes). Ensembl identifiers were the primary identities, with symbols retained as annotations. Task-specific resources were intersected only after this universe was fixed. The primary contextual vector for each gene was the arithmetic mean over all reference cells in which the method retained that gene. Similarities involving an unavailable contextual state received the neutral score zero while retaining the fixed candidate universe.

Within each task, vectors were centred over the eligible genes and normalized to unit length before cosine similarity was calculated. Genes tied at the neutral score were assigned their expected relevance within the tied set.

##### 1.8.3 Protein-interaction and functional-set retrieval

Protein–protein-interaction recovery used STRING v12 physical links with score at least 700. The primary 1:5 analysis contained 40,000 positive edges and 200,000 degree-matched non-interacting pairs; a 1:1 negative ratio was retained as a sensitivity. All annotated positive physical interactions were excluded from the negative pool. The PPI task contained 9,841 genes. Pairs were ranked by gene-vector cosine similarity and evaluated by AUROC and AUPRC. The STRING combined-functional network was secondary because its text-mining, co-expression, database and homology channels can overlap information used to construct or pretrain some representations.

Functional retrieval used Gene Ontology Biological Process, Reactome and KEGG annotations, with 12,423, 9,142 and 6,988 eligible genes, respectively. For each resource, 1,200 query genes were sampled; all other eligible genes were ranked by cosine similarity, and a candidate was relevant if it shared at least one retained annotation with the query. We report mean average precision and precision at 10, 50 and 100. Co-expression, gene2vec and a random representation were included as references. Percentile intervals used 1,000 resamples of gene pairs for PPI and query genes for retrieval.

##### 1.8.4 BEELINE regulatory-edge prioritization

Regulatory-edge recovery was evaluated on the seven experimental BEELINE datasets [11]: hESC, hHep, mDC, mESC, mHSC-E, mHSC-GM and mHSC-L. These datasets represent five independent studies; the three mHSC datasets belong to GSE81682. For each dataset, we retained all transcription factors passing the BEELINE nominal $P < 0.01$ criterion after Bonferroni correction and at most 1,000 significant non-transcription-factor genes ranked by variance. The candidate universe comprised directed transcription-factor-to-target pairs among the selected genes and was identical across methods within a dataset. Static representations and the classical algorithms GENIE3, GRNBoost2 and PIDC were evaluated on all seven datasets. Co-expression and random rankings were additional references.

Contextual representations and native outputs were evaluated on the four datasets with count matrices traceable to the BEELINE cells: hESC, mHSC-E, mHSC-GM and mHSC-L. Models received the complete mapped count matrix; BEELINE gene selection was applied only at extraction and evaluation.

Mouse genes were mapped through one-to-one protein-coding orthology from MGI to HGNC identities. Static and contextual vectors produced symmetric, task-centred cosine rankings. Geneformer native scores were obtained by singleton token deletion through InSilicoPerturber and target contextual-state response; scPRINT used its GNInfer query–key attention network. Both native routes produced directed, unsigned within-method rankings.

The primary outcomes were AUPRC divided by candidate-edge prevalence and the early-precision ratio, defined analogously relative to prevalence. The three GSE81682 datasets were first averaged within study, after which independent studies received equal weight. Static and classical summaries therefore contained five studies, and contextual and native summaries contained two.

###### *1.8.5 Perturbation-response retrieval*

We evaluated retrieval of perturbation-associated response genes using 570 eligible single-transcription-factor perturbations: 43 Norman CRISPRa conditions [12], 249 Replogle K562 CRISPRi conditions and 278 Replogle RPE1 CRISPRi conditions [13]. For each perturbation, positives were the 100 genes with the largest absolute change in normalized mean expression relative to control, excluding the perturbed transcription factor. Static scores came from the exported gene vectors. Contextual scores were computed independently for each dataset from 112 selected control cells; no perturbed cell was supplied during contextual extraction. The final candidate universes contained 16,301 genes for Norman, 7,558 for K562 and 7,867 for RPE1. Undefined contextual similarities received neutral score zero without removing candidates.

We report AUROC and AUPRC divided by positive prevalence. Eligible transcription-factor units were first averaged within each dataset, K562 and RPE1 were then averaged within Replogle 2022, and Norman 2019 and Replogle 2022 received equal weight. This endpoint measures recovery of genes with large observed responses, including both direct and downstream effects.

###### *1.8.6 Curated-regulon recovery*

Curated-regulon recovery used TRRUST, ChEA and DoRothEA ABC. For each transcription factor with at least five recorded targets, all other genes in the fixed 16,508-gene universe were ranked by transcription-factor-to-gene cosine similarity. The final panels contained 319, 272 and 295 eligible transcription factors for TRRUST, ChEA and DoRothEA, respectively. Contextual representations used the all-retained-cell states from the 112-cell pan-tissue reference. Per-transcription-factor AUROC distributions were reported for 11 static representations, nine contextual representations, co-expression and chance.

##### 1.9 Cell-level double-perturbation response prediction

Cell-level perturbation prediction was evaluated on the Norman CRISPRa data using the train–test design of Ahlmann-Eltze *et al.* [14]. All single-gene perturbations and one half of the double perturbations were included in the training set; the remaining double perturbations were held out for evaluation. The final split contained 154 training, 13 validation and 65 test conditions. For each of the 13 scFMs, cell embeddings were averaged within perturbation condition to form condition-level representations. The target was the condition-level mean expression change relative to control for the 1,000 genes with the highest mean control

**Table S4:** Cross-domain transfer inventory. Candidate denotes all constructed directed contrasts; fine and coarse denote eligibility after their respective shared-label rules and before the per-task ten-cell threshold. The shift strata are operational and may contain co-varying factors.

| Shift stratum | Operational source–target pairing | Candidate | Fine | Coarse |
|---|---|---|---|---|
| cross-dataset | TSv1 ↔ TSv2 within tissue | 44 | 44 | 44 |
| cross-donor | largest donor → other donors | 56 | 55 | 55 |
| cross-chemistry | 10× 3′ ↔ 10× 5′ | 20 | 20 | 20 |
| cross-platform | 10× ↔ Smart-seq2 | 48 | 44 | 46 |
| cross-tissue | larger → smaller tissue over shared lineages | 16 | 16 | 16 |
| Total | | 184 | 179 | 181 |

expression, excluding the directly perturbed genes from the primary metric.

Decoder fitting was nested within the training conditions. Each representation was standardized using training-condition means and variances, projected to 30 dimensions using a projection fitted only to the training conditions, and decoded by ridge regression. The ridge penalty was selected on the validation conditions from $\{0.01, 0.1, 1, 10, 100, 1000\}$. The fitted standardization, projection and decoder were then applied to held-out double-perturbation conditions. The additive reference summed the observed mean-expression responses of the two corresponding single-gene perturbations. Performance was summarized by normalized L2 error, $\|\widehat{\Delta} - \Delta\|_2/\|\Delta\|_2$, across held-out double perturbations. Squared Pearson correlation between predicted and observed gene-wise responses was used for the CEBPA+CEBPE illustration.

##### 1.10 Extended methods: cross-domain transfer

**Domain-shift design.** We evaluated transfer across five operational shift strata: dataset release, donor, chemistry, platform and tissue. Donor, assay, chemistry and release can co-vary within a contrast. The candidate inventory comprised 184 directed source-to-target pairs. After label-eligibility filtering, 179 pairs were available for fine-label analysis and 181 for coarse-label analysis (Table S4). Within an eligible task, analysis was restricted to labels represented by at least ten cells in both source and target domains. All transformations and supervised probe fitting for inductive methods used source cells only. Query features, but not query labels, were accessed by the explicitly transductive comparators described below.

**Label harmonization and task construction.** Cell-type labels were harmonized through Cell Ontology identifiers. The fine-grained label was the canonical name associated with `cell_type_ontology_term_id`, with a normalized dataset label used only when an ontology identifier was unavailable; a coarse broad-cell-class label was defined analogously. The eligible label universe was constructed independently for every directed pair. Fine-label analysis required at least three eligible shared labels and coarse-label analysis at least two, followed by the requirement of at least ten source and ten target cells per retained label. The primary closed-set analysis retained only shared labels. A separate open-set view retained target-only labels and measured forced annotation under an incomplete source vocabulary. Novel-type detection was evaluated separately using confidence- and distance-based scores.

**Primary probe and annotation comparators.** For embedding-based methods, the primary annotation probe was unweighted multinomial logistic regression ($L_2$, $C = 1$, `max_iter=2000`) after a source-fit `StandardScaler`. Balanced logistic regression, 15-nearest-neighbour classification with cosine distance, and nearest-prototype classification using cosine similarity were sensitivity analyses. Source training was capped at 200 cells per class by stratified sampling with seed 0, and predictions were made for the full target. CellTypist and SingleR used their native annotation procedures. scANVI was fitted de novo to source and unlabelled query features, with query labels withheld, and returned native predictions; it is therefore a transductive comparator rather than a frozen-embedding probe.

**Variational and PCA baselines.** Pretrained scVI denotes the CELLxGENE Census 2024-07-01 model, aligned through Ensembl identifiers and queried without parameter updating to obtain a 50-dimensional representation. It is distinct from scVI trained de novo as a target-data representation or integration method. scANVI was fitted de novo jointly to source and unlabelled query cells. Inductive PCA used source-only library-size normalization, 2,000 source-selected highly variable genes, scaling and 50 principal components before projection of target cells. PCA(transductive) estimated its feature transform jointly from capped source and unlabelled query features without query labels. It was retained as a feature-transductive comparator in the 19-method summary and interpreted separately from the inductive methods.

**Transfer endpoints and retention.** The principal endpoint was macro-F1 over the direction-specific shared fine-label set; accuracy, balanced accuracy and per-class recall were reported as complementary endpoints. Closed-set reference-scarce recall was calculated over shared labels comprising less than 1% of reference cells, with 5% used as a sensitivity threshold. When no shared class met a threshold, the least frequent shared reference class was used. Retention was defined as target-transfer macro-F1 divided by five-fold stratified within-source cross-validation macro-F1 using the same representation and probe. Ratios were summarized within method and operational shift stratum.

**Label efficiency.** Label efficiency was evaluated separately on the 44 directed cross-dataset tasks. For each shared fine label, we sampled up to $N \in \{1, 5, 10, 50, 100, 200, 500\}$ labelled source cells, using all available cells when a class contained fewer than $N$, and repeated the sampling ten times. Each classifier was evaluated on all target cells in the shared fine-label set at the target's natural frequencies. The method-specific threshold was the smallest tested budget at which mean target macro-F1 reached 90% of that method's mean macro-F1 at $N = 500$. Low-label reference-scarce recall used the source/reference abundance definitions above, calculated before budget sampling. All 44 directed cross-dataset transfers were included in the label-efficiency comparison.

**Novel-type detection and graceful degradation.** The novelty analysis began from the 44 directed cross-dataset tasks. Target cells were known when their harmonized label occurred in the source and novel when it did not. Thirty-seven directions contained both known and novel target cells and supported AUROC, AUPR and FPR95. We compared maximum-softmax, maximum-logit, energy, ReAct, Mahalanobis, k-nearest-neighbour-distance and ViM scores where the required outputs were available. Mahalanobis distance defined the common cross-embedding summary. Embeddings were standardized using source statistics; source class means and a shared pooled covariance matrix with a $10^{-4}$ ridge term defined the minimum squared whitened distance from each target cell to a source class. CellTypist and SingleR lack embeddings for this calculation and contributed their available native confidence or similarity scores.

To describe the biological proximity of forced closed-set assignments, we analysed all 18,391 novel target cells without first applying an OOD-score threshold. We recorded whether each predicted source label shared a broad category, a compartment or an annotated ancestor relation with the target-only label.

##### 1.11 Extended methods: model-specific adaptation

We evaluated five adaptation regimes across 13 scFMs. scGPT, LangCell and CellPLM updated the full backbone. CellFM, Geneformer, scBERT and scFoundation updated selected backbone components. CELLama adapted its source-domain sentence encoder before fitting an MLP classifier. scCello retained its pretrained encoder and trained a linear head. SCimilarity, UCE, scPRINT and GenePT-w retained fixed representations and trained a shared two-layer MLP. No query labels were used for adaptation. Table S5 reports the trainable components, optimization and source-validation rule for each method.

**Table S5:** Model-specific supervised adaptation procedures for cross-dataset cell-type annotation. Validation used labelled source cells only; query labels were withheld throughout.

| Method | Trainable components | Optimization and training budget | Validation and model selection |
|---|---|---|---|
| CellFM | Encoder parameters and native dual annotation head | 30 epochs; batch 8; AdamW, lr $10^{-4}$, weight decay $10^{-5}$; StepLR ($\gamma = 0.95$); class-weighted dual-head cross-entropy | 20% stratified source validation; best validation loss; no early stopping |
| scGPT | Full backbone and classification head | 10 epochs; batch 32; Adam, lr $10^{-4}$; StepLR ($\gamma = 0.9$) | 20% stratified source validation; best validation loss |
| scBERT | Final normalization module, penultimate Performer block and classification head | 100-epoch cap; batch 8; gradient accumulation 23; Adam, lr $10^{-4}$; cosine warm-restart schedule | 20% stratified source validation; best validation accuracy; early-stopping patience 10 |
| LangCell | Full backbone and classification head | 20 epochs, extended when needed to reach 1,000 gradient steps; batch 16; AdamW, lr $5 \times 10^{-5}$, weight decay $10^{-3}$; linear decay with 10% warm-up | 20% stratified source validation; best validation loss |
| Geneformer | Backbone except the first two frozen layers, and classification head | $\max(1000, \text{one-epoch steps})$; batch 4; AdamW, lr $5 \times 10^{-5}$, weight decay $10^{-2}$; cosine decay with 10% warm-up | 20% stratified source validation; best validation macro-F1 |
| scCello | Frozen pretrained encoder and trained linear head | 20 epochs, extended when needed to reach 1,000 gradient steps; batch 24; AdamW, lr $5 \times 10^{-3}$, weight decay $10^{-3}$; linear decay with 10% warm-up | 20% stratified source validation; best validation macro-F1 |
| CellPLM | Full native annotation model | 2,000-epoch cap; lr $5 \times 10^{-3}$, weight decay $10^{-7}$; ReduceLROnPlateau; 3,000 HVGs | 20% stratified source validation; early stopping from epoch 200 with patience 25 |
| scFoundation | Transformer block `[-2]` and max-pool MLP head | 20 epochs; batch 32; AdamW, peak lr $10^{-4}$, weight decay $10^{-2}$; 10% warm-up followed by cosine decay to $10^{-5}$ | 20% stratified source validation; best validation accuracy; early-stopping patience 4 |
| CELLama | Sentence encoder followed by a shared MLP | 20 source-domain encoder-adaptation epochs over sentence configurations with $k \in \{16, 20, 24, 28\}$ and 500 or 1,200 HVGs; batch 32; lr $10^{-5}$; cells then embedded at $k = 30$ and classified by the shared MLP | 1,000 validation examples for encoder adaptation; 20% stratified validation for the MLP |
| SCimilarity | Fixed representation and shared MLP | AdamW, lr $10^{-3}$, weight decay $10^{-4}$; cosine decay with 0.5% warm-up; batch 64; at most 50,000 steps | 20% stratified source validation; best validation loss; early-stopping patience 10 |
| UCE | Fixed representation and shared MLP | AdamW, lr $10^{-3}$, weight decay $10^{-4}$; cosine decay with 0.5% warm-up; batch 64; at most 50,000 steps | 20% stratified source validation; best validation loss; early-stopping patience 10 |
| scPRINT | Fixed representation and shared MLP | AdamW, lr $10^{-3}$, weight decay $10^{-4}$; cosine decay with 0.5% warm-up; batch 64; at most 50,000 steps | 20% stratified source validation; best validation loss; early-stopping patience 10 |
| GenePT-w | Fixed representation and shared MLP | AdamW, lr $10^{-3}$, weight decay $10^{-4}$; cosine decay with 0.5% warm-up; batch 64; at most 50,000 steps | 20% stratified source validation; best validation loss; early-stopping patience 10 |

Table S6 relates each executed procedure to the corresponding author-released training guidance. Exact optimization and validation settings are reported in Table S5.

**Table S6:** Relationship between author-released supervised-training guidance and the adaptation procedures used for cross-dataset annotation.

| Method | Released guidance | Implementation used in this study |
|---|---|---|
| CellFM | MindSpore annotation head using cluster prototypes, a CrossRetentionLayer and dual logits | The annotation head was ported to PyTorch. The 30-epoch budget and no-mask inference were retained; encoder and head parameters defined the trainable boundary. |
| scGPT | Released full-backbone cell-type annotation procedure | The released optimizer, schedule and epoch budget were retained, with the study-wide stratified source-validation split. |
| scBERT | Zheng68K annotation example with a 100-epoch cap, cosine warm-restart schedule and full-backbone updating | The optimizer, schedule and epoch cap were retained; the final normalization module, penultimate Performer block and classification head were updated. |
| LangCell | Hugging Face Trainer procedure with 20 epochs, linear decay and an absolute warm-up budget | Warm-up was set to 10% of the available steps and training was extended to at least 1,000 gradient steps for small reference datasets, retaining the released optimizer and schedule. |
| Geneformer | Step-based Hugging Face Trainer procedure of approximately one epoch with absolute warm-up | The first two layers remained frozen; the step budget was at least 1,000 and warm-up was proportional to the available steps. |
| scCello | Frozen encoder with a linear probe, 20 epochs and an absolute 500-step warm-up | The released linear-probing design was retained, with proportional warm-up and a minimum 1,000-step budget for small reference datasets. |
| CellPLM | Native full-backbone annotation procedure with a 2,000-epoch cap, delayed early stopping and ReduceLROnPlateau | The released annotation, optimization and stopping procedure was retained, with the study-wide stratified source-validation split. |
| scFoundation | Released classifier architecture and trainable scope, without a complete annotation-training loop | The indicated transformer block and max-pool classification route were retained; the optimization schedule and source-validation rule completed the training procedure. |
| CELLama | Source-sentence construction and sentence-encoder adaptation utilities, without an end-to-end cell-type annotation head | The sentence encoder was adapted on source-domain cell sentences before fitting the shared MLP to the resulting source representations. |
| SCimilarity, UCE, scPRINT and GenePT-w | Released cell-representation interfaces without a model-specific supervised annotation procedure used here | The pretrained representations remained fixed and the same two-layer MLP and training rule were applied to all four methods. |

The shared MLP comprised LayerNorm, dropout (0.1), a hidden linear layer from $d$ to $d$ dimensions, GELU, dropout (0.1), and an output layer. For each method and transfer direction, adapted predictions were aligned to the corresponding frozen-representation predictions on common query cells. The adaptation effect was the difference in macro-F1 between the model-specific supervised procedure and an unweighted logistic-regression probe trained on the frozen representation. Method-level effects were averaged over the same 44 directed cross-dataset transfers.

**Adaptation cost.** Resource use was measured from the same prepared 10,000-cell reference input to a serialized fitted model on NVIDIA H200 GPUs. Each method retained its model-specific training procedure and was run in five independent launches. We recorded wall-clock time, peak host resident memory and peak GPU memory and report the median across launches.

End-to-end adaptation time was also recorded for the nine model-specific adaptation methods across all 44 directed TSv1–TSv2 transfers. Each measurement was the duration of a successful Nextflow process on one NVIDIA H200, from method-specific preprocessing through adaptation and validation, query prediction and output serialization. Durations were summarized by their median and observed range across directions and by aggregate runtime.

##### 1.12 Evaluation of representations on spatial transcriptomics

**Datasets and endpoints.** We analysed the prepared spatial-transcriptomics panel used throughout this study: 12 SpatialLIBD Visium sections (47,681 spots) [15], eight HER2ST sections (3,481 spots) [16] and five MERFISH z-sections (28,317 cells) [17], for a total of 25 sections and 79,479 observations. SpatialLIBD and HER2ST were evaluated against their curated spatial-domain annotations after removing observations labelled `Unknown` or `undetermined`, respectively. MERFISH was evaluated against spatial-domain annotations in the primary 25-section panel; cell-type clustering was examined as a separate endpoint and did not enter the primary cross-section rank summary. The five MERFISH sections corresponded to z-coordinates 0.04, 0.09, 0.14, 0.19 and 0.24.

**Representation panel and model roles.** The complete comparison contained 17 methods. Thirteen were frozen scFM representations (CELLama, CellFM, CellPLM, Geneformer, GenePT-w, LangCell, scBERT, scCello, scFoundation, scGPT, SCimilarity, scPRINT and UCE). Pretrained scVI was the 50-dimensional query representation from the CELLxGENE Census 2024-07-01 Homo sapiens model, with no target-data parameter updating. A separate scVI model was trained de novo within each target section using at most 2,000 highly variable genes, a 30-dimensional latent space, two hidden layers and seed 0. HVGs+PCA was fitted within each target section after library-size normalization, log transformation, selection of at most 2,000 highly variable genes and scaling; the first 50 components were retained. NOVAE [18] was included as an SRT-native comparator and, unlike the expression-only representations, used the target spatial coordinates: the brain checkpoint was used for SpatialLIBD and MERFISH, and the human checkpoint with an eight-neighbour grid graph for HER2ST; the MERFISH graph used radius 80.

**Clustering and metrics.** Clustering was performed separately for each section and representation. The annotated class count was used to calibrate the number of clusters or the resolution producing the closest cluster count. Thirteen raw metrics were calculated. The primary summary used 11: ARI, NMI, homogeneity, completeness and FMI for cluster concordance; Acc@kNN and graph connectivity for local structure; 1−CHAOS and 1−PAS for spatial coherence; and Moran's I and 1−Geary's C for spatial-marker autocorrelation. ASW and cLISI were retained as diagnostics but excluded from the primary aggregate.

**Aggregation and statistical analysis.** Within each section and raw metric, the 17 methods were ranked with average ranks for ties. Metric ranks were averaged within the four families above, and the four family ranks were then averaged with equal weight. The primary overall summary averaged section-level aggregate ranks over all 25 sections; the resulting dataset weights were therefore 12/25 for SpatialLIBD, 8/25 for HER2ST and 5/25 for MERFISH rather than equal weight per study. Each of the 16 non-NOVAE methods was compared with NOVAE using a two-sided paired Wilcoxon signed-rank test over section-level aggregate ranks, with Holm correction across the 16 comparisons. Descriptive confidence intervals used 5,000 paired section-bootstrap replicates (seed 20260813). Sections, rather than cells or spots, were the inferential units.

The section-level gallery displays 15 methods for SpatialLIBD and HER2ST (the 13 frozen scFMs, PCA and NOVAE). The MERFISH panels display NOVAE, PCA and 11 frozen scFMs (CELLama, CellFM, CellPLM, GenePT-w, Geneformer, LangCell, SCimilarity, UCE, scBERT, scFoundation and scGPT). The aggregate analysis and source tables include all 17 methods.

##### 1.13 Uniform spatial adaptation with DRIFT

We used the DRIFT graph-diffusion operator [19] to test whether a common coordinate-based transformation changed spatial-domain performance and whether any improvement was confined to scFM representations. The paired analysis comprised the 12 methods with complete compatible outputs in all three datasets: scGPT, scFoundation, Geneformer, CellPLM, CellFM, CELLama, SCimilarity, LangCell, UCE, GenePT-w, scBERT and HVGs+PCA. scCello and scPRINT were excluded because their input paths rejected the non-integer diffused matrix; pretrained and de novo scVI and NOVAE did not have complete paired outputs for the common three-dataset analysis. For a section with $N$ spots or cells and spatial coordinates $\{s_i\}$, we form the dense pairwise Euclidean distance matrix $D_{ij} = \|s_i - s_j\|_2$ and a binary spatial graph by thresholding the flattened distances at their $(k_{\text{graph}}N)$-th smallest value $\tau$, as in the implemented reference operator (a global threshold rather than a node-wise $k$-nearest-neighbour graph):

$$A_{ij} = \mathbb{1}\big[\, D_{ij} \leq \tau \,\big], \qquad A_{ii} = 0, \qquad A \leftarrow \max(A, A^{\top}). \tag{S45}$$

With the combinatorial graph Laplacian $L = \mathrm{diag}(A\mathbf{1}) - A$, we diffuse the raw count matrix $X_{\text{raw}} \in \mathbb{R}^{N\times G}$ with the heat kernel,

$$X_{\text{diff}} = e^{-tL}\, X_{\text{raw}}, \tag{S46}$$

which we evaluate through its action (`scipy.sparse.linalg.expm_multiply`) without forming the $N \times N$ kernel; numerical checks against the dense matrix exponential agreed to approximately $10^{-10}$. Because the symmetric heat kernel has unit row and column sums, it preserves each gene's total count across the section and the global count total. It does not, in general, preserve each individual cell's library size. The final paired table used the evaluated variant $k_{\text{graph}} = 3$ and $t = 5$. The resulting non-negative, non-integer and typically denser count matrix was passed through each method's native preprocessing; consequently, model-specific rounding, tokenization and gene selection remained part of each method.

For the paired sensitivity, each adapted embedding was compared with its unadapted counterpart on the same observations and spatial-domain labels. We used k-means with the number of clusters fixed to the number of annotated domains, ten initializations and random state 1, and took adapted minus unadapted ARI as the sole primary outcome. This estimand is distinct from the 11-metric, 17-method unadapted spatial composite above. We report mean change and the number of positive section-level changes for each method.

### Supplementary Benchmarking Results

#### 2.1 Zero-shot benchmarking of scFMs and baseline representations

Across the fixed 20-representation panel, tissue-level biological-conservation ranks differed among methods (Friedman $\chi^2 = 357.212$, d.f. 19, $P = 3.30 \times 10^{-64}$; Kendall's $W = 0.723$). SCimilarity had the best mean biological-only metric-family rank (3.98), followed by scFoundation (6.15), PCA (6.60), CellPLM (7.14) and Harmony (7.59). In the 19 paired comparisons with PCA, SCimilarity was the only frozen scFM with a significantly better aggregate rank after Holm correction (adjusted $P = 0.0145$); scFoundation and CellPLM did not differ significantly from PCA. Batch mixing was analysed separately in the 19 multi-batch tissues and in the supplementary biological-plus-batch aggregate below; the biological-only aggregate defined the primary zero-shot comparison.

The combined-family summary, tissue stability, PCA-relative metric contrasts and metric-dependence analyses are shown in Figs. S1–S7; the corresponding biological-only tissue and leave-one-tissue-out summaries are shown in Figs. S8 and S9.

##### *2.1.1 Biological-conservation and batch-mixing metric families*

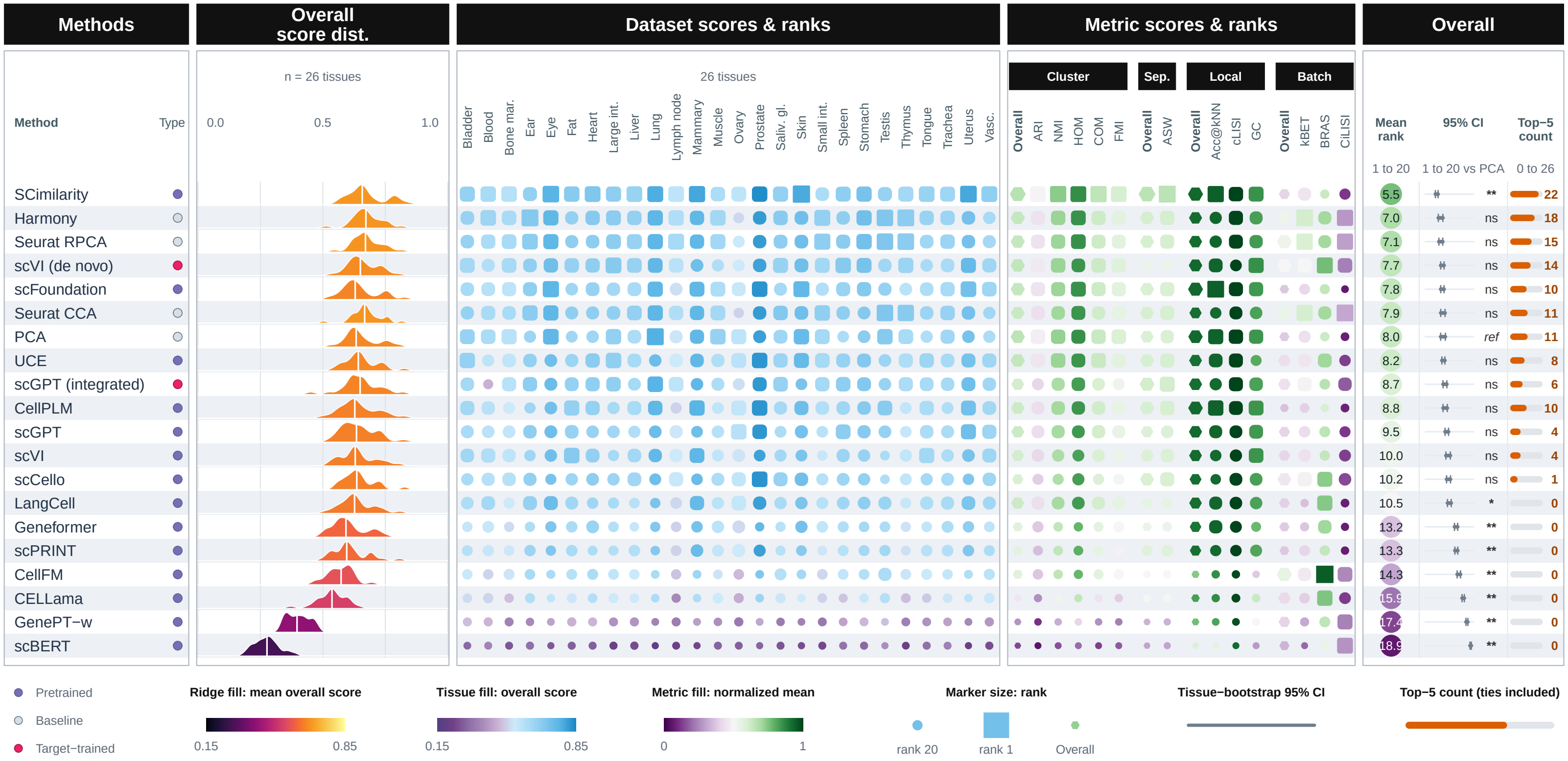

**Figure S1: Tabula Sapiens v2 embedding benchmark summary.** Rows show 20 methods across 26 tissues. Combined scores weight cluster concordance, continuous separation, local neighbourhood structure and batch mixing by 32.5%, 10%, 32.5% and 25%, with renormalization for single-batch tissues. The tissue matrix encodes within-tissue rank and score; the metric matrix shows family summaries and raw metrics. Intervals are 95% tissue-bootstrap intervals. Symbols denote two-sided paired Wilcoxon tests against PCA with Holm correction across 19 comparisons. Top-five counts include ties at the fifth-place boundary.

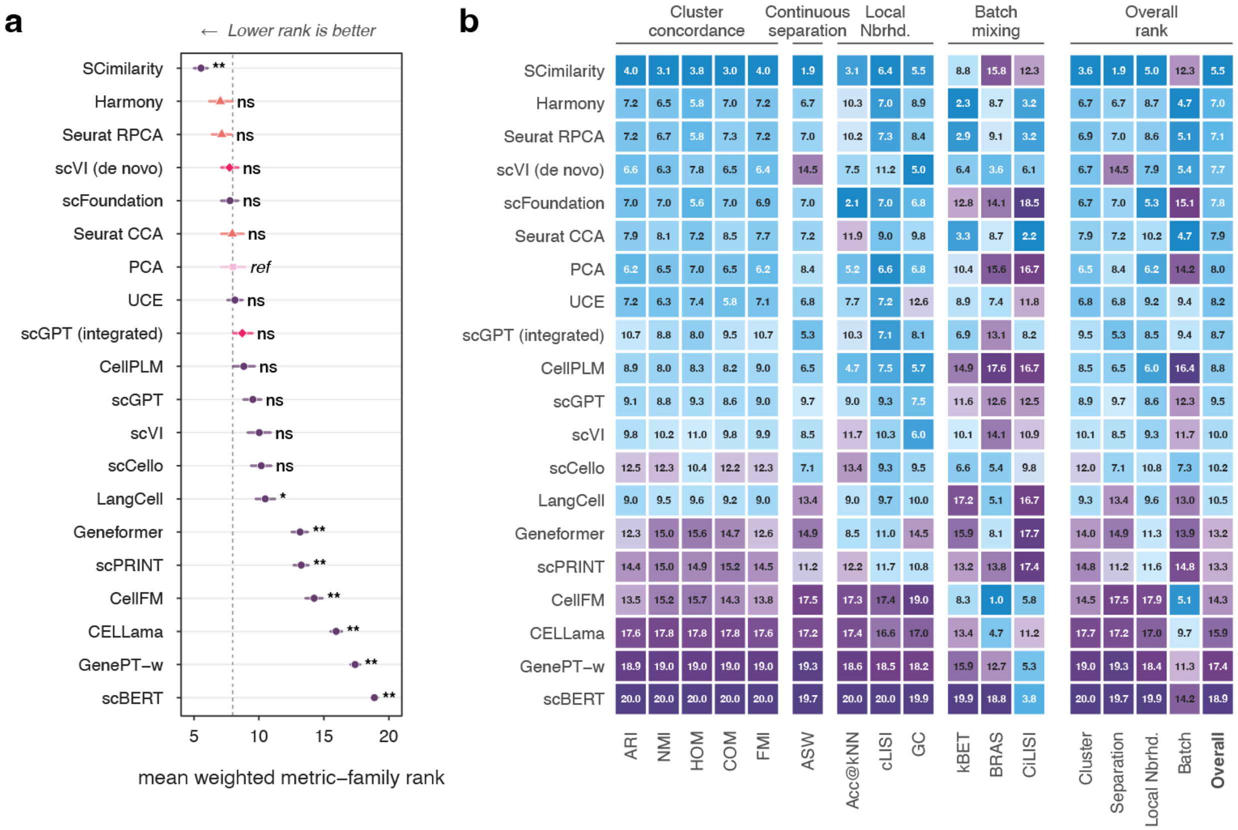

**Figure S2: Across-tissue rank heatmap for the Tabula Sapiens v2 representation benchmark**. Rows show 20 representation methods. Raw metrics are grouped into cluster concordance (ARI, NMI, HOM, COM and FMI), continuous separation (ASW), local neighbourhood structure (Acc@kNN, cLISI and GC), and batch mixing (kBET, BRAS and CiLISI). Within each tissue, methods were first ranked separately for every raw metric (higher is better), with tied values assigned their standard average rank. Cluster concordance, continuous separation, local-neighbourhood structure and batch mixing were weighted by 32.5%, 10%, 32.5% and 25%, respectively; unavailable families were excluded and the remaining weights were renormalized. Family and Overall values were not re-ranked. Raw-metric ranks and rank-derived family and Overall values were computed independently within each of 26 tissues, and cells report their across-tissue means. The final Overall rank panel reports the four family-level average ranks followed by their aggregate Overall value; lower values indicate stronger performance (1 is the best possible value). The panel comprises 13 frozen scFMs, pretrained scVI, PCA, Harmony, Seurat CCA, Seurat RPCA, scVI trained de novo and integrated scGPT. The latter five target-data methods used batch identities but no cell-type labels during fitting.

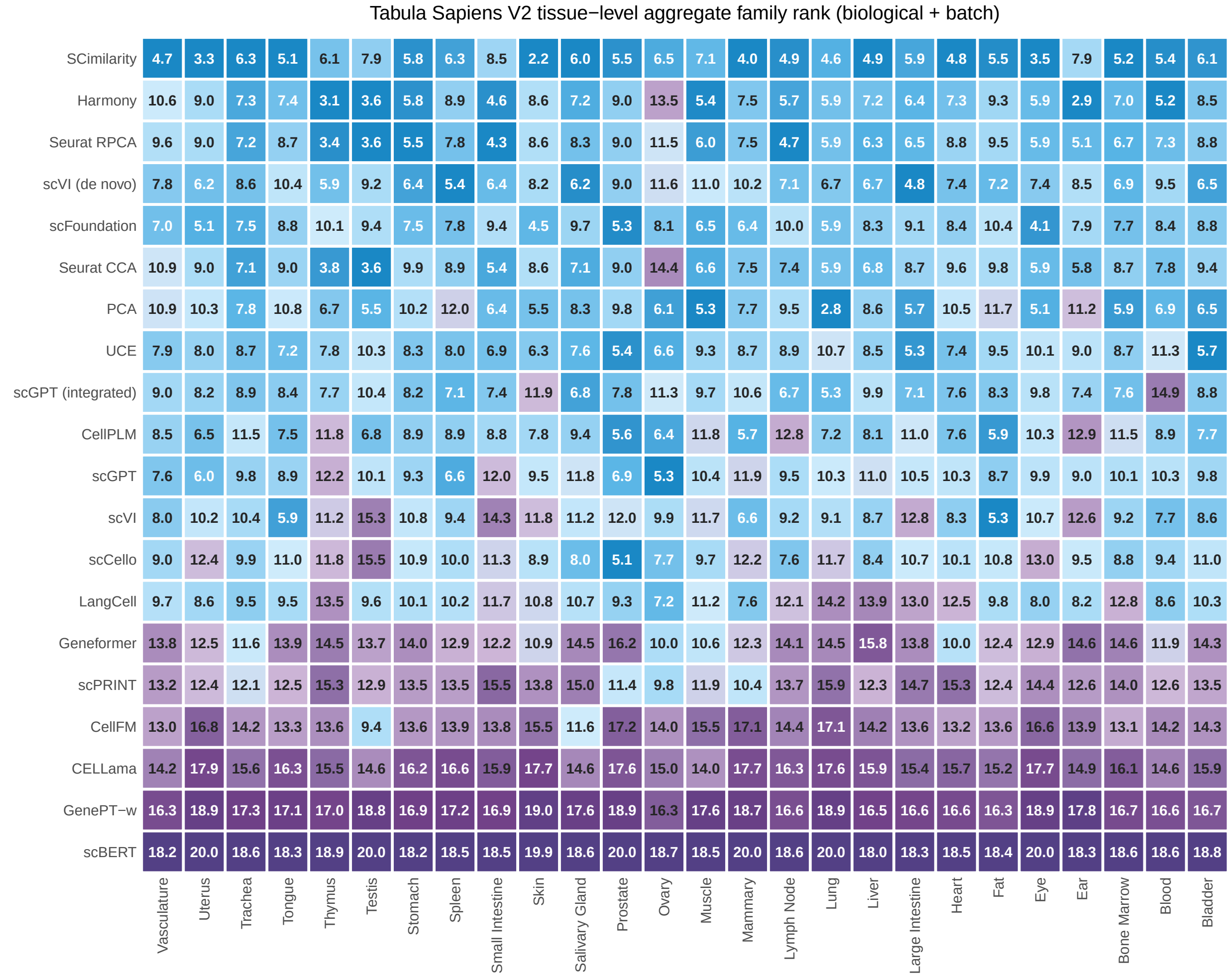

**Figure S3: TSv2 tissue-level method aggregate-rank heatmap.** Rows show 20 embedding methods and columns show 26 tissues. Each cell reports the metric-family aggregate rank for one method within one tissue; lower ranks are better. The aggregate combines cluster-label concordance, continuous cell-type separation, local cell-type neighborhood structure, and batch mixing using the prespecified family weights; weights are renormalized where batch metrics are not applicable. Methods are ordered by the full-data mean metric-family aggregate rank used in the primary TSv2 statistics. Blue indicates better within-tissue rank and purple indicates worse within-tissue rank.

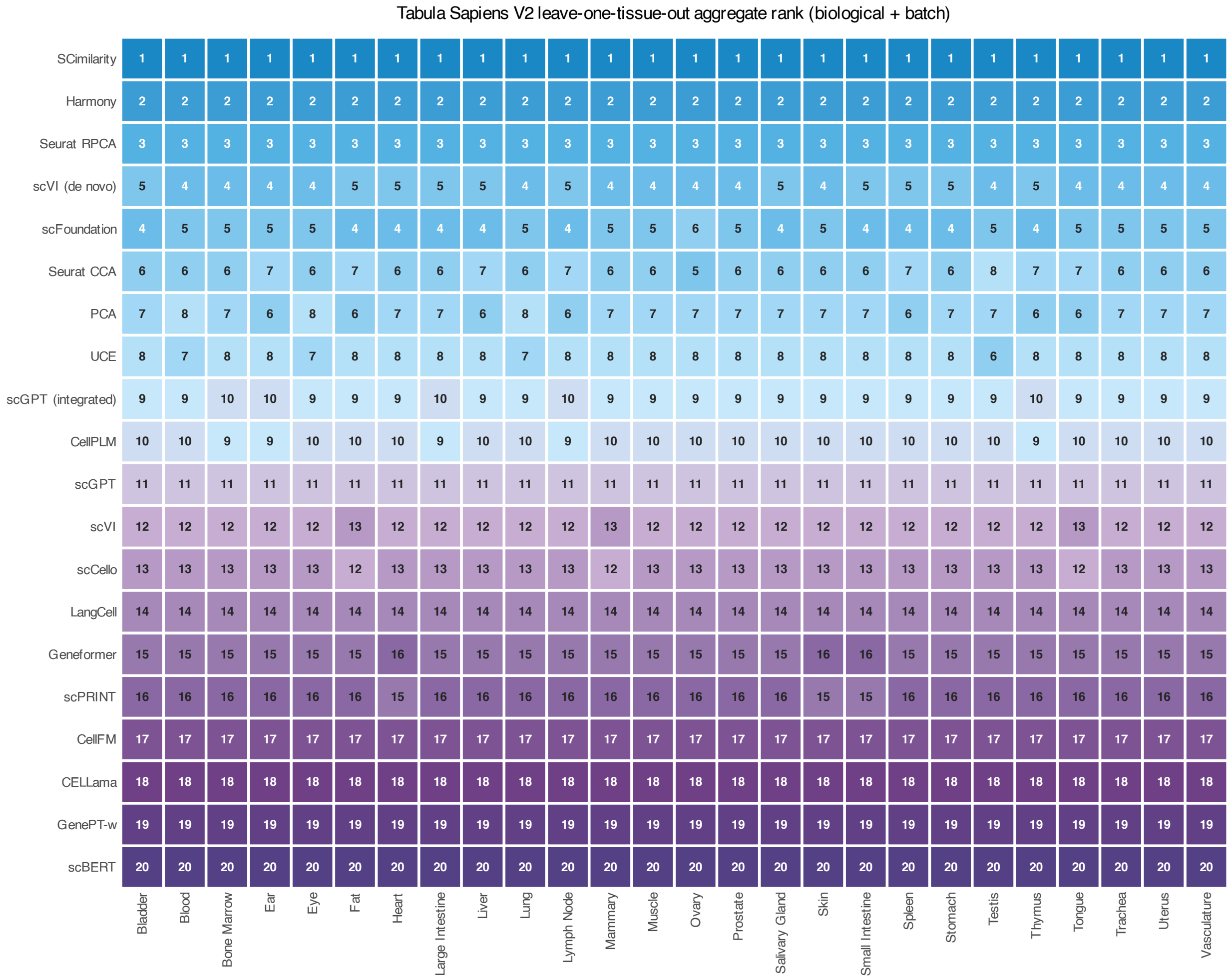

**Figure S4: Tabula Sapiens V2 leave-one-tissue-out aggregate-rank stability heatmap**. Rows show 20 embedding methods and columns show 26 omitted tissues. Each cell reports the method's full benchmark aggregate re-rank after removing the indicated tissue; lower ranks are better. Ranks aggregate the three biological-conservation families and batch mixing using the prespecified family weights, with renormalization where batch metrics are not applicable. Methods are ordered by the full-data mean metric-family aggregate rank used in the primary TSv2 statistics. Blue indicates better rank and purple indicates worse rank.

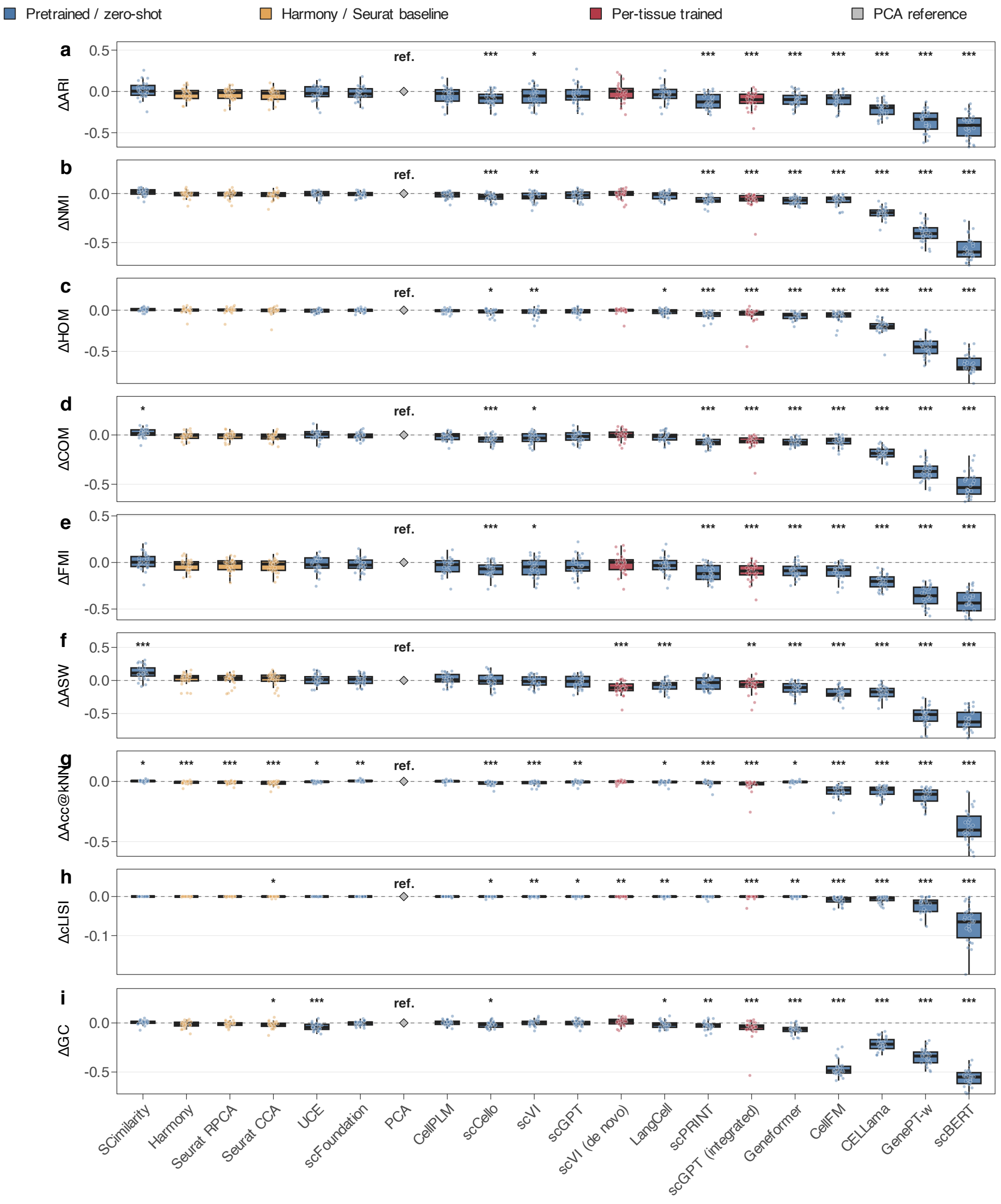

**Figure S5: Paired biological-conservation differences relative to PCA across Tabula Sapiens v2 tissues.** Panels show Leiden cluster concordance metrics (ARI, NMI, homogeneity, completeness and FMI), continuous cell-type separation (ASW), and local-neighbourhood metrics (Acc@kNN, cLISI and graph connectivity). For every tissue and metric, the corresponding PCA score was subtracted from each method's score; each point therefore represents one matched tissue (n=26 tissues per method). Positive values indicate improvement over PCA, boxes show the median and interquartile range, whiskers extend to 1.5 times the interquartile range, and the dashed horizontal line denotes no difference from PCA. Asterisks indicate two-sided paired Wilcoxon signed-rank tests after Benjamini–Hochberg correction across all 228 finite method-by-metric comparisons in the biological-conservation and batch-mixing analyses ((*$P < 0.05$), (**$P < 0.01$), (***$P < 0.001$)).

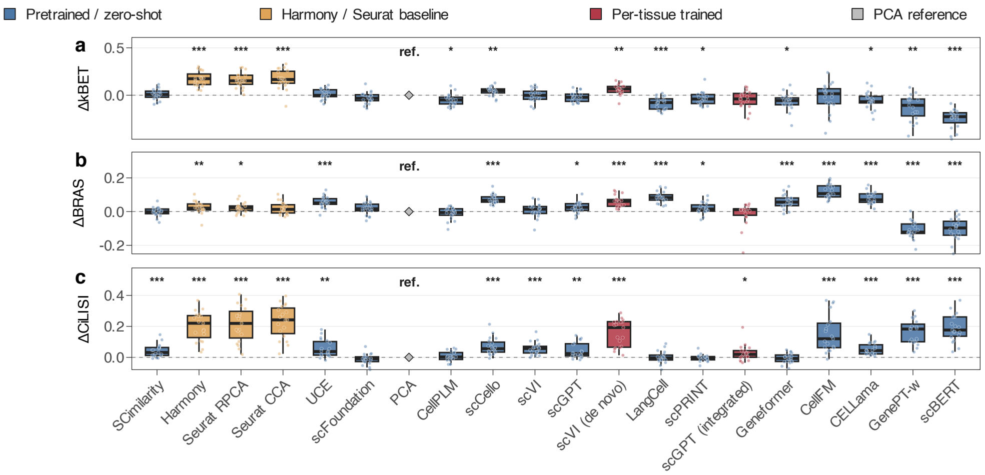

**Figure S6: Paired batch-mixing differences relative to PCA across multi-batch Tabula Sapiens v2 tissues.** Panels show full-cell kBET, BRAS and CiLISI. For every multi-batch tissue and metric, the corresponding PCA score was subtracted from each method's score; each point represents one matched tissue (n=19 tissues per method). Single-batch tissues, for which batch-mixing metrics are not applicable, were excluded. Positive values indicate improvement over PCA, boxes show the median and interquartile range, and the dashed horizontal line denotes no difference from PCA. Asterisks indicate two-sided paired Wilcoxon signed-rank tests after Benjamini–Hochberg correction across all biological-conservation and batch-mixing method-by-metric comparisons ((*$P$ < 0.05), (**$P$ < 0.01), (***$P$ < 0.001)).

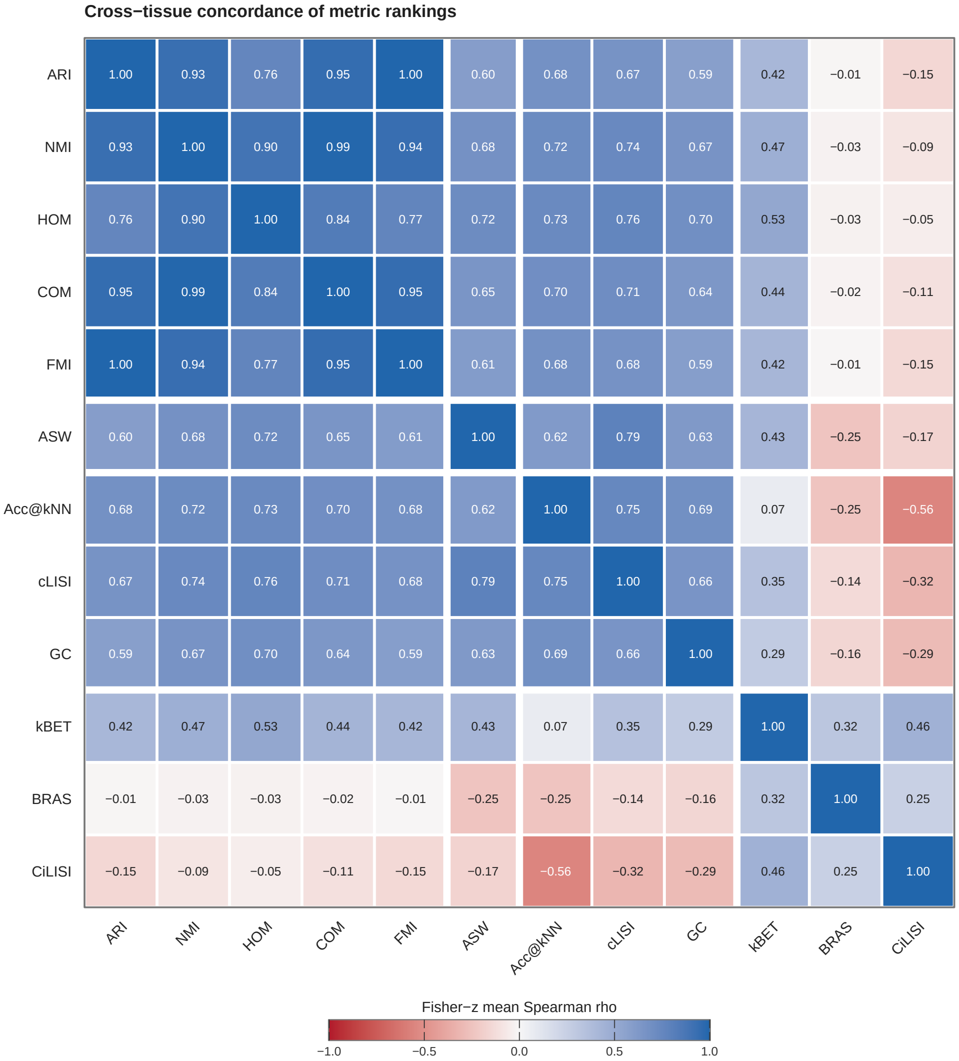

| | ARI | NMI | HOM | COM | FMI | ASW | Acc@kNN | cLISI | GC | kBET | BRAS | CiLISI |
|---|---|---|---|---|---|---|---|---|---|---|---|---|
| ARI | 1.00 | 0.93 | 0.76 | 0.95 | 1.00 | 0.60 | 0.68 | 0.67 | 0.59 | 0.42 | −0.01 | −0.15 |
| NMI | 0.93 | 1.00 | 0.90 | 0.99 | 0.94 | 0.68 | 0.72 | 0.74 | 0.67 | 0.47 | −0.03 | −0.09 |
| HOM | 0.76 | 0.90 | 1.00 | 0.84 | 0.77 | 0.72 | 0.73 | 0.76 | 0.70 | 0.53 | −0.03 | −0.05 |
| COM | 0.95 | 0.99 | 0.84 | 1.00 | 0.95 | 0.65 | 0.70 | 0.71 | 0.64 | 0.44 | −0.02 | −0.11 |
| FMI | 1.00 | 0.94 | 0.77 | 0.95 | 1.00 | 0.61 | 0.68 | 0.68 | 0.59 | 0.42 | −0.01 | −0.15 |
| ASW | 0.60 | 0.68 | 0.72 | 0.65 | 0.61 | 1.00 | 0.62 | 0.79 | 0.63 | 0.43 | −0.25 | −0.17 |
| Acc@kNN | 0.68 | 0.72 | 0.73 | 0.70 | 0.68 | 0.62 | 1.00 | 0.75 | 0.69 | 0.07 | −0.25 | −0.56 |
| cLISI | 0.67 | 0.74 | 0.76 | 0.71 | 0.68 | 0.79 | 0.75 | 1.00 | 0.66 | 0.35 | −0.14 | −0.32 |
| GC | 0.59 | 0.67 | 0.70 | 0.64 | 0.59 | 0.63 | 0.69 | 0.66 | 1.00 | 0.29 | −0.16 | −0.29 |
| kBET | 0.42 | 0.47 | 0.53 | 0.44 | 0.42 | 0.43 | 0.07 | 0.35 | 0.29 | 1.00 | 0.32 | 0.46 |
| BRAS | −0.01 | −0.03 | −0.03 | −0.02 | −0.01 | −0.25 | −0.25 | −0.14 | −0.16 | 0.32 | 1.00 | 0.25 |
| CiLISI | −0.15 | −0.09 | −0.05 | −0.11 | −0.15 | −0.17 | −0.56 | −0.32 | −0.29 | 0.46 | 0.25 | 1.00 |

**Figure S7: Cross-tissue concordance of metric rankings in Tabula Sapiens V2.** Within each tissue, Pearson correlation of the precomputed average-tie ranks across 20 methods was used to obtain Spearman rank correlation for every metric pair. Tissue-specific correlations were combined after Fisher z transformation with weights equal to the number of common methods minus three; cells show the resulting back-transformed mean Spearman rho. Positive values indicate concordant method rankings and negative values indicate opposing rankings. Biological-metric pairs used all available tissues, whereas pairs containing kBET, BRAS, or CiLISI used only tissues for which batch mixing was applicable. Metrics retain the prespecified family order, and white rules delineate cluster concordance, continuous separation, local-neighbourhood structure, and batch mixing.

###### *2.1.2 Biological-conservation metric families*

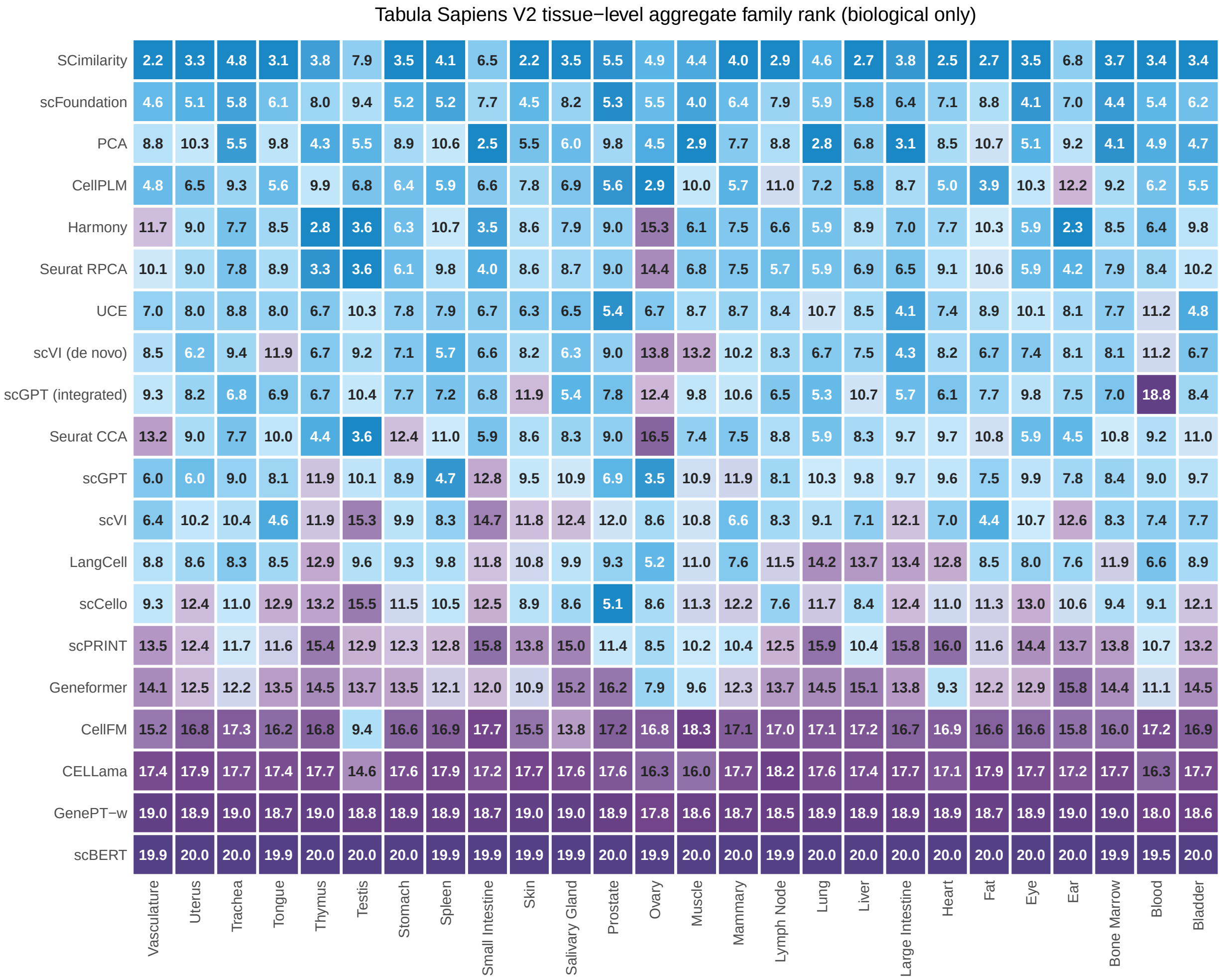

**Figure S8: TSv2 tissue-level method aggregate-rank heatmap.** Rows show 20 embedding methods and columns show 26 tissues. Each cell reports the metric-family aggregate rank for one method within one tissue; lower ranks are better. The aggregate combines the three biological-conservation families using the prespecified family weights renormalized over those families. Methods are ordered by the full-data mean metric-family aggregate rank used in the primary TSv2 statistics. Blue indicates better within-tissue rank and purple indicates worse within-tissue rank.

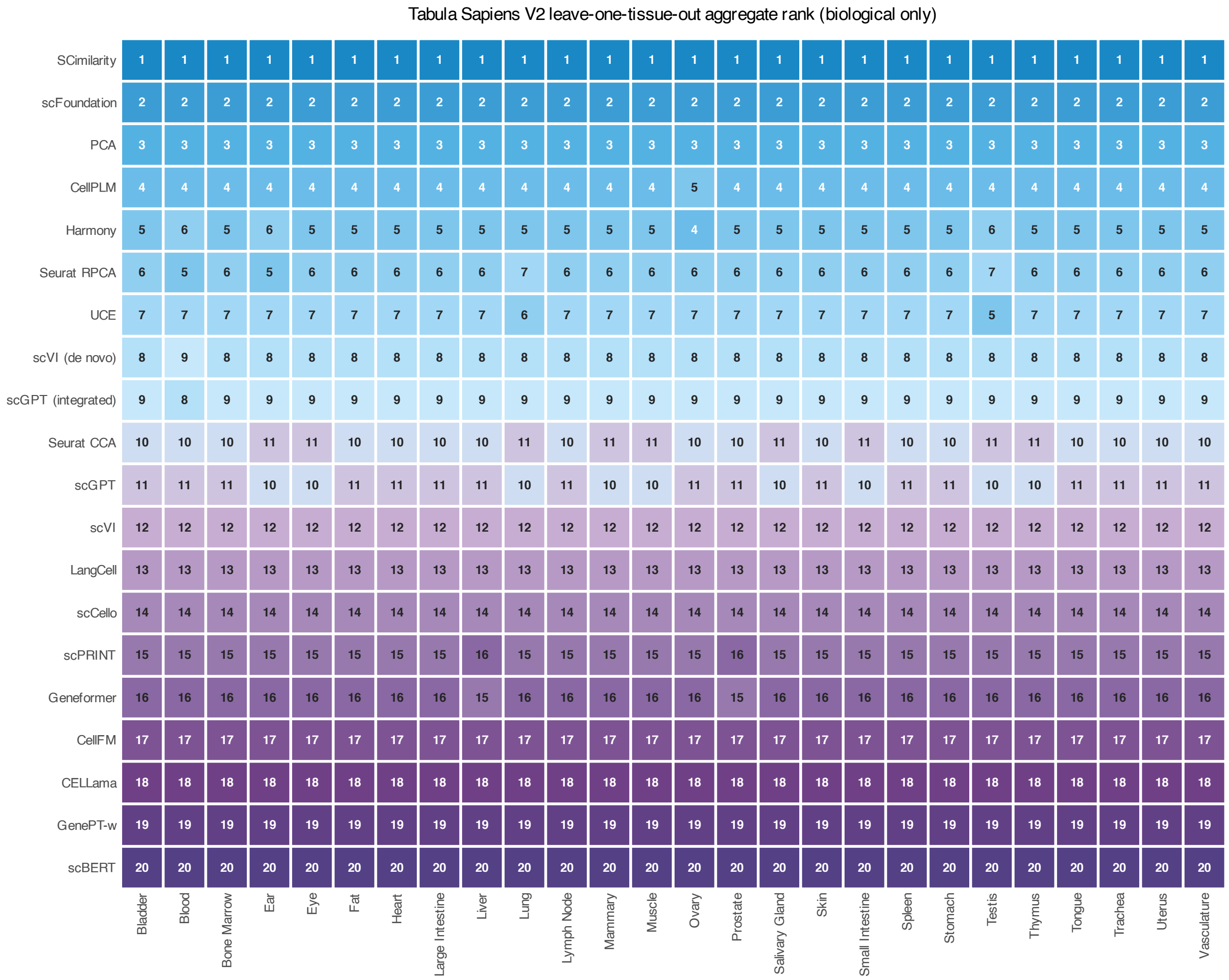

**Figure S9: Tabula Sapiens V2 leave-one-tissue-out aggregate-rank stability heatmap**. Rows show 20 embedding methods and columns show 26 omitted tissues. Each cell reports the method's full benchmark aggregate re-rank after removing the indicated tissue; lower ranks are better. Ranks aggregate the three biological-conservation families using the prespecified family weights. Methods are ordered by the full-data mean metric-family aggregate rank used in the primary TSv2 statistics. Blue indicates better rank and purple indicates worse rank.

##### 2.1.3 *UMAP visualization and detailed metric scores*

The across-tissue metric summary is shown in Fig. S10; the complete tissue gallery then presents paired cell-type and batch UMAPs and the corresponding score table for every analysed tissue (Figs. S11–S88). Cell-type and batch views reuse the same embedding coordinates within each method–tissue panel and differ only in the displayed annotation.

**Tabula Sapiens v2 — Average across tissues**

| | Biological conservation metrics | | | | | | | | | Batch mixing metrics | | | Aggregated scores | | | | |
|---|---|---|---|---|---|---|---|---|---|---|---|---|---|---|---|---|---|
| method | ARI | NMI | HOM | COM | FMI | ASW | Acc@kNN | cLISI | GC | kBET | BRAS | CiLISI | Cluster concordance | Continuous separation | Local structure | Batch mixing | Overall |
| Seurat RPCA | 0.44 | 0.71 | 0.85 | 0.62 | 0.56 | 0.21 | 0.93 | 1.00 | 0.83 | 0.60 | 0.70 | 0.29 | 0.64 | 0.21 | 0.92 | 0.53 | 0.66 |
| Harmony | 0.44 | 0.71 | 0.85 | 0.62 | 0.56 | 0.22 | 0.93 | 1.00 | 0.83 | 0.61 | 0.70 | 0.28 | 0.64 | 0.22 | 0.92 | 0.53 | 0.66 |
| Seurat CCA | 0.44 | 0.70 | 0.85 | 0.61 | 0.56 | 0.21 | 0.93 | 1.00 | 0.82 | 0.61 | 0.70 | 0.31 | 0.63 | 0.21 | 0.92 | 0.54 | 0.66 |
| SCimilarity | 0.49 | 0.73 | 0.86 | 0.65 | 0.60 | 0.32 | 0.94 | 1.00 | 0.85 | 0.45 | 0.63 | 0.12 | 0.67 | 0.32 | 0.93 | 0.40 | 0.65 |
| scVI (de novo) | 0.46 | 0.71 | 0.85 | 0.63 | 0.58 | 0.08 | 0.94 | 1.00 | 0.86 | 0.50 | 0.77 | 0.23 | 0.65 | 0.08 | 0.93 | 0.50 | 0.65 |
| UCE | 0.45 | 0.71 | 0.85 | 0.63 | 0.57 | 0.21 | 0.94 | 1.00 | 0.80 | 0.45 | 0.71 | 0.13 | 0.64 | 0.21 | 0.91 | 0.43 | 0.63 |
| scGPT (integrated) | 0.41 | 0.69 | 0.83 | 0.60 | 0.53 | 0.23 | 0.92 | 1.00 | 0.83 | 0.49 | 0.66 | 0.17 | 0.61 | 0.23 | 0.92 | 0.44 | 0.63 |
| PCA | 0.47 | 0.72 | 0.85 | 0.63 | 0.59 | 0.19 | 0.94 | 1.00 | 0.84 | 0.44 | 0.63 | 0.07 | 0.65 | 0.19 | 0.93 | 0.38 | 0.63 |
| scCello | 0.39 | 0.68 | 0.83 | 0.59 | 0.51 | 0.20 | 0.92 | 1.00 | 0.83 | 0.48 | 0.73 | 0.14 | 0.60 | 0.20 | 0.92 | 0.45 | 0.63 |
| scFoundation | 0.45 | 0.71 | 0.85 | 0.62 | 0.57 | 0.20 | 0.95 | 1.00 | 0.84 | 0.41 | 0.64 | 0.06 | 0.64 | 0.20 | 0.93 | 0.37 | 0.62 |
| scGPT | 0.43 | 0.70 | 0.84 | 0.61 | 0.55 | 0.17 | 0.93 | 1.00 | 0.84 | 0.43 | 0.66 | 0.12 | 0.63 | 0.17 | 0.92 | 0.40 | 0.62 |
| scVI | 0.42 | 0.69 | 0.82 | 0.60 | 0.54 | 0.18 | 0.93 | 1.00 | 0.85 | 0.44 | 0.64 | 0.13 | 0.61 | 0.18 | 0.92 | 0.41 | 0.62 |
| CellPLM | 0.43 | 0.70 | 0.84 | 0.62 | 0.55 | 0.21 | 0.94 | 1.00 | 0.85 | 0.39 | 0.60 | 0.08 | 0.63 | 0.21 | 0.93 | 0.36 | 0.62 |
| LangCell | 0.44 | 0.70 | 0.84 | 0.61 | 0.56 | 0.12 | 0.93 | 1.00 | 0.82 | 0.36 | 0.74 | 0.07 | 0.63 | 0.12 | 0.92 | 0.39 | 0.61 |
| scPRINT | 0.35 | 0.65 | 0.80 | 0.56 | 0.48 | 0.15 | 0.93 | 1.00 | 0.82 | 0.40 | 0.64 | 0.07 | 0.57 | 0.15 | 0.91 | 0.37 | 0.59 |
| Geneformer | 0.38 | 0.65 | 0.78 | 0.56 | 0.50 | 0.08 | 0.94 | 1.00 | 0.78 | 0.37 | 0.70 | 0.07 | 0.57 | 0.08 | 0.90 | 0.38 | 0.58 |
| CellFM | 0.37 | 0.65 | 0.78 | 0.57 | 0.50 | -0.00 | 0.86 | 0.99 | 0.38 | 0.46 | 0.95 | 0.25 | 0.57 | -0.00 | 0.74 | 0.55 | 0.57 |
| CELLama | 0.26 | 0.52 | 0.65 | 0.45 | 0.39 | 0.01 | 0.86 | 0.99 | 0.63 | 0.39 | 0.75 | 0.13 | 0.45 | 0.01 | 0.83 | 0.42 | 0.52 |
| GenePT-w | 0.11 | 0.32 | 0.41 | 0.26 | 0.23 | -0.34 | 0.82 | 0.98 | 0.50 | 0.31 | 0.65 | 0.24 | 0.27 | -0.34 | 0.77 | 0.40 | 0.40 |
| scBERT | 0.05 | 0.15 | 0.20 | 0.13 | 0.17 | -0.40 | 0.56 | 0.93 | 0.28 | 0.19 | 0.55 | 0.26 | 0.14 | -0.40 | 0.59 | 0.33 | 0.28 |

**Figure S10: Average benchmark performance across 26 Tabula Sapiens v2 tissues.** For each method, values are arithmetic means across tissues. Batch-mixing averages include only tissues for which the corresponding metric was defined. The cluster-concordance score is the unweighted mean of ARI, NMI, HOM, COM and FMI; continuous separation is represented by ASW; the local-neighbourhood structure score is the unweighted mean of Acc@kNN, cLISI and GC; and the batch-mixing score is the unweighted mean of kBET, BRAS and CiLISI. The overall score weights cluster concordance, continuous separation, local-neighbourhood structure and batch mixing by 32.5%, 10%, 32.5% and 25%, respectively. scGPT (integrated) was fine-tuned independently on each tissue, whereas scVI (de novo) was trained from random initialization independently on each tissue; both used batch identities but no cell-type labels during model fitting. PCA, Harmony, Seurat CCA, Seurat RPCA and scVI (de novo) were included as conventional baselines. Rows are ordered by the aggregated score; higher values indicate better performance.

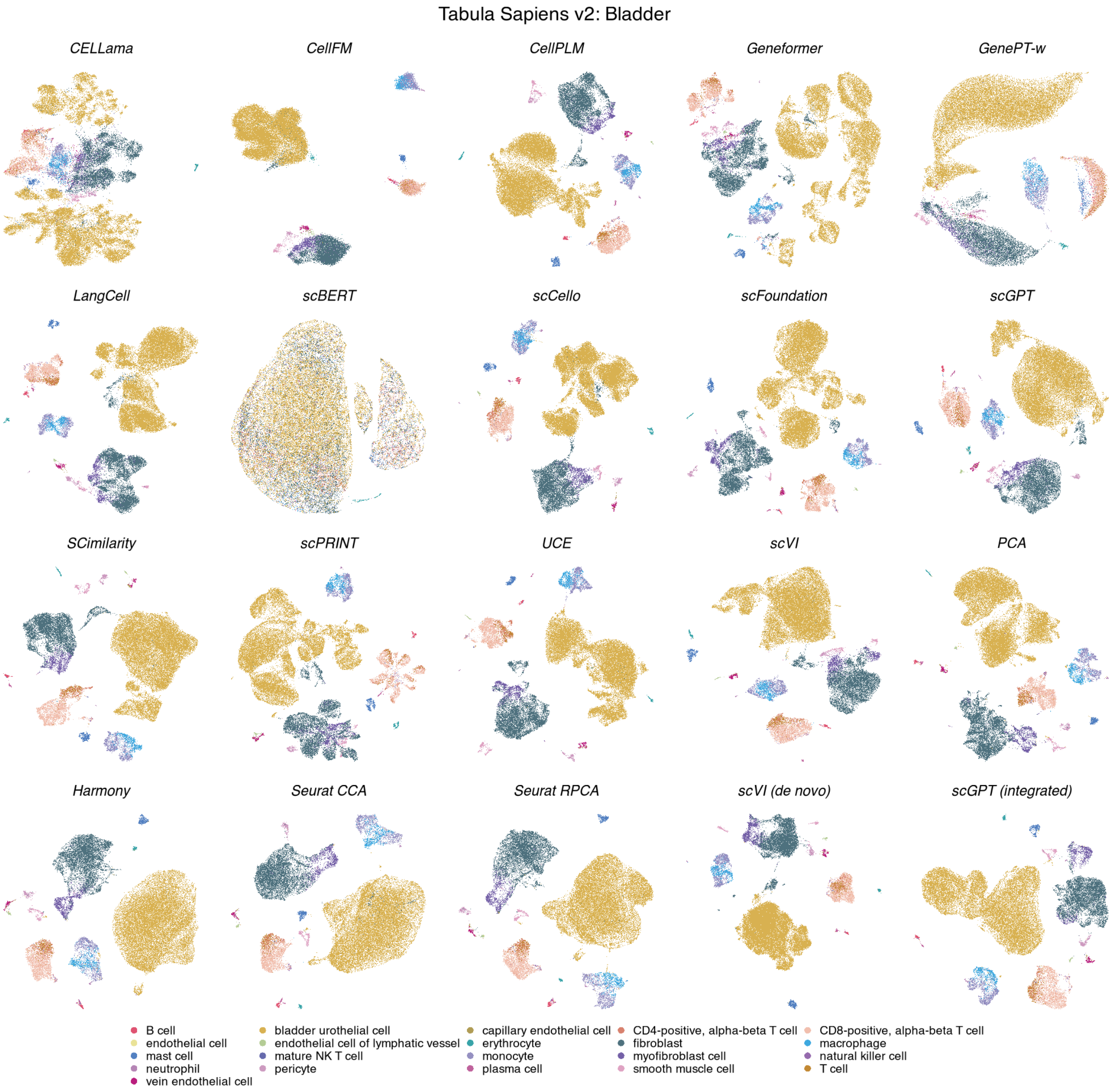

**Figure S11: Cell-type organization in Tabula Sapiens v2 bladder.** UMAPs show the same 36,715 cells across 20 methods, coloured by 21 annotated cell types. Coordinates were computed independently for each method. scGPT (integrated) and scVI (de novo) were fitted within tissue using batch labels but not cell-type labels; the other pretrained representations were evaluated without target-data fitting. PCA, Harmony, Seurat CCA, Seurat RPCA and scVI (de novo) are reference methods.

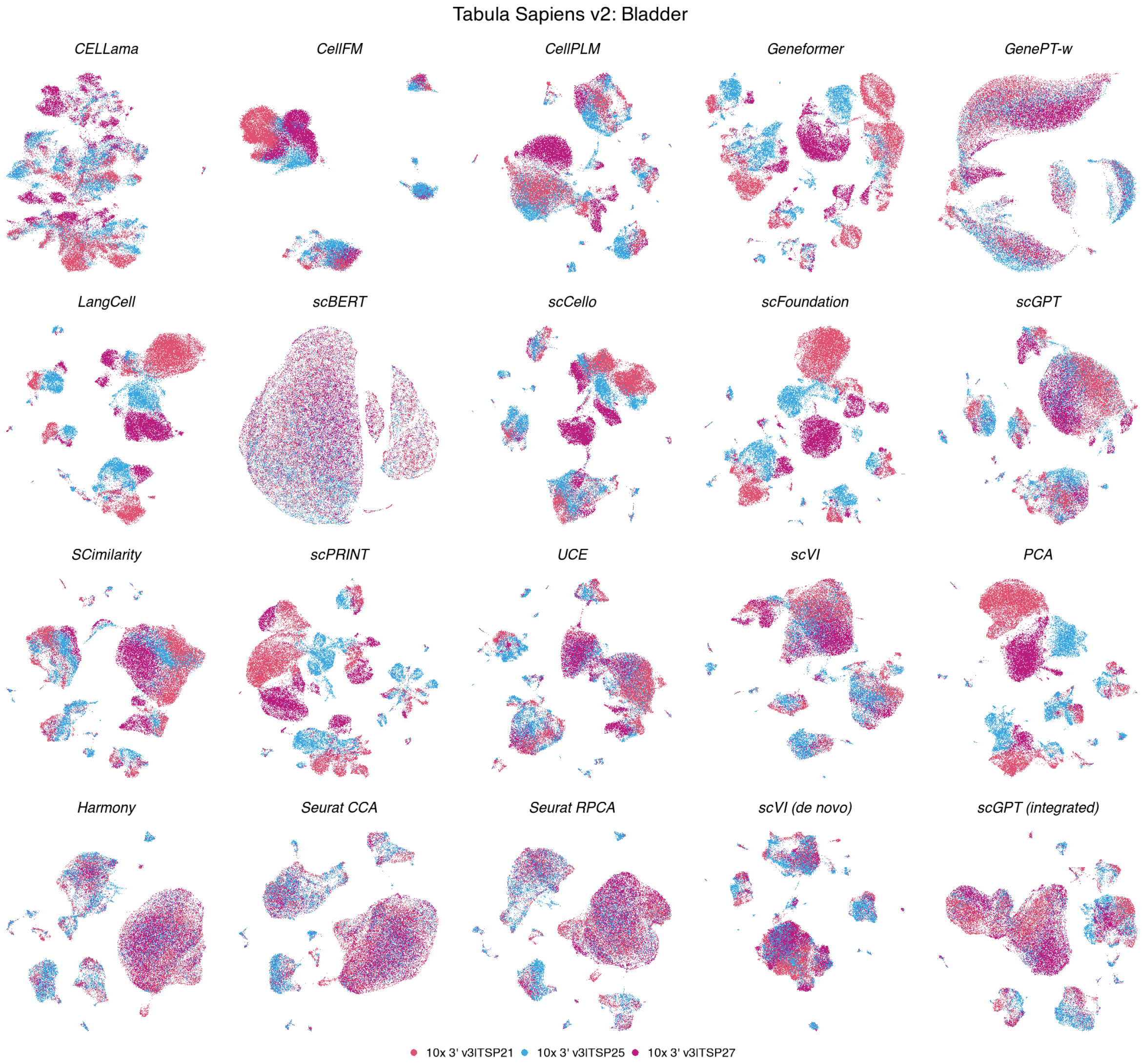

**Figure S12: Batch organization in Tabula Sapiens v2 bladder.** UMAPs show the same 36,715 cells across 20 methods, coloured by 3 experimental batches. Coordinates match the corresponding cell-type view. scGPT (integrated) and scVI (de novo) were fitted within tissue using batch labels but not cell-type labels; the other pretrained representations were evaluated without target-data fitting.

**Tabula Sapiens v2 — Bladder**

| method | ARI | NMI | HOM | COM | FMI | ASW | Acc@kNN | cLISI | GC | kBET | BRAS | CiLISI | Cluster concordance | Continuous separation | Local structure | Batch mixing | Overall |
|---|---|---|---|---|---|---|---|---|---|---|---|---|---|---|---|---|---|
| | Biological conservation metrics | | | | | | | | | Batch mixing metrics | | | Aggregated scores | | | | |
| UCE | 0.40 | 0.67 | 0.88 | 0.55 | 0.60 | 0.30 | 0.97 | 1.00 | 0.88 | 0.43 | 0.76 | 0.19 | 0.62 | 0.30 | 0.95 | 0.46 | 0.66 |
| SCimilarity | 0.33 | 0.66 | 0.88 | 0.53 | 0.54 | 0.47 | 0.97 | 1.00 | 0.91 | 0.43 | 0.71 | 0.13 | 0.59 | 0.47 | 0.96 | 0.42 | 0.66 |
| Seurat CCA | 0.24 | 0.61 | 0.87 | 0.47 | 0.45 | 0.29 | 0.96 | 1.00 | 0.86 | 0.60 | 0.74 | 0.43 | 0.53 | 0.29 | 0.94 | 0.59 | 0.65 |
| PCA | 0.34 | 0.64 | 0.87 | 0.51 | 0.55 | 0.44 | 0.97 | 1.00 | 0.90 | 0.53 | 0.72 | 0.07 | 0.58 | 0.44 | 0.96 | 0.44 | 0.65 |
| Seurat RPCA | 0.25 | 0.61 | 0.87 | 0.47 | 0.46 | 0.25 | 0.97 | 1.00 | 0.88 | 0.58 | 0.75 | 0.44 | 0.53 | 0.25 | 0.95 | 0.59 | 0.65 |
| Harmony | 0.25 | 0.62 | 0.87 | 0.48 | 0.46 | 0.25 | 0.97 | 1.00 | 0.88 | 0.58 | 0.76 | 0.38 | 0.54 | 0.25 | 0.95 | 0.57 | 0.65 |
| scVI | 0.30 | 0.63 | 0.87 | 0.50 | 0.51 | 0.26 | 0.97 | 1.00 | 0.88 | 0.41 | 0.72 | 0.23 | 0.56 | 0.26 | 0.95 | 0.45 | 0.63 |
| CellPLM | 0.28 | 0.63 | 0.87 | 0.49 | 0.50 | 0.30 | 0.97 | 1.00 | 0.93 | 0.44 | 0.70 | 0.11 | 0.55 | 0.30 | 0.97 | 0.42 | 0.63 |
| scVI (de novo) | 0.34 | 0.64 | 0.87 | 0.51 | 0.55 | -0.00 | 0.97 | 1.00 | 0.90 | 0.44 | 0.78 | 0.30 | 0.58 | -0.00 | 0.96 | 0.50 | 0.63 |
| scGPT (integrated) | 0.27 | 0.64 | 0.88 | 0.50 | 0.49 | 0.11 | 0.97 | 1.00 | 0.87 | 0.52 | 0.71 | 0.25 | 0.56 | 0.11 | 0.95 | 0.49 | 0.62 |
| scGPT | 0.24 | 0.61 | 0.86 | 0.47 | 0.46 | 0.22 | 0.97 | 1.00 | 0.92 | 0.46 | 0.72 | 0.16 | 0.53 | 0.22 | 0.96 | 0.45 | 0.62 |
| scFoundation | 0.29 | 0.63 | 0.86 | 0.49 | 0.50 | 0.32 | 0.97 | 1.00 | 0.89 | 0.39 | 0.71 | 0.05 | 0.55 | 0.32 | 0.96 | 0.38 | 0.62 |
| scCello | 0.22 | 0.60 | 0.84 | 0.46 | 0.43 | 0.31 | 0.96 | 1.00 | 0.87 | 0.47 | 0.77 | 0.13 | 0.51 | 0.31 | 0.94 | 0.46 | 0.62 |
| LangCell | 0.31 | 0.61 | 0.83 | 0.49 | 0.52 | 0.19 | 0.97 | 1.00 | 0.89 | 0.33 | 0.76 | 0.06 | 0.55 | 0.19 | 0.95 | 0.38 | 0.60 |
| scPRINT | 0.22 | 0.56 | 0.81 | 0.43 | 0.44 | 0.26 | 0.96 | 1.00 | 0.86 | 0.40 | 0.72 | 0.07 | 0.49 | 0.26 | 0.94 | 0.39 | 0.59 |
| Geneformer | 0.18 | 0.53 | 0.78 | 0.40 | 0.40 | 0.09 | 0.97 | 1.00 | 0.84 | 0.35 | 0.76 | 0.05 | 0.46 | 0.09 | 0.94 | 0.39 | 0.56 |
| CellFM | 0.19 | 0.55 | 0.81 | 0.42 | 0.40 | 0.03 | 0.94 | 1.00 | 0.33 | 0.38 | 0.95 | 0.41 | 0.47 | 0.03 | 0.76 | 0.58 | 0.55 |
| CELLama | 0.15 | 0.48 | 0.72 | 0.36 | 0.35 | 0.02 | 0.93 | 1.00 | 0.59 | 0.34 | 0.82 | 0.16 | 0.41 | 0.02 | 0.84 | 0.44 | 0.52 |
| GenePT-w | 0.10 | 0.39 | 0.62 | 0.28 | 0.28 | -0.11 | 0.91 | 1.00 | 0.49 | 0.29 | 0.76 | 0.33 | 0.34 | -0.11 | 0.80 | 0.46 | 0.47 |
| scBERT | 0.02 | 0.09 | 0.15 | 0.07 | 0.18 | -0.12 | 0.70 | 0.97 | 0.23 | 0.06 | 0.49 | 0.33 | 0.10 | -0.12 | 0.63 | 0.29 | 0.30 |

**Figure S13: Representation performance in Tabula Sapiens v2 bladder.** Scores are shown for 36,715 cells, 21 annotated cell types and 3 experimental batches. Metrics are grouped into cluster concordance, continuous separation, local neighbourhood structure and batch mixing, weighted by 32.5%, 10%, 32.5% and 25%, respectively. Rows are ordered by the resulting aggregate score; higher values indicate better performance.

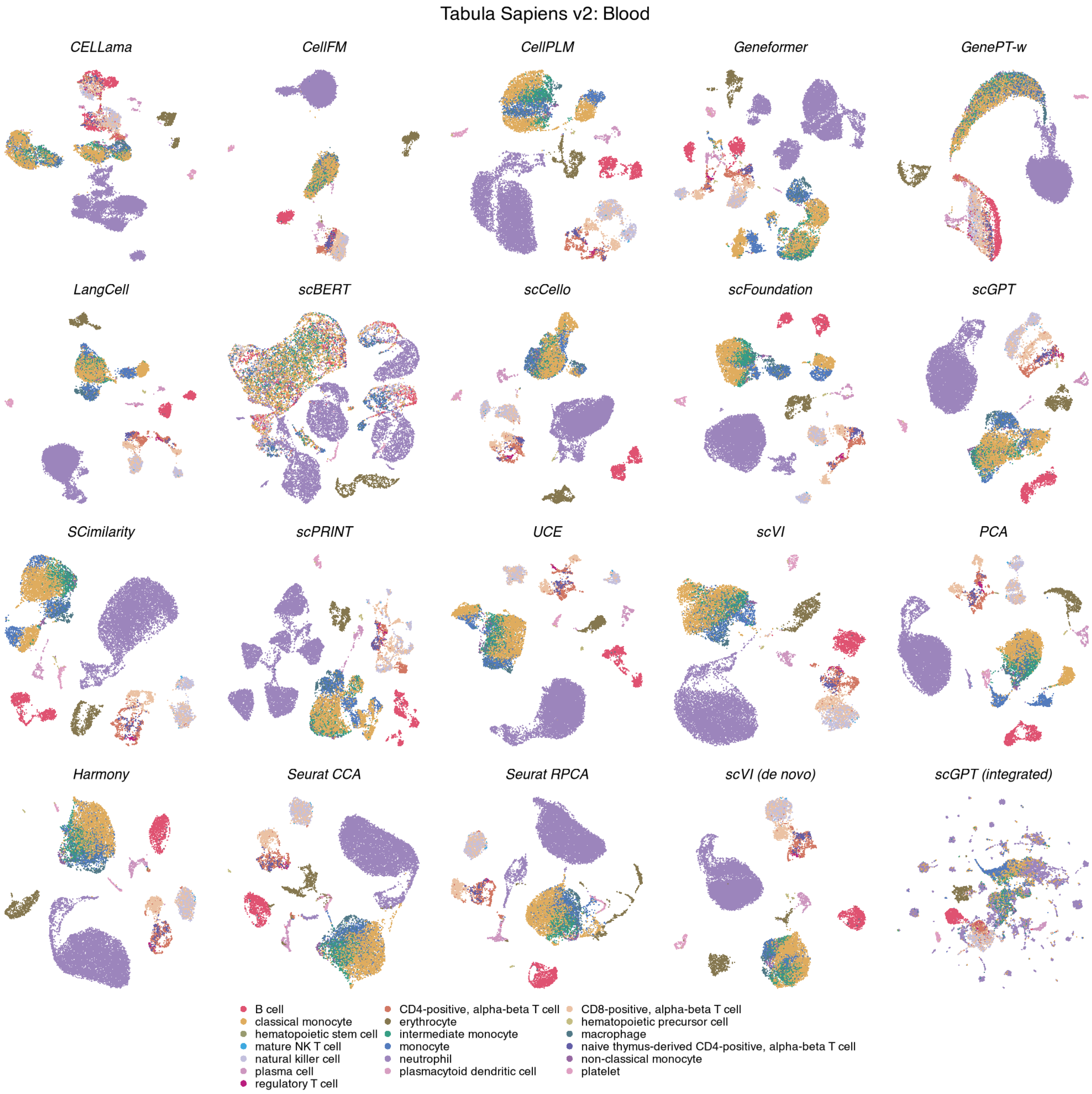

**Figure S14: Cell-type organization in Tabula Sapiens v2 blood.** UMAPs show the same 17,802 cells across 20 methods, coloured by 19 annotated cell types. Display conventions and method categories are as in Fig. S11.

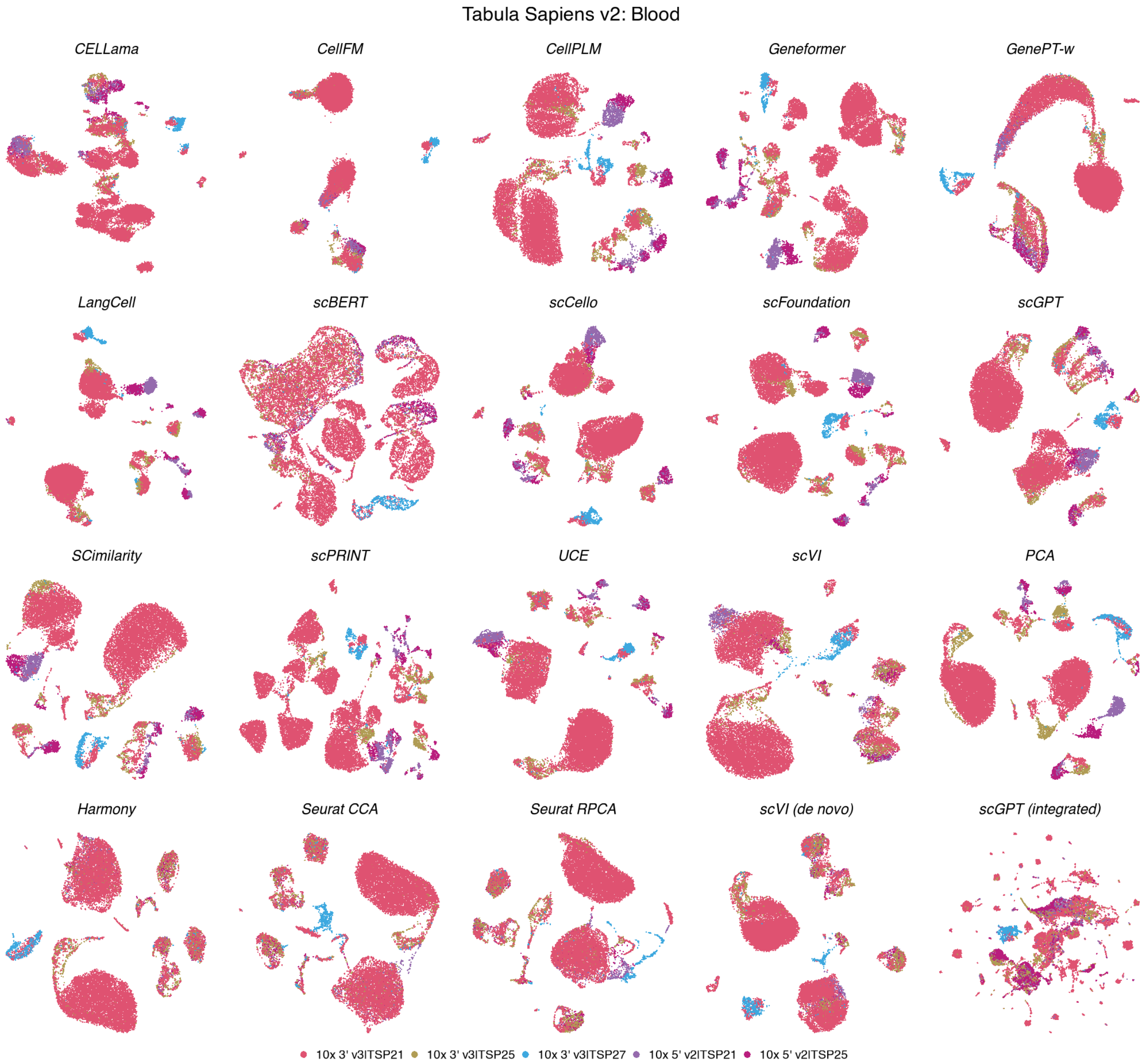

**Figure S15: Batch organization in Tabula Sapiens v2 blood.** UMAPs show the same 17,802 cells across 20 methods, coloured by 5 experimental batches. Coordinates match the corresponding cell-type view; display conventions are as in Fig. S12.

**Tabula Sapiens v2 — Blood**

| method | Biological conservation metrics | | | | | | | | | Batch mixing metrics | | | Aggregated scores | | | | |
|---|---|---|---|---|---|---|---|---|---|---|---|---|---|---|---|---|---|
| | ARI | NMI | HOM | COM | FMI | ASW | Acc@kNN | cLISI | GC | kBET | BRAS | CiLISI | Cluster concordance | Continuous separation | Local structure | Batch mixing | Overall |
| Harmony | 0.32 | 0.67 | 0.79 | 0.58 | 0.46 | 0.36 | 0.87 | 1.00 | 0.83 | 0.63 | 0.75 | 0.12 | 0.56 | 0.36 | 0.90 | 0.50 | 0.64 |
| LangCell | 0.75 | 0.75 | 0.78 | 0.72 | 0.81 | 0.18 | 0.89 | 1.00 | 0.74 | 0.22 | 0.63 | 0.03 | 0.76 | 0.18 | 0.88 | 0.30 | 0.62 |
| SCimilarity | 0.38 | 0.69 | 0.80 | 0.62 | 0.51 | 0.39 | 0.90 | 1.00 | 0.85 | 0.40 | 0.56 | 0.04 | 0.60 | 0.39 | 0.92 | 0.34 | 0.62 |
| Seurat CCA | 0.31 | 0.65 | 0.78 | 0.56 | 0.46 | 0.32 | 0.87 | 1.00 | 0.80 | 0.54 | 0.71 | 0.11 | 0.55 | 0.32 | 0.89 | 0.45 | 0.61 |
| Seurat RPCA | 0.33 | 0.66 | 0.77 | 0.57 | 0.47 | 0.31 | 0.87 | 1.00 | 0.81 | 0.51 | 0.69 | 0.11 | 0.56 | 0.31 | 0.89 | 0.43 | 0.61 |
| PCA | 0.50 | 0.72 | 0.81 | 0.65 | 0.61 | 0.20 | 0.90 | 1.00 | 0.83 | 0.36 | 0.59 | 0.03 | 0.66 | 0.20 | 0.91 | 0.33 | 0.61 |
| scVI | 0.30 | 0.66 | 0.78 | 0.57 | 0.45 | 0.31 | 0.88 | 1.00 | 0.84 | 0.46 | 0.63 | 0.06 | 0.55 | 0.31 | 0.91 | 0.38 | 0.60 |
| scVI (de novo) | 0.29 | 0.65 | 0.77 | 0.57 | 0.44 | 0.05 | 0.87 | 1.00 | 0.87 | 0.51 | 0.79 | 0.09 | 0.54 | 0.05 | 0.91 | 0.46 | 0.59 |
| scCello | 0.44 | 0.66 | 0.74 | 0.60 | 0.56 | 0.26 | 0.86 | 1.00 | 0.75 | 0.35 | 0.63 | 0.05 | 0.60 | 0.26 | 0.87 | 0.35 | 0.59 |
| scFoundation | 0.37 | 0.68 | 0.80 | 0.59 | 0.51 | 0.28 | 0.91 | 1.00 | 0.82 | 0.32 | 0.52 | 0.03 | 0.59 | 0.28 | 0.91 | 0.29 | 0.59 |
| CellPLM | 0.37 | 0.66 | 0.76 | 0.58 | 0.51 | 0.34 | 0.91 | 1.00 | 0.81 | 0.33 | 0.47 | 0.03 | 0.57 | 0.34 | 0.90 | 0.28 | 0.58 |
| scGPT | 0.30 | 0.65 | 0.78 | 0.55 | 0.44 | 0.29 | 0.89 | 1.00 | 0.83 | 0.34 | 0.56 | 0.04 | 0.54 | 0.29 | 0.91 | 0.31 | 0.58 |
| UCE | 0.29 | 0.64 | 0.75 | 0.56 | 0.43 | 0.30 | 0.88 | 1.00 | 0.74 | 0.36 | 0.62 | 0.04 | 0.54 | 0.30 | 0.87 | 0.34 | 0.57 |
| Geneformer | 0.33 | 0.63 | 0.76 | 0.54 | 0.47 | 0.20 | 0.89 | 1.00 | 0.67 | 0.30 | 0.62 | 0.03 | 0.55 | 0.20 | 0.85 | 0.32 | 0.55 |
| scPRINT | 0.28 | 0.65 | 0.77 | 0.56 | 0.42 | 0.26 | 0.89 | 1.00 | 0.75 | 0.25 | 0.52 | 0.03 | 0.53 | 0.26 | 0.88 | 0.27 | 0.55 |
| CellFM | 0.26 | 0.61 | 0.69 | 0.55 | 0.40 | 0.01 | 0.79 | 1.00 | 0.30 | 0.36 | 0.96 | 0.09 | 0.50 | 0.01 | 0.70 | 0.47 | 0.51 |
| CELLama | 0.23 | 0.54 | 0.65 | 0.46 | 0.38 | 0.07 | 0.83 | 1.00 | 0.59 | 0.36 | 0.68 | 0.05 | 0.45 | 0.07 | 0.81 | 0.37 | 0.51 |
| GenePT-w | 0.21 | 0.46 | 0.57 | 0.39 | 0.36 | -0.06 | 0.80 | 0.99 | 0.53 | 0.31 | 0.57 | 0.09 | 0.40 | -0.06 | 0.78 | 0.32 | 0.46 |
| scGPT (integrated) | 0.07 | 0.38 | 0.46 | 0.32 | 0.22 | 0.02 | 0.67 | 0.97 | 0.55 | 0.60 | 0.72 | 0.11 | 0.29 | 0.02 | 0.73 | 0.48 | 0.45 |
| scBERT | 0.16 | 0.34 | 0.40 | 0.29 | 0.30 | -0.19 | 0.71 | 0.96 | 0.34 | 0.20 | 0.51 | 0.09 | 0.30 | -0.19 | 0.67 | 0.26 | 0.36 |

**Figure S16: Representation performance in Tabula Sapiens v2 blood.** Scores are shown for 17,802 cells, 19 annotated cell types and 5 experimental batches. Metrics, family weighting and row ordering are as in Fig. S13.

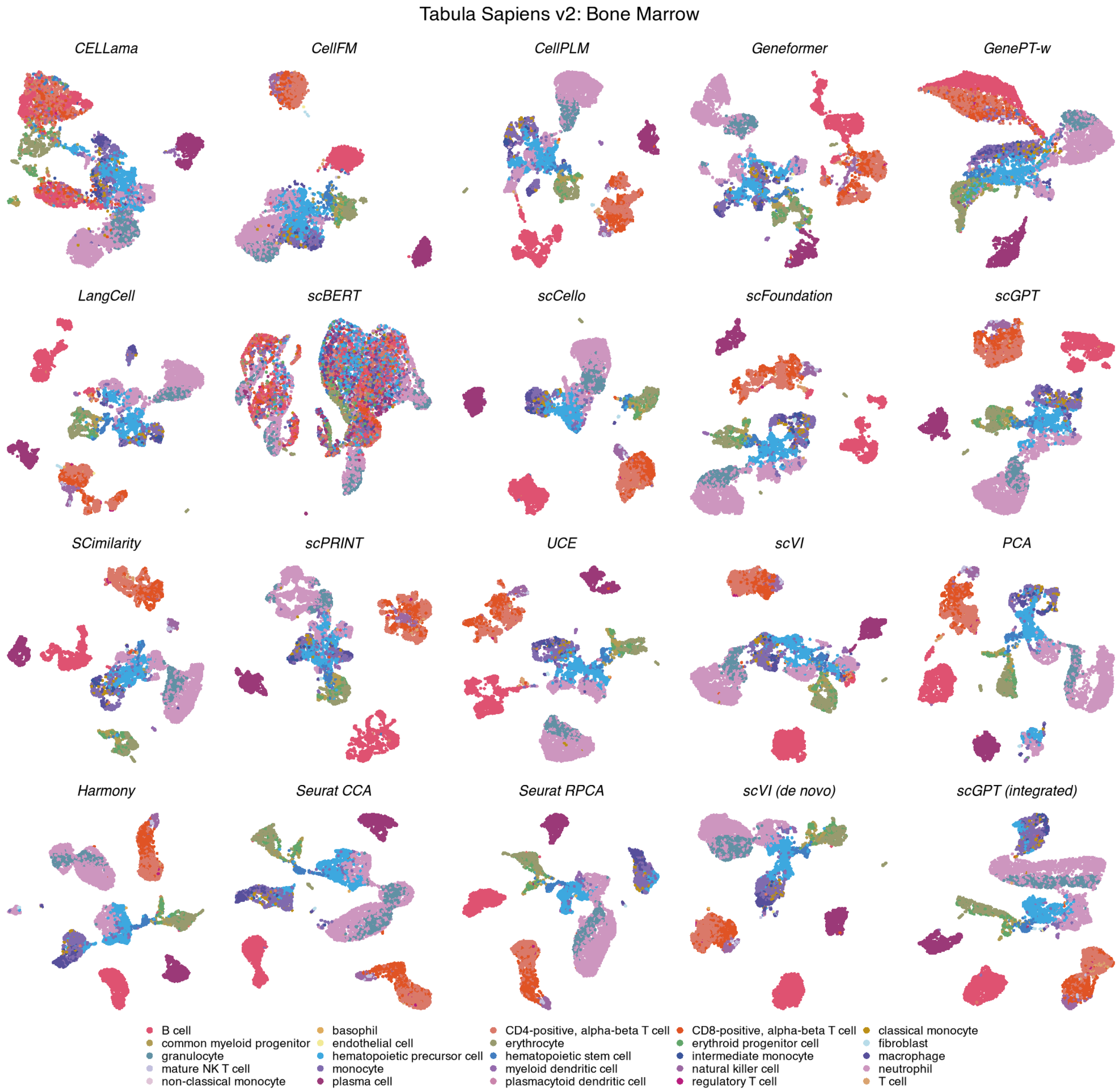

**Figure S17: Cell-type organization in Tabula Sapiens v2 bone marrow.** UMAPs show the same 8,045 cells across 20 methods, coloured by 25 annotated cell types. Display conventions and method categories are as in Fig. S11.

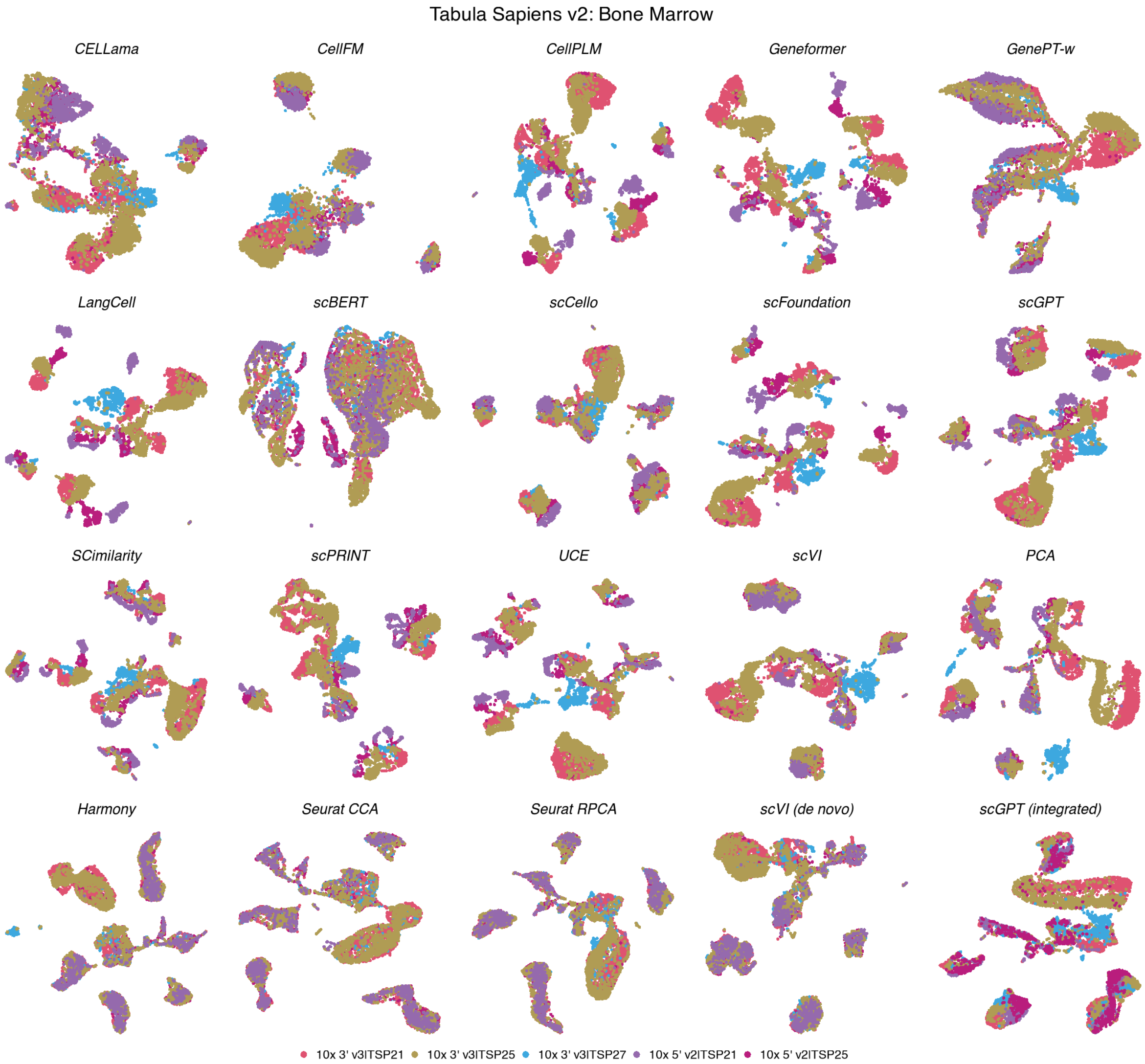

**Figure S18: Batch organization in Tabula Sapiens v2 bone marrow.** UMAPs show the same 8,045 cells across 20 methods, coloured by 5 experimental batches. Coordinates match the corresponding cell-type view; display conventions are as in Fig. S12.

**Tabula Sapiens v2 — Bone Marrow**

| method | Biological conservation metrics | | | | | | | | | Batch mixing metrics | | | Aggregated scores | | | | |
|---|---|---|---|---|---|---|---|---|---|---|---|---|---|---|---|---|---|
| | ARI | NMI | HOM | COM | FMI | ASW | Acc@kNN | cLISI | GC | kBET | BRAS | CiLISI | Cluster concordance | Continuous separation | Local structure | Batch mixing | Overall |
| **scVI (de novo)** | 0.45 | 0.70 | 0.78 | 0.63 | 0.52 | 0.06 | 0.84 | 0.99 | 0.79 | 0.42 | 0.80 | 0.33 | 0.62 | 0.06 | 0.87 | 0.52 | 0.62 |
| **Seurat CCA** | 0.41 | 0.68 | 0.77 | 0.61 | 0.49 | 0.06 | 0.81 | 0.99 | 0.76 | 0.61 | 0.72 | 0.36 | 0.59 | 0.06 | 0.86 | 0.56 | 0.62 |
| **Seurat RPCA** | 0.44 | 0.71 | 0.79 | 0.64 | 0.52 | 0.06 | 0.84 | 0.99 | 0.76 | 0.49 | 0.72 | 0.34 | 0.62 | 0.06 | 0.86 | 0.52 | 0.62 |
| **Harmony** | 0.45 | 0.70 | 0.78 | 0.64 | 0.52 | 0.04 | 0.84 | 0.99 | 0.74 | 0.49 | 0.73 | 0.35 | 0.62 | 0.04 | 0.86 | 0.53 | 0.62 |
| **SCimilarity** | 0.38 | 0.70 | 0.79 | 0.63 | 0.47 | 0.13 | 0.87 | 1.00 | 0.81 | 0.32 | 0.64 | 0.12 | 0.59 | 0.13 | 0.89 | 0.36 | 0.59 |
| **scCello** | 0.40 | 0.68 | 0.75 | 0.61 | 0.48 | 0.09 | 0.83 | 0.99 | 0.80 | 0.35 | 0.71 | 0.17 | 0.58 | 0.09 | 0.88 | 0.41 | 0.59 |
| **scGPT (integrated)** | 0.41 | 0.69 | 0.78 | 0.62 | 0.49 | 0.11 | 0.85 | 1.00 | 0.76 | 0.30 | 0.64 | 0.19 | 0.60 | 0.11 | 0.87 | 0.38 | 0.58 |
| **PCA** | 0.44 | 0.71 | 0.79 | 0.64 | 0.52 | 0.04 | 0.86 | 1.00 | 0.81 | 0.31 | 0.60 | 0.12 | 0.62 | 0.04 | 0.89 | 0.35 | 0.58 |
| **scFoundation** | 0.48 | 0.69 | 0.76 | 0.64 | 0.55 | 0.08 | 0.86 | 1.00 | 0.81 | 0.26 | 0.56 | 0.07 | 0.62 | 0.08 | 0.89 | 0.29 | 0.57 |
| **scVI** | 0.36 | 0.67 | 0.76 | 0.60 | 0.44 | 0.09 | 0.85 | 0.99 | 0.83 | 0.30 | 0.59 | 0.16 | 0.57 | 0.09 | 0.89 | 0.35 | 0.57 |
| **UCE** | 0.38 | 0.69 | 0.77 | 0.62 | 0.46 | 0.09 | 0.86 | 1.00 | 0.76 | 0.26 | 0.65 | 0.12 | 0.58 | 0.09 | 0.87 | 0.34 | 0.57 |
| **scGPT** | 0.41 | 0.68 | 0.76 | 0.62 | 0.48 | 0.07 | 0.85 | 0.99 | 0.79 | 0.25 | 0.60 | 0.11 | 0.59 | 0.07 | 0.88 | 0.32 | 0.57 |
| **LangCell** | 0.39 | 0.66 | 0.75 | 0.60 | 0.47 | 0.04 | 0.85 | 0.99 | 0.74 | 0.15 | 0.64 | 0.07 | 0.57 | 0.04 | 0.86 | 0.29 | 0.54 |
| **CellPLM** | 0.37 | 0.66 | 0.74 | 0.59 | 0.45 | 0.07 | 0.86 | 0.99 | 0.80 | 0.17 | 0.51 | 0.08 | 0.56 | 0.07 | 0.89 | 0.25 | 0.54 |
| **scPRINT** | 0.31 | 0.61 | 0.68 | 0.54 | 0.39 | 0.07 | 0.85 | 0.99 | 0.77 | 0.28 | 0.58 | 0.10 | 0.51 | 0.07 | 0.87 | 0.32 | 0.53 |
| **CellFM** | 0.39 | 0.64 | 0.71 | 0.58 | 0.47 | -0.02 | 0.75 | 0.98 | 0.30 | 0.40 | 0.95 | 0.26 | 0.56 | -0.02 | 0.67 | 0.53 | 0.53 |
| **Geneformer** | 0.33 | 0.62 | 0.69 | 0.56 | 0.41 | 0.01 | 0.85 | 0.99 | 0.68 | 0.18 | 0.63 | 0.09 | 0.52 | 0.01 | 0.84 | 0.30 | 0.52 |
| **CELLama** | 0.26 | 0.52 | 0.59 | 0.46 | 0.35 | -0.04 | 0.76 | 0.97 | 0.54 | 0.24 | 0.70 | 0.14 | 0.44 | -0.04 | 0.76 | 0.36 | 0.47 |
| **GenePT-w** | 0.11 | 0.30 | 0.34 | 0.26 | 0.19 | -0.46 | 0.69 | 0.96 | 0.44 | 0.26 | 0.63 | 0.31 | 0.24 | -0.46 | 0.70 | 0.40 | 0.36 |
| **scBERT** | 0.07 | 0.20 | 0.23 | 0.18 | 0.15 | -0.41 | 0.43 | 0.91 | 0.27 | 0.14 | 0.56 | 0.27 | 0.17 | -0.41 | 0.54 | 0.32 | 0.27 |

**Figure S19: Representation performance in Tabula Sapiens v2 bone marrow.** Scores are shown for 8,045 cells, 25 annotated cell types and 5 experimental batches. Metrics, family weighting and row ordering are as in Fig. S13.

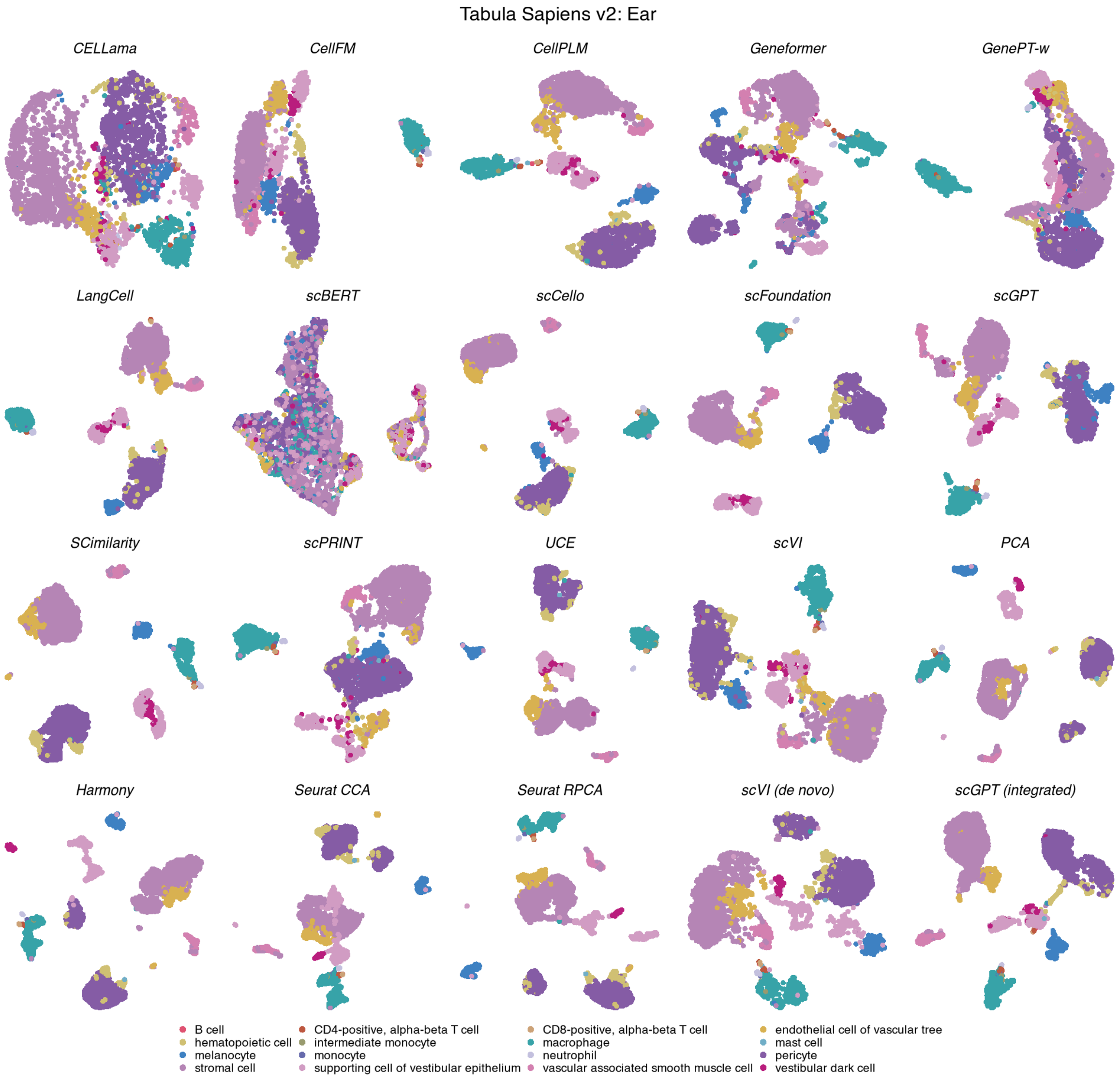

**Figure S20: Cell-type organization in Tabula Sapiens v2 ear.** UMAPs show the same 3,055 cells across 20 methods, coloured by 16 annotated cell types. Display conventions and method categories are as in Fig. S11.

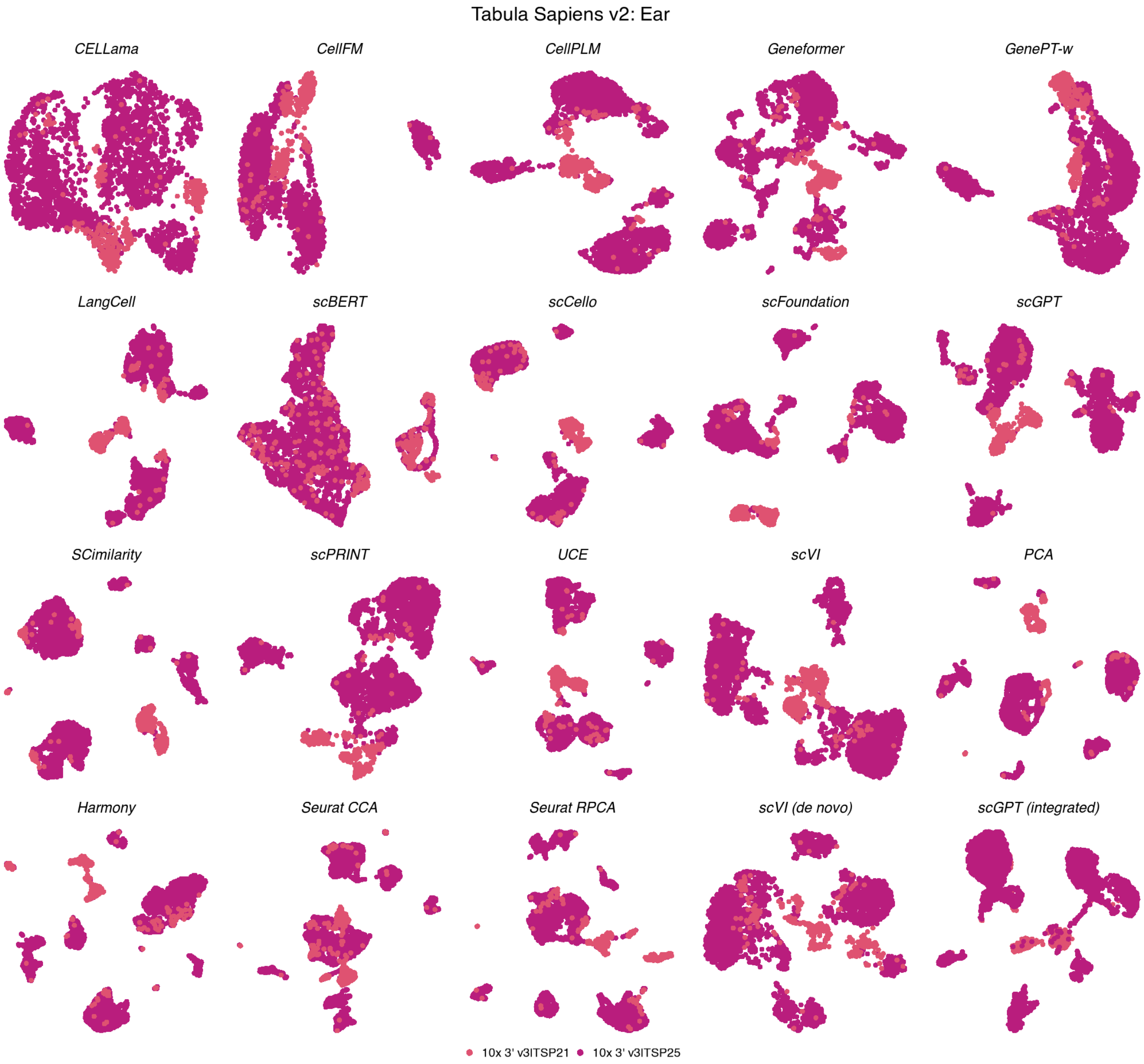

**Figure S21: Batch organization in Tabula Sapiens v2 ear.** UMAPs show the same 3,055 cells across 20 methods, coloured by 2 experimental batches. Coordinates match the corresponding cell-type view; display conventions are as in Fig. S12.

**Tabula Sapiens v2 — Ear**

| method | Biological conservation metrics: ARI | NMI | HOM | COM | FMI | ASW | Acc@kNN | cLISI | GC | Batch mixing metrics: kBET | BRAS | CiLISI | Aggregated scores: Cluster concordance | Continuous separation | Local structure | Batch mixing | Overall |
|---|---|---|---|---|---|---|---|---|---|---|---|---|---|---|---|---|---|
| Harmony | 0.47 | 0.74 | 0.90 | 0.63 | 0.59 | 0.22 | 0.98 | 1.00 | 0.86 | 0.92 | 0.71 | 0.07 | 0.67 | 0.22 | 0.95 | 0.56 | 0.69 |
| Seurat CCA | 0.46 | 0.74 | 0.89 | 0.63 | 0.59 | 0.17 | 0.97 | 1.00 | 0.88 | 0.83 | 0.71 | 0.06 | 0.66 | 0.17 | 0.95 | 0.53 | 0.68 |
| Seurat RPCA | 0.41 | 0.72 | 0.90 | 0.60 | 0.55 | 0.20 | 0.98 | 1.00 | 0.89 | 0.88 | 0.70 | 0.05 | 0.64 | 0.20 | 0.95 | 0.54 | 0.67 |
| scGPT (integrated) | 0.36 | 0.69 | 0.87 | 0.57 | 0.51 | 0.27 | 0.97 | 1.00 | 0.86 | 0.93 | 0.69 | 0.05 | 0.60 | 0.27 | 0.94 | 0.56 | 0.67 |
| scGPT | 0.50 | 0.72 | 0.86 | 0.62 | 0.61 | 0.17 | 0.96 | 1.00 | 0.84 | 0.80 | 0.72 | 0.04 | 0.66 | 0.17 | 0.93 | 0.52 | 0.66 |
| scVI (de novo) | 0.51 | 0.72 | 0.85 | 0.62 | 0.63 | 0.10 | 0.97 | 1.00 | 0.83 | 0.78 | 0.78 | 0.05 | 0.67 | 0.10 | 0.93 | 0.54 | 0.66 |
| scFoundation | 0.43 | 0.71 | 0.88 | 0.60 | 0.56 | 0.19 | 0.97 | 1.00 | 0.83 | 0.85 | 0.72 | 0.04 | 0.64 | 0.19 | 0.93 | 0.54 | 0.66 |
| UCE | 0.41 | 0.70 | 0.85 | 0.59 | 0.55 | 0.19 | 0.96 | 1.00 | 0.87 | 0.84 | 0.74 | 0.04 | 0.62 | 0.19 | 0.94 | 0.54 | 0.66 |
| scCello | 0.40 | 0.69 | 0.85 | 0.58 | 0.54 | 0.19 | 0.95 | 1.00 | 0.84 | 0.86 | 0.76 | 0.05 | 0.61 | 0.19 | 0.93 | 0.56 | 0.66 |
| SCimilarity | 0.39 | 0.69 | 0.86 | 0.58 | 0.53 | 0.26 | 0.97 | 1.00 | 0.83 | 0.84 | 0.67 | 0.05 | 0.61 | 0.26 | 0.93 | 0.52 | 0.66 |
| LangCell | 0.43 | 0.72 | 0.87 | 0.61 | 0.57 | 0.11 | 0.97 | 1.00 | 0.82 | 0.75 | 0.81 | 0.04 | 0.64 | 0.11 | 0.93 | 0.54 | 0.66 |
| scPRINT | 0.38 | 0.67 | 0.84 | 0.56 | 0.52 | 0.09 | 0.95 | 1.00 | 0.82 | 0.90 | 0.73 | 0.04 | 0.59 | 0.09 | 0.92 | 0.56 | 0.64 |
| PCA | 0.36 | 0.68 | 0.84 | 0.57 | 0.50 | 0.24 | 0.97 | 1.00 | 0.83 | 0.73 | 0.68 | 0.04 | 0.59 | 0.24 | 0.93 | 0.48 | 0.64 |
| scVI | 0.37 | 0.65 | 0.81 | 0.54 | 0.51 | 0.19 | 0.95 | 1.00 | 0.85 | 0.84 | 0.67 | 0.05 | 0.57 | 0.19 | 0.93 | 0.52 | 0.64 |
| CellPLM | 0.35 | 0.68 | 0.85 | 0.56 | 0.49 | 0.19 | 0.96 | 1.00 | 0.81 | 0.83 | 0.65 | 0.04 | 0.59 | 0.19 | 0.92 | 0.51 | 0.64 |
| CellFM | 0.39 | 0.66 | 0.81 | 0.55 | 0.52 | -0.00 | 0.93 | 0.99 | 0.56 | 0.89 | 0.95 | 0.03 | 0.59 | -0.00 | 0.83 | 0.63 | 0.62 |
| Geneformer | 0.38 | 0.54 | 0.64 | 0.47 | 0.51 | 0.06 | 0.95 | 1.00 | 0.75 | 0.84 | 0.79 | 0.04 | 0.51 | 0.06 | 0.90 | 0.56 | 0.60 |
| CELLama | 0.30 | 0.58 | 0.72 | 0.48 | 0.44 | 0.07 | 0.92 | 1.00 | 0.74 | 0.86 | 0.81 | 0.04 | 0.50 | 0.07 | 0.89 | 0.57 | 0.60 |
| GenePT-w | 0.10 | 0.24 | 0.31 | 0.20 | 0.23 | -0.40 | 0.77 | 0.96 | 0.49 | 0.55 | 0.61 | 0.07 | 0.22 | -0.40 | 0.74 | 0.41 | 0.37 |
| scBERT | 0.06 | 0.14 | 0.18 | 0.12 | 0.19 | -0.43 | 0.63 | 0.92 | 0.40 | 0.64 | 0.53 | 0.07 | 0.14 | -0.43 | 0.65 | 0.42 | 0.32 |

**Figure S22: Representation performance in Tabula Sapiens v2 ear.** Scores are shown for 3,055 cells, 16 annotated cell types and 2 experimental batches. Metrics, family weighting and row ordering are as in Fig. S13.

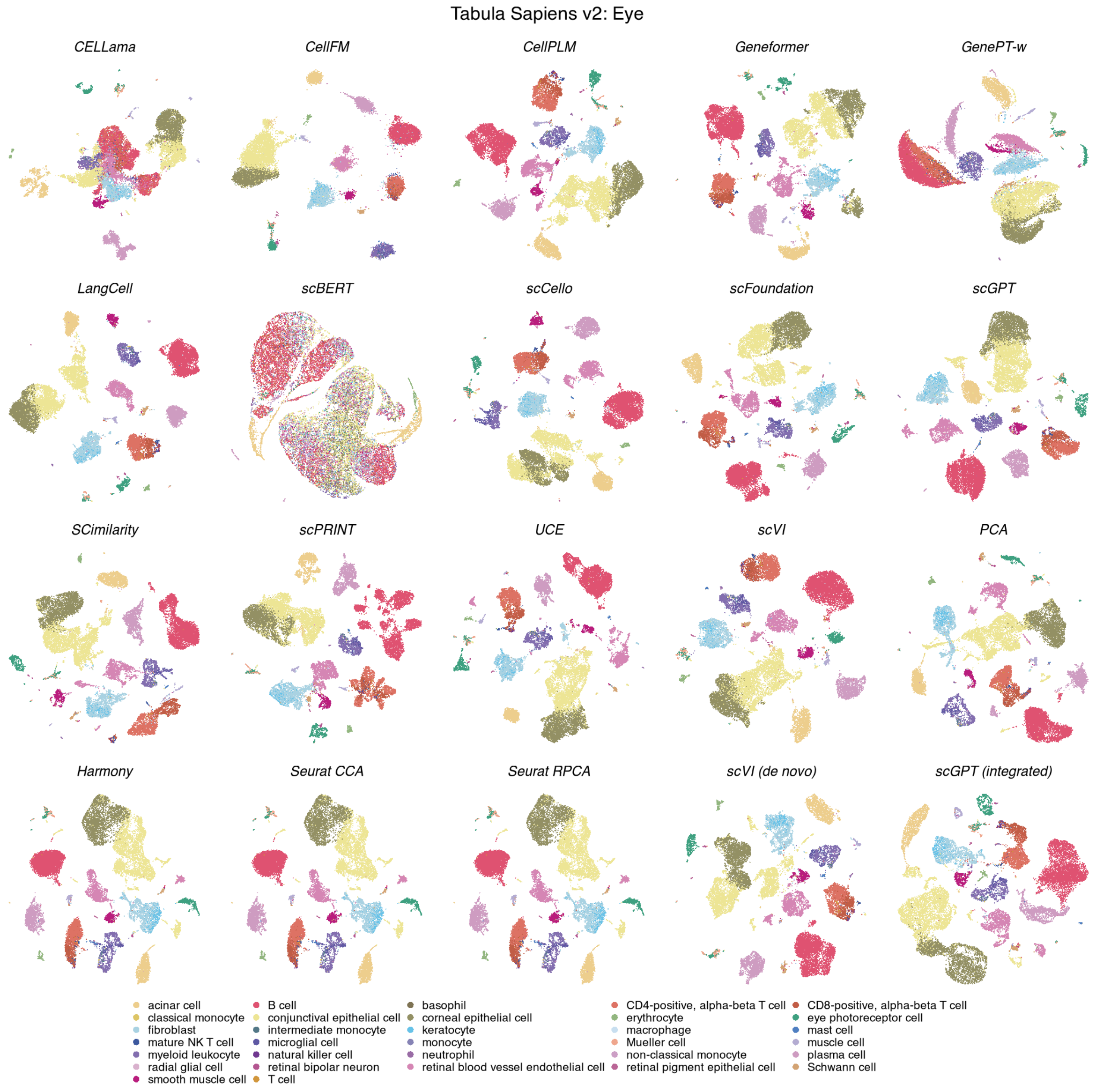

**Figure S23: Cell-type organization in Tabula Sapiens v2 eye.** UMAPs show the same 21,121 cells across 20 methods, coloured by 32 annotated cell types. Display conventions and method categories are as in Fig. S11.

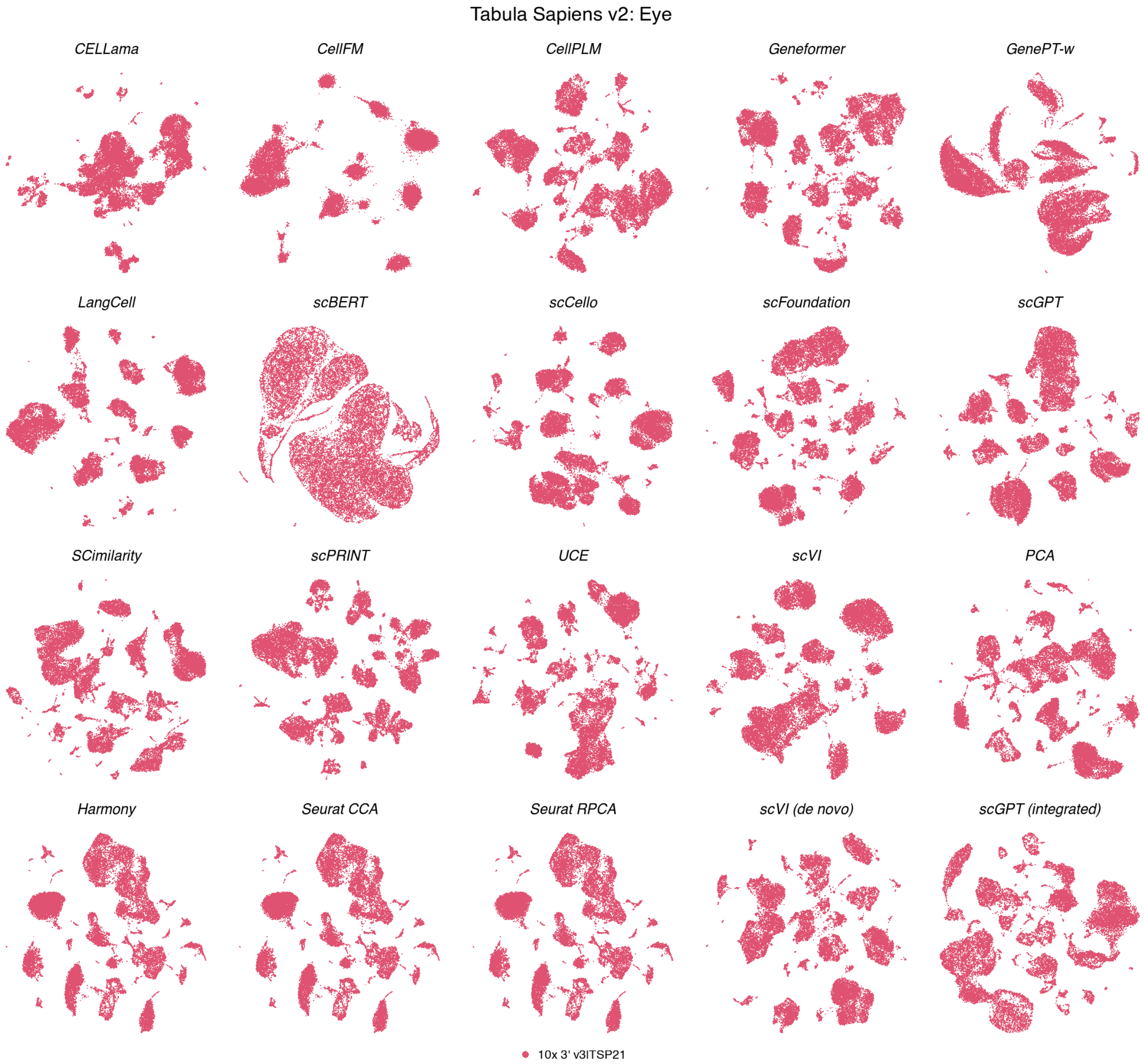

**Figure S24: Batch organization in Tabula Sapiens v2 eye.** UMAPs show the same 21,121 cells across 20 methods, coloured by a single experimental batch. Coordinates match the corresponding cell-type view; display conventions are as in Fig. S12.

**Tabula Sapiens v2 — Eye**

| method | Biological conservation metrics | | | | | | | | | Batch mixing metrics | | | Aggregated scores | | | | |
|---|---|---|---|---|---|---|---|---|---|---|---|---|---|---|---|---|---|
| | ARI | NMI | HOM | COM | FMI | ASW | Acc@kNN | cLISI | GC | kBET | BRAS | CiLISI | Cluster concordance | Continuous separation | Local structure | Batch mixing | Overall |
| SCimilarity | 0.63 | 0.83 | 0.91 | 0.77 | 0.69 | 0.29 | 0.94 | 1.00 | 0.81 | nan | nan | nan | 0.77 | 0.29 | 0.92 | | 0.77 |
| scFoundation | 0.64 | 0.83 | 0.90 | 0.77 | 0.70 | 0.21 | 0.95 | 1.00 | 0.80 | nan | nan | nan | 0.77 | 0.21 | 0.92 | | 0.76 |
| Harmony | 0.68 | 0.81 | 0.89 | 0.75 | 0.73 | 0.25 | 0.94 | 1.00 | 0.76 | nan | nan | nan | 0.77 | 0.25 | 0.90 | | 0.76 |
| Seurat CCA | 0.68 | 0.81 | 0.89 | 0.75 | 0.73 | 0.25 | 0.94 | 1.00 | 0.76 | nan | nan | nan | 0.77 | 0.25 | 0.90 | | 0.76 |
| Seurat RPCA | 0.68 | 0.81 | 0.89 | 0.75 | 0.73 | 0.25 | 0.94 | 1.00 | 0.76 | nan | nan | nan | 0.77 | 0.25 | 0.90 | | 0.76 |
| PCA | 0.65 | 0.82 | 0.90 | 0.76 | 0.70 | 0.20 | 0.94 | 1.00 | 0.80 | nan | nan | nan | 0.77 | 0.20 | 0.91 | | 0.75 |
| LangCell | 0.62 | 0.82 | 0.90 | 0.76 | 0.67 | 0.12 | 0.94 | 1.00 | 0.78 | nan | nan | nan | 0.75 | 0.12 | 0.91 | | 0.74 |
| scGPT (integrated) | 0.59 | 0.80 | 0.88 | 0.73 | 0.65 | 0.23 | 0.94 | 1.00 | 0.75 | nan | nan | nan | 0.73 | 0.23 | 0.89 | | 0.73 |
| scVI (de novo) | 0.56 | 0.81 | 0.88 | 0.74 | 0.62 | 0.12 | 0.94 | 1.00 | 0.83 | nan | nan | nan | 0.72 | 0.12 | 0.92 | | 0.73 |
| scVI | 0.54 | 0.79 | 0.88 | 0.71 | 0.62 | 0.19 | 0.93 | 1.00 | 0.80 | nan | nan | nan | 0.71 | 0.19 | 0.91 | | 0.73 |
| UCE | 0.59 | 0.80 | 0.89 | 0.73 | 0.66 | 0.17 | 0.94 | 1.00 | 0.73 | nan | nan | nan | 0.73 | 0.17 | 0.89 | | 0.73 |
| scGPT | 0.52 | 0.79 | 0.88 | 0.72 | 0.59 | 0.19 | 0.94 | 1.00 | 0.81 | nan | nan | nan | 0.70 | 0.19 | 0.92 | | 0.73 |
| CellPLM | 0.52 | 0.79 | 0.88 | 0.71 | 0.60 | 0.19 | 0.94 | 1.00 | 0.78 | nan | nan | nan | 0.70 | 0.19 | 0.91 | | 0.72 |
| scCello | 0.50 | 0.79 | 0.87 | 0.72 | 0.57 | 0.17 | 0.93 | 1.00 | 0.75 | nan | nan | nan | 0.69 | 0.17 | 0.89 | | 0.71 |
| Geneformer | 0.55 | 0.77 | 0.86 | 0.71 | 0.61 | 0.09 | 0.94 | 1.00 | 0.73 | nan | nan | nan | 0.70 | 0.09 | 0.89 | | 0.70 |
| scPRINT | 0.45 | 0.75 | 0.85 | 0.67 | 0.53 | 0.18 | 0.93 | 1.00 | 0.78 | nan | nan | nan | 0.65 | 0.18 | 0.90 | | 0.70 |
| CellFM | 0.53 | 0.77 | 0.85 | 0.71 | 0.59 | -0.01 | 0.89 | 0.99 | 0.29 | nan | nan | nan | 0.69 | -0.01 | 0.72 | | 0.61 |
| CELLama | 0.30 | 0.60 | 0.68 | 0.53 | 0.38 | 0.02 | 0.82 | 0.99 | 0.59 | nan | nan | nan | 0.50 | 0.02 | 0.80 | | 0.57 |
| GenePT-w | 0.13 | 0.40 | 0.46 | 0.35 | 0.21 | -0.32 | 0.81 | 0.98 | 0.51 | nan | nan | nan | 0.31 | -0.32 | 0.77 | | 0.43 |
| scBERT | 0.09 | 0.23 | 0.26 | 0.20 | 0.16 | -0.37 | 0.47 | 0.94 | 0.25 | nan | nan | nan | 0.19 | -0.37 | 0.55 | | 0.27 |

**Figure S25: Representation performance in Tabula Sapiens v2 eye.** Scores are shown for 21,121 cells, 32 annotated cell types and a single experimental batch. Metrics, family weighting and row ordering are as in Fig. S13. Batch-mixing metrics were unavailable, and the remaining family weights were renormalized.

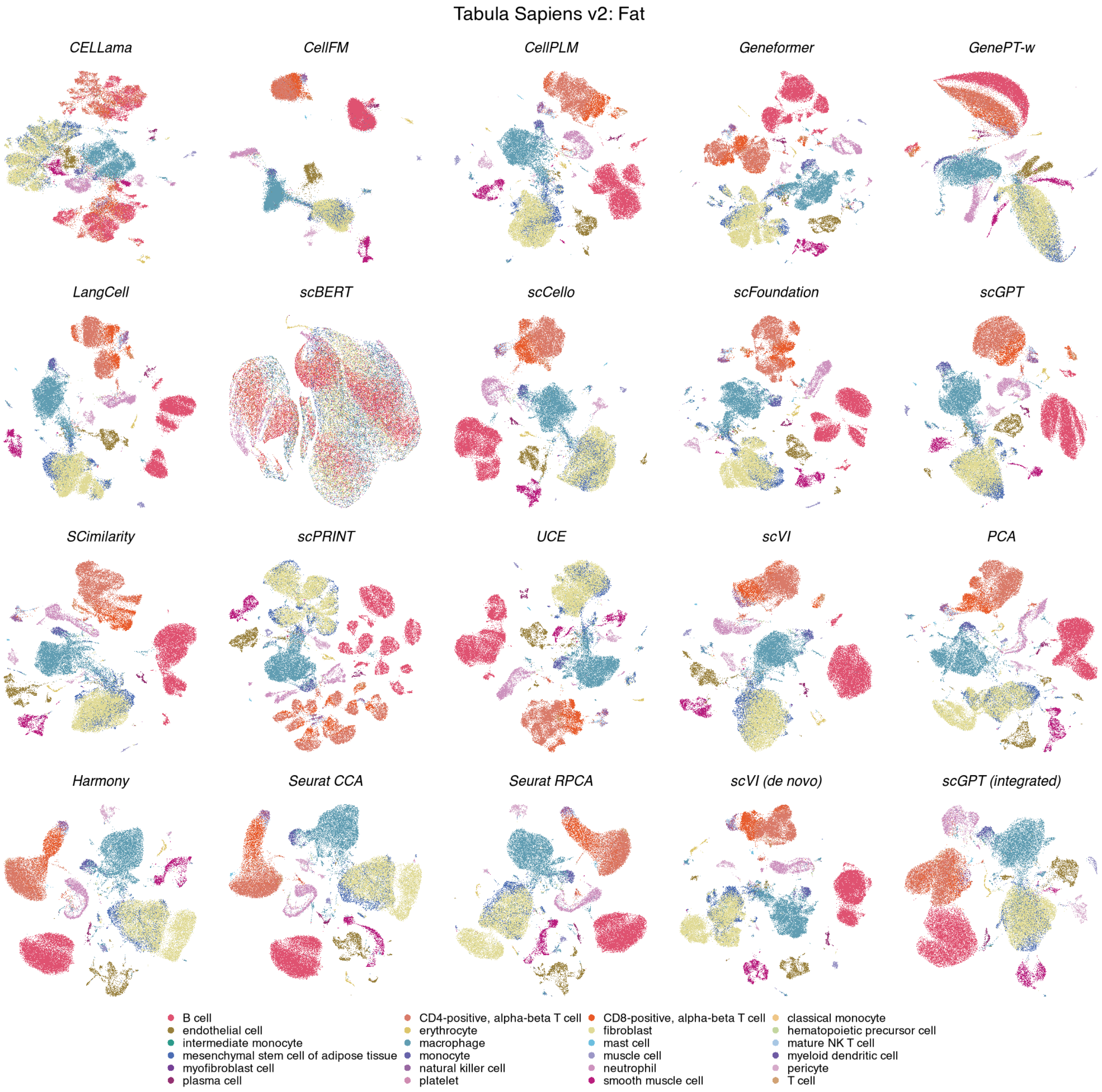

**Figure S26: Cell-type organization in Tabula Sapiens v2 fat.** UMAPs show the same 71,021 cells across 20 methods, coloured by 24 annotated cell types. Display conventions and method categories are as in Fig. S11.

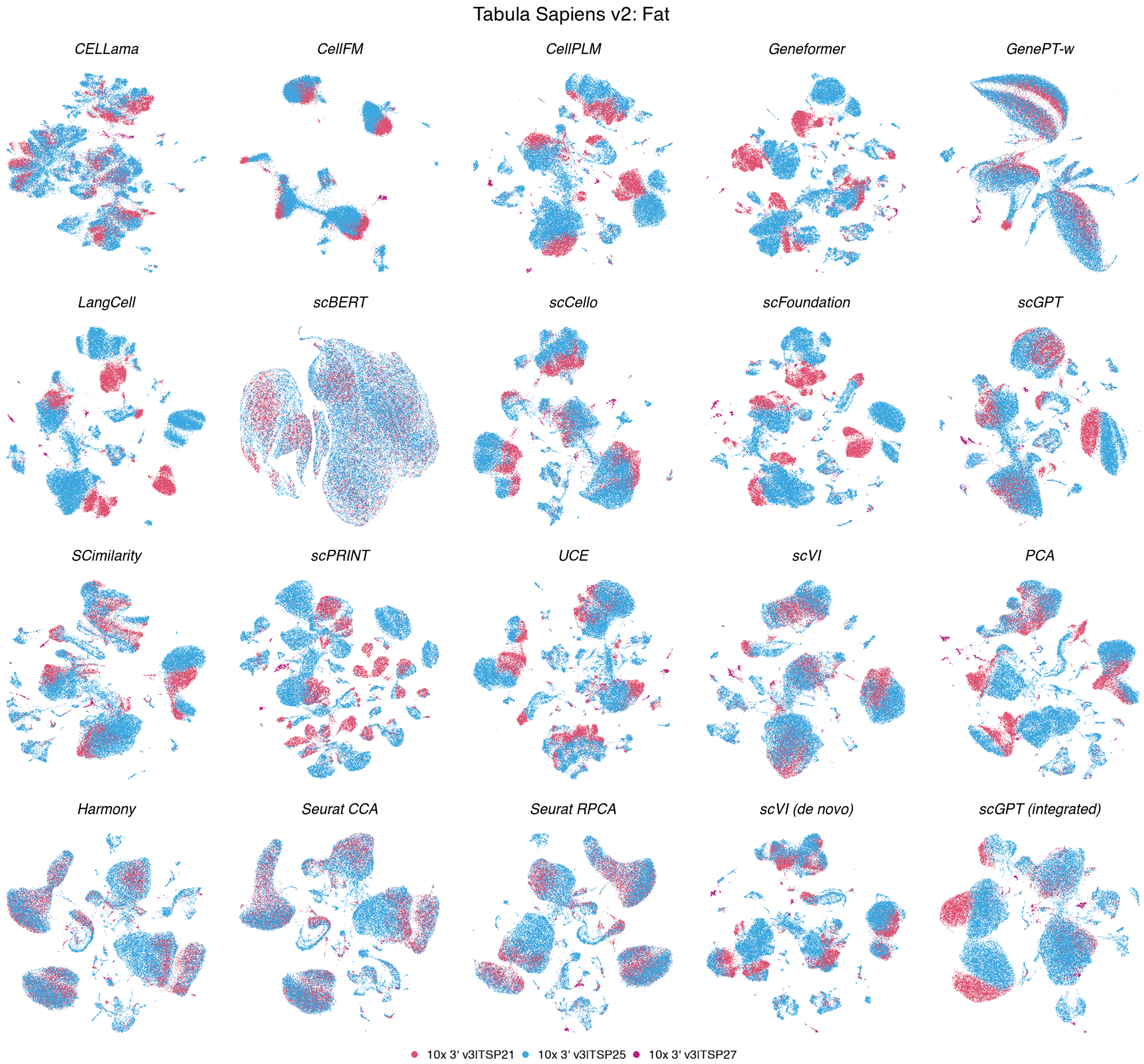

**Figure S27: Batch organization in Tabula Sapiens v2 fat.** UMAPs show the same 71,021 cells across 20 methods, coloured by 3 experimental batches. Coordinates match the corresponding cell-type view; display conventions are as in Fig. S12.

**Tabula Sapiens v2 — Fat**

| method | Biological conservation metrics | | | | | | | | | Batch mixing metrics | | | Aggregated scores | | | | |
|---|---|---|---|---|---|---|---|---|---|---|---|---|---|---|---|---|---|
| | ARI | NMI | HOM | COM | FMI | ASW | Acc@kNN | cLISI | GC | kBET | BRAS | CiLISI | Cluster concordance | Continuous separation | Local structure | Batch mixing | Overall |
| scVI | 0.72 | 0.79 | 0.83 | 0.75 | 0.77 | 0.17 | 0.93 | 1.00 | 0.83 | 0.49 | 0.70 | 0.19 | 0.77 | 0.17 | 0.92 | 0.46 | 0.68 |
| SCimilarity | 0.76 | 0.82 | 0.85 | 0.79 | 0.79 | 0.24 | 0.94 | 1.00 | 0.81 | 0.42 | 0.66 | 0.11 | 0.80 | 0.24 | 0.91 | 0.40 | 0.68 |
| Seurat CCA | 0.62 | 0.75 | 0.82 | 0.69 | 0.68 | 0.17 | 0.92 | 1.00 | 0.75 | 0.55 | 0.69 | 0.31 | 0.71 | 0.17 | 0.89 | 0.52 | 0.67 |
| Seurat RPCA | 0.61 | 0.75 | 0.83 | 0.69 | 0.67 | 0.17 | 0.92 | 1.00 | 0.76 | 0.56 | 0.69 | 0.30 | 0.71 | 0.17 | 0.89 | 0.51 | 0.67 |
| CellPLM | 0.67 | 0.78 | 0.83 | 0.74 | 0.71 | 0.17 | 0.94 | 1.00 | 0.83 | 0.46 | 0.70 | 0.07 | 0.75 | 0.17 | 0.92 | 0.41 | 0.66 |
| Harmony | 0.57 | 0.73 | 0.82 | 0.66 | 0.64 | 0.18 | 0.92 | 1.00 | 0.76 | 0.51 | 0.71 | 0.28 | 0.69 | 0.18 | 0.89 | 0.50 | 0.66 |
| scGPT (integrated) | 0.62 | 0.77 | 0.83 | 0.71 | 0.68 | 0.17 | 0.93 | 1.00 | 0.78 | 0.47 | 0.69 | 0.19 | 0.72 | 0.17 | 0.90 | 0.45 | 0.66 |
| scVI (de novo) | 0.63 | 0.78 | 0.82 | 0.74 | 0.68 | 0.07 | 0.94 | 1.00 | 0.83 | 0.42 | 0.75 | 0.17 | 0.73 | 0.07 | 0.92 | 0.45 | 0.66 |
| scGPT | 0.65 | 0.77 | 0.83 | 0.72 | 0.70 | 0.15 | 0.93 | 1.00 | 0.81 | 0.38 | 0.74 | 0.10 | 0.73 | 0.15 | 0.91 | 0.41 | 0.65 |
| UCE | 0.62 | 0.77 | 0.83 | 0.72 | 0.68 | 0.14 | 0.93 | 1.00 | 0.71 | 0.41 | 0.75 | 0.10 | 0.72 | 0.14 | 0.88 | 0.42 | 0.64 |
| scCello | 0.56 | 0.75 | 0.83 | 0.67 | 0.63 | 0.16 | 0.92 | 1.00 | 0.76 | 0.44 | 0.74 | 0.11 | 0.69 | 0.16 | 0.89 | 0.43 | 0.64 |
| scFoundation | 0.57 | 0.75 | 0.81 | 0.69 | 0.64 | 0.16 | 0.94 | 1.00 | 0.80 | 0.38 | 0.73 | 0.04 | 0.69 | 0.16 | 0.91 | 0.38 | 0.63 |
| LangCell | 0.58 | 0.76 | 0.84 | 0.69 | 0.65 | 0.08 | 0.93 | 1.00 | 0.80 | 0.35 | 0.78 | 0.04 | 0.70 | 0.08 | 0.91 | 0.39 | 0.63 |
| PCA | 0.60 | 0.75 | 0.82 | 0.69 | 0.66 | 0.12 | 0.92 | 1.00 | 0.76 | 0.40 | 0.65 | 0.12 | 0.71 | 0.12 | 0.89 | 0.39 | 0.63 |
| scPRINT | 0.52 | 0.70 | 0.78 | 0.64 | 0.59 | 0.16 | 0.93 | 1.00 | 0.80 | 0.35 | 0.74 | 0.06 | 0.65 | 0.16 | 0.91 | 0.38 | 0.62 |
| Geneformer | 0.51 | 0.71 | 0.81 | 0.64 | 0.59 | 0.08 | 0.93 | 1.00 | 0.77 | 0.35 | 0.79 | 0.05 | 0.65 | 0.08 | 0.90 | 0.39 | 0.61 |
| CellFM | 0.53 | 0.72 | 0.79 | 0.66 | 0.60 | -0.00 | 0.87 | 0.99 | 0.28 | 0.45 | 0.97 | 0.25 | 0.66 | -0.00 | 0.71 | 0.56 | 0.59 |
| CELLama | 0.28 | 0.52 | 0.61 | 0.46 | 0.37 | 0.02 | 0.84 | 0.99 | 0.64 | 0.44 | 0.83 | 0.14 | 0.45 | 0.02 | 0.82 | 0.47 | 0.53 |
| GenePT-w | 0.18 | 0.40 | 0.47 | 0.35 | 0.28 | -0.36 | 0.85 | 0.99 | 0.58 | 0.43 | 0.72 | 0.23 | 0.34 | -0.36 | 0.81 | 0.46 | 0.45 |
| scBERT | 0.06 | 0.15 | 0.18 | 0.13 | 0.15 | -0.46 | 0.55 | 0.94 | 0.27 | 0.19 | 0.62 | 0.31 | 0.13 | -0.46 | 0.59 | 0.37 | 0.28 |

**Figure S28: Representation performance in Tabula Sapiens v2 fat.** Scores are shown for 71,021 cells, 24 annotated cell types and 3 experimental batches. Metrics, family weighting and row ordering are as in Fig. S13.

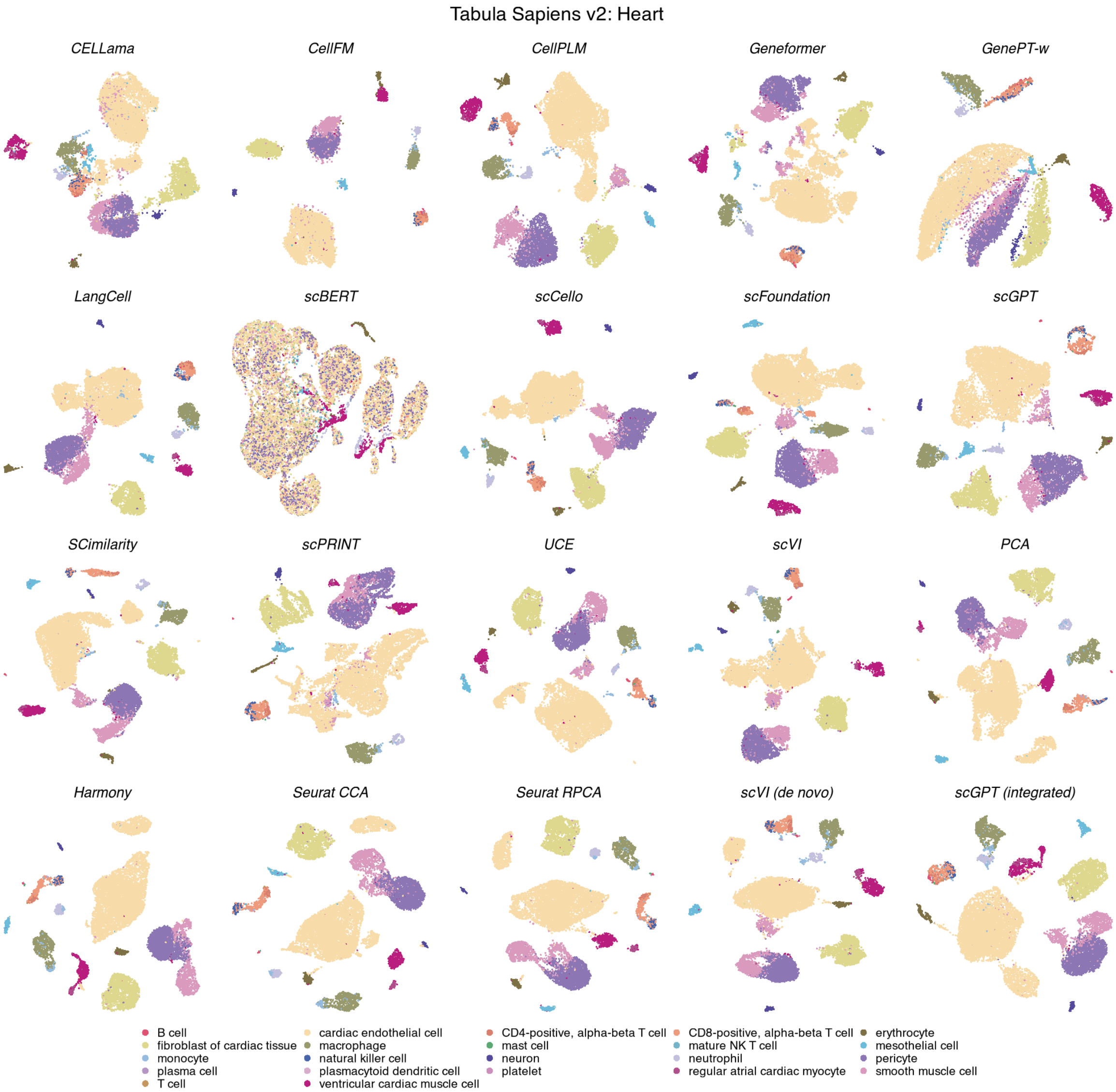

**Figure S29: Cell-type organization in Tabula Sapiens v2 heart.** UMAPs show the same 13,344 cells across 20 methods, coloured by 22 annotated cell types. Display conventions and method categories are as in Fig. S11.

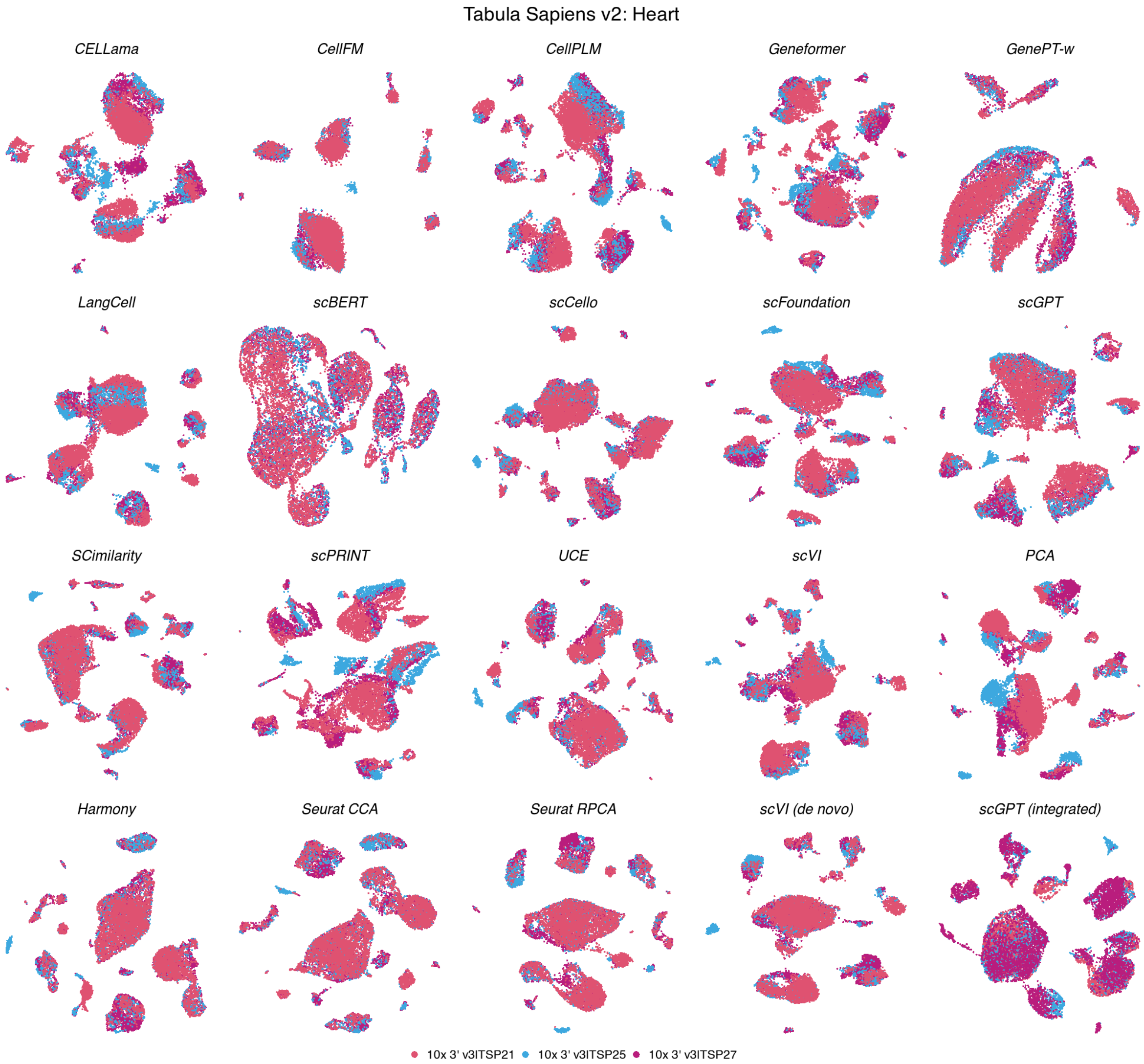

**Figure S30: Batch organization in Tabula Sapiens v2 heart.** UMAPs show the same 13,344 cells across 20 methods, coloured by 3 experimental batches. Coordinates match the corresponding cell-type view; display conventions are as in Fig. S12.

**Tabula Sapiens v2 — Heart**

| method | Biological conservation metrics | | | | | | | | | Batch mixing metrics | | | Aggregated scores | | | | |
|---|---|---|---|---|---|---|---|---|---|---|---|---|---|---|---|---|---|
| | ARI | NMI | HOM | COM | FMI | ASW | Acc@kNN | cLISI | GC | kBET | BRAS | CiLISI | Cluster concordance | Continuous separation | Local structure | Batch mixing | Overall |
| SCimilarity | 0.41 | 0.74 | 0.91 | 0.63 | 0.57 | 0.49 | 0.97 | 1.00 | 0.86 | 0.59 | 0.68 | 0.31 | 0.65 | 0.49 | 0.94 | 0.53 | 0.70 |
| Harmony | 0.37 | 0.71 | 0.90 | 0.58 | 0.53 | 0.24 | 0.97 | 1.00 | 0.85 | 0.68 | 0.71 | 0.44 | 0.62 | 0.24 | 0.94 | 0.61 | 0.68 |
| UCE | 0.37 | 0.71 | 0.90 | 0.59 | 0.53 | 0.29 | 0.97 | 1.00 | 0.80 | 0.56 | 0.80 | 0.35 | 0.62 | 0.29 | 0.92 | 0.57 | 0.67 |
| Seurat RPCA | 0.36 | 0.71 | 0.90 | 0.58 | 0.52 | 0.22 | 0.97 | 1.00 | 0.83 | 0.66 | 0.68 | 0.42 | 0.61 | 0.22 | 0.93 | 0.59 | 0.67 |
| scCello | 0.28 | 0.67 | 0.89 | 0.54 | 0.45 | 0.35 | 0.96 | 1.00 | 0.85 | 0.57 | 0.77 | 0.38 | 0.57 | 0.35 | 0.94 | 0.57 | 0.67 |
| scVI (de novo) | 0.37 | 0.71 | 0.90 | 0.59 | 0.54 | 0.10 | 0.97 | 1.00 | 0.82 | 0.60 | 0.78 | 0.46 | 0.62 | 0.10 | 0.93 | 0.61 | 0.67 |
| Seurat CCA | 0.34 | 0.70 | 0.90 | 0.58 | 0.51 | 0.23 | 0.96 | 1.00 | 0.84 | 0.64 | 0.68 | 0.38 | 0.61 | 0.23 | 0.93 | 0.57 | 0.67 |
| scGPT (integrated) | 0.34 | 0.71 | 0.90 | 0.59 | 0.51 | 0.28 | 0.97 | 1.00 | 0.87 | 0.47 | 0.69 | 0.39 | 0.61 | 0.28 | 0.95 | 0.52 | 0.66 |
| scVI | 0.37 | 0.71 | 0.89 | 0.60 | 0.53 | 0.27 | 0.97 | 1.00 | 0.86 | 0.52 | 0.69 | 0.28 | 0.62 | 0.27 | 0.94 | 0.50 | 0.66 |
| CellPLM | 0.33 | 0.72 | 0.91 | 0.59 | 0.50 | 0.32 | 0.97 | 1.00 | 0.86 | 0.49 | 0.69 | 0.23 | 0.61 | 0.32 | 0.94 | 0.47 | 0.65 |
| scFoundation | 0.31 | 0.70 | 0.91 | 0.57 | 0.49 | 0.29 | 0.97 | 1.00 | 0.86 | 0.51 | 0.74 | 0.24 | 0.59 | 0.29 | 0.94 | 0.50 | 0.65 |
| scGPT | 0.32 | 0.70 | 0.90 | 0.57 | 0.49 | 0.26 | 0.97 | 1.00 | 0.85 | 0.48 | 0.73 | 0.31 | 0.60 | 0.26 | 0.94 | 0.51 | 0.65 |
| Geneformer | 0.41 | 0.69 | 0.86 | 0.57 | 0.56 | 0.11 | 0.97 | 1.00 | 0.84 | 0.50 | 0.80 | 0.22 | 0.62 | 0.11 | 0.93 | 0.51 | 0.64 |
| PCA | 0.37 | 0.69 | 0.88 | 0.57 | 0.53 | 0.18 | 0.97 | 1.00 | 0.85 | 0.51 | 0.66 | 0.17 | 0.61 | 0.18 | 0.94 | 0.45 | 0.63 |
| LangCell | 0.29 | 0.68 | 0.89 | 0.55 | 0.47 | 0.13 | 0.96 | 1.00 | 0.84 | 0.45 | 0.82 | 0.26 | 0.58 | 0.13 | 0.93 | 0.51 | 0.63 |
| scPRINT | 0.24 | 0.62 | 0.82 | 0.50 | 0.41 | 0.20 | 0.95 | 1.00 | 0.82 | 0.53 | 0.71 | 0.18 | 0.52 | 0.20 | 0.92 | 0.47 | 0.61 |
| CellFM | 0.30 | 0.64 | 0.83 | 0.53 | 0.46 | 0.01 | 0.92 | 1.00 | 0.42 | 0.63 | 0.97 | 0.49 | 0.55 | 0.01 | 0.78 | 0.70 | 0.61 |
| CELLama | 0.29 | 0.62 | 0.80 | 0.50 | 0.46 | 0.04 | 0.94 | 1.00 | 0.74 | 0.46 | 0.81 | 0.26 | 0.53 | 0.04 | 0.89 | 0.51 | 0.60 |
| GenePT-w | 0.10 | 0.34 | 0.48 | 0.27 | 0.25 | -0.31 | 0.88 | 0.99 | 0.59 | 0.31 | 0.76 | 0.47 | 0.29 | -0.31 | 0.82 | 0.51 | 0.46 |
| scBERT | 0.02 | 0.10 | 0.13 | 0.08 | 0.16 | -0.54 | 0.52 | 0.93 | 0.33 | 0.27 | 0.58 | 0.49 | 0.10 | -0.54 | 0.59 | 0.45 | 0.28 |

**Figure S31: Representation performance in Tabula Sapiens v2 heart.** Scores are shown for 13,344 cells, 22 annotated cell types and 3 experimental batches. Metrics, family weighting and row ordering are as in Fig. S13.

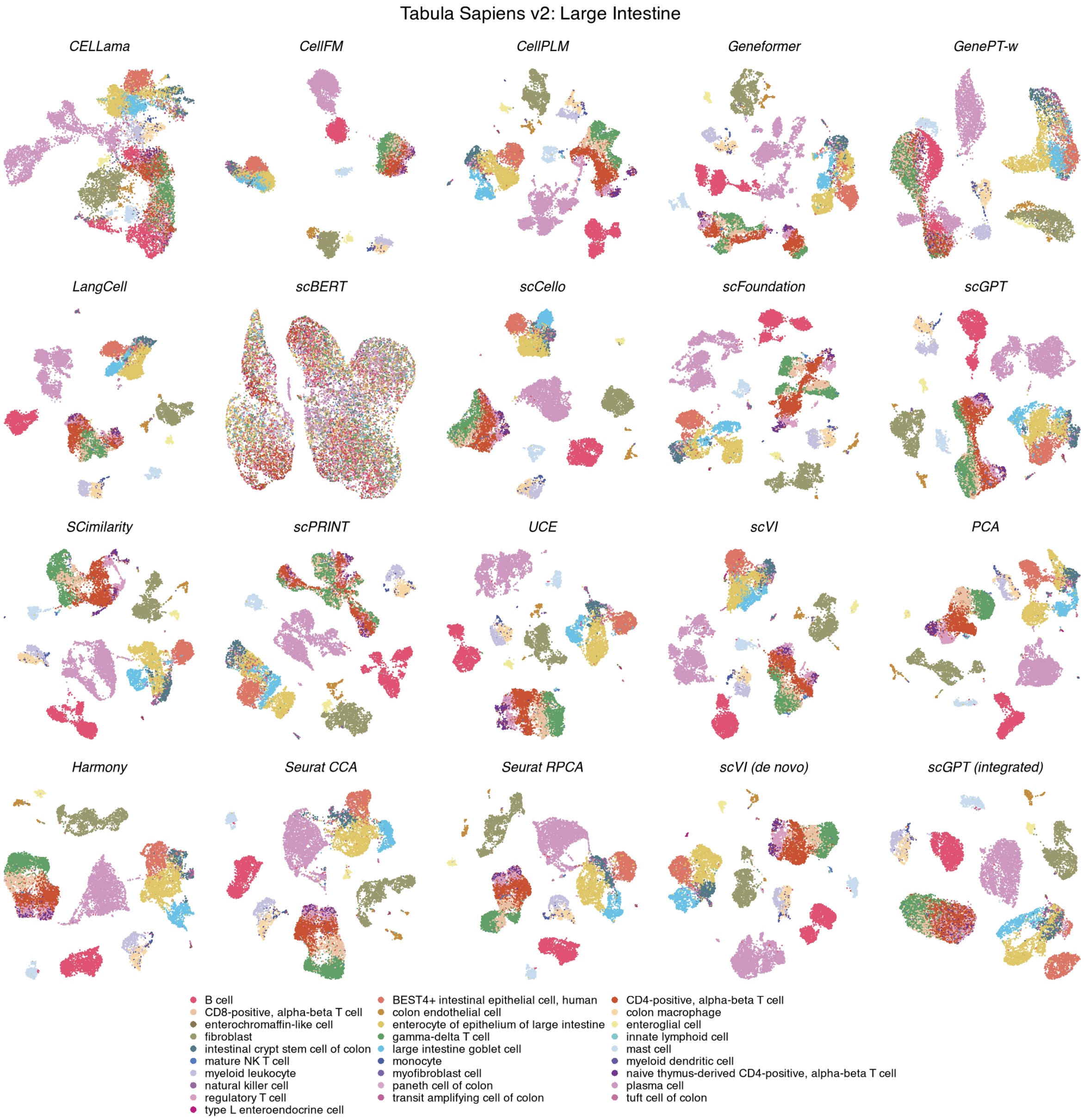

**Figure S32: Cell-type organization in Tabula Sapiens v2 large intestine.** UMAPs show the same 15,231 cells across 20 methods, coloured by 28 annotated cell types. Display conventions and method categories are as in Fig. S11.

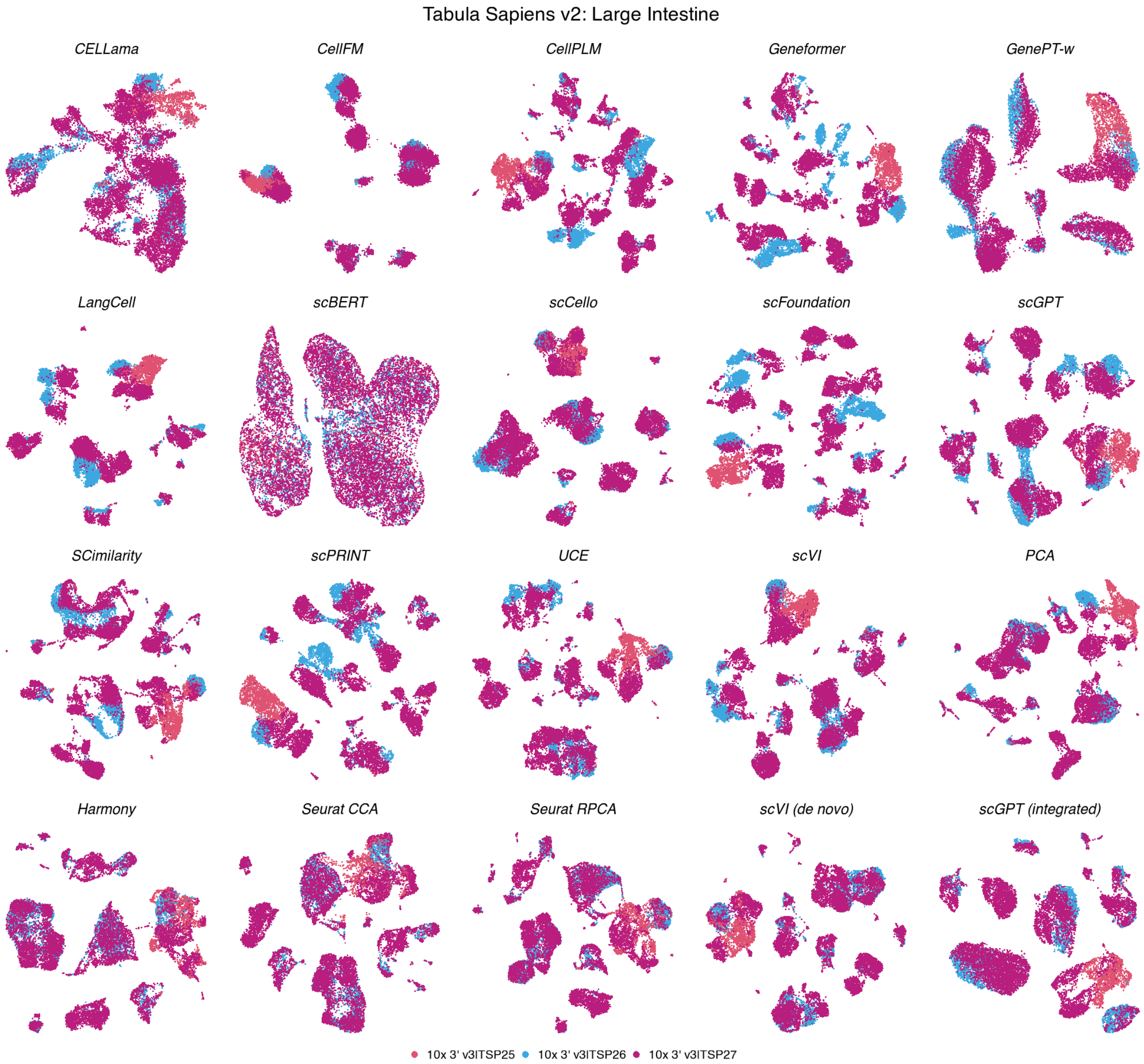

**Figure S33: Batch organization in Tabula Sapiens v2 large intestine.** UMAPs show the same 15,231 cells across 20 methods, coloured by 3 experimental batches. Coordinates match the corresponding cell-type view; display conventions are as in Fig. S12.

**Tabula Sapiens v2 — Large Intestine**

| method | Biological conservation metrics | | | | | | | | | Batch mixing metrics | | | Aggregated scores | | | | |
|---|---|---|---|---|---|---|---|---|---|---|---|---|---|---|---|---|---|
| | ARI | NMI | HOM | COM | FMI | ASW | Acc@kNN | cLISI | GC | kBET | BRAS | CiLISI | Cluster concordance | Continuous separation | Local structure | Batch mixing | Overall |
| scVI (de novo) | 0.67 | 0.82 | 0.87 | 0.78 | 0.71 | 0.11 | 0.94 | 1.00 | 0.86 | 0.47 | 0.75 | 0.24 | 0.77 | 0.11 | 0.93 | 0.48 | 0.69 |
| Harmony | 0.54 | 0.77 | 0.83 | 0.73 | 0.59 | 0.24 | 0.92 | 1.00 | 0.81 | 0.55 | 0.73 | 0.28 | 0.69 | 0.24 | 0.91 | 0.52 | 0.67 |
| Seurat RPCA | 0.53 | 0.77 | 0.83 | 0.71 | 0.59 | 0.26 | 0.92 | 1.00 | 0.83 | 0.55 | 0.68 | 0.28 | 0.69 | 0.26 | 0.92 | 0.50 | 0.67 |
| SCimilarity | 0.57 | 0.79 | 0.84 | 0.75 | 0.62 | 0.34 | 0.94 | 1.00 | 0.83 | 0.41 | 0.66 | 0.13 | 0.71 | 0.34 | 0.92 | 0.40 | 0.67 |
| scGPT (integrated) | 0.53 | 0.77 | 0.85 | 0.71 | 0.59 | 0.37 | 0.92 | 1.00 | 0.83 | 0.40 | 0.65 | 0.21 | 0.69 | 0.37 | 0.92 | 0.42 | 0.66 |
| UCE | 0.57 | 0.80 | 0.85 | 0.76 | 0.62 | 0.22 | 0.94 | 1.00 | 0.84 | 0.39 | 0.75 | 0.15 | 0.72 | 0.22 | 0.92 | 0.43 | 0.66 |
| Seurat CCA | 0.51 | 0.76 | 0.81 | 0.71 | 0.56 | 0.25 | 0.91 | 1.00 | 0.79 | 0.57 | 0.67 | 0.30 | 0.67 | 0.25 | 0.90 | 0.51 | 0.66 |
| PCA | 0.63 | 0.80 | 0.87 | 0.75 | 0.67 | 0.22 | 0.94 | 1.00 | 0.84 | 0.40 | 0.61 | 0.13 | 0.74 | 0.22 | 0.93 | 0.38 | 0.66 |
| scCello | 0.48 | 0.73 | 0.80 | 0.67 | 0.55 | 0.20 | 0.91 | 1.00 | 0.78 | 0.47 | 0.78 | 0.19 | 0.65 | 0.20 | 0.89 | 0.48 | 0.64 |
| scGPT | 0.56 | 0.76 | 0.81 | 0.72 | 0.61 | 0.11 | 0.91 | 1.00 | 0.80 | 0.37 | 0.71 | 0.13 | 0.69 | 0.11 | 0.90 | 0.40 | 0.63 |
| scFoundation | 0.49 | 0.77 | 0.85 | 0.70 | 0.55 | 0.19 | 0.94 | 1.00 | 0.83 | 0.28 | 0.70 | 0.07 | 0.67 | 0.19 | 0.93 | 0.35 | 0.63 |
| CellPLM | 0.47 | 0.75 | 0.82 | 0.70 | 0.53 | 0.20 | 0.94 | 1.00 | 0.84 | 0.32 | 0.64 | 0.09 | 0.65 | 0.20 | 0.92 | 0.35 | 0.62 |
| scVI | 0.49 | 0.75 | 0.80 | 0.71 | 0.54 | 0.17 | 0.90 | 1.00 | 0.81 | 0.35 | 0.66 | 0.13 | 0.66 | 0.17 | 0.90 | 0.38 | 0.62 |
| LangCell | 0.47 | 0.75 | 0.82 | 0.69 | 0.53 | 0.09 | 0.91 | 1.00 | 0.80 | 0.32 | 0.82 | 0.10 | 0.65 | 0.09 | 0.90 | 0.41 | 0.62 |
| scPRINT | 0.43 | 0.71 | 0.77 | 0.65 | 0.49 | 0.10 | 0.89 | 1.00 | 0.73 | 0.39 | 0.74 | 0.11 | 0.61 | 0.10 | 0.87 | 0.41 | 0.60 |
| Geneformer | 0.44 | 0.70 | 0.76 | 0.65 | 0.49 | 0.01 | 0.93 | 1.00 | 0.80 | 0.30 | 0.77 | 0.08 | 0.61 | 0.01 | 0.91 | 0.38 | 0.59 |
| CellFM | 0.46 | 0.71 | 0.78 | 0.65 | 0.52 | 0.00 | 0.81 | 0.98 | 0.32 | 0.41 | 0.96 | 0.26 | 0.63 | 0.00 | 0.70 | 0.54 | 0.57 |
| CELLama | 0.35 | 0.60 | 0.66 | 0.56 | 0.41 | -0.01 | 0.79 | 0.98 | 0.60 | 0.38 | 0.80 | 0.18 | 0.51 | -0.01 | 0.79 | 0.45 | 0.54 |
| GenePT-w | 0.11 | 0.34 | 0.38 | 0.31 | 0.18 | -0.29 | 0.72 | 0.96 | 0.43 | 0.36 | 0.71 | 0.24 | 0.26 | -0.29 | 0.70 | 0.44 | 0.40 |
| scBERT | 0.03 | 0.10 | 0.11 | 0.09 | 0.09 | -0.48 | 0.32 | 0.86 | 0.12 | 0.17 | 0.64 | 0.35 | 0.08 | -0.48 | 0.44 | 0.39 | 0.22 |

**Figure S34: Representation performance in Tabula Sapiens v2 large intestine.** Scores are shown for 15,231 cells, 28 annotated cell types and 3 experimental batches. Metrics, family weighting and row ordering are as in Fig. S13.

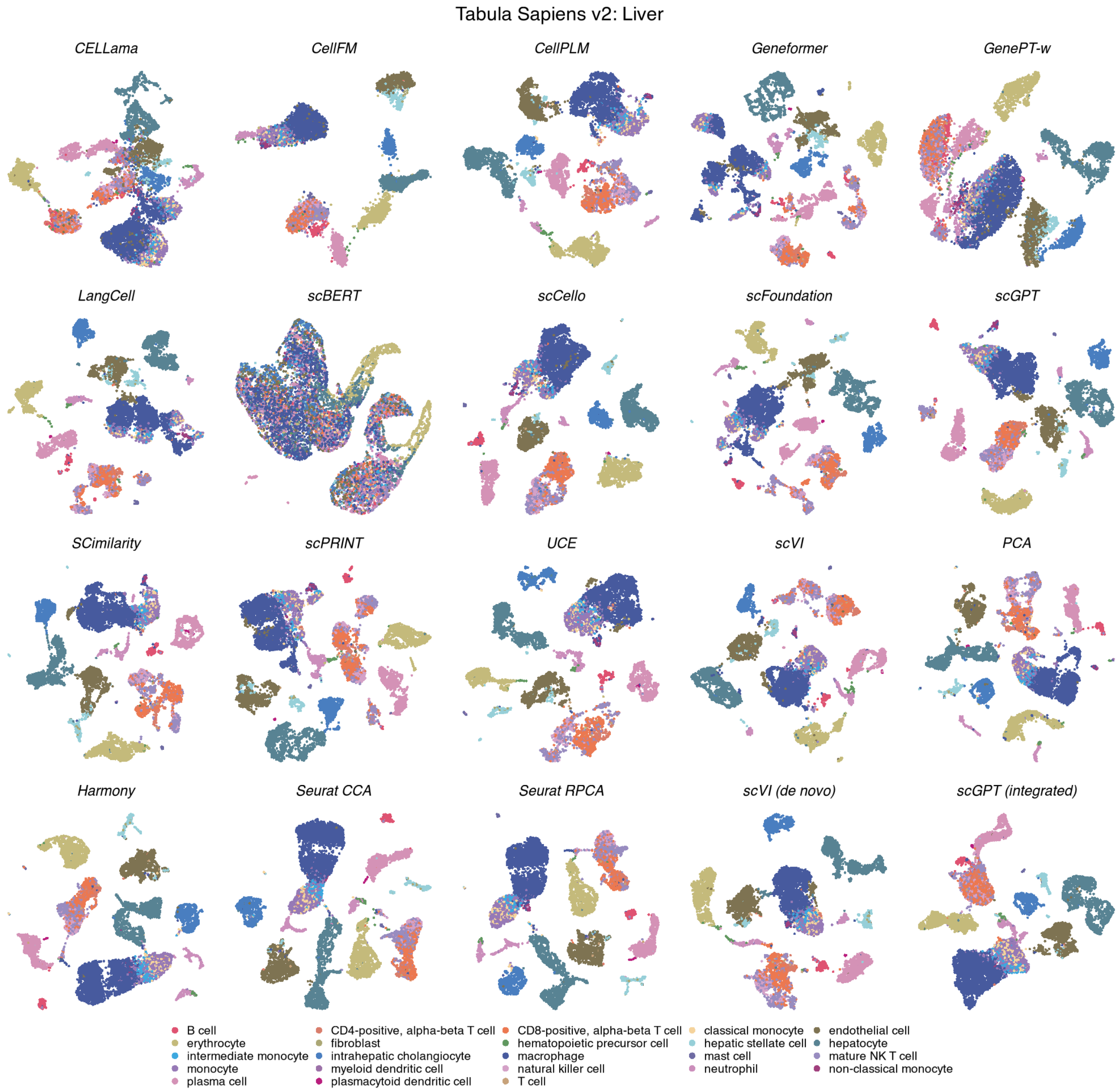

**Figure S35: Cell-type organization in Tabula Sapiens v2 liver.** UMAPs show the same 10,002 cells across 20 methods, coloured by 23 annotated cell types. Display conventions and method categories are as in Fig. S11.

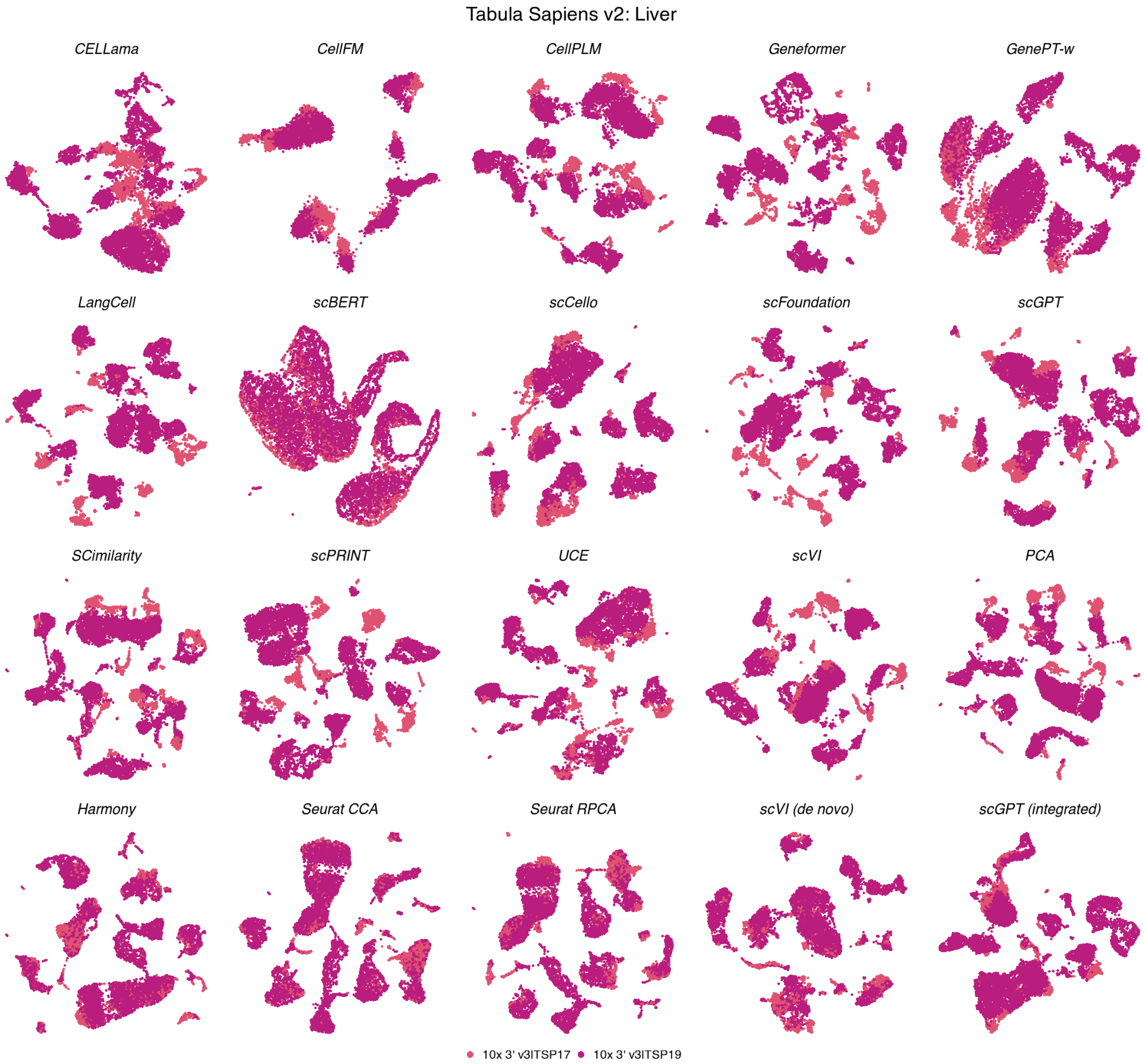

**Figure S36: Batch organization in Tabula Sapiens v2 liver.** UMAPs show the same 10,002 cells across 20 methods, coloured by 2 experimental batches. Coordinates match the corresponding cell-type view; display conventions are as in Fig. S12.

**Tabula Sapiens v2 — Liver**

| | Biological conservation metrics | | | | | | | | | Batch mixing metrics | | | Aggregated scores | | | | |
|---|---|---|---|---|---|---|---|---|---|---|---|---|---|---|---|---|---|
| **method** | ARI | NMI | HOM | COM | FMI | ASW | Acc@kNN | cLISI | GC | kBET | BRAS | CiLISI | Cluster concordance | Continuous separation | Local structure | Batch mixing | Overall |
| **Seurat CCA** | 0.59 | 0.79 | 0.81 | 0.77 | 0.63 | 0.11 | 0.85 | 1.00 | 0.74 | 0.56 | 0.75 | 0.31 | 0.72 | 0.11 | 0.87 | 0.54 | 0.66 |
| **Harmony** | 0.58 | 0.79 | 0.82 | 0.76 | 0.63 | 0.11 | 0.86 | 1.00 | 0.75 | 0.57 | 0.76 | 0.25 | 0.72 | 0.11 | 0.87 | 0.53 | 0.66 |
| **Seurat RPCA** | 0.58 | 0.79 | 0.82 | 0.77 | 0.63 | 0.12 | 0.87 | 1.00 | 0.77 | 0.51 | 0.74 | 0.24 | 0.72 | 0.12 | 0.88 | 0.49 | 0.65 |
| **scVI (de novo)** | 0.58 | 0.78 | 0.83 | 0.74 | 0.63 | 0.11 | 0.87 | 1.00 | 0.81 | 0.42 | 0.79 | 0.20 | 0.71 | 0.11 | 0.90 | 0.47 | 0.65 |
| **SCimilarity** | 0.65 | 0.81 | 0.83 | 0.78 | 0.69 | 0.30 | 0.89 | 1.00 | 0.77 | 0.29 | 0.65 | 0.09 | 0.75 | 0.30 | 0.88 | 0.34 | 0.65 |
| **scCello** | 0.54 | 0.77 | 0.82 | 0.73 | 0.60 | 0.19 | 0.87 | 1.00 | 0.80 | 0.35 | 0.71 | 0.11 | 0.69 | 0.19 | 0.89 | 0.39 | 0.63 |
| **scVI** | 0.68 | 0.78 | 0.79 | 0.76 | 0.71 | 0.16 | 0.87 | 1.00 | 0.78 | 0.30 | 0.61 | 0.08 | 0.75 | 0.16 | 0.88 | 0.33 | 0.63 |
| **scGPT (integrated)** | 0.49 | 0.76 | 0.80 | 0.71 | 0.55 | 0.23 | 0.86 | 1.00 | 0.74 | 0.40 | 0.67 | 0.13 | 0.66 | 0.23 | 0.87 | 0.40 | 0.62 |
| **UCE** | 0.53 | 0.78 | 0.80 | 0.75 | 0.57 | 0.17 | 0.88 | 1.00 | 0.73 | 0.28 | 0.74 | 0.13 | 0.69 | 0.17 | 0.87 | 0.38 | 0.62 |
| **CellPLM** | 0.58 | 0.78 | 0.82 | 0.74 | 0.63 | 0.16 | 0.88 | 1.00 | 0.80 | 0.25 | 0.65 | 0.07 | 0.71 | 0.16 | 0.90 | 0.32 | 0.62 |
| **scFoundation** | 0.57 | 0.77 | 0.83 | 0.72 | 0.63 | 0.17 | 0.88 | 1.00 | 0.78 | 0.27 | 0.64 | 0.05 | 0.70 | 0.17 | 0.89 | 0.32 | 0.61 |
| **PCA** | 0.55 | 0.78 | 0.83 | 0.73 | 0.61 | 0.12 | 0.88 | 1.00 | 0.78 | 0.30 | 0.62 | 0.07 | 0.70 | 0.12 | 0.88 | 0.33 | 0.61 |
| **scGPT** | 0.52 | 0.76 | 0.79 | 0.74 | 0.57 | 0.11 | 0.88 | 1.00 | 0.78 | 0.28 | 0.59 | 0.08 | 0.68 | 0.11 | 0.88 | 0.32 | 0.60 |
| **scPRINT** | 0.54 | 0.74 | 0.80 | 0.70 | 0.59 | 0.17 | 0.87 | 1.00 | 0.76 | 0.26 | 0.59 | 0.04 | 0.67 | 0.17 | 0.88 | 0.30 | 0.60 |
| **LangCell** | 0.48 | 0.71 | 0.77 | 0.67 | 0.54 | 0.09 | 0.87 | 1.00 | 0.76 | 0.24 | 0.73 | 0.05 | 0.63 | 0.09 | 0.88 | 0.34 | 0.58 |
| **Geneformer** | 0.44 | 0.69 | 0.74 | 0.64 | 0.49 | 0.05 | 0.87 | 1.00 | 0.72 | 0.24 | 0.63 | 0.04 | 0.60 | 0.05 | 0.86 | 0.31 | 0.56 |
| **CellFM** | 0.47 | 0.71 | 0.75 | 0.67 | 0.53 | -0.02 | 0.79 | 0.99 | 0.28 | 0.33 | 0.95 | 0.19 | 0.63 | -0.02 | 0.68 | 0.49 | 0.55 |
| **CELLama** | 0.42 | 0.64 | 0.68 | 0.60 | 0.48 | 0.03 | 0.81 | 0.99 | 0.62 | 0.27 | 0.68 | 0.09 | 0.57 | 0.03 | 0.81 | 0.34 | 0.54 |
| **GenePT-w** | 0.14 | 0.37 | 0.41 | 0.34 | 0.21 | -0.33 | 0.74 | 0.96 | 0.45 | 0.38 | 0.58 | 0.27 | 0.29 | -0.33 | 0.72 | 0.41 | 0.40 |
| **scBERT** | 0.08 | 0.19 | 0.21 | 0.18 | 0.15 | -0.51 | 0.45 | 0.88 | 0.18 | 0.11 | 0.65 | 0.23 | 0.16 | -0.51 | 0.50 | 0.33 | 0.25 |

**Figure S37: Representation performance in Tabula Sapiens v2 liver.** Scores are shown for 10,002 cells, 23 annotated cell types and 2 experimental batches. Metrics, family weighting and row ordering are as in Fig. S13.

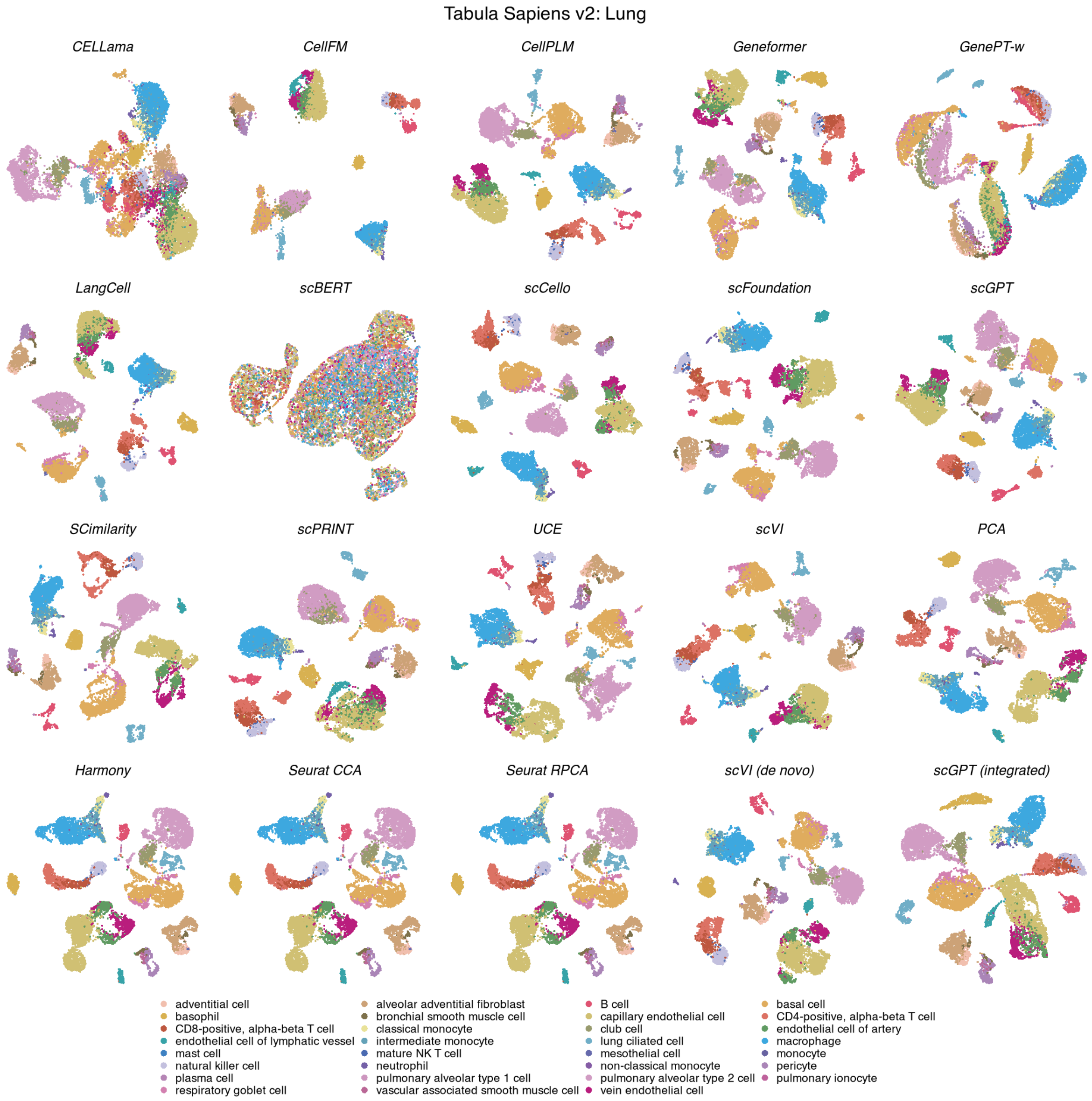

**Figure S38: Cell-type organization in Tabula Sapiens v2 lung.** UMAPs show the same 11,716 cells across 20 methods, coloured by 31 annotated cell types. Display conventions and method categories are as in Fig. S11.

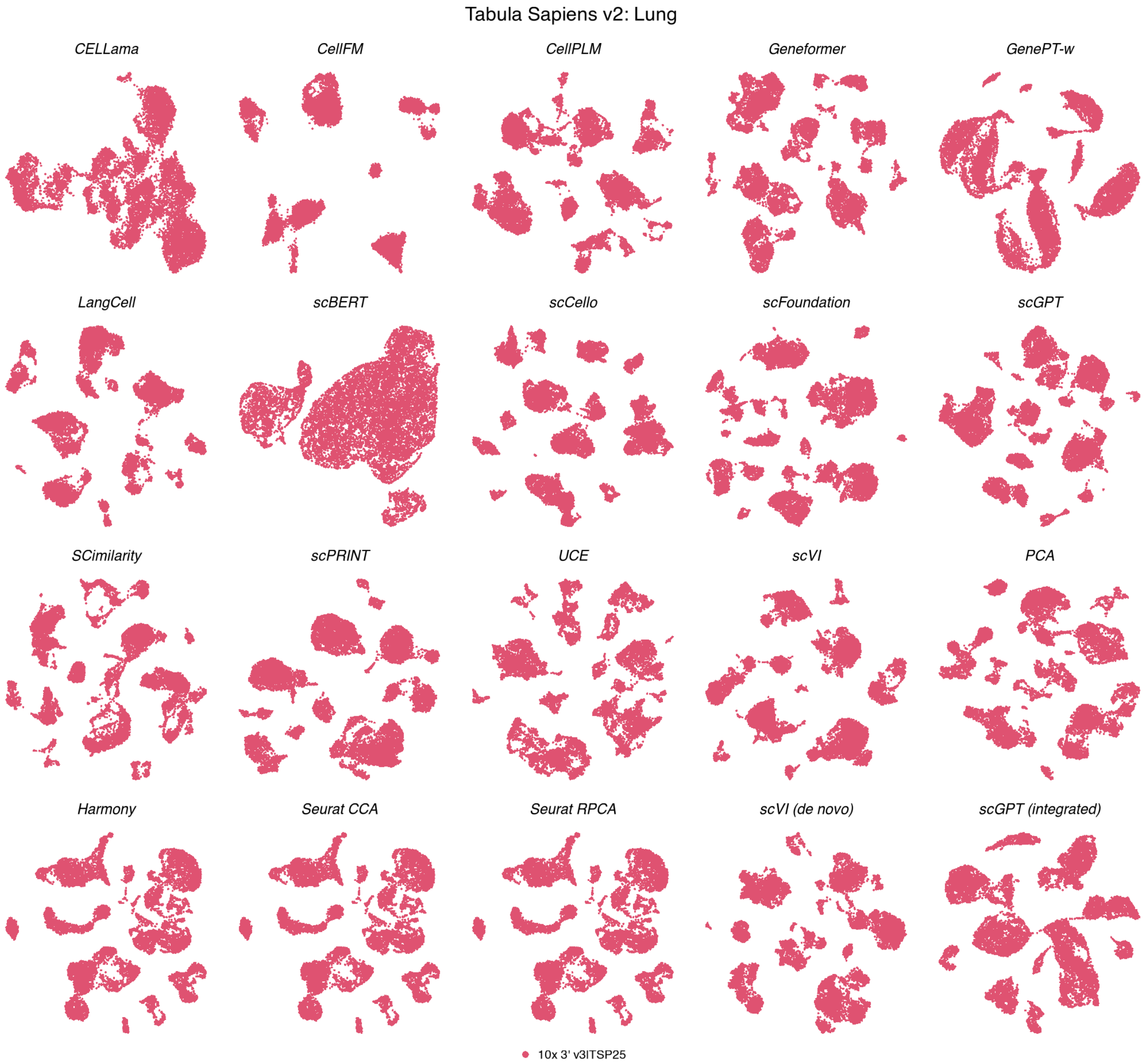

**Figure S39: Batch organization in Tabula Sapiens v2 lung.** UMAPs show the same 11,716 cells across 20 methods, coloured by a single experimental batch. Coordinates match the corresponding cell-type view; display conventions are as in Fig. S12.

**Tabula Sapiens v2 — Lung**

| method | Biological conservation metrics | | | | | | | | | Batch mixing metrics | | | Aggregated scores | | | | |
|---|---|---|---|---|---|---|---|---|---|---|---|---|---|---|---|---|---|
| | ARI | NMI | HOM | COM | FMI | ASW | Acc@kNN | cLISI | GC | kBET | BRAS | CiLISI | Cluster concordance | Continuous separation | Local structure | Batch mixing | Overall |
| SCimilarity | 0.65 | 0.83 | 0.87 | 0.79 | 0.69 | 0.37 | 0.94 | 1.00 | 0.84 | nan | nan | nan | 0.77 | 0.37 | 0.93 | | 0.78 |
| PCA | 0.67 | 0.83 | 0.88 | 0.79 | 0.71 | 0.26 | 0.94 | 1.00 | 0.87 | nan | nan | nan | 0.77 | 0.26 | 0.94 | | 0.78 |
| scGPT (integrated) | 0.60 | 0.82 | 0.89 | 0.77 | 0.65 | 0.32 | 0.94 | 1.00 | 0.85 | nan | nan | nan | 0.75 | 0.32 | 0.93 | | 0.77 |
| Harmony | 0.58 | 0.81 | 0.88 | 0.75 | 0.63 | 0.29 | 0.94 | 1.00 | 0.87 | nan | nan | nan | 0.73 | 0.29 | 0.94 | | 0.76 |
| Seurat CCA | 0.58 | 0.81 | 0.88 | 0.75 | 0.63 | 0.29 | 0.94 | 1.00 | 0.87 | nan | nan | nan | 0.73 | 0.29 | 0.94 | | 0.76 |
| Seurat RPCA | 0.58 | 0.81 | 0.88 | 0.75 | 0.63 | 0.29 | 0.94 | 1.00 | 0.87 | nan | nan | nan | 0.73 | 0.29 | 0.94 | | 0.76 |
| scFoundation | 0.63 | 0.82 | 0.88 | 0.77 | 0.68 | 0.20 | 0.94 | 1.00 | 0.85 | nan | nan | nan | 0.76 | 0.20 | 0.93 | | 0.76 |
| CellPLM | 0.65 | 0.82 | 0.87 | 0.78 | 0.68 | 0.18 | 0.93 | 1.00 | 0.85 | nan | nan | nan | 0.76 | 0.18 | 0.93 | | 0.76 |
| scVI (de novo) | 0.66 | 0.83 | 0.88 | 0.79 | 0.70 | 0.14 | 0.93 | 1.00 | 0.84 | nan | nan | nan | 0.77 | 0.14 | 0.93 | | 0.75 |
| scVI | 0.64 | 0.81 | 0.85 | 0.78 | 0.67 | 0.18 | 0.93 | 1.00 | 0.85 | nan | nan | nan | 0.75 | 0.18 | 0.93 | | 0.75 |
| scGPT | 0.60 | 0.81 | 0.85 | 0.77 | 0.64 | 0.14 | 0.93 | 1.00 | 0.83 | nan | nan | nan | 0.73 | 0.14 | 0.92 | | 0.74 |
| UCE | 0.57 | 0.81 | 0.86 | 0.76 | 0.62 | 0.20 | 0.93 | 1.00 | 0.81 | nan | nan | nan | 0.72 | 0.20 | 0.91 | | 0.74 |
| scCello | 0.53 | 0.80 | 0.86 | 0.74 | 0.59 | 0.21 | 0.92 | 1.00 | 0.82 | nan | nan | nan | 0.70 | 0.21 | 0.91 | | 0.73 |
| LangCell | 0.52 | 0.78 | 0.83 | 0.74 | 0.57 | 0.08 | 0.91 | 1.00 | 0.83 | nan | nan | nan | 0.69 | 0.08 | 0.91 | | 0.71 |
| Geneformer | 0.52 | 0.77 | 0.83 | 0.72 | 0.57 | 0.05 | 0.93 | 1.00 | 0.78 | nan | nan | nan | 0.68 | 0.05 | 0.90 | | 0.69 |
| scPRINT | 0.47 | 0.75 | 0.81 | 0.70 | 0.52 | 0.10 | 0.90 | 1.00 | 0.82 | nan | nan | nan | 0.65 | 0.10 | 0.91 | | 0.69 |
| CellFM | 0.51 | 0.76 | 0.82 | 0.71 | 0.56 | -0.00 | 0.84 | 0.98 | 0.39 | nan | nan | nan | 0.67 | -0.00 | 0.74 | | 0.61 |
| CELLama | 0.40 | 0.65 | 0.71 | 0.61 | 0.45 | 0.04 | 0.83 | 0.99 | 0.67 | nan | nan | nan | 0.56 | 0.04 | 0.83 | | 0.61 |
| GenePT-w | 0.13 | 0.34 | 0.38 | 0.32 | 0.19 | -0.41 | 0.72 | 0.96 | 0.39 | nan | nan | nan | 0.27 | -0.41 | 0.69 | | 0.36 |
| scBERT | 0.04 | 0.12 | 0.13 | 0.11 | 0.10 | -0.44 | 0.35 | 0.89 | 0.17 | nan | nan | nan | 0.10 | -0.44 | 0.47 | | 0.19 |

**Figure S40: Representation performance in Tabula Sapiens v2 lung.** Scores are shown for 11,716 cells, 31 annotated cell types and a single experimental batch. Metrics, family weighting and row ordering are as in Fig. S13. Batch-mixing metrics were unavailable, and the remaining family weights were renormalized.

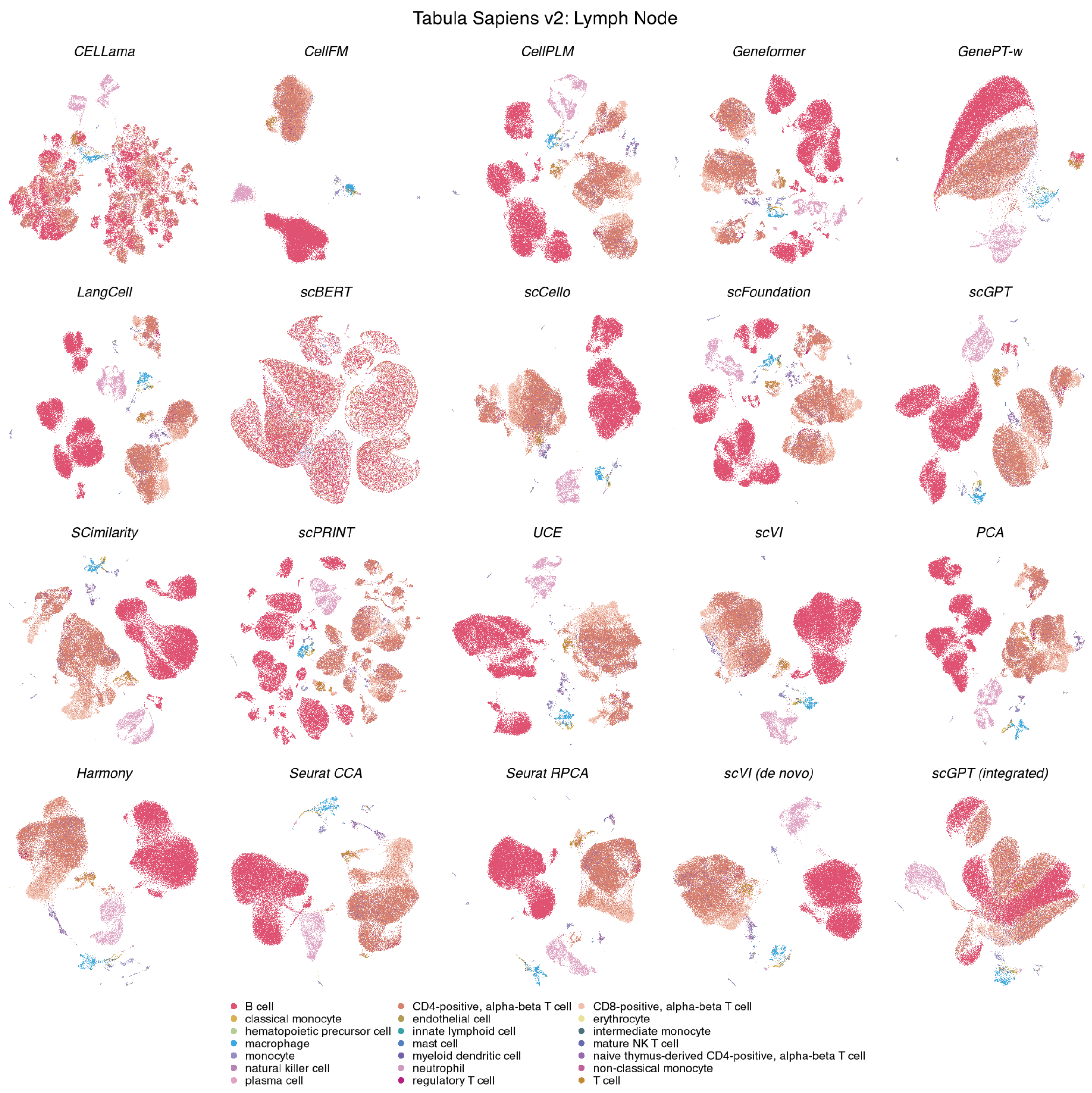

**Figure S41: Cell-type organization in Tabula Sapiens v2 lymph node.** UMAPs show the same 72,290 cells across 20 methods, coloured by 21 annotated cell types. Display conventions and method categories are as in Fig. S11.

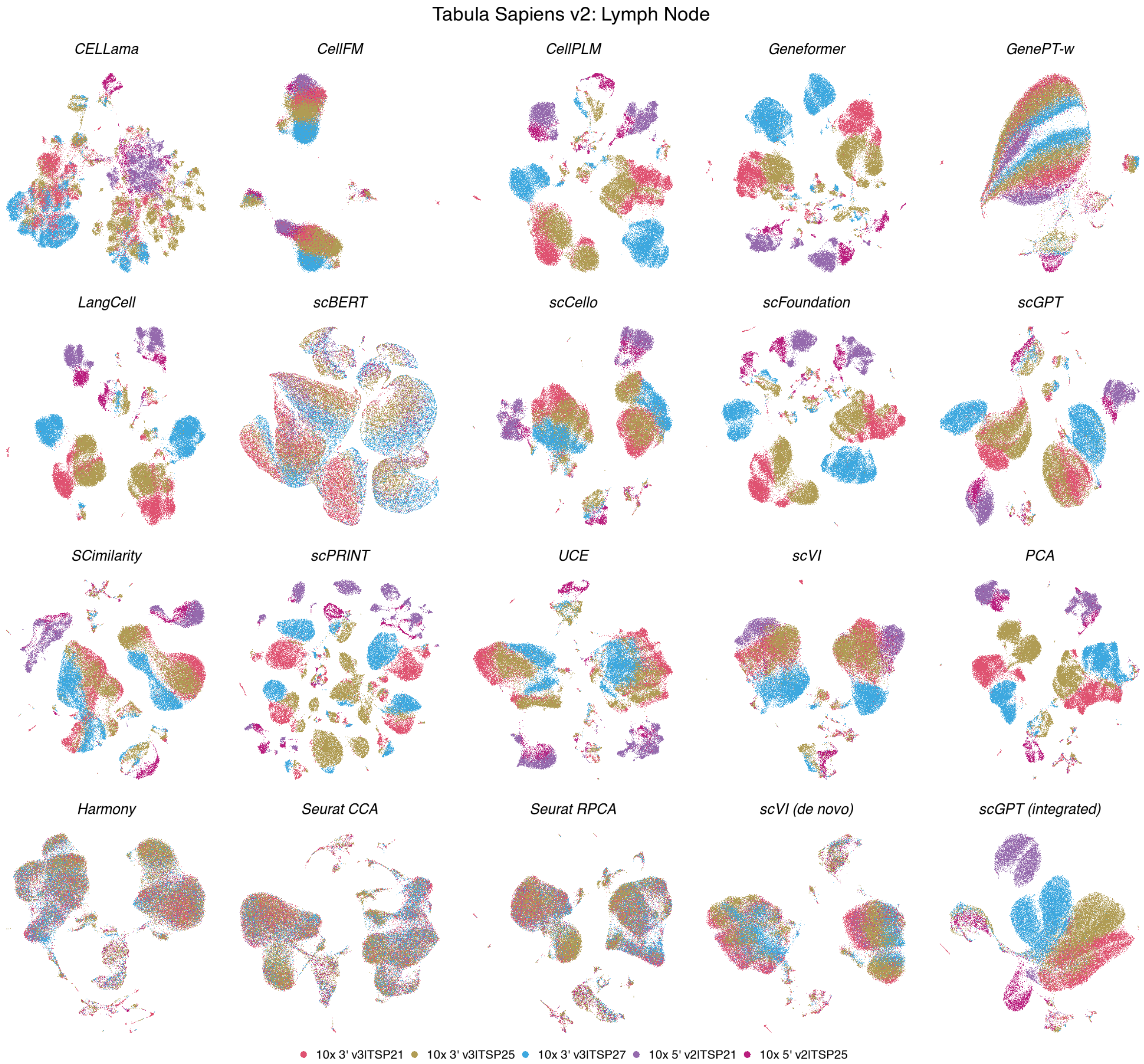

**Figure S42: Batch organization in Tabula Sapiens v2 lymph node.** UMAPs show the same 72,290 cells across 20 methods, coloured by 5 experimental batches. Coordinates match the corresponding cell-type view; display conventions are as in Fig. S12.

**Tabula Sapiens v2 — Lymph Node**

| | Biological conservation metrics | | | | | | | | | Batch mixing metrics | | | Aggregated scores | | | | |
|---|---|---|---|---|---|---|---|---|---|---|---|---|---|---|---|---|---|
| **method** | ARI | NMI | HOM | COM | FMI | ASW | Acc@kNN | cLISI | GC | kBET | BRAS | CiLISI | Cluster concordance | Continuous separation | Local structure | Batch mixing | Overall |
| **Seurat RPCA** | 0.30 | 0.58 | 0.80 | 0.46 | 0.48 | 0.13 | 0.91 | 1.00 | 0.73 | 0.62 | 0.74 | 0.44 | 0.52 | 0.13 | 0.88 | 0.60 | 0.62 |
| **Harmony** | 0.29 | 0.57 | 0.80 | 0.44 | 0.48 | 0.13 | 0.91 | 1.00 | 0.71 | 0.53 | 0.72 | 0.40 | 0.52 | 0.13 | 0.87 | 0.55 | 0.60 |
| **Seurat CCA** | 0.26 | 0.55 | 0.78 | 0.42 | 0.45 | 0.19 | 0.90 | 1.00 | 0.71 | 0.59 | 0.70 | 0.37 | 0.49 | 0.19 | 0.87 | 0.55 | 0.60 |
| **scVI (de novo)** | 0.32 | 0.56 | 0.76 | 0.45 | 0.50 | 0.04 | 0.90 | 1.00 | 0.77 | 0.39 | 0.78 | 0.27 | 0.52 | 0.04 | 0.89 | 0.48 | 0.58 |
| **SCimilarity** | 0.31 | 0.58 | 0.79 | 0.46 | 0.49 | 0.26 | 0.91 | 1.00 | 0.73 | 0.32 | 0.64 | 0.08 | 0.53 | 0.26 | 0.88 | 0.35 | 0.57 |
| **scGPT (integrated)** | 0.28 | 0.55 | 0.77 | 0.43 | 0.47 | 0.15 | 0.91 | 1.00 | 0.75 | 0.39 | 0.67 | 0.18 | 0.50 | 0.15 | 0.88 | 0.42 | 0.57 |
| **scCello** | 0.24 | 0.53 | 0.76 | 0.41 | 0.42 | 0.18 | 0.91 | 1.00 | 0.71 | 0.34 | 0.70 | 0.09 | 0.47 | 0.18 | 0.87 | 0.38 | 0.55 |
| **UCE** | 0.29 | 0.55 | 0.77 | 0.43 | 0.48 | 0.15 | 0.91 | 1.00 | 0.65 | 0.30 | 0.68 | 0.07 | 0.50 | 0.15 | 0.85 | 0.35 | 0.54 |
| **scVI** | 0.26 | 0.53 | 0.74 | 0.41 | 0.44 | 0.16 | 0.91 | 1.00 | 0.74 | 0.32 | 0.57 | 0.09 | 0.47 | 0.16 | 0.88 | 0.33 | 0.54 |
| **scGPT** | 0.24 | 0.53 | 0.73 | 0.41 | 0.43 | 0.17 | 0.91 | 1.00 | 0.71 | 0.25 | 0.65 | 0.06 | 0.47 | 0.17 | 0.87 | 0.32 | 0.53 |
| **PCA** | 0.22 | 0.52 | 0.76 | 0.39 | 0.41 | 0.11 | 0.91 | 1.00 | 0.74 | 0.35 | 0.64 | 0.03 | 0.46 | 0.11 | 0.88 | 0.34 | 0.53 |
| **scFoundation** | 0.23 | 0.53 | 0.77 | 0.40 | 0.42 | 0.14 | 0.91 | 1.00 | 0.71 | 0.25 | 0.61 | 0.02 | 0.47 | 0.14 | 0.87 | 0.30 | 0.52 |
| **LangCell** | 0.22 | 0.52 | 0.75 | 0.40 | 0.41 | 0.06 | 0.91 | 1.00 | 0.67 | 0.18 | 0.69 | 0.03 | 0.46 | 0.06 | 0.86 | 0.30 | 0.51 |
| **scPRINT** | 0.20 | 0.46 | 0.69 | 0.35 | 0.38 | 0.15 | 0.91 | 1.00 | 0.68 | 0.24 | 0.60 | 0.03 | 0.41 | 0.15 | 0.86 | 0.29 | 0.50 |
| **CellPLM** | 0.22 | 0.50 | 0.72 | 0.39 | 0.40 | 0.11 | 0.91 | 1.00 | 0.70 | 0.22 | 0.51 | 0.03 | 0.44 | 0.11 | 0.87 | 0.25 | 0.50 |
| **Geneformer** | 0.20 | 0.48 | 0.69 | 0.37 | 0.38 | 0.06 | 0.91 | 1.00 | 0.67 | 0.25 | 0.65 | 0.02 | 0.42 | 0.06 | 0.86 | 0.31 | 0.50 |
| **CellFM** | 0.20 | 0.47 | 0.69 | 0.36 | 0.38 | -0.02 | 0.86 | 0.99 | 0.22 | 0.25 | 0.95 | 0.25 | 0.42 | -0.02 | 0.69 | 0.49 | 0.48 |
| **CELLama** | 0.02 | 0.15 | 0.22 | 0.11 | 0.19 | -0.02 | 0.78 | 0.98 | 0.47 | 0.28 | 0.69 | 0.07 | 0.14 | -0.02 | 0.75 | 0.35 | 0.37 |
| **GenePT-w** | 0.02 | 0.11 | 0.16 | 0.08 | 0.18 | -0.30 | 0.83 | 0.98 | 0.43 | 0.30 | 0.62 | 0.23 | 0.11 | -0.30 | 0.75 | 0.38 | 0.34 |
| **scBERT** | 0.01 | 0.03 | 0.05 | 0.03 | 0.16 | -0.55 | 0.51 | 0.94 | 0.24 | 0.11 | 0.56 | 0.23 | 0.06 | -0.55 | 0.56 | 0.30 | 0.22 |

**Figure S43: Representation performance in Tabula Sapiens v2 lymph node.** Scores are shown for 72,290 cells, 21 annotated cell types and 5 experimental batches. Metrics, family weighting and row ordering are as in Fig. S13.

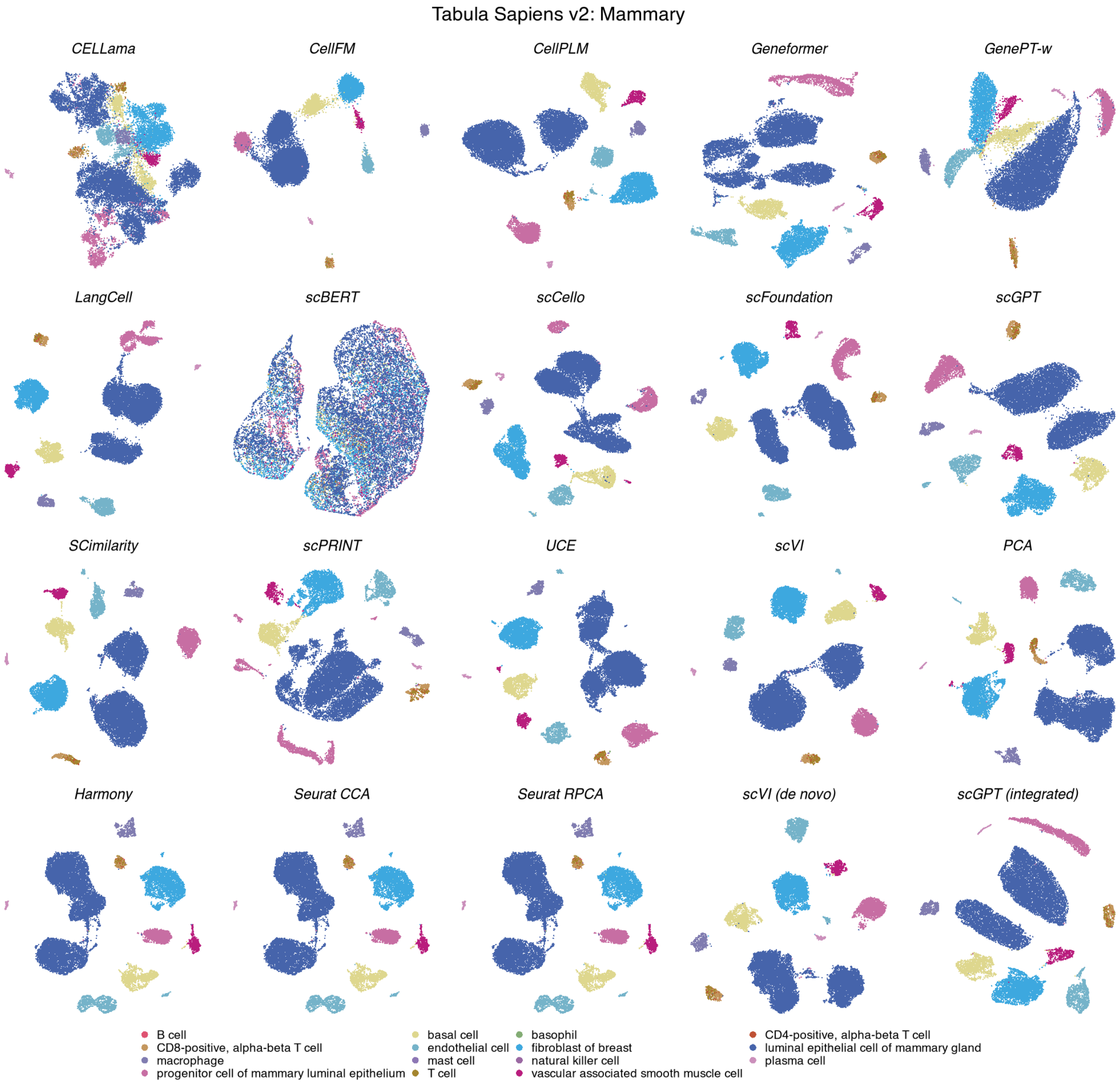

**Figure S44: Cell-type organization in Tabula Sapiens v2 mammary.** UMAPs show the same 18,539 cells across 20 methods, coloured by 15 annotated cell types. Display conventions and method categories are as in Fig. S11.

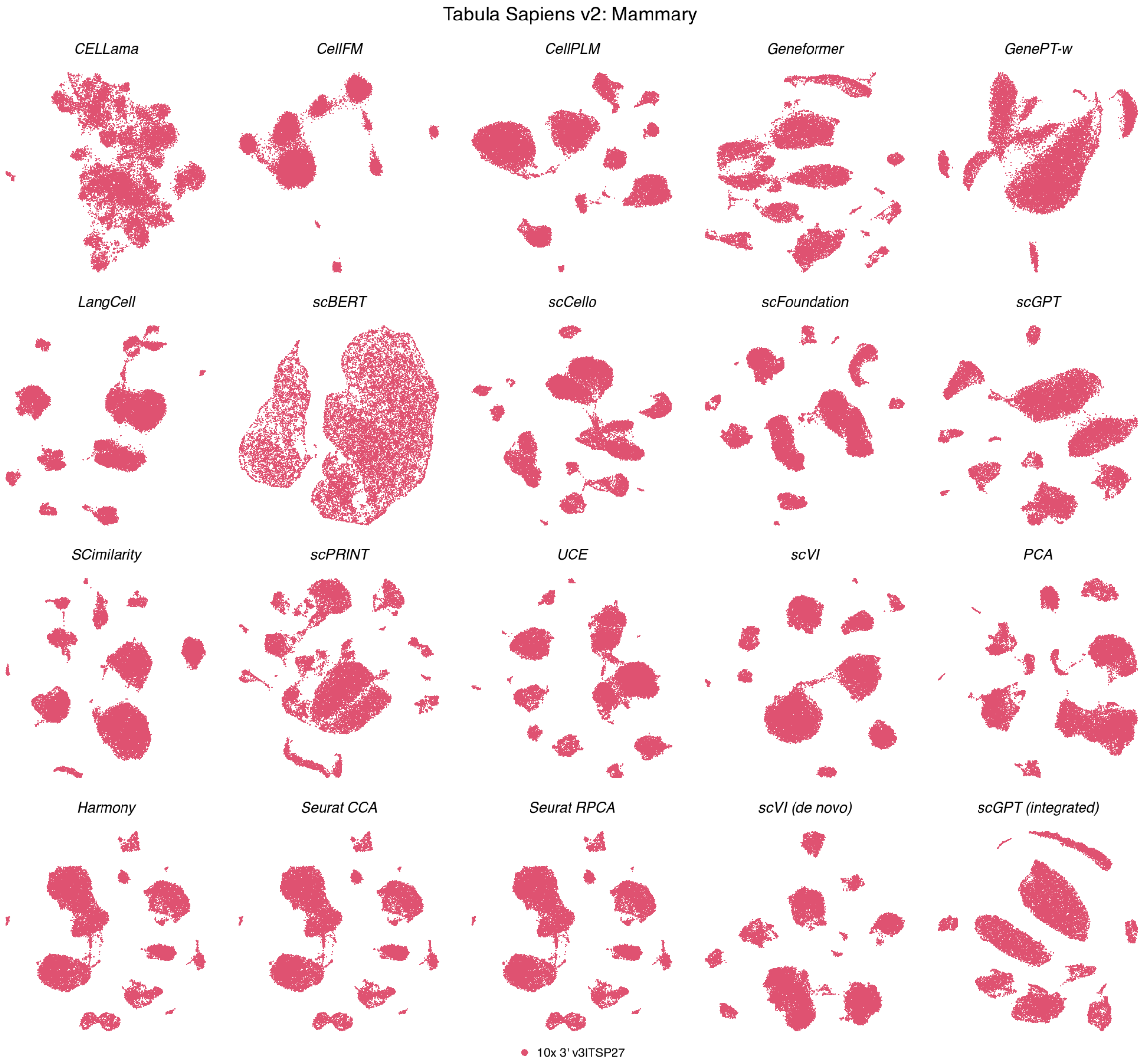

**Figure S45: Batch organization in Tabula Sapiens v2 mammary.** UMAPs show the same 18,539 cells across 20 methods, coloured by a single experimental batch. Coordinates match the corresponding cell-type view; display conventions are as in Fig. S12.

**Tabula Sapiens v2 — Mammary**

| | Biological conservation metrics | | | | | | | | | Batch mixing metrics | | | Aggregated scores | | | | |
|---|---|---|---|---|---|---|---|---|---|---|---|---|---|---|---|---|---|
| method | ARI | NMI | HOM | COM | FMI | ASW | Acc@kNN | cLISI | GC | kBET | BRAS | CiLISI | Cluster concordance | Continuous separation | Local structure | Batch mixing | Overall |
| SCimilarity | 0.46 | 0.80 | 0.98 | 0.68 | 0.63 | 0.49 | 0.99 | 1.00 | 0.93 | nan | nan | nan | 0.71 | 0.49 | 0.97 | | 0.80 |
| CellPLM | 0.46 | 0.79 | 0.97 | 0.67 | 0.63 | 0.29 | 0.99 | 1.00 | 0.94 | nan | nan | nan | 0.70 | 0.29 | 0.98 | | 0.77 |
| scVI | 0.41 | 0.78 | 0.98 | 0.64 | 0.60 | 0.30 | 0.99 | 1.00 | 0.95 | nan | nan | nan | 0.68 | 0.30 | 0.98 | | 0.76 |
| Harmony | 0.48 | 0.80 | 0.98 | 0.68 | 0.65 | 0.19 | 0.99 | 1.00 | 0.93 | nan | nan | nan | 0.72 | 0.19 | 0.97 | | 0.76 |
| Seurat CCA | 0.48 | 0.80 | 0.98 | 0.68 | 0.65 | 0.19 | 0.99 | 1.00 | 0.93 | nan | nan | nan | 0.72 | 0.19 | 0.97 | | 0.76 |
| Seurat RPCA | 0.48 | 0.80 | 0.98 | 0.68 | 0.65 | 0.19 | 0.99 | 1.00 | 0.93 | nan | nan | nan | 0.72 | 0.19 | 0.97 | | 0.76 |
| scFoundation | 0.40 | 0.77 | 0.98 | 0.64 | 0.58 | 0.30 | 0.99 | 1.00 | 0.95 | nan | nan | nan | 0.67 | 0.30 | 0.98 | | 0.76 |
| PCA | 0.47 | 0.79 | 0.97 | 0.67 | 0.64 | 0.21 | 0.99 | 1.00 | 0.92 | nan | nan | nan | 0.71 | 0.21 | 0.97 | | 0.75 |
| scGPT (integrated) | 0.39 | 0.75 | 0.97 | 0.62 | 0.57 | 0.38 | 0.99 | 1.00 | 0.90 | nan | nan | nan | 0.66 | 0.38 | 0.96 | | 0.75 |
| UCE | 0.44 | 0.78 | 0.98 | 0.65 | 0.62 | 0.26 | 0.99 | 1.00 | 0.89 | nan | nan | nan | 0.70 | 0.26 | 0.96 | | 0.75 |
| LangCell | 0.42 | 0.77 | 0.98 | 0.64 | 0.60 | 0.21 | 0.99 | 1.00 | 0.94 | nan | nan | nan | 0.68 | 0.21 | 0.98 | | 0.75 |
| scPRINT | 0.47 | 0.76 | 0.94 | 0.63 | 0.64 | 0.21 | 0.99 | 1.00 | 0.87 | nan | nan | nan | 0.69 | 0.21 | 0.96 | | 0.74 |
| scCello | 0.37 | 0.74 | 0.96 | 0.60 | 0.56 | 0.29 | 0.99 | 1.00 | 0.90 | nan | nan | nan | 0.65 | 0.29 | 0.96 | | 0.74 |
| scGPT | 0.37 | 0.75 | 0.98 | 0.61 | 0.56 | 0.19 | 0.99 | 1.00 | 0.93 | nan | nan | nan | 0.65 | 0.19 | 0.97 | | 0.73 |
| scVI (de novo) | 0.45 | 0.79 | 0.97 | 0.66 | 0.63 | 0.04 | 0.99 | 1.00 | 0.93 | nan | nan | nan | 0.70 | 0.04 | 0.97 | | 0.73 |
| Geneformer | 0.40 | 0.75 | 0.97 | 0.62 | 0.59 | 0.10 | 0.99 | 1.00 | 0.88 | nan | nan | nan | 0.66 | 0.10 | 0.96 | | 0.72 |
| CellFM | 0.37 | 0.75 | 0.95 | 0.61 | 0.56 | 0.01 | 0.98 | 1.00 | 0.46 | nan | nan | nan | 0.65 | 0.01 | 0.81 | | 0.63 |
| CELLama | 0.27 | 0.55 | 0.72 | 0.44 | 0.47 | -0.04 | 0.95 | 1.00 | 0.78 | nan | nan | nan | 0.49 | -0.04 | 0.91 | | 0.60 |
| GenePT-w | 0.05 | 0.22 | 0.30 | 0.18 | 0.24 | -0.38 | 0.94 | 1.00 | 0.71 | nan | nan | nan | 0.20 | -0.38 | 0.88 | | 0.42 |
| scBERT | 0.02 | 0.06 | 0.08 | 0.05 | 0.19 | -0.59 | 0.67 | 0.95 | 0.33 | nan | nan | nan | 0.08 | -0.59 | 0.65 | | 0.24 |

**Figure S46: Representation performance in Tabula Sapiens v2 mammary.** Scores are shown for 18,539 cells, 15 annotated cell types and a single experimental batch. Metrics, family weighting and row ordering are as in Fig. S13. Batch-mixing metrics were unavailable, and the remaining family weights were renormalized.

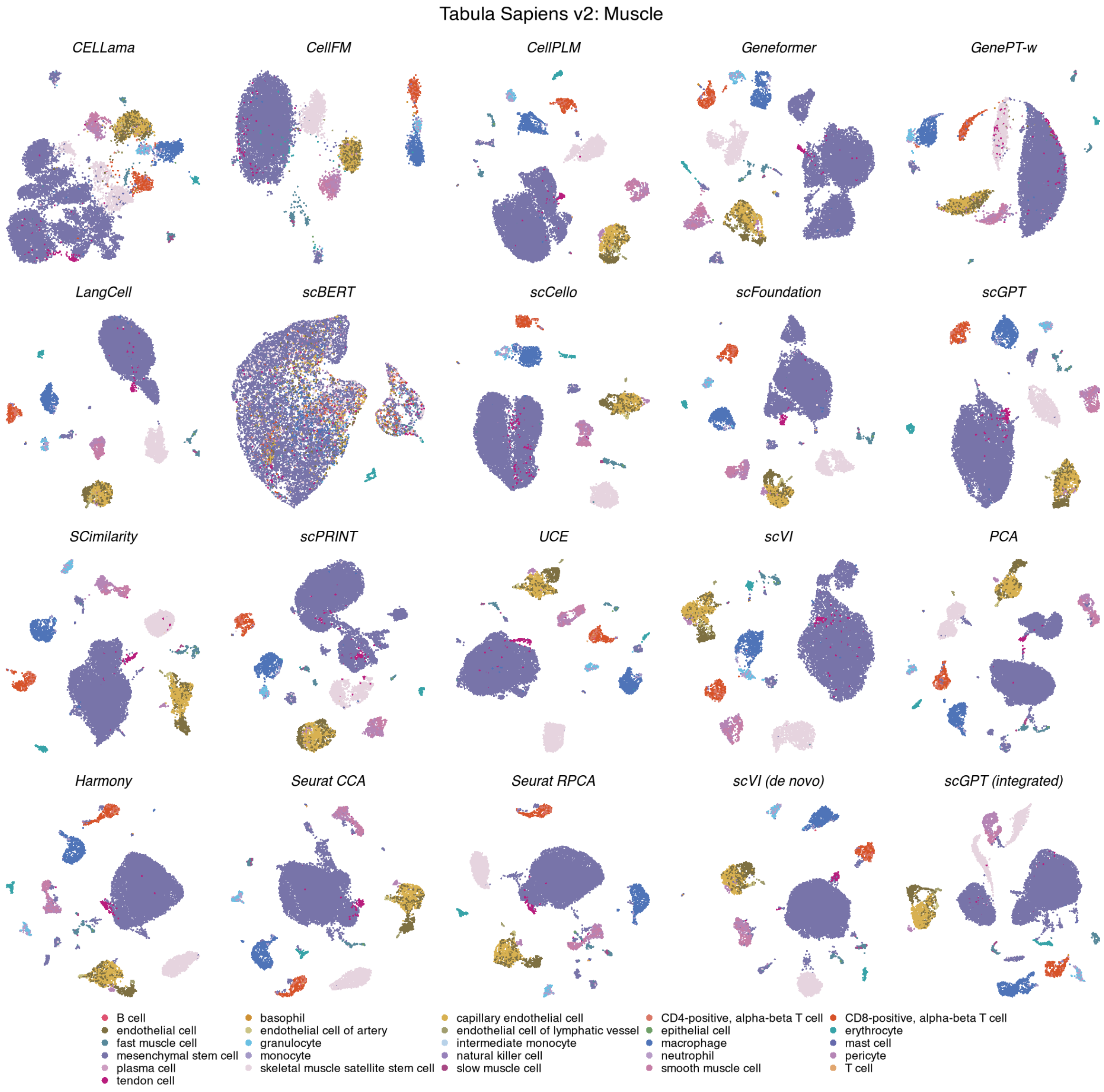

**Figure S47: Cell-type organization in Tabula Sapiens v2 muscle.** UMAPs show the same 13,035 cells across 20 methods, coloured by 26 annotated cell types. Display conventions and method categories are as in Fig. S11.

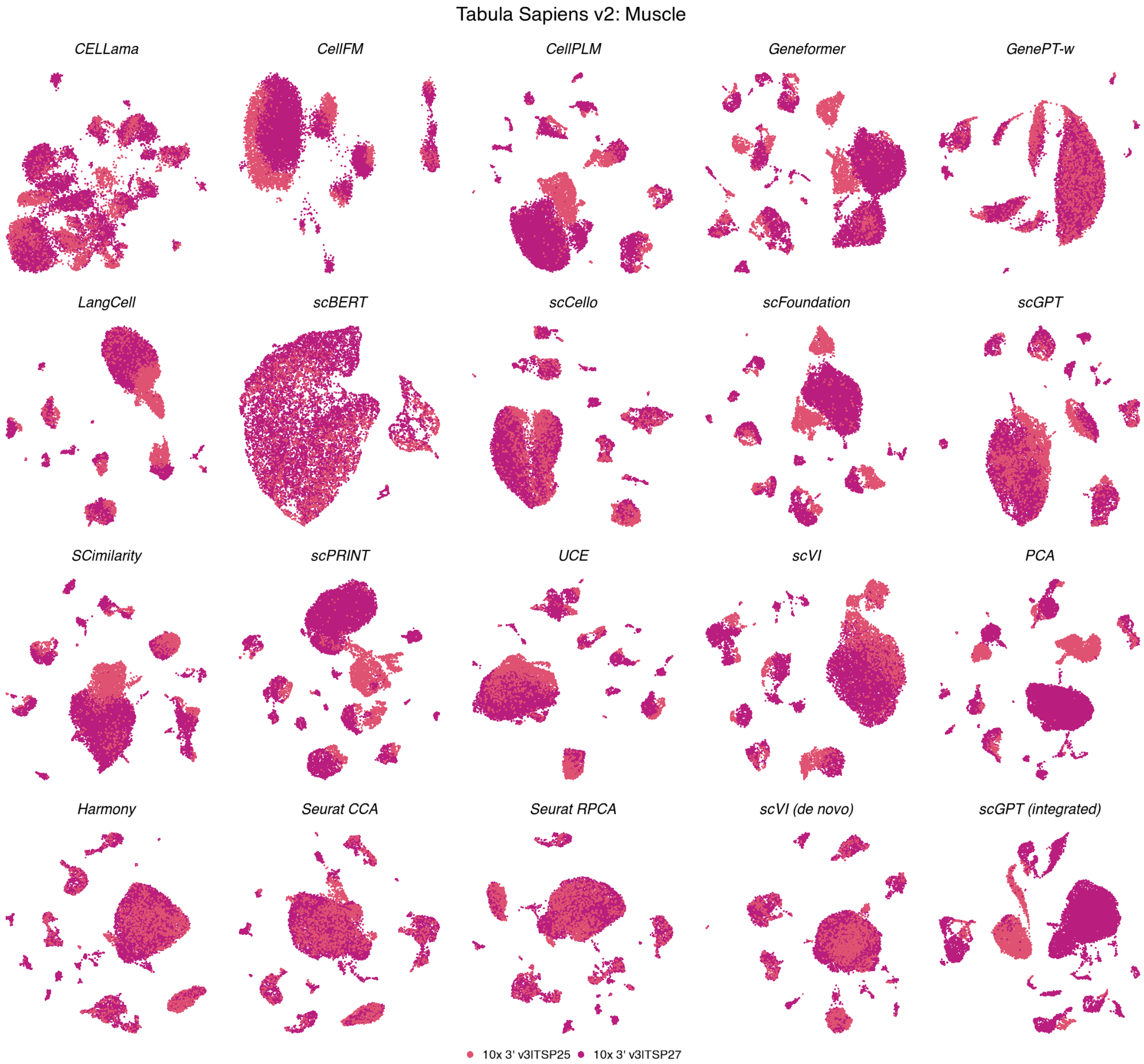

**Figure S48: Batch organization in Tabula Sapiens v2 muscle.** UMAPs show the same 13,035 cells across 20 methods, coloured by 2 experimental batches. Coordinates match the corresponding cell-type view; display conventions are as in Fig. S12.

**Tabula Sapiens v2 — Muscle**

| method | Biological conservation metrics | | | | | | | | | Batch mixing metrics | | | Aggregated scores | | | | |
|---|---|---|---|---|---|---|---|---|---|---|---|---|---|---|---|---|---|
| | ARI | NMI | HOM | COM | FMI | ASW | Acc@kNN | cLISI | GC | kBET | BRAS | CiLISI | Cluster concordance | Continuous separation | Local structure | Batch mixing | Overall |
| PCA | 0.40 | 0.67 | 0.88 | 0.55 | 0.62 | 0.28 | 0.97 | 1.00 | 0.88 | 0.66 | 0.62 | 0.04 | 0.62 | 0.28 | 0.95 | 0.44 | 0.65 |
| Harmony | 0.22 | 0.61 | 0.88 | 0.47 | 0.46 | 0.08 | 0.96 | 1.00 | 0.82 | 0.79 | 0.68 | 0.31 | 0.53 | 0.08 | 0.93 | 0.59 | 0.63 |
| Seurat RPCA | 0.17 | 0.59 | 0.89 | 0.45 | 0.40 | 0.07 | 0.96 | 1.00 | 0.84 | 0.78 | 0.68 | 0.39 | 0.50 | 0.07 | 0.93 | 0.62 | 0.63 |
| Seurat CCA | 0.17 | 0.59 | 0.88 | 0.45 | 0.41 | 0.04 | 0.96 | 1.00 | 0.82 | 0.78 | 0.68 | 0.44 | 0.50 | 0.04 | 0.93 | 0.63 | 0.63 |
| scCello | 0.12 | 0.55 | 0.87 | 0.41 | 0.33 | 0.19 | 0.95 | 1.00 | 0.84 | 0.75 | 0.73 | 0.20 | 0.46 | 0.19 | 0.93 | 0.56 | 0.61 |
| SCimilarity | 0.15 | 0.59 | 0.90 | 0.44 | 0.38 | 0.33 | 0.96 | 1.00 | 0.88 | 0.58 | 0.56 | 0.11 | 0.49 | 0.33 | 0.95 | 0.42 | 0.60 |
| scFoundation | 0.20 | 0.61 | 0.89 | 0.46 | 0.43 | 0.24 | 0.96 | 1.00 | 0.85 | 0.67 | 0.55 | 0.06 | 0.52 | 0.24 | 0.94 | 0.43 | 0.60 |
| scGPT (integrated) | 0.12 | 0.55 | 0.86 | 0.40 | 0.33 | 0.32 | 0.96 | 1.00 | 0.84 | 0.68 | 0.62 | 0.15 | 0.45 | 0.32 | 0.93 | 0.48 | 0.60 |
| UCE | 0.14 | 0.58 | 0.89 | 0.43 | 0.36 | 0.28 | 0.96 | 1.00 | 0.79 | 0.64 | 0.60 | 0.15 | 0.48 | 0.28 | 0.92 | 0.46 | 0.60 |
| scVI (de novo) | 0.12 | 0.56 | 0.88 | 0.41 | 0.33 | 0.04 | 0.96 | 1.00 | 0.81 | 0.75 | 0.75 | 0.25 | 0.46 | 0.04 | 0.92 | 0.58 | 0.60 |
| scGPT | 0.12 | 0.56 | 0.88 | 0.41 | 0.34 | 0.12 | 0.96 | 1.00 | 0.80 | 0.70 | 0.60 | 0.15 | 0.46 | 0.12 | 0.92 | 0.49 | 0.58 |
| LangCell | 0.12 | 0.56 | 0.88 | 0.42 | 0.34 | 0.11 | 0.96 | 1.00 | 0.81 | 0.57 | 0.72 | 0.11 | 0.47 | 0.11 | 0.92 | 0.47 | 0.58 |
| Geneformer | 0.18 | 0.57 | 0.85 | 0.43 | 0.41 | 0.13 | 0.96 | 1.00 | 0.78 | 0.61 | 0.60 | 0.08 | 0.49 | 0.13 | 0.91 | 0.43 | 0.58 |
| CellPLM | 0.12 | 0.56 | 0.88 | 0.41 | 0.33 | 0.18 | 0.96 | 1.00 | 0.82 | 0.61 | 0.50 | 0.07 | 0.46 | 0.18 | 0.93 | 0.39 | 0.57 |
| scVI | 0.13 | 0.56 | 0.87 | 0.41 | 0.35 | 0.11 | 0.96 | 1.00 | 0.80 | 0.61 | 0.53 | 0.12 | 0.46 | 0.11 | 0.92 | 0.42 | 0.57 |
| scPRINT | 0.15 | 0.56 | 0.85 | 0.42 | 0.38 | 0.14 | 0.95 | 1.00 | 0.81 | 0.55 | 0.56 | 0.05 | 0.47 | 0.14 | 0.92 | 0.39 | 0.56 |
| CELLama | 0.11 | 0.50 | 0.79 | 0.36 | 0.32 | -0.01 | 0.93 | 1.00 | 0.65 | 0.64 | 0.73 | 0.19 | 0.42 | -0.01 | 0.86 | 0.52 | 0.54 |
| CellFM | 0.09 | 0.48 | 0.77 | 0.35 | 0.30 | 0.01 | 0.90 | 1.00 | 0.39 | 0.59 | 0.94 | 0.29 | 0.40 | 0.01 | 0.76 | 0.61 | 0.53 |
| GenePT-w | 0.06 | 0.30 | 0.49 | 0.22 | 0.25 | -0.41 | 0.91 | 1.00 | 0.48 | 0.23 | 0.53 | 0.27 | 0.26 | -0.41 | 0.80 | 0.34 | 0.39 |
| scBERT | 0.02 | 0.11 | 0.18 | 0.08 | 0.19 | -0.51 | 0.72 | 0.99 | 0.29 | 0.22 | 0.53 | 0.31 | 0.12 | -0.51 | 0.67 | 0.35 | 0.29 |

**Figure S49: Representation performance in Tabula Sapiens v2 muscle.** Scores are shown for 13,035 cells, 26 annotated cell types and 2 experimental batches. Metrics, family weighting and row ordering are as in Fig. S13.

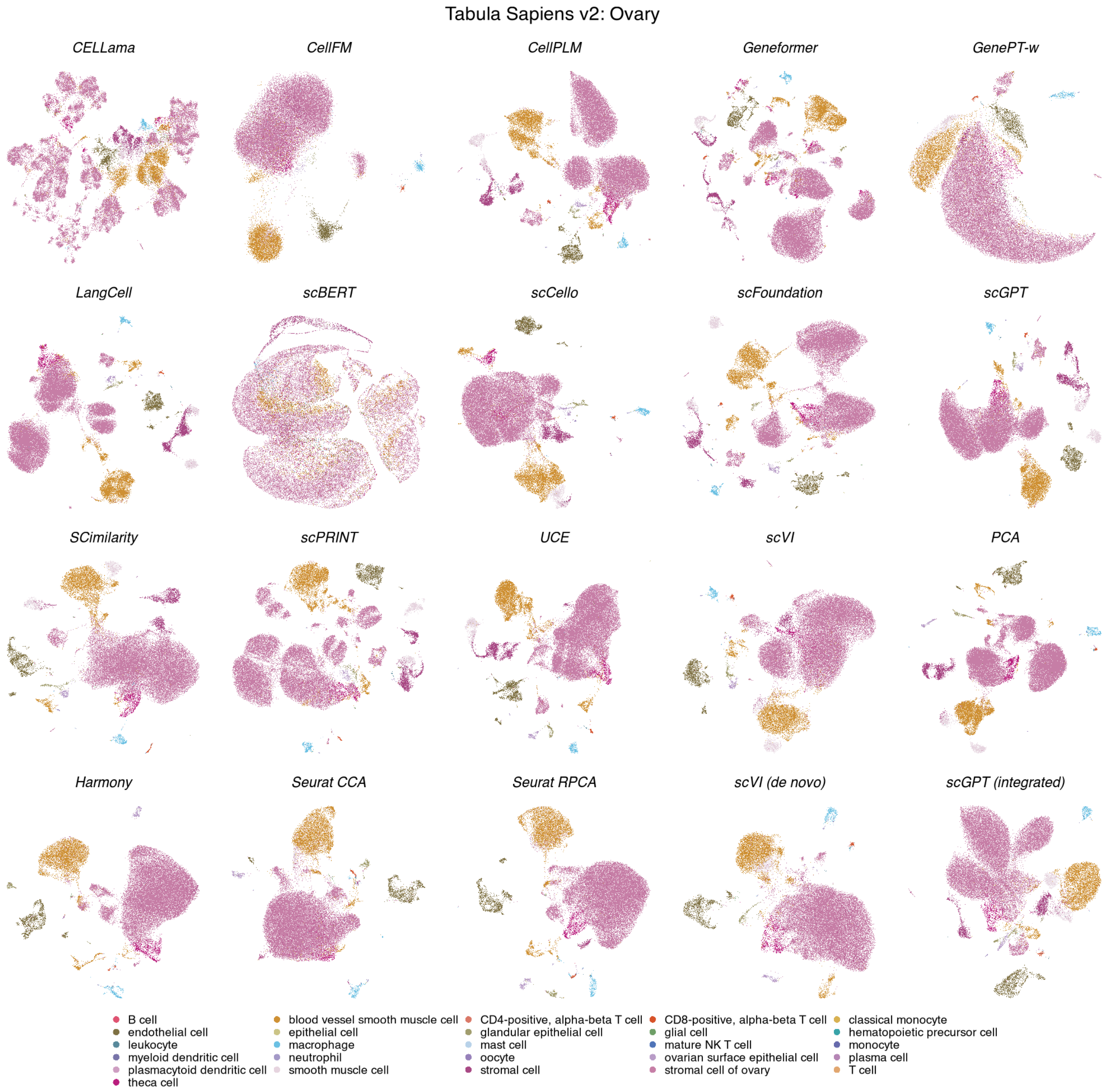

**Figure S50: Cell-type organization in Tabula Sapiens v2 ovary.** UMAPs show the same 48,942 cells across 20 methods, coloured by 26 annotated cell types. Display conventions and method categories are as in Fig. S11.

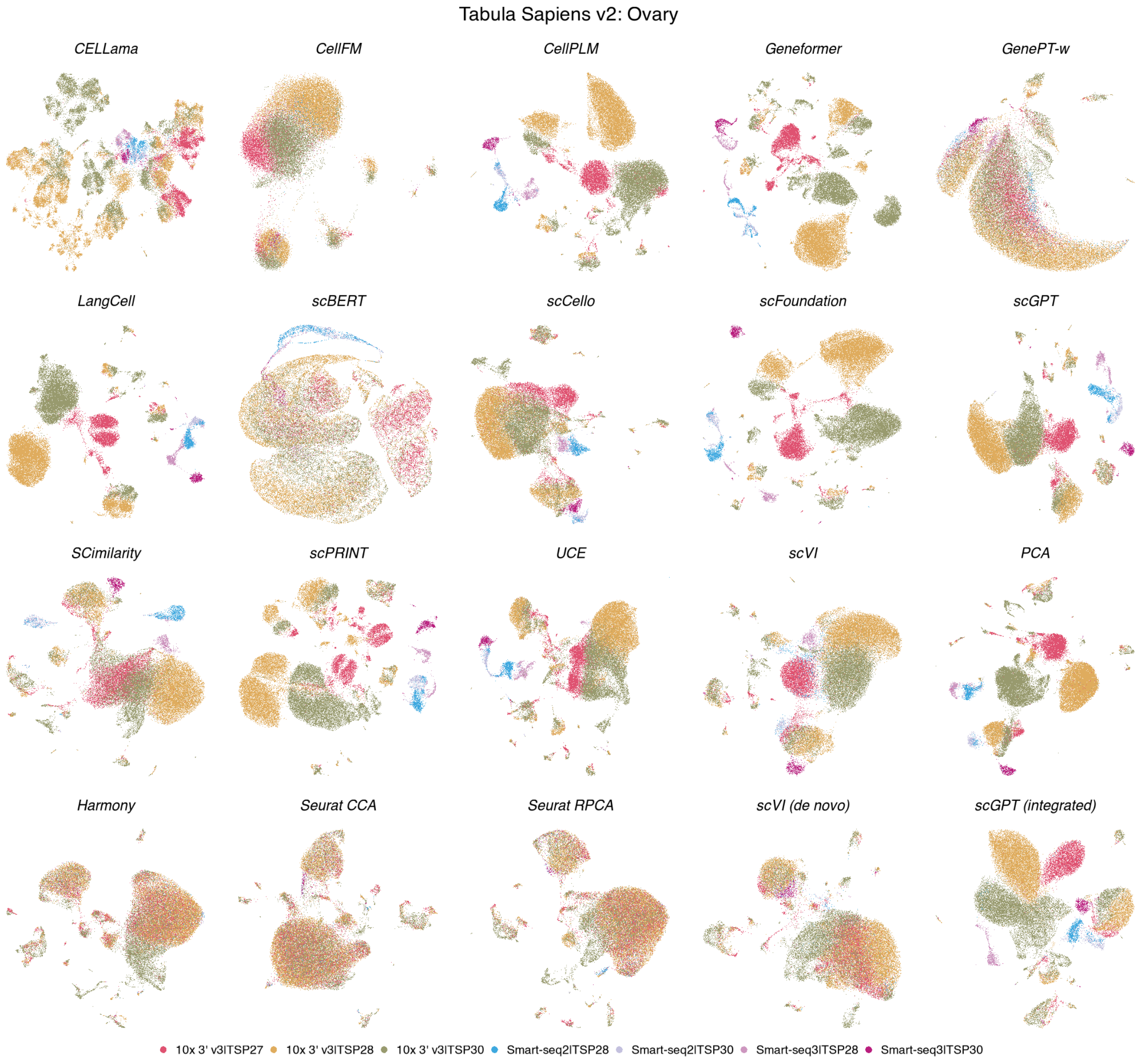

**Figure S51: Batch organization in Tabula Sapiens v2 ovary.** UMAPs show the same 48,942 cells across 20 methods, coloured by 7 experimental batches. Coordinates match the corresponding cell-type view; display conventions are as in Fig. S12.

**Tabula Sapiens v2 — Ovary**

| method | Biological conservation metrics | | | | | | | | | Batch mixing metrics | | | Aggregated scores | | | | |
|---|---|---|---|---|---|---|---|---|---|---|---|---|---|---|---|---|---|
| | ARI | NMI | HOM | COM | FMI | ASW | Acc@kNN | cLISI | GC | kBET | BRAS | CiLISI | Cluster concordance | Continuous separation | Local structure | Batch mixing | Overall |
| **scGPT** | 0.18 | 0.53 | 0.87 | 0.38 | 0.44 | 0.24 | 0.98 | 1.00 | 0.88 | 0.45 | 0.63 | 0.07 | 0.48 | 0.24 | 0.95 | 0.38 | 0.58 |
| **UCE** | 0.12 | 0.47 | 0.84 | 0.33 | 0.36 | 0.26 | 0.98 | 1.00 | 0.84 | 0.52 | 0.69 | 0.10 | 0.42 | 0.26 | 0.94 | 0.44 | 0.58 |
| **PCA** | 0.16 | 0.51 | 0.86 | 0.36 | 0.42 | 0.17 | 0.98 | 1.00 | 0.88 | 0.44 | 0.66 | 0.05 | 0.46 | 0.17 | 0.95 | 0.38 | 0.57 |
| **SCimilarity** | 0.16 | 0.50 | 0.85 | 0.35 | 0.41 | 0.18 | 0.98 | 1.00 | 0.89 | 0.40 | 0.59 | 0.13 | 0.45 | 0.18 | 0.96 | 0.37 | 0.57 |
| **CellPLM** | 0.18 | 0.51 | 0.86 | 0.36 | 0.44 | 0.19 | 0.98 | 1.00 | 0.89 | 0.36 | 0.57 | 0.03 | 0.47 | 0.19 | 0.96 | 0.32 | 0.56 |
| **scCello** | 0.12 | 0.46 | 0.83 | 0.32 | 0.36 | 0.09 | 0.97 | 1.00 | 0.84 | 0.50 | 0.74 | 0.12 | 0.42 | 0.09 | 0.94 | 0.46 | 0.56 |
| **LangCell** | 0.21 | 0.52 | 0.86 | 0.37 | 0.47 | 0.12 | 0.98 | 1.00 | 0.83 | 0.29 | 0.69 | 0.03 | 0.48 | 0.12 | 0.94 | 0.34 | 0.56 |
| **scFoundation** | 0.16 | 0.50 | 0.86 | 0.36 | 0.42 | 0.14 | 0.98 | 1.00 | 0.83 | 0.40 | 0.57 | 0.02 | 0.46 | 0.14 | 0.94 | 0.33 | 0.55 |
| **Seurat RPCA** | 0.09 | 0.38 | 0.69 | 0.26 | 0.32 | 0.01 | 0.93 | 1.00 | 0.81 | 0.52 | 0.71 | 0.39 | 0.35 | 0.01 | 0.91 | 0.54 | 0.55 |
| **scVI (de novo)** | 0.08 | 0.37 | 0.67 | 0.25 | 0.31 | 0.07 | 0.94 | 1.00 | 0.81 | 0.44 | 0.74 | 0.30 | 0.34 | 0.07 | 0.92 | 0.49 | 0.54 |
| **scPRINT** | 0.10 | 0.43 | 0.80 | 0.30 | 0.34 | 0.14 | 0.98 | 1.00 | 0.85 | 0.44 | 0.57 | 0.05 | 0.39 | 0.14 | 0.94 | 0.35 | 0.54 |
| **scVI** | 0.11 | 0.42 | 0.75 | 0.29 | 0.34 | 0.15 | 0.96 | 1.00 | 0.86 | 0.30 | 0.62 | 0.08 | 0.38 | 0.15 | 0.94 | 0.34 | 0.53 |
| **scGPT (integrated)** | 0.10 | 0.39 | 0.70 | 0.27 | 0.33 | -0.02 | 0.96 | 1.00 | 0.83 | 0.45 | 0.64 | 0.21 | 0.36 | -0.02 | 0.93 | 0.43 | 0.52 |
| **Geneformer** | 0.22 | 0.48 | 0.78 | 0.34 | 0.48 | 0.07 | 0.98 | 1.00 | 0.73 | 0.12 | 0.64 | 0.02 | 0.46 | 0.07 | 0.90 | 0.26 | 0.52 |
| **Harmony** | 0.08 | 0.38 | 0.69 | 0.26 | 0.30 | -0.02 | 0.92 | 1.00 | 0.77 | 0.50 | 0.58 | 0.27 | 0.34 | -0.02 | 0.90 | 0.45 | 0.51 |
| **Seurat CCA** | 0.08 | 0.34 | 0.62 | 0.24 | 0.31 | -0.03 | 0.89 | 0.99 | 0.76 | 0.32 | 0.69 | 0.38 | 0.32 | -0.03 | 0.88 | 0.46 | 0.50 |
| **CellFM** | 0.09 | 0.32 | 0.56 | 0.22 | 0.34 | -0.05 | 0.87 | 1.00 | 0.34 | 0.38 | 0.94 | 0.27 | 0.30 | -0.05 | 0.74 | 0.53 | 0.47 |
| **CELLama** | 0.07 | 0.33 | 0.61 | 0.22 | 0.27 | -0.08 | 0.94 | 1.00 | 0.55 | 0.19 | 0.72 | 0.09 | 0.30 | -0.08 | 0.83 | 0.33 | 0.44 |
| **GenePT-w** | 0.04 | 0.21 | 0.40 | 0.14 | 0.22 | -0.68 | 0.93 | 1.00 | 0.47 | 0.16 | 0.67 | 0.15 | 0.20 | -0.68 | 0.80 | 0.33 | 0.34 |
| **scBERT** | 0.02 | 0.15 | 0.29 | 0.10 | 0.19 | -0.59 | 0.81 | 0.99 | 0.33 | 0.10 | 0.41 | 0.21 | 0.15 | -0.59 | 0.71 | 0.24 | 0.28 |

**Figure S52: Representation performance in Tabula Sapiens v2 ovary.** Scores are shown for 48,942 cells, 26 annotated cell types and 7 experimental batches. Metrics, family weighting and row ordering are as in Fig. S13.

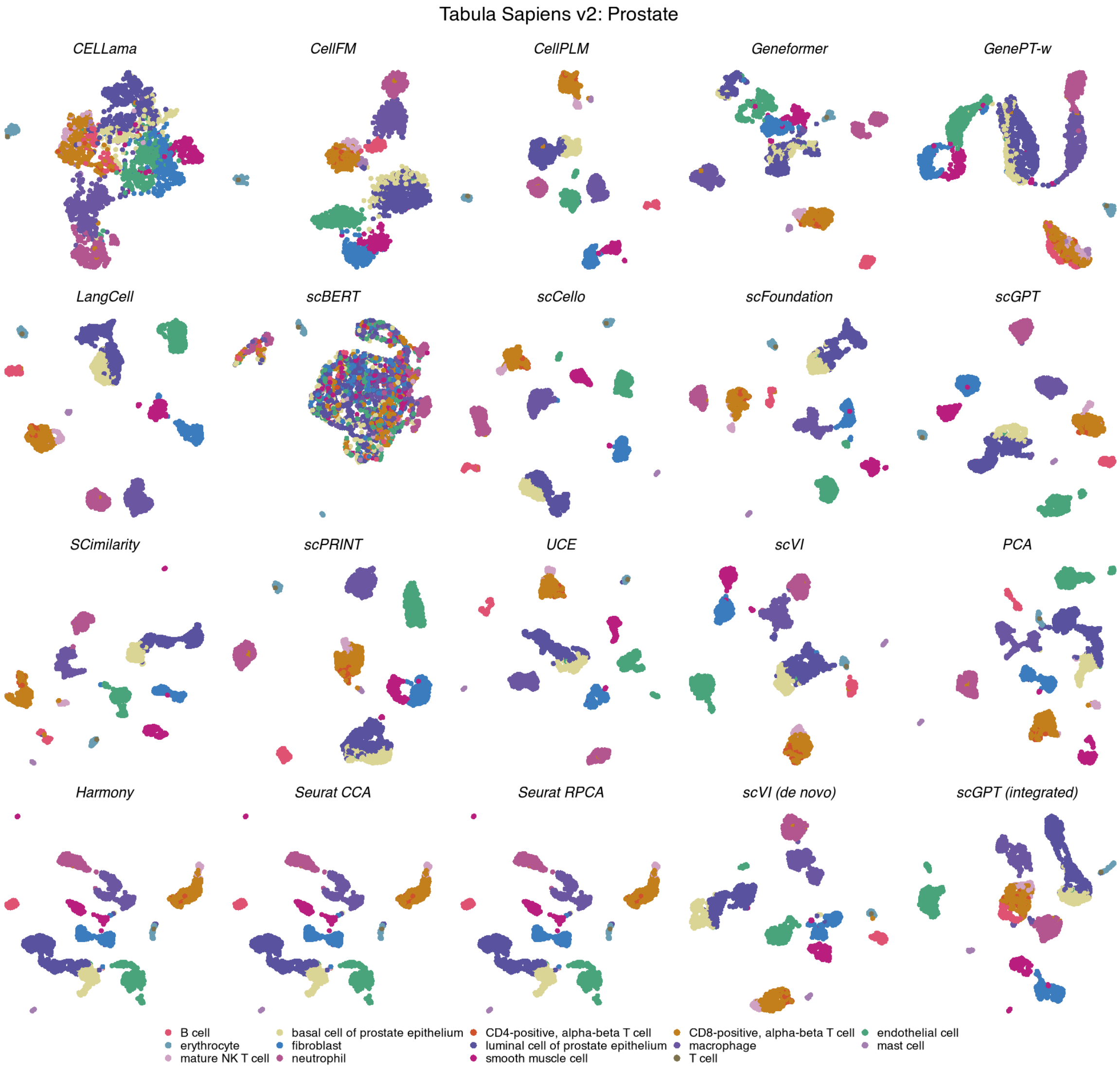

**Figure S53: Cell-type organization in Tabula Sapiens v2 prostate.** UMAPs show the same 2,044 cells across 20 methods, coloured by 14 annotated cell types. Display conventions and method categories are as in Fig. S11.

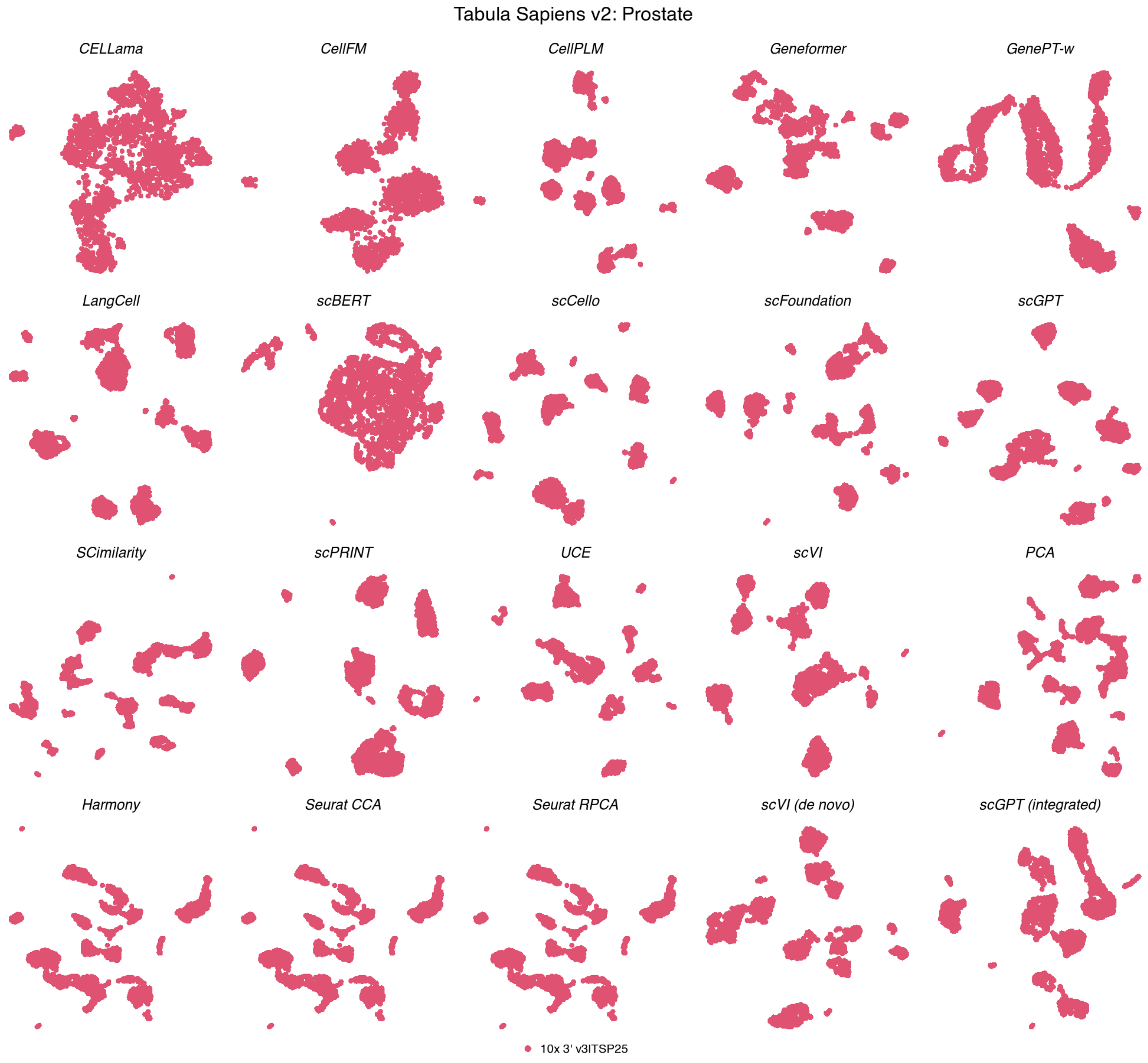

**Figure S54: Batch organization in Tabula Sapiens v2 prostate.** UMAPs show the same 2,044 cells across 20 methods, coloured by a single experimental batch. Coordinates match the corresponding cell-type view; display conventions are as in Fig. S12.

**Tabula Sapiens v2 — Prostate**

| method | Biological conservation metrics | | | | | | | | | Batch mixing metrics | | | Aggregated scores | | | | |
|---|---|---|---|---|---|---|---|---|---|---|---|---|---|---|---|---|---|
| | ARI | NMI | HOM | COM | FMI | ASW | Acc@kNN | cLISI | GC | kBET | BRAS | CiLISI | Cluster concordance | Continuous separation | Local structure | Batch mixing | Overall |
| SCimilarity | 0.77 | 0.89 | 0.90 | 0.89 | 0.79 | 0.45 | 0.97 | 1.00 | 0.87 | nan | nan | nan | 0.85 | 0.45 | 0.95 | | 0.84 |
| scFoundation | 0.79 | 0.91 | 0.91 | 0.91 | 0.81 | 0.28 | 0.97 | 1.00 | 0.91 | nan | nan | nan | 0.86 | 0.28 | 0.96 | | 0.83 |
| scCello | 0.79 | 0.89 | 0.91 | 0.87 | 0.81 | 0.30 | 0.96 | 1.00 | 0.92 | nan | nan | nan | 0.86 | 0.30 | 0.96 | | 0.83 |
| UCE | 0.80 | 0.90 | 0.92 | 0.88 | 0.82 | 0.27 | 0.97 | 1.00 | 0.91 | nan | nan | nan | 0.87 | 0.27 | 0.96 | | 0.83 |
| CellPLM | 0.78 | 0.89 | 0.90 | 0.87 | 0.81 | 0.31 | 0.96 | 1.00 | 0.91 | nan | nan | nan | 0.85 | 0.31 | 0.96 | | 0.82 |
| scGPT | 0.80 | 0.89 | 0.92 | 0.87 | 0.82 | 0.25 | 0.96 | 1.00 | 0.92 | nan | nan | nan | 0.86 | 0.25 | 0.96 | | 0.82 |
| scGPT (integrated) | 0.73 | 0.88 | 0.88 | 0.87 | 0.76 | 0.34 | 0.96 | 1.00 | 0.94 | nan | nan | nan | 0.82 | 0.34 | 0.97 | | 0.82 |
| Harmony | 0.75 | 0.86 | 0.88 | 0.85 | 0.78 | 0.29 | 0.96 | 1.00 | 0.96 | nan | nan | nan | 0.82 | 0.29 | 0.97 | | 0.82 |
| Seurat CCA | 0.75 | 0.86 | 0.88 | 0.85 | 0.78 | 0.29 | 0.96 | 1.00 | 0.96 | nan | nan | nan | 0.82 | 0.29 | 0.97 | | 0.82 |
| Seurat RPCA | 0.75 | 0.86 | 0.88 | 0.85 | 0.78 | 0.29 | 0.96 | 1.00 | 0.96 | nan | nan | nan | 0.82 | 0.29 | 0.97 | | 0.82 |
| LangCell | 0.77 | 0.90 | 0.89 | 0.90 | 0.80 | 0.16 | 0.95 | 1.00 | 0.94 | nan | nan | nan | 0.85 | 0.16 | 0.96 | | 0.81 |
| scPRINT | 0.74 | 0.87 | 0.90 | 0.84 | 0.77 | 0.23 | 0.96 | 1.00 | 0.94 | nan | nan | nan | 0.83 | 0.23 | 0.96 | | 0.81 |
| PCA | 0.74 | 0.87 | 0.90 | 0.84 | 0.77 | 0.22 | 0.96 | 1.00 | 0.94 | nan | nan | nan | 0.83 | 0.22 | 0.97 | | 0.81 |
| scVI | 0.74 | 0.87 | 0.88 | 0.85 | 0.77 | 0.23 | 0.95 | 1.00 | 0.94 | nan | nan | nan | 0.82 | 0.23 | 0.96 | | 0.80 |
| scVI (de novo) | 0.77 | 0.87 | 0.90 | 0.85 | 0.79 | 0.13 | 0.96 | 1.00 | 0.96 | nan | nan | nan | 0.84 | 0.13 | 0.97 | | 0.80 |
| Geneformer | 0.66 | 0.81 | 0.86 | 0.77 | 0.70 | 0.11 | 0.95 | 1.00 | 0.87 | nan | nan | nan | 0.76 | 0.11 | 0.94 | | 0.75 |
| CellFM | 0.70 | 0.83 | 0.85 | 0.81 | 0.74 | 0.01 | 0.89 | 0.97 | 0.59 | nan | nan | nan | 0.79 | 0.01 | 0.82 | | 0.70 |
| CELLama | 0.51 | 0.66 | 0.68 | 0.63 | 0.56 | 0.03 | 0.85 | 0.97 | 0.77 | nan | nan | nan | 0.61 | 0.03 | 0.87 | | 0.64 |
| GenePT-w | 0.15 | 0.32 | 0.33 | 0.30 | 0.23 | -0.10 | 0.75 | 0.92 | 0.61 | nan | nan | nan | 0.27 | -0.10 | 0.76 | | 0.43 |
| scBERT | 0.08 | 0.19 | 0.20 | 0.18 | 0.18 | -0.13 | 0.44 | 0.80 | 0.39 | nan | nan | nan | 0.16 | -0.13 | 0.54 | | 0.29 |

**Figure S55: Representation performance in Tabula Sapiens v2 prostate.** Scores are shown for 2,044 cells, 14 annotated cell types and a single experimental batch. Metrics, family weighting and row ordering are as in Fig. S13. Batch-mixing metrics were unavailable, and the remaining family weights were renormalized.

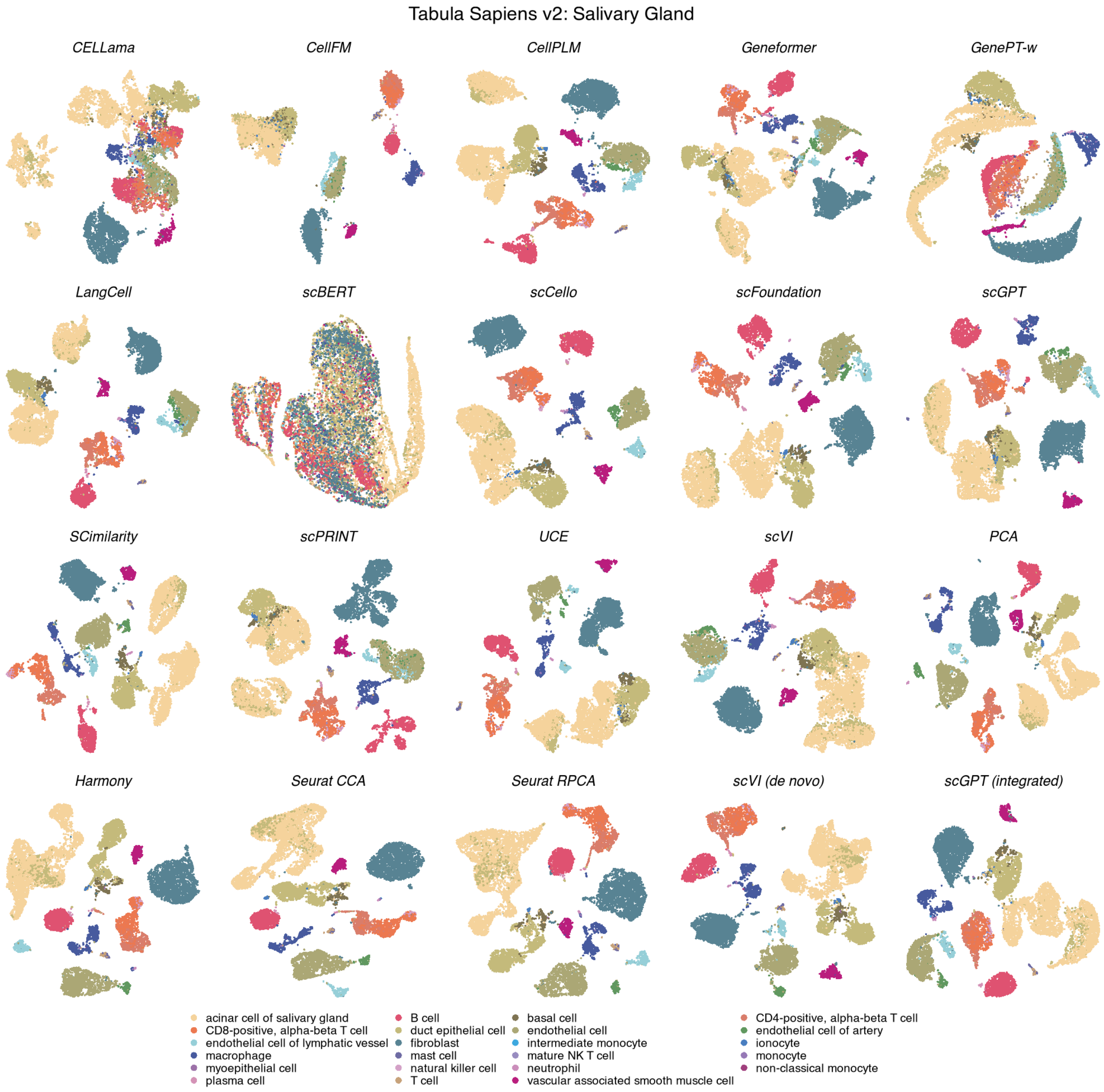

**Figure S56: Cell-type organization in Tabula Sapiens v2 salivary gland.** UMAPs show the same 11,561 cells across 20 methods, coloured by 23 annotated cell types. Display conventions and method categories are as in Fig. S11.

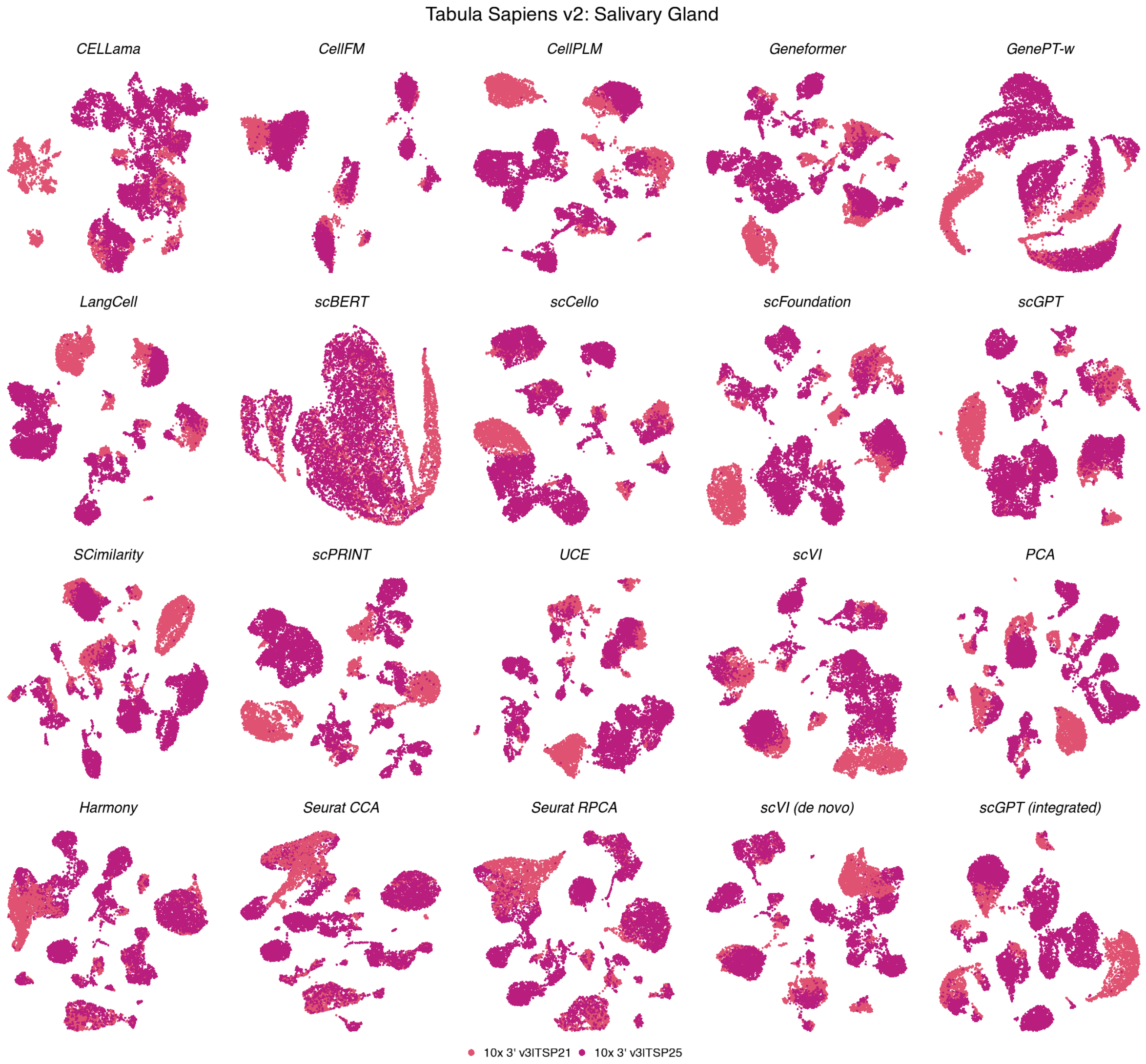

**Figure S57: Batch organization in Tabula Sapiens v2 salivary gland.** UMAPs show the same 11,561 cells across 20 methods, coloured by 2 experimental batches. Coordinates match the corresponding cell-type view; display conventions are as in Fig. S12.

**Tabula Sapiens v2 — Salivary Gland**

| | Biological conservation metrics | | | | | | | | | Batch mixing metrics | | | Aggregated scores | | | | |
|---|---|---|---|---|---|---|---|---|---|---|---|---|---|---|---|---|---|
| method | ARI | NMI | HOM | COM | FMI | ASW | Acc@kNN | cLISI | GC | kBET | BRAS | CiLISI | Cluster concordance | Continuous separation | Local structure | Batch mixing | Overall |
| Seurat CCA | 0.52 | 0.77 | 0.89 | 0.67 | 0.62 | 0.36 | 0.95 | 1.00 | 0.88 | 0.53 | 0.65 | 0.33 | 0.69 | 0.36 | 0.94 | 0.50 | 0.69 |
| Harmony | 0.45 | 0.76 | 0.90 | 0.66 | 0.56 | 0.37 | 0.95 | 1.00 | 0.90 | 0.55 | 0.62 | 0.24 | 0.66 | 0.37 | 0.95 | 0.47 | 0.68 |
| Seurat RPCA | 0.46 | 0.76 | 0.89 | 0.66 | 0.57 | 0.36 | 0.95 | 1.00 | 0.90 | 0.45 | 0.59 | 0.25 | 0.67 | 0.36 | 0.95 | 0.43 | 0.67 |
| SCimilarity | 0.58 | 0.79 | 0.91 | 0.70 | 0.66 | 0.37 | 0.97 | 1.00 | 0.89 | 0.23 | 0.56 | 0.12 | 0.73 | 0.37 | 0.95 | 0.30 | 0.66 |
| scGPT (integrated) | 0.51 | 0.78 | 0.89 | 0.69 | 0.60 | 0.34 | 0.96 | 1.00 | 0.91 | 0.25 | 0.60 | 0.12 | 0.69 | 0.34 | 0.96 | 0.32 | 0.65 |
| scVI (de novo) | 0.56 | 0.77 | 0.88 | 0.68 | 0.64 | 0.11 | 0.96 | 1.00 | 0.91 | 0.26 | 0.72 | 0.20 | 0.71 | 0.11 | 0.96 | 0.39 | 0.65 |
| scCello | 0.46 | 0.76 | 0.89 | 0.66 | 0.56 | 0.26 | 0.95 | 1.00 | 0.90 | 0.30 | 0.70 | 0.16 | 0.66 | 0.26 | 0.95 | 0.39 | 0.65 |
| UCE | 0.54 | 0.79 | 0.90 | 0.70 | 0.63 | 0.26 | 0.96 | 1.00 | 0.85 | 0.24 | 0.63 | 0.11 | 0.71 | 0.26 | 0.93 | 0.32 | 0.64 |
| PCA | 0.52 | 0.78 | 0.90 | 0.69 | 0.61 | 0.28 | 0.96 | 1.00 | 0.87 | 0.24 | 0.53 | 0.07 | 0.70 | 0.28 | 0.94 | 0.28 | 0.63 |
| scVI | 0.46 | 0.74 | 0.86 | 0.65 | 0.56 | 0.20 | 0.95 | 1.00 | 0.89 | 0.28 | 0.65 | 0.13 | 0.65 | 0.20 | 0.95 | 0.35 | 0.63 |
| CellPLM | 0.50 | 0.76 | 0.88 | 0.67 | 0.59 | 0.26 | 0.96 | 1.00 | 0.91 | 0.15 | 0.56 | 0.08 | 0.68 | 0.26 | 0.96 | 0.26 | 0.62 |
| scFoundation | 0.47 | 0.76 | 0.88 | 0.67 | 0.57 | 0.21 | 0.97 | 1.00 | 0.89 | 0.18 | 0.61 | 0.09 | 0.67 | 0.21 | 0.95 | 0.30 | 0.62 |
| LangCell | 0.47 | 0.74 | 0.87 | 0.65 | 0.57 | 0.14 | 0.96 | 1.00 | 0.90 | 0.09 | 0.74 | 0.09 | 0.66 | 0.14 | 0.95 | 0.31 | 0.61 |
| scGPT | 0.42 | 0.73 | 0.86 | 0.63 | 0.53 | 0.17 | 0.95 | 1.00 | 0.92 | 0.16 | 0.60 | 0.11 | 0.63 | 0.17 | 0.96 | 0.29 | 0.61 |
| CellFM | 0.52 | 0.75 | 0.81 | 0.70 | 0.60 | -0.01 | 0.86 | 0.99 | 0.44 | 0.21 | 0.94 | 0.33 | 0.68 | -0.01 | 0.76 | 0.50 | 0.59 |
| scPRINT | 0.37 | 0.69 | 0.83 | 0.60 | 0.48 | 0.12 | 0.94 | 1.00 | 0.89 | 0.20 | 0.61 | 0.06 | 0.59 | 0.12 | 0.94 | 0.29 | 0.59 |
| Geneformer | 0.43 | 0.68 | 0.79 | 0.59 | 0.53 | 0.05 | 0.95 | 1.00 | 0.81 | 0.15 | 0.71 | 0.10 | 0.60 | 0.05 | 0.92 | 0.32 | 0.58 |
| CELLama | 0.28 | 0.58 | 0.70 | 0.50 | 0.39 | -0.01 | 0.91 | 1.00 | 0.67 | 0.26 | 0.74 | 0.14 | 0.49 | -0.01 | 0.86 | 0.38 | 0.53 |
| GenePT-w | 0.09 | 0.30 | 0.36 | 0.25 | 0.19 | -0.57 | 0.84 | 0.98 | 0.50 | 0.20 | 0.49 | 0.19 | 0.24 | -0.57 | 0.77 | 0.29 | 0.34 |
| scBERT | 0.06 | 0.16 | 0.19 | 0.14 | 0.16 | -0.46 | 0.56 | 0.92 | 0.32 | 0.09 | 0.50 | 0.24 | 0.14 | -0.46 | 0.60 | 0.27 | 0.26 |

**Figure S58: Representation performance in Tabula Sapiens v2 salivary gland.** Scores are shown for 11,561 cells, 23 annotated cell types and 2 experimental batches. Metrics, family weighting and row ordering are as in Fig. S13.

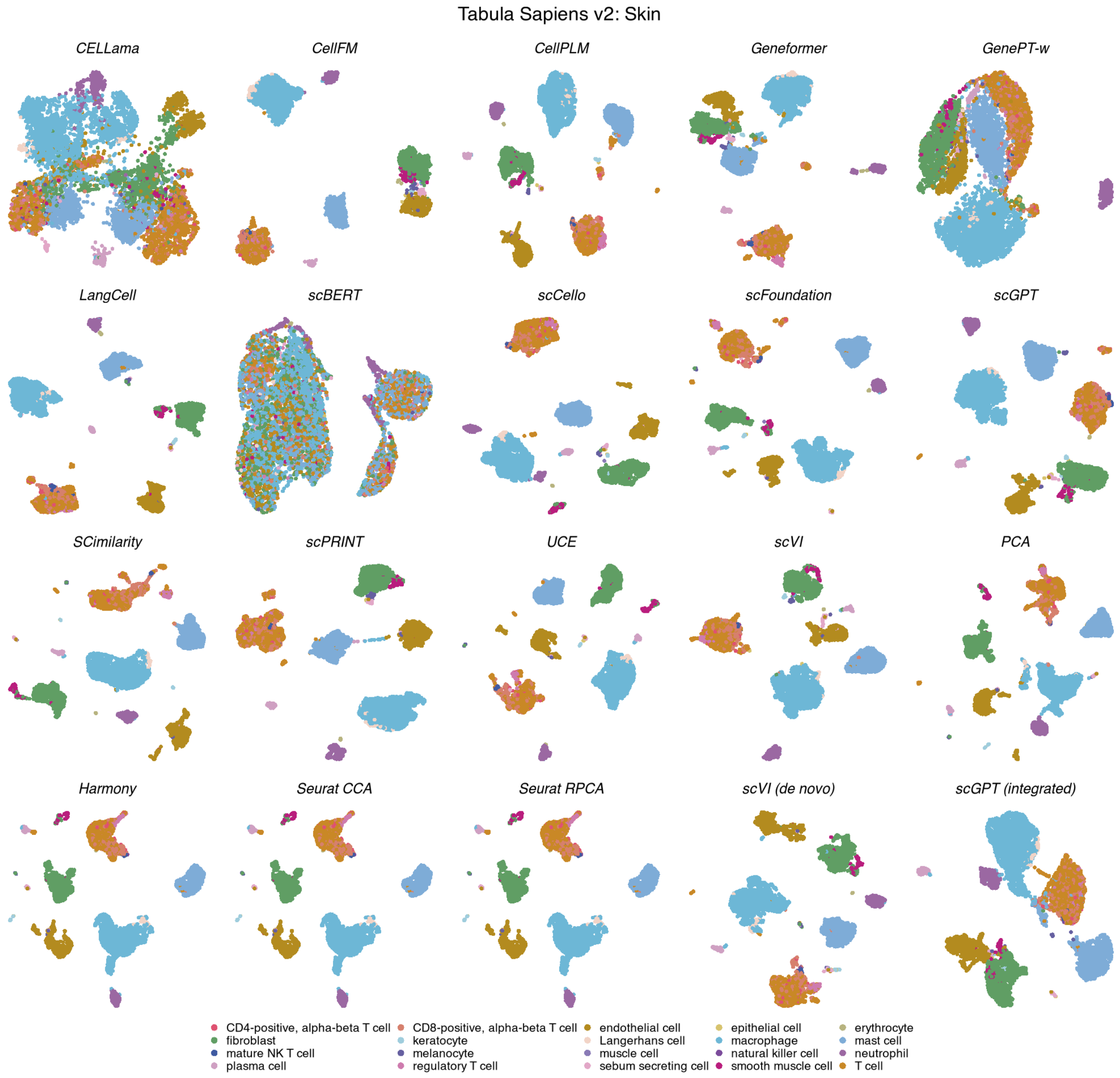

**Figure S59: Cell-type organization in Tabula Sapiens v2 skin.** UMAPs show the same 6,486 cells across 20 methods, coloured by 20 annotated cell types. Display conventions and method categories are as in Fig. S11.

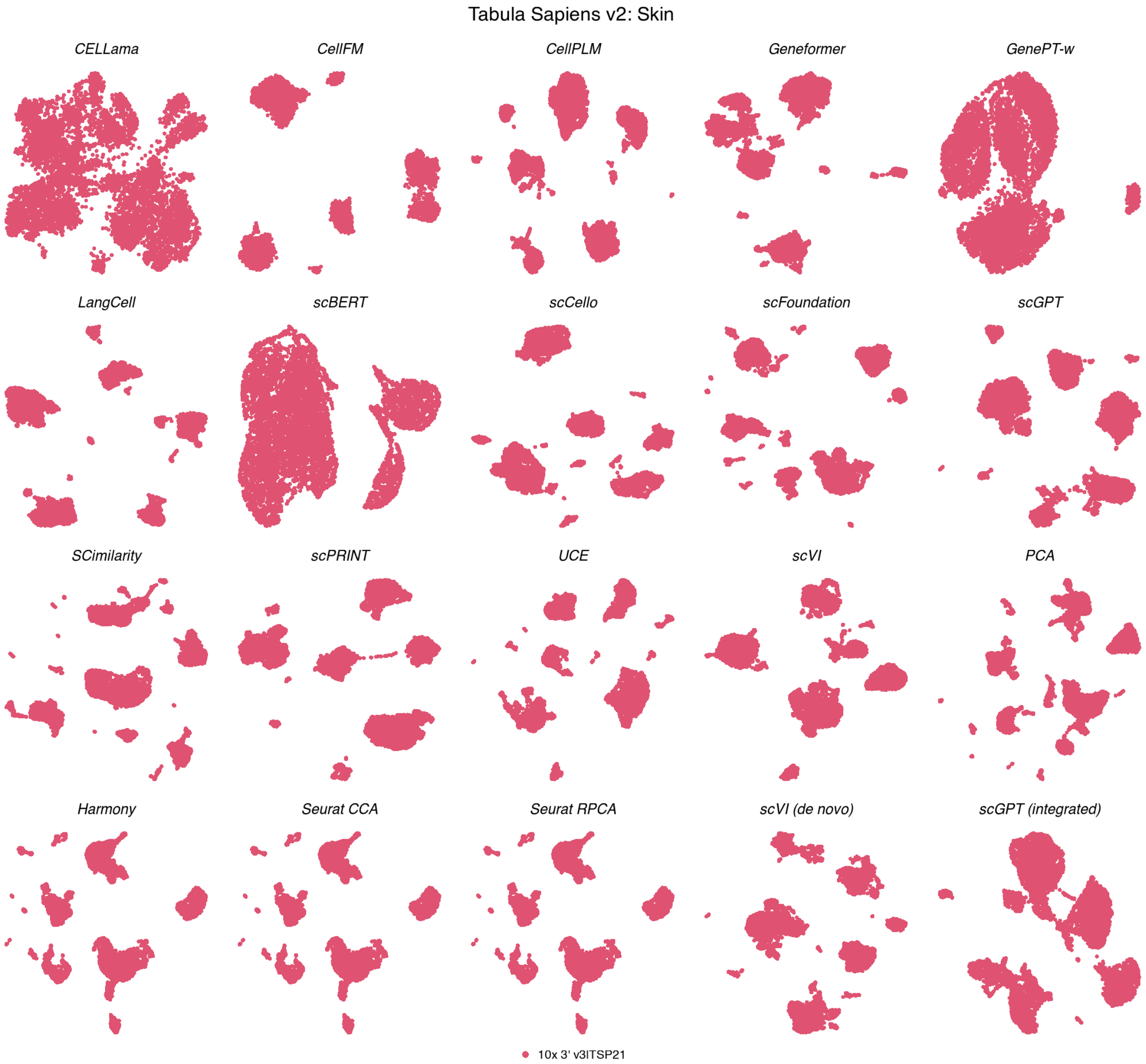

**Figure S60: Batch organization in Tabula Sapiens v2 skin.** UMAPs show the same 6,486 cells across 20 methods, coloured by a single experimental batch. Coordinates match the corresponding cell-type view; display conventions are as in Fig. S12.

**Tabula Sapiens v2 — Skin**

| | Biological conservation metrics | | | | | | | | | Batch mixing metrics | | | Aggregated scores | | | | |
|---|---|---|---|---|---|---|---|---|---|---|---|---|---|---|---|---|---|
| method | ARI | NMI | HOM | COM | FMI | ASW | Acc@kNN | cLISI | GC | kBET | BRAS | CiLISI | Cluster concordance | Continuous separation | Local structure | Batch mixing | Overall |
| SCimilarity | 0.70 | 0.82 | 0.89 | 0.77 | 0.75 | 0.31 | 0.94 | 1.00 | 0.87 | nan | nan | nan | 0.79 | 0.31 | 0.94 | | 0.79 |
| scFoundation | 0.65 | 0.79 | 0.88 | 0.72 | 0.71 | 0.18 | 0.94 | 1.00 | 0.85 | nan | nan | nan | 0.75 | 0.18 | 0.93 | | 0.75 |
| UCE | 0.64 | 0.79 | 0.88 | 0.73 | 0.70 | 0.21 | 0.94 | 1.00 | 0.79 | nan | nan | nan | 0.75 | 0.21 | 0.91 | | 0.75 |
| PCA | 0.69 | 0.80 | 0.88 | 0.74 | 0.75 | 0.05 | 0.94 | 1.00 | 0.85 | nan | nan | nan | 0.77 | 0.05 | 0.93 | | 0.74 |
| CellPLM | 0.56 | 0.77 | 0.87 | 0.70 | 0.64 | 0.15 | 0.93 | 1.00 | 0.87 | nan | nan | nan | 0.71 | 0.15 | 0.93 | | 0.73 |
| scVI (de novo) | 0.62 | 0.78 | 0.88 | 0.70 | 0.70 | 0.06 | 0.93 | 1.00 | 0.86 | nan | nan | nan | 0.73 | 0.06 | 0.93 | | 0.73 |
| Harmony | 0.58 | 0.77 | 0.87 | 0.69 | 0.66 | 0.13 | 0.93 | 1.00 | 0.85 | nan | nan | nan | 0.72 | 0.13 | 0.92 | | 0.73 |
| Seurat CCA | 0.58 | 0.77 | 0.87 | 0.69 | 0.66 | 0.13 | 0.93 | 1.00 | 0.85 | nan | nan | nan | 0.72 | 0.13 | 0.92 | | 0.73 |
| Seurat RPCA | 0.58 | 0.77 | 0.87 | 0.69 | 0.66 | 0.13 | 0.93 | 1.00 | 0.85 | nan | nan | nan | 0.72 | 0.13 | 0.92 | | 0.73 |
| scCello | 0.49 | 0.74 | 0.87 | 0.64 | 0.59 | 0.25 | 0.94 | 1.00 | 0.84 | nan | nan | nan | 0.67 | 0.25 | 0.93 | | 0.72 |
| Geneformer | 0.65 | 0.77 | 0.84 | 0.71 | 0.71 | 0.06 | 0.93 | 1.00 | 0.79 | nan | nan | nan | 0.73 | 0.06 | 0.91 | | 0.72 |
| scGPT | 0.54 | 0.75 | 0.86 | 0.67 | 0.62 | 0.15 | 0.93 | 1.00 | 0.84 | nan | nan | nan | 0.69 | 0.15 | 0.92 | | 0.72 |
| scGPT (integrated) | 0.51 | 0.75 | 0.86 | 0.67 | 0.60 | 0.17 | 0.92 | 1.00 | 0.80 | nan | nan | nan | 0.68 | 0.17 | 0.91 | | 0.71 |
| scVI | 0.52 | 0.75 | 0.86 | 0.66 | 0.61 | 0.13 | 0.93 | 1.00 | 0.81 | nan | nan | nan | 0.68 | 0.13 | 0.91 | | 0.71 |
| LangCell | 0.49 | 0.74 | 0.86 | 0.65 | 0.59 | 0.12 | 0.94 | 1.00 | 0.84 | nan | nan | nan | 0.67 | 0.12 | 0.93 | | 0.71 |
| scPRINT | 0.44 | 0.71 | 0.85 | 0.62 | 0.55 | 0.12 | 0.93 | 1.00 | 0.81 | nan | nan | nan | 0.63 | 0.12 | 0.91 | | 0.69 |
| CellFM | 0.55 | 0.75 | 0.84 | 0.68 | 0.63 | -0.01 | 0.90 | 1.00 | 0.37 | nan | nan | nan | 0.69 | -0.01 | 0.76 | | 0.63 |
| CELLama | 0.30 | 0.51 | 0.60 | 0.44 | 0.41 | -0.01 | 0.83 | 0.98 | 0.58 | nan | nan | nan | 0.45 | -0.01 | 0.80 | | 0.54 |
| GenePT-w | 0.07 | 0.21 | 0.26 | 0.18 | 0.17 | -0.33 | 0.80 | 0.96 | 0.48 | nan | nan | nan | 0.18 | -0.33 | 0.75 | | 0.36 |
| scBERT | 0.04 | 0.12 | 0.15 | 0.11 | 0.14 | -0.27 | 0.45 | 0.88 | 0.30 | nan | nan | nan | 0.11 | -0.27 | 0.55 | | 0.25 |

**Figure S61: Representation performance in Tabula Sapiens v2 skin.** Scores are shown for 6,486 cells, 20 annotated cell types and a single experimental batch. Metrics, family weighting and row ordering are as in Fig. S13. Batch-mixing metrics were unavailable, and the remaining family weights were renormalized.

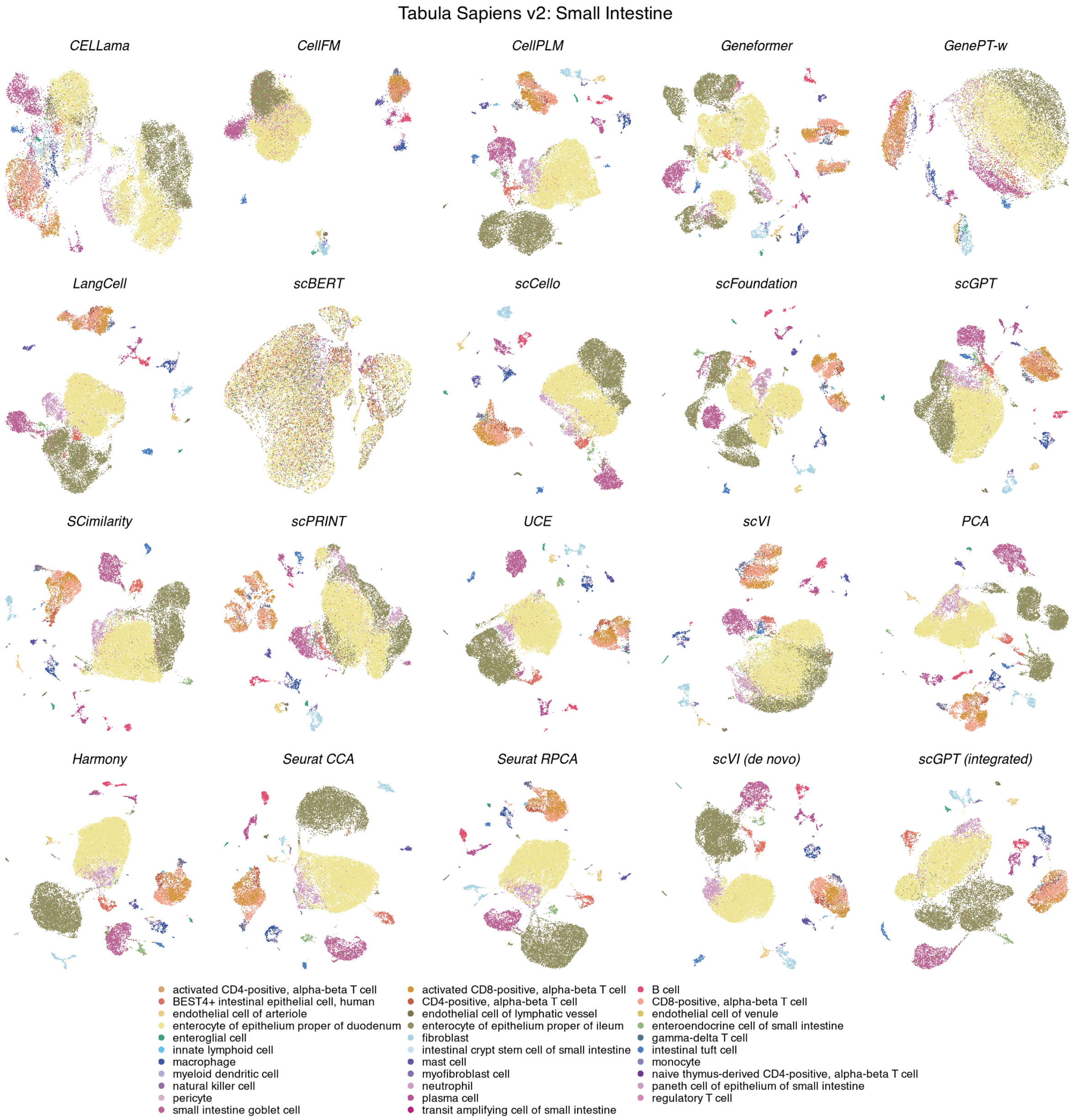

**Figure S62: Cell-type organization in Tabula Sapiens v2 small intestine.** UMAPs show the same 25,834 cells across 20 methods, coloured by 32 annotated cell types. Display conventions and method categories are as in Fig. S11.

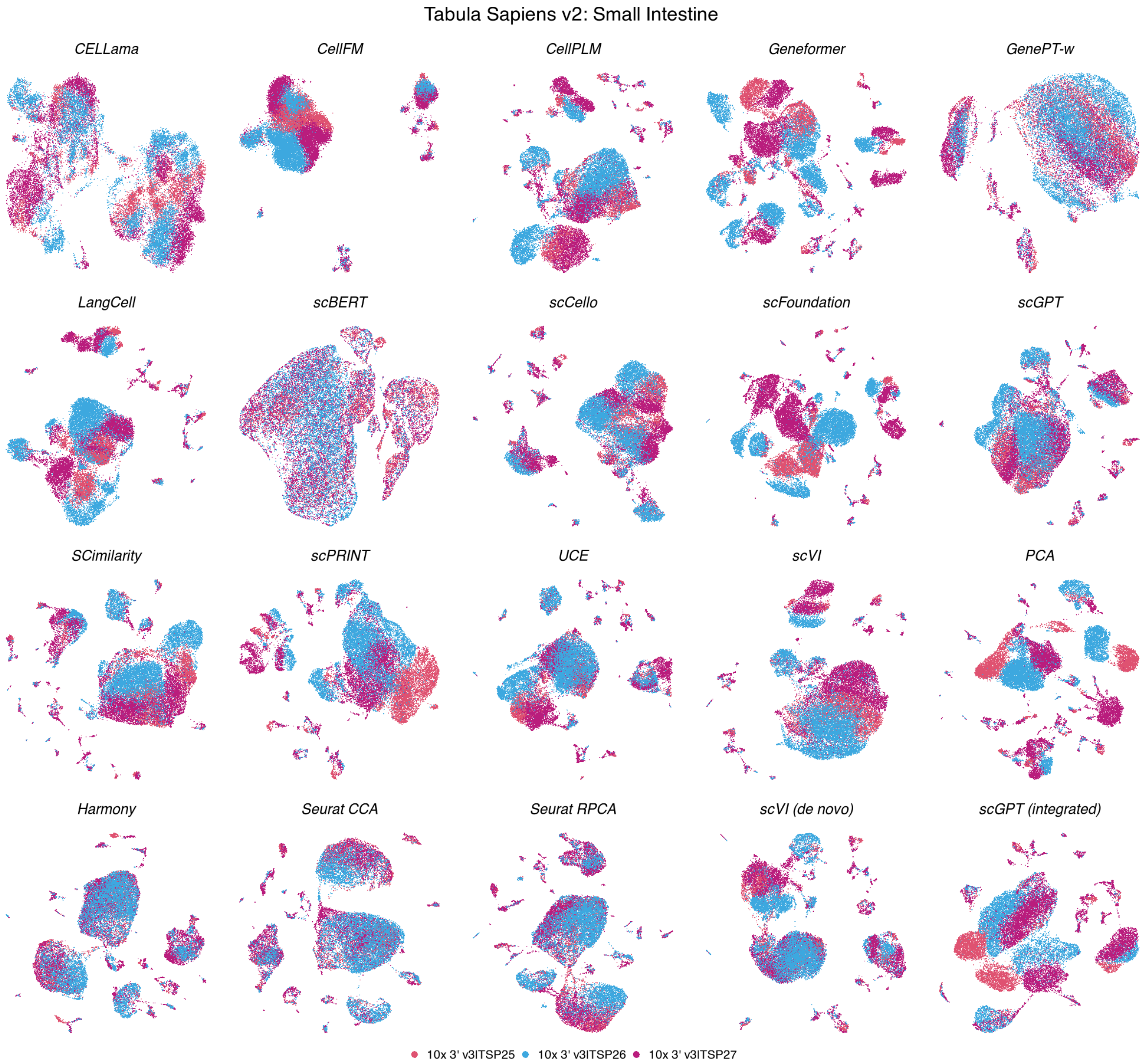

**Figure S63: Batch organization in Tabula Sapiens v2 small intestine.** UMAPs show the same 25,834 cells across 20 methods, coloured by 3 experimental batches. Coordinates match the corresponding cell-type view; display conventions are as in Fig. S12.

**Tabula Sapiens v2 — Small Intestine**

| | Biological conservation metrics | | | | | | | | | Batch mixing metrics | | | Aggregated scores | | | | |
|---|---|---|---|---|---|---|---|---|---|---|---|---|---|---|---|---|---|
| method | ARI | NMI | HOM | COM | FMI | ASW | Acc@kNN | cLISI | GC | kBET | BRAS | CiLISI | Cluster concordance | Continuous separation | Local structure | Batch mixing | Overall |
| Seurat RPCA | 0.28 | 0.68 | 0.86 | 0.56 | 0.44 | 0.34 | 0.94 | 1.00 | 0.87 | 0.67 | 0.73 | 0.36 | 0.56 | 0.34 | 0.94 | 0.59 | 0.67 |
| Seurat CCA | 0.25 | 0.65 | 0.85 | 0.53 | 0.41 | 0.37 | 0.94 | 1.00 | 0.87 | 0.63 | 0.74 | 0.40 | 0.54 | 0.37 | 0.93 | 0.59 | 0.66 |
| Harmony | 0.29 | 0.68 | 0.86 | 0.56 | 0.45 | 0.34 | 0.94 | 1.00 | 0.87 | 0.56 | 0.72 | 0.36 | 0.57 | 0.34 | 0.94 | 0.54 | 0.66 |
| scVI (de novo) | 0.27 | 0.67 | 0.86 | 0.54 | 0.44 | 0.06 | 0.96 | 1.00 | 0.86 | 0.52 | 0.78 | 0.32 | 0.56 | 0.06 | 0.94 | 0.54 | 0.63 |
| scGPT (integrated) | 0.24 | 0.65 | 0.84 | 0.53 | 0.41 | 0.21 | 0.94 | 1.00 | 0.87 | 0.54 | 0.72 | 0.24 | 0.53 | 0.21 | 0.94 | 0.50 | 0.62 |
| PCA | 0.34 | 0.68 | 0.85 | 0.57 | 0.50 | 0.21 | 0.95 | 1.00 | 0.88 | 0.45 | 0.69 | 0.09 | 0.59 | 0.21 | 0.94 | 0.41 | 0.62 |
| UCE | 0.28 | 0.65 | 0.81 | 0.54 | 0.44 | 0.14 | 0.94 | 1.00 | 0.86 | 0.53 | 0.78 | 0.21 | 0.55 | 0.14 | 0.93 | 0.50 | 0.62 |
| SCimilarity | 0.28 | 0.65 | 0.81 | 0.54 | 0.44 | 0.12 | 0.95 | 1.00 | 0.87 | 0.50 | 0.68 | 0.15 | 0.54 | 0.12 | 0.94 | 0.44 | 0.61 |
| CellPLM | 0.21 | 0.63 | 0.84 | 0.51 | 0.38 | 0.11 | 0.95 | 1.00 | 0.88 | 0.46 | 0.71 | 0.14 | 0.51 | 0.11 | 0.94 | 0.44 | 0.59 |
| scFoundation | 0.27 | 0.63 | 0.81 | 0.52 | 0.44 | 0.08 | 0.95 | 1.00 | 0.86 | 0.48 | 0.73 | 0.07 | 0.54 | 0.08 | 0.94 | 0.43 | 0.59 |
| LangCell | 0.23 | 0.61 | 0.80 | 0.49 | 0.40 | 0.05 | 0.93 | 1.00 | 0.84 | 0.46 | 0.83 | 0.12 | 0.51 | 0.05 | 0.92 | 0.47 | 0.59 |
| scCello | 0.21 | 0.59 | 0.76 | 0.48 | 0.37 | 0.09 | 0.91 | 1.00 | 0.82 | 0.51 | 0.81 | 0.21 | 0.48 | 0.09 | 0.91 | 0.51 | 0.59 |
| scGPT | 0.20 | 0.57 | 0.74 | 0.47 | 0.36 | 0.08 | 0.92 | 1.00 | 0.86 | 0.52 | 0.73 | 0.21 | 0.47 | 0.08 | 0.92 | 0.49 | 0.58 |
| Geneformer | 0.21 | 0.57 | 0.75 | 0.46 | 0.37 | 0.04 | 0.95 | 1.00 | 0.78 | 0.47 | 0.75 | 0.10 | 0.47 | 0.04 | 0.91 | 0.44 | 0.56 |
| scVI | 0.17 | 0.51 | 0.66 | 0.41 | 0.33 | 0.08 | 0.88 | 1.00 | 0.84 | 0.51 | 0.70 | 0.16 | 0.42 | 0.08 | 0.91 | 0.46 | 0.55 |
| scPRINT | 0.16 | 0.50 | 0.67 | 0.40 | 0.32 | 0.04 | 0.91 | 1.00 | 0.75 | 0.46 | 0.72 | 0.11 | 0.41 | 0.04 | 0.88 | 0.43 | 0.53 |
| CellFM | 0.18 | 0.48 | 0.61 | 0.40 | 0.33 | 0.00 | 0.69 | 0.97 | 0.37 | 0.57 | 0.97 | 0.40 | 0.40 | 0.00 | 0.68 | 0.65 | 0.51 |
| CELLama | 0.16 | 0.48 | 0.63 | 0.39 | 0.31 | 0.01 | 0.87 | 1.00 | 0.55 | 0.38 | 0.77 | 0.19 | 0.39 | 0.01 | 0.81 | 0.44 | 0.50 |
| GenePT-w | 0.04 | 0.25 | 0.34 | 0.19 | 0.16 | -0.49 | 0.76 | 0.98 | 0.38 | 0.13 | 0.75 | 0.28 | 0.20 | -0.49 | 0.71 | 0.39 | 0.34 |
| scBERT | 0.01 | 0.07 | 0.09 | 0.05 | 0.11 | -0.46 | 0.53 | 0.95 | 0.15 | 0.04 | 0.63 | 0.38 | 0.06 | -0.46 | 0.55 | 0.35 | 0.24 |

**Figure S64: Representation performance in Tabula Sapiens v2 small intestine.** Scores are shown for 25,834 cells, 32 annotated cell types and 3 experimental batches. Metrics, family weighting and row ordering are as in Fig. S13.

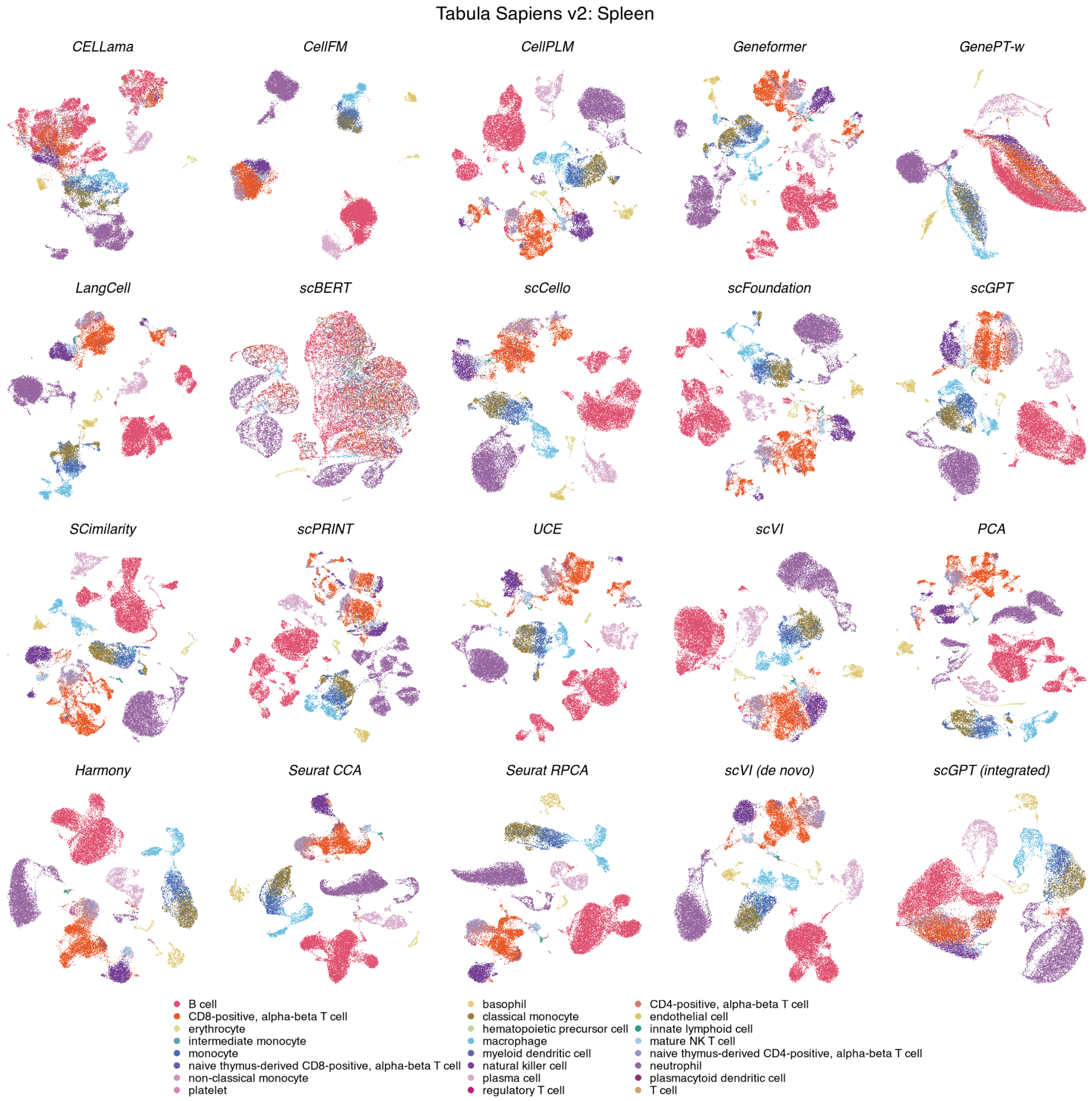

**Figure S65: Cell-type organization in Tabula Sapiens v2 spleen.** UMAPs show the same 30,690 cells across 20 methods, coloured by 24 annotated cell types. Display conventions and method categories are as in Fig. S11.

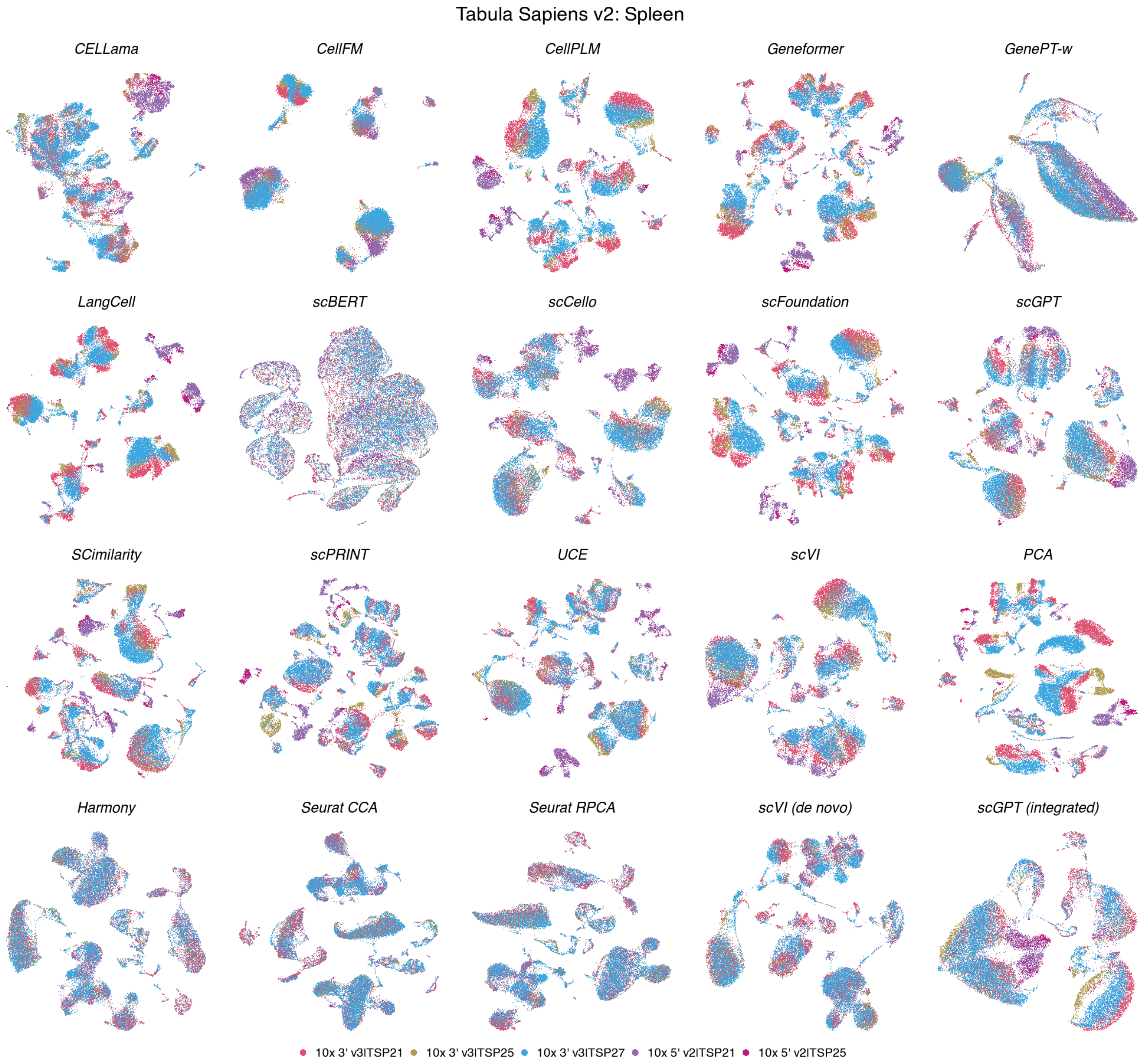

**Figure S66: Batch organization in Tabula Sapiens v2 spleen.** UMAPs show the same 30,690 cells across 20 methods, coloured by 5 experimental batches. Coordinates match the corresponding cell-type view; display conventions are as in Fig. S12.

**Tabula Sapiens v2 — Spleen**

| | Biological conservation metrics | | | | | | | | | Batch mixing metrics | | | Aggregated scores | | | | |
|---|---|---|---|---|---|---|---|---|---|---|---|---|---|---|---|---|---|
| method | ARI | NMI | HOM | COM | FMI | ASW | Acc@kNN | cLISI | GC | kBET | BRAS | CiLISI | Cluster concordance | Continuous separation | Local structure | Batch mixing | Overall |
| scVI (de novo) | 0.66 | 0.79 | 0.86 | 0.74 | 0.71 | 0.10 | 0.93 | 1.00 | 0.88 | 0.41 | 0.80 | 0.30 | 0.75 | 0.10 | 0.94 | 0.50 | 0.68 |
| Seurat RPCA | 0.47 | 0.75 | 0.87 | 0.66 | 0.57 | 0.18 | 0.93 | 1.00 | 0.78 | 0.63 | 0.76 | 0.38 | 0.67 | 0.18 | 0.90 | 0.59 | 0.68 |
| scGPT | 0.72 | 0.79 | 0.84 | 0.75 | 0.77 | 0.23 | 0.94 | 1.00 | 0.84 | 0.35 | 0.66 | 0.15 | 0.77 | 0.23 | 0.93 | 0.38 | 0.67 |
| Harmony | 0.46 | 0.74 | 0.85 | 0.66 | 0.55 | 0.22 | 0.93 | 1.00 | 0.78 | 0.60 | 0.75 | 0.37 | 0.65 | 0.22 | 0.90 | 0.57 | 0.67 |
| Seurat CCA | 0.46 | 0.75 | 0.86 | 0.66 | 0.56 | 0.20 | 0.93 | 1.00 | 0.75 | 0.62 | 0.76 | 0.36 | 0.66 | 0.20 | 0.89 | 0.58 | 0.67 |
| scGPT (integrated) | 0.52 | 0.77 | 0.86 | 0.69 | 0.61 | 0.28 | 0.93 | 1.00 | 0.79 | 0.46 | 0.73 | 0.27 | 0.69 | 0.28 | 0.91 | 0.48 | 0.67 |
| SCimilarity | 0.62 | 0.77 | 0.84 | 0.70 | 0.69 | 0.37 | 0.95 | 1.00 | 0.83 | 0.34 | 0.66 | 0.13 | 0.72 | 0.37 | 0.93 | 0.38 | 0.67 |
| UCE | 0.59 | 0.76 | 0.85 | 0.69 | 0.66 | 0.24 | 0.94 | 1.00 | 0.74 | 0.37 | 0.74 | 0.16 | 0.71 | 0.24 | 0.89 | 0.42 | 0.65 |
| scFoundation | 0.63 | 0.77 | 0.84 | 0.70 | 0.69 | 0.21 | 0.95 | 1.00 | 0.82 | 0.34 | 0.64 | 0.08 | 0.73 | 0.21 | 0.92 | 0.36 | 0.65 |
| scVI | 0.52 | 0.75 | 0.84 | 0.68 | 0.60 | 0.20 | 0.93 | 1.00 | 0.85 | 0.31 | 0.69 | 0.17 | 0.68 | 0.20 | 0.93 | 0.39 | 0.64 |
| scCello | 0.50 | 0.73 | 0.83 | 0.66 | 0.59 | 0.23 | 0.92 | 1.00 | 0.80 | 0.36 | 0.72 | 0.19 | 0.66 | 0.23 | 0.91 | 0.42 | 0.64 |
| CellPLM | 0.62 | 0.76 | 0.83 | 0.70 | 0.68 | 0.23 | 0.94 | 1.00 | 0.82 | 0.32 | 0.55 | 0.09 | 0.72 | 0.23 | 0.92 | 0.32 | 0.64 |
| LangCell | 0.61 | 0.76 | 0.83 | 0.70 | 0.68 | 0.09 | 0.93 | 1.00 | 0.77 | 0.34 | 0.75 | 0.11 | 0.72 | 0.09 | 0.90 | 0.40 | 0.63 |
| PCA | 0.45 | 0.73 | 0.84 | 0.65 | 0.55 | 0.10 | 0.94 | 1.00 | 0.81 | 0.34 | 0.65 | 0.06 | 0.64 | 0.10 | 0.92 | 0.35 | 0.61 |
| Geneformer | 0.43 | 0.69 | 0.79 | 0.61 | 0.53 | 0.10 | 0.94 | 1.00 | 0.77 | 0.30 | 0.71 | 0.10 | 0.61 | 0.10 | 0.90 | 0.37 | 0.59 |
| scPRINT | 0.38 | 0.65 | 0.74 | 0.58 | 0.47 | 0.20 | 0.93 | 1.00 | 0.78 | 0.32 | 0.63 | 0.12 | 0.56 | 0.20 | 0.91 | 0.36 | 0.59 |
| CellFM | 0.40 | 0.69 | 0.80 | 0.61 | 0.50 | -0.01 | 0.85 | 1.00 | 0.31 | 0.36 | 0.97 | 0.31 | 0.60 | -0.01 | 0.72 | 0.55 | 0.56 |
| CELLama | 0.23 | 0.51 | 0.59 | 0.45 | 0.33 | -0.02 | 0.82 | 0.99 | 0.54 | 0.22 | 0.72 | 0.17 | 0.42 | -0.02 | 0.79 | 0.37 | 0.48 |
| GenePT-w | 0.14 | 0.38 | 0.46 | 0.33 | 0.24 | -0.27 | 0.81 | 0.99 | 0.51 | 0.21 | 0.68 | 0.32 | 0.31 | -0.27 | 0.77 | 0.40 | 0.43 |
| scBERT | 0.10 | 0.25 | 0.30 | 0.22 | 0.20 | -0.38 | 0.60 | 0.94 | 0.33 | 0.14 | 0.60 | 0.32 | 0.21 | -0.38 | 0.62 | 0.35 | 0.32 |

**Figure S67: Representation performance in Tabula Sapiens v2 spleen.** Scores are shown for 30,690 cells, 24 annotated cell types and 5 experimental batches. Metrics, family weighting and row ordering are as in Fig. S13.

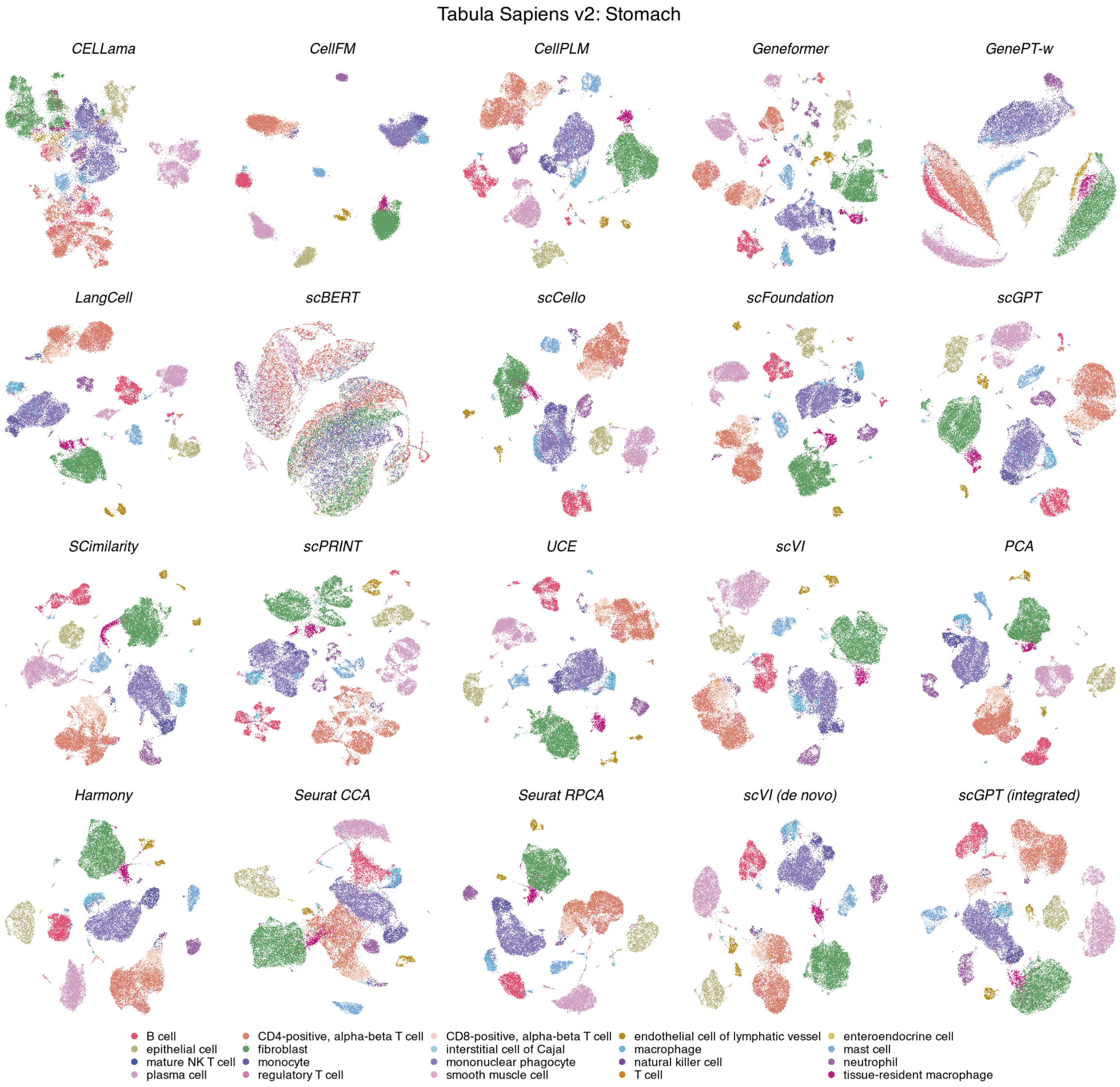

**Figure S68: Cell-type organization in Tabula Sapiens v2 stomach.** UMAPs show the same 33,064 cells across 20 methods, coloured by 20 annotated cell types. Display conventions and method categories are as in Fig. S11.

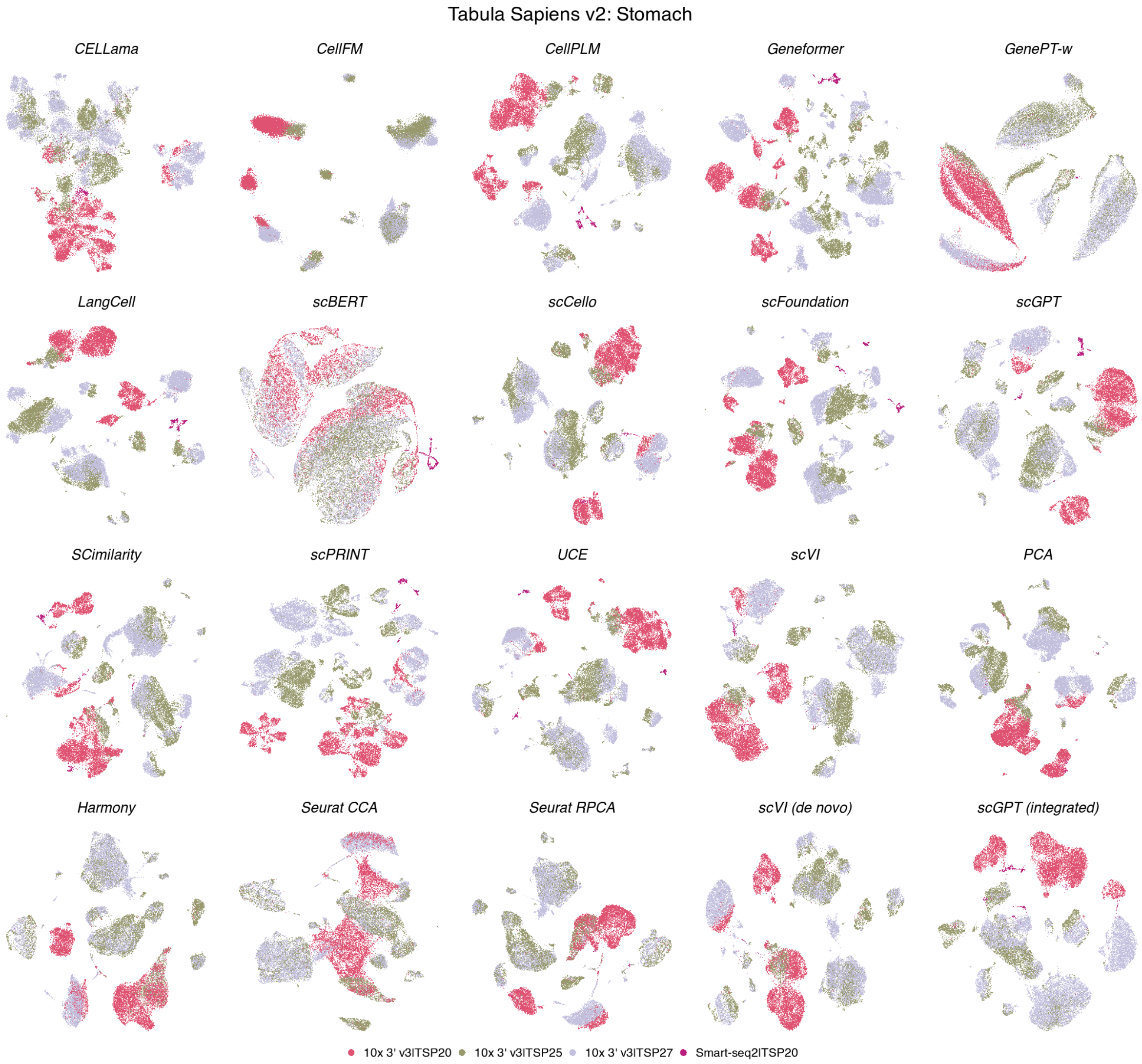

**Figure S69: Batch organization in Tabula Sapiens v2 stomach.** UMAPs show the same 33,064 cells across 20 methods, coloured by 4 experimental batches. Coordinates match the corresponding cell-type view; display conventions are as in Fig. S12.

**Tabula Sapiens v2 — Stomach**

| method | ARI | NMI | HOM | COM | FMI | ASW | Acc@kNN | cLISI | GC | kBET | BRAS | CiLISI | Cluster concordance | Continuous separation | Local structure | Batch mixing | Overall |
|---|---|---|---|---|---|---|---|---|---|---|---|---|---|---|---|---|---|
| | Biological conservation metrics | | | | | | | | | Batch mixing metrics | | | Aggregated scores | | | | |
| **Harmony** | 0.83 | 0.86 | 0.87 | 0.86 | 0.85 | 0.26 | 0.94 | 1.00 | 0.83 | 0.59 | 0.69 | 0.16 | 0.86 | 0.26 | 0.92 | 0.48 | 0.72 |
| **Seurat RPCA** | 0.83 | 0.86 | 0.86 | 0.86 | 0.85 | 0.23 | 0.94 | 1.00 | 0.85 | 0.52 | 0.74 | 0.15 | 0.85 | 0.23 | 0.93 | 0.47 | 0.72 |
| **scVI (de novo)** | 0.82 | 0.86 | 0.87 | 0.85 | 0.84 | 0.10 | 0.95 | 1.00 | 0.86 | 0.57 | 0.78 | 0.12 | 0.85 | 0.10 | 0.94 | 0.49 | 0.71 |
| **SCimilarity** | 0.83 | 0.87 | 0.89 | 0.86 | 0.85 | 0.34 | 0.95 | 1.00 | 0.85 | 0.50 | 0.57 | 0.07 | 0.86 | 0.34 | 0.93 | 0.38 | 0.71 |
| **scGPT (integrated)** | 0.81 | 0.86 | 0.87 | 0.85 | 0.84 | 0.20 | 0.95 | 1.00 | 0.81 | 0.48 | 0.63 | 0.10 | 0.85 | 0.20 | 0.92 | 0.40 | 0.69 |
| **UCE** | 0.80 | 0.87 | 0.83 | 0.92 | 0.84 | 0.20 | 0.95 | 1.00 | 0.73 | 0.48 | 0.68 | 0.08 | 0.85 | 0.20 | 0.89 | 0.41 | 0.69 |
| **Seurat CCA** | 0.65 | 0.81 | 0.87 | 0.76 | 0.70 | 0.19 | 0.94 | 1.00 | 0.83 | 0.60 | 0.71 | 0.19 | 0.76 | 0.19 | 0.92 | 0.50 | 0.69 |
| **scFoundation** | 0.81 | 0.85 | 0.87 | 0.83 | 0.83 | 0.20 | 0.95 | 1.00 | 0.85 | 0.46 | 0.61 | 0.04 | 0.84 | 0.20 | 0.93 | 0.37 | 0.69 |
| **CellPLM** | 0.81 | 0.85 | 0.87 | 0.83 | 0.83 | 0.19 | 0.95 | 1.00 | 0.85 | 0.41 | 0.59 | 0.05 | 0.84 | 0.19 | 0.93 | 0.35 | 0.68 |
| **scGPT** | 0.76 | 0.83 | 0.84 | 0.83 | 0.79 | 0.19 | 0.95 | 1.00 | 0.84 | 0.50 | 0.61 | 0.09 | 0.81 | 0.19 | 0.93 | 0.40 | 0.68 |
| **PCA** | 0.75 | 0.83 | 0.87 | 0.80 | 0.78 | 0.21 | 0.94 | 1.00 | 0.85 | 0.41 | 0.63 | 0.05 | 0.81 | 0.21 | 0.93 | 0.37 | 0.68 |
| **LangCell** | 0.73 | 0.83 | 0.87 | 0.80 | 0.77 | 0.10 | 0.95 | 1.00 | 0.85 | 0.39 | 0.73 | 0.05 | 0.80 | 0.10 | 0.93 | 0.39 | 0.67 |
| **scCello** | 0.59 | 0.79 | 0.86 | 0.73 | 0.64 | 0.22 | 0.94 | 1.00 | 0.83 | 0.45 | 0.70 | 0.09 | 0.72 | 0.22 | 0.92 | 0.41 | 0.66 |
| **scVI** | 0.62 | 0.79 | 0.84 | 0.76 | 0.67 | 0.17 | 0.95 | 1.00 | 0.86 | 0.44 | 0.63 | 0.06 | 0.74 | 0.17 | 0.93 | 0.38 | 0.65 |
| **scPRINT** | 0.51 | 0.75 | 0.83 | 0.68 | 0.58 | 0.17 | 0.94 | 1.00 | 0.85 | 0.42 | 0.59 | 0.04 | 0.67 | 0.17 | 0.93 | 0.35 | 0.63 |
| **Geneformer** | 0.59 | 0.76 | 0.82 | 0.70 | 0.65 | 0.09 | 0.95 | 1.00 | 0.77 | 0.40 | 0.65 | 0.04 | 0.70 | 0.09 | 0.90 | 0.36 | 0.62 |
| **CellFM** | 0.58 | 0.78 | 0.84 | 0.73 | 0.64 | -0.02 | 0.90 | 0.99 | 0.26 | 0.48 | 0.94 | 0.14 | 0.71 | -0.02 | 0.72 | 0.52 | 0.59 |
| **CELLama** | 0.42 | 0.63 | 0.69 | 0.58 | 0.49 | 0.03 | 0.88 | 1.00 | 0.65 | 0.34 | 0.71 | 0.08 | 0.56 | 0.03 | 0.84 | 0.38 | 0.55 |
| **GenePT-w** | 0.23 | 0.45 | 0.49 | 0.41 | 0.31 | -0.23 | 0.86 | 0.99 | 0.58 | 0.38 | 0.66 | 0.15 | 0.38 | -0.23 | 0.81 | 0.40 | 0.46 |
| **scBERT** | 0.08 | 0.19 | 0.21 | 0.18 | 0.17 | -0.28 | 0.58 | 0.93 | 0.25 | 0.26 | 0.60 | 0.16 | 0.17 | -0.28 | 0.59 | 0.34 | 0.30 |

**Figure S70: Representation performance in Tabula Sapiens v2 stomach.** Scores are shown for 33,064 cells, 20 annotated cell types and 4 experimental batches. Metrics, family weighting and row ordering are as in Fig. S13.

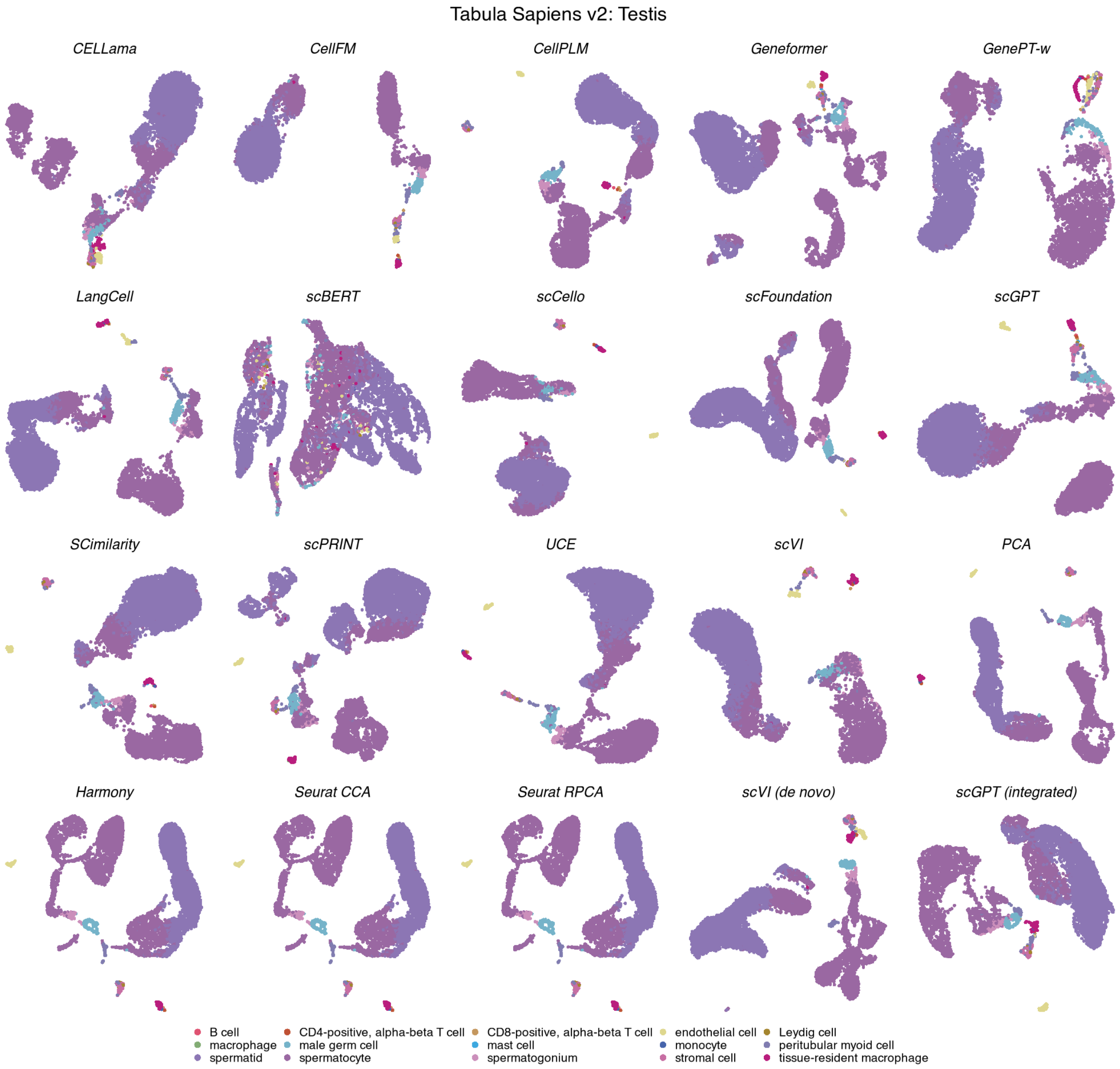

**Figure S71: Cell-type organization in Tabula Sapiens v2 testis.** UMAPs show the same 7,513 cells across 20 methods, coloured by 15 annotated cell types. Display conventions and method categories are as in Fig. S11.

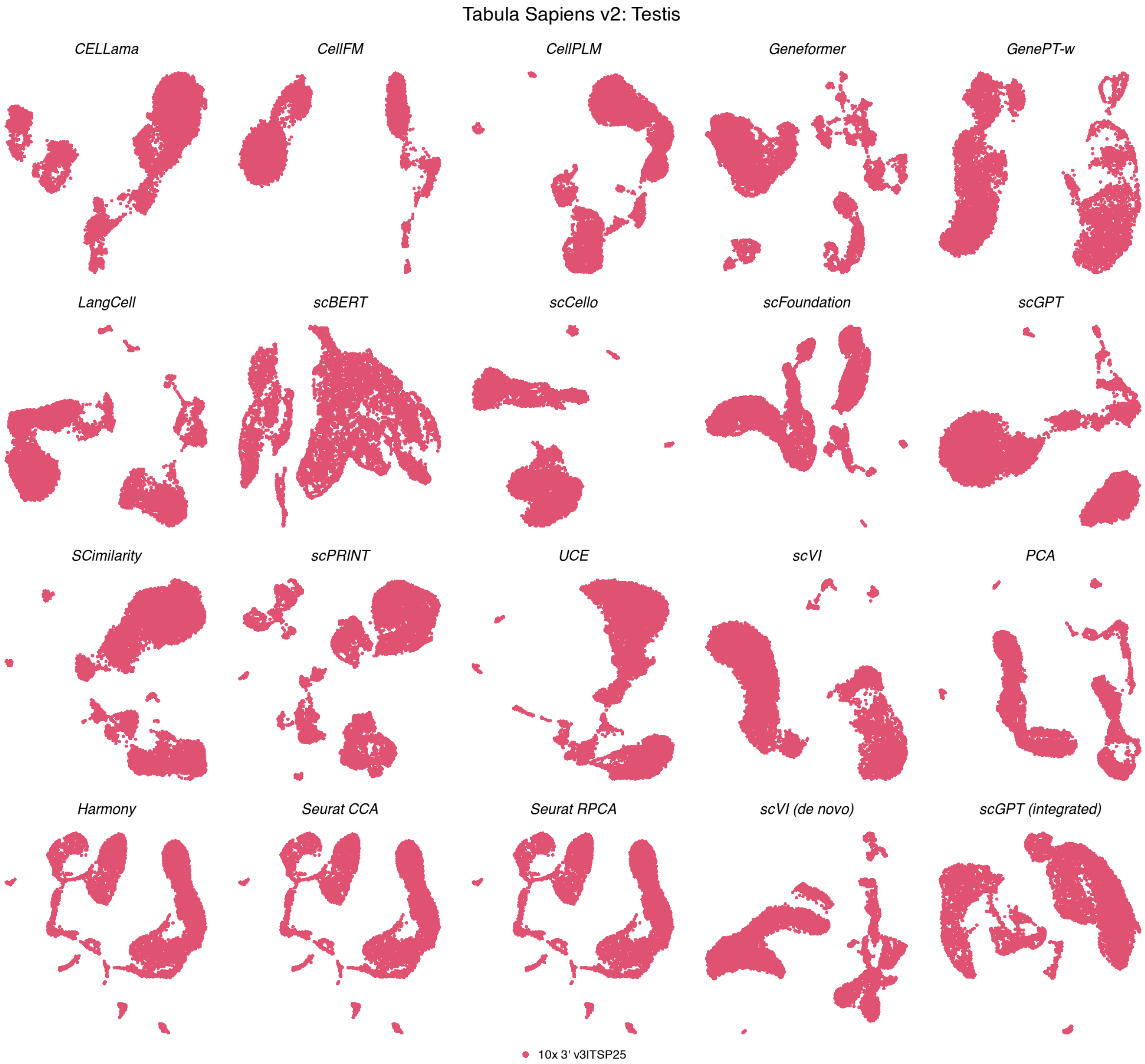

**Figure S72: Batch organization in Tabula Sapiens v2 testis.** UMAPs show the same 7,513 cells across 20 methods, coloured by a single experimental batch. Coordinates match the corresponding cell-type view; display conventions are as in Fig. S12.

**Tabula Sapiens v2 — Testis**

| | Biological conservation metrics | | | | | | | | | Batch mixing metrics | | | Aggregated scores | | | | |
|---|---|---|---|---|---|---|---|---|---|---|---|---|---|---|---|---|---|
| method | ARI | NMI | HOM | COM | FMI | ASW | Acc@kNN | cLISI | GC | kBET | BRAS | CiLISI | Cluster concordance | Continuous separation | Local structure | Batch mixing | Overall |
| Harmony | 0.32 | 0.53 | 0.87 | 0.38 | 0.54 | 0.39 | 0.98 | 1.00 | 0.91 | nan | nan | nan | 0.53 | 0.39 | 0.96 | | 0.70 |
| Seurat CCA | 0.32 | 0.53 | 0.87 | 0.38 | 0.54 | 0.39 | 0.98 | 1.00 | 0.91 | nan | nan | nan | 0.53 | 0.39 | 0.96 | | 0.70 |
| Seurat RPCA | 0.32 | 0.53 | 0.87 | 0.38 | 0.54 | 0.39 | 0.98 | 1.00 | 0.91 | nan | nan | nan | 0.53 | 0.39 | 0.96 | | 0.70 |
| PCA | 0.33 | 0.53 | 0.85 | 0.39 | 0.56 | 0.30 | 0.98 | 1.00 | 0.88 | nan | nan | nan | 0.53 | 0.30 | 0.95 | | 0.68 |
| CellPLM | 0.25 | 0.50 | 0.85 | 0.36 | 0.48 | 0.42 | 0.97 | 1.00 | 0.89 | nan | nan | nan | 0.49 | 0.42 | 0.95 | | 0.68 |
| scGPT | 0.24 | 0.48 | 0.81 | 0.34 | 0.47 | 0.32 | 0.96 | 1.00 | 0.87 | nan | nan | nan | 0.47 | 0.32 | 0.94 | | 0.65 |
| scFoundation | 0.20 | 0.47 | 0.82 | 0.33 | 0.43 | 0.33 | 0.97 | 1.00 | 0.89 | nan | nan | nan | 0.45 | 0.33 | 0.96 | | 0.65 |
| SCimilarity | 0.26 | 0.49 | 0.81 | 0.35 | 0.49 | 0.24 | 0.97 | 1.00 | 0.91 | nan | nan | nan | 0.48 | 0.24 | 0.96 | | 0.65 |
| scGPT (integrated) | 0.20 | 0.46 | 0.80 | 0.33 | 0.43 | 0.29 | 0.97 | 1.00 | 0.90 | nan | nan | nan | 0.45 | 0.29 | 0.96 | | 0.65 |
| LangCell | 0.25 | 0.47 | 0.77 | 0.33 | 0.48 | 0.24 | 0.97 | 1.00 | 0.89 | nan | nan | nan | 0.46 | 0.24 | 0.96 | | 0.64 |
| UCE | 0.21 | 0.47 | 0.81 | 0.33 | 0.44 | 0.24 | 0.96 | 1.00 | 0.90 | nan | nan | nan | 0.45 | 0.24 | 0.95 | | 0.64 |
| scVI (de novo) | 0.24 | 0.48 | 0.83 | 0.33 | 0.47 | 0.08 | 0.98 | 1.00 | 0.89 | nan | nan | nan | 0.47 | 0.08 | 0.96 | | 0.63 |
| scPRINT | 0.21 | 0.44 | 0.76 | 0.31 | 0.44 | 0.24 | 0.96 | 1.00 | 0.86 | nan | nan | nan | 0.43 | 0.24 | 0.94 | | 0.63 |
| scVI | 0.17 | 0.42 | 0.73 | 0.29 | 0.39 | 0.21 | 0.94 | 1.00 | 0.90 | nan | nan | nan | 0.40 | 0.21 | 0.95 | | 0.61 |
| CellFM | 0.33 | 0.53 | 0.82 | 0.39 | 0.55 | 0.03 | 0.97 | 1.00 | 0.64 | nan | nan | nan | 0.52 | 0.03 | 0.87 | | 0.61 |
| Geneformer | 0.19 | 0.41 | 0.73 | 0.29 | 0.42 | 0.22 | 0.97 | 1.00 | 0.81 | nan | nan | nan | 0.41 | 0.22 | 0.93 | | 0.61 |
| scCello | 0.21 | 0.42 | 0.73 | 0.30 | 0.44 | 0.09 | 0.95 | 1.00 | 0.88 | nan | nan | nan | 0.42 | 0.09 | 0.94 | | 0.60 |
| CELLama | 0.22 | 0.40 | 0.68 | 0.28 | 0.45 | 0.19 | 0.95 | 1.00 | 0.76 | nan | nan | nan | 0.40 | 0.19 | 0.90 | | 0.59 |
| GenePT-w | 0.14 | 0.33 | 0.59 | 0.23 | 0.36 | -0.24 | 0.94 | 1.00 | 0.60 | nan | nan | nan | 0.33 | -0.24 | 0.85 | | 0.48 |
| scBERT | 0.12 | 0.25 | 0.45 | 0.18 | 0.34 | -0.57 | 0.90 | 1.00 | 0.50 | nan | nan | nan | 0.27 | -0.57 | 0.80 | | 0.39 |

**Figure S73: Representation performance in Tabula Sapiens v2 testis.** Scores are shown for 7,513 cells, 15 annotated cell types and a single experimental batch. Metrics, family weighting and row ordering are as in Fig. S13. Batch-mixing metrics were unavailable, and the remaining family weights were renormalized.

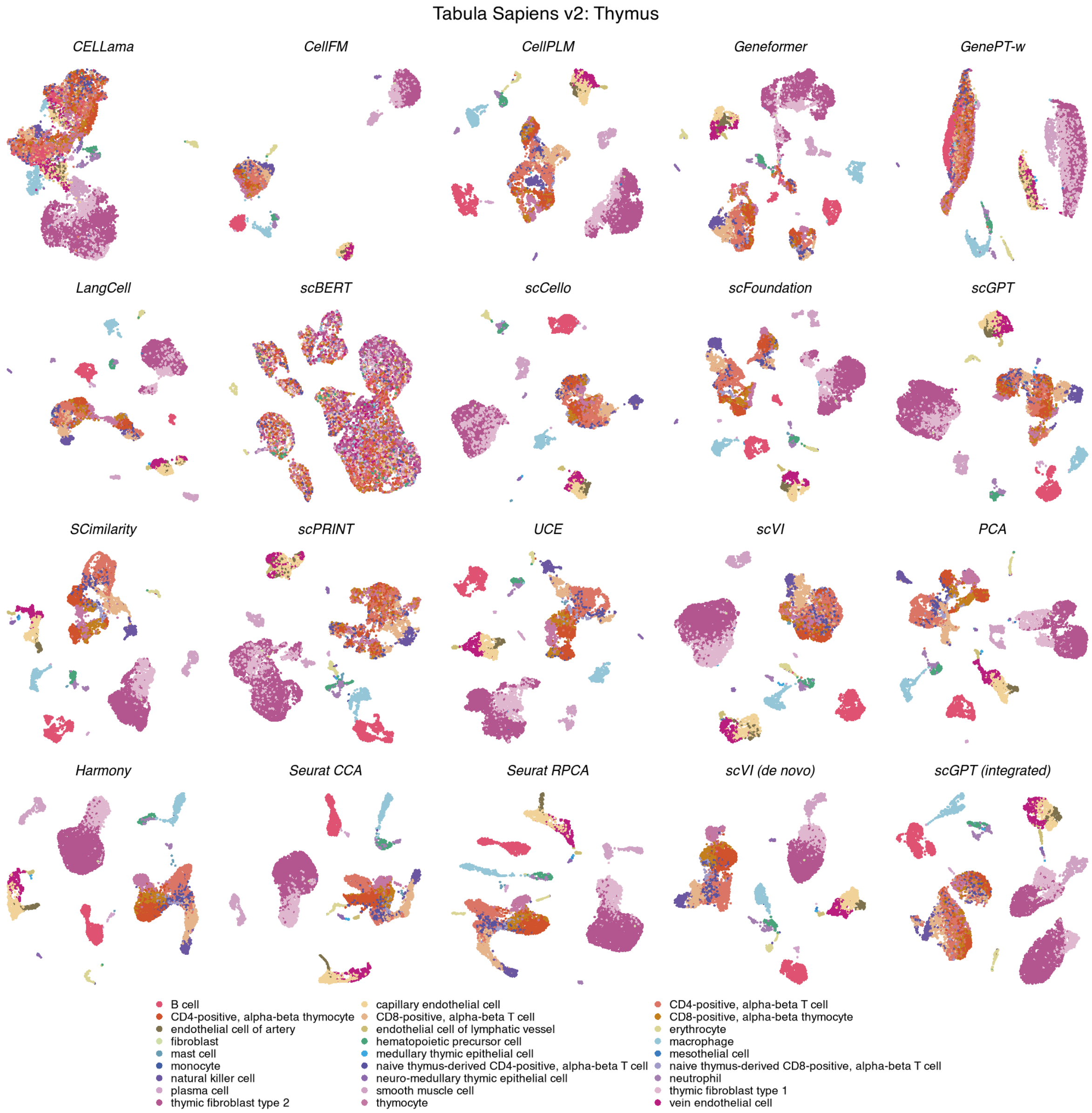

**Figure S74: Cell-type organization in Tabula Sapiens v2 thymus.** UMAPs show the same 9,933 cells across 20 methods, coloured by 27 annotated cell types. Display conventions and method categories are as in Fig. S11.

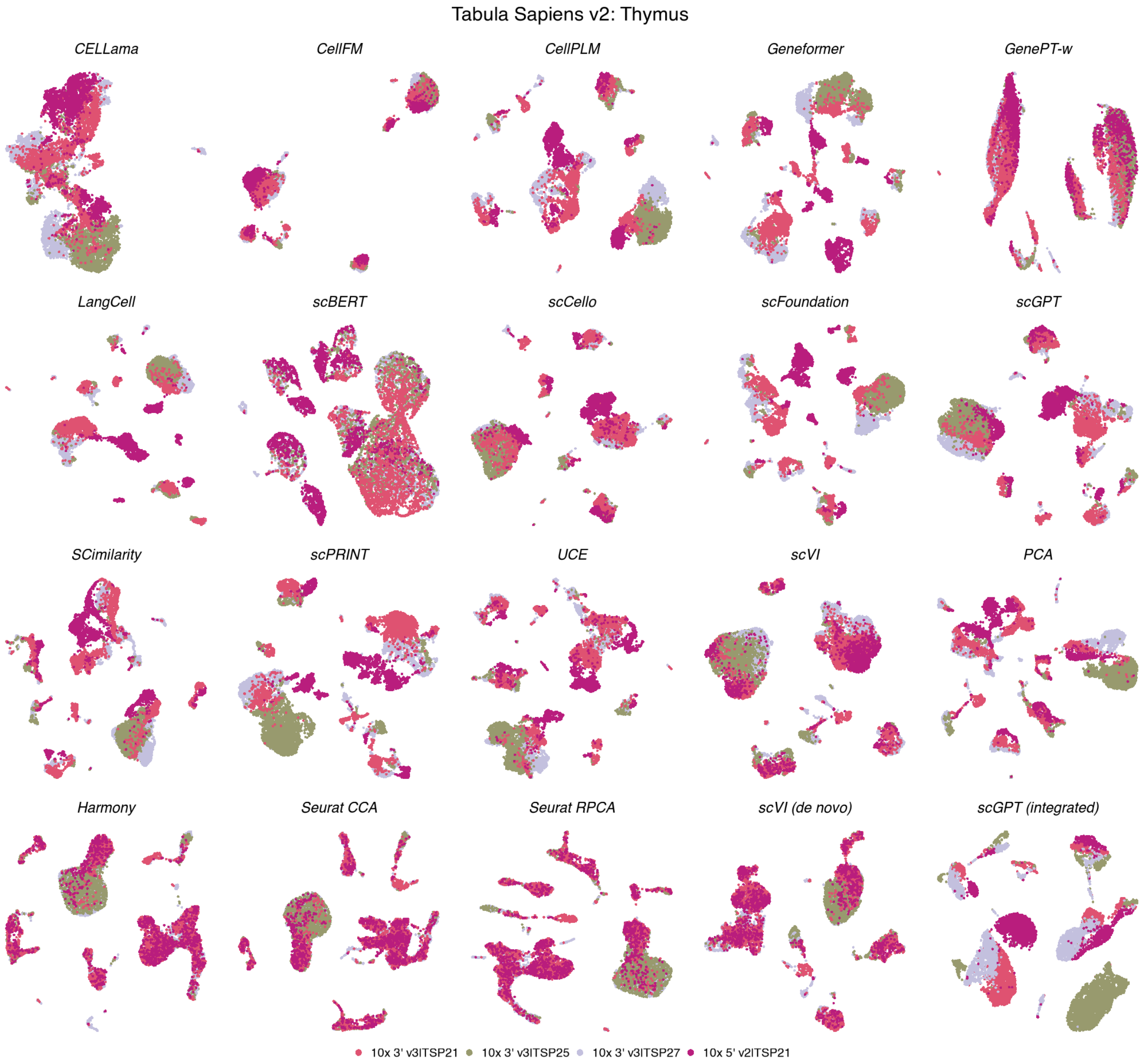

**Figure S75: Batch organization in Tabula Sapiens v2 thymus.** UMAPs show the same 9,933 cells across 20 methods, coloured by 4 experimental batches. Coordinates match the corresponding cell-type view; display conventions are as in Fig. S12.

**Tabula Sapiens v2 — Thymus**

| method | Biological conservation metrics | | | | | | | | | Batch mixing metrics | | | Aggregated scores | | | | |
|---|---|---|---|---|---|---|---|---|---|---|---|---|---|---|---|---|---|
| | ARI | NMI | HOM | COM | FMI | ASW | Acc@kNN | cLISI | GC | kBET | BRAS | CiLISI | Cluster concordance | Continuous separation | Local structure | Batch mixing | Overall |
| **Seurat CCA** | 0.49 | 0.76 | 0.83 | 0.70 | 0.57 | 0.15 | 0.89 | 1.00 | 0.88 | 0.68 | 0.72 | 0.37 | 0.67 | 0.15 | 0.92 | 0.59 | 0.68 |
| **Harmony** | 0.45 | 0.77 | 0.84 | 0.70 | 0.54 | 0.18 | 0.90 | 1.00 | 0.88 | 0.63 | 0.71 | 0.35 | 0.66 | 0.18 | 0.93 | 0.56 | 0.67 |
| **Seurat RPCA** | 0.47 | 0.76 | 0.83 | 0.70 | 0.55 | 0.16 | 0.89 | 1.00 | 0.88 | 0.64 | 0.70 | 0.36 | 0.66 | 0.16 | 0.93 | 0.57 | 0.67 |
| **scVI (de novo)** | 0.41 | 0.75 | 0.83 | 0.68 | 0.49 | 0.07 | 0.90 | 1.00 | 0.89 | 0.45 | 0.75 | 0.32 | 0.63 | 0.07 | 0.93 | 0.51 | 0.64 |
| **SCimilarity** | 0.55 | 0.75 | 0.81 | 0.69 | 0.61 | 0.20 | 0.91 | 1.00 | 0.88 | 0.28 | 0.57 | 0.11 | 0.68 | 0.20 | 0.93 | 0.32 | 0.62 |
| **PCA** | 0.56 | 0.75 | 0.82 | 0.69 | 0.62 | 0.12 | 0.92 | 1.00 | 0.86 | 0.35 | 0.53 | 0.10 | 0.69 | 0.12 | 0.93 | 0.33 | 0.62 |
| **scGPT (integrated)** | 0.39 | 0.72 | 0.79 | 0.65 | 0.48 | 0.23 | 0.88 | 1.00 | 0.88 | 0.39 | 0.55 | 0.15 | 0.61 | 0.23 | 0.92 | 0.36 | 0.61 |
| **UCE** | 0.45 | 0.73 | 0.79 | 0.68 | 0.52 | 0.14 | 0.89 | 1.00 | 0.86 | 0.29 | 0.62 | 0.11 | 0.64 | 0.14 | 0.92 | 0.34 | 0.60 |
| **scFoundation** | 0.44 | 0.70 | 0.76 | 0.65 | 0.51 | 0.14 | 0.91 | 1.00 | 0.81 | 0.26 | 0.55 | 0.06 | 0.61 | 0.14 | 0.90 | 0.29 | 0.58 |
| **scVI** | 0.30 | 0.65 | 0.73 | 0.59 | 0.39 | 0.11 | 0.85 | 0.99 | 0.85 | 0.32 | 0.60 | 0.16 | 0.53 | 0.11 | 0.90 | 0.36 | 0.57 |
| **scCello** | 0.28 | 0.64 | 0.71 | 0.58 | 0.37 | 0.09 | 0.83 | 0.99 | 0.86 | 0.30 | 0.71 | 0.16 | 0.52 | 0.09 | 0.90 | 0.39 | 0.57 |
| **CellPLM** | 0.37 | 0.68 | 0.74 | 0.63 | 0.45 | 0.10 | 0.88 | 1.00 | 0.85 | 0.24 | 0.55 | 0.08 | 0.57 | 0.10 | 0.91 | 0.29 | 0.56 |
| **LangCell** | 0.36 | 0.66 | 0.72 | 0.61 | 0.44 | 0.06 | 0.85 | 0.99 | 0.81 | 0.23 | 0.68 | 0.07 | 0.56 | 0.06 | 0.89 | 0.32 | 0.56 |
| **scGPT** | 0.32 | 0.66 | 0.73 | 0.61 | 0.41 | 0.08 | 0.86 | 0.99 | 0.84 | 0.27 | 0.59 | 0.10 | 0.55 | 0.08 | 0.90 | 0.32 | 0.56 |
| **CellFM** | 0.27 | 0.62 | 0.68 | 0.56 | 0.35 | 0.01 | 0.76 | 0.97 | 0.42 | 0.36 | 0.96 | 0.30 | 0.50 | 0.01 | 0.72 | 0.54 | 0.53 |
| **Geneformer** | 0.29 | 0.61 | 0.68 | 0.55 | 0.37 | 0.04 | 0.87 | 0.99 | 0.70 | 0.25 | 0.66 | 0.07 | 0.50 | 0.04 | 0.85 | 0.33 | 0.53 |
| **scPRINT** | 0.27 | 0.59 | 0.65 | 0.54 | 0.36 | 0.07 | 0.81 | 0.99 | 0.83 | 0.25 | 0.58 | 0.08 | 0.48 | 0.07 | 0.87 | 0.30 | 0.52 |
| **CELLama** | 0.20 | 0.49 | 0.55 | 0.45 | 0.28 | -0.03 | 0.72 | 0.97 | 0.63 | 0.27 | 0.73 | 0.14 | 0.39 | -0.03 | 0.78 | 0.38 | 0.47 |
| **GenePT-w** | 0.09 | 0.31 | 0.35 | 0.27 | 0.17 | -0.35 | 0.64 | 0.94 | 0.41 | 0.24 | 0.63 | 0.28 | 0.24 | -0.35 | 0.66 | 0.38 | 0.35 |
| **scBERT** | 0.02 | 0.10 | 0.12 | 0.09 | 0.10 | -0.47 | 0.34 | 0.87 | 0.21 | 0.10 | 0.53 | 0.27 | 0.09 | -0.47 | 0.48 | 0.30 | 0.21 |

**Figure S76: Representation performance in Tabula Sapiens v2 thymus.** Scores are shown for 9,933 cells, 27 annotated cell types and 4 experimental batches. Metrics, family weighting and row ordering are as in Fig. S13.

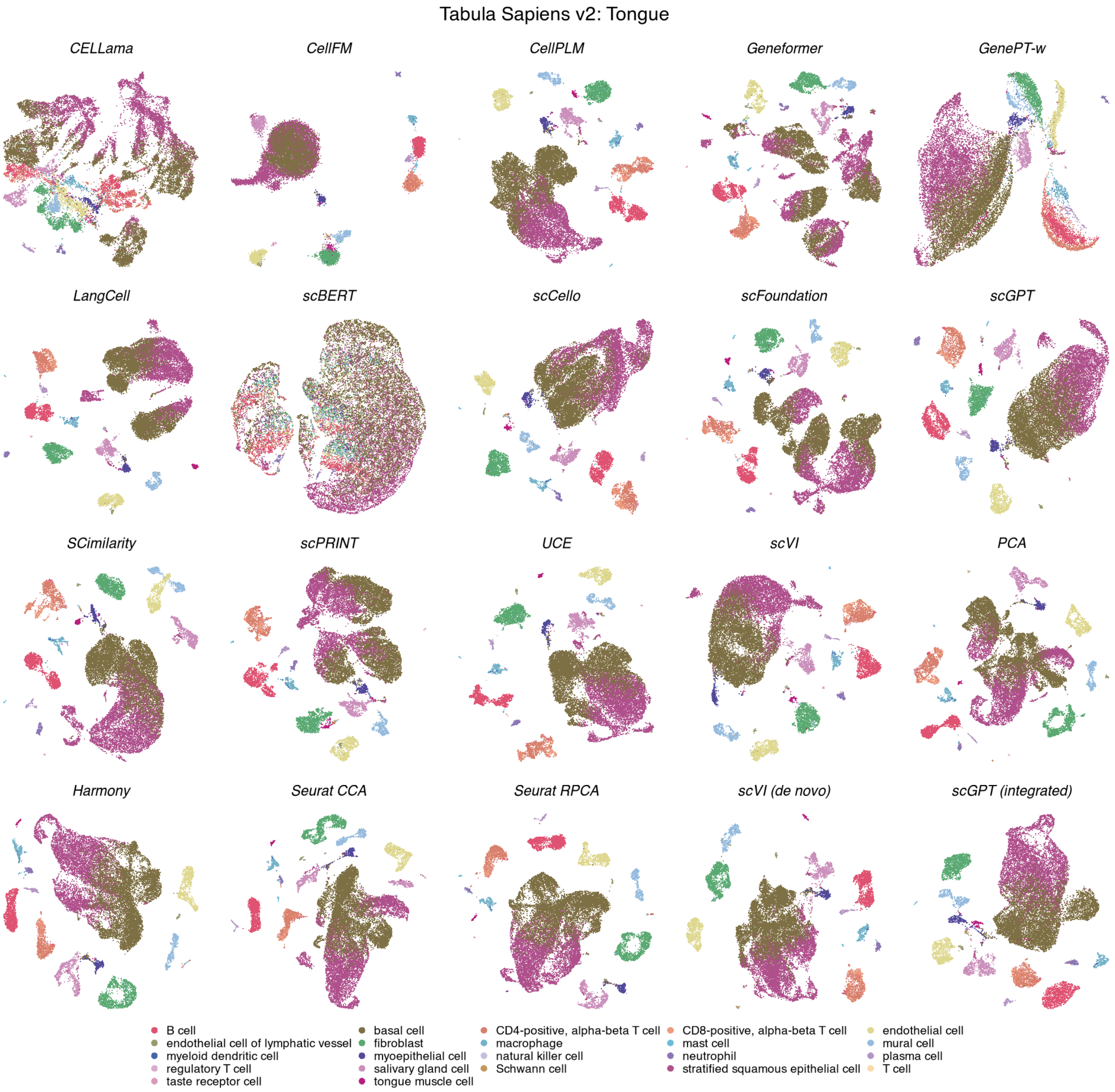

**Figure S77: Cell-type organization in Tabula Sapiens v2 tongue.** UMAPs show the same 21,499 cells across 20 methods, coloured by 22 annotated cell types. Display conventions and method categories are as in Fig. S11.

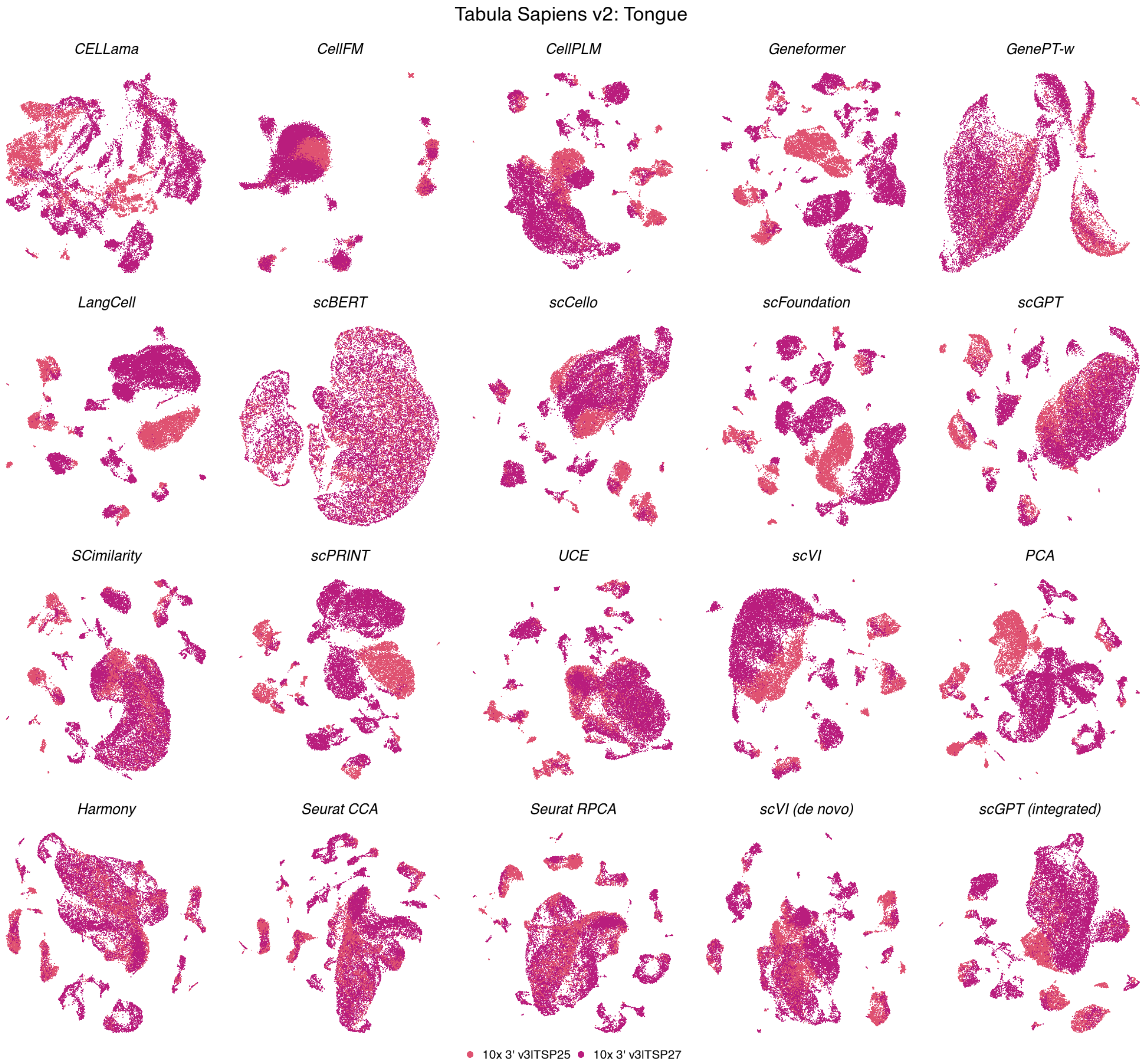

**Figure S78: Batch organization in Tabula Sapiens v2 tongue.** UMAPs show the same 21,499 cells across 20 methods, coloured by 2 experimental batches. Coordinates match the corresponding cell-type view; display conventions are as in Fig. S12.

**Tabula Sapiens v2 — Tongue**

| | Biological conservation metrics | | | | | | | | | Batch mixing metrics | | | Aggregated scores | | | | |
|---|---|---|---|---|---|---|---|---|---|---|---|---|---|---|---|---|---|
| method | ARI | NMI | HOM | COM | FMI | ASW | Acc@kNN | cLISI | GC | kBET | BRAS | CiLISI | Cluster concordance | Continuous separation | Local structure | Batch mixing | Overall |
| Harmony | 0.27 | 0.66 | 0.82 | 0.55 | 0.43 | 0.28 | 0.93 | 1.00 | 0.82 | 0.60 | 0.77 | 0.34 | 0.55 | 0.28 | 0.91 | 0.57 | 0.65 |
| SCimilarity | 0.35 | 0.69 | 0.83 | 0.59 | 0.49 | 0.32 | 0.94 | 1.00 | 0.87 | 0.53 | 0.69 | 0.18 | 0.59 | 0.32 | 0.94 | 0.47 | 0.65 |
| Seurat CCA | 0.26 | 0.65 | 0.82 | 0.54 | 0.42 | 0.28 | 0.92 | 1.00 | 0.82 | 0.53 | 0.73 | 0.41 | 0.54 | 0.28 | 0.92 | 0.56 | 0.64 |
| UCE | 0.35 | 0.68 | 0.79 | 0.59 | 0.49 | 0.22 | 0.93 | 1.00 | 0.81 | 0.54 | 0.80 | 0.22 | 0.58 | 0.22 | 0.91 | 0.52 | 0.64 |
| Seurat RPCA | 0.28 | 0.66 | 0.82 | 0.55 | 0.43 | 0.27 | 0.92 | 1.00 | 0.82 | 0.48 | 0.72 | 0.34 | 0.55 | 0.27 | 0.92 | 0.51 | 0.63 |
| scVI | 0.35 | 0.68 | 0.83 | 0.58 | 0.50 | 0.18 | 0.93 | 1.00 | 0.89 | 0.49 | 0.74 | 0.12 | 0.59 | 0.18 | 0.94 | 0.45 | 0.63 |
| scCello | 0.25 | 0.64 | 0.80 | 0.53 | 0.41 | 0.17 | 0.91 | 1.00 | 0.88 | 0.58 | 0.81 | 0.17 | 0.53 | 0.17 | 0.93 | 0.52 | 0.62 |
| scVI (de novo) | 0.27 | 0.65 | 0.80 | 0.54 | 0.42 | 0.08 | 0.93 | 1.00 | 0.86 | 0.52 | 0.79 | 0.26 | 0.54 | 0.08 | 0.93 | 0.52 | 0.61 |
| scGPT (integrated) | 0.30 | 0.67 | 0.84 | 0.55 | 0.45 | 0.19 | 0.92 | 1.00 | 0.87 | 0.44 | 0.70 | 0.19 | 0.56 | 0.19 | 0.93 | 0.44 | 0.61 |
| CellPLM | 0.28 | 0.66 | 0.81 | 0.55 | 0.43 | 0.23 | 0.94 | 1.00 | 0.88 | 0.43 | 0.72 | 0.12 | 0.55 | 0.23 | 0.94 | 0.42 | 0.61 |
| scGPT | 0.27 | 0.66 | 0.83 | 0.55 | 0.43 | 0.15 | 0.94 | 1.00 | 0.86 | 0.48 | 0.70 | 0.18 | 0.55 | 0.15 | 0.93 | 0.45 | 0.61 |
| scFoundation | 0.29 | 0.66 | 0.80 | 0.55 | 0.44 | 0.21 | 0.95 | 1.00 | 0.86 | 0.47 | 0.69 | 0.04 | 0.55 | 0.21 | 0.94 | 0.40 | 0.60 |
| LangCell | 0.28 | 0.66 | 0.81 | 0.55 | 0.43 | 0.13 | 0.93 | 1.00 | 0.91 | 0.40 | 0.80 | 0.05 | 0.55 | 0.13 | 0.95 | 0.41 | 0.60 |
| PCA | 0.29 | 0.65 | 0.78 | 0.56 | 0.44 | 0.22 | 0.92 | 1.00 | 0.84 | 0.47 | 0.70 | 0.06 | 0.54 | 0.22 | 0.92 | 0.41 | 0.60 |
| scPRINT | 0.26 | 0.61 | 0.76 | 0.51 | 0.41 | 0.18 | 0.93 | 1.00 | 0.89 | 0.40 | 0.70 | 0.05 | 0.51 | 0.18 | 0.94 | 0.39 | 0.59 |
| Geneformer | 0.22 | 0.59 | 0.73 | 0.50 | 0.37 | 0.06 | 0.94 | 1.00 | 0.86 | 0.37 | 0.75 | 0.04 | 0.48 | 0.06 | 0.93 | 0.39 | 0.56 |
| CellFM | 0.28 | 0.61 | 0.77 | 0.51 | 0.43 | 0.01 | 0.80 | 0.99 | 0.36 | 0.62 | 0.96 | 0.13 | 0.52 | 0.01 | 0.71 | 0.57 | 0.54 |
| CELLama | 0.12 | 0.42 | 0.56 | 0.34 | 0.27 | 0.06 | 0.87 | 1.00 | 0.62 | 0.40 | 0.74 | 0.08 | 0.34 | 0.06 | 0.83 | 0.41 | 0.49 |
| GenePT-w | 0.06 | 0.25 | 0.33 | 0.20 | 0.19 | -0.34 | 0.86 | 0.99 | 0.48 | 0.29 | 0.71 | 0.32 | 0.21 | -0.34 | 0.78 | 0.44 | 0.40 |
| scBERT | 0.04 | 0.17 | 0.23 | 0.13 | 0.17 | -0.13 | 0.64 | 0.96 | 0.21 | 0.24 | 0.35 | 0.43 | 0.15 | -0.13 | 0.60 | 0.34 | 0.32 |

**Figure S79: Representation performance in Tabula Sapiens v2 tongue.** Scores are shown for 21,499 cells, 22 annotated cell types and 2 experimental batches. Metrics, family weighting and row ordering are as in Fig. S13.

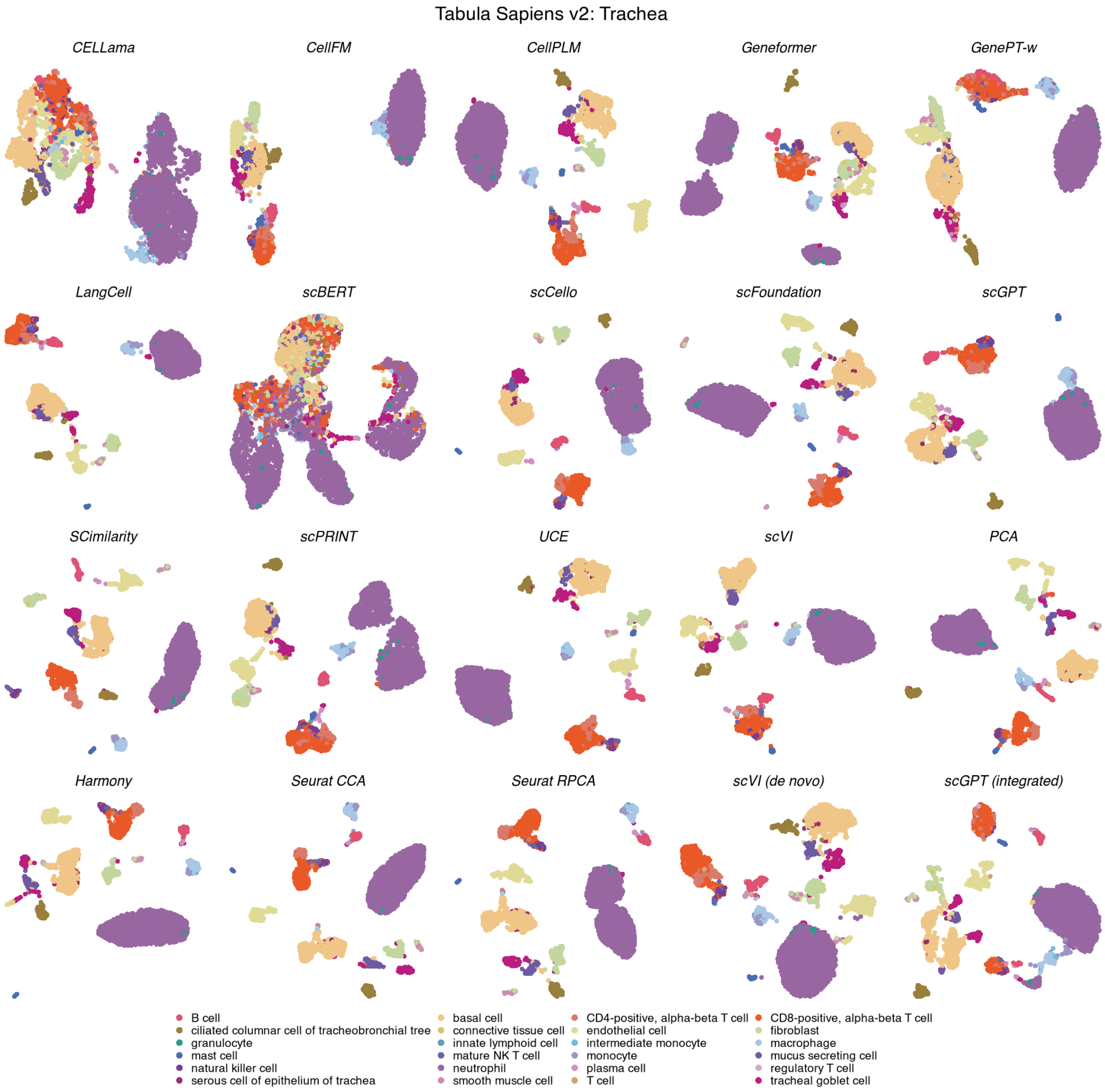

**Figure S80: Cell-type organization in Tabula Sapiens v2 trachea.** UMAPs show the same 5,346 cells across 20 methods, coloured by 24 annotated cell types. Display conventions and method categories are as in Fig. S11.

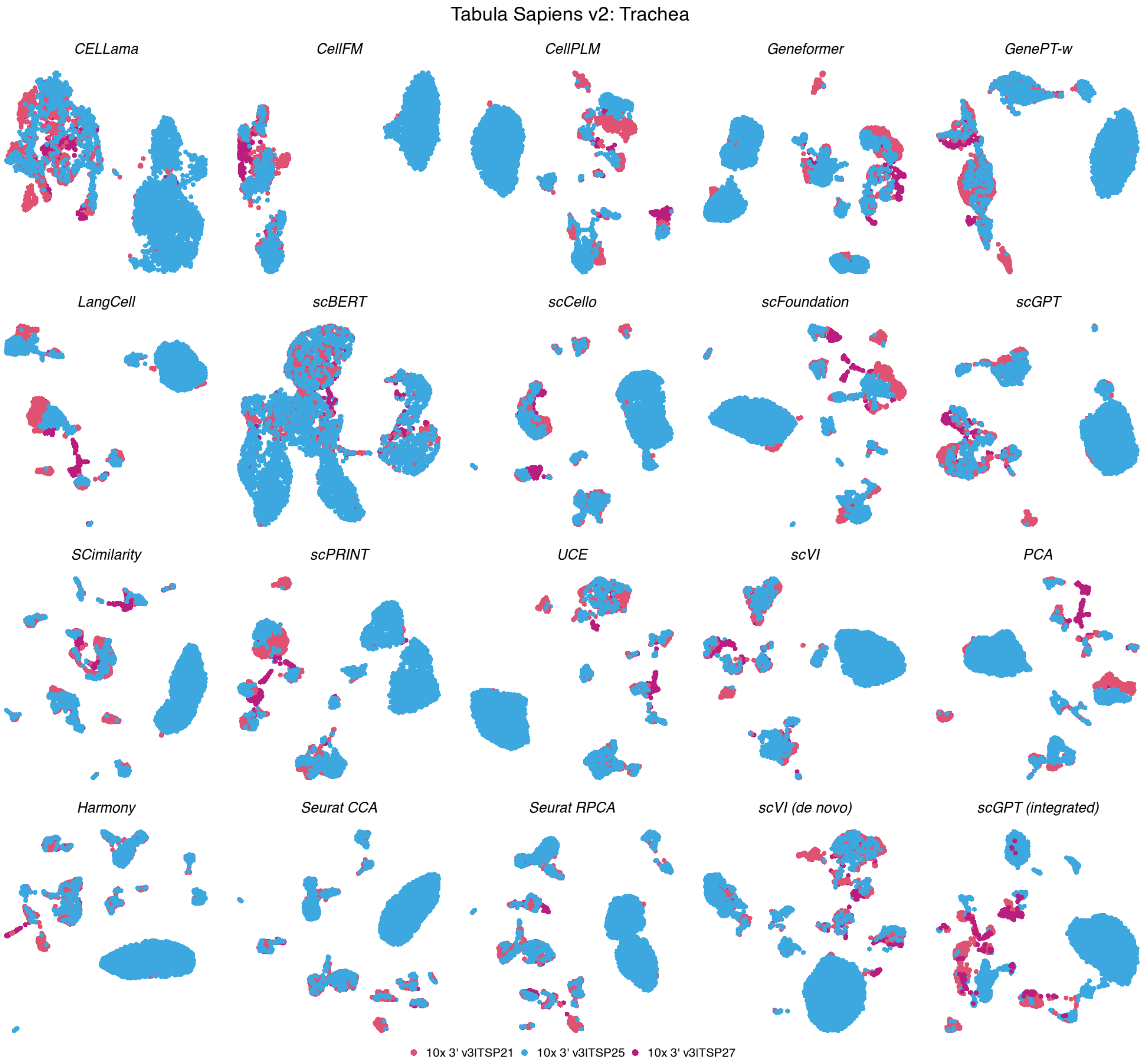

**Figure S81: Batch organization in Tabula Sapiens v2 trachea.** UMAPs show the same 5,346 cells across 20 methods, coloured by 3 experimental batches. Coordinates match the corresponding cell-type view; display conventions are as in Fig. S12.

**Tabula Sapiens v2 — Trachea**

| method | Biological conservation metrics | | | | | | | | | Batch mixing metrics | | | Aggregated scores | | | | |
|---|---|---|---|---|---|---|---|---|---|---|---|---|---|---|---|---|---|
| | ARI | NMI | HOM | COM | FMI | ASW | Acc@kNN | cLISI | GC | kBET | BRAS | CiLISI | Cluster concordance | Continuous separation | Local structure | Batch mixing | Overall |
| Seurat CCA | 0.33 | 0.71 | 0.89 | 0.60 | 0.52 | 0.19 | 0.96 | 1.00 | 0.76 | 0.78 | 0.68 | 0.19 | 0.61 | 0.19 | 0.91 | 0.55 | 0.65 |
| Seurat RPCA | 0.28 | 0.69 | 0.90 | 0.56 | 0.47 | 0.19 | 0.97 | 1.00 | 0.75 | 0.82 | 0.68 | 0.17 | 0.58 | 0.19 | 0.91 | 0.56 | 0.64 |
| Harmony | 0.28 | 0.70 | 0.90 | 0.57 | 0.48 | 0.25 | 0.96 | 1.00 | 0.75 | 0.75 | 0.67 | 0.19 | 0.58 | 0.25 | 0.90 | 0.53 | 0.64 |
| SCimilarity | 0.28 | 0.71 | 0.92 | 0.58 | 0.48 | 0.26 | 0.97 | 1.00 | 0.78 | 0.66 | 0.65 | 0.12 | 0.59 | 0.26 | 0.92 | 0.47 | 0.63 |
| PCA | 0.34 | 0.69 | 0.89 | 0.57 | 0.53 | 0.30 | 0.97 | 1.00 | 0.77 | 0.65 | 0.56 | 0.09 | 0.60 | 0.30 | 0.91 | 0.43 | 0.63 |
| UCE | 0.24 | 0.68 | 0.91 | 0.54 | 0.43 | 0.16 | 0.97 | 1.00 | 0.76 | 0.65 | 0.75 | 0.12 | 0.56 | 0.16 | 0.91 | 0.51 | 0.62 |
| scGPT (integrated) | 0.24 | 0.68 | 0.90 | 0.55 | 0.44 | 0.27 | 0.97 | 1.00 | 0.80 | 0.55 | 0.64 | 0.11 | 0.56 | 0.27 | 0.92 | 0.44 | 0.62 |
| scFoundation | 0.26 | 0.69 | 0.91 | 0.55 | 0.45 | 0.17 | 0.97 | 1.00 | 0.80 | 0.59 | 0.69 | 0.09 | 0.57 | 0.17 | 0.92 | 0.46 | 0.62 |
| scCello | 0.20 | 0.66 | 0.90 | 0.52 | 0.40 | 0.11 | 0.97 | 1.00 | 0.78 | 0.70 | 0.77 | 0.12 | 0.54 | 0.11 | 0.91 | 0.53 | 0.62 |
| LangCell | 0.27 | 0.69 | 0.90 | 0.56 | 0.47 | 0.09 | 0.97 | 1.00 | 0.81 | 0.58 | 0.77 | 0.07 | 0.58 | 0.09 | 0.92 | 0.47 | 0.61 |
| scVI (de novo) | 0.20 | 0.66 | 0.90 | 0.52 | 0.40 | 0.07 | 0.97 | 1.00 | 0.83 | 0.65 | 0.76 | 0.15 | 0.54 | 0.07 | 0.93 | 0.52 | 0.61 |
| scGPT | 0.24 | 0.68 | 0.89 | 0.54 | 0.44 | 0.15 | 0.97 | 1.00 | 0.79 | 0.57 | 0.72 | 0.11 | 0.56 | 0.15 | 0.92 | 0.47 | 0.61 |
| scVI | 0.20 | 0.65 | 0.89 | 0.52 | 0.40 | 0.17 | 0.97 | 1.00 | 0.79 | 0.63 | 0.68 | 0.14 | 0.53 | 0.17 | 0.92 | 0.48 | 0.61 |
| Geneformer | 0.26 | 0.65 | 0.85 | 0.53 | 0.45 | 0.06 | 0.96 | 1.00 | 0.77 | 0.62 | 0.78 | 0.11 | 0.55 | 0.06 | 0.91 | 0.50 | 0.60 |
| CellPLM | 0.25 | 0.67 | 0.89 | 0.54 | 0.44 | 0.17 | 0.97 | 1.00 | 0.77 | 0.53 | 0.63 | 0.08 | 0.56 | 0.17 | 0.91 | 0.41 | 0.60 |
| scPRINT | 0.19 | 0.64 | 0.87 | 0.51 | 0.39 | 0.11 | 0.97 | 1.00 | 0.81 | 0.58 | 0.68 | 0.09 | 0.52 | 0.11 | 0.92 | 0.45 | 0.59 |
| CellFM | 0.19 | 0.60 | 0.81 | 0.48 | 0.38 | -0.03 | 0.92 | 1.00 | 0.44 | 0.64 | 0.95 | 0.18 | 0.49 | -0.03 | 0.79 | 0.59 | 0.56 |
| CELLama | 0.18 | 0.53 | 0.71 | 0.43 | 0.36 | -0.00 | 0.91 | 1.00 | 0.56 | 0.49 | 0.80 | 0.15 | 0.44 | -0.00 | 0.82 | 0.48 | 0.53 |
| GenePT-w | 0.09 | 0.36 | 0.50 | 0.29 | 0.26 | -0.25 | 0.87 | 1.00 | 0.43 | 0.35 | 0.67 | 0.18 | 0.30 | -0.25 | 0.77 | 0.40 | 0.42 |
| scBERT | 0.07 | 0.30 | 0.41 | 0.23 | 0.23 | -0.33 | 0.76 | 0.99 | 0.28 | 0.32 | 0.52 | 0.19 | 0.25 | -0.33 | 0.67 | 0.34 | 0.35 |

**Figure S82: Representation performance in Tabula Sapiens v2 trachea.** Scores are shown for 5,346 cells, 24 annotated cell types and 3 experimental batches. Metrics, family weighting and row ordering are as in Fig. S13.

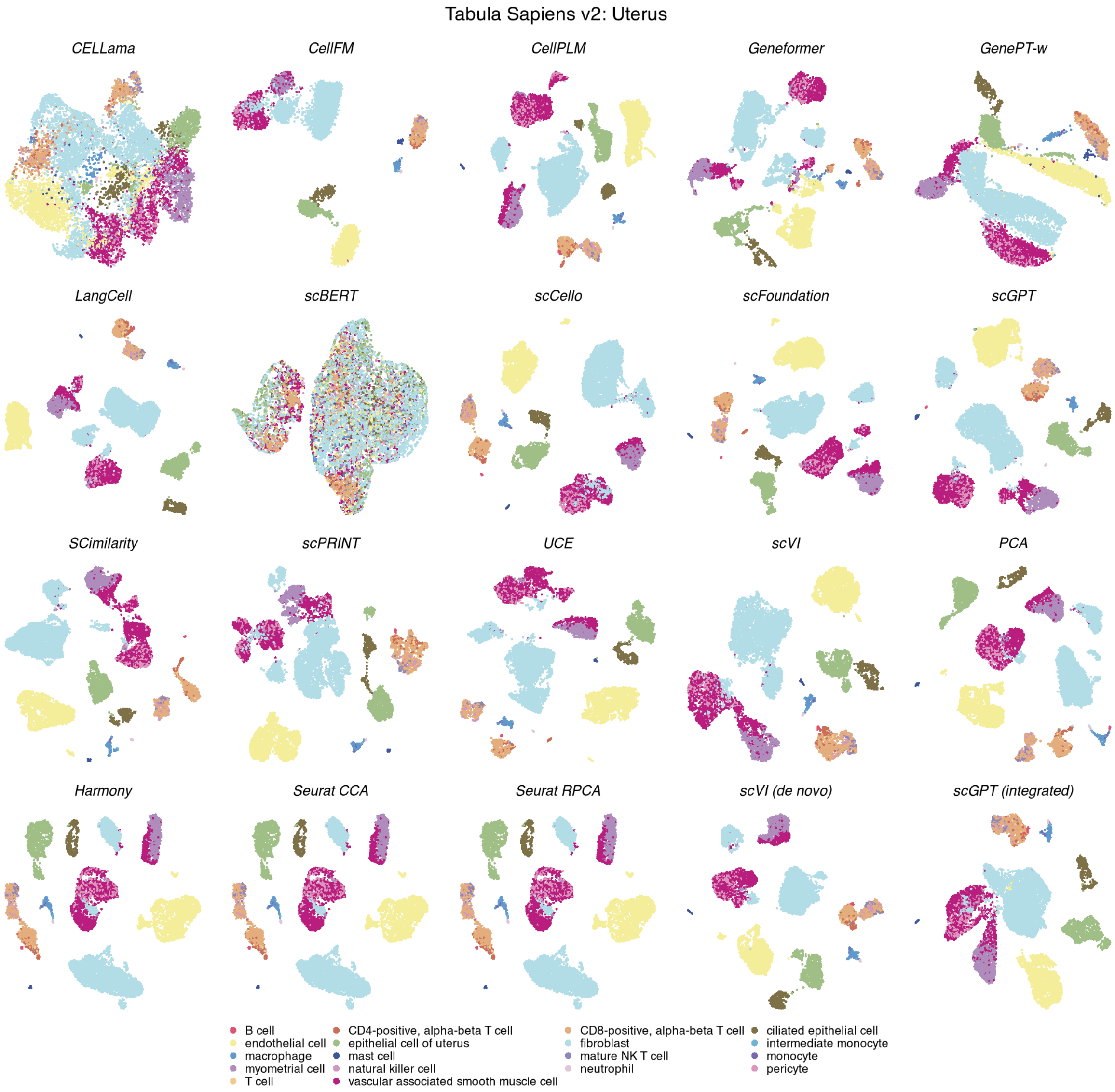

**Figure S83: Cell-type organization in Tabula Sapiens v2 uterus.** UMAPs show the same 10,580 cells across 20 methods, coloured by 18 annotated cell types. Display conventions and method categories are as in Fig. S11.

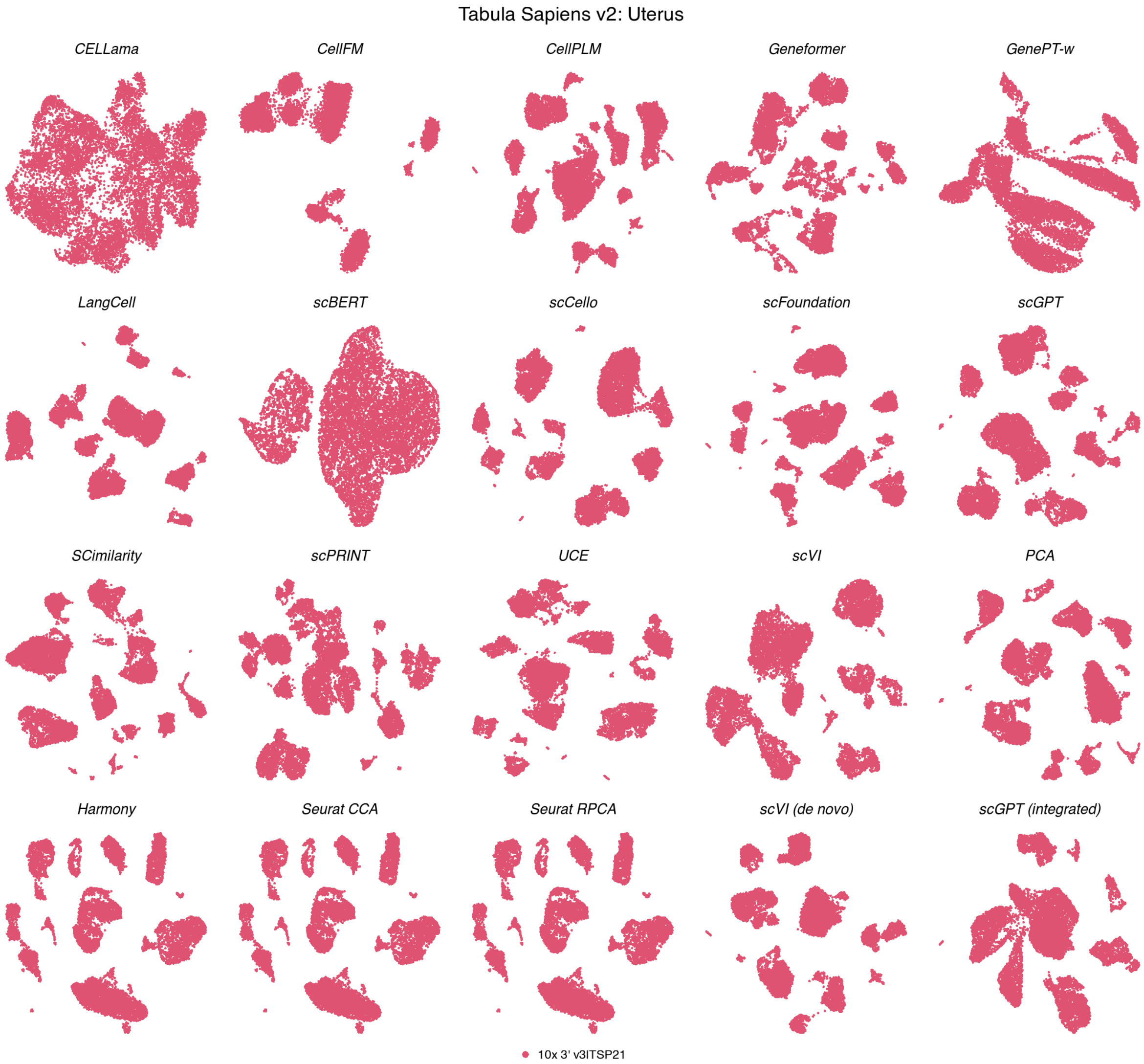

**Figure S84: Batch organization in Tabula Sapiens v2 uterus.** UMAPs show the same 10,580 cells across 20 methods, coloured by a single experimental batch. Coordinates match the corresponding cell-type view; display conventions are as in Fig. S12.

**Tabula Sapiens v2 — Uterus**

| method | Biological conservation metrics | | | | | | | | | Batch mixing metrics | | | Aggregated scores | | | | |
|---|---|---|---|---|---|---|---|---|---|---|---|---|---|---|---|---|---|
| | ARI | NMI | HOM | COM | FMI | ASW | Acc@kNN | cLISI | GC | kBET | BRAS | CiLISI | Cluster concordance | Continuous separation | Local structure | Batch mixing | Overall |
| SCimilarity | 0.72 | 0.80 | 0.86 | 0.75 | 0.78 | 0.41 | 0.93 | 1.00 | 0.84 | nan | nan | nan | 0.78 | 0.41 | 0.92 | | 0.79 |
| scVI (de novo) | 0.70 | 0.79 | 0.85 | 0.74 | 0.76 | 0.07 | 0.92 | 1.00 | 0.88 | nan | nan | nan | 0.77 | 0.07 | 0.93 | | 0.75 |
| scGPT | 0.58 | 0.78 | 0.88 | 0.70 | 0.67 | 0.14 | 0.92 | 1.00 | 0.83 | nan | nan | nan | 0.72 | 0.14 | 0.92 | | 0.73 |
| scGPT (integrated) | 0.48 | 0.75 | 0.86 | 0.66 | 0.58 | 0.29 | 0.92 | 1.00 | 0.85 | nan | nan | nan | 0.67 | 0.29 | 0.92 | | 0.73 |
| scFoundation | 0.52 | 0.76 | 0.86 | 0.68 | 0.62 | 0.17 | 0.93 | 1.00 | 0.84 | nan | nan | nan | 0.69 | 0.17 | 0.92 | | 0.72 |
| CellPLM | 0.49 | 0.75 | 0.87 | 0.67 | 0.59 | 0.23 | 0.92 | 1.00 | 0.84 | nan | nan | nan | 0.67 | 0.23 | 0.92 | | 0.72 |
| Harmony | 0.49 | 0.75 | 0.86 | 0.66 | 0.60 | 0.24 | 0.92 | 1.00 | 0.82 | nan | nan | nan | 0.67 | 0.24 | 0.91 | | 0.72 |
| Seurat CCA | 0.49 | 0.75 | 0.86 | 0.66 | 0.60 | 0.24 | 0.92 | 1.00 | 0.82 | nan | nan | nan | 0.67 | 0.24 | 0.91 | | 0.72 |
| Seurat RPCA | 0.49 | 0.75 | 0.86 | 0.66 | 0.60 | 0.24 | 0.92 | 1.00 | 0.82 | nan | nan | nan | 0.67 | 0.24 | 0.91 | | 0.72 |
| UCE | 0.52 | 0.75 | 0.86 | 0.67 | 0.62 | 0.17 | 0.92 | 1.00 | 0.82 | nan | nan | nan | 0.69 | 0.17 | 0.91 | | 0.72 |
| scVI | 0.49 | 0.74 | 0.86 | 0.66 | 0.60 | 0.19 | 0.92 | 1.00 | 0.84 | nan | nan | nan | 0.67 | 0.19 | 0.92 | | 0.71 |
| PCA | 0.47 | 0.74 | 0.86 | 0.66 | 0.57 | 0.22 | 0.92 | 1.00 | 0.83 | nan | nan | nan | 0.66 | 0.22 | 0.92 | | 0.71 |
| LangCell | 0.50 | 0.75 | 0.86 | 0.66 | 0.60 | 0.12 | 0.93 | 1.00 | 0.81 | nan | nan | nan | 0.67 | 0.12 | 0.91 | | 0.70 |
| scCello | 0.41 | 0.72 | 0.86 | 0.62 | 0.53 | 0.22 | 0.91 | 1.00 | 0.83 | nan | nan | nan | 0.63 | 0.22 | 0.91 | | 0.70 |
| scPRINT | 0.50 | 0.74 | 0.85 | 0.65 | 0.60 | 0.09 | 0.92 | 1.00 | 0.81 | nan | nan | nan | 0.67 | 0.09 | 0.91 | | 0.70 |
| Geneformer | 0.50 | 0.69 | 0.79 | 0.61 | 0.59 | 0.03 | 0.92 | 1.00 | 0.77 | nan | nan | nan | 0.64 | 0.03 | 0.90 | | 0.67 |
| CellFM | 0.42 | 0.72 | 0.84 | 0.63 | 0.53 | 0.01 | 0.87 | 0.99 | 0.38 | nan | nan | nan | 0.63 | 0.01 | 0.75 | | 0.60 |
| CELLama | 0.33 | 0.57 | 0.68 | 0.49 | 0.45 | -0.04 | 0.86 | 0.99 | 0.66 | nan | nan | nan | 0.50 | -0.04 | 0.84 | | 0.57 |
| GenePT-w | 0.08 | 0.25 | 0.31 | 0.21 | 0.19 | -0.40 | 0.84 | 0.97 | 0.52 | nan | nan | nan | 0.21 | -0.40 | 0.78 | | 0.37 |
| scBERT | 0.02 | 0.07 | 0.09 | 0.06 | 0.13 | -0.44 | 0.47 | 0.88 | 0.26 | nan | nan | nan | 0.07 | -0.44 | 0.54 | | 0.21 |

**Figure S85: Representation performance in Tabula Sapiens v2 uterus.** Scores are shown for 10,580 cells, 18 annotated cell types and a single experimental batch. Metrics, family weighting and row ordering are as in Fig. S13. Batch-mixing metrics were unavailable, and the remaining family weights were renormalized.

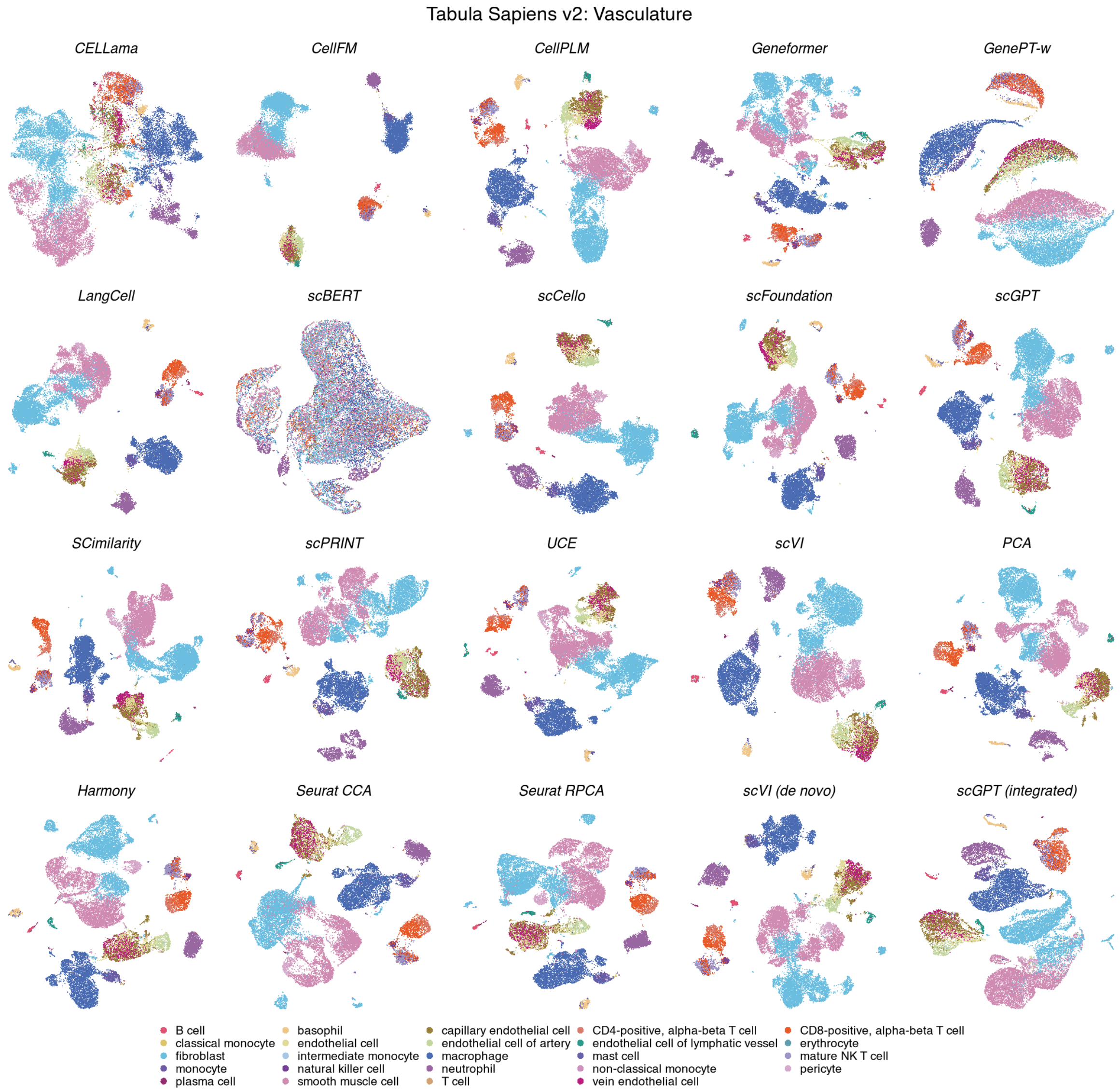

**Figure S86: Cell-type organization in Tabula Sapiens v2 vasculature.** UMAPs show the same 23,569 cells across 20 methods, coloured by 24 annotated cell types. Display conventions and method categories are as in Fig. S11.

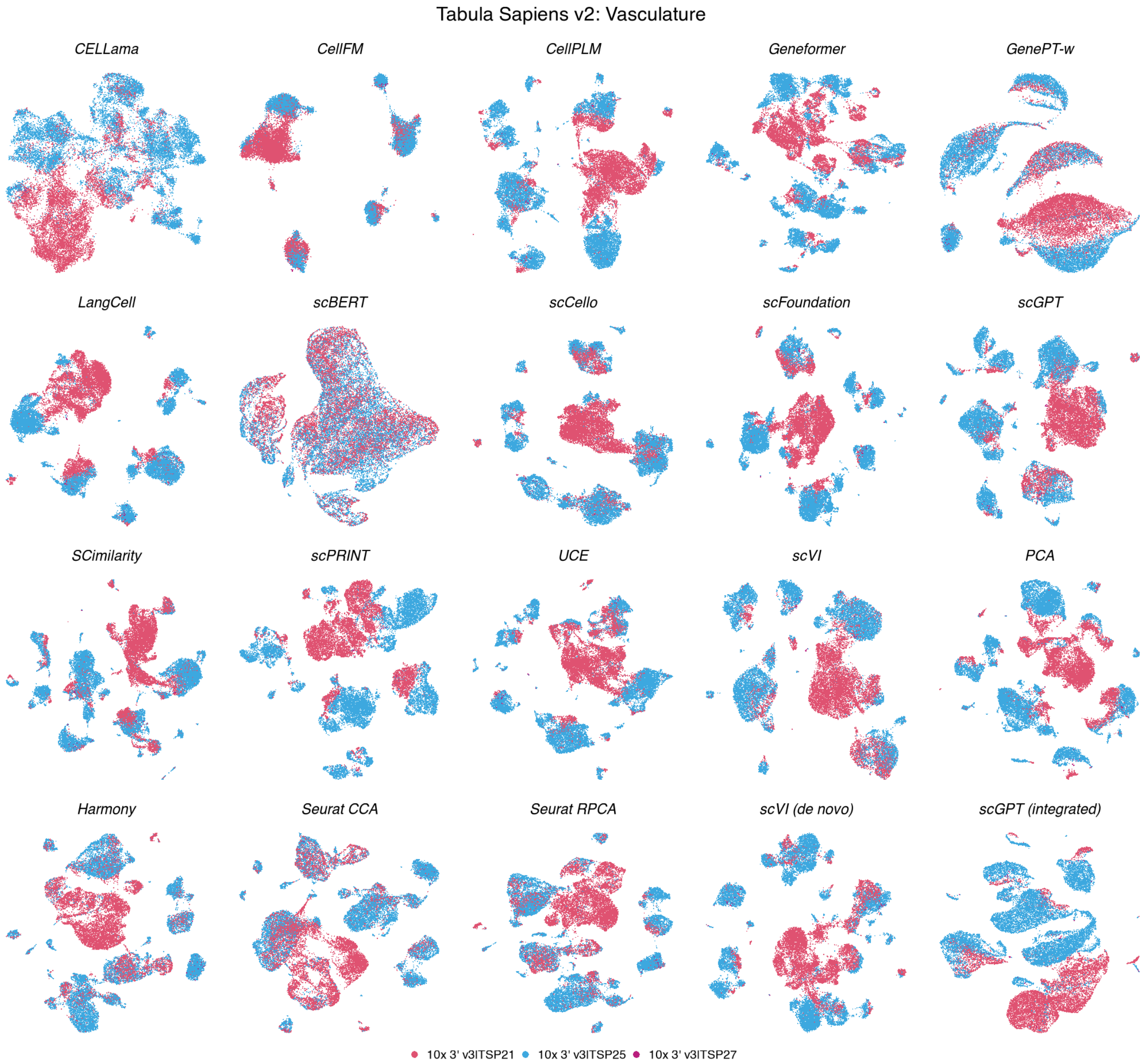

**Figure S87: Batch organization in Tabula Sapiens v2 vasculature.** UMAPs show the same 23,569 cells across 20 methods, coloured by 3 experimental batches. Coordinates match the corresponding cell-type view; display conventions are as in Fig. S12.

##### Tabula Sapiens v2 — Vasculature

| | Biological conservation metrics | | | | | | | | | Batch mixing metrics | | | Aggregated scores | | | | |
|---|---|---|---|---|---|---|---|---|---|---|---|---|---|---|---|---|---|
| method | ARI | NMI | HOM | COM | FMI | ASW | Acc@kNN | cLISI | GC | kBET | BRAS | CiLISI | Cluster concordance | Continuous separation | Local structure | Batch mixing | Overall |
| SCimilarity | 0.61 | 0.77 | 0.83 | 0.71 | 0.68 | 0.30 | 0.94 | 1.00 | 0.87 | 0.50 | 0.59 | 0.05 | 0.72 | 0.30 | 0.94 | 0.38 | 0.66 |
| scGPT | 0.57 | 0.75 | 0.83 | 0.69 | 0.65 | 0.14 | 0.92 | 1.00 | 0.87 | 0.47 | 0.63 | 0.05 | 0.70 | 0.14 | 0.93 | 0.38 | 0.64 |
| scFoundation | 0.55 | 0.75 | 0.83 | 0.68 | 0.63 | 0.17 | 0.94 | 1.00 | 0.86 | 0.47 | 0.62 | 0.03 | 0.69 | 0.17 | 0.93 | 0.38 | 0.64 |
| scVI | 0.57 | 0.75 | 0.83 | 0.69 | 0.65 | 0.15 | 0.91 | 1.00 | 0.87 | 0.49 | 0.57 | 0.05 | 0.70 | 0.15 | 0.93 | 0.37 | 0.64 |
| UCE | 0.54 | 0.75 | 0.82 | 0.69 | 0.62 | 0.19 | 0.92 | 1.00 | 0.80 | 0.50 | 0.66 | 0.04 | 0.68 | 0.19 | 0.91 | 0.40 | 0.64 |
| scCello | 0.49 | 0.74 | 0.82 | 0.67 | 0.58 | 0.21 | 0.91 | 1.00 | 0.82 | 0.51 | 0.68 | 0.05 | 0.66 | 0.21 | 0.91 | 0.41 | 0.63 |
| scVI (de novo) | 0.51 | 0.74 | 0.83 | 0.67 | 0.59 | 0.05 | 0.93 | 1.00 | 0.87 | 0.54 | 0.69 | 0.07 | 0.67 | 0.05 | 0.93 | 0.43 | 0.63 |
| scGPT (integrated) | 0.47 | 0.73 | 0.82 | 0.66 | 0.56 | 0.14 | 0.91 | 1.00 | 0.86 | 0.55 | 0.62 | 0.07 | 0.65 | 0.14 | 0.92 | 0.41 | 0.63 |
| Seurat CCA | 0.48 | 0.70 | 0.80 | 0.63 | 0.57 | 0.12 | 0.90 | 1.00 | 0.84 | 0.59 | 0.64 | 0.13 | 0.64 | 0.12 | 0.91 | 0.46 | 0.63 |
| LangCell | 0.55 | 0.75 | 0.81 | 0.69 | 0.62 | 0.09 | 0.92 | 1.00 | 0.84 | 0.45 | 0.71 | 0.03 | 0.68 | 0.09 | 0.92 | 0.40 | 0.63 |
| CellPLM | 0.52 | 0.74 | 0.83 | 0.67 | 0.60 | 0.17 | 0.93 | 1.00 | 0.89 | 0.43 | 0.56 | 0.03 | 0.67 | 0.17 | 0.94 | 0.34 | 0.63 |
| Seurat RPCA | 0.48 | 0.73 | 0.83 | 0.65 | 0.57 | 0.12 | 0.91 | 1.00 | 0.84 | 0.53 | 0.62 | 0.09 | 0.65 | 0.12 | 0.92 | 0.41 | 0.63 |
| Harmony | 0.46 | 0.72 | 0.82 | 0.64 | 0.55 | 0.12 | 0.91 | 1.00 | 0.82 | 0.58 | 0.61 | 0.08 | 0.64 | 0.12 | 0.91 | 0.42 | 0.62 |
| PCA | 0.48 | 0.73 | 0.82 | 0.65 | 0.57 | 0.03 | 0.93 | 1.00 | 0.88 | 0.46 | 0.59 | 0.03 | 0.65 | 0.03 | 0.94 | 0.36 | 0.61 |
| scPRINT | 0.42 | 0.69 | 0.80 | 0.61 | 0.52 | 0.12 | 0.91 | 1.00 | 0.86 | 0.48 | 0.64 | 0.04 | 0.61 | 0.12 | 0.93 | 0.39 | 0.61 |
| CellFM | 0.51 | 0.72 | 0.78 | 0.66 | 0.59 | 0.00 | 0.85 | 0.99 | 0.39 | 0.47 | 0.95 | 0.08 | 0.65 | 0.00 | 0.74 | 0.50 | 0.58 |
| Geneformer | 0.32 | 0.63 | 0.72 | 0.56 | 0.42 | 0.05 | 0.93 | 1.00 | 0.82 | 0.44 | 0.69 | 0.04 | 0.53 | 0.05 | 0.92 | 0.39 | 0.57 |
| CELLama | 0.31 | 0.59 | 0.69 | 0.51 | 0.42 | 0.02 | 0.86 | 1.00 | 0.64 | 0.56 | 0.76 | 0.06 | 0.50 | 0.02 | 0.83 | 0.46 | 0.55 |
| GenePT-w | 0.14 | 0.38 | 0.45 | 0.33 | 0.24 | -0.46 | 0.82 | 0.99 | 0.45 | 0.47 | 0.67 | 0.11 | 0.31 | -0.46 | 0.75 | 0.42 | 0.40 |
| scBERT | 0.03 | 0.11 | 0.13 | 0.09 | 0.12 | -0.38 | 0.47 | 0.92 | 0.26 | 0.25 | 0.56 | 0.16 | 0.10 | -0.38 | 0.55 | 0.32 | 0.25 |

**Figure S88: Representation performance in Tabula Sapiens v2 vasculature.** Scores are shown for 23,569 cells, 24 annotated cell types and 3 experimental batches. Metrics, family weighting and row ordering are as in Fig. S13.

#### 2.2 Cross-release robustness of the cell-representation benchmark

The biological-conservation hierarchy was strongly concordant between TSv1 and TSv2 across the fixed 20-representation panel (Spearman $\rho = 0.884$, $P = 2.32 \times 10^{-7}$; Kendall $\tau = 0.779$, $P = 5.84 \times 10^{-8}$; Fig. S89). The same ordering was recovered for the biological-plus-batch aggregate ($\rho = 0.913$; $\tau = 0.779$), for which the three highest-ranked methods were identical across releases. None of the 19 non-PCA methods showed a Holm-significant cross-release change in aggregate PCA-relative rank.

The metric-level analyses gave the same broad result while separating changes in absolute difference from changes in ratio. For the primary raw-score difference from PCA, 167 of 171 biological method–metric comparisons showed no Holm-significant cross-release difference; the four supported changes were lower TSv2 values for CellFM graph connectivity and for scBERT ASW, FMI and graph connectivity. For the supplementary method-to-PCA ratio, 169 of 171 comparisons showed no Holm-significant cross-release difference; Geneformer ASW was higher and scBERT graph connectivity was lower in TSv2 (Fig. S90). Differences in the biological-plus-batch analysis were concentrated in BRAS and CiLISI and generally favoured TSv2. The matched TSv2 tissues typically contained fewer batches than their TSv1 counterparts, consistent with the greater cross-release sensitivity of batch-mixing metrics than of biological conservation.

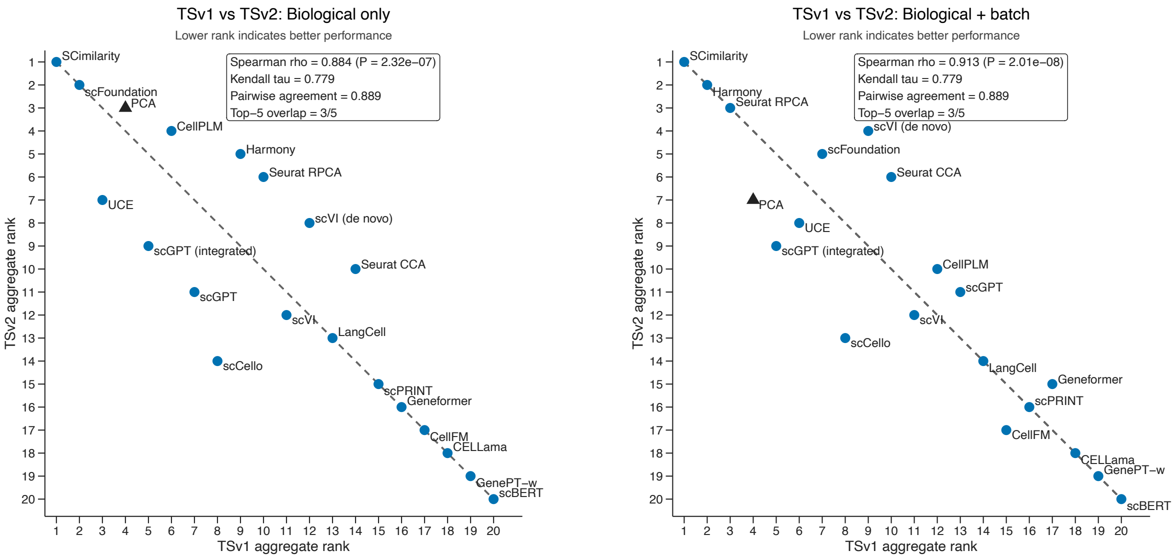

**Figure S89: Concordance of method rankings between Tabula Sapiens releases.** Points show the atlas-wide rank of each of the 20 methods in TSv1 and TSv2 using biological-conservation metrics alone (left) or the biological-plus-batch aggregate (right); lower ranks indicate stronger performance. The dashed diagonal denotes identical rank positions.

#### 2.3 Representation geometry and biological performance

The organization of the emitted embeddings separated the strongest and weakest representations more clearly than broad model-design categories. Greater spectral anisotropy was associated with poorer biological conservation across the 20 methods (Spearman $\rho = -0.591$, nominal $P = 0.00607$), whereas a larger within-minus-between cell-type anisotropy contrast was associated with better performance ($\rho = 0.687$, nominal $P = 8.15 \times 10^{-4}$; Fig. S91). PCA and the established target-data methods had low global anisotropy. Among frozen scFMs, SCimilarity showed the strongest separation between within- and between-cell-type directional coherence, whereas GenePT-w, Geneformer, scBERT and scPRINT showed pronounced global

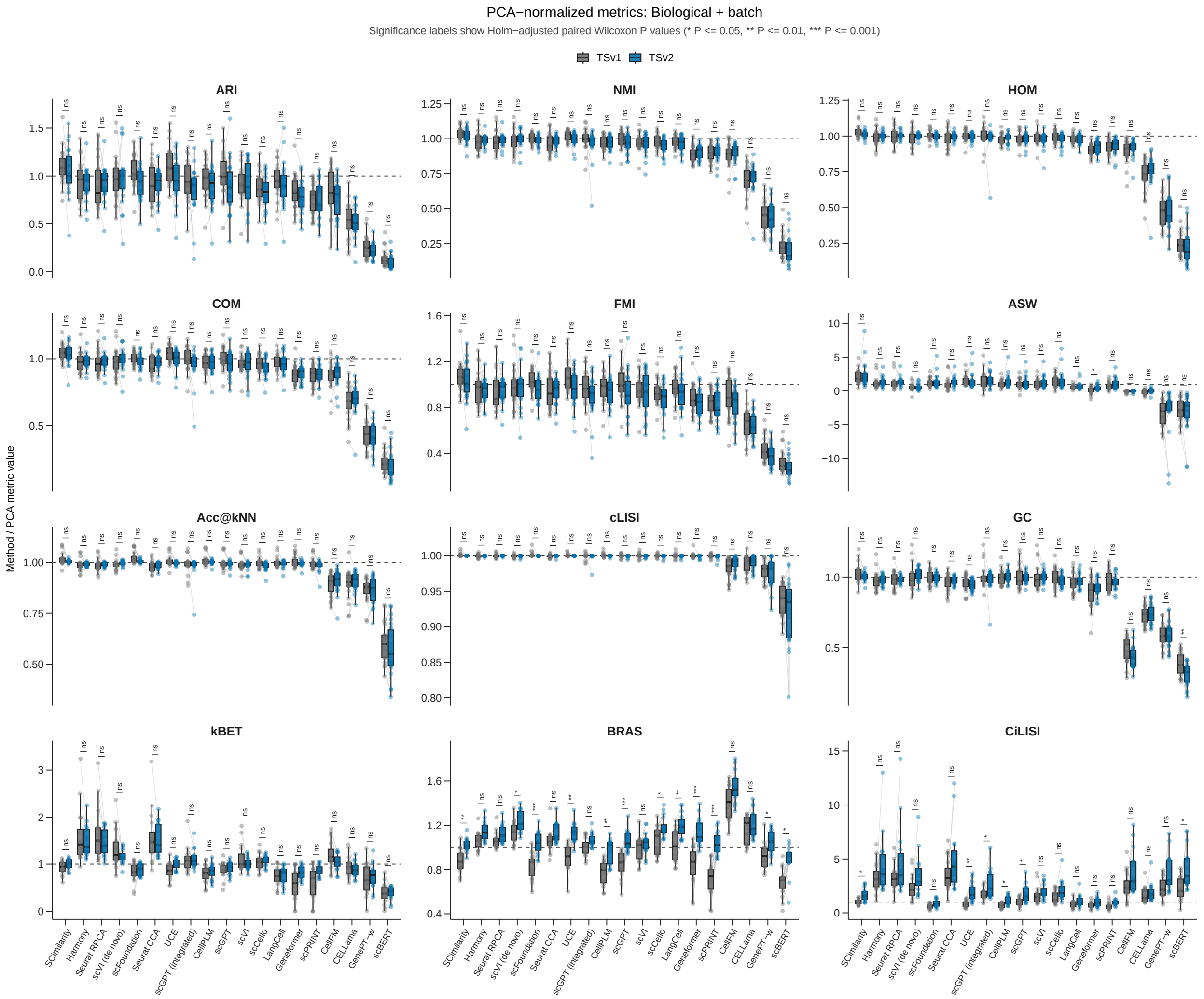

**Figure S90: PCA-normalized performance across Tabula Sapiens releases.** Panels show nine biological-conservation and three batch-mixing metrics for the 19 non-PCA methods. Values are method scores divided by the corresponding PCA score, and lines pair the same tissue between TSv1 and TSv2. Biological metrics use up to 22 shared tissues; batch metrics use the 16 shared tissues that were multi-batch in both releases. Symbols denote two-sided paired Wilcoxon tests with Holm correction across methods within each metric.

concentration.

Local expression-neighbourhood preservation provided the strongest performance association. Across all 20 representations, mean chance-corrected $R_{NX}$ was associated with the biological-conservation outcome excluding Acc@kNN, cLISI and graph connectivity ($\rho = 0.798$, nominal $P = 2.44 \times 10^{-5}$); the association was stronger among the 13 frozen scFMs ($\rho = 0.901$, nominal $P = 2.61 \times 10^{-5}$). Established methods, including PCA, Harmony and scVI trained de novo, retained comparatively high local fidelity. scFoundation and SCimilarity were among the strongest frozen representations, whereas scBERT, GenePT-w and CellFM substantially altered local expression structure.

Effective dimensionality identified distinct geometric failure modes (Fig. S92). GenePT-w and scBERT concentrated their standardized variance into approximately one to two effective dimensions (mean participation ratios 1.36 and 1.99). CellFM occupied the opposite extreme, with a mean participation ratio of approximately 357 and a raw-space TwoNN estimate of approximately 159, but this high-dimensional variation did not resolve correspondingly strong biological organization. Low effective rank and high-dimensional but weakly organized variation therefore represent different warning signs of embedding failure.

Cell-type partial $\eta^2$ was highest for SCimilarity (mean approximately 0.535) and lower for CellFM and scBERT (approximately 0.074 and 0.166, respectively). CellFM also had low batch partial $\eta^2$ (0.0065), showing that weak batch association alone did not imply preservation of biological structure. Cell-type and batch-associated organization were therefore evaluated as separate properties.

Broad model descriptors were less informative (Fig. S93). Reported pretraining corpus size ($\rho = -0.237$, permutation $P = 0.480$) and runtime parameter count ($\rho = 0.392$, $P = 0.211$) were not associated with performance. Model family explained a moderate descriptive fraction of between-model variation ($R^2 = 0.374$, exact $P = 0.095$), whereas retention of expression magnitude explained little ($R^2 = 0.018$, exact $P = 0.653$). None of the four model-characteristic tests remained supported after Holm correction (all adjusted $P \geq 0.379$).

##### 2.4 Dataset-level moderators and shared failures in rare populations

None of the six dataset properties was associated with the scFM–PCA performance difference in any of the three biological-score families after joint adjustment (all 18 Holm-adjusted block-permutation $P \geq 0.617$; Fig. S94a). For example, the mutually adjusted $\log_{10}$ cell-count slopes were 0.0134 for cluster–label concordance (95% CI, −0.0056 to 0.0401), 0.0251 for ASW (−0.0184 to 0.0692), and 0.0009 for local-neighbourhood structure (−0.0077 to 0.0114). Cell-type count, normalized cell-type entropy, median detected genes, rare-cell burden and reference-expression effective rank likewise showed no supported association.

Rare populations nevertheless formed a recurrent cell-level failure mode. In the complete 20-representation readout, none of the 19 non-PCA representations significantly outperformed PCA for macro-averaged rare-type accuracy after Holm correction. Among the frozen scFMs, SCimilarity, scFoundation, UCE and CellPLM did not differ significantly from PCA, whereas the other nine had lower rare-type accuracy (Fig. S94b).

Of 548,977 TSv2 cells, 548,826 supported the complete label-out-of-fold readout. At the threshold of at least 16 incorrect readouts in the 20-method panel, the tissue-equal consensus-hard fraction was 18.7% for cells from rare types, 5.39% for intermediate types and 0.684% for common types (Fig. S94c). In the complementary PCA-plus-13-scFM analysis, 15,050 cells met the corresponding threshold of at least 12 of 14 incorrect readouts. Rare cells comprised 3.2% of all evaluable cells but 19.5% of consensus-hard cells. The adjusted odds ratio was 57.21 for rare versus common cells (95% CI, 28.42–115.18; Holm-adjusted

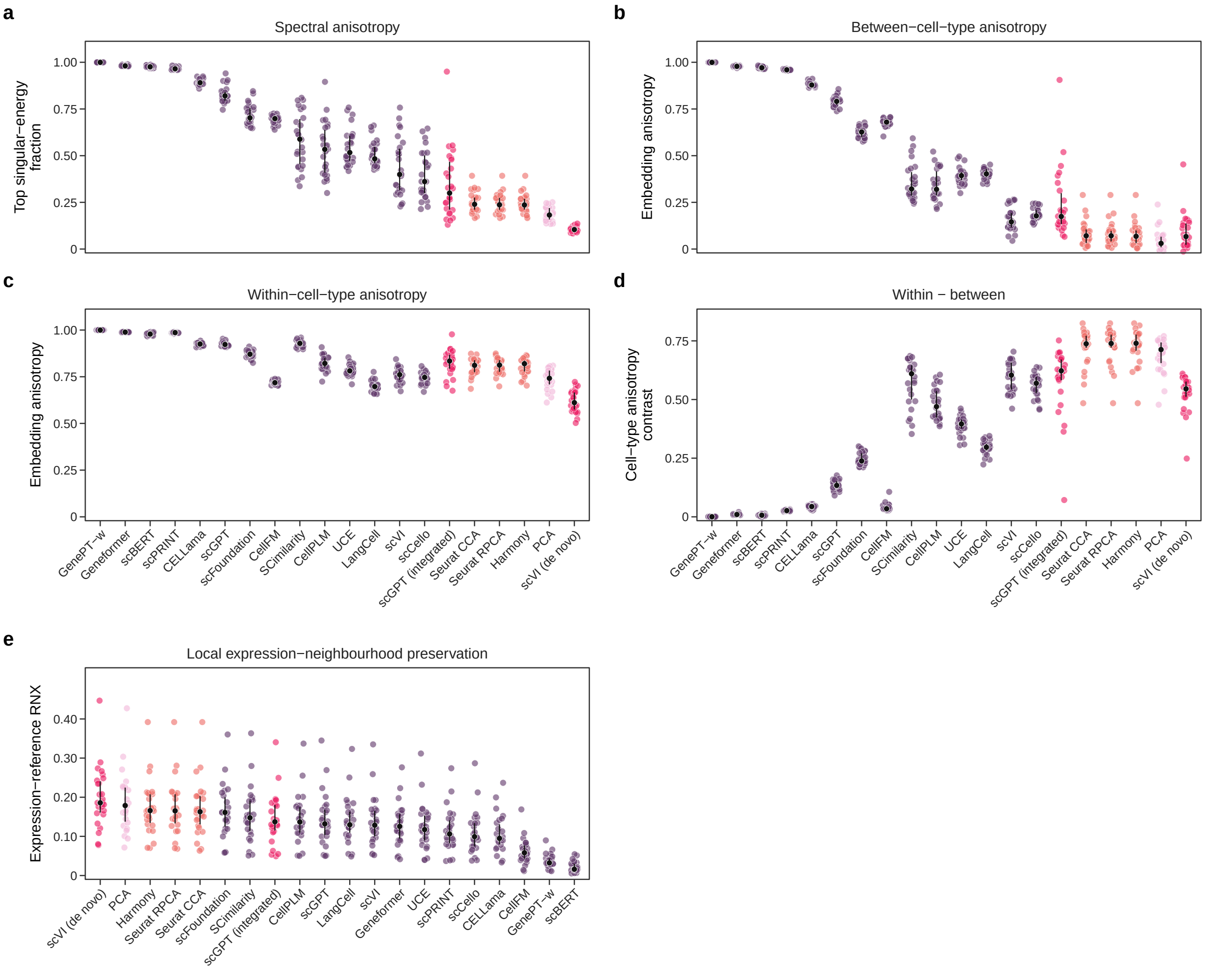

**Figure S91: Embedding anisotropy and local expression-neighbourhood preservation. a–d,** Spectral, between-cell-type and within-cell-type anisotropy and their within-minus-between contrast across TSv2 tissues. **e,** Mean chance-corrected $R_{NX}$ against biological-conservation performance after excluding local-neighbourhood endpoints. Points in **a–d** are tissues; black markers and lines show medians and interquartile ranges.

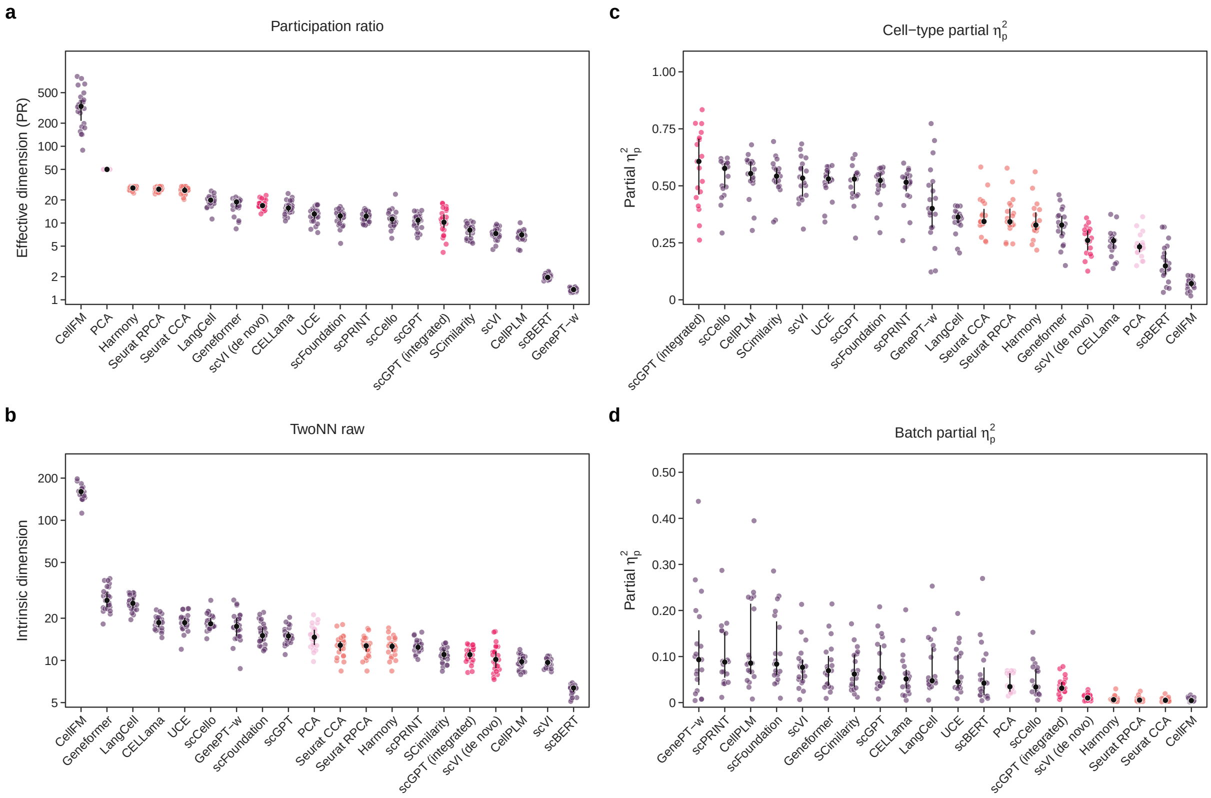

**Figure S92: Complementary embedding-geometry diagnostics. a,b,** Participation ratio and TwoNN intrinsic dimension. **c,d,** Cell-type- and batch-associated partial $\eta^2$ across multi-batch tissues. Points are tissues; black markers and lines show medians and interquartile ranges.

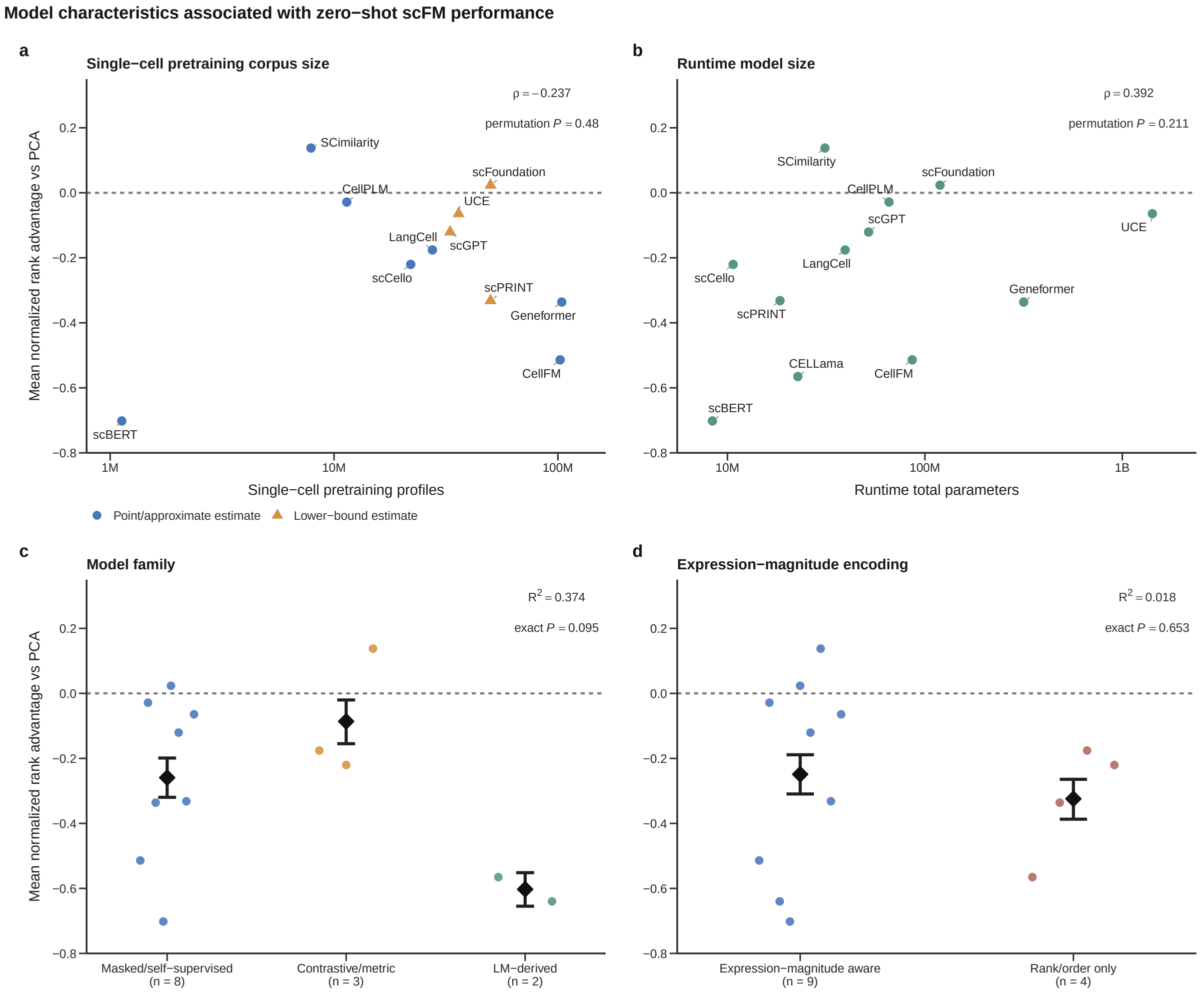

**Figure S93: Model characteristics and zero-shot biological performance. a,b,** Associations with reported pretraining corpus size and runtime parameter count. **c,d,** Performance grouped by model family and retention of expression magnitude. The dashed line denotes parity with PCA.

$P = 1.78 \times 10^{-29}$) and 8.13 for intermediate versus common cells (4.05–16.32; $P = 3.81 \times 10^{-9}$; Fig. S94d).

#### 2.5 Trajectory inference across frozen and target-fitted representations

Method rankings varied across the four inference backends, although their mean ranks remained positively correlated (pairwise Spearman $\rho = 0.471$–$0.814$). After averaging backends within each dataset and then the nine datasets, scFoundation, CellPLM and PCA had the three lowest mean aggregate ranks (6.52, 7.21 and 7.27, respectively). The 17 rank profiles differed overall (Friedman $\chi^2 = 49.89$, d.f. 16; blockwise method-label permutation $P = 1.00 \times 10^{-5}$; Kendall's $W = 0.346$), but none of the 16 paired comparisons with PCA remained significant after Holm correction. A common-denominator analysis using the same four metric families for every backend gave a similar ordering, with scFoundation, PCA and CellPLM ranked first (6.07), second (6.71) and third (6.79).

The canonical and backend-specific summaries are reported in Figs. S95–S100. Complete cross-backend trajectory maps and dataset–backend metric-score panels for all nine developmental systems follow in Figs. S101–S145; dataset-specific pseudotime ribbons are shown in Figs. S146–S154.

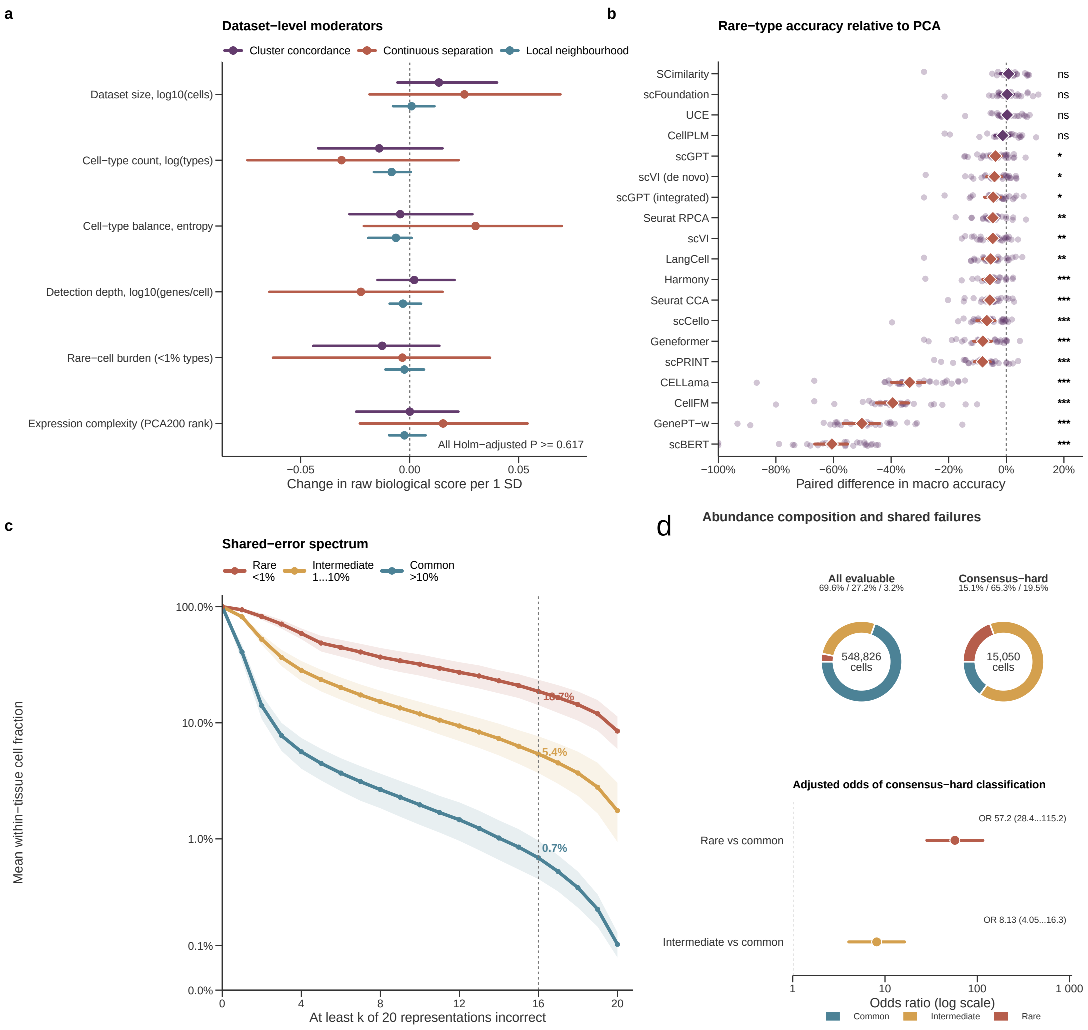

**Figure S94: Dataset-level moderation and shared failures in rare populations. a,** Mutually adjusted associations between six standardized dataset properties and paired raw biological-family score differences (scFM minus PCA). Points and intervals show coefficients and 95% confidence intervals from 5,000 tissue-block bootstrap resamples; Freedman–Lane tissue-block permutation $P$ values were Holm-adjusted across the 18 family–moderator tests. **b,** Paired differences from PCA in rare-type macro accuracy for the 19 non-PCA representations. Small points denote tissues; diamonds and intervals show tissue means and 95% tissue-bootstrap confidence intervals; labels report paired Wilcoxon tests after Holm correction. **c,** Tissue-equal cumulative sharing of cell-level errors across the complete 20-readout panel. The vertical line marks the threshold of at least 16 incorrect readouts. **d,** Pooled composition of all 548,826 evaluable cells and the 15,050 consensus-hard cells by within-tissue abundance, using PCA and the 13 frozen scFMs and a threshold of at least 12 of 14 incorrect readouts. Points and intervals below the rings show adjusted odds ratios and 95% confidence intervals. Cell types represented by fewer than five cells were excluded.

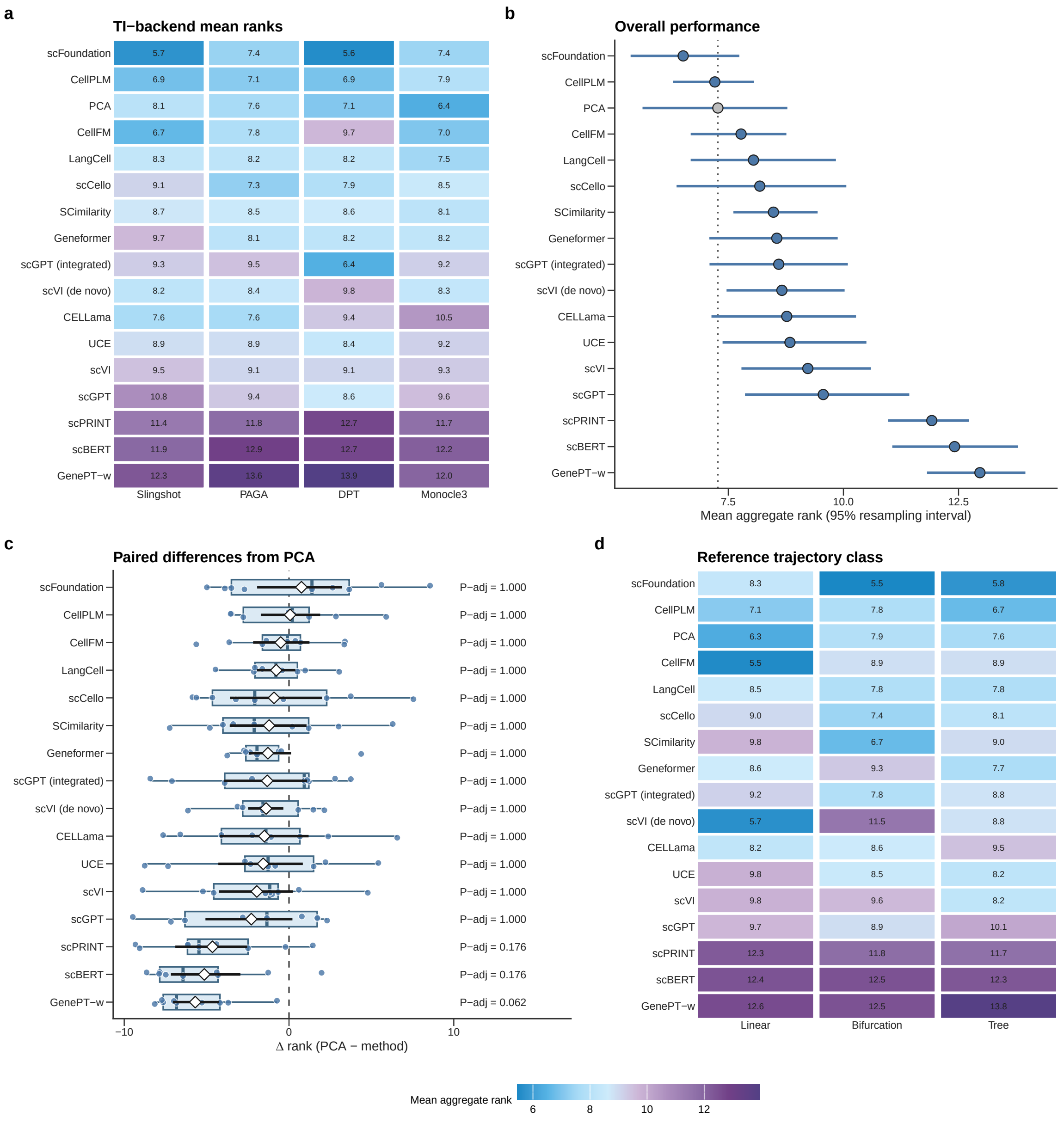

**Figure S95: Canonical trajectory-inference rank summary. a,** Mean metric-family aggregate rank for each representation and trajectory-inference backend, averaged across nine trajectory datasets; lower values indicate better performance. **b,** Overall mean aggregate rank after first averaging the four backends within each dataset. Points show dataset means and intervals show 95% percentile intervals from 10,000 trajectory-class-stratified bootstrap replicates (seed 20260723); the dotted line marks the PCA mean. **c,** Paired PCA-relative differences across datasets, defined as PCA rank minus method rank. Positive values favour the evaluated representation. Labels report two-sided exact sign-flip $P$ values after Holm correction across all 16 non-PCA comparisons. **d,** Mean aggregate ranks by reference trajectory class. The backend-applicable specification uses all metric families that each backend can estimate. DPT contributes no topology family, and its assignment family contains milestone F1 only, with weights renormalized over available families.

Local preservation of expression structure generalized to the continuous trajectory task. Across all 17 representations, mean direct-expression $R_{NX}$ was positively associated with the PCA-relative continuous outcome (Spearman $\rho = 0.775$; 95% trajectory-class-stratified bootstrap stability interval, 0.350–0.895; two-sided whole-method-label permutation $P = 4.50 \times 10^{-4}$). The association was stronger within the 13 frozen scFMs ($\rho = 0.874$; interval, 0.522–0.945; $P = 2.70 \times 10^{-4}$). It remained positive after removing the complete reference-position-recoverability family from the performance outcome (all 17, $\rho = 0.625$, $P = 0.00910$). In contrast, spectral anisotropy showed a weaker negative relationship with trajectory recovery across the 17 representations ($\rho = -0.446$, permutation $P = 0.0759$). Local preservation of expression neighbourhoods therefore provided the more consistent geometric signature of trajectory recovery.

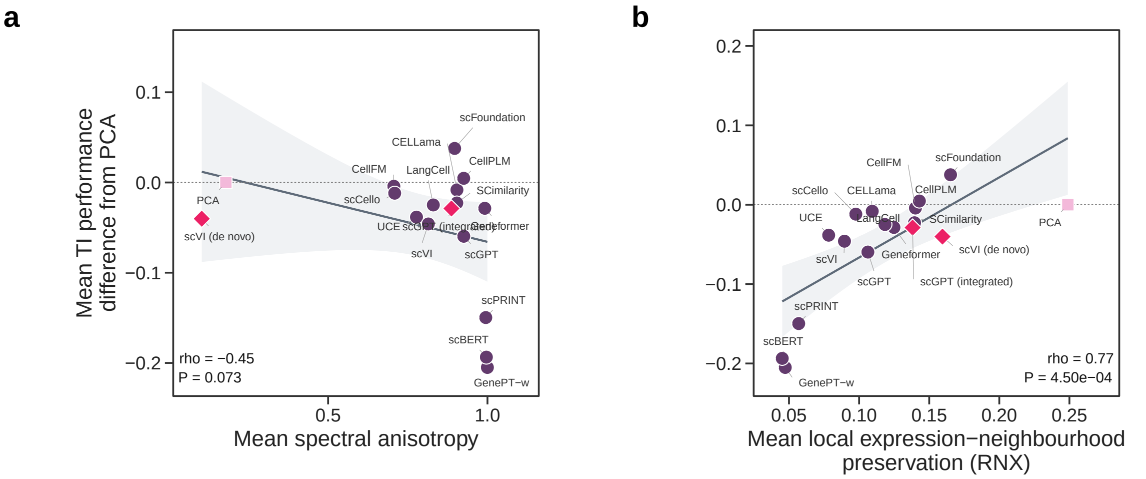

**Figure S96: Embedding geometry is associated with transfer to continuous developmental trajectories. a,** Model-level mean spectral anisotropy, defined as the fraction of raw, uncentred embedding energy in the leading singular direction, plotted against PCA-relative continuous trajectory-inference performance for 17 representations. **b,** Mean local expression-neighbourhood preservation, quantified as chance-corrected $R_{NX}$ averaged equally over $k = 5$, 10 and 15, plotted against the same outcome. Reference neighbours in **b** were computed directly in analytic-Pearson-residual gene space. Geometry and performance were averaged across the same nine developmental systems. Performance averages direction-aligned differences from PCA within metric family, across applicable families, across four inference backends and across datasets. Panel annotations report Spearman correlations and two-sided whole-method-label permutation $P$ values; intervals are from 10,000 trajectory-class-stratified dataset-bootstrap replicates. Each point is one representation.

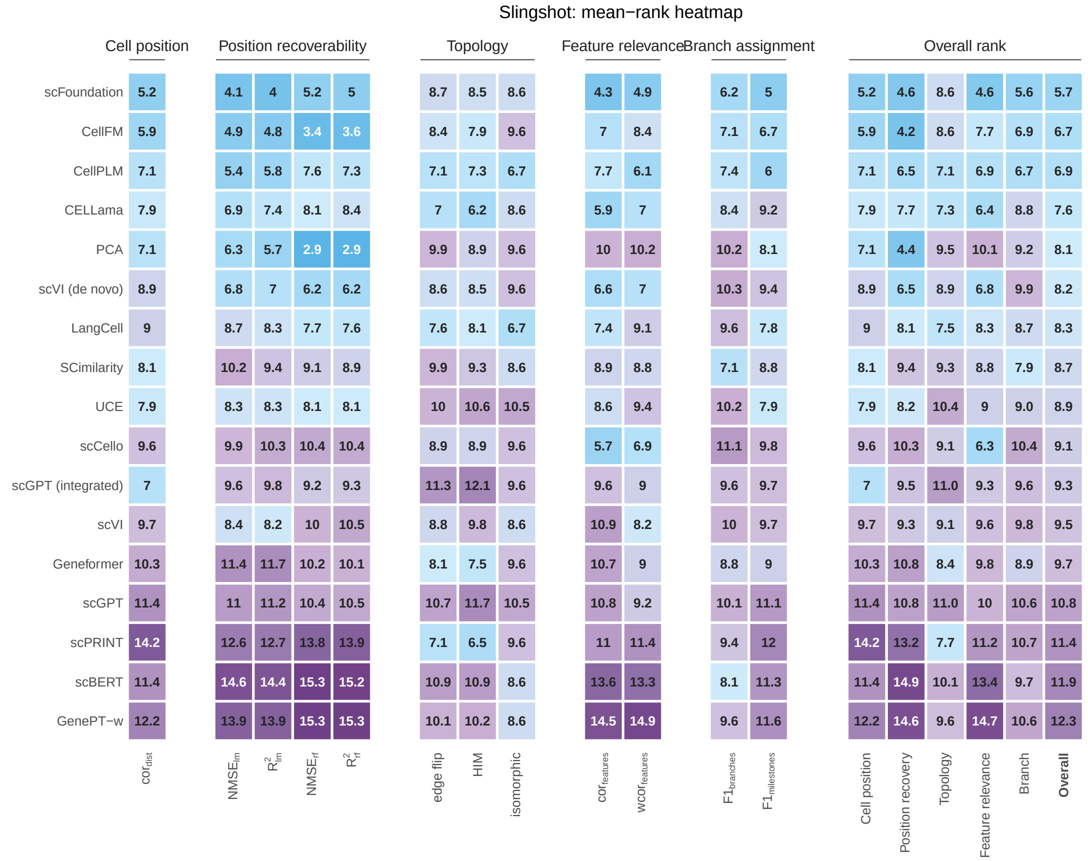

**Figure S97: Trajectory-inference mean-rank heatmap for Slingshot.** Rows show 17 representations evaluated with Slingshot. Within each dataset and backend, methods were ranked separately for each of the 12 retained `dyneval` metrics, with average ranks for ties. Metric ranks were averaged within five prespecified families: cell position, reference-position recoverability, topology, feature relevance and branch assignment. The final Overall panel reports the family ranks followed by their equal-weight aggregate; family and Overall values were not re-ranked. Rank-derived values were then averaged across nine trajectory datasets without further re-ranking. All five metric families contribute equal weight to Overall. Lower values indicate better performance, all panels use the same rank scale, and grey blanks denote structurally unavailable comparisons rather than observed zero performance.

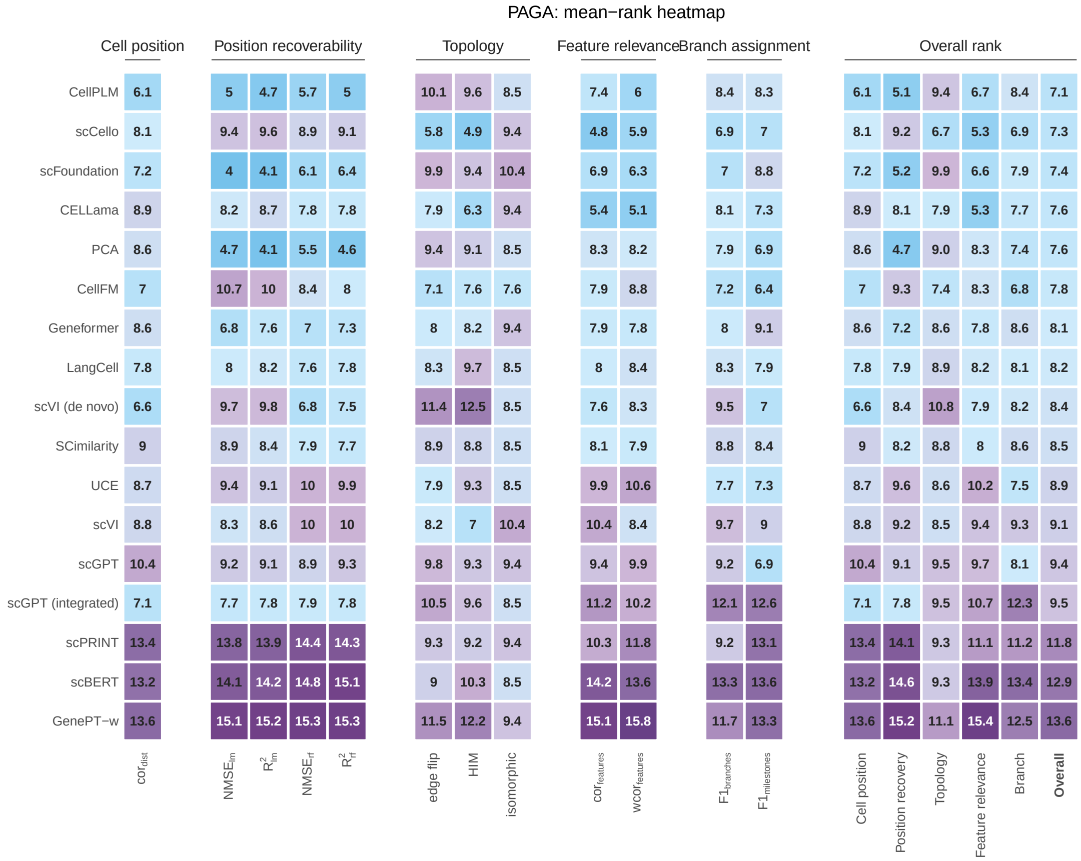

**Figure S98: Trajectory-inference mean-rank heatmap for PAGA.** Rows show 17 representations averaged across nine trajectory datasets. Ranking, family aggregation and display conventions are as in Fig. S97.

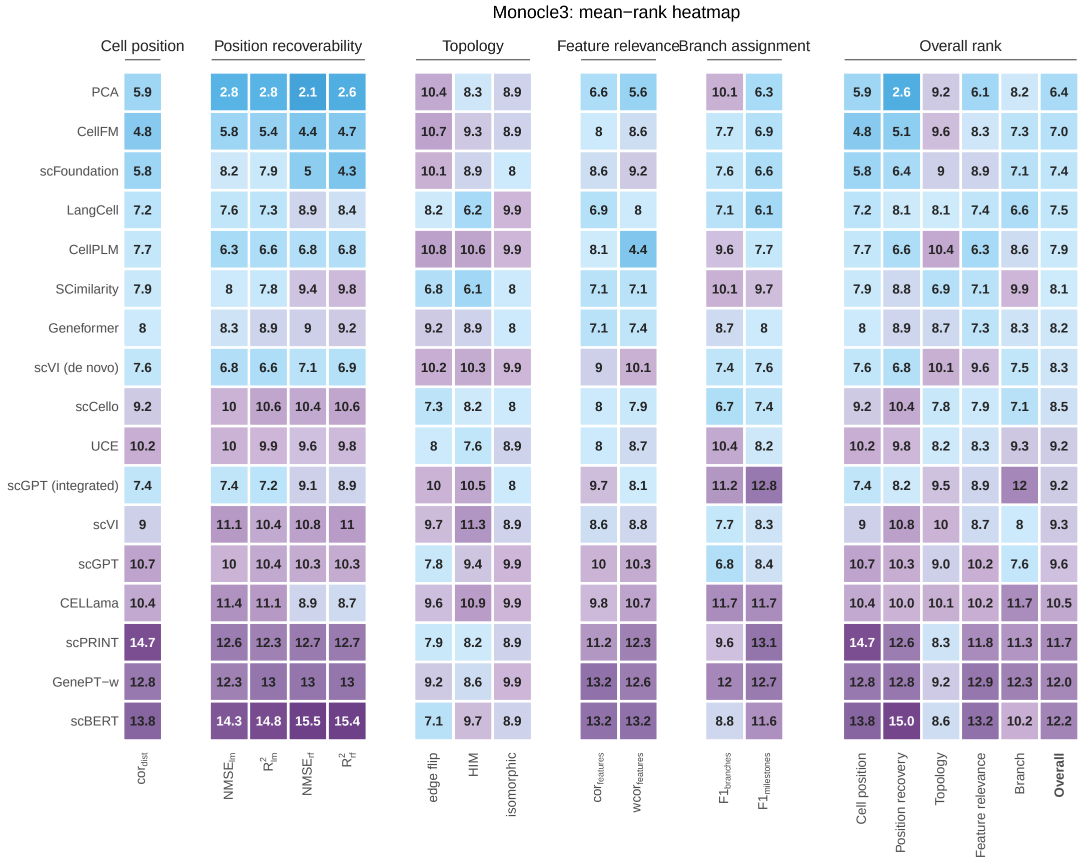

**Figure S99: Trajectory-inference mean-rank heatmap for Monocle 3.** Rows show 17 representations averaged across nine trajectory datasets. Ranking, family aggregation and display conventions are as in Fig. S97.

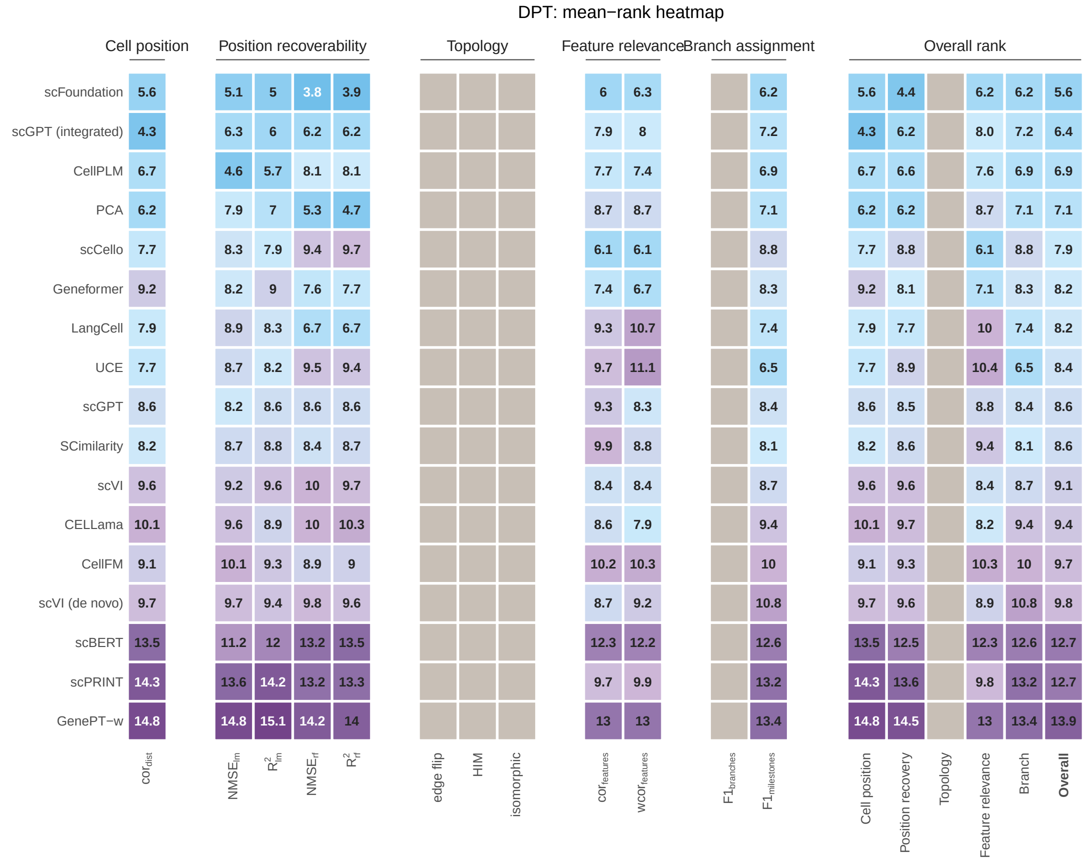

**Figure S100: Trajectory-inference mean-rank heatmap for DPT.** Rows show 17 representations averaged across nine trajectory datasets. Ranking and display conventions are as in Fig. S97. DPT contributes no topology family; branch assignment is represented by milestone F1, and weights are renormalized over the four available families.

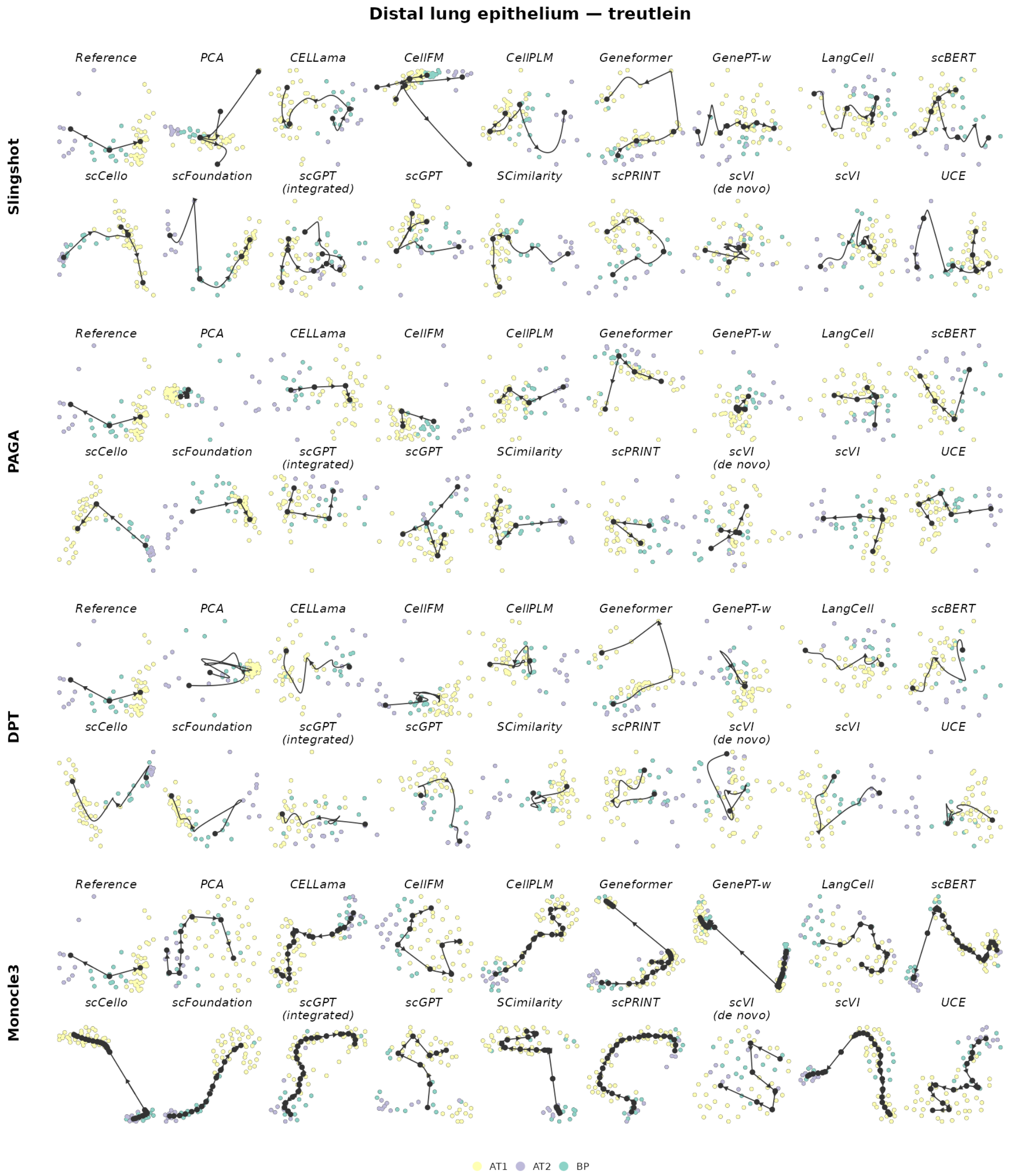

**Figure S101: Trajectory inference on distal lung epithelium.** The reference trajectory is bifurcating. Rows show Slingshot, PAGA, DPT and Monocle 3; within each row, the reference is followed by the same 17 representations. Cells are coloured by the reference milestone or state. Black nodes, edges and arrows show the inferred milestone graph and its orientation. Layouts and trajectories were reconstructed independently for each representation and backend.

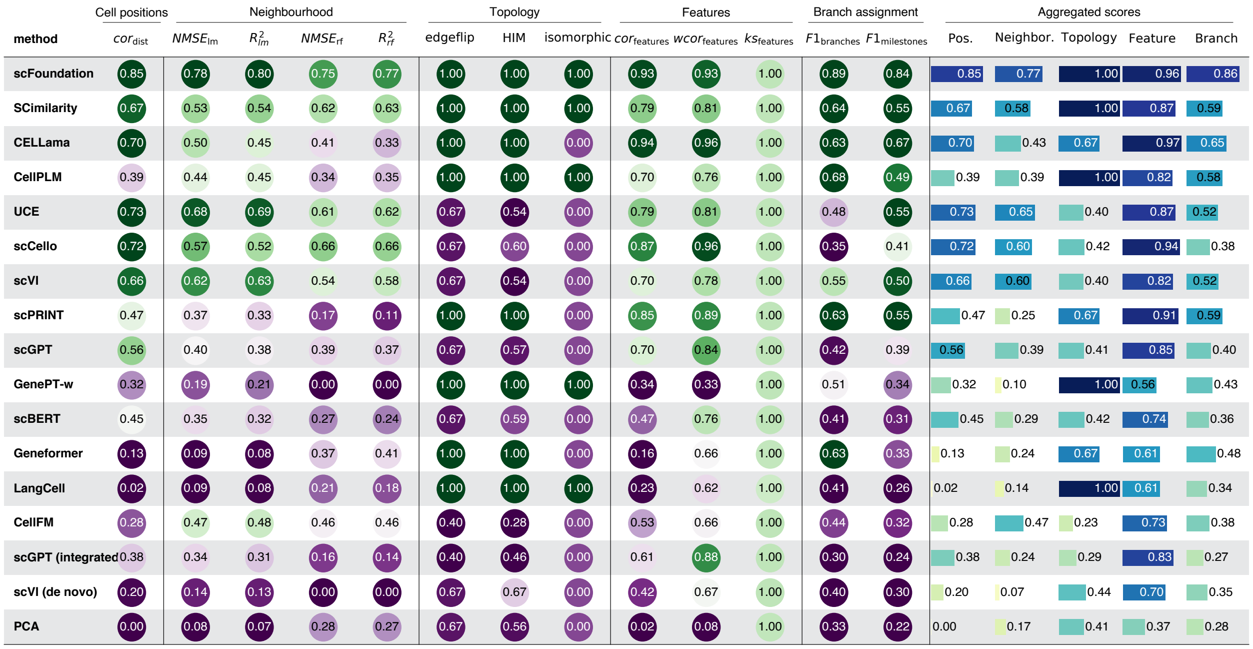

**Figure S102: Trajectory-inference metric scores for distal lung epithelium using Slingshot.** Rows show 17 representations; circles report individual `dyneval` scores grouped into cell position, reference-position recoverability, topology, feature relevance and branch assignment. Horizontal bars show the mean of the available scores within each family, and rows are ordered by the mean of the family summaries. Blank entries indicate unavailable metrics.

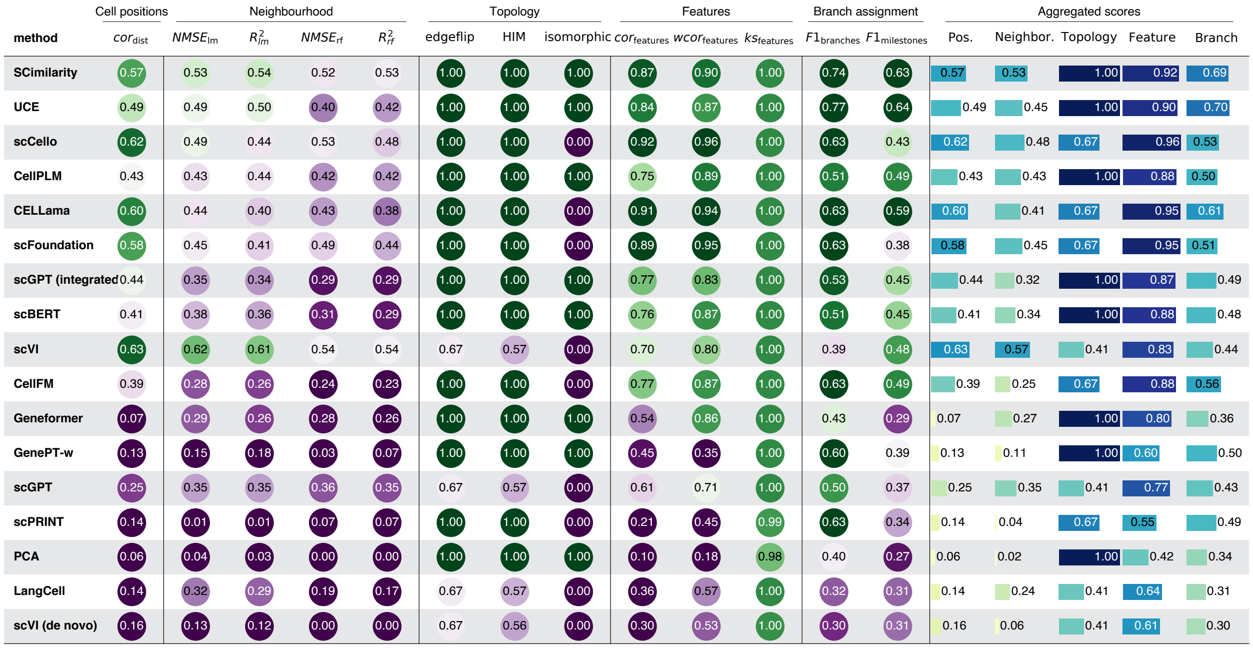

**Figure S103: Trajectory-inference metric scores for distal lung epithelium using PAGA.** Rows show 17 representations; circles report individual `dyneval` scores grouped into cell position, reference-position recoverability, topology, feature relevance and branch assignment. Horizontal bars show the mean of the available scores within each family, and rows are ordered by the mean of the family summaries. Blank entries indicate unavailable metrics.

| | Cell positions | Neighbourhood | | | | Topology | | | Features | | | Branch assignment | | Aggregated scores | | | | |
|---|---|---|---|---|---|---|---|---|---|---|---|---|---|---|---|---|---|---|
| method | $cor_{dist}$ | $NMSE_{lm}$ | $R^2_{lm}$ | $NMSE_{rf}$ | $R^2_{rf}$ | edgeflip | HIM | isomorphic | $cor_{features}$ | $wcor_{features}$ | $ks_{features}$ | $F1_{branches}$ | $F1_{milestones}$ | Pos. | Neighbor. | Topology | Feature | Branch |
| scFoundation | 0.87 | 0.81 | 0.81 | 0.83 | 0.84 | 1.00 | 1.00 | 1.00 | 0.96 | 0.97 | 1.00 | 0.77 | 0.81 | 0.87 | 0.82 | 1.00 | 0.98 | 0.79 |
| scCello | 0.72 | 0.68 | 0.65 | 0.59 | 0.55 | 1.00 | 1.00 | 1.00 | 0.93 | 0.93 | 1.00 | 0.73 | 0.69 | 0.72 | 0.62 | 1.00 | 0.95 | 0.71 |
| PCA | 0.65 | 0.37 | 0.37 | 0.31 | 0.31 | 1.00 | 1.00 | 1.00 | 0.83 | 0.85 | 1.00 | 0.79 | 0.46 | 0.65 | 0.34 | 1.00 | 0.89 | 0.63 |
| UCE | 0.63 | 0.46 | 0.42 | 0.47 | 0.47 | 1.00 | 1.00 | 0.00 | 0.92 | 0.95 | 1.00 | 0.63 | 0.55 | 0.63 | 0.45 | 0.67 | 0.96 | 0.59 |
| SCimilarity | 0.62 | 0.43 | 0.39 | 0.40 | 0.37 | 1.00 | 1.00 | 0.00 | 0.93 | 0.95 | 1.00 | 0.63 | 0.66 | 0.62 | 0.40 | 0.67 | 0.96 | 0.64 |
| scVI | 0.53 | 0.35 | 0.36 | 0.53 | 0.51 | 1.00 | 1.00 | 1.00 | 0.72 | 0.69 | 1.00 | 0.42 | 0.50 | 0.53 | 0.44 | 1.00 | 0.80 | 0.46 |
| CELLama | 0.59 | 0.43 | 0.39 | 0.45 | 0.41 | 1.00 | 1.00 | 0.00 | 0.91 | 0.93 | 1.00 | 0.63 | 0.59 | 0.59 | 0.42 | 0.67 | 0.94 | 0.61 |
| scBERT | 0.39 | 0.37 | 0.34 | 0.38 | 0.35 | 1.00 | 1.00 | 1.00 | 0.81 | 0.89 | 1.00 | 0.66 | 0.44 | 0.39 | 0.36 | 1.00 | 0.90 | 0.55 |
| scGPT (integrated | 0.37 | 0.36 | 0.33 | 0.46 | 0.45 | 1.00 | 1.00 | 1.00 | 0.80 | 0.88 | 1.00 | 0.63 | 0.39 | 0.37 | 0.40 | 1.00 | 0.89 | 0.51 |
| CellPLM | 0.34 | 0.33 | 0.30 | 0.26 | 0.24 | 1.00 | 1.00 | 0.00 | 0.79 | 0.89 | 1.00 | 0.63 | 0.49 | 0.34 | 0.28 | 0.67 | 0.89 | 0.56 |
| CellFM | 0.00 | 0.26 | 0.25 | 0.27 | 0.26 | 1.00 | 1.00 | 1.00 | 0.51 | 0.78 | 1.00 | 0.37 | 0.27 | 0.00 | 0.26 | 1.00 | 0.76 | 0.32 |
| scPRINT | 0.27 | 0.00 | 0.00 | 0.37 | 0.34 | 1.00 | 1.00 | 0.00 | 0.36 | 0.74 | 1.00 | 0.63 | 0.31 | 0.27 | 0.18 | 0.67 | 0.70 | 0.47 |
| GenePT-w | 0.32 | 0.11 | 0.11 | 0.05 | 0.02 | 1.00 | 1.00 | 0.00 | 0.46 | 0.47 | 1.00 | 0.63 | 0.38 | 0.32 | 0.07 | 0.67 | 0.64 | 0.50 |
| scGPT | 0.08 | 0.19 | 0.17 | 0.09 | 0.10 | 1.00 | 1.00 | 0.00 | 0.54 | 0.65 | 1.00 | 0.63 | 0.38 | 0.08 | 0.14 | 0.67 | 0.73 | 0.51 |
| Geneformer | 0.16 | 0.31 | 0.28 | 0.20 | 0.18 | 0.67 | 0.65 | 0.00 | 0.55 | 0.79 | 1.00 | 0.32 | 0.31 | 0.16 | 0.24 | 0.44 | 0.78 | 0.32 |
| LangCell | 0.00 | 0.09 | 0.08 | 0.03 | 0.04 | 1.00 | 1.00 | 0.00 | 0.17 | 0.30 | 1.00 | 0.63 | 0.33 | 0.00 | 0.06 | 0.67 | 0.49 | 0.48 |
| scVI (de novo) | 0.00 | 0.00 | 0.00 | 0.00 | 0.00 | 1.00 | 1.00 | 0.00 | 0.08 | 0.04 | 1.00 | 0.63 | 0.31 | 0.00 | 0.00 | 0.67 | 0.37 | 0.47 |

**Figure S104: Trajectory-inference metric scores for distal lung epithelium using Monocle 3.** Rows show 17 representations; circles report individual `dyneval` scores grouped into cell position, reference-position recoverability, topology, feature relevance and branch assignment. Horizontal bars show the mean of the available scores within each family, and rows are ordered by the mean of the family summaries. Blank entries indicate unavailable metrics.

| | Cell positions | Neighbourhood | | | | Topology | | | Features | | | Branch assignment | | Aggregated scores | | | | |
|---|---|---|---|---|---|---|---|---|---|---|---|---|---|---|---|---|---|---|
| method | $cor_{dist}$ | $NMSE_{lm}$ | $R^2_{lm}$ | $NMSE_{rf}$ | $R^2_{rf}$ | edgeflip | HIM | isomorphic | $cor_{features}$ | $wcor_{features}$ | $ks_{features}$ | $F1_{branches}$ | $F1_{milestones}$ | Pos. | Neighbor. | Topology | Feature | Branch |
| scFoundation | 0.71 | 0.54 | 0.48 | 0.74 | 0.72 | | | | 0.91 | 0.93 | 1.00 | | 0.70 | 0.71 | 0.62 | | 0.95 | 0.70 |
| scCello | 0.75 | 0.60 | 0.54 | 0.48 | 0.41 | | | | 0.93 | 0.95 | 1.00 | | 0.65 | 0.75 | 0.51 | | 0.96 | 0.65 |
| CELLama | 0.67 | 0.51 | 0.46 | 0.24 | 0.16 | | | | 0.92 | 0.94 | 1.00 | | 0.63 | 0.67 | 0.34 | | 0.96 | 0.63 |
| scVI | 0.70 | 0.30 | 0.34 | 0.37 | 0.43 | | | | 0.82 | 0.74 | 1.00 | | 0.66 | 0.70 | 0.36 | | 0.85 | 0.66 |
| scGPT | 0.49 | 0.14 | 0.15 | 0.03 | 0.06 | | | | 0.70 | 0.65 | 1.00 | | 0.54 | 0.49 | 0.09 | | 0.78 | 0.54 |
| scGPT (integrated | 0.32 | 0.25 | 0.22 | 0.02 | 0.01 | | | | 0.77 | 0.90 | 1.00 | | 0.54 | 0.32 | 0.12 | | 0.89 | 0.54 |
| SCimilarity | 0.24 | 0.34 | 0.31 | 0.32 | 0.32 | | | | 0.42 | 0.67 | 1.00 | | 0.57 | 0.24 | 0.32 | | 0.70 | 0.57 |
| UCE | 0.20 | 0.26 | 0.24 | 0.00 | 0.00 | | | | 0.54 | 0.80 | 1.00 | | 0.58 | 0.20 | 0.13 | | 0.78 | 0.58 |
| CellFM | 0.23 | 0.12 | 0.13 | 0.00 | 0.00 | | | | 0.30 | 0.55 | 0.99 | | 0.39 | 0.23 | 0.06 | | 0.62 | 0.39 |
| PCA | 0.28 | 0.00 | 0.00 | 0.03 | 0.03 | | | | 0.37 | 0.59 | 1.00 | | 0.34 | 0.28 | 0.02 | | 0.65 | 0.34 |
| Geneformer | 0.11 | 0.08 | 0.08 | 0.20 | 0.20 | | | | 0.10 | 0.52 | 1.00 | | 0.33 | 0.11 | 0.14 | | 0.54 | 0.33 |
| scBERT | 0.07 | 0.13 | 0.12 | 0.00 | 0.00 | | | | 0.29 | 0.43 | 1.00 | | 0.40 | 0.07 | 0.06 | | 0.57 | 0.40 |
| CellPLM | 0.11 | 0.14 | 0.14 | 0.00 | 0.00 | | | | 0.22 | 0.26 | 1.00 | | 0.43 | 0.11 | 0.07 | | 0.49 | 0.43 |
| GenePT-w | 0.01 | 0.05 | 0.06 | 0.00 | 0.00 | | | | 0.20 | 0.25 | 1.00 | | 0.35 | 0.01 | 0.03 | | 0.48 | 0.35 |
| LangCell | 0.04 | 0.09 | 0.08 | 0.00 | 0.00 | | | | 0.14 | 0.22 | 1.00 | | 0.32 | 0.04 | 0.04 | | 0.45 | 0.32 |
| scPRINT | 0.10 | 0.00 | 0.00 | 0.00 | 0.00 | | | | 0.11 | 0.25 | 0.98 | | 0.31 | 0.10 | 0.00 | | 0.45 | 0.31 |
| scVI (de novo) | 0.00 | 0.01 | 0.01 | 0.00 | 0.00 | | | | 0.04 | 0.00 | 0.95 | | 0.31 | 0.00 | 0.01 | | 0.33 | 0.31 |

**Figure S105: Trajectory-inference metric scores for distal lung epithelium using DPT.** Rows show 17 representations; circles report individual `dyneval` scores grouped into cell position, reference-position recoverability, topology, feature relevance and branch assignment. Horizontal bars show the mean of the available scores within each family, and rows are ordered by the mean of the family summaries. DPT yields a directed linear trajectory, so topology scores and branch-level F1 are unavailable and shown as blanks.

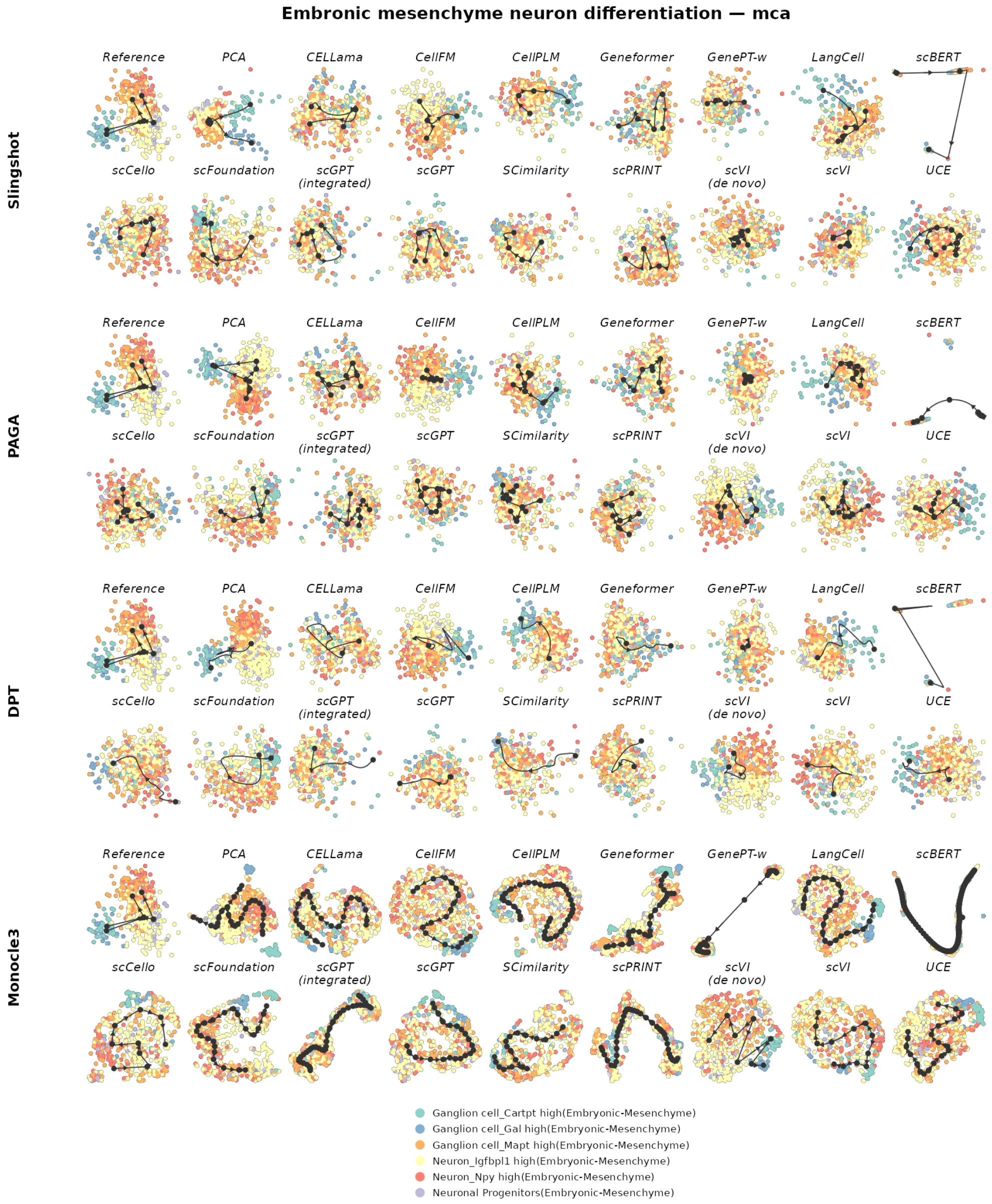

**Figure S106: Trajectory inference on embryonic mesenchyme-to-neuron differentiation.** The reference trajectory is tree-structured. Rows show Slingshot, PAGA, DPT and Monocle 3; within each row, the reference is followed by the same 17 representations. Cells are coloured by the reference milestone or state. Black nodes, edges and arrows show the inferred milestone graph and its orientation. Layouts and trajectories were reconstructed independently for each representation and backend.

| | Cell positions | Neighbourhood | | | | Topology | | | Features | | | Branch assignment | | Aggregated scores | | | | |
|---|---|---|---|---|---|---|---|---|---|---|---|---|---|---|---|---|---|---|
| method | $cor_{dist}$ | $NMSE_{lm}$ | $R^2_{lm}$ | $NMSE_{rf}$ | $R^2_{rf}$ | edgeflip | HIM | isomorphic | $cor_{features}$ | $wcor_{features}$ | $ks_{features}$ | $F1_{branches}$ | $F1_{milestones}$ | Pos. | Neighbor. | Topology | Feature | Branch |
| PCA | 0.21 | 0.21 | 0.30 | 0.21 | 0.31 | 0.75 | 0.62 | 0.00 | 0.49 | 0.42 | 1.00 | 0.28 | 0.37 | 0.21 | 0.26 | 0.46 | 0.64 | 0.32 |
| scVI | 0.26 | 0.11 | 0.08 | 0.00 | 0.00 | 0.80 | 0.68 | 0.00 | 0.55 | 0.73 | 1.00 | 0.25 | 0.23 | 0.26 | 0.05 | 0.49 | 0.76 | 0.24 |
| LangCell | 0.09 | 0.13 | 0.16 | 0.10 | 0.15 | 1.00 | 0.94 | 0.00 | 0.50 | 0.35 | 1.00 | 0.28 | 0.28 | 0.09 | 0.14 | 0.65 | 0.62 | 0.28 |
| scFoundation | 0.14 | 0.14 | 0.17 | 0.13 | 0.20 | 0.80 | 0.75 | 0.00 | 0.52 | 0.46 | 1.00 | 0.30 | 0.27 | 0.14 | 0.16 | 0.52 | 0.66 | 0.28 |
| UCE | 0.12 | 0.20 | 0.22 | 0.16 | 0.19 | 0.75 | 0.67 | 0.00 | 0.45 | 0.57 | 1.00 | 0.26 | 0.28 | 0.12 | 0.19 | 0.47 | 0.67 | 0.27 |
| CellFM | 0.10 | 0.14 | 0.14 | 0.10 | 0.09 | 0.80 | 0.69 | 0.00 | 0.56 | 0.54 | 1.00 | 0.22 | 0.26 | 0.10 | 0.12 | 0.50 | 0.70 | 0.24 |
| scVI (de novo) | 0.10 | 0.14 | 0.14 | 0.14 | 0.19 | 0.67 | 0.63 | 0.00 | 0.54 | 0.64 | 1.00 | 0.17 | 0.22 | 0.10 | 0.15 | 0.43 | 0.73 | 0.19 |
| CellPLM | 0.15 | 0.07 | 0.09 | 0.00 | 0.00 | 0.50 | 0.39 | 0.00 | 0.44 | 0.32 | 1.00 | 0.29 | 0.27 | 0.15 | 0.04 | 0.30 | 0.59 | 0.28 |
| scCello | 0.08 | 0.03 | 0.04 | 0.00 | 0.00 | 0.50 | 0.39 | 0.00 | 0.51 | 0.57 | 1.00 | 0.29 | 0.22 | 0.08 | 0.02 | 0.30 | 0.70 | 0.25 |
| CELLama | 0.04 | 0.02 | 0.02 | 0.00 | 0.00 | 0.80 | 0.70 | 0.00 | 0.34 | 0.35 | 1.00 | 0.20 | 0.17 | 0.04 | 0.01 | 0.50 | 0.56 | 0.19 |
| Geneformer | 0.10 | 0.05 | 0.07 | 0.04 | 0.06 | 0.50 | 0.39 | 0.00 | 0.43 | 0.34 | 1.00 | 0.24 | 0.25 | 0.10 | 0.05 | 0.30 | 0.59 | 0.25 |
| scGPT | 0.00 | 0.02 | 0.02 | 0.00 | 0.00 | 0.80 | 0.69 | 0.00 | 0.32 | 0.38 | 1.00 | 0.21 | 0.16 | 0.00 | 0.01 | 0.50 | 0.57 | 0.18 |
| scBERT | 0.13 | 0.02 | 0.03 | 0.00 | 0.00 | 0.50 | 0.39 | 0.00 | 0.24 | 0.11 | 1.00 | 0.29 | 0.20 | 0.13 | 0.01 | 0.30 | 0.45 | 0.25 |
| GenePT-w | 0.04 | 0.02 | 0.02 | 0.00 | 0.00 | 0.50 | 0.39 | 0.00 | 0.29 | 0.23 | 1.00 | 0.26 | 0.20 | 0.04 | 0.01 | 0.30 | 0.51 | 0.23 |
| SCimilarity | 0.01 | 0.01 | 0.01 | 0.00 | 0.00 | 0.50 | 0.39 | 0.00 | 0.32 | 0.19 | 1.00 | 0.29 | 0.20 | 0.01 | 0.01 | 0.30 | 0.50 | 0.24 |
| scGPT (integrated) | 0.03 | 0.01 | 0.01 | 0.00 | 0.00 | 0.50 | 0.39 | 0.00 | 0.30 | 0.23 | 1.00 | 0.23 | 0.18 | 0.03 | 0.01 | 0.30 | 0.51 | 0.20 |
| scPRINT | 0.03 | 0.01 | 0.01 | 0.00 | 0.00 | 0.50 | 0.39 | 0.00 | 0.28 | 0.18 | 1.00 | 0.23 | 0.17 | 0.03 | 0.00 | 0.30 | 0.49 | 0.20 |

**Figure S107: Trajectory-inference metric scores for embryonic mesenchyme-to-neuron differentiation using Slingshot.** Display and scoring conventions are as in Fig. S102.

| | Cell positions | Neighbourhood | | | | Topology | | | Features | | | Branch assignment | | Aggregated scores | | | | |
|---|---|---|---|---|---|---|---|---|---|---|---|---|---|---|---|---|---|---|
| method | $cor_{dist}$ | $NMSE_{lm}$ | $R^2_{lm}$ | $NMSE_{rf}$ | $R^2_{rf}$ | edgeflip | HIM | isomorphic | $cor_{features}$ | $wcor_{features}$ | $ks_{features}$ | $F1_{branches}$ | $F1_{milestones}$ | Pos. | Neighbor. | Topology | Feature | Branch |
| PCA | 0.27 | 0.24 | 0.32 | 0.27 | 0.36 | 1.00 | 0.88 | 1.00 | 0.71 | 0.85 | 1.00 | 0.41 | 0.37 | 0.27 | 0.30 | 0.96 | 0.85 | 0.39 |
| scVI | 0.07 | 0.13 | 0.16 | 0.19 | 0.22 | 1.00 | 0.91 | 0.00 | 0.57 | 0.64 | 1.00 | 0.29 | 0.31 | 0.07 | 0.18 | 0.64 | 0.74 | 0.30 |
| scCello | 0.11 | 0.07 | 0.09 | 0.10 | 0.12 | 1.00 | 0.88 | 0.00 | 0.55 | 0.68 | 1.00 | 0.21 | 0.23 | 0.11 | 0.09 | 0.63 | 0.74 | 0.22 |
| scVI (de novo) | 0.18 | 0.10 | 0.15 | 0.19 | 0.26 | 0.50 | 0.39 | 0.00 | 0.52 | 0.44 | 1.00 | 0.35 | 0.39 | 0.18 | 0.17 | 0.30 | 0.65 | 0.37 |
| UCE | 0.12 | 0.12 | 0.14 | 0.14 | 0.21 | 0.80 | 0.70 | 0.00 | 0.39 | 0.41 | 1.00 | 0.28 | 0.27 | 0.12 | 0.15 | 0.50 | 0.60 | 0.28 |
| Geneformer | 0.10 | 0.14 | 0.15 | 0.14 | 0.16 | 0.80 | 0.69 | 0.00 | 0.51 | 0.44 | 1.00 | 0.27 | 0.25 | 0.10 | 0.15 | 0.50 | 0.65 | 0.26 |
| scFoundation | 0.01 | 0.14 | 0.17 | 0.13 | 0.16 | 0.80 | 0.72 | 0.00 | 0.50 | 0.34 | 1.00 | 0.36 | 0.30 | 0.01 | 0.15 | 0.51 | 0.61 | 0.33 |
| CellPLM | 0.08 | 0.12 | 0.15 | 0.13 | 0.18 | 0.57 | 0.68 | 0.00 | 0.50 | 0.53 | 1.00 | 0.29 | 0.22 | 0.08 | 0.14 | 0.42 | 0.68 | 0.26 |
| CELLama | 0.06 | 0.07 | 0.07 | 0.06 | 0.06 | 0.57 | 0.73 | 0.00 | 0.59 | 0.77 | 1.00 | 0.19 | 0.17 | 0.06 | 0.06 | 0.43 | 0.79 | 0.18 |
| LangCell | 0.04 | 0.16 | 0.20 | 0.15 | 0.20 | 0.57 | 0.68 | 0.00 | 0.48 | 0.35 | 1.00 | 0.30 | 0.27 | 0.04 | 0.18 | 0.42 | 0.61 | 0.28 |
| SCimilarity | 0.00 | 0.07 | 0.07 | 0.05 | 0.05 | 0.75 | 0.65 | 0.00 | 0.33 | 0.36 | 1.00 | 0.19 | 0.16 | 0.00 | 0.06 | 0.47 | 0.56 | 0.18 |
| scGPT | 0.03 | 0.09 | 0.14 | 0.07 | 0.11 | 0.44 | 0.64 | 0.00 | 0.39 | 0.26 | 1.00 | 0.20 | 0.22 | 0.03 | 0.10 | 0.36 | 0.55 | 0.21 |
| CellFM | 0.01 | 0.03 | 0.05 | 0.01 | 0.03 | 0.57 | 0.72 | 0.00 | 0.39 | 0.37 | 1.00 | 0.22 | 0.18 | 0.01 | 0.03 | 0.43 | 0.59 | 0.20 |
| scBERT | 0.03 | 0.02 | 0.03 | 0.00 | 0.00 | 0.80 | 0.61 | 0.00 | 0.29 | 0.26 | 1.00 | 0.23 | 0.17 | 0.03 | 0.01 | 0.47 | 0.52 | 0.20 |
| scPRINT | 0.01 | 0.01 | 0.02 | 0.00 | 0.00 | 0.80 | 0.68 | 0.00 | 0.32 | 0.32 | 1.00 | 0.18 | 0.14 | 0.01 | 0.01 | 0.49 | 0.55 | 0.16 |
| scGPT (integrated) | 0.01 | 0.03 | 0.04 | 0.01 | 0.02 | 0.57 | 0.73 | 0.00 | 0.32 | 0.19 | 1.00 | 0.20 | 0.16 | 0.01 | 0.02 | 0.43 | 0.50 | 0.18 |
| GenePT-w | 0.04 | 0.02 | 0.02 | 0.00 | 0.00 | 0.50 | 0.39 | 0.00 | 0.29 | 0.18 | 1.00 | 0.25 | 0.20 | 0.04 | 0.01 | 0.30 | 0.49 | 0.22 |

**Figure S108: Trajectory-inference metric scores for embryonic mesenchyme-to-neuron differentiation using PAGA.** Display and scoring conventions are as in Fig. S103.

| | Cell positions | Neighbourhood | | | | Topology | | | Features | | | Branch assignment | | Aggregated scores | | | | |
|---|---|---|---|---|---|---|---|---|---|---|---|---|---|---|---|---|---|---|
| method | $cor_{dist}$ | $NMSE_{lm}$ | $R^2_{lm}$ | $NMSE_{rf}$ | $R^2_{rf}$ | edgeflip | HIM | isomorphic | $cor_{features}$ | $wcor_{features}$ | $ks_{features}$ | $F1_{branches}$ | $F1_{milestones}$ | Pos. | Neighbor. | Topology | Feature | Branch |
| PCA | 0.11 | 0.15 | 0.16 | 0.20 | 0.20 | 0.80 | 0.67 | 0.00 | 0.63 | 0.68 | 1.00 | 0.25 | 0.26 | 0.11 | 0.18 | 0.49 | 0.77 | 0.25 |
| scVI | 0.10 | 0.08 | 0.09 | 0.13 | 0.17 | 0.50 | 0.39 | 0.00 | 0.57 | 0.65 | 1.00 | 0.29 | 0.29 | 0.10 | 0.12 | 0.30 | 0.74 | 0.29 |
| Geneformer | 0.08 | 0.10 | 0.11 | 0.15 | 0.15 | 0.80 | 0.59 | 0.00 | 0.44 | 0.31 | 1.00 | 0.27 | 0.23 | 0.08 | 0.13 | 0.46 | 0.58 | 0.25 |
| scVI (de novo) | 0.13 | 0.07 | 0.12 | 0.15 | 0.22 | 0.50 | 0.39 | 0.00 | 0.44 | 0.31 | 1.00 | 0.29 | 0.32 | 0.13 | 0.14 | 0.30 | 0.58 | 0.30 |
| LangCell | 0.13 | 0.09 | 0.11 | 0.20 | 0.28 | 0.50 | 0.39 | 0.00 | 0.47 | 0.34 | 1.00 | 0.23 | 0.27 | 0.13 | 0.17 | 0.30 | 0.60 | 0.25 |
| CellFM | 0.11 | 0.06 | 0.08 | 0.08 | 0.13 | 0.50 | 0.39 | 0.00 | 0.42 | 0.38 | 1.00 | 0.25 | 0.29 | 0.11 | 0.09 | 0.30 | 0.60 | 0.27 |
| scFoundation | 0.07 | 0.07 | 0.10 | 0.13 | 0.19 | 0.50 | 0.39 | 0.00 | 0.42 | 0.23 | 1.00 | 0.35 | 0.30 | 0.07 | 0.12 | 0.30 | 0.55 | 0.33 |
| SCimilarity | 0.05 | 0.03 | 0.05 | 0.03 | 0.04 | 0.80 | 0.57 | 0.00 | 0.38 | 0.30 | 1.00 | 0.26 | 0.23 | 0.05 | 0.04 | 0.46 | 0.56 | 0.25 |
| scGPT | 0.10 | 0.04 | 0.05 | 0.08 | 0.13 | 0.50 | 0.39 | 0.00 | 0.40 | 0.25 | 1.00 | 0.29 | 0.24 | 0.10 | 0.07 | 0.30 | 0.55 | 0.27 |
| scCello | 0.06 | 0.01 | 0.02 | 0.05 | 0.06 | 0.50 | 0.39 | 0.00 | 0.33 | 0.50 | 1.00 | 0.29 | 0.22 | 0.06 | 0.03 | 0.30 | 0.61 | 0.25 |
| CELLama | 0.03 | 0.02 | 0.03 | 0.06 | 0.08 | 0.50 | 0.39 | 0.00 | 0.44 | 0.39 | 1.00 | 0.29 | 0.21 | 0.03 | 0.05 | 0.30 | 0.61 | 0.25 |
| UCE | 0.00 | 0.07 | 0.06 | 0.12 | 0.13 | 0.50 | 0.39 | 0.00 | 0.36 | 0.45 | 1.00 | 0.24 | 0.19 | 0.00 | 0.10 | 0.30 | 0.60 | 0.22 |
| GenePT-w | 0.01 | 0.03 | 0.03 | 0.03 | 0.05 | 0.80 | 0.51 | 0.00 | 0.33 | 0.28 | 1.00 | 0.20 | 0.15 | 0.01 | 0.04 | 0.44 | 0.54 | 0.18 |
| CellPLM | 0.03 | 0.03 | 0.03 | 0.07 | 0.11 | 0.50 | 0.39 | 0.00 | 0.37 | 0.35 | 1.00 | 0.24 | 0.22 | 0.03 | 0.06 | 0.30 | 0.57 | 0.23 |
| scGPT (integrated | 0.02 | 0.03 | 0.04 | 0.00 | 0.00 | 0.80 | 0.48 | 0.00 | 0.29 | 0.23 | 1.00 | 0.23 | 0.17 | 0.02 | 0.02 | 0.43 | 0.50 | 0.20 |
| scBERT | 0.03 | 0.02 | 0.03 | 0.00 | 0.00 | 0.80 | 0.50 | 0.00 | 0.29 | 0.22 | 1.00 | 0.18 | 0.16 | 0.03 | 0.01 | 0.43 | 0.50 | 0.17 |
| scPRINT | 0.01 | 0.02 | 0.04 | 0.03 | 0.06 | 0.50 | 0.39 | 0.00 | 0.30 | 0.21 | 1.00 | 0.28 | 0.22 | 0.01 | 0.04 | 0.30 | 0.50 | 0.25 |

**Figure S109: Trajectory-inference metric scores for embryonic mesenchyme-to-neuron differentiation using Monocle 3.** Display and scoring conventions are as in Fig. S104.

| | Cell positions | Neighbourhood | | | | Topology | | | Features | | | Branch assignment | | Aggregated scores | | | | |
|---|---|---|---|---|---|---|---|---|---|---|---|---|---|---|---|---|---|---|
| method | $cor_{dist}$ | $NMSE_{lm}$ | $R^2_{lm}$ | $NMSE_{rf}$ | $R^2_{rf}$ | edgeflip | HIM | isomorphic | $cor_{features}$ | $wcor_{features}$ | $ks_{features}$ | $F1_{branches}$ | $F1_{milestones}$ | Pos. | Neighbor. | Topology | Feature | Branch |
| scCello | 0.20 | 0.05 | 0.05 | 0.00 | 0.00 | | | | 0.57 | 0.71 | 1.00 | | 0.27 | 0.20 | 0.02 | | 0.76 | 0.27 |
| scVI | 0.16 | 0.03 | 0.04 | 0.00 | 0.00 | | | | 0.59 | 0.68 | 1.00 | | 0.25 | 0.16 | 0.02 | | 0.76 | 0.25 |
| UCE | 0.22 | 0.08 | 0.12 | 0.00 | 0.00 | | | | 0.35 | 0.16 | 1.00 | | 0.38 | 0.22 | 0.05 | | 0.50 | 0.38 |
| CellPLM | 0.19 | 0.10 | 0.13 | 0.00 | 0.00 | | | | 0.37 | 0.22 | 1.00 | | 0.35 | 0.19 | 0.06 | | 0.53 | 0.35 |
| LangCell | 0.20 | 0.10 | 0.13 | 0.00 | 0.00 | | | | 0.31 | 0.14 | 0.89 | | 0.36 | 0.20 | 0.06 | | 0.45 | 0.36 |
| Geneformer | 0.16 | 0.07 | 0.07 | 0.00 | 0.00 | | | | 0.37 | 0.25 | 1.00 | | 0.30 | 0.16 | 0.04 | | 0.54 | 0.30 |
| PCA | 0.17 | 0.07 | 0.13 | 0.00 | 0.00 | | | | 0.28 | 0.12 | 0.82 | | 0.34 | 0.17 | 0.05 | | 0.41 | 0.34 |
| scFoundation | 0.06 | 0.05 | 0.06 | 0.00 | 0.00 | | | | 0.35 | 0.19 | 1.00 | | 0.27 | 0.06 | 0.03 | | 0.51 | 0.27 |
| scGPT | 0.09 | 0.01 | 0.01 | 0.00 | 0.00 | | | | 0.30 | 0.35 | 1.00 | | 0.22 | 0.09 | 0.00 | | 0.55 | 0.22 |
| scGPT (integrated | 0.09 | 0.01 | 0.02 | 0.00 | 0.00 | | | | 0.28 | 0.17 | 1.00 | | 0.24 | 0.09 | 0.01 | | 0.48 | 0.24 |
| scVI (de novo) | 0.01 | 0.02 | 0.03 | 0.00 | 0.00 | | | | 0.37 | 0.30 | 1.00 | | 0.22 | 0.01 | 0.01 | | 0.55 | 0.22 |
| SCimilarity | 0.10 | 0.01 | 0.02 | 0.00 | 0.00 | | | | 0.18 | 0.23 | 1.00 | | 0.21 | 0.10 | 0.01 | | 0.47 | 0.21 |
| GenePT-w | 0.05 | 0.01 | 0.01 | 0.00 | 0.00 | | | | 0.28 | 0.24 | 1.00 | | 0.20 | 0.05 | 0.01 | | 0.51 | 0.20 |
| scPRINT | 0.00 | 0.01 | 0.01 | 0.00 | 0.00 | | | | 0.28 | 0.21 | 1.00 | | 0.21 | 0.00 | 0.00 | | 0.50 | 0.21 |
| scBERT | 0.00 | 0.00 | 0.00 | 0.00 | 0.00 | | | | 0.24 | 0.16 | 1.00 | | 0.20 | 0.00 | 0.00 | | 0.47 | 0.20 |
| CellFM | 0.05 | 0.01 | 0.01 | 0.00 | 0.00 | | | | 0.13 | 0.12 | 1.00 | | 0.20 | 0.05 | 0.00 | | 0.42 | 0.20 |
| CELLama | 0.02 | 0.01 | 0.01 | 0.00 | 0.00 | | | | 0.05 | 0.13 | 1.00 | | 0.24 | 0.02 | 0.00 | | 0.39 | 0.24 |

**Figure S110: Trajectory-inference metric scores for embryonic mesenchyme-to-neuron differentiation using DPT.** Display and scoring conventions are as in Fig. S105.

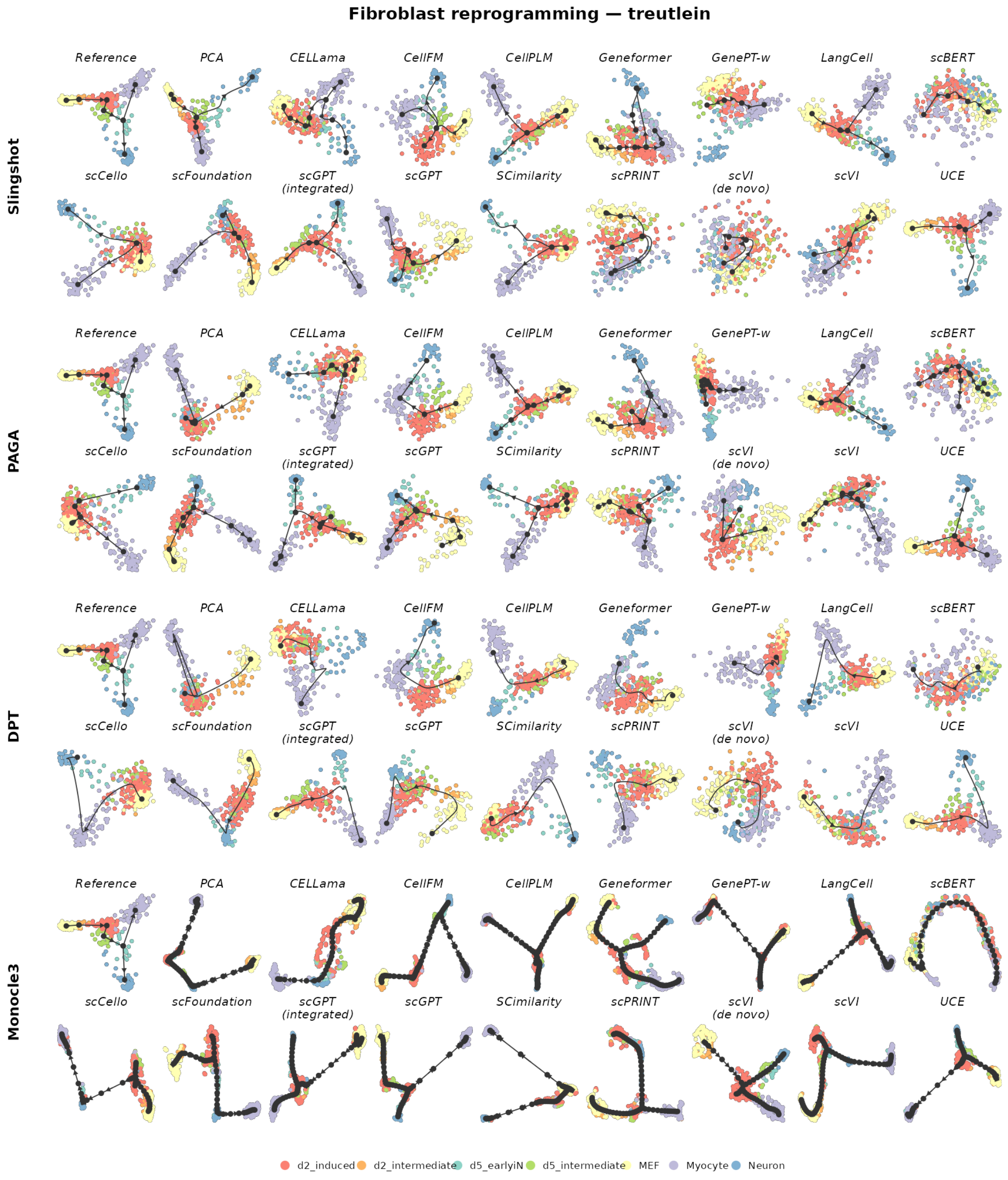

**Figure S111: Trajectory inference on fibroblast reprogramming.** The reference trajectory is bifurcating. Rows show Slingshot, PAGA, DPT and Monocle 3; within each row, the reference is followed by the same 17 representations. Cells are coloured by the reference milestone or state. Black nodes, edges and arrows show the inferred milestone graph and its orientation. Layouts and trajectories were reconstructed independently for each representation and backend.

| | Cell positions | Neighbourhood | | | | Topology | | | Features | | | Branch assignment | | Aggregated scores | | | | |
|---|---|---|---|---|---|---|---|---|---|---|---|---|---|---|---|---|---|---|
| method | $cor_{dist}$ | $NMSE_{lm}$ | $R^2_{lm}$ | $NMSE_{rf}$ | $R^2_{rf}$ | edgeflip | HIM | isomorphic | $cor_{features}$ | $wcor_{features}$ | $ks_{features}$ | $F1_{branches}$ | $F1_{milestones}$ | Pos. | Neighbor. | Topology | Feature | Branch |
| LangCell | 0.82 | 0.85 | 0.84 | 0.91 | 0.90 | 1.00 | 0.70 | 1.00 | 0.91 | 0.90 | 0.99 | 0.74 | 0.65 | 0.82 | 0.87 | 0.90 | 0.93 | 0.69 |
| UCE | 0.81 | 0.85 | 0.83 | 0.91 | 0.90 | 1.00 | 0.71 | 1.00 | 0.90 | 0.87 | 1.00 | 0.71 | 0.63 | 0.81 | 0.87 | 0.90 | 0.92 | 0.67 |
| scFoundation | 0.81 | 0.84 | 0.82 | 0.85 | 0.83 | 1.00 | 0.68 | 1.00 | 0.91 | 0.87 | 1.00 | 0.66 | 0.63 | 0.81 | 0.83 | 0.89 | 0.93 | 0.64 |
| scGPT (integrated | 0.79 | 0.82 | 0.79 | 0.84 | 0.82 | 1.00 | 0.68 | 1.00 | 0.90 | 0.89 | 1.00 | 0.71 | 0.60 | 0.79 | 0.82 | 0.89 | 0.93 | 0.66 |
| CellPLM | 0.76 | 0.81 | 0.77 | 0.86 | 0.84 | 1.00 | 0.68 | 1.00 | 0.87 | 0.89 | 1.00 | 0.64 | 0.60 | 0.76 | 0.82 | 0.89 | 0.92 | 0.62 |
| SCimilarity | 0.65 | 0.76 | 0.72 | 0.80 | 0.78 | 1.00 | 0.90 | 1.00 | 0.91 | 0.91 | 0.93 | 0.69 | 0.61 | 0.65 | 0.77 | 0.97 | 0.92 | 0.65 |
| scCello | 0.72 | 0.77 | 0.75 | 0.81 | 0.79 | 1.00 | 0.73 | 1.00 | 0.90 | 0.87 | 1.00 | 0.60 | 0.61 | 0.72 | 0.78 | 0.91 | 0.92 | 0.61 |
| PCA | 0.69 | 0.79 | 0.79 | 0.88 | 0.87 | 1.00 | 0.72 | 1.00 | 0.86 | 0.85 | 1.00 | 0.57 | 0.65 | 0.69 | 0.83 | 0.91 | 0.90 | 0.61 |
| scVI | 0.65 | 0.77 | 0.74 | 0.83 | 0.80 | 1.00 | 0.84 | 1.00 | 0.79 | 0.89 | 1.00 | 0.63 | 0.61 | 0.65 | 0.79 | 0.95 | 0.89 | 0.62 |
| scVI (de novo) | 0.72 | 0.71 | 0.66 | 0.80 | 0.78 | 1.00 | 0.69 | 1.00 | 0.83 | 0.84 | 1.00 | 0.48 | 0.56 | 0.72 | 0.74 | 0.90 | 0.89 | 0.52 |
| scGPT | 0.72 | 0.78 | 0.74 | 0.84 | 0.81 | 1.00 | 0.68 | 0.00 | 0.91 | 0.91 | 1.00 | 0.73 | 0.60 | 0.72 | 0.79 | 0.56 | 0.94 | 0.67 |
| CellFM | 0.79 | 0.85 | 0.83 | 0.88 | 0.87 | 0.80 | 0.73 | 0.00 | 0.87 | 0.83 | 1.00 | 0.62 | 0.59 | 0.79 | 0.86 | 0.51 | 0.90 | 0.61 |
| CELLama | 0.72 | 0.61 | 0.58 | 0.82 | 0.79 | 1.00 | 0.80 | 1.00 | 0.81 | 0.66 | 0.50 | 0.62 | 0.58 | 0.72 | 0.70 | 0.93 | 0.66 | 0.60 |
| Geneformer | 0.46 | 0.60 | 0.52 | 0.78 | 0.76 | 1.00 | 0.76 | 1.00 | 0.78 | 0.84 | 1.00 | 0.60 | 0.43 | 0.46 | 0.66 | 0.92 | 0.87 | 0.51 |
| scPRINT | 0.64 | 0.69 | 0.65 | 0.72 | 0.67 | 0.80 | 0.73 | 0.00 | 0.88 | 0.87 | 1.00 | 0.58 | 0.46 | 0.64 | 0.68 | 0.51 | 0.92 | 0.52 |
| GenePT-w | 0.29 | 0.27 | 0.22 | 0.25 | 0.20 | 0.67 | 0.52 | 0.00 | 0.59 | 0.74 | 1.00 | 0.37 | 0.37 | 0.29 | 0.24 | 0.40 | 0.78 | 0.37 |
| scBERT | 0.20 | 0.12 | 0.11 | 0.00 | 0.00 | 0.67 | 0.52 | 0.00 | 0.26 | 0.04 | 1.00 | 0.45 | 0.29 | 0.20 | 0.06 | 0.40 | 0.43 | 0.37 |

**Figure S112: Trajectory-inference metric scores for fibroblast reprogramming using Slingshot.** Display and scoring conventions are as in Fig. S102.

| | Cell positions | Neighbourhood | | | | Topology | | | Features | | | Branch assignment | | Aggregated scores | | | | |
|---|---|---|---|---|---|---|---|---|---|---|---|---|---|---|---|---|---|---|
| method | $cor_{dist}$ | $NMSE_{lm}$ | $R^2_{lm}$ | $NMSE_{rf}$ | $R^2_{rf}$ | edgeflip | HIM | isomorphic | $cor_{features}$ | $wcor_{features}$ | $ks_{features}$ | $F1_{branches}$ | $F1_{milestones}$ | Pos. | Neighbor. | Topology | Feature | Branch |
| LangCell | 0.75 | 0.83 | 0.79 | 0.87 | 0.86 | 1.00 | 0.76 | 1.00 | 0.92 | 0.91 | 1.00 | 0.77 | 0.66 | 0.75 | 0.84 | 0.92 | 0.94 | 0.72 |
| CellPLM | 0.75 | 0.78 | 0.76 | 0.89 | 0.88 | 1.00 | 0.77 | 1.00 | 0.84 | 0.79 | 1.00 | 0.84 | 0.48 | 0.75 | 0.83 | 0.92 | 0.88 | 0.66 |
| scGPT | 0.71 | 0.74 | 0.70 | 0.83 | 0.78 | 1.00 | 0.80 | 1.00 | 0.87 | 0.83 | 1.00 | 0.66 | 0.61 | 0.71 | 0.76 | 0.93 | 0.90 | 0.63 |
| scVI (de novo) | 0.74 | 0.75 | 0.68 | 0.74 | 0.67 | 1.00 | 0.68 | 1.00 | 0.85 | 0.84 | 1.00 | 0.75 | 0.59 | 0.74 | 0.71 | 0.89 | 0.90 | 0.67 |
| UCE | 0.66 | 0.71 | 0.69 | 0.82 | 0.78 | 1.00 | 0.76 | 1.00 | 0.81 | 0.73 | 1.00 | 0.58 | 0.62 | 0.66 | 0.75 | 0.92 | 0.85 | 0.60 |
| scCello | 0.57 | 0.73 | 0.67 | 0.75 | 0.71 | 1.00 | 0.88 | 1.00 | 0.89 | 0.87 | 1.00 | 0.61 | 0.56 | 0.57 | 0.71 | 0.96 | 0.92 | 0.59 |
| scPRINT | 0.61 | 0.64 | 0.58 | 0.68 | 0.63 | 1.00 | 0.81 | 1.00 | 0.92 | 0.90 | 1.00 | 0.59 | 0.52 | 0.61 | 0.63 | 0.94 | 0.94 | 0.56 |
| scFoundation | 0.77 | 0.80 | 0.78 | 0.84 | 0.82 | 0.80 | 0.76 | 0.00 | 0.90 | 0.88 | 1.00 | 0.75 | 0.52 | 0.77 | 0.81 | 0.52 | 0.93 | 0.64 |
| SCimilarity | 0.76 | 0.73 | 0.68 | 0.81 | 0.76 | 1.00 | 0.86 | 0.00 | 0.87 | 0.89 | 1.00 | 0.70 | 0.47 | 0.76 | 0.74 | 0.62 | 0.92 | 0.58 |
| Geneformer | 0.80 | 0.84 | 0.82 | 0.88 | 0.86 | 0.67 | 0.58 | 0.00 | 0.88 | 0.95 | 1.00 | 0.68 | 0.53 | 0.80 | 0.85 | 0.41 | 0.94 | 0.60 |
| CELLama | 0.45 | 0.67 | 0.61 | 0.76 | 0.74 | 1.00 | 0.84 | 1.00 | 0.91 | 0.90 | 1.00 | 0.57 | 0.51 | 0.45 | 0.70 | 0.95 | 0.94 | 0.54 |
| scVI | 0.82 | 0.80 | 0.77 | 0.82 | 0.80 | 0.67 | 0.62 | 0.00 | 0.84 | 0.91 | 1.00 | 0.64 | 0.54 | 0.82 | 0.80 | 0.43 | 0.92 | 0.59 |
| PCA | 0.61 | 0.87 | 0.85 | 0.89 | 0.88 | 0.67 | 0.60 | 0.00 | 0.86 | 0.91 | 0.99 | 0.67 | 0.67 | 0.61 | 0.87 | 0.42 | 0.92 | 0.67 |
| scGPT (integrated | 0.80 | 0.85 | 0.84 | 0.90 | 0.88 | 0.44 | 0.47 | 0.00 | 0.74 | 0.82 | 1.00 | 0.48 | 0.42 | 0.80 | 0.87 | 0.31 | 0.85 | 0.45 |
| CellFM | 0.78 | 0.34 | 0.31 | 0.74 | 0.70 | 0.67 | 0.52 | 0.00 | 0.64 | 0.41 | 1.00 | 0.45 | 0.56 | 0.78 | 0.52 | 0.40 | 0.68 | 0.50 |
| GenePT-w | 0.28 | 0.27 | 0.22 | 0.36 | 0.34 | 0.67 | 0.52 | 0.00 | 0.62 | 0.75 | 1.00 | 0.38 | 0.38 | 0.28 | 0.30 | 0.40 | 0.79 | 0.38 |
| scBERT | 0.27 | 0.31 | 0.27 | 0.34 | 0.30 | 0.67 | 0.63 | 0.00 | 0.56 | 0.68 | 1.00 | 0.27 | 0.25 | 0.27 | 0.31 | 0.43 | 0.74 | 0.26 |

**Figure S113: Trajectory-inference metric scores for fibroblast reprogramming using PAGA.** Display and scoring conventions are as in Fig. S103.

| | Cell positions | Neighbourhood | | | | Topology | | | Features | | | Branch assignment | | Aggregated scores | | | | |
|---|---|---|---|---|---|---|---|---|---|---|---|---|---|---|---|---|---|---|
| method | $cor_{dist}$ | $NMSE_{lm}$ | $R^2_{lm}$ | $NMSE_{rf}$ | $R^2_{rf}$ | edgeflip | HIM | isomorphic | $cor_{features}$ | $wcor_{features}$ | $ks_{features}$ | $F1_{branches}$ | $F1_{milestones}$ | Pos. | Neighbor. | Topology | Feature | Branch |
| scFoundation | 0.80 | 0.68 | 0.60 | 0.91 | 0.90 | 1.00 | 0.77 | 1.00 | 0.84 | 0.82 | 0.99 | 0.49 | 0.49 | 0.80 | 0.77 | 0.92 | 0.89 | 0.49 |
| scGPT | 0.81 | 0.83 | 0.80 | 0.91 | 0.89 | 1.00 | 0.75 | 0.00 | 0.95 | 0.95 | 1.00 | 0.54 | 0.54 | 0.81 | 0.86 | 0.58 | 0.97 | 0.54 |
| SCimilarity | 0.57 | 0.71 | 0.63 | 0.90 | 0.88 | 1.00 | 0.74 | 1.00 | 0.91 | 0.93 | 1.00 | 0.46 | 0.52 | 0.57 | 0.78 | 0.91 | 0.95 | 0.49 |
| CellFM | 0.89 | 0.87 | 0.86 | 0.94 | 0.94 | 0.67 | 0.62 | 0.00 | 0.90 | 0.87 | 1.00 | 0.53 | 0.57 | 0.89 | 0.90 | 0.43 | 0.92 | 0.55 |
| UCE | 0.75 | 0.84 | 0.82 | 0.92 | 0.92 | 0.67 | 0.74 | 0.00 | 0.90 | 0.91 | 1.00 | 0.38 | 0.52 | 0.75 | 0.88 | 0.47 | 0.94 | 0.45 |
| LangCell | 0.77 | 0.85 | 0.83 | 0.89 | 0.89 | 0.67 | 0.68 | 0.00 | 0.86 | 0.88 | 1.00 | 0.32 | 0.50 | 0.77 | 0.87 | 0.45 | 0.91 | 0.41 |
| scGPT (integrated | 0.74 | 0.83 | 0.81 | 0.91 | 0.89 | 0.67 | 0.65 | 0.00 | 0.87 | 0.91 | 1.00 | 0.35 | 0.49 | 0.74 | 0.86 | 0.44 | 0.93 | 0.42 |
| scVI (de novo) | 0.74 | 0.83 | 0.81 | 0.91 | 0.91 | 0.67 | 0.61 | 0.00 | 0.82 | 0.73 | 0.95 | 0.44 | 0.51 | 0.74 | 0.86 | 0.43 | 0.84 | 0.48 |
| Geneformer | 0.68 | 0.75 | 0.73 | 0.88 | 0.87 | 0.67 | 0.61 | 0.00 | 0.87 | 0.89 | 1.00 | 0.31 | 0.42 | 0.68 | 0.81 | 0.43 | 0.92 | 0.37 |
| scVI | 0.71 | 0.72 | 0.64 | 0.87 | 0.84 | 0.67 | 0.62 | 0.00 | 0.88 | 0.90 | 0.99 | 0.34 | 0.35 | 0.71 | 0.77 | 0.43 | 0.92 | 0.35 |
| CellPLM | 0.67 | 0.81 | 0.77 | 0.89 | 0.88 | 0.50 | 0.59 | 0.00 | 0.89 | 0.91 | 1.00 | 0.35 | 0.38 | 0.67 | 0.84 | 0.36 | 0.93 | 0.36 |
| scCello | 0.77 | 0.66 | 0.58 | 0.75 | 0.70 | 0.67 | 0.71 | 0.00 | 0.84 | 0.87 | 1.00 | 0.28 | 0.38 | 0.77 | 0.67 | 0.46 | 0.90 | 0.33 |
| PCA | 0.57 | 0.66 | 0.59 | 0.93 | 0.93 | 0.67 | 0.66 | 0.00 | 0.85 | 0.89 | 1.00 | 0.32 | 0.43 | 0.57 | 0.78 | 0.44 | 0.91 | 0.38 |
| CELLama | 0.74 | 0.52 | 0.44 | 0.81 | 0.78 | 1.00 | 0.63 | 0.00 | 0.80 | 0.79 | 1.00 | 0.23 | 0.28 | 0.74 | 0.64 | 0.54 | 0.86 | 0.26 |
| scPRINT | 0.40 | 0.56 | 0.49 | 0.75 | 0.70 | 1.00 | 0.76 | 0.00 | 0.84 | 0.81 | 1.00 | 0.48 | 0.31 | 0.40 | 0.63 | 0.59 | 0.88 | 0.40 |
| GenePT-w | 0.54 | 0.60 | 0.59 | 0.75 | 0.72 | 0.67 | 0.73 | 0.00 | 0.65 | 0.84 | 0.99 | 0.35 | 0.39 | 0.54 | 0.66 | 0.46 | 0.83 | 0.37 |
| scBERT | 0.21 | 0.13 | 0.11 | 0.33 | 0.30 | 0.67 | 0.52 | 0.00 | 0.35 | 0.35 | 1.00 | 0.34 | 0.26 | 0.21 | 0.22 | 0.40 | 0.57 | 0.30 |

**Figure S114: Trajectory-inference metric scores for fibroblast reprogramming using Monocle 3.** Display and scoring conventions are as in Fig. S104.

| | Cell positions | Neighbourhood | | | | Topology | | | Features | | | Branch assignment | | Aggregated scores | | | | |
|---|---|---|---|---|---|---|---|---|---|---|---|---|---|---|---|---|---|---|
| method | $cor_{dist}$ | $NMSE_{lm}$ | $R^2_{lm}$ | $NMSE_{rf}$ | $R^2_{rf}$ | edgeflip | HIM | isomorphic | $cor_{features}$ | $wcor_{features}$ | $ks_{features}$ | $F1_{branches}$ | $F1_{milestones}$ | Pos. | Neighbor. | Topology | Feature | Branch |
| scFoundation | 0.81 | 0.47 | 0.37 | 0.83 | 0.81 | | | | 0.88 | 0.89 | 1.00 | | 0.58 | 0.81 | 0.62 | | 0.92 | 0.58 |
| LangCell | 0.79 | 0.37 | 0.33 | 0.80 | 0.79 | | | | 0.61 | 0.31 | 1.00 | | 0.57 | 0.79 | 0.57 | | 0.64 | 0.57 |
| CELLama | 0.73 | 0.43 | 0.35 | 0.48 | 0.44 | | | | 0.81 | 0.82 | 0.92 | | 0.55 | 0.73 | 0.42 | | 0.85 | 0.55 |
| scGPT (integrated | 0.81 | 0.44 | 0.36 | 0.70 | 0.66 | | | | 0.68 | 0.54 | 1.00 | | 0.43 | 0.81 | 0.54 | | 0.74 | 0.43 |
| PCA | 0.71 | 0.30 | 0.32 | 0.84 | 0.83 | | | | 0.58 | 0.37 | 1.00 | | 0.58 | 0.71 | 0.57 | | 0.65 | 0.58 |
| scCello | 0.75 | 0.35 | 0.34 | 0.73 | 0.70 | | | | 0.70 | 0.60 | 0.99 | | 0.46 | 0.75 | 0.53 | | 0.76 | 0.46 |
| scGPT | 0.79 | 0.44 | 0.35 | 0.80 | 0.77 | | | | 0.70 | 0.58 | 1.00 | | 0.35 | 0.79 | 0.59 | | 0.76 | 0.35 |
| SCimilarity | 0.79 | 0.40 | 0.35 | 0.66 | 0.64 | | | | 0.57 | 0.34 | 1.00 | | 0.54 | 0.79 | 0.51 | | 0.64 | 0.54 |
| UCE | 0.79 | 0.36 | 0.33 | 0.82 | 0.81 | | | | 0.57 | 0.26 | 1.00 | | 0.49 | 0.79 | 0.58 | | 0.61 | 0.49 |
| CellFM | 0.63 | 0.28 | 0.30 | 0.54 | 0.54 | | | | 0.60 | 0.49 | 1.00 | | 0.55 | 0.63 | 0.42 | | 0.70 | 0.55 |
| CellPLM | 0.80 | 0.42 | 0.35 | 0.53 | 0.48 | | | | 0.60 | 0.41 | 1.00 | | 0.36 | 0.80 | 0.44 | | 0.67 | 0.36 |
| scPRINT | 0.68 | 0.39 | 0.31 | 0.51 | 0.45 | | | | 0.66 | 0.50 | 1.00 | | 0.35 | 0.68 | 0.42 | | 0.72 | 0.35 |
| scVI | 0.67 | 0.38 | 0.30 | 0.76 | 0.73 | | | | 0.46 | 0.36 | 1.00 | | 0.33 | 0.67 | 0.54 | | 0.61 | 0.33 |
| scVI (de novo) | 0.74 | 0.37 | 0.32 | 0.56 | 0.54 | | | | 0.52 | 0.23 | 1.00 | | 0.34 | 0.74 | 0.45 | | 0.58 | 0.34 |
| Geneformer | 0.67 | 0.36 | 0.30 | 0.57 | 0.51 | | | | 0.48 | 0.20 | 1.00 | | 0.32 | 0.67 | 0.43 | | 0.56 | 0.32 |
| GenePT-w | 0.29 | 0.26 | 0.20 | 0.06 | 0.00 | | | | 0.68 | 0.76 | 0.79 | | 0.46 | 0.29 | 0.13 | | 0.75 | 0.46 |
| scBERT | 0.14 | 0.10 | 0.09 | 0.00 | 0.00 | | | | 0.25 | 0.05 | 0.99 | | 0.30 | 0.14 | 0.05 | | 0.43 | 0.30 |

**Figure S115: Trajectory-inference metric scores for fibroblast reprogramming using DPT.** Display and scoring conventions are as in Fig. S105.

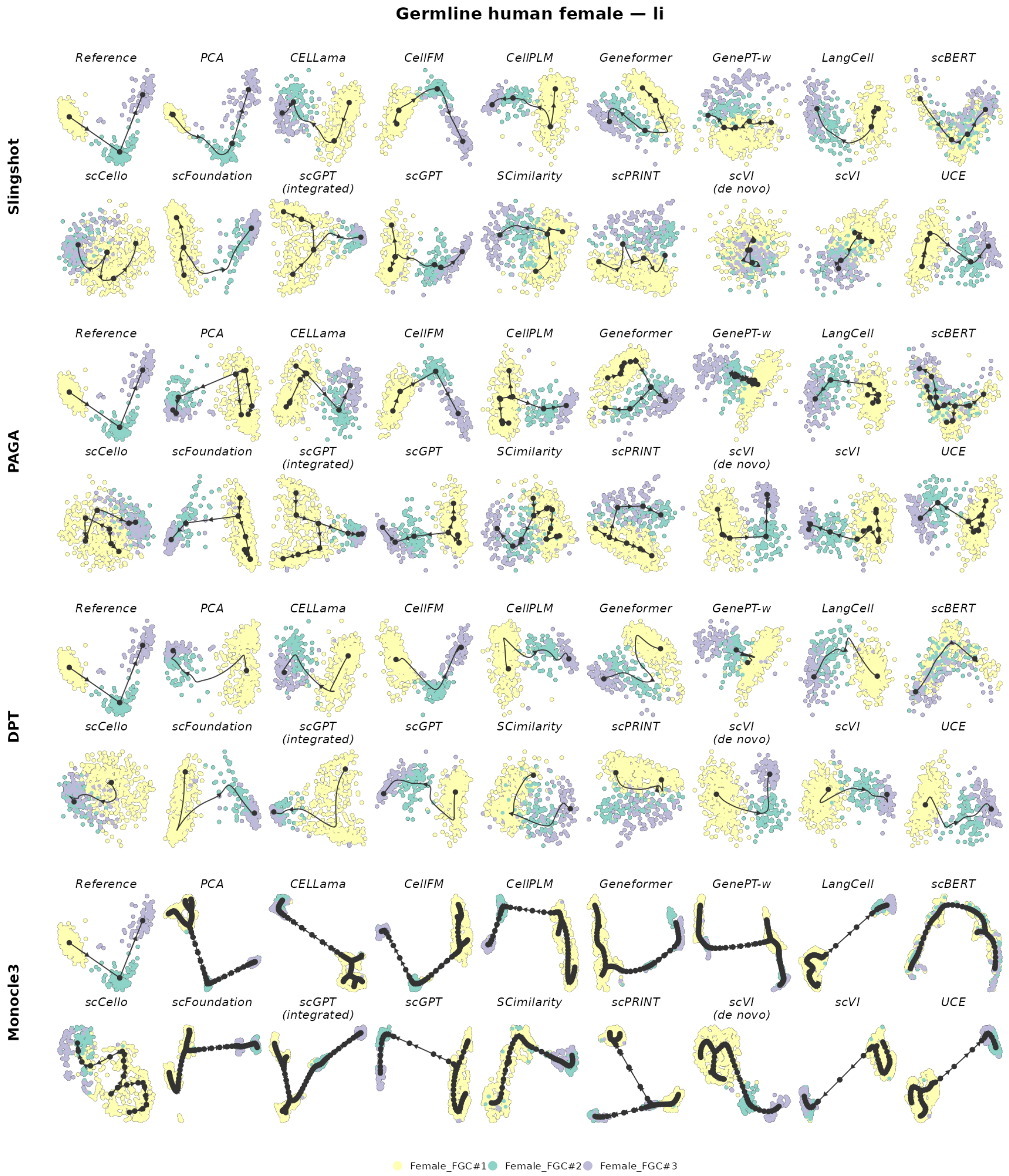

**Figure S116: Trajectory inference on human female germline development.** The reference trajectory is linear. Rows show Slingshot, PAGA, DPT and Monocle 3; within each row, the reference is followed by the same 17 representations. Cells are coloured by the reference milestone or state. Black nodes, edges and arrows show the inferred milestone graph and its orientation. Layouts and trajectories were reconstructed independently for each representation and backend.

| | Cell positions | Neighbourhood | | | | Topology | | | Features | | | Branch assignment | | Aggregated scores | | | | |
|---|---|---|---|---|---|---|---|---|---|---|---|---|---|---|---|---|---|---|
| method | $cor_{dist}$ | $NMSE_{lm}$ | $R^2_{lm}$ | $NMSE_{rf}$ | $R^2_{rf}$ | edgeflip | HIM | isomorphic | $cor_{features}$ | $wcor_{features}$ | $ks_{features}$ | $F1_{branches}$ | $F1_{milestones}$ | Pos. | Neighbor. | Topology | Feature | Branch |
| CellFM | 0.88 | 0.90 | 0.90 | 0.96 | 0.96 | 1.00 | 1.00 | 1.00 | 0.96 | 0.93 | 1.00 | 1.00 | 0.67 | 0.88 | 0.93 | 1.00 | 0.96 | 0.83 |
| scVI (de novo) | 0.86 | 0.86 | 0.86 | 0.95 | 0.95 | 1.00 | 1.00 | 1.00 | 0.95 | 0.92 | 1.00 | 1.00 | 0.65 | 0.86 | 0.90 | 1.00 | 0.96 | 0.83 |
| CELLama | 0.79 | 0.78 | 0.78 | 0.85 | 0.85 | 1.00 | 1.00 | 1.00 | 0.91 | 0.88 | 1.00 | 1.00 | 0.64 | 0.79 | 0.81 | 1.00 | 0.93 | 0.82 |
| LangCell | 0.76 | 0.78 | 0.78 | 0.87 | 0.87 | 1.00 | 1.00 | 1.00 | 0.90 | 0.85 | 0.97 | 1.00 | 0.62 | 0.76 | 0.83 | 1.00 | 0.91 | 0.81 |
| CellPLM | 0.83 | 0.91 | 0.91 | 0.91 | 0.91 | 1.00 | 1.00 | 0.00 | 0.63 | 0.87 | 0.98 | 0.51 | 0.45 | 0.83 | 0.91 | 0.67 | 0.83 | 0.48 |
| UCE | 0.78 | 0.84 | 0.84 | 0.89 | 0.89 | 1.00 | 1.00 | 0.00 | 0.79 | 0.84 | 0.92 | 0.60 | 0.48 | 0.78 | 0.86 | 0.67 | 0.85 | 0.54 |
| PCA | 0.85 | 0.90 | 0.90 | 0.94 | 0.94 | 1.00 | 1.00 | 0.00 | 0.63 | 0.86 | 0.78 | 0.53 | 0.44 | 0.85 | 0.92 | 0.67 | 0.76 | 0.49 |
| Geneformer | 0.80 | 0.82 | 0.82 | 0.84 | 0.84 | 1.00 | 1.00 | 0.00 | 0.82 | 0.91 | 0.63 | 0.52 | 0.49 | 0.80 | 0.83 | 0.67 | 0.79 | 0.51 |
| scGPT | 0.85 | 0.91 | 0.91 | 0.94 | 0.94 | 0.67 | 0.52 | 0.00 | 0.65 | 0.86 | 0.23 | 0.38 | 0.38 | 0.85 | 0.92 | 0.40 | 0.58 | 0.38 |
| scFoundation | 0.77 | 0.86 | 0.86 | 0.86 | 0.87 | 0.67 | 0.53 | 0.00 | 0.65 | 0.84 | 0.64 | 0.38 | 0.38 | 0.77 | 0.86 | 0.40 | 0.71 | 0.38 |
| scVI | 0.84 | 0.91 | 0.91 | 0.90 | 0.90 | 0.67 | 0.49 | 0.00 | 0.71 | 0.91 | 0.01 | 0.40 | 0.41 | 0.84 | 0.90 | 0.38 | 0.55 | 0.41 |
| scGPT (integrated | 0.65 | 0.85 | 0.85 | 0.88 | 0.88 | 0.67 | 0.60 | 0.00 | 0.72 | 0.83 | 0.38 | 0.35 | 0.35 | 0.65 | 0.86 | 0.42 | 0.64 | 0.35 |
| SCimilarity | 0.58 | 0.74 | 0.74 | 0.71 | 0.71 | 0.67 | 0.57 | 0.00 | 0.53 | 0.81 | 0.80 | 0.36 | 0.40 | 0.58 | 0.72 | 0.41 | 0.71 | 0.38 |
| scCello | 0.19 | 0.40 | 0.40 | 0.47 | 0.47 | 1.00 | 1.00 | 0.00 | 0.78 | 0.77 | 0.80 | 0.52 | 0.35 | 0.19 | 0.43 | 0.67 | 0.79 | 0.43 |
| GenePT-w | 0.08 | 0.01 | 0.01 | 0.00 | 0.00 | 1.00 | 1.00 | 1.00 | 0.00 | 0.00 | 0.97 | 1.00 | 0.36 | 0.08 | 0.01 | 1.00 | 0.32 | 0.68 |
| scBERT | 0.06 | 0.03 | 0.03 | 0.00 | 0.00 | 1.00 | 1.00 | 1.00 | 0.00 | 0.00 | 0.73 | 1.00 | 0.40 | 0.06 | 0.02 | 1.00 | 0.24 | 0.70 |
| scPRINT | 0.00 | 0.05 | 0.05 | 0.29 | 0.29 | 1.00 | 1.00 | 0.00 | 0.24 | 0.46 | 0.90 | 0.58 | 0.33 | 0.00 | 0.17 | 0.67 | 0.53 | 0.46 |

**Figure S117: Trajectory-inference metric scores for human female germline development using Slingshot.** Display and scoring conventions are as in Fig. S102.

| | Cell positions | Neighbourhood | | | | Topology | | | Features | | | Branch assignment | | Aggregated scores | | | | |
|---|---|---|---|---|---|---|---|---|---|---|---|---|---|---|---|---|---|---|
| method | $cor_{dist}$ | $NMSE_{lm}$ | $R^2_{lm}$ | $NMSE_{rf}$ | $R^2_{rf}$ | edgeflip | HIM | isomorphic | $cor_{features}$ | $wcor_{features}$ | $ks_{features}$ | $F1_{branches}$ | $F1_{milestones}$ | Pos. | Neighbor. | Topology | Feature | Branch |
| CellFM | 0.82 | 0.82 | 0.82 | 0.98 | 0.98 | 1.00 | 1.00 | 1.00 | 0.96 | 0.94 | 1.00 | 1.00 | 0.66 | 0.82 | 0.90 | 1.00 | 0.96 | 0.83 |
| scVI (de novo) | 0.73 | 0.74 | 0.74 | 0.96 | 0.96 | 1.00 | 1.00 | 1.00 | 0.94 | 0.90 | 1.00 | 1.00 | 0.64 | 0.73 | 0.85 | 1.00 | 0.95 | 0.82 |
| CELLama | 0.61 | 0.94 | 0.94 | 0.93 | 0.93 | 0.67 | 0.61 | 0.00 | 0.91 | 0.91 | 0.46 | 0.44 | 0.56 | 0.61 | 0.93 | 0.42 | 0.76 | 0.50 |
| scVI | 0.58 | 0.91 | 0.91 | 0.91 | 0.91 | 0.67 | 0.66 | 0.00 | 0.67 | 0.89 | 0.43 | 0.38 | 0.54 | 0.58 | 0.91 | 0.44 | 0.66 | 0.46 |
| scGPT | 0.54 | 0.91 | 0.91 | 0.95 | 0.95 | 0.67 | 0.56 | 0.00 | 0.54 | 0.84 | 0.67 | 0.37 | 0.57 | 0.54 | 0.93 | 0.41 | 0.68 | 0.47 |
| scGPT (integrated | 0.57 | 0.92 | 0.92 | 0.94 | 0.94 | 0.67 | 0.57 | 0.00 | 0.64 | 0.85 | 0.33 | 0.35 | 0.41 | 0.57 | 0.93 | 0.41 | 0.60 | 0.38 |
| SCimilarity | 0.45 | 0.81 | 0.81 | 0.85 | 0.85 | 0.67 | 0.63 | 0.00 | 0.56 | 0.82 | 0.54 | 0.42 | 0.55 | 0.45 | 0.83 | 0.43 | 0.64 | 0.49 |
| scCello | 0.39 | 0.46 | 0.46 | 0.75 | 0.76 | 1.00 | 1.00 | 0.00 | 0.83 | 0.80 | 0.33 | 0.59 | 0.43 | 0.39 | 0.61 | 0.67 | 0.66 | 0.51 |
| PCA | 0.50 | 0.91 | 0.91 | 0.93 | 0.93 | 0.67 | 0.63 | 0.00 | 0.56 | 0.83 | 0.03 | 0.27 | 0.49 | 0.50 | 0.92 | 0.43 | 0.47 | 0.38 |
| LangCell | 0.64 | 0.87 | 0.87 | 0.93 | 0.93 | 0.50 | 0.41 | 0.00 | 0.66 | 0.87 | 0.00 | 0.33 | 0.31 | 0.64 | 0.90 | 0.30 | 0.51 | 0.32 |
| UCE | 0.47 | 0.95 | 0.95 | 0.92 | 0.92 | 0.40 | 0.42 | 0.00 | 0.72 | 0.91 | 0.00 | 0.24 | 0.43 | 0.47 | 0.93 | 0.27 | 0.55 | 0.33 |
| scPRINT | 0.46 | 0.52 | 0.52 | 0.78 | 0.78 | 0.67 | 0.63 | 0.00 | 0.81 | 0.79 | 0.27 | 0.40 | 0.30 | 0.46 | 0.65 | 0.43 | 0.63 | 0.35 |
| scFoundation | 0.48 | 0.94 | 0.94 | 0.92 | 0.92 | 0.40 | 0.39 | 0.00 | 0.62 | 0.89 | 0.00 | 0.24 | 0.43 | 0.48 | 0.93 | 0.26 | 0.50 | 0.33 |
| CellPLM | 0.44 | 0.97 | 0.97 | 0.92 | 0.92 | 0.29 | 0.33 | 0.00 | 0.72 | 0.92 | 0.00 | 0.20 | 0.36 | 0.44 | 0.94 | 0.21 | 0.55 | 0.28 |
| Geneformer | 0.42 | 0.93 | 0.93 | 0.93 | 0.93 | 0.29 | 0.33 | 0.00 | 0.66 | 0.69 | 0.00 | 0.18 | 0.34 | 0.42 | 0.93 | 0.21 | 0.45 | 0.26 |
| GenePT-w | 0.00 | 0.24 | 0.24 | 0.50 | 0.50 | 0.40 | 0.47 | 0.00 | 0.22 | 0.49 | 0.95 | 0.29 | 0.22 | 0.00 | 0.37 | 0.29 | 0.55 | 0.26 |
| scBERT | 0.13 | 0.21 | 0.21 | 0.22 | 0.22 | 0.40 | 0.38 | 0.00 | 0.21 | 0.48 | 0.01 | 0.20 | 0.23 | 0.13 | 0.21 | 0.26 | 0.23 | 0.21 |

**Figure S118: Trajectory-inference metric scores for human female germline development using PAGA.** Display and scoring conventions are as in Fig. S103.

| | Cell positions | Neighbourhood | | | | Topology | | | Features | | | Branch assignment | | Aggregated scores | | | | |
|---|---|---|---|---|---|---|---|---|---|---|---|---|---|---|---|---|---|---|
| method | $cor_{dist}$ | $NMSE_{lm}$ | $R^2_{lm}$ | $NMSE_{rf}$ | $R^2_{rf}$ | edgeflip | HIM | isomorphic | $cor_{features}$ | $wcor_{features}$ | $ks_{features}$ | $F1_{branches}$ | $F1_{milestones}$ | Pos. | Neighbor. | Topology | Feature | Branch |
| CellPLM | 0.67 | 0.93 | 0.93 | 0.96 | 0.96 | 0.67 | 0.68 | 0.00 | 0.78 | 0.92 | 0.16 | 0.38 | 0.38 | 0.67 | 0.95 | 0.45 | 0.62 | 0.38 |
| scGPT | 0.66 | 0.91 | 0.91 | 0.93 | 0.93 | 0.67 | 0.62 | 0.00 | 0.68 | 0.86 | 0.20 | 0.32 | 0.33 | 0.66 | 0.92 | 0.43 | 0.58 | 0.32 |
| CellFM | 0.78 | 0.93 | 0.93 | 0.94 | 0.94 | 0.40 | 0.48 | 0.00 | 0.79 | 0.89 | 0.00 | 0.25 | 0.30 | 0.78 | 0.94 | 0.29 | 0.56 | 0.27 |
| scVI (de novo) | 0.67 | 0.88 | 0.88 | 0.95 | 0.95 | 0.40 | 0.51 | 0.00 | 0.89 | 0.91 | 0.11 | 0.30 | 0.30 | 0.67 | 0.91 | 0.30 | 0.64 | 0.30 |
| scVI | 0.61 | 0.86 | 0.86 | 0.86 | 0.86 | 0.67 | 0.73 | 0.00 | 0.79 | 0.85 | 0.05 | 0.33 | 0.32 | 0.61 | 0.86 | 0.47 | 0.56 | 0.33 |
| PCA | 0.63 | 0.97 | 0.97 | 0.97 | 0.97 | 0.40 | 0.45 | 0.00 | 0.77 | 0.88 | 0.04 | 0.23 | 0.40 | 0.63 | 0.97 | 0.28 | 0.56 | 0.31 |
| SCimilarity | 0.61 | 0.59 | 0.59 | 0.81 | 0.81 | 0.67 | 0.59 | 0.00 | 0.54 | 0.77 | 0.80 | 0.30 | 0.26 | 0.61 | 0.70 | 0.42 | 0.70 | 0.28 |
| LangCell | 0.51 | 0.65 | 0.65 | 0.83 | 0.83 | 0.67 | 0.68 | 0.00 | 0.81 | 0.84 | 0.22 | 0.47 | 0.29 | 0.51 | 0.74 | 0.45 | 0.63 | 0.38 |
| UCE | 0.59 | 0.60 | 0.60 | 0.94 | 0.94 | 0.67 | 0.57 | 0.00 | 0.83 | 0.86 | 0.08 | 0.35 | 0.33 | 0.59 | 0.77 | 0.41 | 0.59 | 0.34 |
| Geneformer | 0.67 | 0.81 | 0.81 | 0.88 | 0.88 | 0.40 | 0.48 | 0.00 | 0.77 | 0.87 | 0.00 | 0.27 | 0.31 | 0.67 | 0.85 | 0.29 | 0.55 | 0.29 |
| scGPT (integrated | 0.77 | 0.92 | 0.92 | 0.92 | 0.92 | 0.00 | 0.43 | 0.00 | 0.76 | 0.85 | 0.00 | 0.18 | 0.21 | 0.77 | 0.92 | 0.14 | 0.54 | 0.19 |
| scFoundation | 0.59 | 0.63 | 0.63 | 0.92 | 0.92 | 0.40 | 0.53 | 0.00 | 0.61 | 0.79 | 0.01 | 0.23 | 0.22 | 0.59 | 0.78 | 0.31 | 0.47 | 0.23 |
| CELLama | 0.58 | 0.63 | 0.63 | 0.91 | 0.91 | 0.29 | 0.36 | 0.00 | 0.76 | 0.80 | 0.00 | 0.22 | 0.20 | 0.58 | 0.77 | 0.22 | 0.52 | 0.21 |
| scCello | 0.32 | 0.57 | 0.57 | 0.63 | 0.63 | 0.67 | 0.73 | 0.00 | 0.82 | 0.80 | 0.00 | 0.41 | 0.30 | 0.32 | 0.60 | 0.47 | 0.54 | 0.36 |
| scPRINT | 0.21 | 0.84 | 0.84 | 0.87 | 0.87 | 0.40 | 0.41 | 0.00 | 0.86 | 0.87 | 0.05 | 0.21 | 0.24 | 0.21 | 0.86 | 0.27 | 0.59 | 0.23 |
| GenePT-w | 0.27 | 0.46 | 0.46 | 0.85 | 0.85 | 0.40 | 0.60 | 0.00 | 0.57 | 0.78 | 0.01 | 0.22 | 0.23 | 0.27 | 0.65 | 0.33 | 0.45 | 0.23 |
| scBERT | 0.12 | 0.20 | 0.20 | 0.22 | 0.22 | 0.67 | 0.55 | 0.00 | 0.35 | 0.43 | 1.00 | 0.44 | 0.33 | 0.12 | 0.21 | 0.40 | 0.59 | 0.38 |

**Figure S119: Trajectory-inference metric scores for human female germline development using Monocle 3.** Display and scoring conventions are as in Fig. S104.

| | Cell positions | Neighbourhood | | | | Topology | | | Features | | | Branch assignment | | Aggregated scores | | | | |
|---|---|---|---|---|---|---|---|---|---|---|---|---|---|---|---|---|---|---|
| method | $cor_{dist}$ | $NMSE_{lm}$ | $R^2_{lm}$ | $NMSE_{rf}$ | $R^2_{rf}$ | edgeflip | HIM | isomorphic | $cor_{features}$ | $wcor_{features}$ | $ks_{features}$ | $F1_{branches}$ | $F1_{milestones}$ | Pos. | Neighbor. | Topology | Feature | Branch |
| CellFM | 0.90 | 0.92 | 0.92 | 0.97 | 0.97 | | | | 0.95 | 0.92 | 1.00 | | 0.97 | 0.90 | 0.95 | | 0.96 | 0.97 |
| scVI | 0.85 | 0.90 | 0.90 | 0.88 | 0.88 | | | | 0.95 | 0.90 | 1.00 | | 0.77 | 0.85 | 0.89 | | 0.95 | 0.77 |
| PCA | 0.89 | 0.93 | 0.93 | 0.94 | 0.94 | | | | 0.94 | 0.91 | 1.00 | | 0.65 | 0.89 | 0.94 | | 0.95 | 0.65 |
| CellPLM | 0.89 | 0.94 | 0.94 | 0.96 | 0.96 | | | | 0.96 | 0.93 | 1.00 | | 0.64 | 0.89 | 0.95 | | 0.96 | 0.64 |
| scVI (de novo) | 0.88 | 0.90 | 0.90 | 0.93 | 0.93 | | | | 0.96 | 0.93 | 1.00 | | 0.65 | 0.88 | 0.92 | | 0.96 | 0.65 |
| scGPT (integrated | 0.89 | 0.89 | 0.89 | 0.91 | 0.91 | | | | 0.94 | 0.90 | 1.00 | | 0.65 | 0.89 | 0.90 | | 0.95 | 0.65 |
| UCE | 0.89 | 0.89 | 0.89 | 0.90 | 0.90 | | | | 0.93 | 0.90 | 1.00 | | 0.66 | 0.89 | 0.90 | | 0.94 | 0.66 |
| LangCell | 0.88 | 0.89 | 0.89 | 0.91 | 0.91 | | | | 0.93 | 0.89 | 1.00 | | 0.65 | 0.88 | 0.90 | | 0.94 | 0.65 |
| scGPT | 0.87 | 0.90 | 0.90 | 0.92 | 0.92 | | | | 0.92 | 0.89 | 1.00 | | 0.64 | 0.87 | 0.91 | | 0.94 | 0.64 |
| scFoundation | 0.84 | 0.85 | 0.85 | 0.91 | 0.91 | | | | 0.94 | 0.91 | 1.00 | | 0.64 | 0.84 | 0.88 | | 0.95 | 0.64 |
| Geneformer | 0.82 | 0.84 | 0.84 | 0.87 | 0.87 | | | | 0.96 | 0.94 | 1.00 | | 0.62 | 0.82 | 0.86 | | 0.97 | 0.62 |
| CELLama | 0.81 | 0.82 | 0.82 | 0.87 | 0.87 | | | | 0.91 | 0.88 | 1.00 | | 0.65 | 0.81 | 0.84 | | 0.93 | 0.65 |
| scCello | 0.62 | 0.62 | 0.62 | 0.58 | 0.58 | | | | 0.90 | 0.86 | 1.00 | | 0.60 | 0.62 | 0.60 | | 0.92 | 0.60 |
| SCimilarity | 0.47 | 0.47 | 0.47 | 0.50 | 0.50 | | | | 0.58 | 0.31 | 1.00 | | 0.68 | 0.47 | 0.48 | | 0.63 | 0.68 |
| GenePT-w | 0.04 | 0.01 | 0.01 | 0.00 | 0.00 | | | | 0.00 | 0.00 | 1.00 | | 0.43 | 0.04 | 0.00 | | 0.33 | 0.43 |
| scBERT | 0.06 | 0.03 | 0.03 | 0.00 | 0.00 | | | | 0.00 | 0.00 | 0.67 | | 0.42 | 0.06 | 0.01 | | 0.22 | 0.42 |
| scPRINT | 0.00 | 0.00 | 0.00 | 0.00 | 0.00 | | | | 0.09 | 0.22 | 0.38 | | 0.43 | 0.00 | 0.00 | | 0.23 | 0.43 |

**Figure S120: Trajectory-inference metric scores for human female germline development using DPT.** Display and scoring conventions are as in Fig. S105.

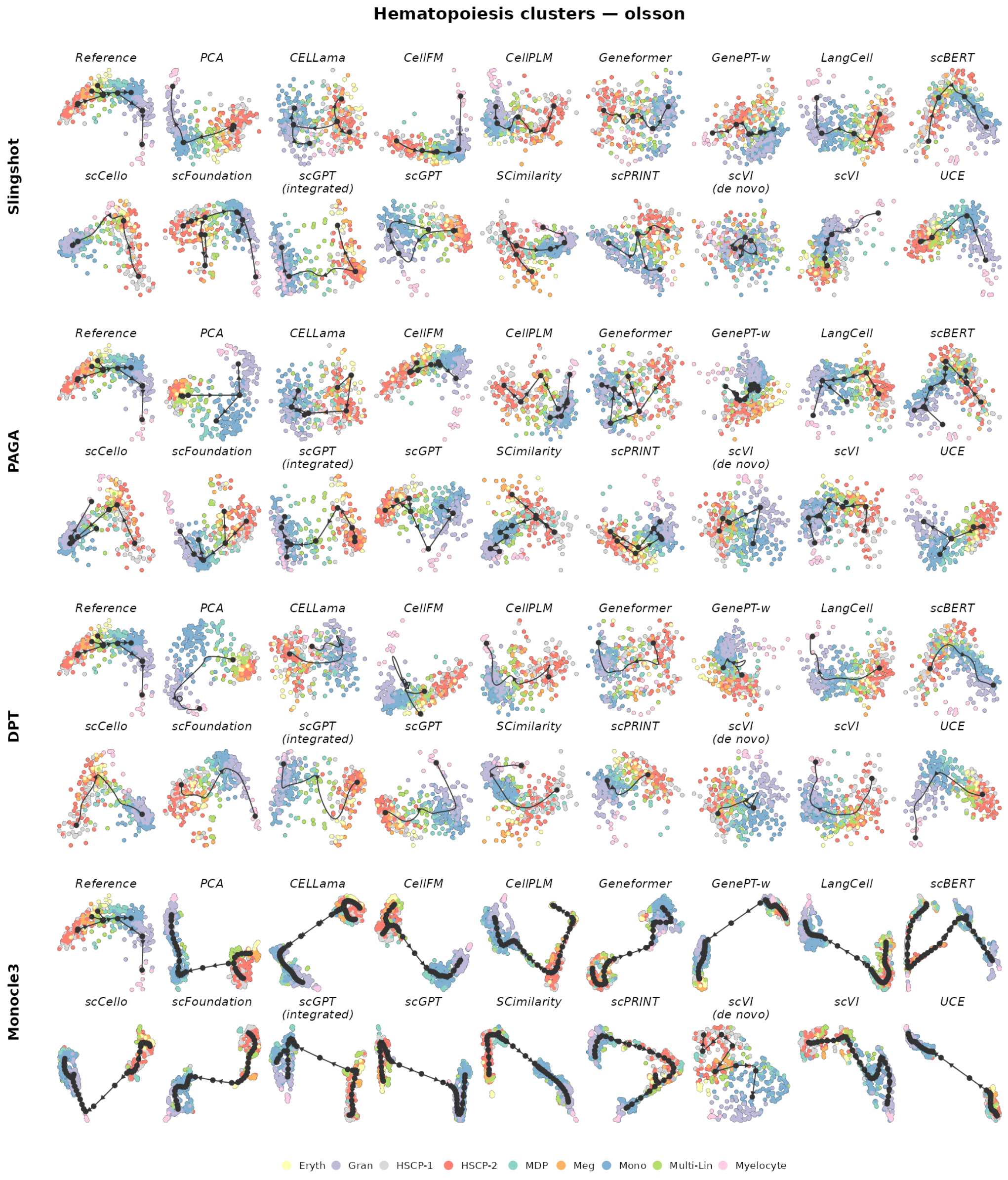

**Figure S121: Trajectory inference on haematopoiesis.** The reference trajectory is tree-structured. Rows show Slingshot, PAGA, DPT and Monocle 3; within each row, the reference is followed by the same 17 representations. Cells are coloured by the reference milestone or state. Black nodes, edges and arrows show the inferred milestone graph and its orientation. Layouts and trajectories were reconstructed independently for each representation and backend.

| | Cell positions | Neighbourhood | | | | Topology | | | Features | | | Branch assignment | | Aggregated scores | | | | |
|---|---|---|---|---|---|---|---|---|---|---|---|---|---|---|---|---|---|---|
| method | $cor_{dist}$ | $NMSE_{lm}$ | $R^2_{lm}$ | $NMSE_{rf}$ | $R^2_{rf}$ | edgeflip | HIM | isomorphic | $cor_{features}$ | $wcor_{features}$ | $ks_{features}$ | $F1_{branches}$ | $F1_{milestones}$ | Pos. | Neighbor. | Topology | Feature | Branch |
| scFoundation | 0.55 | 0.37 | 0.37 | 0.49 | 0.50 | 0.89 | 0.85 | 0.00 | 0.83 | 0.91 | 1.00 | 0.34 | 0.40 | 0.55 | 0.44 | 0.58 | 0.92 | 0.37 |
| CELLama | 0.59 | 0.31 | 0.27 | 0.33 | 0.29 | 0.89 | 0.87 | 0.00 | 0.81 | 0.86 | 1.00 | 0.35 | 0.27 | 0.59 | 0.30 | 0.59 | 0.89 | 0.31 |
| CellFM | 0.63 | 0.31 | 0.30 | 0.45 | 0.42 | 0.57 | 0.55 | 0.00 | 0.79 | 0.87 | 1.00 | 0.38 | 0.36 | 0.63 | 0.37 | 0.37 | 0.89 | 0.37 |
| UCE | 0.74 | 0.24 | 0.20 | 0.34 | 0.29 | 0.33 | 0.34 | 0.00 | 0.84 | 0.88 | 1.00 | 0.23 | 0.36 | 0.74 | 0.27 | 0.22 | 0.91 | 0.30 |
| SCimilarity | 0.61 | 0.23 | 0.25 | 0.44 | 0.46 | 0.33 | 0.34 | 0.00 | 0.81 | 0.88 | 1.00 | 0.29 | 0.33 | 0.61 | 0.34 | 0.22 | 0.89 | 0.31 |
| scGPT (integrated | 0.67 | 0.27 | 0.22 | 0.35 | 0.29 | 0.33 | 0.34 | 0.00 | 0.78 | 0.88 | 1.00 | 0.26 | 0.37 | 0.67 | 0.28 | 0.22 | 0.89 | 0.31 |
| PCA | 0.66 | 0.17 | 0.19 | 0.58 | 0.51 | 0.33 | 0.34 | 0.00 | 0.77 | 0.81 | 1.00 | 0.23 | 0.29 | 0.66 | 0.36 | 0.22 | 0.86 | 0.26 |
| scVI (de novo) | 0.52 | 0.27 | 0.24 | 0.22 | 0.18 | 0.57 | 0.51 | 0.00 | 0.83 | 0.88 | 1.00 | 0.32 | 0.31 | 0.52 | 0.23 | 0.36 | 0.90 | 0.31 |
| LangCell | 0.68 | 0.21 | 0.18 | 0.25 | 0.16 | 0.33 | 0.34 | 0.00 | 0.82 | 0.87 | 1.00 | 0.23 | 0.33 | 0.68 | 0.20 | 0.22 | 0.89 | 0.28 |
| scCello | 0.66 | 0.25 | 0.17 | 0.22 | 0.10 | 0.33 | 0.34 | 0.00 | 0.84 | 0.90 | 1.00 | 0.23 | 0.37 | 0.66 | 0.18 | 0.22 | 0.91 | 0.30 |
| CellPLM | 0.65 | 0.23 | 0.18 | 0.21 | 0.14 | 0.33 | 0.34 | 0.00 | 0.80 | 0.88 | 1.00 | 0.23 | 0.37 | 0.65 | 0.19 | 0.22 | 0.89 | 0.30 |
| scVI | 0.56 | 0.15 | 0.16 | 0.14 | 0.06 | 0.33 | 0.34 | 0.00 | 0.65 | 0.67 | 1.00 | 0.23 | 0.25 | 0.56 | 0.13 | 0.22 | 0.77 | 0.24 |
| scPRINT | 0.30 | 0.17 | 0.13 | 0.12 | 0.06 | 0.57 | 0.51 | 0.00 | 0.76 | 0.85 | 1.00 | 0.31 | 0.26 | 0.30 | 0.12 | 0.36 | 0.87 | 0.28 |
| Geneformer | 0.49 | 0.19 | 0.12 | 0.00 | 0.00 | 0.33 | 0.34 | 0.00 | 0.73 | 0.85 | 1.00 | 0.23 | 0.31 | 0.49 | 0.08 | 0.22 | 0.86 | 0.27 |
| scGPT | 0.35 | 0.11 | 0.09 | 0.29 | 0.26 | 0.33 | 0.34 | 0.00 | 0.70 | 0.72 | 1.00 | 0.23 | 0.31 | 0.35 | 0.19 | 0.22 | 0.81 | 0.27 |
| scBERT | 0.34 | 0.13 | 0.09 | 0.00 | 0.00 | 0.33 | 0.34 | 0.00 | 0.64 | 0.77 | 1.00 | 0.23 | 0.30 | 0.34 | 0.05 | 0.22 | 0.80 | 0.27 |
| GenePT-w | 0.17 | 0.09 | 0.07 | 0.05 | 0.02 | 0.33 | 0.34 | 0.00 | 0.26 | 0.22 | 1.00 | 0.20 | 0.27 | 0.17 | 0.06 | 0.22 | 0.49 | 0.23 |

**Figure S122: Trajectory-inference metric scores for haematopoiesis using Slingshot.** Display and scoring conventions are as in Fig. S102.

| | Cell positions | Neighbourhood | | | | Topology | | | Features | | | Branch assignment | | Aggregated scores | | | | |
|---|---|---|---|---|---|---|---|---|---|---|---|---|---|---|---|---|---|---|
| method | $cor_{dist}$ | $NMSE_{lm}$ | $R^2_{lm}$ | $NMSE_{rf}$ | $R^2_{rf}$ | edgeflip | HIM | isomorphic | $cor_{features}$ | $wcor_{features}$ | $ks_{features}$ | $F1_{branches}$ | $F1_{milestones}$ | Pos. | Neighbor. | Topology | Feature | Branch |
| scFoundation | 0.60 | 0.38 | 0.36 | 0.36 | 0.33 | 0.89 | 0.87 | 0.00 | 0.80 | 0.91 | 1.00 | 0.46 | 0.32 | 0.60 | 0.36 | 0.58 | 0.90 | 0.39 |
| scCello | 0.70 | 0.38 | 0.32 | 0.45 | 0.39 | 0.57 | 0.54 | 0.00 | 0.92 | 0.94 | 1.00 | 0.40 | 0.42 | 0.70 | 0.38 | 0.37 | 0.95 | 0.41 |
| scVI | 0.62 | 0.34 | 0.29 | 0.34 | 0.30 | 0.89 | 0.88 | 0.00 | 0.83 | 0.90 | 1.00 | 0.36 | 0.35 | 0.62 | 0.32 | 0.59 | 0.91 | 0.35 |
| scGPT | 0.64 | 0.34 | 0.28 | 0.38 | 0.35 | 0.57 | 0.54 | 0.00 | 0.87 | 0.91 | 0.99 | 0.44 | 0.43 | 0.64 | 0.34 | 0.37 | 0.92 | 0.44 |
| UCE | 0.57 | 0.40 | 0.37 | 0.38 | 0.35 | 0.57 | 0.52 | 0.00 | 0.88 | 0.92 | 1.00 | 0.43 | 0.39 | 0.57 | 0.37 | 0.37 | 0.93 | 0.41 |
| Geneformer | 0.62 | 0.28 | 0.24 | 0.26 | 0.22 | 0.89 | 0.85 | 0.00 | 0.75 | 0.86 | 1.00 | 0.31 | 0.30 | 0.62 | 0.25 | 0.58 | 0.87 | 0.31 |
| scGPT (integrated | 0.64 | 0.33 | 0.28 | 0.45 | 0.42 | 0.57 | 0.54 | 0.00 | 0.83 | 0.91 | 1.00 | 0.27 | 0.33 | 0.64 | 0.37 | 0.37 | 0.92 | 0.30 |
| CellPLM | 0.63 | 0.30 | 0.26 | 0.35 | 0.36 | 0.57 | 0.54 | 0.00 | 0.72 | 0.86 | 1.00 | 0.26 | 0.40 | 0.63 | 0.32 | 0.37 | 0.86 | 0.33 |
| PCA | 0.45 | 0.29 | 0.31 | 0.31 | 0.38 | 0.57 | 0.54 | 0.00 | 0.83 | 0.85 | 1.00 | 0.37 | 0.43 | 0.45 | 0.32 | 0.37 | 0.89 | 0.40 |
| LangCell | 0.57 | 0.25 | 0.22 | 0.28 | 0.24 | 0.57 | 0.54 | 0.00 | 0.85 | 0.88 | 0.97 | 0.22 | 0.40 | 0.57 | 0.25 | 0.37 | 0.90 | 0.31 |
| SCimilarity | 0.57 | 0.21 | 0.24 | 0.44 | 0.42 | 0.33 | 0.34 | 0.00 | 0.83 | 0.87 | 1.00 | 0.29 | 0.38 | 0.57 | 0.33 | 0.22 | 0.90 | 0.33 |
| CELLama | 0.47 | 0.22 | 0.20 | 0.29 | 0.32 | 0.57 | 0.54 | 0.00 | 0.83 | 0.88 | 1.00 | 0.33 | 0.31 | 0.47 | 0.26 | 0.37 | 0.90 | 0.32 |
| scBERT | 0.44 | 0.21 | 0.19 | 0.23 | 0.20 | 0.89 | 0.73 | 0.00 | 0.67 | 0.79 | 1.00 | 0.23 | 0.33 | 0.44 | 0.21 | 0.54 | 0.82 | 0.28 |
| scVI (de novo) | 0.53 | 0.21 | 0.14 | 0.35 | 0.30 | 0.33 | 0.34 | 0.00 | 0.87 | 0.90 | 1.00 | 0.23 | 0.34 | 0.53 | 0.25 | 0.22 | 0.92 | 0.29 |
| CellFM | 0.37 | 0.15 | 0.20 | 0.24 | 0.28 | 0.57 | 0.54 | 0.00 | 0.76 | 0.79 | 0.98 | 0.28 | 0.29 | 0.37 | 0.22 | 0.37 | 0.84 | 0.28 |
| scPRINT | 0.19 | 0.12 | 0.09 | 0.23 | 0.18 | 0.33 | 0.34 | 0.00 | 0.77 | 0.86 | 1.00 | 0.27 | 0.23 | 0.19 | 0.15 | 0.22 | 0.88 | 0.25 |
| GenePT-w | 0.17 | 0.09 | 0.07 | 0.09 | 0.08 | 0.33 | 0.34 | 0.00 | 0.27 | 0.23 | 1.00 | 0.20 | 0.25 | 0.17 | 0.08 | 0.22 | 0.50 | 0.22 |

**Figure S123: Trajectory-inference metric scores for haematopoiesis using PAGA.** Display and scoring conventions are as in Fig. S103.

| | Cell positions | Neighbourhood | | | | Topology | | | Features | | | Branch assignment | | Aggregated scores | | | | |
|---|---|---|---|---|---|---|---|---|---|---|---|---|---|---|---|---|---|---|
| method | $cor_{dist}$ | $NMSE_{lm}$ | $R^2_{lm}$ | $NMSE_{rf}$ | $R^2_{rf}$ | edgeflip | HIM | isomorphic | $cor_{features}$ | $wcor_{features}$ | $ks_{features}$ | $F1_{branches}$ | $F1_{milestones}$ | Pos. | Neighbor. | Topology | Feature | Branch |
| PCA | 0.69 | 0.38 | 0.35 | 0.52 | 0.47 | 0.57 | 0.54 | 0.00 | 0.87 | 0.92 | 1.00 | 0.31 | 0.31 | 0.69 | 0.43 | 0.37 | 0.93 | 0.31 |
| scGPT (integrated) | 0.71 | 0.34 | 0.28 | 0.36 | 0.32 | 0.57 | 0.53 | 0.00 | 0.87 | 0.93 | 1.00 | 0.32 | 0.27 | 0.71 | 0.33 | 0.37 | 0.93 | 0.30 |
| scCello | 0.58 | 0.31 | 0.26 | 0.50 | 0.49 | 0.57 | 0.48 | 0.00 | 0.90 | 0.94 | 1.00 | 0.38 | 0.30 | 0.58 | 0.39 | 0.35 | 0.95 | 0.34 |
| CellFM | 0.67 | 0.29 | 0.28 | 0.47 | 0.41 | 0.57 | 0.54 | 0.00 | 0.83 | 0.87 | 0.99 | 0.27 | 0.28 | 0.67 | 0.36 | 0.37 | 0.90 | 0.27 |
| SCimilarity | 0.58 | 0.29 | 0.27 | 0.40 | 0.35 | 0.57 | 0.47 | 0.00 | 0.86 | 0.89 | 1.00 | 0.27 | 0.26 | 0.58 | 0.33 | 0.35 | 0.92 | 0.27 |
| scGPT | 0.57 | 0.28 | 0.22 | 0.43 | 0.40 | 0.57 | 0.41 | 0.00 | 0.85 | 0.87 | 1.00 | 0.31 | 0.29 | 0.57 | 0.33 | 0.33 | 0.91 | 0.30 |
| CellPLM | 0.59 | 0.27 | 0.27 | 0.38 | 0.39 | 0.57 | 0.42 | 0.00 | 0.74 | 0.88 | 1.00 | 0.23 | 0.28 | 0.59 | 0.33 | 0.33 | 0.87 | 0.26 |
| scFoundation | 0.62 | 0.21 | 0.17 | 0.44 | 0.47 | 0.33 | 0.34 | 0.00 | 0.82 | 0.89 | 1.00 | 0.23 | 0.36 | 0.62 | 0.32 | 0.22 | 0.90 | 0.30 |
| LangCell | 0.59 | 0.23 | 0.20 | 0.36 | 0.29 | 0.33 | 0.34 | 0.00 | 0.85 | 0.91 | 1.00 | 0.34 | 0.33 | 0.59 | 0.27 | 0.22 | 0.92 | 0.34 |
| UCE | 0.69 | 0.22 | 0.19 | 0.37 | 0.34 | 0.33 | 0.34 | 0.00 | 0.83 | 0.87 | 1.00 | 0.19 | 0.31 | 0.69 | 0.28 | 0.22 | 0.90 | 0.25 |
| CELLama | 0.55 | 0.21 | 0.18 | 0.36 | 0.34 | 0.57 | 0.53 | 0.00 | 0.82 | 0.86 | 0.93 | 0.26 | 0.23 | 0.55 | 0.27 | 0.37 | 0.87 | 0.24 |
| scVI (de novo) | 0.60 | 0.24 | 0.18 | 0.28 | 0.23 | 0.33 | 0.34 | 0.00 | 0.85 | 0.89 | 1.00 | 0.23 | 0.30 | 0.60 | 0.23 | 0.22 | 0.91 | 0.27 |
| scVI | 0.59 | 0.19 | 0.17 | 0.37 | 0.32 | 0.33 | 0.34 | 0.00 | 0.81 | 0.86 | 1.00 | 0.20 | 0.28 | 0.59 | 0.26 | 0.22 | 0.89 | 0.24 |
| Geneformer | 0.47 | 0.21 | 0.14 | 0.29 | 0.23 | 0.33 | 0.34 | 0.00 | 0.79 | 0.88 | 0.99 | 0.27 | 0.28 | 0.47 | 0.22 | 0.22 | 0.89 | 0.27 |
| scPRINT | 0.22 | 0.23 | 0.19 | 0.31 | 0.28 | 0.57 | 0.53 | 0.00 | 0.80 | 0.86 | 1.00 | 0.27 | 0.22 | 0.22 | 0.25 | 0.37 | 0.89 | 0.25 |
| scBERT | 0.33 | 0.18 | 0.14 | 0.26 | 0.20 | 0.57 | 0.47 | 0.00 | 0.67 | 0.79 | 1.00 | 0.27 | 0.23 | 0.33 | 0.19 | 0.35 | 0.82 | 0.25 |
| GenePT-w | 0.24 | 0.21 | 0.15 | 0.28 | 0.23 | 0.57 | 0.48 | 0.00 | 0.65 | 0.82 | 0.99 | 0.18 | 0.23 | 0.24 | 0.22 | 0.35 | 0.82 | 0.20 |

**Figure S124: Trajectory-inference metric scores for haematopoiesis using Monocle 3.** Display and scoring conventions are as in Fig. S104.

| | Cell positions | Neighbourhood | | | | Topology | | | Features | | | Branch assignment | | Aggregated scores | | | | |
|---|---|---|---|---|---|---|---|---|---|---|---|---|---|---|---|---|---|---|
| method | $cor_{dist}$ | $NMSE_{lm}$ | $R^2_{lm}$ | $NMSE_{rf}$ | $R^2_{rf}$ | edgeflip | HIM | isomorphic | $cor_{features}$ | $wcor_{features}$ | $ks_{features}$ | $F1_{branches}$ | $F1_{milestones}$ | Pos. | Neighbor. | Topology | Feature | Branch |
| scGPT (integrated) | 0.79 | 0.26 | 0.20 | 0.23 | 0.20 | | | | 0.83 | 0.88 | 1.00 | | 0.37 | 0.79 | 0.23 | | 0.91 | 0.37 |
| scGPT | 0.74 | 0.25 | 0.19 | 0.21 | 0.16 | | | | 0.83 | 0.89 | 1.00 | | 0.36 | 0.74 | 0.20 | | 0.91 | 0.36 |
| scCello | 0.67 | 0.24 | 0.18 | 0.14 | 0.05 | | | | 0.85 | 0.90 | 1.00 | | 0.36 | 0.67 | 0.15 | | 0.91 | 0.36 |
| scFoundation | 0.67 | 0.20 | 0.18 | 0.33 | 0.26 | | | | 0.81 | 0.85 | 1.00 | | 0.29 | 0.67 | 0.24 | | 0.89 | 0.29 |
| UCE | 0.70 | 0.21 | 0.20 | 0.25 | 0.22 | | | | 0.79 | 0.82 | 1.00 | | 0.29 | 0.70 | 0.22 | | 0.87 | 0.29 |
| LangCell | 0.61 | 0.17 | 0.17 | 0.23 | 0.11 | | | | 0.73 | 0.78 | 1.00 | | 0.29 | 0.61 | 0.17 | | 0.84 | 0.29 |
| SCimilarity | 0.60 | 0.18 | 0.16 | 0.17 | 0.11 | | | | 0.80 | 0.84 | 1.00 | | 0.24 | 0.60 | 0.16 | | 0.88 | 0.24 |
| CellPLM | 0.58 | 0.20 | 0.16 | 0.04 | 0.00 | | | | 0.70 | 0.84 | 1.00 | | 0.33 | 0.58 | 0.10 | | 0.85 | 0.33 |
| scVI | 0.57 | 0.18 | 0.16 | 0.15 | 0.09 | | | | 0.79 | 0.85 | 1.00 | | 0.25 | 0.57 | 0.14 | | 0.88 | 0.25 |
| Geneformer | 0.51 | 0.20 | 0.14 | 0.00 | 0.00 | | | | 0.78 | 0.86 | 0.99 | | 0.33 | 0.51 | 0.08 | | 0.88 | 0.33 |
| CELLama | 0.46 | 0.13 | 0.14 | 0.03 | 0.00 | | | | 0.60 | 0.60 | 1.00 | | 0.27 | 0.46 | 0.08 | | 0.73 | 0.27 |
| scBERT | 0.34 | 0.13 | 0.10 | 0.00 | 0.00 | | | | 0.65 | 0.76 | 1.00 | | 0.28 | 0.34 | 0.06 | | 0.80 | 0.28 |
| scVI (de novo) | 0.34 | 0.12 | 0.10 | 0.00 | 0.00 | | | | 0.72 | 0.73 | 1.00 | | 0.26 | 0.34 | 0.05 | | 0.82 | 0.26 |
| scPRINT | 0.19 | 0.10 | 0.08 | 0.00 | 0.00 | | | | 0.74 | 0.83 | 1.00 | | 0.21 | 0.19 | 0.04 | | 0.86 | 0.21 |
| PCA | 0.30 | 0.11 | 0.14 | 0.00 | 0.06 | | | | 0.45 | 0.36 | 1.00 | | 0.26 | 0.30 | 0.08 | | 0.60 | 0.26 |
| CellFM | 0.18 | 0.05 | 0.07 | 0.00 | 0.00 | | | | 0.37 | 0.24 | 1.00 | | 0.22 | 0.18 | 0.03 | | 0.54 | 0.22 |
| GenePT-w | 0.16 | 0.07 | 0.04 | 0.00 | 0.00 | | | | 0.18 | 0.13 | 1.00 | | 0.25 | 0.16 | 0.03 | | 0.44 | 0.25 |

**Figure S125: Trajectory-inference metric scores for haematopoiesis using DPT.** Display and scoring conventions are as in Fig. S105.

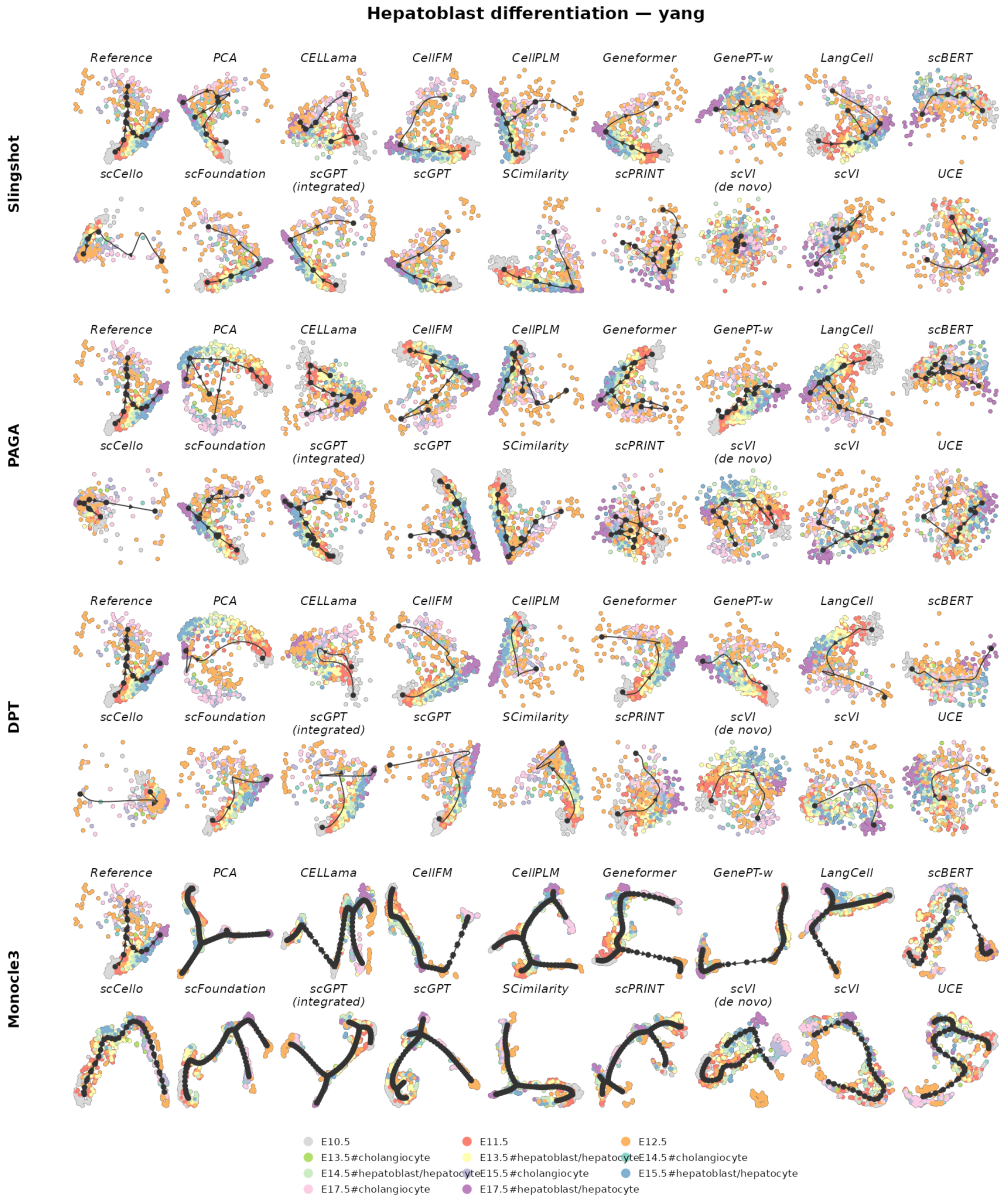

**Figure S126: Trajectory inference on hepatoblast differentiation.** The reference trajectory is bifurcating. Rows show Slingshot, PAGA, DPT and Monocle 3; within each row, the reference is followed by the same 17 representations. Cells are coloured by the reference milestone or state. Black nodes, edges and arrows show the inferred milestone graph and its orientation. Layouts and trajectories were reconstructed independently for each representation and backend.

| | Cell positions | Neighbourhood | | | | Topology | | | Features | | | Branch assignment | | Aggregated scores | | | | |
|---|---|---|---|---|---|---|---|---|---|---|---|---|---|---|---|---|---|---|
| **method** | $cor_{dist}$ | $NMSE_{lm}$ | $R^2_{lm}$ | $NMSE_{rf}$ | $R^2_{rf}$ | edgeflip | HIM | isomorphic | $cor_{features}$ | $wcor_{features}$ | $ks_{features}$ | $F1_{branches}$ | $F1_{milestones}$ | Pos. | Neighbor. | Topology | Feature | Branch |
| **LangCell** | 0.34 | 0.45 | 0.45 | 0.62 | 0.62 | 1.00 | 0.84 | 1.00 | 0.73 | 0.56 | 1.00 | 0.38 | 0.48 | 0.34 | 0.54 | 0.95 | 0.76 | 0.43 |
| **PCA** | 0.37 | 0.46 | 0.48 | 0.69 | 0.72 | 0.80 | 0.73 | 0.00 | 0.75 | 0.58 | 1.00 | 0.37 | 0.38 | 0.37 | 0.59 | 0.51 | 0.78 | 0.37 |
| **CellPLM** | 0.20 | 0.33 | 0.32 | 0.46 | 0.46 | 1.00 | 0.84 | 1.00 | 0.50 | 0.47 | 1.00 | 0.42 | 0.37 | 0.20 | 0.39 | 0.95 | 0.66 | 0.39 |
| **scVI** | 0.32 | 0.23 | 0.25 | 0.36 | 0.37 | 1.00 | 0.72 | 1.00 | 0.36 | 0.26 | 1.00 | 0.33 | 0.33 | 0.32 | 0.30 | 0.91 | 0.54 | 0.33 |
| **Geneformer** | 0.33 | 0.19 | 0.19 | 0.53 | 0.53 | 0.67 | 0.62 | 0.00 | 0.53 | 0.41 | 1.00 | 0.39 | 0.36 | 0.33 | 0.36 | 0.43 | 0.65 | 0.37 |
| **scGPT (integrated** | 0.37 | 0.15 | 0.16 | 0.46 | 0.45 | 0.67 | 0.62 | 0.00 | 0.46 | 0.32 | 1.00 | 0.39 | 0.33 | 0.37 | 0.30 | 0.43 | 0.59 | 0.36 |
| **SCimilarity** | 0.37 | 0.15 | 0.16 | 0.45 | 0.45 | 0.67 | 0.62 | 0.00 | 0.46 | 0.31 | 1.00 | 0.39 | 0.34 | 0.37 | 0.30 | 0.43 | 0.59 | 0.36 |
| **scFoundation** | 0.34 | 0.16 | 0.17 | 0.41 | 0.42 | 0.67 | 0.62 | 0.00 | 0.51 | 0.36 | 1.00 | 0.39 | 0.34 | 0.34 | 0.29 | 0.43 | 0.62 | 0.36 |
| **scVI (de novo)** | 0.30 | 0.20 | 0.21 | 0.48 | 0.50 | 0.67 | 0.62 | 0.00 | 0.53 | 0.37 | 1.00 | 0.30 | 0.33 | 0.30 | 0.35 | 0.43 | 0.63 | 0.31 |
| **CellFM** | 0.35 | 0.15 | 0.16 | 0.52 | 0.54 | 0.67 | 0.62 | 0.00 | 0.40 | 0.17 | 1.00 | 0.39 | 0.34 | 0.35 | 0.34 | 0.43 | 0.52 | 0.37 |
| **scGPT** | 0.29 | 0.16 | 0.16 | 0.39 | 0.39 | 0.67 | 0.62 | 0.00 | 0.46 | 0.34 | 1.00 | 0.39 | 0.34 | 0.29 | 0.28 | 0.43 | 0.60 | 0.36 |
| **CELLama** | 0.11 | 0.25 | 0.24 | 0.45 | 0.44 | 0.67 | 0.62 | 0.00 | 0.63 | 0.48 | 1.00 | 0.30 | 0.32 | 0.11 | 0.34 | 0.43 | 0.70 | 0.31 |
| **scCello** | 0.27 | 0.09 | 0.10 | 0.32 | 0.33 | 0.67 | 0.62 | 0.00 | 0.53 | 0.34 | 1.00 | 0.35 | 0.28 | 0.27 | 0.21 | 0.43 | 0.63 | 0.31 |
| **scPRINT** | 0.06 | 0.20 | 0.20 | 0.35 | 0.33 | 1.00 | 0.73 | 0.00 | 0.64 | 0.44 | 1.00 | 0.25 | 0.25 | 0.06 | 0.27 | 0.58 | 0.69 | 0.25 |
| **GenePT-w** | 0.30 | 0.16 | 0.16 | 0.11 | 0.11 | 0.67 | 0.62 | 0.00 | 0.48 | 0.30 | 1.00 | 0.39 | 0.33 | 0.30 | 0.13 | 0.43 | 0.59 | 0.36 |
| **UCE** | 0.22 | 0.14 | 0.15 | 0.33 | 0.36 | 0.67 | 0.62 | 0.00 | 0.39 | 0.09 | 1.00 | 0.31 | 0.31 | 0.22 | 0.24 | 0.43 | 0.49 | 0.31 |
| **scBERT** | 0.20 | 0.12 | 0.12 | 0.06 | 0.05 | 0.67 | 0.62 | 0.00 | 0.48 | 0.33 | 1.00 | 0.39 | 0.32 | 0.20 | 0.09 | 0.43 | 0.61 | 0.35 |

**Figure S127: Trajectory-inference metric scores for hepatoblast differentiation using Slingshot.** Display and scoring conventions are as in Fig. S102.

| | Cell positions | Neighbourhood | | | | Topology | | | Features | | | Branch assignment | | Aggregated scores | | | | |
|---|---|---|---|---|---|---|---|---|---|---|---|---|---|---|---|---|---|---|
| **method** | $cor_{dist}$ | $NMSE_{lm}$ | $R^2_{lm}$ | $NMSE_{rf}$ | $R^2_{rf}$ | edgeflip | HIM | isomorphic | $cor_{features}$ | $wcor_{features}$ | $ks_{features}$ | $F1_{branches}$ | $F1_{milestones}$ | Pos. | Neighbor. | Topology | Feature | Branch |
| **CellFM** | 0.52 | 0.50 | 0.53 | 0.64 | 0.67 | 1.00 | 0.94 | 1.00 | 0.63 | 0.46 | 1.00 | 0.42 | 0.52 | 0.52 | 0.58 | 0.98 | 0.70 | 0.47 |
| **PCA** | 0.43 | 0.66 | 0.67 | 0.69 | 0.70 | 0.67 | 0.63 | 0.00 | 0.77 | 0.65 | 1.00 | 0.48 | 0.43 | 0.43 | 0.68 | 0.43 | 0.81 | 0.45 |
| **CellPLM** | 0.41 | 0.58 | 0.58 | 0.62 | 0.63 | 0.67 | 0.63 | 0.00 | 0.75 | 0.61 | 1.00 | 0.46 | 0.42 | 0.41 | 0.60 | 0.43 | 0.79 | 0.44 |
| **LangCell** | 0.40 | 0.55 | 0.55 | 0.70 | 0.71 | 0.67 | 0.60 | 0.00 | 0.77 | 0.65 | 1.00 | 0.46 | 0.33 | 0.40 | 0.63 | 0.42 | 0.81 | 0.40 |
| **scCello** | 0.41 | 0.38 | 0.39 | 0.41 | 0.43 | 1.00 | 0.93 | 0.00 | 0.73 | 0.56 | 1.00 | 0.44 | 0.41 | 0.41 | 0.40 | 0.64 | 0.76 | 0.42 |
| **scBERT** | 0.23 | 0.35 | 0.33 | 0.42 | 0.40 | 1.00 | 0.86 | 1.00 | 0.71 | 0.55 | 1.00 | 0.30 | 0.33 | 0.23 | 0.38 | 0.95 | 0.75 | 0.32 |
| **Geneformer** | 0.28 | 0.56 | 0.54 | 0.72 | 0.72 | 0.67 | 0.65 | 0.00 | 0.76 | 0.64 | 1.00 | 0.42 | 0.40 | 0.28 | 0.64 | 0.44 | 0.80 | 0.41 |
| **SCimilarity** | 0.34 | 0.56 | 0.58 | 0.68 | 0.68 | 0.67 | 0.66 | 0.00 | 0.72 | 0.58 | 1.00 | 0.37 | 0.42 | 0.34 | 0.62 | 0.44 | 0.77 | 0.39 |
| **CELLama** | 0.29 | 0.39 | 0.40 | 0.52 | 0.52 | 1.00 | 0.86 | 0.00 | 0.71 | 0.55 | 1.00 | 0.38 | 0.43 | 0.29 | 0.46 | 0.62 | 0.75 | 0.41 |
| **scGPT (integrated** | 0.31 | 0.58 | 0.59 | 0.61 | 0.61 | 0.50 | 0.55 | 0.00 | 0.72 | 0.61 | 1.00 | 0.32 | 0.35 | 0.31 | 0.60 | 0.35 | 0.77 | 0.33 |
| **scGPT** | 0.21 | 0.65 | 0.66 | 0.64 | 0.65 | 0.50 | 0.54 | 0.00 | 0.74 | 0.63 | 1.00 | 0.34 | 0.38 | 0.21 | 0.65 | 0.35 | 0.79 | 0.36 |
| **UCE** | 0.36 | 0.26 | 0.29 | 0.40 | 0.42 | 0.67 | 0.62 | 0.00 | 0.55 | 0.35 | 1.00 | 0.46 | 0.37 | 0.36 | 0.34 | 0.43 | 0.63 | 0.42 |
| **scFoundation** | 0.17 | 0.51 | 0.51 | 0.48 | 0.48 | 0.50 | 0.52 | 0.00 | 0.72 | 0.60 | 1.00 | 0.32 | 0.34 | 0.17 | 0.49 | 0.34 | 0.77 | 0.33 |
| **scVI (de novo)** | 0.31 | 0.17 | 0.19 | 0.55 | 0.57 | 0.67 | 0.62 | 0.00 | 0.32 | 0.17 | 1.00 | 0.34 | 0.29 | 0.31 | 0.37 | 0.43 | 0.50 | 0.32 |
| **scVI** | 0.07 | 0.27 | 0.28 | 0.37 | 0.38 | 0.80 | 0.74 | 0.00 | 0.49 | 0.36 | 1.00 | 0.30 | 0.28 | 0.07 | 0.33 | 0.51 | 0.62 | 0.29 |
| **GenePT-w** | 0.26 | 0.17 | 0.17 | 0.26 | 0.27 | 0.67 | 0.62 | 0.00 | 0.49 | 0.28 | 1.00 | 0.32 | 0.29 | 0.26 | 0.22 | 0.43 | 0.59 | 0.30 |
| **scPRINT** | 0.21 | 0.40 | 0.43 | 0.37 | 0.40 | 0.00 | 0.43 | 0.00 | 0.72 | 0.49 | 1.00 | 0.30 | 0.27 | 0.21 | 0.40 | 0.14 | 0.74 | 0.28 |

**Figure S128: Trajectory-inference metric scores for hepatoblast differentiation using PAGA.** Display and scoring conventions are as in Fig. S103.

| | Cell positions | Neighbourhood | | | | Topology | | | Features | | | Branch assignment | | Aggregated scores | | | | |
|---|---|---|---|---|---|---|---|---|---|---|---|---|---|---|---|---|---|---|
| method | $cor_{dist}$ | $NMSE_{lm}$ | $R^2_{lm}$ | $NMSE_{rf}$ | $R^2_{rf}$ | edgeflip | HIM | isomorphic | $cor_{features}$ | $wcor_{features}$ | $ks_{features}$ | $F1_{branches}$ | $F1_{milestones}$ | Pos. | Neighbor. | Topology | Feature | Branch |
| Geneformer | 0.42 | 0.49 | 0.46 | 0.76 | 0.77 | 1.00 | 0.85 | 1.00 | 0.73 | 0.58 | 1.00 | 0.39 | 0.33 | 0.42 | 0.62 | 0.95 | 0.77 | 0.36 |
| PCA | 0.43 | 0.69 | 0.71 | 0.86 | 0.87 | 0.67 | 0.68 | 0.00 | 0.71 | 0.52 | 1.00 | 0.31 | 0.38 | 0.43 | 0.78 | 0.45 | 0.75 | 0.34 |
| LangCell | 0.31 | 0.50 | 0.49 | 0.73 | 0.72 | 1.00 | 0.87 | 0.00 | 0.75 | 0.61 | 1.00 | 0.38 | 0.39 | 0.31 | 0.61 | 0.62 | 0.79 | 0.39 |
| CellPLM | 0.34 | 0.68 | 0.68 | 0.73 | 0.73 | 0.50 | 0.55 | 0.00 | 0.72 | 0.69 | 1.00 | 0.33 | 0.34 | 0.34 | 0.70 | 0.35 | 0.80 | 0.33 |
| scGPT (integrated | 0.31 | 0.54 | 0.53 | 0.75 | 0.75 | 0.67 | 0.65 | 0.00 | 0.63 | 0.51 | 1.00 | 0.36 | 0.31 | 0.31 | 0.64 | 0.44 | 0.71 | 0.34 |
| SCimilarity | 0.37 | 0.56 | 0.56 | 0.65 | 0.65 | 0.67 | 0.78 | 0.00 | 0.57 | 0.40 | 1.00 | 0.30 | 0.32 | 0.37 | 0.61 | 0.48 | 0.65 | 0.31 |
| CellFM | 0.40 | 0.18 | 0.19 | 0.77 | 0.77 | 0.67 | 0.62 | 0.00 | 0.47 | 0.38 | 1.00 | 0.39 | 0.37 | 0.40 | 0.48 | 0.43 | 0.62 | 0.38 |
| scFoundation | 0.25 | 0.44 | 0.45 | 0.65 | 0.65 | 0.67 | 0.74 | 0.00 | 0.59 | 0.41 | 1.00 | 0.36 | 0.32 | 0.25 | 0.55 | 0.47 | 0.67 | 0.34 |
| scVI (de novo) | 0.18 | 0.37 | 0.38 | 0.59 | 0.59 | 1.00 | 0.76 | 0.00 | 0.48 | 0.25 | 1.00 | 0.50 | 0.33 | 0.18 | 0.48 | 0.59 | 0.58 | 0.42 |
| scCello | 0.34 | 0.27 | 0.26 | 0.48 | 0.49 | 0.67 | 0.62 | 0.00 | 0.53 | 0.35 | 1.00 | 0.34 | 0.36 | 0.34 | 0.38 | 0.43 | 0.63 | 0.35 |
| CELLama | 0.17 | 0.31 | 0.32 | 0.64 | 0.64 | 0.67 | 0.70 | 0.00 | 0.63 | 0.36 | 1.00 | 0.31 | 0.32 | 0.17 | 0.48 | 0.46 | 0.66 | 0.31 |
| GenePT-w | 0.26 | 0.26 | 0.24 | 0.49 | 0.50 | 0.67 | 0.62 | 0.00 | 0.59 | 0.45 | 1.00 | 0.33 | 0.30 | 0.26 | 0.37 | 0.43 | 0.68 | 0.31 |
| UCE | 0.21 | 0.25 | 0.27 | 0.46 | 0.47 | 1.00 | 0.86 | 0.00 | 0.52 | 0.22 | 1.00 | 0.24 | 0.29 | 0.21 | 0.36 | 0.62 | 0.58 | 0.27 |
| scGPT | 0.19 | 0.45 | 0.45 | 0.68 | 0.68 | 0.50 | 0.60 | 0.00 | 0.55 | 0.41 | 1.00 | 0.25 | 0.25 | 0.19 | 0.56 | 0.37 | 0.65 | 0.25 |
| scBERT | 0.22 | 0.19 | 0.18 | 0.35 | 0.34 | 0.67 | 0.62 | 0.00 | 0.56 | 0.41 | 1.00 | 0.34 | 0.29 | 0.22 | 0.26 | 0.43 | 0.66 | 0.32 |
| scPRINT | 0.02 | 0.37 | 0.38 | 0.50 | 0.52 | 0.50 | 0.61 | 0.00 | 0.66 | 0.40 | 1.00 | 0.24 | 0.26 | 0.02 | 0.45 | 0.37 | 0.69 | 0.25 |
| scVI | 0.17 | 0.14 | 0.14 | 0.39 | 0.40 | 0.67 | 0.62 | 0.00 | 0.31 | 0.30 | 1.00 | 0.39 | 0.30 | 0.17 | 0.27 | 0.43 | 0.54 | 0.34 |

**Figure S129: Trajectory-inference metric scores for hepatoblast differentiation using Monocle 3.** Display and scoring conventions are as in Fig. S104.

| | Cell positions | Neighbourhood | | | | Topology | | | Features | | | Branch assignment | | Aggregated scores | | | | |
|---|---|---|---|---|---|---|---|---|---|---|---|---|---|---|---|---|---|---|
| method | $cor_{dist}$ | $NMSE_{lm}$ | $R^2_{lm}$ | $NMSE_{rf}$ | $R^2_{rf}$ | edgeflip | HIM | isomorphic | $cor_{features}$ | $wcor_{features}$ | $ks_{features}$ | $F1_{branches}$ | $F1_{milestones}$ | Pos. | Neighbor. | Topology | Feature | Branch |
| Geneformer | 0.30 | 0.27 | 0.26 | 0.52 | 0.50 | | | | 0.64 | 0.59 | 1.00 | | 0.46 | 0.30 | 0.39 | | 0.74 | 0.46 |
| PCA | 0.43 | 0.22 | 0.23 | 0.46 | 0.42 | | | | 0.62 | 0.50 | 1.00 | | 0.40 | 0.43 | 0.33 | | 0.71 | 0.40 |
| LangCell | 0.30 | 0.27 | 0.25 | 0.44 | 0.42 | | | | 0.65 | 0.58 | 1.00 | | 0.44 | 0.30 | 0.34 | | 0.74 | 0.44 |
| CellFM | 0.50 | 0.19 | 0.20 | 0.41 | 0.42 | | | | 0.50 | 0.32 | 1.00 | | 0.36 | 0.50 | 0.30 | | 0.61 | 0.36 |
| scGPT (integrated | 0.38 | 0.20 | 0.20 | 0.40 | 0.38 | | | | 0.63 | 0.50 | 1.00 | | 0.37 | 0.38 | 0.29 | | 0.71 | 0.37 |
| scFoundation | 0.28 | 0.23 | 0.23 | 0.37 | 0.36 | | | | 0.59 | 0.50 | 1.00 | | 0.45 | 0.28 | 0.30 | | 0.69 | 0.45 |
| SCimilarity | 0.37 | 0.20 | 0.20 | 0.39 | 0.38 | | | | 0.55 | 0.51 | 1.00 | | 0.34 | 0.37 | 0.29 | | 0.69 | 0.34 |
| scVI (de novo) | 0.37 | 0.19 | 0.20 | 0.26 | 0.27 | | | | 0.52 | 0.40 | 1.00 | | 0.35 | 0.37 | 0.23 | | 0.64 | 0.35 |
| CellPLM | 0.25 | 0.20 | 0.19 | 0.21 | 0.18 | | | | 0.54 | 0.53 | 1.00 | | 0.41 | 0.25 | 0.19 | | 0.69 | 0.41 |
| scGPT | 0.28 | 0.16 | 0.16 | 0.28 | 0.25 | | | | 0.48 | 0.43 | 1.00 | | 0.33 | 0.28 | 0.21 | | 0.64 | 0.33 |
| GenePT-w | 0.34 | 0.16 | 0.17 | 0.13 | 0.13 | | | | 0.50 | 0.34 | 1.00 | | 0.35 | 0.34 | 0.15 | | 0.61 | 0.35 |
| scCello | 0.30 | 0.12 | 0.12 | 0.13 | 0.13 | | | | 0.51 | 0.37 | 1.00 | | 0.38 | 0.30 | 0.13 | | 0.63 | 0.38 |
| scBERT | 0.24 | 0.20 | 0.19 | 0.14 | 0.11 | | | | 0.55 | 0.42 | 1.00 | | 0.34 | 0.24 | 0.16 | | 0.66 | 0.34 |
| UCE | 0.24 | 0.15 | 0.15 | 0.15 | 0.16 | | | | 0.51 | 0.34 | 1.00 | | 0.33 | 0.24 | 0.15 | | 0.61 | 0.33 |
| scPRINT | 0.21 | 0.10 | 0.10 | 0.03 | 0.02 | | | | 0.61 | 0.44 | 1.00 | | 0.33 | 0.21 | 0.06 | | 0.68 | 0.33 |
| scVI | 0.31 | 0.11 | 0.12 | 0.11 | 0.12 | | | | 0.33 | 0.29 | 1.00 | | 0.32 | 0.31 | 0.11 | | 0.54 | 0.32 |
| CELLama | 0.04 | 0.02 | 0.02 | 0.04 | 0.03 | | | | 0.51 | 0.40 | 1.00 | | 0.31 | 0.04 | 0.03 | | 0.64 | 0.31 |

**Figure S130: Trajectory-inference metric scores for hepatoblast differentiation using DPT.** Display and scoring conventions are as in Fig. S105.

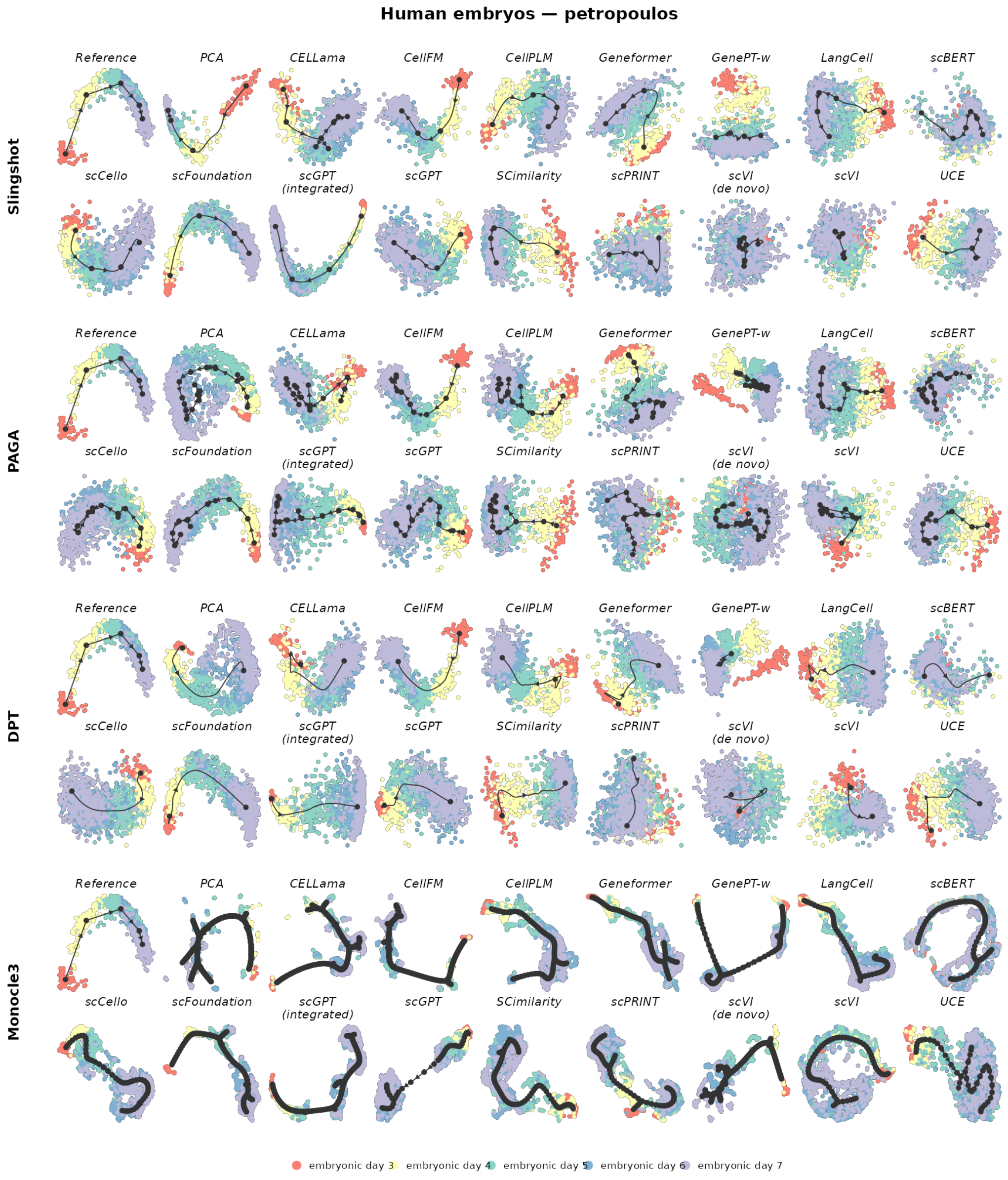

**Figure S131: Trajectory inference on human preimplantation embryogenesis.** The reference trajectory is linear. Rows show Slingshot, PAGA, DPT and Monocle 3; within each row, the reference is followed by the same 17 representations. Cells are coloured by the reference milestone or state. Black nodes, edges and arrows show the inferred milestone graph and its orientation. Layouts and trajectories were reconstructed independently for each representation and backend.

| | Cell positions | Neighbourhood | | | | Topology | | | Features | | | Branch assignment | | Aggregated scores | | | | |
|---|---|---|---|---|---|---|---|---|---|---|---|---|---|---|---|---|---|---|
| method | $cor_{dist}$ | $NMSE_{lm}$ | $R^2_{lm}$ | $NMSE_{rf}$ | $R^2_{rf}$ | edgeflip | HIM | isomorphic | $cor_{features}$ | $wcor_{features}$ | $ks_{features}$ | $F1_{branches}$ | $F1_{milestones}$ | Pos. | Neighbor. | Topology | Feature | Branch |
| scGPT (integrated | 0.74 | 0.81 | 0.81 | 0.78 | 0.78 | 1.00 | 1.00 | 1.00 | 0.91 | 0.88 | 1.00 | 1.00 | 0.77 | 0.74 | 0.80 | 1.00 | 0.93 | 0.89 |
| scFoundation | 0.72 | 0.80 | 0.80 | 0.77 | 0.77 | 1.00 | 1.00 | 1.00 | 0.88 | 0.82 | 1.00 | 1.00 | 0.71 | 0.72 | 0.79 | 1.00 | 0.90 | 0.85 |
| scCello | 0.70 | 0.74 | 0.74 | 0.71 | 0.71 | 1.00 | 1.00 | 1.00 | 0.88 | 0.82 | 1.00 | 1.00 | 0.78 | 0.70 | 0.73 | 1.00 | 0.90 | 0.89 |
| CellPLM | 0.68 | 0.75 | 0.75 | 0.74 | 0.74 | 1.00 | 1.00 | 1.00 | 0.85 | 0.73 | 1.00 | 1.00 | 0.69 | 0.68 | 0.75 | 1.00 | 0.86 | 0.85 |
| CellFM | 0.74 | 0.75 | 0.75 | 0.79 | 0.79 | 1.00 | 1.00 | 1.00 | 0.65 | 0.47 | 1.00 | 1.00 | 0.45 | 0.74 | 0.77 | 1.00 | 0.71 | 0.73 |
| scGPT | 0.57 | 0.64 | 0.64 | 0.60 | 0.60 | 1.00 | 1.00 | 1.00 | 0.83 | 0.76 | 1.00 | 1.00 | 0.72 | 0.57 | 0.62 | 1.00 | 0.86 | 0.86 |
| Geneformer | 0.54 | 0.56 | 0.56 | 0.57 | 0.57 | 1.00 | 1.00 | 1.00 | 0.82 | 0.71 | 1.00 | 1.00 | 0.71 | 0.54 | 0.56 | 1.00 | 0.84 | 0.85 |
| PCA | 0.72 | 0.64 | 0.64 | 0.83 | 0.83 | 1.00 | 1.00 | 1.00 | 0.46 | 0.23 | 1.00 | 1.00 | 0.43 | 0.72 | 0.73 | 1.00 | 0.56 | 0.72 |
| LangCell | 0.51 | 0.43 | 0.43 | 0.69 | 0.69 | 1.00 | 1.00 | 1.00 | 0.64 | 0.38 | 1.00 | 1.00 | 0.64 | 0.51 | 0.56 | 1.00 | 0.67 | 0.82 |
| SCimilarity | 0.52 | 0.50 | 0.50 | 0.57 | 0.57 | 1.00 | 1.00 | 1.00 | 0.62 | 0.39 | 1.00 | 1.00 | 0.63 | 0.52 | 0.53 | 1.00 | 0.67 | 0.81 |
| scVI | 0.34 | 0.13 | 0.13 | 0.36 | 0.36 | 1.00 | 1.00 | 1.00 | 0.54 | 0.51 | 1.00 | 1.00 | 0.66 | 0.34 | 0.24 | 1.00 | 0.68 | 0.83 |
| UCE | 0.50 | 0.67 | 0.67 | 0.66 | 0.66 | 0.67 | 0.55 | 0.00 | 0.70 | 0.52 | 1.00 | 0.36 | 0.38 | 0.50 | 0.66 | 0.41 | 0.74 | 0.37 |
| CELLama | 0.59 | 0.75 | 0.75 | 0.77 | 0.77 | 0.33 | 0.39 | 0.00 | 0.68 | 0.63 | 1.00 | 0.22 | 0.28 | 0.59 | 0.76 | 0.24 | 0.77 | 0.25 |
| scVI (de novo) | 0.49 | 0.61 | 0.61 | 0.84 | 0.84 | 0.50 | 0.53 | 0.00 | 0.39 | 0.35 | 1.00 | 0.33 | 0.27 | 0.49 | 0.72 | 0.34 | 0.58 | 0.30 |
| scPRINT | 0.04 | 0.01 | 0.01 | 0.00 | 0.00 | 1.00 | 1.00 | 1.00 | 0.17 | 0.00 | 1.00 | 1.00 | 0.38 | 0.04 | 0.00 | 1.00 | 0.39 | 0.69 |
| scBERT | 0.02 | 0.00 | 0.00 | 0.00 | 0.00 | 1.00 | 1.00 | 1.00 | 0.00 | 0.00 | 1.00 | 1.00 | 0.40 | 0.02 | 0.00 | 1.00 | 0.33 | 0.70 |
| GenePT-w | 0.02 | 0.00 | 0.00 | 0.00 | 0.00 | 1.00 | 1.00 | 0.00 | 0.00 | 0.00 | 1.00 | 0.54 | 0.29 | 0.02 | 0.00 | 0.67 | 0.33 | 0.42 |

**Figure S132: Trajectory-inference metric scores for human preimplantation embryogenesis using Slingshot.** Display and scoring conventions are as in Fig. S102.

| | Cell positions | Neighbourhood | | | | Topology | | | Features | | | Branch assignment | | Aggregated scores | | | | |
|---|---|---|---|---|---|---|---|---|---|---|---|---|---|---|---|---|---|---|
| method | $cor_{dist}$ | $NMSE_{lm}$ | $R^2_{lm}$ | $NMSE_{rf}$ | $R^2_{rf}$ | edgeflip | HIM | isomorphic | $cor_{features}$ | $wcor_{features}$ | $ks_{features}$ | $F1_{branches}$ | $F1_{milestones}$ | Pos. | Neighbor. | Topology | Feature | Branch |
| CellFM | 0.71 | 0.79 | 0.79 | 0.91 | 0.91 | 1.00 | 1.00 | 1.00 | 0.86 | 0.82 | 1.00 | 1.00 | 0.80 | 0.71 | 0.85 | 1.00 | 0.89 | 0.90 |
| scGPT | 0.57 | 0.69 | 0.69 | 0.75 | 0.75 | 0.67 | 0.62 | 0.00 | 0.84 | 0.82 | 1.00 | 0.43 | 0.48 | 0.57 | 0.72 | 0.43 | 0.89 | 0.46 |
| scFoundation | 0.65 | 0.84 | 0.84 | 0.85 | 0.85 | 0.40 | 0.42 | 0.00 | 0.77 | 0.80 | 1.00 | 0.26 | 0.36 | 0.65 | 0.84 | 0.27 | 0.85 | 0.31 |
| scCello | 0.59 | 0.78 | 0.78 | 0.80 | 0.80 | 0.40 | 0.40 | 0.00 | 0.83 | 0.83 | 1.00 | 0.24 | 0.37 | 0.59 | 0.79 | 0.27 | 0.89 | 0.31 |
| scVI (de novo) | 0.71 | 0.91 | 0.91 | 0.91 | 0.91 | 0.00 | 0.36 | 0.00 | 0.77 | 0.81 | 1.00 | 0.17 | 0.24 | 0.71 | 0.91 | 0.12 | 0.86 | 0.21 |
| Geneformer | 0.51 | 0.79 | 0.79 | 0.78 | 0.78 | 0.40 | 0.40 | 0.00 | 0.73 | 0.69 | 1.00 | 0.25 | 0.31 | 0.51 | 0.79 | 0.27 | 0.81 | 0.28 |
| scGPT (integrated | 0.67 | 0.84 | 0.84 | 0.79 | 0.79 | 0.29 | 0.38 | 0.00 | 0.57 | 0.57 | 1.00 | 0.20 | 0.24 | 0.67 | 0.82 | 0.22 | 0.71 | 0.22 |
| LangCell | 0.49 | 0.80 | 0.80 | 0.80 | 0.80 | 0.40 | 0.40 | 0.00 | 0.66 | 0.68 | 1.00 | 0.24 | 0.33 | 0.49 | 0.80 | 0.27 | 0.78 | 0.28 |
| CellPLM | 0.58 | 0.83 | 0.83 | 0.84 | 0.84 | 0.00 | 0.32 | 0.00 | 0.74 | 0.75 | 1.00 | 0.17 | 0.26 | 0.58 | 0.84 | 0.11 | 0.83 | 0.22 |
| PCA | 0.59 | 0.91 | 0.91 | 0.93 | 0.93 | 0.00 | 0.31 | 0.00 | 0.64 | 0.63 | 1.00 | 0.14 | 0.20 | 0.59 | 0.92 | 0.10 | 0.75 | 0.17 |
| UCE | 0.42 | 0.56 | 0.56 | 0.65 | 0.65 | 0.67 | 0.45 | 0.00 | 0.66 | 0.56 | 1.00 | 0.37 | 0.34 | 0.42 | 0.60 | 0.37 | 0.74 | 0.35 |
| SCimilarity | 0.46 | 0.78 | 0.78 | 0.77 | 0.77 | 0.29 | 0.36 | 0.00 | 0.59 | 0.43 | 1.00 | 0.18 | 0.25 | 0.46 | 0.78 | 0.21 | 0.67 | 0.21 |
| CELLama | 0.42 | 0.77 | 0.77 | 0.79 | 0.79 | 0.00 | 0.35 | 0.00 | 0.65 | 0.63 | 1.00 | 0.16 | 0.21 | 0.42 | 0.78 | 0.12 | 0.76 | 0.18 |
| scPRINT | 0.16 | 0.46 | 0.46 | 0.53 | 0.53 | 0.67 | 0.57 | 0.00 | 0.55 | 0.30 | 1.00 | 0.43 | 0.27 | 0.16 | 0.49 | 0.41 | 0.62 | 0.35 |
| scVI | 0.08 | 0.50 | 0.50 | 0.60 | 0.60 | 0.29 | 0.35 | 0.00 | 0.54 | 0.49 | 1.00 | 0.20 | 0.21 | 0.08 | 0.55 | 0.21 | 0.68 | 0.21 |
| GenePT-w | 0.00 | 0.34 | 0.34 | 0.43 | 0.43 | 0.00 | 0.32 | 0.00 | 0.28 | 0.27 | 1.00 | 0.14 | 0.15 | 0.00 | 0.39 | 0.11 | 0.52 | 0.15 |
| scBERT | 0.02 | 0.11 | 0.11 | 0.17 | 0.17 | 0.40 | 0.44 | 0.00 | 0.27 | 0.15 | 1.00 | 0.26 | 0.20 | 0.02 | 0.14 | 0.28 | 0.47 | 0.23 |

**Figure S133: Trajectory-inference metric scores for human preimplantation embryogenesis using PAGA.** Display and scoring conventions are as in Fig. S103.

| | Cell positions | Neighbourhood | | | | Topology | | | Features | | | Branch assignment | | Aggregated scores | | | | |
|---|---|---|---|---|---|---|---|---|---|---|---|---|---|---|---|---|---|---|
| **method** | $cor_{dist}$ | $NMSE_{lm}$ | $R^2_{lm}$ | $NMSE_{rf}$ | $R^2_{rf}$ | edgeflip | HIM | isomorphic | $cor_{features}$ | $wcor_{features}$ | $ks_{features}$ | $F1_{branches}$ | $F1_{milestones}$ | Pos. | Neighbor. | Topology | Feature | Branch |
| **LangCell** | 0.57 | 0.82 | 0.82 | 0.87 | 0.87 | 0.67 | 0.70 | 0.00 | 0.80 | 0.79 | 1.00 | 0.33 | 0.41 | 0.57 | 0.85 | 0.46 | 0.86 | 0.37 |
| **CellPLM** | 0.56 | 0.80 | 0.80 | 0.84 | 0.84 | 0.67 | 0.63 | 0.00 | 0.74 | 0.76 | 1.00 | 0.34 | 0.44 | 0.56 | 0.82 | 0.43 | 0.83 | 0.39 |
| **scCello** | 0.53 | 0.75 | 0.75 | 0.82 | 0.82 | 0.67 | 0.54 | 0.00 | 0.78 | 0.77 | 1.00 | 0.38 | 0.46 | 0.53 | 0.78 | 0.40 | 0.85 | 0.42 |
| **scGPT** | 0.52 | 0.67 | 0.67 | 0.73 | 0.73 | 0.67 | 0.51 | 0.00 | 0.76 | 0.75 | 1.00 | 0.34 | 0.38 | 0.52 | 0.70 | 0.39 | 0.84 | 0.36 |
| **SCimilarity** | 0.53 | 0.71 | 0.71 | 0.76 | 0.76 | 0.40 | 0.54 | 0.00 | 0.73 | 0.71 | 1.00 | 0.27 | 0.33 | 0.53 | 0.73 | 0.31 | 0.81 | 0.30 |
| **scGPT (integrated** | 0.66 | 0.84 | 0.84 | 0.85 | 0.85 | 0.00 | 0.40 | 0.00 | 0.75 | 0.82 | 1.00 | 0.15 | 0.22 | 0.66 | 0.84 | 0.13 | 0.86 | 0.18 |
| **scFoundation** | 0.69 | 0.85 | 0.85 | 0.87 | 0.87 | 0.00 | 0.42 | 0.00 | 0.71 | 0.71 | 1.00 | 0.15 | 0.22 | 0.69 | 0.86 | 0.14 | 0.81 | 0.18 |
| **scVI** | 0.48 | 0.58 | 0.58 | 0.63 | 0.63 | 0.67 | 0.42 | 0.00 | 0.71 | 0.72 | 1.00 | 0.36 | 0.38 | 0.48 | 0.61 | 0.36 | 0.81 | 0.37 |
| **Geneformer** | 0.60 | 0.78 | 0.78 | 0.81 | 0.81 | 0.29 | 0.43 | 0.00 | 0.66 | 0.69 | 1.00 | 0.18 | 0.22 | 0.60 | 0.79 | 0.24 | 0.78 | 0.20 |
| **CellFM** | 0.70 | 0.87 | 0.87 | 0.92 | 0.92 | 0.00 | 0.34 | 0.00 | 0.65 | 0.63 | 1.00 | 0.09 | 0.15 | 0.70 | 0.89 | 0.11 | 0.76 | 0.12 |
| **UCE** | 0.36 | 0.66 | 0.66 | 0.68 | 0.68 | 0.40 | 0.53 | 0.00 | 0.82 | 0.78 | 1.00 | 0.28 | 0.34 | 0.36 | 0.67 | 0.31 | 0.87 | 0.31 |
| **scVI (de novo)** | 0.69 | 0.93 | 0.93 | 0.94 | 0.94 | 0.00 | 0.34 | 0.00 | 0.49 | 0.51 | 1.00 | 0.06 | 0.09 | 0.69 | 0.93 | 0.11 | 0.67 | 0.08 |
| **PCA** | 0.63 | 0.94 | 0.94 | 0.94 | 0.94 | 0.00 | 0.35 | 0.00 | 0.48 | 0.43 | 1.00 | 0.05 | 0.10 | 0.63 | 0.94 | 0.12 | 0.63 | 0.08 |
| **GenePT-w** | 0.27 | 0.76 | 0.76 | 0.82 | 0.82 | 0.40 | 0.44 | 0.00 | 0.62 | 0.67 | 1.00 | 0.22 | 0.29 | 0.27 | 0.79 | 0.28 | 0.76 | 0.26 |
| **CELLama** | 0.45 | 0.82 | 0.82 | 0.85 | 0.85 | 0.00 | 0.34 | 0.00 | 0.50 | 0.49 | 1.00 | 0.07 | 0.11 | 0.45 | 0.83 | 0.11 | 0.67 | 0.09 |
| **scPRINT** | 0.06 | 0.49 | 0.49 | 0.61 | 0.61 | 0.40 | 0.42 | 0.00 | 0.58 | 0.63 | 1.00 | 0.22 | 0.22 | 0.06 | 0.55 | 0.27 | 0.74 | 0.22 |
| **scBERT** | 0.03 | 0.09 | 0.09 | 0.20 | 0.20 | 0.67 | 0.51 | 0.00 | 0.22 | 0.12 | 1.00 | 0.29 | 0.23 | 0.03 | 0.14 | 0.39 | 0.45 | 0.26 |

**Figure S134: Trajectory-inference metric scores for human preimplantation embryogenesis using Monocle 3.** Display and scoring conventions are as in Fig. S104.

| | Cell positions | Neighbourhood | | | | Topology | | | Features | | | Branch assignment | | Aggregated scores | | | | |
|---|---|---|---|---|---|---|---|---|---|---|---|---|---|---|---|---|---|---|
| **method** | $cor_{dist}$ | $NMSE_{lm}$ | $R^2_{lm}$ | $NMSE_{rf}$ | $R^2_{rf}$ | edgeflip | HIM | isomorphic | $cor_{features}$ | $wcor_{features}$ | $ks_{features}$ | $F1_{branches}$ | $F1_{milestones}$ | Pos. | Neighbor. | Topology | Feature | Branch |
| **PCA** | 0.77 | 0.79 | 0.79 | 0.88 | 0.88 | | | | 0.81 | 0.69 | 1.00 | | 0.53 | 0.77 | 0.84 | | 0.83 | 0.53 |
| **scCello** | 0.71 | 0.76 | 0.76 | 0.74 | 0.74 | | | | 0.79 | 0.67 | 1.00 | | 0.51 | 0.71 | 0.75 | | 0.82 | 0.51 |
| **CellPLM** | 0.67 | 0.71 | 0.71 | 0.76 | 0.76 | | | | 0.75 | 0.58 | 1.00 | | 0.56 | 0.67 | 0.73 | | 0.78 | 0.56 |
| **Geneformer** | 0.65 | 0.73 | 0.73 | 0.74 | 0.74 | | | | 0.74 | 0.57 | 1.00 | | 0.52 | 0.65 | 0.73 | | 0.77 | 0.52 |
| **scFoundation** | 0.68 | 0.74 | 0.74 | 0.78 | 0.79 | | | | 0.68 | 0.45 | 1.00 | | 0.50 | 0.68 | 0.76 | | 0.71 | 0.50 |
| **scVI (de novo)** | 0.70 | 0.56 | 0.56 | 0.65 | 0.65 | | | | 0.83 | 0.86 | 1.00 | | 0.43 | 0.70 | 0.60 | | 0.90 | 0.43 |
| **scGPT** | 0.61 | 0.71 | 0.71 | 0.65 | 0.65 | | | | 0.76 | 0.59 | 1.00 | | 0.55 | 0.61 | 0.68 | | 0.78 | 0.55 |
| **scGPT (integrated** | 0.71 | 0.72 | 0.72 | 0.80 | 0.80 | | | | 0.63 | 0.43 | 1.00 | | 0.45 | 0.71 | 0.76 | | 0.69 | 0.45 |
| **SCimilarity** | 0.60 | 0.65 | 0.65 | 0.64 | 0.64 | | | | 0.65 | 0.39 | 1.00 | | 0.54 | 0.60 | 0.65 | | 0.68 | 0.54 |
| **UCE** | 0.57 | 0.64 | 0.64 | 0.59 | 0.59 | | | | 0.72 | 0.51 | 1.00 | | 0.50 | 0.57 | 0.61 | | 0.74 | 0.50 |
| **LangCell** | 0.66 | 0.63 | 0.63 | 0.78 | 0.78 | | | | 0.48 | 0.22 | 1.00 | | 0.44 | 0.66 | 0.70 | | 0.57 | 0.44 |
| **CELLama** | 0.59 | 0.59 | 0.59 | 0.63 | 0.63 | | | | 0.69 | 0.46 | 1.00 | | 0.40 | 0.59 | 0.61 | | 0.72 | 0.40 |
| **CellFM** | 0.60 | 0.60 | 0.60 | 0.65 | 0.65 | | | | 0.42 | 0.21 | 1.00 | | 0.43 | 0.60 | 0.62 | | 0.54 | 0.43 |
| **scVI** | 0.21 | 0.17 | 0.17 | 0.00 | 0.00 | | | | 0.41 | 0.10 | 1.00 | | 0.37 | 0.21 | 0.09 | | 0.50 | 0.37 |
| **scPRINT** | 0.01 | 0.00 | 0.00 | 0.00 | 0.00 | | | | 0.11 | 0.02 | 1.00 | | 0.39 | 0.01 | 0.00 | | 0.38 | 0.39 |
| **GenePT-w** | 0.00 | 0.00 | 0.00 | 0.00 | 0.00 | | | | 0.10 | 0.00 | 1.00 | | 0.39 | 0.00 | 0.00 | | 0.37 | 0.39 |
| **scBERT** | 0.02 | 0.00 | 0.00 | 0.00 | 0.00 | | | | 0.02 | 0.00 | 1.00 | | 0.39 | 0.02 | 0.00 | | 0.34 | 0.39 |

**Figure S135: Trajectory-inference metric scores for human preimplantation embryogenesis using DPT.** Display and scoring conventions are as in Fig. S105.

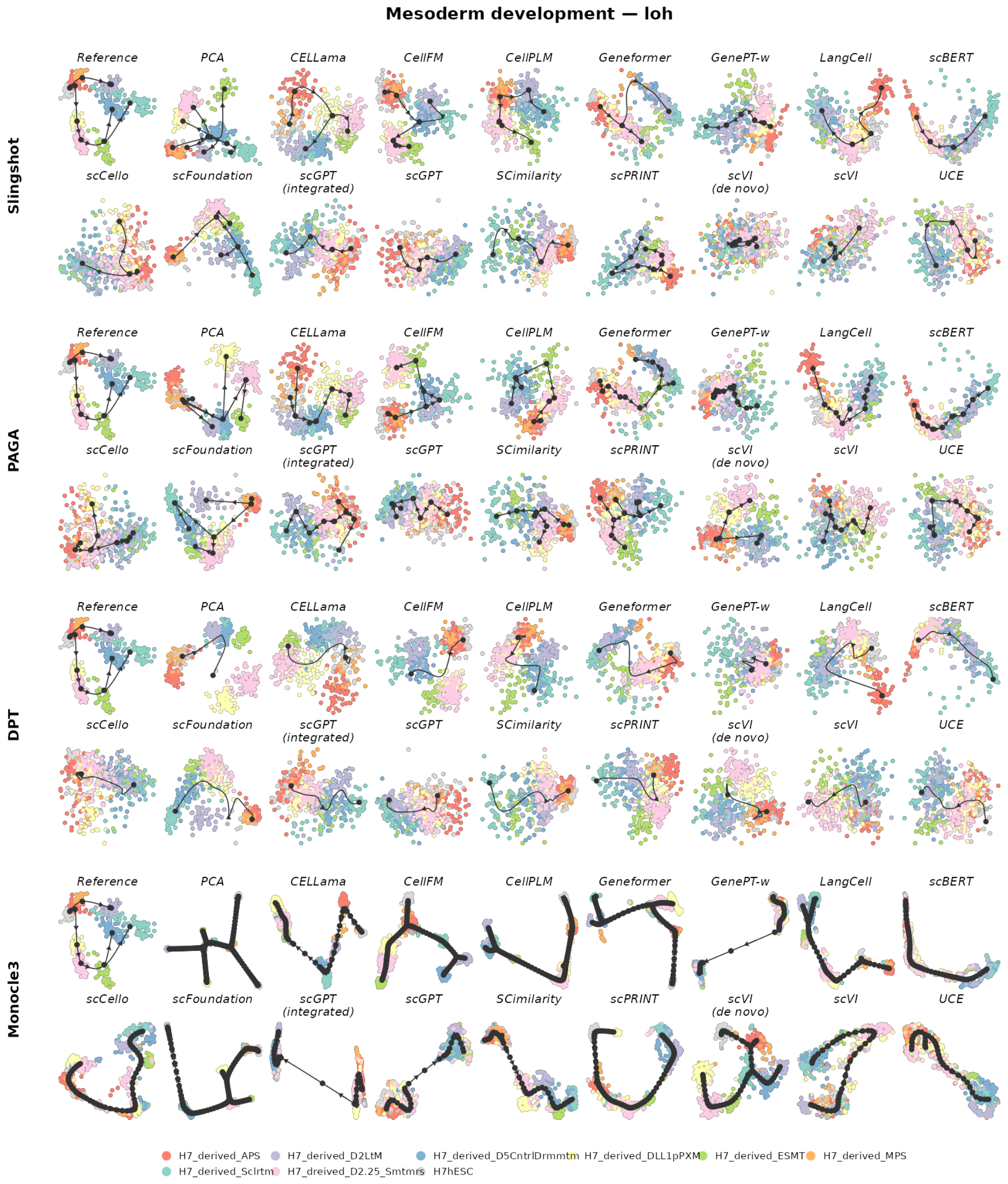

**Figure S136: Trajectory inference on mesoderm development.** The reference trajectory is tree-structured. Rows show Slingshot, PAGA, DPT and Monocle 3; within each row, the reference is followed by the same 17 representations. Cells are coloured by the reference milestone or state. Black nodes, edges and arrows show the inferred milestone graph and its orientation. Layouts and trajectories were reconstructed independently for each representation and backend.

| | Cell positions | Neighbourhood | | | | Topology | | | Features | | | Branch assignment | | Aggregated scores | | | | |
|---|---|---|---|---|---|---|---|---|---|---|---|---|---|---|---|---|---|---|
| method | $cor_{dist}$ | $NMSE_{lm}$ | $R^2_{lm}$ | $NMSE_{rf}$ | $R^2_{rf}$ | edgeflip | HIM | isomorphic | $cor_{features}$ | $wcor_{features}$ | $ks_{features}$ | $F1_{branches}$ | $F1_{milestones}$ | Pos. | Neighbor. | Topology | Feature | Branch |
| scFoundation | 0.65 | 0.80 | 0.80 | 0.82 | 0.83 | 0.67 | 0.55 | 0.00 | 0.86 | 0.85 | 1.00 | 0.48 | 0.67 | 0.65 | 0.81 | 0.41 | 0.90 | 0.57 |
| CellPLM | 0.43 | 0.53 | 0.47 | 0.65 | 0.65 | 1.00 | 0.77 | 1.00 | 0.80 | 0.78 | 1.00 | 0.33 | 0.41 | 0.43 | 0.58 | 0.92 | 0.86 | 0.37 |
| CellFM | 0.38 | 0.51 | 0.53 | 0.74 | 0.75 | 1.00 | 0.50 | 0.00 | 0.79 | 0.79 | 1.00 | 0.40 | 0.58 | 0.38 | 0.63 | 0.50 | 0.86 | 0.49 |
| scVI (de novo) | 0.38 | 0.53 | 0.51 | 0.71 | 0.69 | 1.00 | 0.67 | 0.00 | 0.76 | 0.73 | 1.00 | 0.43 | 0.48 | 0.38 | 0.61 | 0.56 | 0.83 | 0.46 |
| CELLama | 0.43 | 0.52 | 0.47 | 0.56 | 0.50 | 1.00 | 0.73 | 0.00 | 0.77 | 0.73 | 1.00 | 0.36 | 0.50 | 0.43 | 0.51 | 0.58 | 0.83 | 0.43 |
| PCA | 0.27 | 0.82 | 0.82 | 0.94 | 0.94 | 0.44 | 0.51 | 0.00 | 0.73 | 0.75 | 1.00 | 0.42 | 0.54 | 0.27 | 0.88 | 0.32 | 0.83 | 0.48 |
| Geneformer | 0.43 | 0.34 | 0.32 | 0.43 | 0.39 | 1.00 | 0.74 | 0.00 | 0.64 | 0.58 | 1.00 | 0.36 | 0.35 | 0.43 | 0.37 | 0.58 | 0.74 | 0.36 |
| scPRINT | 0.38 | 0.35 | 0.32 | 0.53 | 0.50 | 1.00 | 0.60 | 0.00 | 0.64 | 0.64 | 1.00 | 0.30 | 0.41 | 0.38 | 0.43 | 0.53 | 0.76 | 0.36 |
| SCimilarity | 0.53 | 0.27 | 0.24 | 0.31 | 0.25 | 0.67 | 0.45 | 0.00 | 0.75 | 0.77 | 1.00 | 0.36 | 0.38 | 0.53 | 0.27 | 0.37 | 0.84 | 0.37 |
| scVI | 0.26 | 0.32 | 0.28 | 0.26 | 0.24 | 1.00 | 0.49 | 0.00 | 0.75 | 0.75 | 1.00 | 0.38 | 0.40 | 0.26 | 0.27 | 0.50 | 0.83 | 0.39 |
| scGPT (integrated | 0.47 | 0.22 | 0.21 | 0.23 | 0.20 | 0.67 | 0.45 | 0.00 | 0.52 | 0.46 | 1.00 | 0.36 | 0.40 | 0.47 | 0.22 | 0.37 | 0.66 | 0.38 |
| LangCell | 0.34 | 0.25 | 0.23 | 0.43 | 0.40 | 0.67 | 0.45 | 0.00 | 0.62 | 0.61 | 1.00 | 0.28 | 0.32 | 0.34 | 0.33 | 0.37 | 0.74 | 0.30 |
| scBERT | 0.43 | 0.23 | 0.21 | 0.22 | 0.20 | 0.67 | 0.45 | 0.00 | 0.46 | 0.40 | 1.00 | 0.36 | 0.39 | 0.43 | 0.21 | 0.37 | 0.62 | 0.38 |
| scCello | 0.25 | 0.24 | 0.22 | 0.24 | 0.21 | 1.00 | 0.54 | 0.00 | 0.61 | 0.58 | 1.00 | 0.29 | 0.27 | 0.25 | 0.23 | 0.51 | 0.73 | 0.28 |
| UCE | 0.39 | 0.20 | 0.19 | 0.17 | 0.14 | 0.67 | 0.45 | 0.00 | 0.52 | 0.48 | 1.00 | 0.36 | 0.38 | 0.39 | 0.18 | 0.37 | 0.67 | 0.37 |
| GenePT-w | 0.40 | 0.22 | 0.21 | 0.12 | 0.11 | 0.67 | 0.45 | 0.00 | 0.41 | 0.38 | 1.00 | 0.36 | 0.39 | 0.40 | 0.16 | 0.37 | 0.60 | 0.37 |
| scGPT | 0.26 | 0.15 | 0.14 | 0.02 | 0.00 | 0.67 | 0.45 | 0.00 | 0.43 | 0.36 | 1.00 | 0.36 | 0.31 | 0.26 | 0.08 | 0.37 | 0.60 | 0.34 |

**Figure S137: Trajectory-inference metric scores for mesoderm development using Slingshot.** Display and scoring conventions are as in Fig. S102.

| | Cell positions | Neighbourhood | | | | Topology | | | Features | | | Branch assignment | | Aggregated scores | | | | |
|---|---|---|---|---|---|---|---|---|---|---|---|---|---|---|---|---|---|---|
| method | $cor_{dist}$ | $NMSE_{lm}$ | $R^2_{lm}$ | $NMSE_{rf}$ | $R^2_{rf}$ | edgeflip | HIM | isomorphic | $cor_{features}$ | $wcor_{features}$ | $ks_{features}$ | $F1_{branches}$ | $F1_{milestones}$ | Pos. | Neighbor. | Topology | Feature | Branch |
| CellPLM | 0.68 | 0.65 | 0.68 | 0.89 | 0.89 | 1.00 | 0.83 | 0.00 | 0.85 | 0.83 | 1.00 | 0.69 | 0.79 | 0.68 | 0.78 | 0.61 | 0.90 | 0.74 |
| Geneformer | 0.53 | 0.56 | 0.51 | 0.83 | 0.84 | 1.00 | 0.73 | 0.00 | 0.84 | 0.83 | 1.00 | 0.66 | 0.68 | 0.53 | 0.68 | 0.58 | 0.89 | 0.67 |
| LangCell | 0.56 | 0.50 | 0.44 | 0.82 | 0.82 | 1.00 | 0.65 | 1.00 | 0.74 | 0.78 | 1.00 | 0.31 | 0.47 | 0.56 | 0.65 | 0.88 | 0.84 | 0.39 |
| scFoundation | 0.71 | 0.72 | 0.74 | 0.88 | 0.88 | 0.67 | 0.55 | 0.00 | 0.79 | 0.78 | 1.00 | 0.47 | 0.55 | 0.71 | 0.81 | 0.41 | 0.86 | 0.51 |
| SCimilarity | 0.52 | 0.47 | 0.42 | 0.63 | 0.62 | 1.00 | 0.57 | 1.00 | 0.83 | 0.85 | 1.00 | 0.36 | 0.43 | 0.52 | 0.54 | 0.86 | 0.89 | 0.39 |
| CELLama | 0.43 | 0.76 | 0.77 | 0.84 | 0.84 | 0.67 | 0.59 | 0.00 | 0.82 | 0.86 | 1.00 | 0.63 | 0.63 | 0.43 | 0.81 | 0.42 | 0.90 | 0.63 |
| scGPT (integrated | 0.49 | 0.32 | 0.30 | 0.54 | 0.56 | 1.00 | 0.88 | 1.00 | 0.67 | 0.66 | 1.00 | 0.25 | 0.28 | 0.49 | 0.43 | 0.96 | 0.78 | 0.26 |
| CellFM | 0.35 | 0.53 | 0.52 | 0.80 | 0.80 | 1.00 | 0.48 | 0.00 | 0.82 | 0.79 | 1.00 | 0.45 | 0.54 | 0.35 | 0.66 | 0.49 | 0.87 | 0.49 |
| scPRINT | 0.41 | 0.64 | 0.59 | 0.67 | 0.65 | 0.50 | 0.48 | 0.00 | 0.81 | 0.85 | 1.00 | 0.42 | 0.37 | 0.41 | 0.64 | 0.33 | 0.89 | 0.40 |
| scVI (de novo) | 0.38 | 0.50 | 0.45 | 0.69 | 0.64 | 0.67 | 0.45 | 0.00 | 0.75 | 0.68 | 1.00 | 0.40 | 0.64 | 0.38 | 0.57 | 0.37 | 0.81 | 0.52 |
| PCA | 0.28 | 0.63 | 0.66 | 0.66 | 0.69 | 0.57 | 0.52 | 0.00 | 0.73 | 0.75 | 1.00 | 0.42 | 0.43 | 0.28 | 0.66 | 0.36 | 0.83 | 0.43 |
| scGPT | 0.37 | 0.34 | 0.36 | 0.41 | 0.43 | 1.00 | 0.76 | 0.00 | 0.67 | 0.66 | 1.00 | 0.36 | 0.48 | 0.37 | 0.38 | 0.59 | 0.78 | 0.42 |
| scCello | 0.32 | 0.50 | 0.45 | 0.55 | 0.53 | 0.67 | 0.53 | 0.00 | 0.76 | 0.76 | 1.00 | 0.41 | 0.40 | 0.32 | 0.51 | 0.40 | 0.84 | 0.40 |
| scVI | 0.18 | 0.36 | 0.36 | 0.38 | 0.37 | 0.80 | 0.76 | 0.00 | 0.77 | 0.76 | 1.00 | 0.43 | 0.42 | 0.18 | 0.37 | 0.52 | 0.84 | 0.42 |
| UCE | 0.43 | 0.20 | 0.20 | 0.42 | 0.43 | 0.67 | 0.45 | 0.00 | 0.55 | 0.48 | 1.00 | 0.28 | 0.33 | 0.43 | 0.31 | 0.37 | 0.68 | 0.30 |
| scBERT | 0.38 | 0.24 | 0.22 | 0.36 | 0.34 | 0.67 | 0.45 | 0.00 | 0.53 | 0.50 | 1.00 | 0.28 | 0.34 | 0.38 | 0.29 | 0.37 | 0.68 | 0.31 |
| GenePT-w | 0.36 | 0.22 | 0.21 | 0.31 | 0.30 | 0.67 | 0.45 | 0.00 | 0.47 | 0.44 | 1.00 | 0.28 | 0.32 | 0.36 | 0.26 | 0.37 | 0.63 | 0.30 |

**Figure S138: Trajectory-inference metric scores for mesoderm development using PAGA.** Display and scoring conventions are as in Fig. S103.

| | Cell positions | Neighbourhood | | | | Topology | | | Features | | | Branch assignment | | Aggregated scores | | | | |
|---|---|---|---|---|---|---|---|---|---|---|---|---|---|---|---|---|---|---|
| method | $cor_{dist}$ | $NMSE_{lm}$ | $R^2_{lm}$ | $NMSE_{rf}$ | $R^2_{rf}$ | edgeflip | HIM | isomorphic | $cor_{features}$ | $wcor_{features}$ | $ks_{features}$ | $F1_{branches}$ | $F1_{milestones}$ | Pos. | Neighbor. | Topology | Feature | Branch |
| CellFM | 0.47 | 0.90 | 0.89 | 0.94 | 0.94 | 0.50 | 0.64 | 0.00 | 0.85 | 0.85 | 1.00 | 0.52 | 0.62 | 0.47 | 0.92 | 0.38 | 0.90 | 0.57 |
| LangCell | 0.58 | 0.62 | 0.65 | 0.80 | 0.82 | 0.67 | 0.67 | 0.00 | 0.72 | 0.77 | 1.00 | 0.42 | 0.54 | 0.58 | 0.72 | 0.45 | 0.83 | 0.48 |
| CellPLM | 0.68 | 0.64 | 0.64 | 0.86 | 0.85 | 0.50 | 0.54 | 0.00 | 0.80 | 0.78 | 1.00 | 0.41 | 0.37 | 0.68 | 0.75 | 0.35 | 0.86 | 0.39 |
| Geneformer | 0.46 | 0.67 | 0.69 | 0.85 | 0.85 | 0.67 | 0.58 | 0.00 | 0.79 | 0.80 | 1.00 | 0.44 | 0.52 | 0.46 | 0.77 | 0.41 | 0.86 | 0.48 |
| scFoundation | 0.69 | 0.53 | 0.53 | 0.89 | 0.90 | 0.67 | 0.60 | 0.00 | 0.76 | 0.73 | 1.00 | 0.24 | 0.36 | 0.69 | 0.71 | 0.42 | 0.83 | 0.30 |
| SCimilarity | 0.47 | 0.30 | 0.27 | 0.68 | 0.66 | 1.00 | 0.67 | 1.00 | 0.74 | 0.76 | 1.00 | 0.21 | 0.25 | 0.47 | 0.48 | 0.89 | 0.83 | 0.23 |
| scPRINT | 0.48 | 0.32 | 0.29 | 0.62 | 0.62 | 1.00 | 0.66 | 1.00 | 0.67 | 0.61 | 1.00 | 0.23 | 0.25 | 0.48 | 0.46 | 0.89 | 0.76 | 0.24 |
| scGPT (integrated | 0.51 | 0.22 | 0.21 | 0.57 | 0.59 | 1.00 | 0.48 | 1.00 | 0.57 | 0.52 | 1.00 | 0.20 | 0.25 | 0.51 | 0.40 | 0.83 | 0.70 | 0.23 |
| scVI (de novo) | 0.35 | 0.60 | 0.59 | 0.85 | 0.84 | 0.50 | 0.56 | 0.00 | 0.76 | 0.76 | 1.00 | 0.31 | 0.34 | 0.35 | 0.72 | 0.35 | 0.84 | 0.32 |
| UCE | 0.37 | 0.28 | 0.26 | 0.42 | 0.39 | 1.00 | 0.72 | 1.00 | 0.61 | 0.56 | 1.00 | 0.22 | 0.25 | 0.37 | 0.34 | 0.91 | 0.72 | 0.23 |
| PCA | 0.21 | 0.93 | 0.93 | 0.95 | 0.96 | 0.00 | 0.48 | 0.00 | 0.74 | 0.77 | 1.00 | 0.32 | 0.50 | 0.21 | 0.94 | 0.16 | 0.84 | 0.41 |
| scVI | 0.59 | 0.22 | 0.26 | 0.46 | 0.46 | 0.67 | 0.45 | 0.00 | 0.73 | 0.72 | 1.00 | 0.44 | 0.43 | 0.59 | 0.35 | 0.37 | 0.82 | 0.44 |
| scCello | 0.25 | 0.31 | 0.27 | 0.53 | 0.53 | 1.00 | 0.52 | 1.00 | 0.64 | 0.62 | 1.00 | 0.23 | 0.28 | 0.25 | 0.41 | 0.84 | 0.76 | 0.26 |
| CELLama | 0.38 | 0.21 | 0.19 | 0.74 | 0.74 | 0.67 | 0.45 | 0.00 | 0.68 | 0.67 | 1.00 | 0.28 | 0.32 | 0.38 | 0.47 | 0.37 | 0.78 | 0.30 |
| GenePT-w | 0.43 | 0.23 | 0.21 | 0.46 | 0.45 | 0.67 | 0.45 | 0.00 | 0.55 | 0.54 | 1.00 | 0.28 | 0.32 | 0.43 | 0.33 | 0.37 | 0.70 | 0.30 |
| scBERT | 0.34 | 0.24 | 0.22 | 0.42 | 0.39 | 0.67 | 0.45 | 0.00 | 0.54 | 0.51 | 1.00 | 0.28 | 0.30 | 0.34 | 0.32 | 0.37 | 0.68 | 0.29 |
| scGPT | 0.34 | 0.17 | 0.16 | 0.35 | 0.36 | 0.67 | 0.45 | 0.00 | 0.53 | 0.50 | 1.00 | 0.28 | 0.31 | 0.34 | 0.26 | 0.37 | 0.67 | 0.29 |

**Figure S139: Trajectory-inference metric scores for mesoderm development using Monocle 3.** Display and scoring conventions are as in Fig. S104.

| | Cell positions | Neighbourhood | | | | Topology | | | Features | | | Branch assignment | | Aggregated scores | | | | |
|---|---|---|---|---|---|---|---|---|---|---|---|---|---|---|---|---|---|---|
| method | $cor_{dist}$ | $NMSE_{lm}$ | $R^2_{lm}$ | $NMSE_{rf}$ | $R^2_{rf}$ | edgeflip | HIM | isomorphic | $cor_{features}$ | $wcor_{features}$ | $ks_{features}$ | $F1_{branches}$ | $F1_{milestones}$ | Pos. | Neighbor. | Topology | Feature | Branch |
| scFoundation | 0.49 | 0.25 | 0.22 | 0.52 | 0.44 | | | | 0.60 | 0.54 | 1.00 | | 0.37 | 0.49 | 0.36 | | 0.71 | 0.37 |
| scGPT (integrated | 0.50 | 0.23 | 0.22 | 0.33 | 0.32 | | | | 0.52 | 0.49 | 1.00 | | 0.40 | 0.50 | 0.28 | | 0.67 | 0.40 |
| CellPLM | 0.49 | 0.26 | 0.24 | 0.21 | 0.18 | | | | 0.64 | 0.59 | 1.00 | | 0.38 | 0.49 | 0.22 | | 0.74 | 0.38 |
| SCimilarity | 0.44 | 0.25 | 0.22 | 0.36 | 0.29 | | | | 0.64 | 0.60 | 1.00 | | 0.36 | 0.44 | 0.28 | | 0.75 | 0.36 |
| Geneformer | 0.45 | 0.22 | 0.21 | 0.39 | 0.36 | | | | 0.57 | 0.49 | 1.00 | | 0.40 | 0.45 | 0.29 | | 0.69 | 0.40 |
| scBERT | 0.37 | 0.23 | 0.21 | 0.17 | 0.14 | | | | 0.50 | 0.49 | 1.00 | | 0.39 | 0.37 | 0.19 | | 0.66 | 0.39 |
| UCE | 0.42 | 0.19 | 0.19 | 0.06 | 0.04 | | | | 0.48 | 0.44 | 1.00 | | 0.39 | 0.42 | 0.12 | | 0.64 | 0.39 |
| scPRINT | 0.27 | 0.18 | 0.16 | 0.22 | 0.17 | | | | 0.61 | 0.60 | 1.00 | | 0.34 | 0.27 | 0.18 | | 0.73 | 0.34 |
| CELLama | 0.34 | 0.20 | 0.17 | 0.30 | 0.33 | | | | 0.55 | 0.55 | 1.00 | | 0.24 | 0.34 | 0.25 | | 0.70 | 0.24 |
| CellFM | 0.30 | 0.22 | 0.19 | 0.05 | 0.00 | | | | 0.58 | 0.55 | 1.00 | | 0.29 | 0.30 | 0.11 | | 0.71 | 0.29 |
| scGPT | 0.34 | 0.16 | 0.15 | 0.00 | 0.00 | | | | 0.41 | 0.36 | 1.00 | | 0.36 | 0.34 | 0.08 | | 0.59 | 0.36 |
| scVI | 0.10 | 0.17 | 0.15 | 0.00 | 0.00 | | | | 0.56 | 0.50 | 1.00 | | 0.42 | 0.10 | 0.08 | | 0.69 | 0.42 |
| LangCell | 0.26 | 0.07 | 0.06 | 0.11 | 0.08 | | | | 0.51 | 0.49 | 1.00 | | 0.25 | 0.26 | 0.08 | | 0.67 | 0.25 |
| PCA | 0.20 | 0.11 | 0.09 | 0.11 | 0.09 | | | | 0.45 | 0.48 | 1.00 | | 0.23 | 0.20 | 0.10 | | 0.64 | 0.23 |
| scVI (de novo) | 0.18 | 0.19 | 0.16 | 0.00 | 0.00 | | | | 0.51 | 0.48 | 1.00 | | 0.24 | 0.18 | 0.09 | | 0.66 | 0.24 |
| scCello | 0.13 | 0.14 | 0.12 | 0.03 | 0.00 | | | | 0.53 | 0.53 | 1.00 | | 0.23 | 0.13 | 0.07 | | 0.69 | 0.23 |
| GenePT-w | 0.04 | 0.03 | 0.03 | 0.00 | 0.00 | | | | 0.28 | 0.27 | 1.00 | | 0.25 | 0.04 | 0.01 | | 0.52 | 0.25 |

**Figure S140: Trajectory-inference metric scores for mesoderm development using DPT.** Display and scoring conventions are as in Fig. S105.

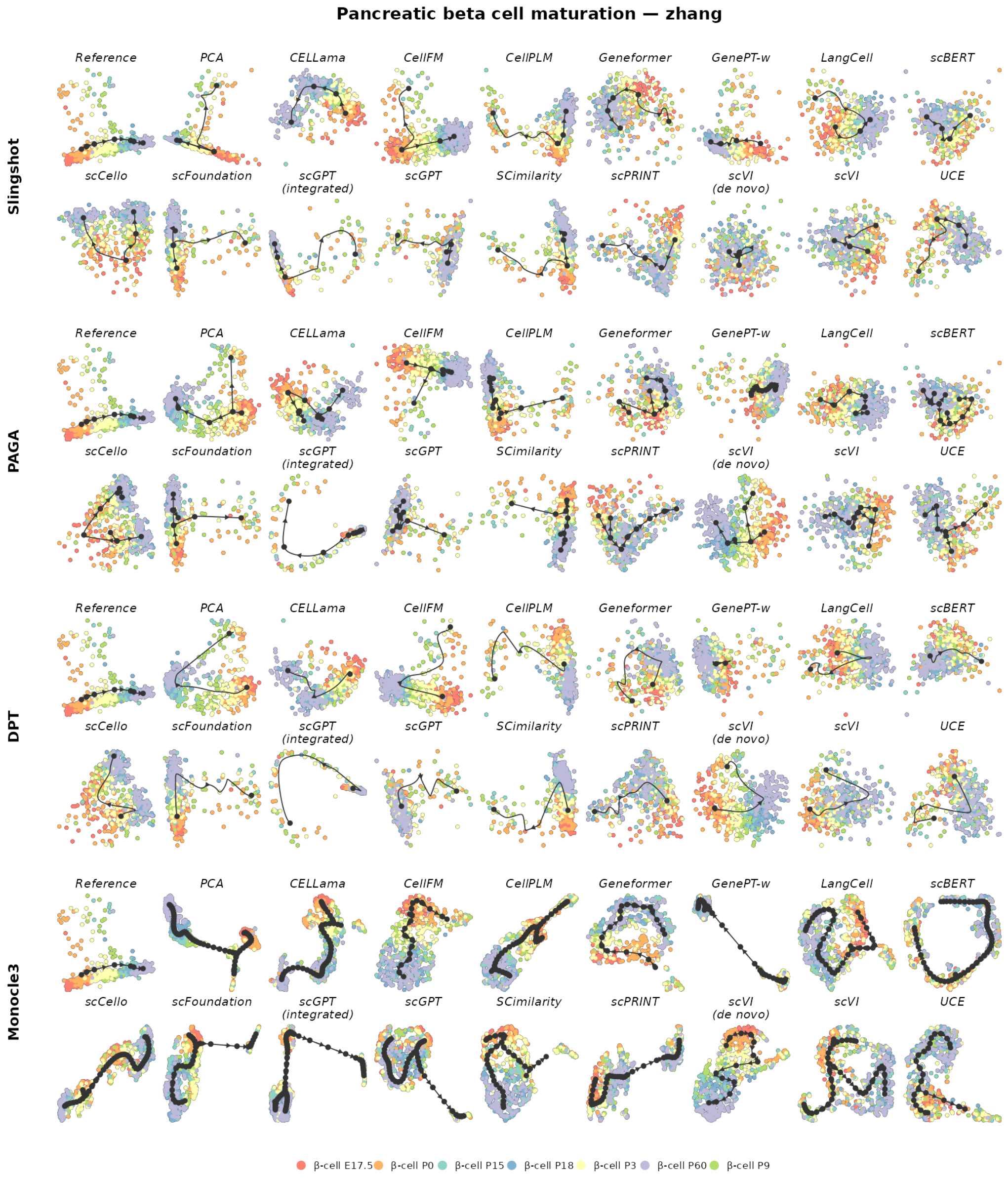

**Figure S141: Trajectory inference on pancreatic beta-cell maturation.** The reference trajectory is linear. Rows show Slingshot, PAGA, DPT and Monocle 3; within each row, the reference is followed by the same 17 representations. Cells are coloured by the reference milestone or state. Black nodes, edges and arrows show the inferred milestone graph and its orientation. Layouts and trajectories were reconstructed independently for each representation and backend.

| | Cell positions | Neighbourhood | | | | Topology | | | Features | | | Branch assignment | | Aggregated scores | | | | |
|---|---|---|---|---|---|---|---|---|---|---|---|---|---|---|---|---|---|---|
| method | $cor_{dist}$ | $NMSE_{lm}$ | $R^2_{lm}$ | $NMSE_{rf}$ | $R^2_{rf}$ | edgeflip | HIM | isomorphic | $cor_{features}$ | $wcor_{features}$ | $ks_{features}$ | $F1_{branches}$ | $F1_{milestones}$ | Pos. | Neighbor. | Topology | Feature | Branch |
| CELLama | 0.50 | 0.45 | 0.45 | 0.44 | 0.44 | 1.00 | 1.00 | 1.00 | 0.83 | 0.82 | 1.00 | 1.00 | 0.62 | 0.50 | 0.45 | 1.00 | 0.88 | 0.81 |
| scVI (de novo) | 0.54 | 0.56 | 0.56 | 0.58 | 0.58 | 1.00 | 1.00 | 0.00 | 0.84 | 0.84 | 1.00 | 0.64 | 0.45 | 0.54 | 0.57 | 0.67 | 0.89 | 0.55 |
| GenePT-w | 0.40 | 0.35 | 0.35 | 0.19 | 0.19 | 1.00 | 1.00 | 1.00 | 0.71 | 0.66 | 1.00 | 1.00 | 0.45 | 0.40 | 0.27 | 1.00 | 0.79 | 0.72 |
| scBERT | 0.24 | 0.22 | 0.22 | 0.05 | 0.05 | 1.00 | 1.00 | 1.00 | 0.80 | 0.84 | 1.00 | 1.00 | 0.44 | 0.24 | 0.13 | 1.00 | 0.88 | 0.72 |
| CellFM | 0.38 | 0.53 | 0.53 | 0.53 | 0.54 | 1.00 | 1.00 | 0.00 | 0.73 | 0.78 | 1.00 | 0.62 | 0.38 | 0.38 | 0.53 | 0.67 | 0.84 | 0.50 |
| SCimilarity | 0.28 | 0.43 | 0.43 | 0.42 | 0.42 | 1.00 | 1.00 | 0.00 | 0.71 | 0.77 | 1.00 | 0.63 | 0.41 | 0.28 | 0.42 | 0.67 | 0.83 | 0.52 |
| CellPLM | 0.23 | 0.43 | 0.43 | 0.42 | 0.42 | 1.00 | 1.00 | 0.00 | 0.70 | 0.79 | 0.98 | 0.63 | 0.40 | 0.23 | 0.42 | 0.67 | 0.83 | 0.51 |
| scCello | 0.18 | 0.33 | 0.33 | 0.35 | 0.35 | 1.00 | 1.00 | 0.00 | 0.63 | 0.70 | 1.00 | 0.54 | 0.32 | 0.18 | 0.34 | 0.67 | 0.78 | 0.43 |
| scPRINT | 0.08 | 0.03 | 0.03 | 0.00 | 0.00 | 1.00 | 1.00 | 1.00 | 0.24 | 0.42 | 1.00 | 1.00 | 0.43 | 0.08 | 0.01 | 1.00 | 0.56 | 0.71 |
| scGPT (integrated) | 0.19 | 0.28 | 0.28 | 0.26 | 0.26 | 1.00 | 1.00 | 0.00 | 0.53 | 0.62 | 1.00 | 0.63 | 0.35 | 0.19 | 0.27 | 0.67 | 0.72 | 0.49 |
| PCA | 0.20 | 0.43 | 0.43 | 0.55 | 0.55 | 0.67 | 0.56 | 0.00 | 0.66 | 0.61 | 1.00 | 0.44 | 0.30 | 0.20 | 0.49 | 0.41 | 0.76 | 0.37 |
| Geneformer | 0.12 | 0.17 | 0.17 | 0.27 | 0.27 | 1.00 | 1.00 | 0.00 | 0.48 | 0.72 | 0.95 | 0.62 | 0.32 | 0.12 | 0.22 | 0.67 | 0.72 | 0.47 |
| scFoundation | 0.22 | 0.37 | 0.37 | 0.42 | 0.42 | 0.67 | 0.51 | 0.00 | 0.72 | 0.78 | 1.00 | 0.39 | 0.32 | 0.22 | 0.39 | 0.39 | 0.83 | 0.36 |
| UCE | 0.07 | 0.19 | 0.19 | 0.18 | 0.19 | 1.00 | 1.00 | 0.00 | 0.49 | 0.64 | 1.00 | 0.62 | 0.34 | 0.07 | 0.19 | 0.67 | 0.71 | 0.48 |
| LangCell | 0.17 | 0.35 | 0.35 | 0.25 | 0.26 | 0.67 | 0.51 | 0.00 | 0.61 | 0.64 | 1.00 | 0.35 | 0.31 | 0.17 | 0.30 | 0.39 | 0.75 | 0.33 |
| scVI | 0.11 | 0.25 | 0.25 | 0.22 | 0.22 | 0.67 | 0.56 | 0.00 | 0.49 | 0.66 | 1.00 | 0.41 | 0.31 | 0.11 | 0.23 | 0.41 | 0.72 | 0.36 |
| scGPT | 0.02 | 0.13 | 0.13 | 0.13 | 0.13 | 0.67 | 0.44 | 0.00 | 0.20 | 0.62 | 0.99 | 0.42 | 0.29 | 0.02 | 0.13 | 0.37 | 0.60 | 0.35 |

**Figure S142: Trajectory-inference metric scores for pancreatic beta-cell maturation using Slingshot.** Display and scoring conventions are as in Fig. S102.

| | Cell positions | Neighbourhood | | | | Topology | | | Features | | | Branch assignment | | Aggregated scores | | | | |
|---|---|---|---|---|---|---|---|---|---|---|---|---|---|---|---|---|---|---|
| method | $cor_{dist}$ | $NMSE_{lm}$ | $R^2_{lm}$ | $NMSE_{rf}$ | $R^2_{rf}$ | edgeflip | HIM | isomorphic | $cor_{features}$ | $wcor_{features}$ | $ks_{features}$ | $F1_{branches}$ | $F1_{milestones}$ | Pos. | Neighbor. | Topology | Feature | Branch |
| PCA | 0.68 | 0.74 | 0.74 | 0.89 | 0.89 | 0.67 | 0.54 | 0.00 | 0.72 | 0.75 | 1.00 | 0.43 | 0.31 | 0.68 | 0.82 | 0.40 | 0.83 | 0.37 |
| CELLama | 0.36 | 0.35 | 0.35 | 0.44 | 0.44 | 1.00 | 1.00 | 0.00 | 0.63 | 0.78 | 1.00 | 0.54 | 0.44 | 0.36 | 0.39 | 0.67 | 0.80 | 0.49 |
| GenePT-w | 0.37 | 0.33 | 0.33 | 0.34 | 0.34 | 1.00 | 1.00 | 0.00 | 0.62 | 0.57 | 1.00 | 0.64 | 0.39 | 0.37 | 0.33 | 0.67 | 0.73 | 0.51 |
| scVI (de novo) | 0.46 | 0.62 | 0.62 | 0.61 | 0.61 | 0.40 | 0.40 | 0.00 | 0.79 | 0.87 | 1.00 | 0.25 | 0.34 | 0.46 | 0.62 | 0.27 | 0.89 | 0.29 |
| LangCell | 0.27 | 0.25 | 0.25 | 0.28 | 0.28 | 1.00 | 1.00 | 0.00 | 0.48 | 0.57 | 1.00 | 0.62 | 0.36 | 0.27 | 0.26 | 0.67 | 0.68 | 0.49 |
| CellPLM | 0.41 | 0.45 | 0.45 | 0.53 | 0.53 | 0.40 | 0.40 | 0.00 | 0.65 | 0.86 | 1.00 | 0.24 | 0.23 | 0.41 | 0.49 | 0.27 | 0.84 | 0.24 |
| Geneformer | 0.32 | 0.32 | 0.32 | 0.37 | 0.37 | 0.67 | 0.59 | 0.00 | 0.56 | 0.65 | 1.00 | 0.45 | 0.29 | 0.32 | 0.34 | 0.42 | 0.74 | 0.37 |
| scFoundation | 0.38 | 0.51 | 0.51 | 0.48 | 0.48 | 0.33 | 0.33 | 0.00 | 0.63 | 0.77 | 1.00 | 0.23 | 0.25 | 0.38 | 0.49 | 0.22 | 0.80 | 0.24 |
| SCimilarity | 0.32 | 0.42 | 0.42 | 0.44 | 0.44 | 0.40 | 0.43 | 0.00 | 0.72 | 0.83 | 1.00 | 0.27 | 0.26 | 0.32 | 0.43 | 0.28 | 0.85 | 0.27 |
| CellFM | 0.39 | 0.37 | 0.37 | 0.36 | 0.36 | 0.40 | 0.38 | 0.00 | 0.69 | 0.69 | 1.00 | 0.27 | 0.25 | 0.39 | 0.36 | 0.26 | 0.79 | 0.26 |
| scCello | 0.12 | 0.29 | 0.29 | 0.33 | 0.33 | 0.67 | 0.49 | 0.00 | 0.56 | 0.70 | 1.00 | 0.43 | 0.31 | 0.12 | 0.31 | 0.39 | 0.76 | 0.37 |
| scVI | 0.20 | 0.30 | 0.30 | 0.31 | 0.31 | 0.50 | 0.44 | 0.00 | 0.45 | 0.69 | 1.00 | 0.25 | 0.26 | 0.20 | 0.31 | 0.31 | 0.71 | 0.25 |
| scGPT (integrated) | 0.18 | 0.29 | 0.29 | 0.30 | 0.30 | 0.40 | 0.42 | 0.00 | 0.46 | 0.65 | 1.00 | 0.26 | 0.25 | 0.18 | 0.30 | 0.27 | 0.70 | 0.25 |
| UCE | 0.07 | 0.18 | 0.18 | 0.20 | 0.20 | 0.67 | 0.56 | 0.00 | 0.41 | 0.55 | 0.99 | 0.33 | 0.24 | 0.07 | 0.19 | 0.41 | 0.65 | 0.28 |
| scBERT | 0.19 | 0.27 | 0.27 | 0.25 | 0.25 | 0.29 | 0.35 | 0.00 | 0.61 | 0.66 | 1.00 | 0.18 | 0.18 | 0.19 | 0.26 | 0.21 | 0.76 | 0.18 |
| scPRINT | 0.07 | 0.11 | 0.11 | 0.12 | 0.13 | 0.67 | 0.47 | 0.00 | 0.37 | 0.53 | 0.95 | 0.43 | 0.28 | 0.07 | 0.12 | 0.38 | 0.62 | 0.36 |
| scGPT | 0.03 | 0.13 | 0.13 | 0.11 | 0.11 | 0.40 | 0.41 | 0.00 | 0.14 | 0.43 | 1.00 | 0.23 | 0.18 | 0.03 | 0.12 | 0.27 | 0.52 | 0.21 |

**Figure S143: Trajectory-inference metric scores for pancreatic beta-cell maturation using PAGA.** Display and scoring conventions are as in Fig. S103.

| | Cell positions | Neighbourhood | | | | Topology | | | Features | | | Branch assignment | | Aggregated scores | | | | |
|---|---|---|---|---|---|---|---|---|---|---|---|---|---|---|---|---|---|---|
| method | $cor_{dist}$ | $NMSE_{lm}$ | $R^2_{lm}$ | $NMSE_{rf}$ | $R^2_{rf}$ | edgeflip | HIM | isomorphic | $cor_{features}$ | $wcor_{features}$ | $ks_{features}$ | $F1_{branches}$ | $F1_{milestones}$ | Pos. | Neighbor. | Topology | Feature | Branch |
| Geneformer | 0.32 | 0.26 | 0.26 | 0.39 | 0.39 | 1.00 | 1.00 | 1.00 | 0.72 | 0.69 | 1.00 | 1.00 | 0.48 | 0.32 | 0.33 | 1.00 | 0.80 | 0.74 |
| scVI (de novo) | 0.50 | 0.54 | 0.54 | 0.56 | 0.56 | 1.00 | 1.00 | 0.00 | 0.80 | 0.76 | 1.00 | 0.64 | 0.41 | 0.50 | 0.55 | 0.67 | 0.85 | 0.53 |
| PCA | 0.73 | 0.76 | 0.76 | 0.85 | 0.85 | 0.50 | 0.61 | 0.00 | 0.72 | 0.85 | 1.00 | 0.30 | 0.29 | 0.73 | 0.81 | 0.37 | 0.86 | 0.29 |
| CellFM | 0.35 | 0.49 | 0.49 | 0.48 | 0.48 | 0.67 | 0.71 | 0.00 | 0.70 | 0.73 | 1.00 | 0.45 | 0.33 | 0.35 | 0.49 | 0.46 | 0.81 | 0.39 |
| CELLama | 0.45 | 0.41 | 0.41 | 0.47 | 0.47 | 0.67 | 0.60 | 0.00 | 0.64 | 0.75 | 1.00 | 0.36 | 0.30 | 0.45 | 0.44 | 0.42 | 0.79 | 0.33 |
| SCimilarity | 0.41 | 0.49 | 0.49 | 0.51 | 0.51 | 0.40 | 0.51 | 0.00 | 0.67 | 0.80 | 1.00 | 0.24 | 0.25 | 0.41 | 0.50 | 0.30 | 0.82 | 0.24 |
| scFoundation | 0.29 | 0.33 | 0.33 | 0.54 | 0.54 | 0.67 | 0.60 | 0.00 | 0.54 | 0.73 | 0.98 | 0.37 | 0.30 | 0.29 | 0.43 | 0.42 | 0.75 | 0.34 |
| LangCell | 0.32 | 0.32 | 0.32 | 0.41 | 0.41 | 0.67 | 0.62 | 0.00 | 0.59 | 0.57 | 1.00 | 0.36 | 0.25 | 0.32 | 0.36 | 0.43 | 0.72 | 0.31 |
| scCello | 0.21 | 0.35 | 0.35 | 0.38 | 0.38 | 0.67 | 0.52 | 0.00 | 0.66 | 0.71 | 1.00 | 0.37 | 0.28 | 0.21 | 0.37 | 0.40 | 0.79 | 0.32 |
| CellPLM | 0.31 | 0.47 | 0.47 | 0.51 | 0.51 | 0.40 | 0.46 | 0.00 | 0.45 | 0.79 | 1.00 | 0.22 | 0.24 | 0.31 | 0.49 | 0.29 | 0.74 | 0.23 |
| scBERT | 0.21 | 0.28 | 0.28 | 0.29 | 0.29 | 0.67 | 0.65 | 0.00 | 0.75 | 0.76 | 1.00 | 0.31 | 0.29 | 0.21 | 0.28 | 0.44 | 0.84 | 0.30 |
| GenePT-w | 0.31 | 0.40 | 0.40 | 0.41 | 0.41 | 0.40 | 0.56 | 0.00 | 0.66 | 0.66 | 1.00 | 0.24 | 0.25 | 0.31 | 0.41 | 0.32 | 0.77 | 0.24 |
| scPRINT | 0.08 | 0.07 | 0.07 | 0.20 | 0.20 | 1.00 | 1.00 | 0.00 | 0.37 | 0.49 | 1.00 | 0.64 | 0.32 | 0.08 | 0.14 | 0.67 | 0.62 | 0.48 |
| scVI | 0.23 | 0.34 | 0.34 | 0.35 | 0.35 | 0.40 | 0.50 | 0.00 | 0.66 | 0.79 | 1.00 | 0.25 | 0.22 | 0.23 | 0.35 | 0.30 | 0.82 | 0.23 |
| UCE | 0.18 | 0.24 | 0.24 | 0.22 | 0.23 | 0.67 | 0.56 | 0.00 | 0.47 | 0.60 | 1.00 | 0.40 | 0.27 | 0.18 | 0.23 | 0.41 | 0.69 | 0.34 |
| scGPT (integrated | 0.18 | 0.34 | 0.34 | 0.32 | 0.33 | 0.29 | 0.47 | 0.00 | 0.42 | 0.63 | 1.00 | 0.17 | 0.19 | 0.18 | 0.33 | 0.25 | 0.68 | 0.18 |
| scGPT | 0.01 | 0.10 | 0.10 | 0.18 | 0.18 | 0.67 | 0.61 | 0.00 | 0.10 | 0.34 | 1.00 | 0.43 | 0.26 | 0.01 | 0.14 | 0.42 | 0.48 | 0.35 |

**Figure S144: Trajectory-inference metric scores for pancreatic beta-cell maturation using Monocle 3.** Display and scoring conventions are as in Fig. S104.

| | Cell positions | Neighbourhood | | | | Topology | | | Features | | | Branch assignment | | Aggregated scores | | | | |
|---|---|---|---|---|---|---|---|---|---|---|---|---|---|---|---|---|---|---|
| method | $cor_{dist}$ | $NMSE_{lm}$ | $R^2_{lm}$ | $NMSE_{rf}$ | $R^2_{rf}$ | edgeflip | HIM | isomorphic | $cor_{features}$ | $wcor_{features}$ | $ks_{features}$ | $F1_{branches}$ | $F1_{milestones}$ | Pos. | Neighbor. | Topology | Feature | Branch |
| PCA | 0.54 | 0.38 | 0.38 | 0.70 | 0.71 | | | | 0.71 | 0.82 | 1.00 | | 0.67 | 0.54 | 0.54 | | 0.84 | 0.67 |
| CELLama | 0.34 | 0.32 | 0.32 | 0.18 | 0.18 | | | | 0.73 | 0.79 | 1.00 | | 0.54 | 0.34 | 0.25 | | 0.84 | 0.54 |
| scVI (de novo) | 0.30 | 0.12 | 0.12 | 0.38 | 0.38 | | | | 0.39 | 0.72 | 1.00 | | 0.54 | 0.30 | 0.25 | | 0.70 | 0.54 |
| CellFM | 0.24 | 0.12 | 0.12 | 0.15 | 0.15 | | | | 0.53 | 0.68 | 1.00 | | 0.43 | 0.24 | 0.13 | | 0.74 | 0.43 |
| scBERT | 0.12 | 0.15 | 0.15 | 0.00 | 0.00 | | | | 0.74 | 0.80 | 0.99 | | 0.41 | 0.12 | 0.07 | | 0.84 | 0.41 |
| scFoundation | 0.33 | 0.01 | 0.01 | 0.41 | 0.41 | | | | 0.05 | 0.33 | 1.00 | | 0.43 | 0.33 | 0.21 | | 0.46 | 0.43 |
| scVI | 0.18 | 0.10 | 0.10 | 0.06 | 0.06 | | | | 0.30 | 0.65 | 1.00 | | 0.48 | 0.18 | 0.08 | | 0.65 | 0.48 |
| LangCell | 0.25 | 0.01 | 0.01 | 0.21 | 0.21 | | | | 0.06 | 0.34 | 1.00 | | 0.43 | 0.25 | 0.11 | | 0.47 | 0.43 |
| Geneformer | 0.17 | 0.01 | 0.01 | 0.11 | 0.11 | | | | 0.08 | 0.50 | 1.00 | | 0.43 | 0.17 | 0.06 | | 0.53 | 0.43 |
| CellPLM | 0.23 | 0.02 | 0.02 | 0.28 | 0.28 | | | | 0.00 | 0.00 | 1.00 | | 0.43 | 0.23 | 0.15 | | 0.33 | 0.43 |
| scPRINT | 0.07 | 0.01 | 0.01 | 0.00 | 0.00 | | | | 0.09 | 0.24 | 0.93 | | 0.43 | 0.07 | 0.01 | | 0.42 | 0.43 |
| scGPT (integrated | 0.11 | 0.01 | 0.01 | 0.03 | 0.03 | | | | 0.00 | 0.07 | 0.99 | | 0.43 | 0.11 | 0.02 | | 0.35 | 0.43 |
| GenePT-w | 0.00 | 0.00 | 0.00 | 0.00 | 0.00 | | | | 0.24 | 0.21 | 0.97 | | 0.38 | 0.00 | 0.00 | | 0.48 | 0.38 |
| UCE | 0.06 | 0.00 | 0.00 | 0.00 | 0.00 | | | | 0.01 | 0.07 | 1.00 | | 0.43 | 0.06 | 0.00 | | 0.36 | 0.43 |
| SCimilarity | 0.06 | 0.00 | 0.00 | 0.00 | 0.00 | | | | 0.00 | 0.03 | 1.00 | | 0.43 | 0.06 | 0.00 | | 0.34 | 0.43 |
| scCello | 0.03 | 0.02 | 0.02 | 0.00 | 0.00 | | | | 0.06 | 0.15 | 0.98 | | 0.39 | 0.03 | 0.01 | | 0.40 | 0.39 |
| scGPT | 0.00 | 0.01 | 0.01 | 0.00 | 0.00 | | | | 0.00 | 0.00 | 0.99 | | 0.43 | 0.00 | 0.00 | | 0.33 | 0.43 |

**Figure S145: Trajectory-inference metric scores for pancreatic beta-cell maturation using DPT.** Display and scoring conventions are as in Fig. S105.

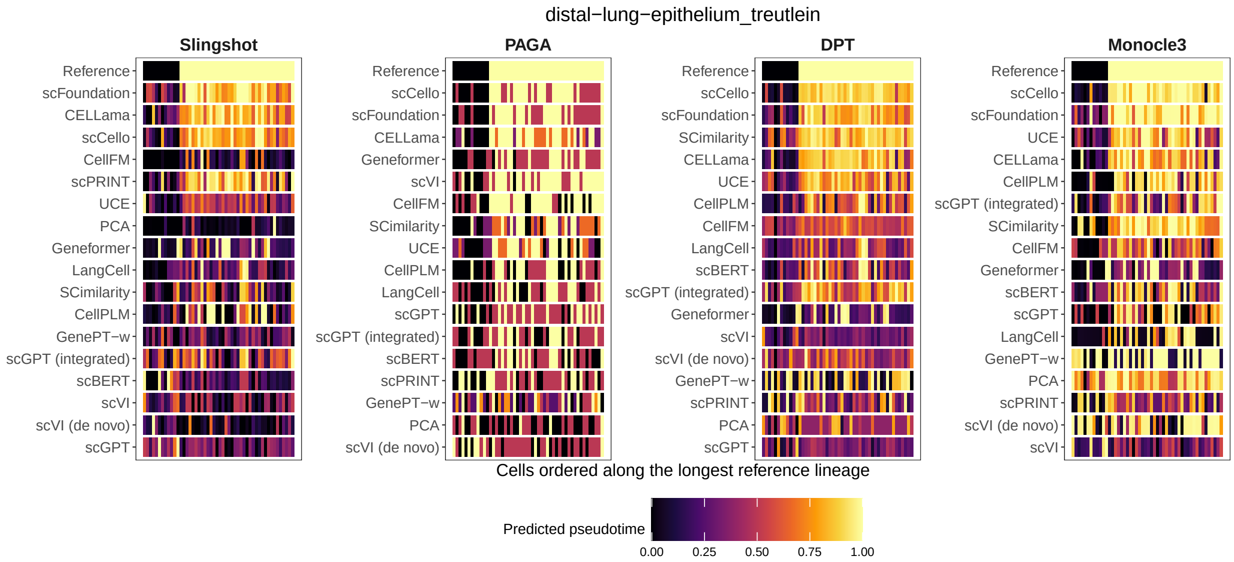

**Figure S146: Pseudotime ribbon plot for distal lung epithelium.** Columns show four trajectory-inference backends and rows show the reference progression followed by the 13 scFM representations and PCA. For this display, cells were restricted to the longest reference lineage, BP → AT1, and ordered by reference progression along that path. Each cell is represented by one borderless tile; colour encodes predicted pseudotime after min–max normalization within each dataset–backend–representation panel. Monocle 3 and DPT use their exported cell-level pseudotime, whereas Slingshot and PAGA use pseudotime derived from the inferred Dynwrap progressions. The reference row contains 50 cells and is shown on the same 0–1 colour scale to indicate the ordering used for visual comparison.

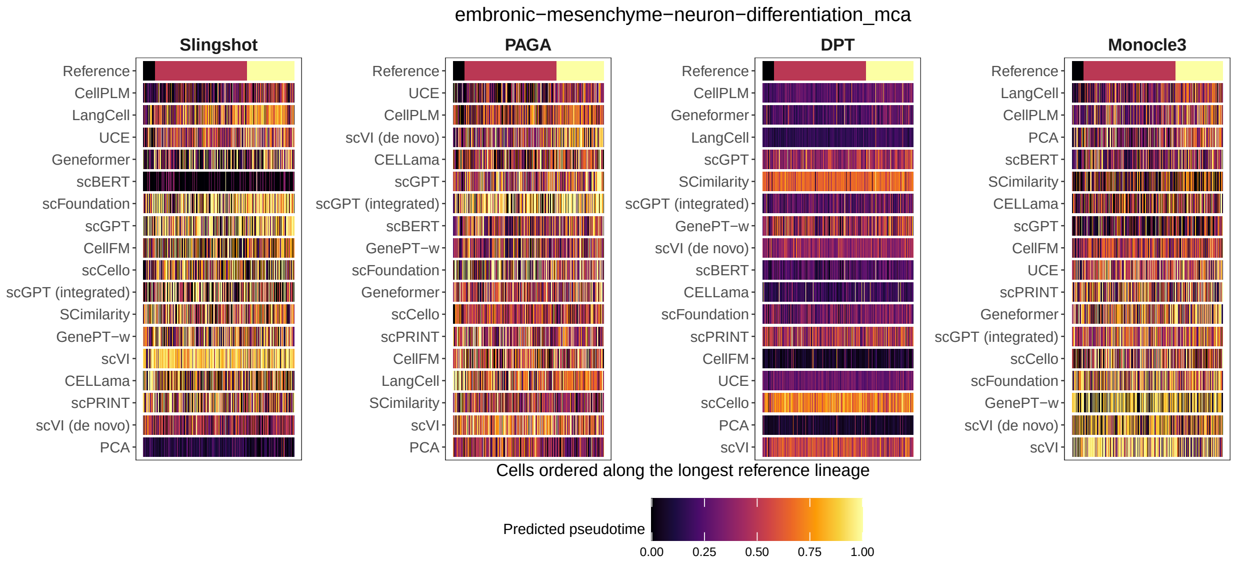

**Figure S147: Pseudotime ribbon plot for embryonic mesenchyme-to-neuron differentiation.** Columns show four trajectory-inference backends and rows show the reference progression followed by the 13 scFM representations and PCA. For this display, cells were restricted to the longest reference lineage, neuronal progenitors → Igfbpl1-high neurons → Mapt-high ganglion cells, and ordered by reference progression along that path. Each cell is represented by one borderless tile; colour encodes predicted pseudotime after min–max normalization within each dataset–backend–representation panel. Monocle 3 and DPT use their exported cell-level pseudotime, whereas Slingshot and PAGA use pseudotime derived from the inferred Dynwrap progressions. The reference row contains 350 cells and is shown on the same 0–1 colour scale to indicate the ordering used for visual comparison.

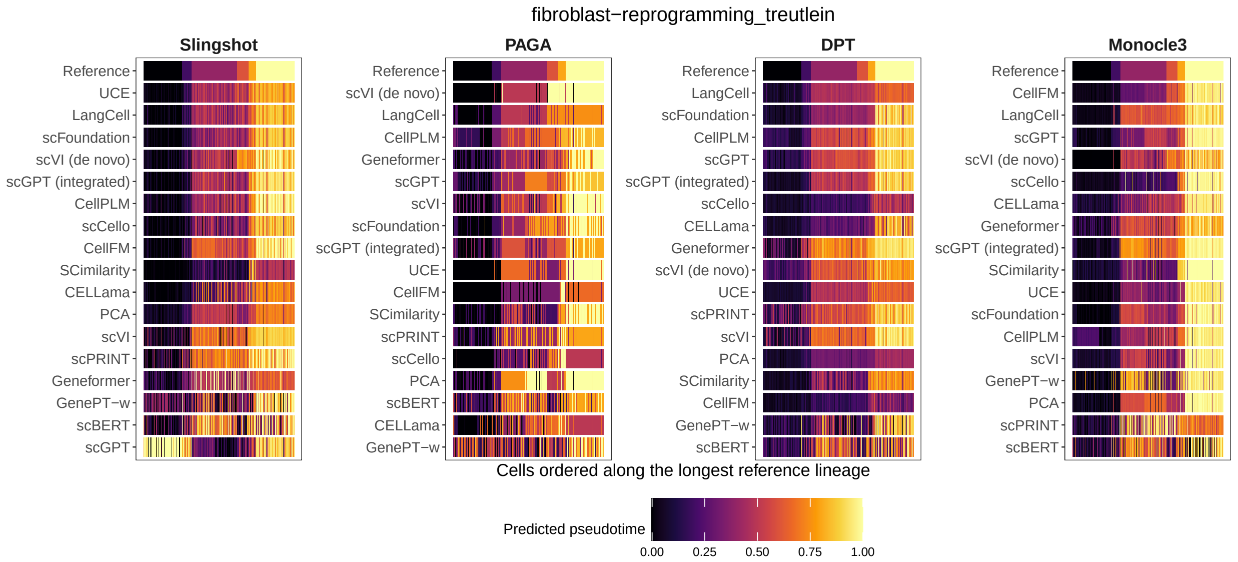

**Figure S148: Pseudotime ribbon plot for fibroblast reprogramming.** Columns show four trajectory-inference backends and rows show the reference progression followed by the 13 scFM representations and PCA. For this display, cells were restricted to the longest reference lineage, MEF → day-2 intermediate → day-2 induced → day-5 intermediate → day-5 early induced neuron → myocyte, and ordered by reference progression along that path. Each cell is represented by one borderless tile; colour encodes predicted pseudotime after min–max normalization within each dataset–backend–representation panel. Monocle 3 and DPT use their exported cell-level pseudotime, whereas Slingshot and PAGA use pseudotime derived from the inferred Dynwrap progressions. The reference row contains 323 cells and is shown on the same 0–1 colour scale to indicate the ordering used for visual comparison.

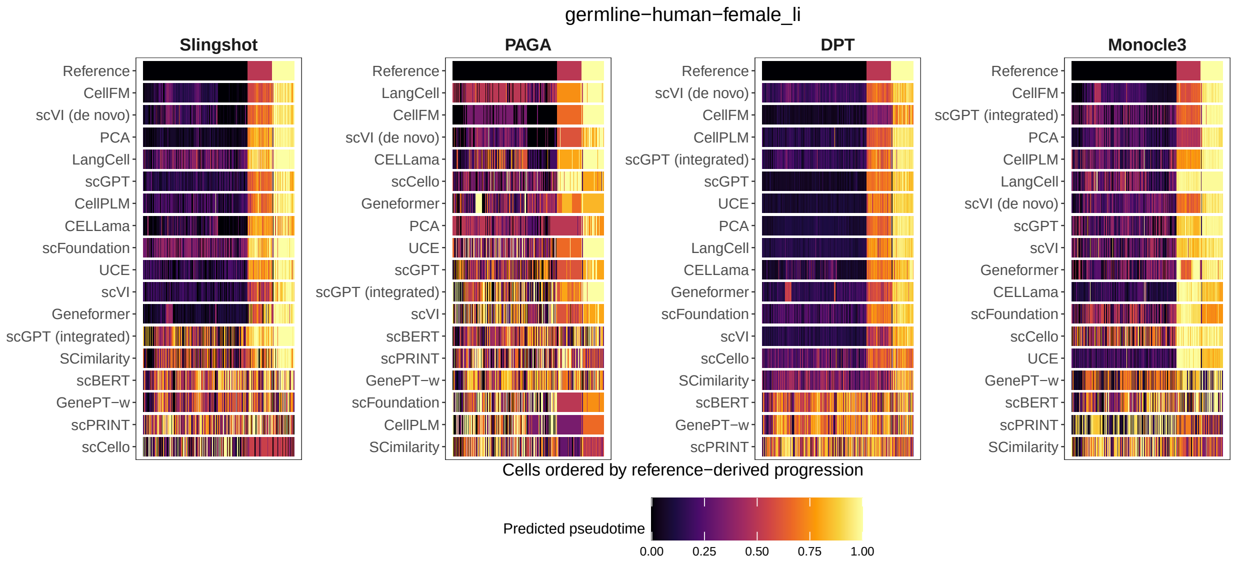

**Figure S149: Pseudotime ribbon plot for human female germline development.** Columns show four trajectory-inference backends and rows show the reference progression followed by the 13 scFM representations and PCA. For this display, cells were ordered by the complete reference-derived progression for this linear trajectory. Each cell is represented by one borderless tile; colour encodes predicted pseudotime after min–max normalization within each dataset–backend–representation panel. Monocle 3 and DPT use their exported cell-level pseudotime, whereas Slingshot and PAGA use pseudotime derived from the inferred Dynwrap progressions. The reference row contains 621 cells and is shown on the same 0–1 colour scale to indicate the ordering used for visual comparison.

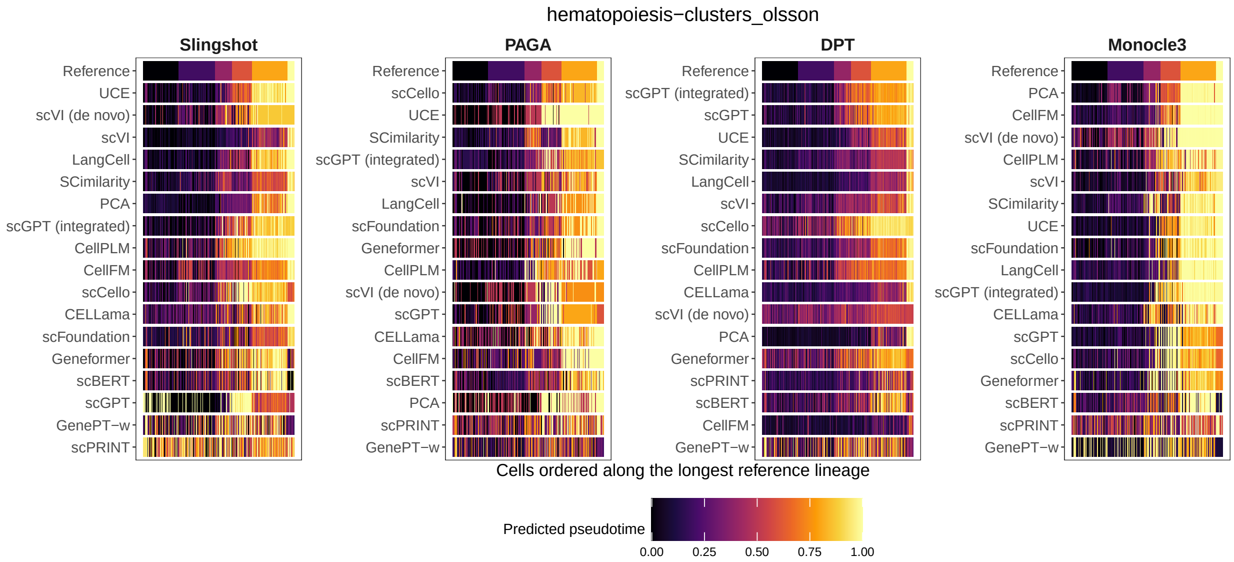

**Figure S150: Pseudotime ribbon plot for haematopoiesis.** Columns show four trajectory-inference backends and rows show the reference progression followed by the 13 scFM representations and PCA. For this display, cells were restricted to the longest reference lineage, HSCP-1 → HSCP-2 → multi-lineage progenitor → MDP → granulocyte → myelocyte, and ordered by reference progression along that path. Each cell is represented by one borderless tile; colour encodes predicted pseudotime after min–max normalization within each dataset–backend–representation panel. Monocle 3 and DPT use their exported cell-level pseudotime, whereas Slingshot and PAGA use pseudotime derived from the inferred Dynwrap progressions. The reference row contains 250 cells and is shown on the same 0–1 colour scale to indicate the ordering used for visual comparison.

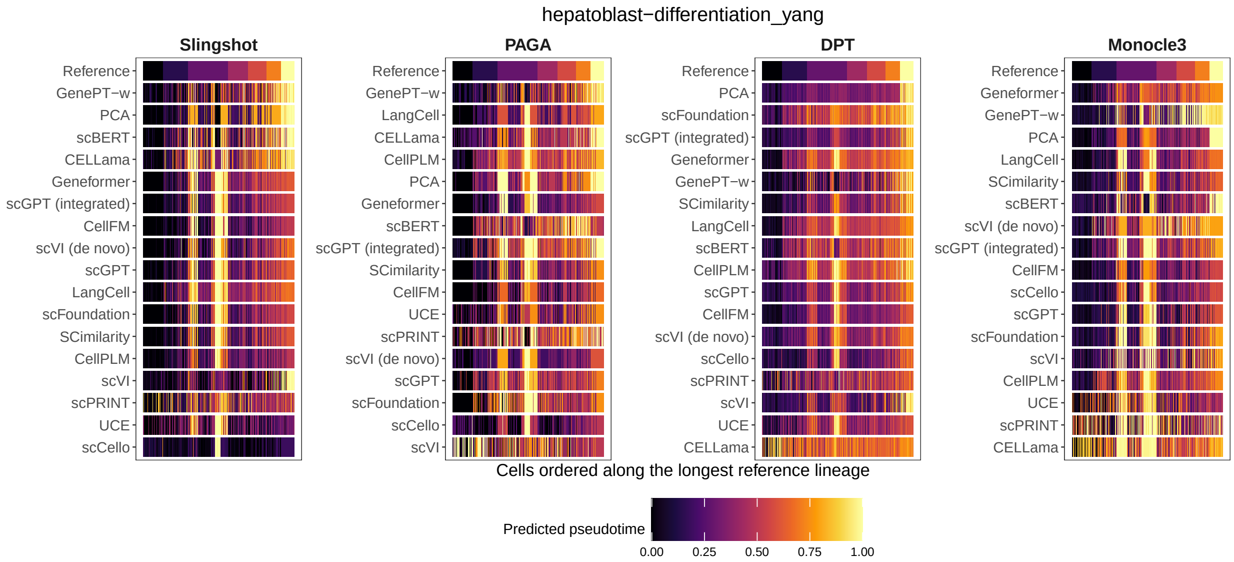

**Figure S151: Pseudotime ribbon plot for hepatoblast differentiation.** Columns show four trajectory-inference backends and rows show the reference progression followed by the 13 scFM representations and PCA. For this display, cells were restricted to the longest reference hepatoblast/hepatocyte lineage from E10.5 through E17.5 and ordered by reference progression along that path. Each cell is represented by one borderless tile; colour encodes predicted pseudotime after min–max normalization within each dataset–backend–representation panel. Monocle 3 and DPT use their exported cell-level pseudotime, whereas Slingshot and PAGA use pseudotime derived from the inferred Dynwrap progressions. The reference row contains 417 cells and is shown on the same 0–1 colour scale to indicate the ordering used for visual comparison.

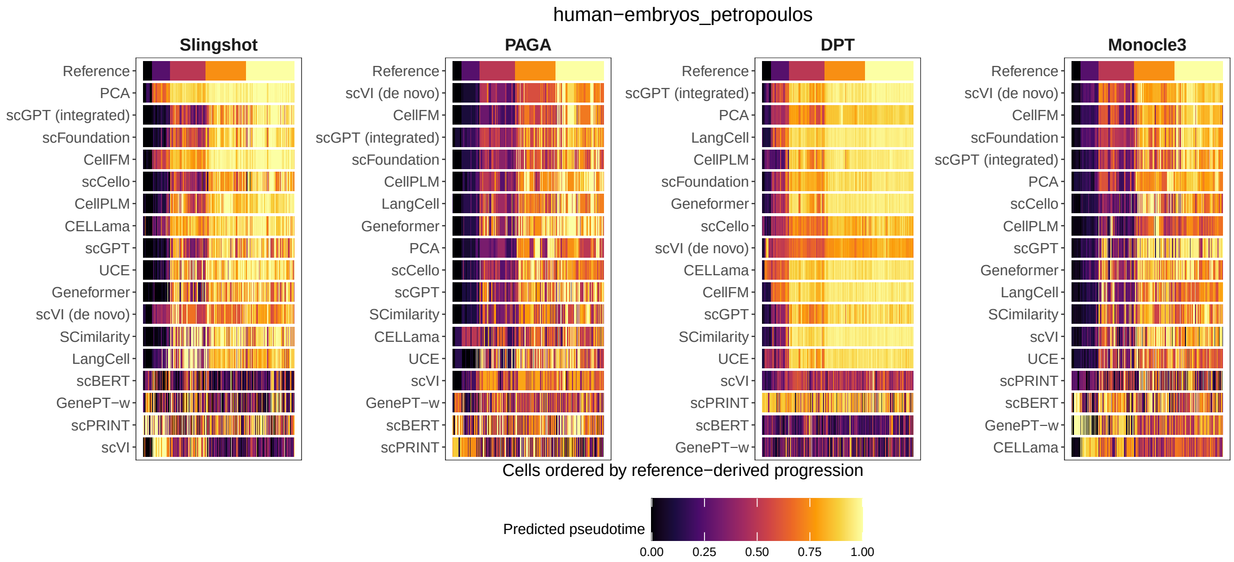

**Figure S152: Pseudotime ribbon plot for human preimplantation embryogenesis.** Columns show four trajectory-inference backends and rows show the reference progression followed by the 13 scFM representations and PCA. For this display, cells were ordered by the complete reference-derived progression for this linear trajectory. Each cell is represented by one borderless tile; colour encodes predicted pseudotime after min–max normalization within each dataset–backend–representation panel. Monocle 3 and DPT use their exported cell-level pseudotime, whereas Slingshot and PAGA use pseudotime derived from the inferred Dynwrap progressions. The reference row contains 1,289 cells and is shown on the same 0–1 colour scale to indicate the ordering used for visual comparison.

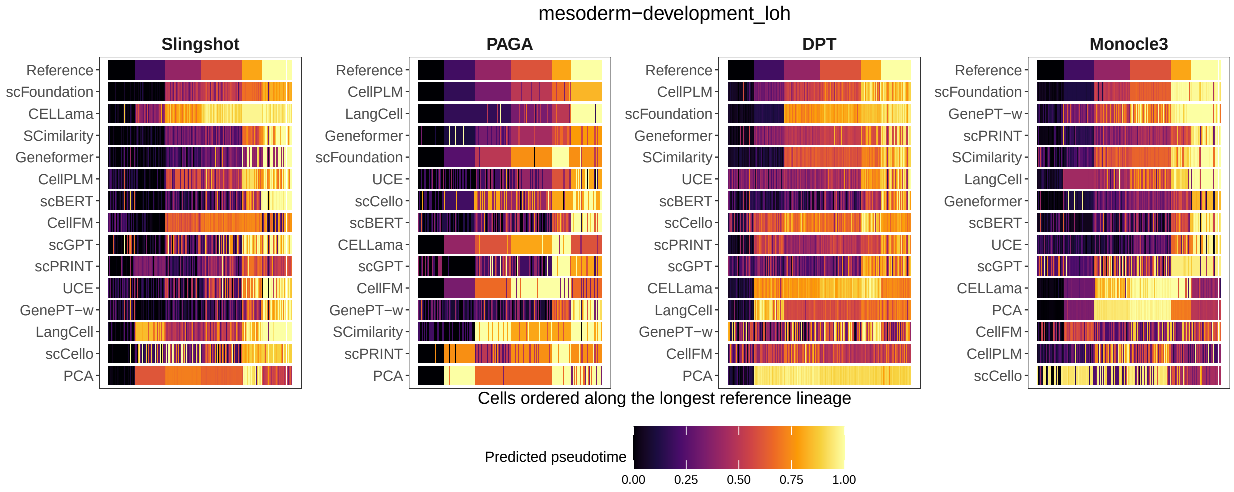

**Figure S153: Pseudotime ribbon plot for mesoderm development.** Columns show four trajectory-inference backends and rows show the reference progression followed by the 13 scFM representations and PCA. For this display, cells were restricted to the longest reference lineage, H7 hESC → anterior primitive streak → presomitic mesoderm → somitomere → early sclerotome → sclerotome, and ordered by reference progression along that path. Each cell is represented by one borderless tile; colour encodes predicted pseudotime after min–max normalization within each dataset–backend–representation panel. Monocle 3 and DPT use their exported cell-level pseudotime, whereas Slingshot and PAGA use pseudotime derived from the inferred Dynwrap progressions. The reference row contains 375 cells and is shown on the same 0–1 colour scale to indicate the ordering used for visual comparison.

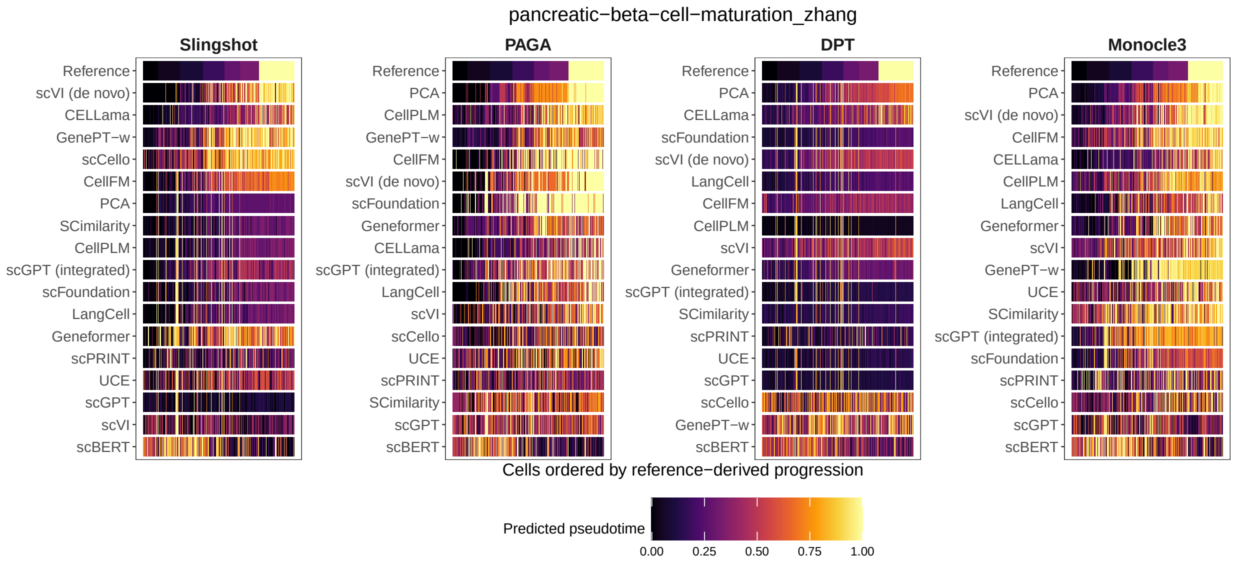

**Figure S154: Pseudotime ribbon plot for pancreatic beta-cell maturation.** Columns show four trajectory-inference backends and rows show the reference progression followed by the 13 scFM representations and PCA. For this display, cells were ordered by the complete reference-derived progression for this linear trajectory. Each cell is represented by one borderless tile; colour encodes predicted pseudotime after min–max normalization within each dataset–backend–representation panel. Monocle 3 and DPT use their exported cell-level pseudotime, whereas Slingshot and PAGA use pseudotime derived from the inferred Dynwrap progressions. The reference row contains 562 cells and is shown on the same 0–1 colour scale to indicate the ordering used for visual comparison.

##### 2.6 Cross-domain transfer with frozen representations and native annotators

Across the common set of 179 fine-label transfer directions, the primary unweighted-probe comparison showed substantial method and shift heterogeneity. Median macro-F1 was 0.749 for CellTypist, 0.729 for SingleR, 0.720 for scFoundation, 0.710 for SCimilarity, 0.708 for UCE, 0.707 for pretrained scVI, 0.707 for scCello, 0.704 for inductive PCA and 0.679 for scANVI. Pretrained scVI therefore ranked sixth of 19 methods. scANVI ranked tenth of 19 by overall macro-F1 and achieved a full-label recall of 0.728 for reference-scarce types, compared with 0.682 for pretrained scVI.

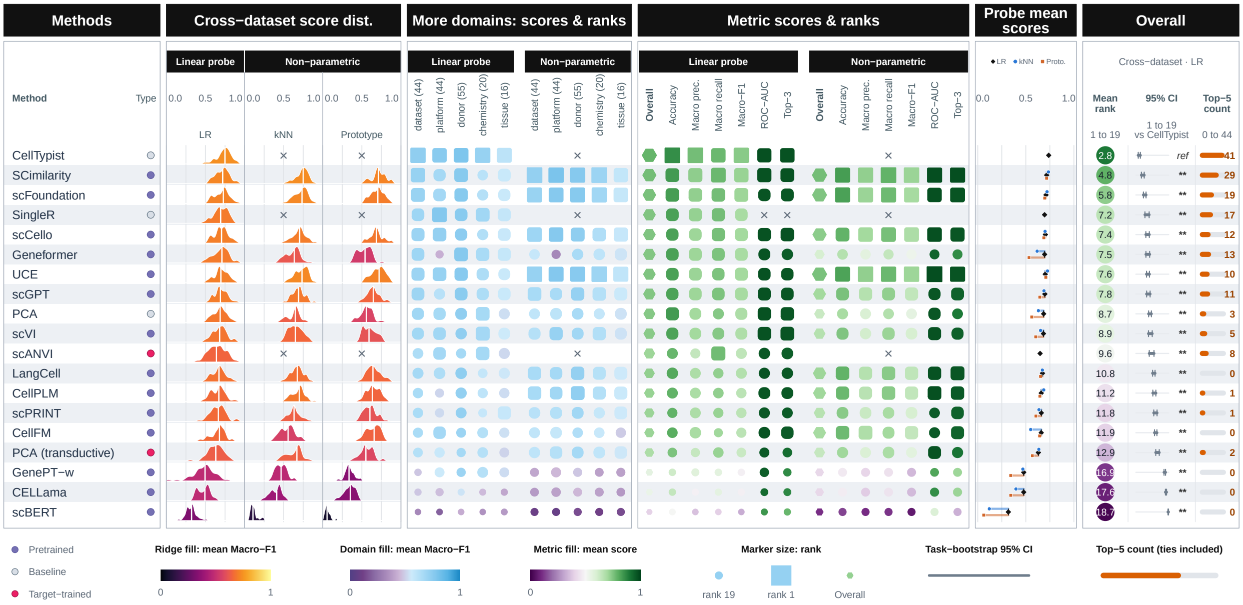

**Figure S155: Closed-set supervised transfer summary.** Rows are ordered by mean unweighted logistic-regression macro-F1 rank across the 44 cross-dataset directions. Ridges show task-level macro-F1 distributions for logistic-regression, $k$NN and prototype probes. The domain matrix extends the unweighted logistic-regression comparison to the five operational shift strata, with the eligible direction count printed for each stratum. The metric matrix reports six cross-dataset annotation metrics; Overall values average accuracy, macro precision, macro recall and macro-F1 ranks. Intervals use 2,000 cross-dataset task-bootstrap replicates, and symbols report two-sided paired Wilcoxon tests against CellTypist with Holm correction. Embedding methods use source-trained probes, CellTypist and SingleR use native annotation, and scANVI is a transductive native-prediction comparator. PCA(transductive) uses unlabelled query features and is interpreted separately from the inductive methods.

The method ordering was strongly concordant between fine and coarse label granularity (Spearman $\rho = 0.965$). Mean macro-F1 was higher at the coarse level for every method, with a mean method-level increase of 0.135. CellTypist remained the leading method (mean macro-F1 0.871), followed by UCE (0.852), SingleR (0.841), scCello (0.836) and scFoundation (0.835; Fig. S156).

Retention relative to the same-probe within-source reference also differed by shift. The median of method-level mean retention values was 0.855 for donor, 0.800 for chemistry, 0.799 for dataset, 0.751 for platform and 0.688 for tissue transfer.

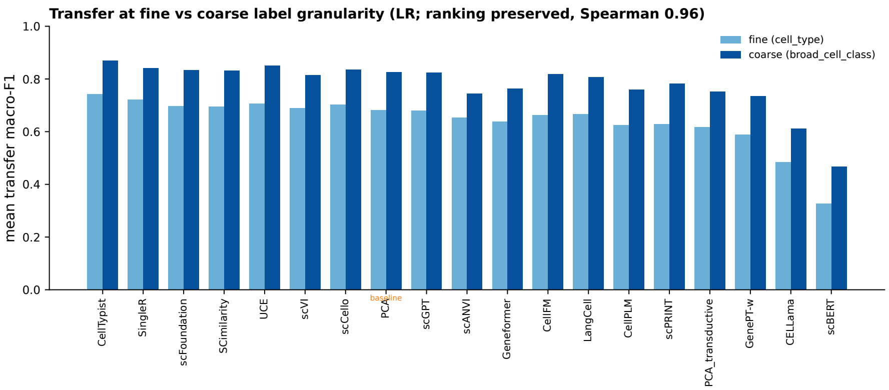

**Figure S156: Cross-domain transfer at fine and coarse label granularity.** Each point is one method. Axes show mean macro-F1 across the eligible fine-label (179 directions) and coarse-label (181 directions) analyses; the diagonal denotes equal performance at the two levels.

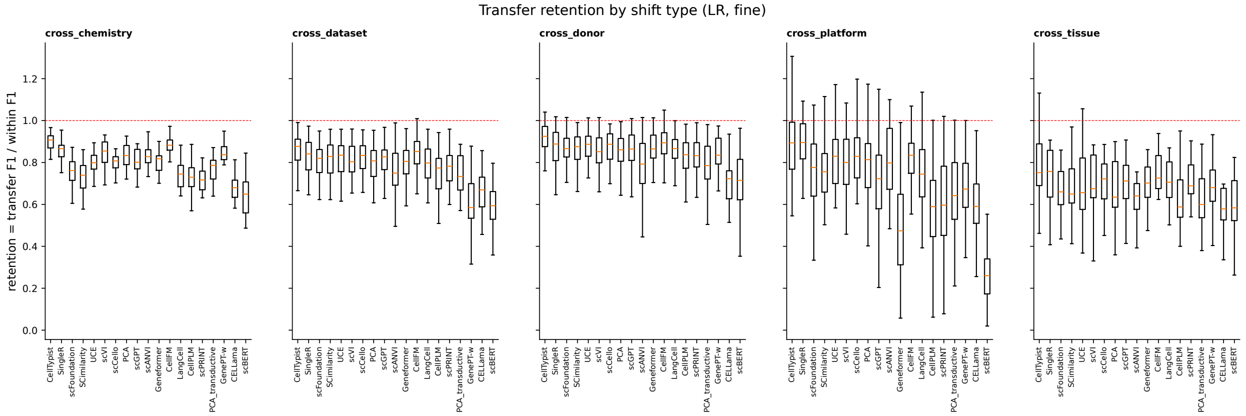

**Figure S157: Transfer retention by operational shift stratum.** Retention is target-transfer macro-F1 divided by five-fold within-source macro-F1 for the same representation and primary unweighted logistic-regression probe. Panels separate chemistry, dataset, donor, platform and tissue shifts. Boxes show the task-level distribution for each method; centre lines denote medians, boxes the interquartile range and whiskers 1.5 times that range. The dashed line at one denotes equal target and within-source macro-F1. Ratios can exceed one.

The aggregate closed-set summary and retention analysis are shown in Figs. S155 and S157. The complete 44-direction, tissue-paired closed-set gallery is retained in Figs. S160–S182.

In the separate label-efficiency analysis, scFoundation, scCello and UCE first reached 90% of their own $N = 500$ mean macro-F1 by $N = 5$ labelled source cells per class; CellTypist, SCimilarity, scGPT, SingleR, LangCell and scPRINT did so by $N = 10$; Geneformer, inductive PCA, pretrained scVI, CellPLM and CellFM by $N = 50$; CELLama by $N = 100$; and GenePT-w and scBERT by $N = 200$. No method reached this internal reference at $N = 1$. At one label per class, recall for reference-scarce types was 0.680 for UCE, 0.651 for scFoundation and 0.628 for scCello, compared with 0.607 for PCA, 0.530 for pretrained scVI and 0.484 for CellTypist.

Model-specific supervised adaptation increased mean macro-F1 for 11 of the 13 scFMs, with a median method-level increase of 0.025 across the 44 directed cross-dataset transfers (Table S7). The largest gain was observed for scBERT (+0.292). Geneformer, CellPLM, scGPT and CELLama improved by 0.039–0.047.

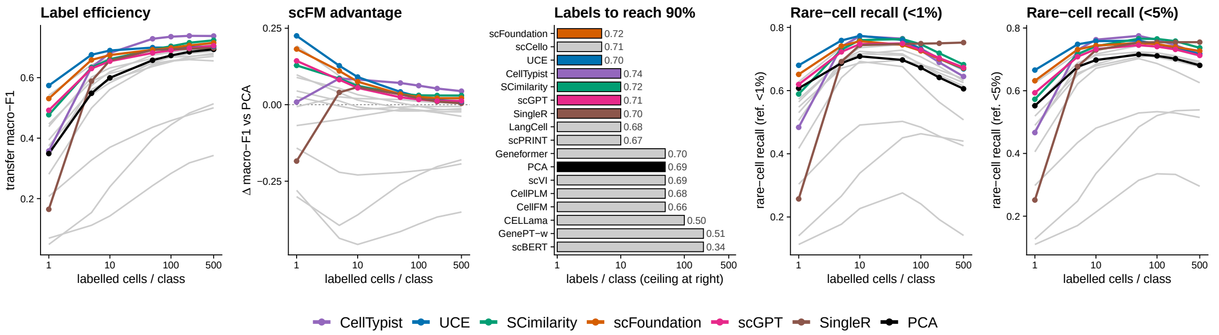

**Figure S158: Label efficiency and recovery of reference-scarce types under domain shift. a,** Transfer macro-F1. **b,** Macro-F1 difference from PCA. **c,** Labels per class required to reach 90% of each method's own macro-F1 at $N = 500$. **d,e,** Recall for types comprising less than 1% or 5% of the shared reference population. Results are averaged over 44 directed cross-dataset transfers and ten label draws.

scFoundation increased from 0.607 to 0.609, whereas CellFM and GenePT-w decreased by 0.016 and 0.147, respectively. The direction and magnitude of the effect therefore depended on the model-specific training procedure.

**Table S7:** Effect of model-specific supervised adaptation. Values are mean macro-F1 over the same 44 directed cross-dataset transfers with the complete query population retained. Difference is adaptation minus the fixed-representation logistic-regression readout.

| Method | Adaptation regime | Fixed + LR | Adaptation | Difference |
|---|---|---|---|---|
| scBERT | Partial backbone | 0.249 | 0.541 | +0.292 |
| Geneformer | Partial backbone | 0.587 | 0.635 | +0.047 |
| CellPLM | Full backbone | 0.566 | 0.612 | +0.046 |
| scGPT | Full backbone | 0.598 | 0.639 | +0.040 |
| CELLama | Sentence encoder + MLP | 0.380 | 0.419 | +0.039 |
| SCimilarity | Fixed + shared MLP | 0.609 | 0.634 | +0.026 |
| scPRINT | Fixed + shared MLP | 0.567 | 0.592 | +0.025 |
| LangCell | Full backbone | 0.567 | 0.591 | +0.025 |
| UCE | Fixed + shared MLP | 0.599 | 0.619 | +0.021 |
| scCello | Frozen + linear head | 0.593 | 0.608 | +0.014 |
| scFoundation | Partial backbone | 0.607 | 0.609 | +0.002 |
| CellFM | Partial backbone | 0.529 | 0.513 | −0.016 |
| GenePT-w | Fixed + shared MLP | 0.387 | 0.240 | −0.147 |

The matched training-cost experiment separated the methods by more than 150-fold (Table S8; Fig. S159). CellPLM completed training in a median of 65 s, followed by SCimilarity (84 s), GenePT-w (114 s) and scPRINT (129 s). scBERT and scFoundation required 9,097 and 10,112 s, respectively; scFoundation also had the largest median peak GPU memory use (142,287 MiB). Trainable scope alone did not determine resource use: CellPLM was the fastest method despite updating its full backbone.

**Table S8:** Resource use for supervised adaptation on the matched 10,000-cell reference. Values are medians over five independent launches.

| Method | Wall time (s) | Peak GPU (MiB) | Peak host RSS (MiB) |
|---|---|---|---|
| CELLama | 1,656 | 8,215 | 4,096.0 |
| CellFM | 1,397 | 7,891 | 5,120.0 |
| CellPLM | 65 | 5,297 | 3,072.0 |
| GenePT-w | 114 | 1,263 | 10,035.2 |
| Geneformer | 4,580 | 34,127 | 60,825.6 |
| LangCell | 3,825 | 78,297 | 10,137.6 |
| SCimilarity | 84 | 1,201 | 12,288.0 |
| UCE | 514 | 11,707 | 12,390.4 |
| scBERT | 9,097 | 27,295 | 3,481.6 |
| scCello | 454 | 18,991 | 31,744.0 |
| scFoundation | 10,112 | 142,287 | 9,318.4 |
| scGPT | 717 | 25,993 | 47,513.6 |
| scPRINT | 129 | 5,921 | 37,888.0 |

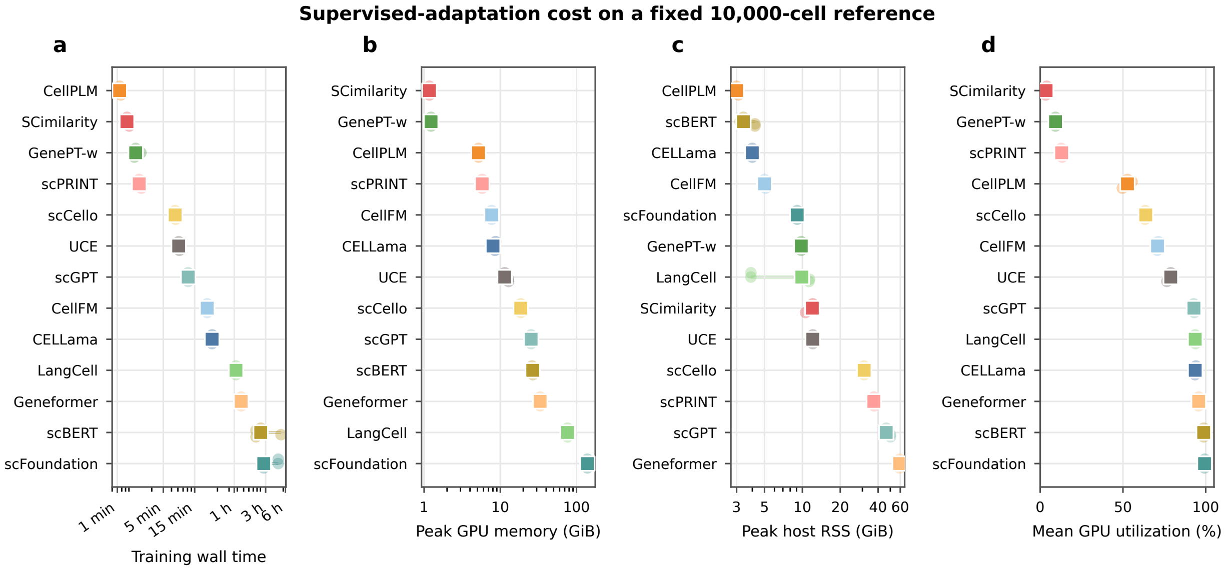

**Figure S159:** Matched supervised-adaptation resource profile. Wall time and peak host and GPU memory were recorded for five launches of each method on the same prepared 10,000-cell reference.

The observed end-to-end runtimes across the 44 real transfer directions are reported in Table S9. Median duration ranged from 40 s for CellPLM to 1 h 13 min for scFoundation. Across all directions, CellPLM and scGPT required 0.5 and 3.6 GPU-hours, respectively, whereas Geneformer and scFoundation required 49.6 and 56.1 GPU-hours.

**Table S9:** Observed end-to-end supervised-adaptation runtime across 44 real-data cross-dataset directions.

| Method | Adaptation regime | Directions | Median | Observed range | Overall |
|---|---|---|---|---|---|
| CellPLM | Native fit | 44 | 40 s | 30 s–1 m 04 s | 0.5 h |
| scGPT | Full backbone | 44 | 4 m 58 s | 2 m 33 s–8 m 03 s | 3.6 h |
| scCello | Frozen backbone + linear head | 44 | 6 m 12 s | 3 m 37 s–14 m 07 s | 5.0 h |
| CellFM | Partial encoder/head | 44 | 8 m 45 s | 4 m 26 s–14 m 42 s | 6.4 h |
| LangCell | Full sequence classifier | 44 | 23 m 54 s | 12 m 01 s–39 m 37 s | 17.9 h |
| CELLama | Sentence encoder + shared MLP | 44 | 28 m 16 s | 22 m 33 s–34 m 51 s | 20.6 h |
| scBERT | Partial backbone + head | 44 | 56 m 06 s | 19 m 33 s–1 h 29 m 08 s | 42.5 h |
| Geneformer | Partial backbone | 44 | 57 m 29 s | 32 m 30 s–2 h 44 m 38 s | 49.6 h |
| scFoundation | Transformer block [−2] + MLP head | 44 | 1 h 13 m 06 s | 18 m 41 s–2 h 41 m 02 s | 56.1 h |

Runtime is the duration of each successful process on one NVIDIA H200 and includes method-specific preprocessing, adaptation and validation, query prediction and output serialization. Fixed-embedding shared-MLP methods are reported in the controlled 10,000-cell training-cost experiment above.

**All 44 tissue x direction pairs (per-pair mean)**

| method | Annotation metrics | | | | | | Aggregated |
|---|---|---|---|---|---|---|---|
| | *Accuracy* | *Prec*$_{\text{Macro}}$ | *Recall*$_{\text{Macro}}$ | *F1*$_{\text{Macro}}$ | *AUC*$_{\text{ROC}}$ | *Acc*$_{\text{top3}}$ | *Annotation* |
| **CellTypist** | 0.86 | 0.76 | 0.78 | 0.74 | 0.98 | 0.97 | 0.78 |
| **SCimilarity** | 0.84 | 0.73 | 0.78 | 0.72 | 0.97 | 0.97 | 0.76 |
| **scFoundation** | 0.83 | 0.72 | 0.76 | 0.71 | 0.97 | 0.97 | 0.76 |
| **SingleR** | 0.83 | 0.72 | 0.76 | 0.69 | n/a | n/a | 0.75 |
| **scCello** | 0.82 | 0.71 | 0.76 | 0.70 | 0.97 | 0.97 | 0.75 |
| **UCE** | 0.82 | 0.71 | 0.75 | 0.70 | 0.97 | 0.96 | 0.75 |
| **Geneformer** | 0.82 | 0.71 | 0.75 | 0.70 | 0.97 | 0.95 | 0.75 |
| **scGPT** | 0.82 | 0.71 | 0.75 | 0.70 | 0.97 | 0.96 | 0.75 |
| **PCA** | 0.83 | 0.70 | 0.74 | 0.69 | 0.97 | 0.96 | 0.74 |
| **scVI** | 0.82 | 0.70 | 0.74 | 0.68 | 0.97 | 0.97 | 0.74 |
| **LangCell** | 0.81 | 0.69 | 0.73 | 0.67 | 0.97 | 0.96 | 0.72 |
| **CellPLM** | 0.79 | 0.69 | 0.73 | 0.66 | 0.97 | 0.96 | 0.72 |
| **CellFM** | 0.80 | 0.70 | 0.71 | 0.66 | 0.97 | 0.96 | 0.72 |
| **scPRINT** | 0.79 | 0.69 | 0.72 | 0.66 | 0.96 | 0.95 | 0.71 |
| **scANVI** | 0.78 | 0.64 | 0.77 | 0.65 | 0.96 | 0.95 | 0.71 |
| **GenePT-w** | 0.62 | 0.58 | 0.56 | 0.48 | 0.95 | 0.87 | 0.56 |
| **CELLama** | 0.64 | 0.48 | 0.56 | 0.48 | 0.93 | 0.87 | 0.54 |
| **scBERT** | 0.51 | 0.36 | 0.39 | 0.32 | 0.84 | 0.78 | 0.39 |

**Figure S160: Closed-set cross-dataset annotation across 44 directed Tabula Sapiens v1–v2 tasks.** Evaluation was restricted separately in each direction to harmonized fine labels represented by at least ten cells in both source and target. The panel comprises 860,906 evaluated target cells and a mean of approximately 16 shared classes per direction; shared labels were defined separately for each direction. Embedding methods use the primary unweighted source-trained logistic-regression probe, CellTypist and SingleR use native annotation outputs, and scANVI is a transductive native-prediction comparator. Columns report accuracy, macro precision, macro recall, macro-F1, macro one-versus-rest ROC-AUC and top-3 accuracy. SingleR does not return class probabilities, so its ROC-AUC and top-3 entries are unavailable. Values are means over the 44 directions.

#### Bladder

reference: v1, query: v2

| method | Annotation metrics: $Accuracy$ | $Prec_{Macro}$ | $Recall_{Macro}$ | $F1_{Macro}$ | $AUC_{ROC}$ | $Acc_{top3}$ | Aggregated: $Annotation$ |
|---|---|---|---|---|---|---|---|
| CellTypist | 0.95 | 0.82 | 0.85 | 0.83 | 1.00 | 0.99 | 0.86 |
| SingleR | 0.95 | 0.78 | 0.88 | 0.80 | n/a | n/a | 0.85 |
| SCimilarity | 0.95 | 0.78 | 0.84 | 0.80 | 1.00 | 0.99 | 0.84 |
| Geneformer | 0.95 | 0.77 | 0.86 | 0.79 | 1.00 | 0.99 | 0.84 |
| scFoundation | 0.95 | 0.78 | 0.85 | 0.79 | 1.00 | 0.99 | 0.84 |
| scGPT | 0.94 | 0.79 | 0.83 | 0.79 | 1.00 | 0.99 | 0.84 |
| CellPLM | 0.95 | 0.77 | 0.84 | 0.78 | 1.00 | 0.99 | 0.83 |
| UCE | 0.93 | 0.77 | 0.83 | 0.79 | 1.00 | 0.98 | 0.83 |
| scVI | 0.94 | 0.75 | 0.82 | 0.77 | 1.00 | 0.99 | 0.82 |
| CellFM | 0.93 | 0.77 | 0.80 | 0.77 | 1.00 | 0.99 | 0.82 |
| scCello | 0.94 | 0.74 | 0.83 | 0.76 | 1.00 | 0.99 | 0.82 |
| LangCell | 0.93 | 0.74 | 0.82 | 0.75 | 0.99 | 0.98 | 0.81 |
| scPRINT | 0.93 | 0.71 | 0.83 | 0.74 | 1.00 | 0.98 | 0.80 |
| PCA | 0.94 | 0.70 | 0.79 | 0.72 | 1.00 | 0.99 | 0.79 |
| GenePT-w | 0.92 | 0.69 | 0.73 | 0.69 | 0.99 | 0.99 | 0.76 |
| scANVI | 0.80 | 0.49 | 0.83 | 0.52 | 0.98 | 0.97 | 0.66 |
| CELLama | 0.75 | 0.35 | 0.58 | 0.39 | 0.94 | 0.91 | 0.52 |
| scBERT | 0.75 | 0.34 | 0.44 | 0.33 | 0.86 | 0.88 | 0.47 |

reference: v2, query: v1

| method | Annotation metrics: $Accuracy$ | $Prec_{Macro}$ | $Recall_{Macro}$ | $F1_{Macro}$ | $AUC_{ROC}$ | $Acc_{top3}$ | Aggregated: $Annotation$ |
|---|---|---|---|---|---|---|---|
| CellTypist | 0.90 | 0.84 | 0.80 | 0.79 | 0.99 | 0.99 | 0.84 |
| scANVI | 0.89 | 0.78 | 0.84 | 0.81 | 0.99 | 0.99 | 0.83 |
| scFoundation | 0.88 | 0.78 | 0.84 | 0.79 | 0.99 | 0.98 | 0.82 |
| SCimilarity | 0.88 | 0.77 | 0.84 | 0.80 | 0.99 | 0.98 | 0.82 |
| UCE | 0.89 | 0.79 | 0.81 | 0.79 | 0.99 | 0.98 | 0.82 |
| scCello | 0.88 | 0.78 | 0.81 | 0.79 | 0.99 | 0.98 | 0.82 |
| scGPT | 0.87 | 0.77 | 0.82 | 0.78 | 0.99 | 0.98 | 0.81 |
| Geneformer | 0.87 | 0.79 | 0.79 | 0.78 | 0.98 | 0.97 | 0.81 |
| SingleR | 0.84 | 0.80 | 0.79 | 0.77 | n/a | n/a | 0.80 |
| PCA | 0.89 | 0.76 | 0.78 | 0.76 | 0.99 | 0.98 | 0.80 |
| CellFM | 0.87 | 0.80 | 0.74 | 0.75 | 0.99 | 0.98 | 0.79 |
| scPRINT | 0.85 | 0.77 | 0.76 | 0.75 | 0.99 | 0.98 | 0.78 |
| scVI | 0.85 | 0.70 | 0.81 | 0.73 | 0.99 | 0.98 | 0.77 |
| LangCell | 0.83 | 0.74 | 0.78 | 0.74 | 0.99 | 0.97 | 0.77 |
| CellPLM | 0.83 | 0.69 | 0.82 | 0.73 | 0.98 | 0.97 | 0.77 |
| GenePT-w | 0.76 | 0.66 | 0.63 | 0.56 | 0.98 | 0.92 | 0.65 |
| CELLama | 0.74 | 0.51 | 0.58 | 0.52 | 0.97 | 0.91 | 0.59 |
| scBERT | 0.66 | 0.38 | 0.37 | 0.35 | 0.90 | 0.88 | 0.44 |

**Figure S161: Closed-set cross-dataset annotation for bladder. a,** TSv1 to TSv2 (36,609 target cells; 20 shared cell types). **b,** TSv2 to TSv1 (23,792 target cells; 20 shared cell types). Shared labels were defined separately in each direction and required at least ten cells in both source and target. Embedding methods use source-trained logistic regression; CellTypist and SingleR use native predictions; scANVI is fitted transductively. Columns report accuracy, macro precision, macro recall, macro-F1, macro one-versus-rest AUROC and top-3 accuracy. Probability-based metrics are unavailable for SingleR.

**Blood**

reference: v1, query: v2

| method | Annotation metrics: *Accuracy* | *Prec*$_{Macro}$ | *Recall*$_{Macro}$ | *F1*$_{Macro}$ | *AUC*$_{ROC}$ | *Acc*$_{top3}$ | Aggregated: *Annotation* |
|---|---|---|---|---|---|---|---|
| SCimilarity | 0.78 | 0.70 | 0.69 | 0.68 | 0.97 | 0.95 | 0.71 |
| Geneformer | 0.76 | 0.70 | 0.67 | 0.67 | 0.96 | 0.95 | 0.70 |
| scGPT | 0.77 | 0.70 | 0.67 | 0.67 | 0.96 | 0.96 | 0.70 |
| CellPLM | 0.79 | 0.69 | 0.66 | 0.65 | 0.97 | 0.97 | 0.70 |
| scVI | 0.74 | 0.71 | 0.68 | 0.65 | 0.96 | 0.94 | 0.69 |
| CellTypist | 0.78 | 0.68 | 0.66 | 0.65 | 0.95 | 0.94 | 0.69 |
| LangCell | 0.77 | 0.68 | 0.66 | 0.66 | 0.97 | 0.96 | 0.69 |
| scFoundation | 0.74 | 0.69 | 0.66 | 0.66 | 0.96 | 0.95 | 0.69 |
| PCA | 0.76 | 0.67 | 0.66 | 0.66 | 0.96 | 0.94 | 0.69 |
| scCello | 0.76 | 0.66 | 0.66 | 0.65 | 0.96 | 0.95 | 0.68 |
| UCE | 0.74 | 0.65 | 0.64 | 0.64 | 0.96 | 0.93 | 0.67 |
| SingleR | 0.76 | 0.63 | 0.63 | 0.61 | n/a | n/a | 0.66 |
| CellFM | 0.74 | 0.63 | 0.64 | 0.62 | 0.96 | 0.93 | 0.66 |
| GenePT-w | 0.74 | 0.63 | 0.63 | 0.59 | 0.96 | 0.93 | 0.65 |
| scANVI | 0.66 | 0.62 | 0.68 | 0.61 | 0.96 | 0.93 | 0.65 |
| scPRINT | 0.70 | 0.64 | 0.61 | 0.61 | 0.95 | 0.94 | 0.64 |
| CELLama | 0.69 | 0.54 | 0.60 | 0.54 | 0.95 | 0.90 | 0.59 |
| scBERT | 0.65 | 0.42 | 0.49 | 0.39 | 0.91 | 0.84 | 0.49 |

reference: v2, query: v1

| method | Annotation metrics: *Accuracy* | *Prec*$_{Macro}$ | *Recall*$_{Macro}$ | *F1*$_{Macro}$ | *AUC*$_{ROC}$ | *Acc*$_{top3}$ | Aggregated: *Annotation* |
|---|---|---|---|---|---|---|---|
| scANVI | 0.68 | 0.60 | 0.71 | 0.60 | 0.95 | 0.91 | 0.65 |
| SCimilarity | 0.68 | 0.60 | 0.68 | 0.60 | 0.95 | 0.90 | 0.64 |
| CellTypist | 0.65 | 0.61 | 0.68 | 0.61 | 0.95 | 0.90 | 0.64 |
| scGPT | 0.66 | 0.61 | 0.68 | 0.60 | 0.95 | 0.91 | 0.64 |
| scVI | 0.67 | 0.57 | 0.70 | 0.58 | 0.95 | 0.93 | 0.63 |
| PCA | 0.67 | 0.59 | 0.67 | 0.59 | 0.95 | 0.87 | 0.63 |
| CellPLM | 0.68 | 0.58 | 0.66 | 0.56 | 0.94 | 0.94 | 0.62 |
| scFoundation | 0.66 | 0.58 | 0.66 | 0.57 | 0.95 | 0.91 | 0.62 |
| UCE | 0.61 | 0.59 | 0.67 | 0.58 | 0.94 | 0.85 | 0.61 |
| LangCell | 0.65 | 0.57 | 0.67 | 0.57 | 0.95 | 0.89 | 0.61 |
| CellFM | 0.65 | 0.59 | 0.62 | 0.57 | 0.95 | 0.90 | 0.61 |
| Geneformer | 0.64 | 0.55 | 0.67 | 0.55 | 0.94 | 0.88 | 0.60 |
| scCello | 0.63 | 0.55 | 0.67 | 0.55 | 0.95 | 0.91 | 0.60 |
| SingleR | 0.58 | 0.56 | 0.66 | 0.53 | n/a | n/a | 0.58 |
| scPRINT | 0.61 | 0.55 | 0.60 | 0.54 | 0.93 | 0.90 | 0.58 |
| CELLama | 0.60 | 0.49 | 0.55 | 0.49 | 0.94 | 0.89 | 0.53 |
| scBERT | 0.49 | 0.40 | 0.48 | 0.36 | 0.88 | 0.81 | 0.43 |
| GenePT-w | 0.50 | 0.44 | 0.37 | 0.30 | 0.94 | 0.63 | 0.40 |

**Figure S162: Closed-set cross-dataset annotation for blood. a,** TSv1 to TSv2 (17,762 target cells; 18 shared cell types). **b,** TSv2 to TSv1 (46,411 target cells; 18 shared cell types). Label selection, methods and metrics are as in Fig. S161.

**Bone Marrow**

reference: v1, query: v2

| method | Annotation metrics: *Accuracy* | *Prec*Macro | *Recall*Macro | *F1*Macro | *AUC*ROC | *Acc*top3 | Aggregated: *Annotation* |
|---|---|---|---|---|---|---|---|
| CellTypist | 0.78 | 0.64 | 0.63 | 0.60 | 0.91 | 0.89 | 0.66 |
| scANVI | 0.74 | 0.61 | 0.64 | 0.59 | 0.94 | 0.90 | 0.65 |
| Geneformer | 0.75 | 0.59 | 0.64 | 0.58 | 0.90 | 0.89 | 0.64 |
| scGPT | 0.76 | 0.58 | 0.62 | 0.58 | 0.89 | 0.90 | 0.63 |
| PCA | 0.75 | 0.59 | 0.61 | 0.57 | 0.92 | 0.89 | 0.63 |
| UCE | 0.75 | 0.58 | 0.62 | 0.57 | 0.90 | 0.90 | 0.63 |
| SCimilarity | 0.73 | 0.58 | 0.63 | 0.56 | 0.93 | 0.89 | 0.63 |
| scFoundation | 0.75 | 0.57 | 0.62 | 0.57 | 0.90 | 0.90 | 0.63 |
| scVI | 0.73 | 0.60 | 0.59 | 0.54 | 0.95 | 0.89 | 0.61 |
| CellPLM | 0.72 | 0.57 | 0.61 | 0.55 | 0.93 | 0.90 | 0.61 |
| scCello | 0.75 | 0.55 | 0.60 | 0.55 | 0.95 | 0.89 | 0.61 |
| SingleR | 0.74 | 0.58 | 0.58 | 0.53 | n/a | n/a | 0.61 |
| LangCell | 0.74 | 0.56 | 0.58 | 0.54 | 0.91 | 0.90 | 0.61 |
| scPRINT | 0.73 | 0.55 | 0.60 | 0.54 | 0.91 | 0.87 | 0.60 |
| CellFM | 0.73 | 0.58 | 0.55 | 0.52 | 0.90 | 0.88 | 0.60 |
| GenePT-w | 0.54 | 0.44 | 0.51 | 0.39 | 0.88 | 0.79 | 0.47 |
| CELLama | 0.59 | 0.42 | 0.43 | 0.39 | 0.87 | 0.82 | 0.46 |
| scBERT | 0.52 | 0.35 | 0.38 | 0.33 | 0.85 | 0.76 | 0.40 |

reference: v2, query: v1

| method | Annotation metrics: *Accuracy* | *Prec*Macro | *Recall*Macro | *F1*Macro | *AUC*ROC | *Acc*top3 | Aggregated: *Annotation* |
|---|---|---|---|---|---|---|---|
| scVI | 0.81 | 0.62 | 0.66 | 0.62 | 0.94 | 0.96 | 0.68 |
| PCA | 0.82 | 0.61 | 0.66 | 0.61 | 0.95 | 0.96 | 0.68 |
| CellTypist | 0.83 | 0.63 | 0.63 | 0.60 | 0.96 | 0.96 | 0.67 |
| SCimilarity | 0.79 | 0.60 | 0.70 | 0.60 | 0.96 | 0.95 | 0.67 |
| scANVI | 0.78 | 0.59 | 0.67 | 0.58 | 0.93 | 0.96 | 0.66 |
| scCello | 0.77 | 0.58 | 0.66 | 0.59 | 0.94 | 0.96 | 0.65 |
| scGPT | 0.76 | 0.59 | 0.66 | 0.58 | 0.92 | 0.93 | 0.65 |
| scFoundation | 0.78 | 0.57 | 0.65 | 0.57 | 0.95 | 0.95 | 0.64 |
| UCE | 0.75 | 0.58 | 0.65 | 0.57 | 0.96 | 0.94 | 0.64 |
| LangCell | 0.76 | 0.58 | 0.63 | 0.58 | 0.96 | 0.93 | 0.64 |
| SingleR | 0.75 | 0.57 | 0.62 | 0.54 | n/a | n/a | 0.62 |
| CellPLM | 0.74 | 0.54 | 0.62 | 0.55 | 0.92 | 0.94 | 0.61 |
| scPRINT | 0.72 | 0.55 | 0.60 | 0.55 | 0.96 | 0.92 | 0.60 |
| Geneformer | 0.71 | 0.54 | 0.62 | 0.53 | 0.96 | 0.87 | 0.60 |
| CellFM | 0.74 | 0.52 | 0.55 | 0.52 | 0.94 | 0.94 | 0.58 |
| CELLama | 0.61 | 0.43 | 0.49 | 0.42 | 0.91 | 0.84 | 0.49 |
| scBERT | 0.50 | 0.34 | 0.34 | 0.30 | 0.85 | 0.78 | 0.37 |
| GenePT-w | 0.47 | 0.36 | 0.32 | 0.25 | 0.93 | 0.71 | 0.35 |

**Figure S163: Closed-set cross-dataset annotation for bone marrow. a,** TSv1 to TSv2 (8,023 target cells; 22 shared cell types). **b,** TSv2 to TSv1 (11,901 target cells; 22 shared cell types). Label selection, methods and metrics are as in Fig. S161.

#### Eye

reference: v1, query: v2

| method | Annotation metrics: $Accuracy$ | $Prec_{Macro}$ | $Recall_{Macro}$ | $F1_{Macro}$ | $AUC_{ROC}$ | $Acc_{top3}$ | Aggregated: $Annotation$ |
|---|---|---|---|---|---|---|---|
| CellTypist | 0.89 | 0.80 | 0.77 | 0.75 | 0.98 | 0.98 | 0.80 |
| SCimilarity | 0.89 | 0.74 | 0.81 | 0.75 | 0.99 | 0.99 | 0.80 |
| scFoundation | 0.88 | 0.75 | 0.78 | 0.73 | 0.99 | 0.97 | 0.78 |
| scGPT | 0.88 | 0.75 | 0.77 | 0.73 | 0.99 | 0.98 | 0.78 |
| Geneformer | 0.86 | 0.74 | 0.77 | 0.73 | 0.98 | 0.98 | 0.78 |
| SingleR | 0.87 | 0.74 | 0.77 | 0.71 | n/a | n/a | 0.77 |
| UCE | 0.86 | 0.74 | 0.77 | 0.71 | 0.98 | 0.97 | 0.77 |
| scCello | 0.86 | 0.71 | 0.78 | 0.71 | 0.99 | 0.98 | 0.77 |
| PCA | 0.88 | 0.73 | 0.74 | 0.71 | 0.99 | 0.98 | 0.77 |
| LangCell | 0.87 | 0.72 | 0.76 | 0.70 | 0.98 | 0.98 | 0.76 |
| CellPLM | 0.87 | 0.71 | 0.76 | 0.69 | 0.99 | 0.98 | 0.76 |
| scPRINT | 0.88 | 0.69 | 0.76 | 0.69 | 0.99 | 0.98 | 0.76 |
| scANVI | 0.88 | 0.66 | 0.78 | 0.67 | 0.99 | 0.98 | 0.75 |
| CellFM | 0.84 | 0.74 | 0.72 | 0.68 | 0.98 | 0.98 | 0.75 |
| scVI | 0.87 | 0.69 | 0.73 | 0.66 | 0.99 | 0.98 | 0.74 |
| GenePT-w | 0.83 | 0.65 | 0.75 | 0.64 | 0.98 | 0.97 | 0.72 |
| CELLama | 0.72 | 0.56 | 0.65 | 0.57 | 0.95 | 0.92 | 0.62 |
| scBERT | 0.37 | 0.32 | 0.34 | 0.26 | 0.82 | 0.66 | 0.32 |

reference: v2, query: v1

| method | Annotation metrics: $Accuracy$ | $Prec_{Macro}$ | $Recall_{Macro}$ | $F1_{Macro}$ | $AUC_{ROC}$ | $Acc_{top3}$ | Aggregated: $Annotation$ |
|---|---|---|---|---|---|---|---|
| CellTypist | 0.91 | 0.79 | 0.80 | 0.76 | 0.98 | 0.99 | 0.81 |
| SCimilarity | 0.89 | 0.78 | 0.80 | 0.76 | 0.99 | 0.98 | 0.80 |
| scFoundation | 0.89 | 0.77 | 0.78 | 0.75 | 0.97 | 0.98 | 0.80 |
| scCello | 0.85 | 0.78 | 0.78 | 0.76 | 0.98 | 0.98 | 0.79 |
| scGPT | 0.87 | 0.78 | 0.76 | 0.75 | 0.99 | 0.98 | 0.79 |
| UCE | 0.86 | 0.77 | 0.75 | 0.74 | 0.97 | 0.98 | 0.78 |
| PCA | 0.88 | 0.74 | 0.73 | 0.72 | 0.99 | 0.99 | 0.77 |
| Geneformer | 0.87 | 0.77 | 0.70 | 0.71 | 0.98 | 0.97 | 0.76 |
| scVI | 0.86 | 0.73 | 0.74 | 0.71 | 0.95 | 0.97 | 0.76 |
| LangCell | 0.84 | 0.74 | 0.73 | 0.71 | 0.98 | 0.97 | 0.76 |
| CellPLM | 0.85 | 0.72 | 0.75 | 0.69 | 0.96 | 0.97 | 0.75 |
| SingleR | 0.88 | 0.70 | 0.75 | 0.69 | n/a | n/a | 0.75 |
| CellFM | 0.85 | 0.74 | 0.70 | 0.70 | 0.98 | 0.98 | 0.75 |
| scPRINT | 0.83 | 0.74 | 0.70 | 0.70 | 0.97 | 0.97 | 0.74 |
| scANVI | 0.85 | 0.65 | 0.77 | 0.68 | 0.96 | 0.97 | 0.74 |
| CELLama | 0.70 | 0.54 | 0.59 | 0.53 | 0.94 | 0.91 | 0.59 |
| GenePT-w | 0.45 | 0.67 | 0.51 | 0.45 | 0.93 | 0.69 | 0.52 |
| scBERT | 0.51 | 0.28 | 0.32 | 0.27 | 0.88 | 0.84 | 0.35 |

**Figure S164: Closed-set cross-dataset annotation for eye. a,** TSv1 to TSv2 (21,117 target cells; 31 shared cell types). **b,** TSv2 to TSv1 (9,190 target cells; 31 shared cell types). Label selection, methods and metrics are as in Fig. S161.

**Fat**

reference: v1, query: v2

| method | Annotation metrics: $Accuracy$ | $Prec_{Macro}$ | $Recall_{Macro}$ | $F1_{Macro}$ | $AUC_{ROC}$ | $Acc_{top3}$ | Aggregated: $Annotation$ |
|---|---|---|---|---|---|---|---|
| scGPT | 0.82 | 0.60 | 0.69 | 0.61 | 0.96 | 0.98 | 0.68 |
| scVI | 0.84 | 0.60 | 0.68 | 0.59 | 0.98 | 0.98 | 0.68 |
| scCello | 0.83 | 0.59 | 0.70 | 0.58 | 0.96 | 0.98 | 0.67 |
| scANVI | 0.78 | 0.58 | 0.67 | 0.57 | 0.94 | 0.97 | 0.65 |
| UCE | 0.75 | 0.58 | 0.66 | 0.57 | 0.95 | 0.97 | 0.64 |
| SCimilarity | 0.72 | 0.58 | 0.67 | 0.55 | 0.95 | 0.95 | 0.63 |
| LangCell | 0.76 | 0.54 | 0.67 | 0.53 | 0.95 | 0.97 | 0.62 |
| PCA | 0.67 | 0.56 | 0.63 | 0.53 | 0.95 | 0.91 | 0.60 |
| scFoundation | 0.64 | 0.58 | 0.62 | 0.54 | 0.95 | 0.97 | 0.60 |
| CellTypist | 0.68 | 0.55 | 0.64 | 0.48 | 0.97 | 0.91 | 0.59 |
| scPRINT | 0.62 | 0.59 | 0.63 | 0.50 | 0.96 | 0.88 | 0.58 |
| CellPLM | 0.64 | 0.59 | 0.61 | 0.50 | 0.94 | 0.95 | 0.58 |
| Geneformer | 0.62 | 0.54 | 0.65 | 0.49 | 0.94 | 0.90 | 0.57 |
| SingleR | 0.60 | 0.55 | 0.66 | 0.48 | n/a | n/a | 0.57 |
| CellFM | 0.62 | 0.49 | 0.60 | 0.45 | 0.94 | 0.78 | 0.54 |
| GenePT-w | 0.55 | 0.50 | 0.55 | 0.42 | 0.94 | 0.76 | 0.51 |
| CELLama | 0.50 | 0.38 | 0.51 | 0.39 | 0.88 | 0.72 | 0.45 |
| scBERT | 0.38 | 0.33 | 0.35 | 0.26 | 0.78 | 0.67 | 0.33 |

reference: v2, query: v1

| method | Annotation metrics: $Accuracy$ | $Prec_{Macro}$ | $Recall_{Macro}$ | $F1_{Macro}$ | $AUC_{ROC}$ | $Acc_{top3}$ | Aggregated: $Annotation$ |
|---|---|---|---|---|---|---|---|
| SCimilarity | 0.85 | 0.64 | 0.70 | 0.59 | 0.97 | 0.99 | 0.69 |
| SingleR | 0.88 | 0.62 | 0.67 | 0.60 | n/a | n/a | 0.69 |
| CellPLM | 0.83 | 0.60 | 0.70 | 0.56 | 0.94 | 0.98 | 0.67 |
| scFoundation | 0.84 | 0.62 | 0.67 | 0.56 | 0.98 | 0.99 | 0.67 |
| scGPT | 0.82 | 0.61 | 0.70 | 0.53 | 0.98 | 0.98 | 0.66 |
| scVI | 0.76 | 0.60 | 0.67 | 0.55 | 0.96 | 0.99 | 0.65 |
| CellTypist | 0.82 | 0.58 | 0.65 | 0.52 | 0.98 | 0.99 | 0.64 |
| scCello | 0.75 | 0.60 | 0.68 | 0.54 | 0.97 | 0.99 | 0.64 |
| PCA | 0.82 | 0.55 | 0.67 | 0.52 | 0.99 | 0.99 | 0.64 |
| UCE | 0.76 | 0.60 | 0.66 | 0.51 | 0.98 | 0.98 | 0.64 |
| scPRINT | 0.74 | 0.61 | 0.64 | 0.50 | 0.97 | 0.98 | 0.62 |
| LangCell | 0.80 | 0.52 | 0.63 | 0.49 | 0.96 | 0.98 | 0.61 |
| CellFM | 0.71 | 0.59 | 0.62 | 0.50 | 0.95 | 0.98 | 0.61 |
| Geneformer | 0.78 | 0.52 | 0.62 | 0.48 | 0.96 | 0.97 | 0.60 |
| scANVI | 0.64 | 0.51 | 0.68 | 0.46 | 0.96 | 0.97 | 0.57 |
| GenePT-w | 0.50 | 0.57 | 0.56 | 0.39 | 0.97 | 0.93 | 0.51 |
| CELLama | 0.61 | 0.42 | 0.52 | 0.38 | 0.92 | 0.91 | 0.48 |
| scBERT | 0.65 | 0.32 | 0.43 | 0.29 | 0.85 | 0.88 | 0.42 |

**Figure S165: Closed-set cross-dataset annotation for fat. a,** TSv1 to TSv2 (69,595 target cells; 18 shared cell types). **b,** TSv2 to TSv1 (19,535 target cells; 18 shared cell types). Label selection, methods and metrics are as in Fig. S161.

#### Heart

reference: v1, query: v2

| method | Accuracy | $Prec_{Macro}$ | $Recall_{Macro}$ | $F1_{Macro}$ | $AUC_{ROC}$ | $Acc_{top3}$ | Annotation (Aggregated) |
|---|---|---|---|---|---|---|---|
| Geneformer | 0.97 | 0.83 | 0.78 | 0.79 | 0.97 | 0.99 | 0.84 |
| scFoundation | 0.97 | 0.83 | 0.77 | 0.78 | 0.95 | 1.00 | 0.83 |
| CellTypist | 0.97 | 0.83 | 0.74 | 0.76 | 0.98 | 0.99 | 0.83 |
| scGPT | 0.97 | 0.81 | 0.75 | 0.77 | 0.96 | 1.00 | 0.82 |
| scPRINT | 0.95 | 0.78 | 0.76 | 0.74 | 0.95 | 0.99 | 0.81 |
| scVI | 0.95 | 0.79 | 0.73 | 0.73 | 0.96 | 1.00 | 0.80 |
| CellPLM | 0.94 | 0.77 | 0.74 | 0.73 | 0.99 | 0.99 | 0.80 |
| scCello | 0.95 | 0.80 | 0.73 | 0.68 | 0.96 | 0.99 | 0.79 |
| UCE | 0.94 | 0.78 | 0.73 | 0.69 | 0.93 | 0.98 | 0.79 |
| LangCell | 0.95 | 0.71 | 0.74 | 0.72 | 0.94 | 0.99 | 0.78 |
| CellFM | 0.94 | 0.81 | 0.69 | 0.68 | 0.96 | 0.99 | 0.78 |
| scANVI | 0.88 | 0.73 | 0.76 | 0.68 | 0.93 | 0.95 | 0.76 |
| SCimilarity | 0.93 | 0.73 | 0.71 | 0.66 | 0.97 | 1.00 | 0.76 |
| PCA | 0.93 | 0.68 | 0.71 | 0.69 | 0.94 | 0.99 | 0.75 |
| SingleR | 0.93 | 0.68 | 0.66 | 0.66 | n/a | n/a | 0.73 |
| GenePT-w | 0.74 | 0.60 | 0.71 | 0.52 | 0.97 | 0.95 | 0.64 |
| CELLama | 0.71 | 0.52 | 0.65 | 0.54 | 0.94 | 0.92 | 0.61 |
| scBERT | 0.58 | 0.62 | 0.47 | 0.43 | 0.85 | 0.97 | 0.53 |

reference: v2, query: v1

| method | Accuracy | $Prec_{Macro}$ | $Recall_{Macro}$ | $F1_{Macro}$ | $AUC_{ROC}$ | $Acc_{top3}$ | Annotation (Aggregated) |
|---|---|---|---|---|---|---|---|
| CellTypist | 0.73 | 0.72 | 0.78 | 0.73 | 0.92 | 0.98 | 0.74 |
| scVI | 0.73 | 0.69 | 0.79 | 0.69 | 0.93 | 0.99 | 0.73 |
| UCE | 0.59 | 0.69 | 0.76 | 0.68 | 0.91 | 0.98 | 0.68 |
| scGPT | 0.62 | 0.64 | 0.78 | 0.66 | 0.90 | 0.95 | 0.67 |
| SingleR | 0.73 | 0.62 | 0.68 | 0.62 | n/a | n/a | 0.66 |
| PCA | 0.65 | 0.62 | 0.70 | 0.59 | 0.90 | 0.86 | 0.64 |
| scCello | 0.55 | 0.63 | 0.73 | 0.61 | 0.90 | 0.98 | 0.63 |
| Geneformer | 0.70 | 0.53 | 0.71 | 0.55 | 0.88 | 0.92 | 0.62 |
| scFoundation | 0.47 | 0.65 | 0.73 | 0.59 | 0.90 | 0.94 | 0.61 |
| CELLama | 0.65 | 0.54 | 0.65 | 0.52 | 0.89 | 0.92 | 0.59 |
| LangCell | 0.52 | 0.59 | 0.69 | 0.55 | 0.90 | 0.96 | 0.59 |
| SCimilarity | 0.46 | 0.59 | 0.68 | 0.55 | 0.85 | 0.96 | 0.57 |
| CellPLM | 0.45 | 0.58 | 0.70 | 0.47 | 0.92 | 0.84 | 0.55 |
| CellFM | 0.49 | 0.40 | 0.70 | 0.42 | 0.91 | 0.83 | 0.50 |
| scANVI | 0.46 | 0.43 | 0.70 | 0.40 | 0.88 | 0.77 | 0.50 |
| scPRINT | 0.39 | 0.48 | 0.64 | 0.42 | 0.85 | 0.77 | 0.48 |
| GenePT-w | 0.37 | 0.62 | 0.36 | 0.34 | 0.83 | 0.81 | 0.42 |
| scBERT | 0.33 | 0.48 | 0.48 | 0.35 | 0.74 | 0.70 | 0.41 |

**Figure S166: Closed-set cross-dataset annotation for heart. a,** TSv1 to TSv2 (11,135 target cells; 14 shared cell types). **b,** TSv2 to TSv1 (10,581 target cells; 14 shared cell types). Label selection, methods and metrics are as in Fig. S161.

#### Large Intestine

reference: v1, query: v2

| method | Annotation metrics: $Accuracy$ | $Prec_{Macro}$ | $Recall_{Macro}$ | $F1_{Macro}$ | $AUC_{ROC}$ | $Acc_{top3}$ | Aggregated: $Annotation$ |
|---|---|---|---|---|---|---|---|
| scANVI | 0.85 | 0.85 | 0.82 | 0.80 | 0.99 | 0.98 | 0.83 |
| CellTypist | 0.84 | 0.83 | 0.80 | 0.77 | 0.98 | 0.93 | 0.81 |
| SCimilarity | 0.82 | 0.78 | 0.81 | 0.77 | 0.99 | 0.96 | 0.79 |
| scCello | 0.83 | 0.74 | 0.78 | 0.74 | 0.99 | 0.97 | 0.77 |
| scFoundation | 0.82 | 0.74 | 0.79 | 0.73 | 0.99 | 0.93 | 0.77 |
| SingleR | 0.78 | 0.77 | 0.77 | 0.68 | n/a | n/a | 0.75 |
| CellFM | 0.76 | 0.80 | 0.74 | 0.69 | 0.98 | 0.96 | 0.75 |
| PCA | 0.81 | 0.69 | 0.78 | 0.69 | 0.98 | 0.92 | 0.74 |
| scGPT | 0.79 | 0.72 | 0.74 | 0.69 | 0.97 | 0.93 | 0.73 |
| Geneformer | 0.77 | 0.73 | 0.75 | 0.69 | 0.97 | 0.92 | 0.73 |
| UCE | 0.74 | 0.72 | 0.77 | 0.67 | 0.98 | 0.91 | 0.72 |
| scVI | 0.77 | 0.70 | 0.74 | 0.67 | 0.98 | 0.96 | 0.72 |
| LangCell | 0.77 | 0.71 | 0.69 | 0.66 | 0.98 | 0.96 | 0.71 |
| CellPLM | 0.72 | 0.70 | 0.71 | 0.67 | 0.97 | 0.97 | 0.70 |
| scPRINT | 0.77 | 0.70 | 0.65 | 0.63 | 0.97 | 0.95 | 0.68 |
| GenePT-w | 0.62 | 0.55 | 0.59 | 0.48 | 0.94 | 0.77 | 0.56 |
| CELLama | 0.56 | 0.46 | 0.50 | 0.44 | 0.89 | 0.76 | 0.49 |
| scBERT | 0.28 | 0.21 | 0.23 | 0.16 | 0.70 | 0.51 | 0.22 |

reference: v2, query: v1

| method | Annotation metrics: $Accuracy$ | $Prec_{Macro}$ | $Recall_{Macro}$ | $F1_{Macro}$ | $AUC_{ROC}$ | $Acc_{top3}$ | Aggregated: $Annotation$ |
|---|---|---|---|---|---|---|---|
| CellTypist | 0.85 | 0.81 | 0.82 | 0.81 | 0.99 | 0.98 | 0.82 |
| SingleR | 0.77 | 0.80 | 0.83 | 0.76 | n/a | n/a | 0.79 |
| UCE | 0.82 | 0.73 | 0.84 | 0.77 | 0.97 | 0.95 | 0.79 |
| PCA | 0.83 | 0.76 | 0.81 | 0.76 | 0.98 | 0.98 | 0.79 |
| SCimilarity | 0.86 | 0.72 | 0.83 | 0.75 | 0.98 | 0.96 | 0.79 |
| scFoundation | 0.80 | 0.75 | 0.84 | 0.76 | 0.96 | 0.95 | 0.79 |
| scGPT | 0.80 | 0.75 | 0.83 | 0.76 | 0.98 | 0.96 | 0.79 |
| scCello | 0.80 | 0.72 | 0.83 | 0.74 | 0.97 | 0.95 | 0.77 |
| Geneformer | 0.82 | 0.69 | 0.78 | 0.70 | 0.98 | 0.92 | 0.75 |
| scANVI | 0.79 | 0.65 | 0.77 | 0.68 | 0.96 | 0.93 | 0.72 |
| CellFM | 0.71 | 0.68 | 0.78 | 0.70 | 0.97 | 0.93 | 0.72 |
| scVI | 0.76 | 0.66 | 0.78 | 0.67 | 0.96 | 0.96 | 0.72 |
| LangCell | 0.70 | 0.66 | 0.75 | 0.64 | 0.96 | 0.90 | 0.69 |
| scPRINT | 0.73 | 0.65 | 0.69 | 0.63 | 0.91 | 0.95 | 0.67 |
| CellPLM | 0.65 | 0.63 | 0.73 | 0.61 | 0.96 | 0.93 | 0.66 |
| GenePT-w | 0.43 | 0.61 | 0.44 | 0.40 | 0.88 | 0.76 | 0.47 |
| CELLama | 0.38 | 0.40 | 0.48 | 0.40 | 0.87 | 0.65 | 0.42 |
| scBERT | 0.25 | 0.29 | 0.27 | 0.22 | 0.75 | 0.51 | 0.26 |

**Figure S167: Closed-set cross-dataset annotation for large intestine. a,** TSv1 to TSv2 (14,104 target cells; 24 shared cell types). **b,** TSv2 to TSv1 (7,651 target cells; 24 shared cell types). Label selection, methods and metrics are as in Fig. S161.

#### Liver

reference: v1, query: v2

| method | Annotation metrics | | | | | | Aggregated |
|---|---|---|---|---|---|---|---|
| | $Accuracy$ | $Prec_{Macro}$ | $Recall_{Macro}$ | $F1_{Macro}$ | $AUC_{ROC}$ | $Acc_{top3}$ | $Annotation$ |
| SCimilarity | 0.82 | 0.72 | 0.75 | 0.72 | 0.98 | 0.97 | 0.75 |
| UCE | 0.82 | 0.70 | 0.74 | 0.70 | 0.98 | 0.96 | 0.74 |
| scANVI | 0.78 | 0.71 | 0.76 | 0.71 | 0.97 | 0.96 | 0.74 |
| CellTypist | 0.81 | 0.70 | 0.72 | 0.69 | 0.97 | 0.95 | 0.73 |
| PCA | 0.79 | 0.70 | 0.70 | 0.67 | 0.98 | 0.96 | 0.72 |
| scCello | 0.77 | 0.69 | 0.72 | 0.67 | 0.98 | 0.95 | 0.72 |
| scFoundation | 0.78 | 0.68 | 0.68 | 0.66 | 0.96 | 0.94 | 0.70 |
| scVI | 0.79 | 0.66 | 0.68 | 0.64 | 0.98 | 0.96 | 0.69 |
| SingleR | 0.78 | 0.65 | 0.70 | 0.63 | n/a | n/a | 0.69 |
| LangCell | 0.78 | 0.69 | 0.64 | 0.62 | 0.97 | 0.95 | 0.68 |
| Geneformer | 0.77 | 0.67 | 0.66 | 0.61 | 0.97 | 0.93 | 0.68 |
| CellFM | 0.76 | 0.66 | 0.65 | 0.62 | 0.97 | 0.95 | 0.67 |
| scGPT | 0.76 | 0.65 | 0.65 | 0.63 | 0.97 | 0.94 | 0.67 |
| scPRINT | 0.79 | 0.62 | 0.65 | 0.62 | 0.97 | 0.93 | 0.67 |
| CellPLM | 0.75 | 0.64 | 0.63 | 0.60 | 0.97 | 0.92 | 0.65 |
| GenePT-w | 0.64 | 0.52 | 0.50 | 0.43 | 0.95 | 0.89 | 0.52 |
| CELLama | 0.60 | 0.48 | 0.47 | 0.45 | 0.91 | 0.84 | 0.50 |
| scBERT | 0.44 | 0.33 | 0.28 | 0.27 | 0.80 | 0.71 | 0.33 |

reference: v2, query: v1

| method | Annotation metrics | | | | | | Aggregated |
|---|---|---|---|---|---|---|---|
| | $Accuracy$ | $Prec_{Macro}$ | $Recall_{Macro}$ | $F1_{Macro}$ | $AUC_{ROC}$ | $Acc_{top3}$ | $Annotation$ |
| SCimilarity | 0.74 | 0.76 | 0.69 | 0.69 | 0.98 | 0.97 | 0.72 |
| scCello | 0.77 | 0.71 | 0.70 | 0.69 | 0.97 | 0.97 | 0.72 |
| CellTypist | 0.75 | 0.69 | 0.73 | 0.69 | 0.97 | 0.95 | 0.72 |
| UCE | 0.74 | 0.71 | 0.69 | 0.68 | 0.97 | 0.96 | 0.70 |
| SingleR | 0.74 | 0.72 | 0.70 | 0.66 | n/a | n/a | 0.70 |
| scVI | 0.76 | 0.69 | 0.70 | 0.66 | 0.98 | 0.96 | 0.70 |
| PCA | 0.76 | 0.71 | 0.68 | 0.66 | 0.98 | 0.97 | 0.70 |
| scGPT | 0.72 | 0.70 | 0.69 | 0.68 | 0.98 | 0.97 | 0.70 |
| scFoundation | 0.76 | 0.70 | 0.66 | 0.65 | 0.97 | 0.96 | 0.69 |
| scANVI | 0.73 | 0.64 | 0.72 | 0.65 | 0.96 | 0.93 | 0.69 |
| Geneformer | 0.70 | 0.68 | 0.69 | 0.65 | 0.96 | 0.92 | 0.68 |
| CellFM | 0.70 | 0.69 | 0.66 | 0.65 | 0.96 | 0.93 | 0.68 |
| LangCell | 0.73 | 0.68 | 0.65 | 0.62 | 0.97 | 0.94 | 0.67 |
| CellPLM | 0.73 | 0.68 | 0.62 | 0.61 | 0.97 | 0.97 | 0.66 |
| scPRINT | 0.71 | 0.67 | 0.62 | 0.60 | 0.97 | 0.94 | 0.65 |
| GenePT-w | 0.63 | 0.43 | 0.54 | 0.42 | 0.95 | 0.82 | 0.51 |
| CELLama | 0.45 | 0.38 | 0.48 | 0.38 | 0.89 | 0.75 | 0.42 |
| scBERT | 0.40 | 0.32 | 0.36 | 0.27 | 0.83 | 0.77 | 0.34 |

**Figure S168: Closed-set cross-dataset annotation for liver. a,** TSv1 to TSv2 (9,793 target cells; 22 shared cell types). **b,** TSv2 to TSv1 (4,590 target cells; 22 shared cell types). Label selection, methods and metrics are as in Fig. S161.

#### Lung

reference: v1, query: v2

| method | Accuracy | $Prec_{Macro}$ | $Recall_{Macro}$ | $F1_{Macro}$ | $AUC_{ROC}$ | $Acc_{top3}$ | Aggregated Annotation |
|---|---|---|---|---|---|---|---|
| scANVI | 0.87 | 0.76 | 0.82 | 0.77 | 0.99 | 0.98 | 0.80 |
| SCimilarity | 0.87 | 0.76 | 0.79 | 0.76 | 0.99 | 0.98 | 0.80 |
| SingleR | 0.88 | 0.77 | 0.79 | 0.75 | n/a | n/a | 0.80 |
| CellTypist | 0.90 | 0.75 | 0.78 | 0.75 | 0.99 | 0.98 | 0.80 |
| scCello | 0.87 | 0.75 | 0.79 | 0.75 | 0.99 | 0.98 | 0.79 |
| Geneformer | 0.87 | 0.74 | 0.78 | 0.74 | 0.99 | 0.98 | 0.78 |
| scFoundation | 0.83 | 0.72 | 0.79 | 0.73 | 0.99 | 0.95 | 0.76 |
| scGPT | 0.83 | 0.72 | 0.77 | 0.72 | 0.99 | 0.96 | 0.76 |
| scVI | 0.83 | 0.73 | 0.75 | 0.72 | 0.99 | 0.97 | 0.76 |
| PCA | 0.84 | 0.71 | 0.77 | 0.71 | 0.99 | 0.96 | 0.76 |
| UCE | 0.83 | 0.73 | 0.75 | 0.72 | 0.99 | 0.96 | 0.76 |
| CellPLM | 0.83 | 0.71 | 0.72 | 0.69 | 0.99 | 0.97 | 0.74 |
| LangCell | 0.82 | 0.70 | 0.73 | 0.70 | 0.98 | 0.96 | 0.74 |
| CellFM | 0.84 | 0.71 | 0.71 | 0.69 | 0.99 | 0.98 | 0.74 |
| scPRINT | 0.80 | 0.69 | 0.72 | 0.68 | 0.98 | 0.95 | 0.72 |
| GenePT-w | 0.72 | 0.64 | 0.66 | 0.61 | 0.98 | 0.94 | 0.66 |
| CELLama | 0.65 | 0.51 | 0.57 | 0.50 | 0.96 | 0.85 | 0.56 |
| scBERT | 0.52 | 0.39 | 0.41 | 0.35 | 0.91 | 0.76 | 0.42 |

reference: v2, query: v1

| method | Accuracy | $Prec_{Macro}$ | $Recall_{Macro}$ | $F1_{Macro}$ | $AUC_{ROC}$ | $Acc_{top3}$ | Aggregated Annotation |
|---|---|---|---|---|---|---|---|
| CellTypist | 0.92 | 0.78 | 0.77 | 0.75 | 0.99 | 0.98 | 0.81 |
| scFoundation | 0.90 | 0.76 | 0.78 | 0.76 | 0.99 | 0.98 | 0.80 |
| SCimilarity | 0.89 | 0.74 | 0.79 | 0.75 | 0.99 | 0.97 | 0.79 |
| SingleR | 0.89 | 0.75 | 0.77 | 0.73 | n/a | n/a | 0.78 |
| Geneformer | 0.89 | 0.76 | 0.74 | 0.73 | 0.99 | 0.96 | 0.78 |
| scCello | 0.89 | 0.73 | 0.76 | 0.74 | 0.99 | 0.97 | 0.78 |
| UCE | 0.90 | 0.73 | 0.76 | 0.72 | 0.99 | 0.98 | 0.78 |
| scGPT | 0.88 | 0.74 | 0.75 | 0.73 | 0.99 | 0.97 | 0.78 |
| scVI | 0.89 | 0.74 | 0.75 | 0.72 | 0.99 | 0.97 | 0.78 |
| PCA | 0.89 | 0.72 | 0.75 | 0.71 | 0.99 | 0.97 | 0.77 |
| scANVI | 0.86 | 0.70 | 0.76 | 0.71 | 0.98 | 0.97 | 0.76 |
| CellFM | 0.87 | 0.71 | 0.71 | 0.68 | 0.99 | 0.97 | 0.74 |
| CellPLM | 0.84 | 0.70 | 0.72 | 0.68 | 0.98 | 0.95 | 0.74 |
| LangCell | 0.87 | 0.67 | 0.70 | 0.66 | 0.98 | 0.95 | 0.72 |
| scPRINT | 0.85 | 0.66 | 0.68 | 0.65 | 0.98 | 0.95 | 0.71 |
| GenePT-w | 0.80 | 0.62 | 0.60 | 0.51 | 0.99 | 0.89 | 0.63 |
| CELLama | 0.81 | 0.56 | 0.59 | 0.55 | 0.97 | 0.93 | 0.63 |
| scBERT | 0.65 | 0.35 | 0.40 | 0.34 | 0.91 | 0.88 | 0.44 |

**Figure S169: Closed-set cross-dataset annotation for lung. a,** TSv1 to TSv2 (11,716 target cells; 31 shared cell types). **b,** TSv2 to TSv1 (34,342 target cells; 31 shared cell types). Label selection, methods and metrics are as in Fig. S161.

#### Lymph Node

reference: v1, query: v2

| method | Annotation metrics: *Accuracy* | *Prec*$_{Macro}$ | *Recall*$_{Macro}$ | *F1*$_{Macro}$ | *AUC*$_{ROC}$ | *Acc*$_{top3}$ | Aggregated: *Annotation* |
|---|---|---|---|---|---|---|---|
| CellTypist | 0.78 | 0.64 | 0.76 | 0.62 | 0.98 | 0.96 | 0.70 |
| SCimilarity | 0.77 | 0.60 | 0.78 | 0.62 | 0.98 | 0.99 | 0.69 |
| SingleR | 0.75 | 0.64 | 0.73 | 0.61 | n/a | n/a | 0.68 |
| Geneformer | 0.76 | 0.59 | 0.77 | 0.60 | 0.98 | 0.97 | 0.68 |
| scPRINT | 0.83 | 0.58 | 0.70 | 0.58 | 0.98 | 0.98 | 0.67 |
| scFoundation | 0.77 | 0.57 | 0.73 | 0.58 | 0.98 | 0.98 | 0.66 |
| scGPT | 0.78 | 0.57 | 0.72 | 0.58 | 0.98 | 0.98 | 0.66 |
| scVI | 0.73 | 0.58 | 0.72 | 0.59 | 0.97 | 0.98 | 0.66 |
| PCA | 0.80 | 0.52 | 0.74 | 0.55 | 0.97 | 0.98 | 0.65 |
| scCello | 0.75 | 0.54 | 0.73 | 0.56 | 0.98 | 0.97 | 0.65 |
| CellPLM | 0.75 | 0.53 | 0.71 | 0.53 | 0.97 | 0.99 | 0.63 |
| UCE | 0.76 | 0.52 | 0.69 | 0.53 | 0.97 | 0.95 | 0.63 |
| LangCell | 0.78 | 0.51 | 0.66 | 0.54 | 0.97 | 0.98 | 0.62 |
| CellFM | 0.73 | 0.56 | 0.63 | 0.52 | 0.97 | 0.94 | 0.61 |
| GenePT-w | 0.68 | 0.50 | 0.63 | 0.47 | 0.95 | 0.93 | 0.57 |
| scANVI | 0.63 | 0.39 | 0.72 | 0.42 | 0.95 | 0.93 | 0.54 |
| CELLama | 0.54 | 0.34 | 0.51 | 0.36 | 0.88 | 0.85 | 0.44 |
| scBERT | 0.48 | 0.35 | 0.47 | 0.34 | 0.85 | 0.70 | 0.41 |

reference: v2, query: v1

| method | Annotation metrics: *Accuracy* | *Prec*$_{Macro}$ | *Recall*$_{Macro}$ | *F1*$_{Macro}$ | *AUC*$_{ROC}$ | *Acc*$_{top3}$ | Aggregated: *Annotation* |
|---|---|---|---|---|---|---|---|
| SingleR | 0.86 | 0.71 | 0.69 | 0.67 | n/a | n/a | 0.73 |
| SCimilarity | 0.81 | 0.68 | 0.68 | 0.64 | 0.98 | 0.99 | 0.70 |
| CellTypist | 0.79 | 0.69 | 0.69 | 0.63 | 0.96 | 0.97 | 0.70 |
| scFoundation | 0.80 | 0.60 | 0.70 | 0.62 | 0.97 | 0.97 | 0.68 |
| Geneformer | 0.77 | 0.63 | 0.64 | 0.60 | 0.96 | 0.95 | 0.66 |
| UCE | 0.71 | 0.62 | 0.66 | 0.59 | 0.97 | 0.97 | 0.65 |
| scGPT | 0.74 | 0.62 | 0.64 | 0.59 | 0.96 | 0.97 | 0.64 |
| scCello | 0.73 | 0.62 | 0.65 | 0.58 | 0.96 | 0.98 | 0.64 |
| scVI | 0.73 | 0.60 | 0.66 | 0.58 | 0.96 | 0.98 | 0.64 |
| CellPLM | 0.76 | 0.58 | 0.65 | 0.57 | 0.97 | 0.98 | 0.64 |
| scPRINT | 0.71 | 0.60 | 0.65 | 0.58 | 0.97 | 0.96 | 0.63 |
| LangCell | 0.74 | 0.60 | 0.63 | 0.56 | 0.96 | 0.97 | 0.63 |
| PCA | 0.75 | 0.59 | 0.63 | 0.55 | 0.97 | 0.98 | 0.63 |
| scANVI | 0.70 | 0.55 | 0.68 | 0.55 | 0.96 | 0.95 | 0.62 |
| CellFM | 0.73 | 0.61 | 0.59 | 0.54 | 0.95 | 0.94 | 0.62 |
| GenePT-w | 0.64 | 0.44 | 0.55 | 0.40 | 0.95 | 0.83 | 0.51 |
| CELLama | 0.49 | 0.43 | 0.51 | 0.41 | 0.89 | 0.78 | 0.46 |
| scBERT | 0.48 | 0.27 | 0.36 | 0.26 | 0.84 | 0.77 | 0.34 |

**Figure S170: Closed-set cross-dataset annotation for lymph node. a,** TSv1 to TSv2 (72,290 target cells; 21 shared cell types). **b,** TSv2 to TSv1 (50,884 target cells; 21 shared cell types). Label selection, methods and metrics are as in Fig. S161.

#### Mammary

reference: v1, query: v2

| method | Accuracy | $Prec_{Macro}$ | $Recall_{Macro}$ | $F1_{Macro}$ | $AUC_{ROC}$ | $Acc_{top3}$ | Aggregated Annotation |
|---|---|---|---|---|---|---|---|
| scVI | 0.98 | 0.87 | 0.87 | 0.86 | 1.00 | 1.00 | 0.89 |
| CellTypist | 0.98 | 0.88 | 0.88 | 0.82 | 1.00 | 1.00 | 0.89 |
| scCello | 0.98 | 0.85 | 0.88 | 0.85 | 0.99 | 1.00 | 0.89 |
| SCimilarity | 0.97 | 0.84 | 0.87 | 0.82 | 1.00 | 1.00 | 0.87 |
| scFoundation | 0.97 | 0.82 | 0.86 | 0.82 | 0.98 | 1.00 | 0.87 |
| UCE | 0.97 | 0.81 | 0.86 | 0.82 | 0.99 | 1.00 | 0.86 |
| SingleR | 0.97 | 0.83 | 0.85 | 0.79 | n/a | n/a | 0.86 |
| PCA | 0.96 | 0.81 | 0.85 | 0.80 | 0.99 | 1.00 | 0.85 |
| GenePT-w | 0.96 | 0.82 | 0.82 | 0.78 | 1.00 | 1.00 | 0.85 |
| LangCell | 0.95 | 0.78 | 0.85 | 0.79 | 0.98 | 1.00 | 0.84 |
| Geneformer | 0.94 | 0.77 | 0.86 | 0.79 | 0.98 | 1.00 | 0.84 |
| CellFM | 0.93 | 0.80 | 0.83 | 0.76 | 0.99 | 1.00 | 0.83 |
| scGPT | 0.88 | 0.77 | 0.84 | 0.77 | 0.99 | 1.00 | 0.82 |
| scPRINT | 0.90 | 0.74 | 0.85 | 0.77 | 0.98 | 1.00 | 0.82 |
| CellPLM | 0.81 | 0.73 | 0.81 | 0.72 | 0.99 | 1.00 | 0.77 |
| CELLama | 0.79 | 0.50 | 0.71 | 0.56 | 0.96 | 0.94 | 0.64 |
| scANVI | 0.67 | 0.54 | 0.75 | 0.56 | 0.93 | 0.95 | 0.63 |
| scBERT | 0.38 | 0.45 | 0.45 | 0.33 | 0.85 | 0.61 | 0.40 |

reference: v2, query: v1

| method | Accuracy | $Prec_{Macro}$ | $Recall_{Macro}$ | $F1_{Macro}$ | $AUC_{ROC}$ | $Acc_{top3}$ | Aggregated Annotation |
|---|---|---|---|---|---|---|---|
| scFoundation | 0.93 | 0.85 | 0.88 | 0.84 | 0.99 | 0.99 | 0.88 |
| scCello | 0.94 | 0.84 | 0.87 | 0.84 | 0.99 | 0.99 | 0.87 |
| SingleR | 0.94 | 0.85 | 0.85 | 0.84 | n/a | n/a | 0.87 |
| CellTypist | 0.94 | 0.86 | 0.86 | 0.81 | 0.99 | 0.99 | 0.87 |
| SCimilarity | 0.92 | 0.81 | 0.84 | 0.81 | 0.99 | 0.99 | 0.85 |
| LangCell | 0.92 | 0.84 | 0.81 | 0.81 | 0.98 | 0.99 | 0.85 |
| scVI | 0.92 | 0.79 | 0.82 | 0.78 | 0.99 | 0.99 | 0.83 |
| PCA | 0.93 | 0.82 | 0.78 | 0.77 | 0.98 | 0.99 | 0.82 |
| Geneformer | 0.90 | 0.79 | 0.83 | 0.78 | 0.97 | 0.98 | 0.82 |
| scANVI | 0.88 | 0.76 | 0.86 | 0.77 | 0.97 | 0.99 | 0.82 |
| CellFM | 0.90 | 0.81 | 0.78 | 0.76 | 0.98 | 0.99 | 0.81 |
| scPRINT | 0.89 | 0.79 | 0.79 | 0.78 | 0.97 | 0.98 | 0.81 |
| UCE | 0.86 | 0.78 | 0.79 | 0.76 | 0.98 | 0.99 | 0.80 |
| scGPT | 0.81 | 0.79 | 0.78 | 0.74 | 0.98 | 0.99 | 0.78 |
| CellPLM | 0.83 | 0.74 | 0.79 | 0.73 | 0.95 | 0.99 | 0.77 |
| CELLama | 0.76 | 0.55 | 0.63 | 0.55 | 0.93 | 0.93 | 0.62 |
| scBERT | 0.52 | 0.37 | 0.39 | 0.33 | 0.82 | 0.80 | 0.40 |
| GenePT-w | 0.25 | 0.66 | 0.30 | 0.29 | 0.92 | 0.90 | 0.37 |

**Figure S171: Closed-set cross-dataset annotation for mammary. a,** TSv1 to TSv2 (18,537 target cells; 14 shared cell types). **b,** TSv2 to TSv1 (10,697 target cells; 14 shared cell types). Label selection, methods and metrics are as in Fig. S161.

#### Muscle

reference: v1, query: v2

| method | Annotation metrics: $Accuracy$ | $Prec_{Macro}$ | $Recall_{Macro}$ | $F1_{Macro}$ | $AUC_{ROC}$ | $Acc_{top3}$ | Aggregated: $Annotation$ |
|---|---|---|---|---|---|---|---|
| CellTypist | 0.94 | 0.84 | 0.82 | 0.77 | 1.00 | 0.99 | 0.84 |
| SingleR | 0.93 | 0.83 | 0.78 | 0.73 | n/a | n/a | 0.82 |
| Geneformer | 0.93 | 0.73 | 0.82 | 0.74 | 0.99 | 0.98 | 0.80 |
| scANVI | 0.89 | 0.70 | 0.82 | 0.71 | 0.99 | 0.98 | 0.78 |
| scFoundation | 0.91 | 0.68 | 0.82 | 0.69 | 0.99 | 0.97 | 0.77 |
| UCE | 0.92 | 0.71 | 0.74 | 0.67 | 0.98 | 0.97 | 0.76 |
| scGPT | 0.88 | 0.72 | 0.74 | 0.68 | 0.99 | 0.97 | 0.76 |
| scPRINT | 0.87 | 0.71 | 0.76 | 0.67 | 0.99 | 0.98 | 0.76 |
| CellPLM | 0.91 | 0.70 | 0.72 | 0.67 | 0.99 | 0.97 | 0.75 |
| SCimilarity | 0.92 | 0.70 | 0.73 | 0.64 | 0.99 | 0.97 | 0.75 |
| CellFM | 0.90 | 0.76 | 0.68 | 0.64 | 0.98 | 0.98 | 0.75 |
| PCA | 0.91 | 0.69 | 0.72 | 0.62 | 0.98 | 0.98 | 0.73 |
| scCello | 0.88 | 0.67 | 0.74 | 0.63 | 0.99 | 0.98 | 0.73 |
| scVI | 0.91 | 0.68 | 0.69 | 0.60 | 0.99 | 0.98 | 0.72 |
| LangCell | 0.86 | 0.67 | 0.70 | 0.62 | 0.99 | 0.98 | 0.71 |
| GenePT-w | 0.87 | 0.60 | 0.65 | 0.55 | 0.99 | 0.97 | 0.67 |
| CELLama | 0.83 | 0.56 | 0.62 | 0.52 | 0.98 | 0.96 | 0.63 |
| scBERT | 0.30 | 0.39 | 0.33 | 0.27 | 0.85 | 0.82 | 0.32 |

reference: v2, query: v1

| method | Annotation metrics: $Accuracy$ | $Prec_{Macro}$ | $Recall_{Macro}$ | $F1_{Macro}$ | $AUC_{ROC}$ | $Acc_{top3}$ | Aggregated: $Annotation$ |
|---|---|---|---|---|---|---|---|
| CellTypist | 0.90 | 0.72 | 0.76 | 0.68 | 0.98 | 0.97 | 0.76 |
| scCello | 0.89 | 0.65 | 0.80 | 0.67 | 0.98 | 0.98 | 0.75 |
| SingleR | 0.89 | 0.67 | 0.78 | 0.65 | n/a | n/a | 0.75 |
| scFoundation | 0.88 | 0.66 | 0.75 | 0.64 | 0.97 | 0.97 | 0.74 |
| SCimilarity | 0.82 | 0.68 | 0.77 | 0.64 | 0.97 | 0.96 | 0.73 |
| UCE | 0.87 | 0.62 | 0.75 | 0.62 | 0.97 | 0.98 | 0.71 |
| Geneformer | 0.83 | 0.71 | 0.68 | 0.63 | 0.95 | 0.93 | 0.71 |
| scVI | 0.85 | 0.60 | 0.76 | 0.61 | 0.96 | 0.97 | 0.71 |
| PCA | 0.89 | 0.59 | 0.75 | 0.59 | 0.98 | 0.97 | 0.71 |
| scANVI | 0.82 | 0.58 | 0.77 | 0.58 | 0.96 | 0.97 | 0.69 |
| CellFM | 0.87 | 0.63 | 0.67 | 0.57 | 0.97 | 0.98 | 0.69 |
| scPRINT | 0.83 | 0.63 | 0.68 | 0.58 | 0.95 | 0.94 | 0.68 |
| CellPLM | 0.83 | 0.63 | 0.68 | 0.58 | 0.90 | 0.94 | 0.68 |
| scGPT | 0.83 | 0.62 | 0.68 | 0.57 | 0.94 | 0.93 | 0.67 |
| LangCell | 0.83 | 0.64 | 0.63 | 0.58 | 0.96 | 0.95 | 0.67 |
| CELLama | 0.67 | 0.36 | 0.50 | 0.35 | 0.91 | 0.88 | 0.47 |
| scBERT | 0.45 | 0.31 | 0.34 | 0.23 | 0.73 | 0.73 | 0.33 |
| GenePT-w | 0.08 | 0.43 | 0.18 | 0.14 | 0.84 | 0.46 | 0.21 |

**Figure S172: Closed-set cross-dataset annotation for muscle. a,** TSv1 to TSv2 (12,941 target cells; 25 shared cell types). **b,** TSv2 to TSv1 (29,260 target cells; 25 shared cell types). Label selection, methods and metrics are as in Fig. S161.

**Prostate**

reference: v1, query: v2

| method | Annotation metrics | | | | | | Aggregated |
|---|---|---|---|---|---|---|---|
| | *Accuracy* | $Prec_{Macro}$ | $Recall_{Macro}$ | $F1_{Macro}$ | $AUC_{ROC}$ | $Acc_{top3}$ | *Annotation* |
| scFoundation | 0.94 | 0.90 | 0.94 | 0.91 | 1.00 | 0.99 | 0.92 |
| SCimilarity | 0.92 | 0.90 | 0.95 | 0.91 | 1.00 | 0.99 | 0.92 |
| CellTypist | 0.92 | 0.89 | 0.94 | 0.90 | 1.00 | 1.00 | 0.91 |
| scGPT | 0.93 | 0.89 | 0.94 | 0.90 | 1.00 | 1.00 | 0.91 |
| scPRINT | 0.93 | 0.88 | 0.93 | 0.90 | 1.00 | 0.99 | 0.91 |
| Geneformer | 0.91 | 0.89 | 0.94 | 0.90 | 1.00 | 0.99 | 0.91 |
| CellPLM | 0.92 | 0.89 | 0.93 | 0.89 | 1.00 | 0.99 | 0.91 |
| scCello | 0.92 | 0.88 | 0.93 | 0.89 | 0.99 | 0.99 | 0.91 |
| UCE | 0.90 | 0.87 | 0.93 | 0.88 | 1.00 | 0.99 | 0.90 |
| scANVI | 0.90 | 0.87 | 0.93 | 0.88 | 1.00 | 1.00 | 0.89 |
| LangCell | 0.91 | 0.87 | 0.90 | 0.88 | 0.99 | 0.99 | 0.89 |
| scVI | 0.90 | 0.87 | 0.91 | 0.88 | 1.00 | 0.99 | 0.89 |
| CellFM | 0.89 | 0.85 | 0.87 | 0.85 | 0.99 | 1.00 | 0.86 |
| PCA | 0.90 | 0.85 | 0.86 | 0.83 | 1.00 | 0.99 | 0.86 |
| SingleR | 0.81 | 0.86 | 0.91 | 0.84 | n/a | n/a | 0.86 |
| CELLama | 0.67 | 0.64 | 0.64 | 0.62 | 0.95 | 0.91 | 0.65 |
| GenePT-w | 0.59 | 0.68 | 0.59 | 0.52 | 0.99 | 0.90 | 0.60 |
| scBERT | 0.63 | 0.47 | 0.49 | 0.46 | 0.90 | 0.88 | 0.51 |

reference: v2, query: v1

| method | Annotation metrics | | | | | | Aggregated |
|---|---|---|---|---|---|---|---|
| | *Accuracy* | $Prec_{Macro}$ | $Recall_{Macro}$ | $F1_{Macro}$ | $AUC_{ROC}$ | $Acc_{top3}$ | *Annotation* |
| UCE | 0.92 | 0.88 | 0.78 | 0.79 | 0.94 | 0.99 | 0.84 |
| scFoundation | 0.93 | 0.87 | 0.78 | 0.78 | 0.94 | 0.98 | 0.84 |
| CellTypist | 0.94 | 0.88 | 0.78 | 0.75 | 0.98 | 0.99 | 0.84 |
| Geneformer | 0.92 | 0.83 | 0.79 | 0.76 | 0.95 | 0.99 | 0.82 |
| scGPT | 0.92 | 0.80 | 0.79 | 0.77 | 0.97 | 0.98 | 0.82 |
| scCello | 0.89 | 0.83 | 0.76 | 0.77 | 0.91 | 0.98 | 0.81 |
| LangCell | 0.91 | 0.80 | 0.75 | 0.74 | 0.94 | 0.98 | 0.80 |
| SCimilarity | 0.89 | 0.77 | 0.78 | 0.74 | 0.96 | 0.98 | 0.80 |
| scVI | 0.89 | 0.80 | 0.74 | 0.74 | 0.96 | 0.98 | 0.79 |
| scPRINT | 0.87 | 0.77 | 0.79 | 0.73 | 0.95 | 0.97 | 0.79 |
| PCA | 0.87 | 0.77 | 0.77 | 0.72 | 0.95 | 0.98 | 0.79 |
| CellFM | 0.88 | 0.76 | 0.76 | 0.72 | 0.92 | 0.99 | 0.78 |
| SingleR | 0.81 | 0.66 | 0.83 | 0.68 | n/a | n/a | 0.75 |
| CellPLM | 0.55 | 0.81 | 0.66 | 0.62 | 0.96 | 0.94 | 0.66 |
| scANVI | 0.50 | 0.53 | 0.73 | 0.50 | 0.92 | 0.74 | 0.56 |
| CELLama | 0.46 | 0.37 | 0.49 | 0.38 | 0.88 | 0.81 | 0.42 |
| scBERT | 0.51 | 0.34 | 0.47 | 0.34 | 0.84 | 0.84 | 0.42 |
| GenePT-w | 0.02 | 0.18 | 0.14 | 0.03 | 0.88 | 0.69 | 0.09 |

**Figure S173: Closed-set cross-dataset annotation for prostate. a,** TSv1 to TSv2 (2,044 target cells; 14 shared cell types). **b,** TSv2 to TSv1 (15,798 target cells; 14 shared cell types). Label selection, methods and metrics are as in Fig. S161.

**Salivary Gland**

reference: v1, query: v2

| method | Annotation metrics | | | | | | Aggregated |
|---|---|---|---|---|---|---|---|
| | *Accuracy* | $Prec_{Macro}$ | $Recall_{Macro}$ | $F1_{Macro}$ | $AUC_{ROC}$ | $Acc_{top3}$ | *Annotation* |
| CellTypist | 0.80 | 0.72 | 0.83 | 0.75 | 0.97 | 0.97 | 0.77 |
| scCello | 0.74 | 0.69 | 0.79 | 0.69 | 0.98 | 0.98 | 0.73 |
| PCA | 0.76 | 0.66 | 0.79 | 0.67 | 0.98 | 0.96 | 0.72 |
| Geneformer | 0.71 | 0.68 | 0.78 | 0.67 | 0.98 | 0.97 | 0.71 |
| UCE | 0.76 | 0.64 | 0.77 | 0.66 | 0.98 | 0.97 | 0.71 |
| scGPT | 0.74 | 0.64 | 0.79 | 0.66 | 0.98 | 0.95 | 0.71 |
| scFoundation | 0.73 | 0.66 | 0.77 | 0.67 | 0.97 | 0.97 | 0.71 |
| SCimilarity | 0.71 | 0.68 | 0.78 | 0.66 | 0.99 | 0.97 | 0.71 |
| SingleR | 0.73 | 0.68 | 0.77 | 0.62 | n/a | n/a | 0.70 |
| CellFM | 0.73 | 0.65 | 0.75 | 0.66 | 0.98 | 0.98 | 0.70 |
| scANVI | 0.74 | 0.59 | 0.80 | 0.60 | 0.98 | 0.96 | 0.68 |
| scPRINT | 0.66 | 0.66 | 0.74 | 0.63 | 0.98 | 0.96 | 0.67 |
| CellPLM | 0.71 | 0.61 | 0.71 | 0.60 | 0.98 | 0.98 | 0.66 |
| scVI | 0.68 | 0.61 | 0.74 | 0.59 | 0.98 | 0.97 | 0.66 |
| LangCell | 0.67 | 0.65 | 0.69 | 0.61 | 0.97 | 0.96 | 0.65 |
| GenePT-w | 0.61 | 0.59 | 0.64 | 0.53 | 0.96 | 0.89 | 0.59 |
| CELLama | 0.62 | 0.51 | 0.61 | 0.50 | 0.94 | 0.90 | 0.56 |
| scBERT | 0.56 | 0.43 | 0.51 | 0.40 | 0.83 | 0.82 | 0.47 |

reference: v2, query: v1

| method | Annotation metrics | | | | | | Aggregated |
|---|---|---|---|---|---|---|---|
| | *Accuracy* | $Prec_{Macro}$ | $Recall_{Macro}$ | $F1_{Macro}$ | $AUC_{ROC}$ | $Acc_{top3}$ | *Annotation* |
| SCimilarity | 0.87 | 0.77 | 0.74 | 0.72 | 0.97 | 0.95 | 0.77 |
| CellTypist | 0.87 | 0.77 | 0.71 | 0.70 | 0.96 | 0.99 | 0.76 |
| scFoundation | 0.86 | 0.77 | 0.69 | 0.69 | 0.97 | 0.97 | 0.76 |
| Geneformer | 0.85 | 0.77 | 0.69 | 0.70 | 0.96 | 0.96 | 0.75 |
| PCA | 0.85 | 0.76 | 0.69 | 0.68 | 0.95 | 0.93 | 0.74 |
| LangCell | 0.84 | 0.72 | 0.69 | 0.68 | 0.96 | 0.96 | 0.73 |
| SingleR | 0.82 | 0.69 | 0.75 | 0.65 | n/a | n/a | 0.73 |
| scCello | 0.83 | 0.69 | 0.71 | 0.67 | 0.97 | 0.96 | 0.73 |
| UCE | 0.86 | 0.71 | 0.67 | 0.66 | 0.97 | 0.96 | 0.73 |
| scPRINT | 0.84 | 0.74 | 0.67 | 0.65 | 0.95 | 0.94 | 0.72 |
| scGPT | 0.82 | 0.71 | 0.70 | 0.66 | 0.95 | 0.94 | 0.72 |
| CellFM | 0.83 | 0.71 | 0.63 | 0.63 | 0.97 | 0.98 | 0.70 |
| scANVI | 0.74 | 0.64 | 0.71 | 0.62 | 0.96 | 0.96 | 0.68 |
| CellPLM | 0.74 | 0.71 | 0.58 | 0.56 | 0.96 | 0.92 | 0.65 |
| scVI | 0.67 | 0.65 | 0.56 | 0.49 | 0.93 | 0.89 | 0.59 |
| GenePT-w | 0.66 | 0.57 | 0.52 | 0.48 | 0.96 | 0.96 | 0.56 |
| CELLama | 0.56 | 0.53 | 0.46 | 0.45 | 0.91 | 0.83 | 0.50 |
| scBERT | 0.59 | 0.32 | 0.31 | 0.28 | 0.89 | 0.87 | 0.37 |

**Figure S174: Closed-set cross-dataset annotation for salivary gland. a,** TSv1 to TSv2 (11,420 target cells; 20 shared cell types). **b,** TSv2 to TSv1 (21,517 target cells; 20 shared cell types). Label selection, methods and metrics are as in Fig. S161.

#### Skin

**reference: v1, query: v2**

| method | Annotation metrics | | | | | | Aggregated |
|---|---|---|---|---|---|---|---|
| | *Accuracy* | *Prec*$_{Macro}$ | *Recall*$_{Macro}$ | *F1*$_{Macro}$ | *AUC*$_{ROC}$ | *Acc*$_{top3}$ | *Annotation* |
| **SCimilarity** | 0.90 | 0.74 | 0.78 | 0.75 | 0.98 | 0.99 | 0.79 |
| **scFoundation** | 0.90 | 0.73 | 0.79 | 0.74 | 0.97 | 0.99 | 0.79 |
| **SingleR** | 0.90 | 0.70 | 0.81 | 0.73 | n/a | n/a | 0.78 |
| **CellTypist** | 0.91 | 0.74 | 0.74 | 0.73 | 0.99 | 0.99 | 0.78 |
| **scVI** | 0.90 | 0.72 | 0.74 | 0.72 | 0.98 | 0.99 | 0.77 |
| **scCello** | 0.89 | 0.73 | 0.74 | 0.72 | 0.98 | 0.99 | 0.77 |
| **LangCell** | 0.90 | 0.71 | 0.74 | 0.72 | 0.98 | 0.99 | 0.77 |
| **scPRINT** | 0.90 | 0.71 | 0.75 | 0.71 | 0.98 | 0.99 | 0.77 |
| **Geneformer** | 0.90 | 0.72 | 0.72 | 0.71 | 0.98 | 0.99 | 0.76 |
| **CellFM** | 0.87 | 0.74 | 0.72 | 0.71 | 0.97 | 0.99 | 0.76 |
| **scANVI** | 0.87 | 0.68 | 0.76 | 0.70 | 0.97 | 0.99 | 0.75 |
| **scGPT** | 0.88 | 0.71 | 0.72 | 0.70 | 0.98 | 0.99 | 0.75 |
| **PCA** | 0.89 | 0.69 | 0.72 | 0.69 | 0.98 | 0.99 | 0.75 |
| **UCE** | 0.88 | 0.68 | 0.70 | 0.68 | 0.98 | 0.99 | 0.73 |
| **CellPLM** | 0.83 | 0.69 | 0.71 | 0.67 | 0.98 | 0.99 | 0.72 |
| **GenePT-w** | 0.79 | 0.63 | 0.69 | 0.63 | 0.96 | 0.97 | 0.68 |
| **CELLama** | 0.72 | 0.52 | 0.58 | 0.52 | 0.93 | 0.92 | 0.58 |
| **scBERT** | 0.60 | 0.34 | 0.37 | 0.33 | 0.84 | 0.83 | 0.41 |

**reference: v2, query: v1**

| method | Annotation metrics | | | | | | Aggregated |
|---|---|---|---|---|---|---|---|
| | *Accuracy* | *Prec*$_{Macro}$ | *Recall*$_{Macro}$ | *F1*$_{Macro}$ | *AUC*$_{ROC}$ | *Acc*$_{top3}$ | *Annotation* |
| **CellTypist** | 0.90 | 0.76 | 0.82 | 0.78 | 0.98 | 0.98 | 0.81 |
| **scFoundation** | 0.86 | 0.74 | 0.78 | 0.74 | 0.96 | 0.97 | 0.78 |
| **SingleR** | 0.85 | 0.70 | 0.81 | 0.72 | n/a | n/a | 0.77 |
| **CellFM** | 0.84 | 0.77 | 0.75 | 0.72 | 0.97 | 0.97 | 0.77 |
| **UCE** | 0.84 | 0.73 | 0.76 | 0.73 | 0.96 | 0.97 | 0.76 |
| **scCello** | 0.86 | 0.71 | 0.76 | 0.72 | 0.97 | 0.98 | 0.76 |
| **SCimilarity** | 0.84 | 0.70 | 0.76 | 0.71 | 0.98 | 0.99 | 0.76 |
| **scANVI** | 0.83 | 0.69 | 0.77 | 0.72 | 0.95 | 0.97 | 0.75 |
| **LangCell** | 0.85 | 0.68 | 0.76 | 0.70 | 0.97 | 0.96 | 0.75 |
| **PCA** | 0.82 | 0.71 | 0.72 | 0.71 | 0.96 | 0.99 | 0.74 |
| **CellPLM** | 0.81 | 0.70 | 0.74 | 0.68 | 0.95 | 0.96 | 0.73 |
| **scVI** | 0.84 | 0.66 | 0.74 | 0.68 | 0.97 | 0.98 | 0.73 |
| **Geneformer** | 0.80 | 0.72 | 0.68 | 0.68 | 0.97 | 0.94 | 0.72 |
| **scGPT** | 0.78 | 0.71 | 0.68 | 0.67 | 0.95 | 0.97 | 0.71 |
| **scPRINT** | 0.78 | 0.71 | 0.67 | 0.66 | 0.95 | 0.96 | 0.71 |
| **CELLama** | 0.71 | 0.50 | 0.60 | 0.53 | 0.93 | 0.90 | 0.58 |
| **GenePT-w** | 0.47 | 0.45 | 0.38 | 0.31 | 0.92 | 0.86 | 0.40 |
| **scBERT** | 0.62 | 0.34 | 0.32 | 0.30 | 0.79 | 0.85 | 0.39 |

**Figure S175: Closed-set cross-dataset annotation for skin. a,** TSv1 to TSv2 (6,466 target cells; 17 shared cell types). **b,** TSv2 to TSv1 (9,685 target cells; 17 shared cell types). Label selection, methods and metrics are as in Fig. S161.

#### Small Intestine

reference: v1, query: v2

| method | Annotation metrics | | | | | | Aggregated |
|---|---|---|---|---|---|---|---|
| | *Accuracy* | $Prec_{Macro}$ | $Recall_{Macro}$ | $F1_{Macro}$ | $AUC_{ROC}$ | $Acc_{top3}$ | *Annotation* |
| CellTypist | 0.88 | 0.84 | 0.84 | 0.81 | 0.99 | 0.99 | 0.84 |
| SingleR | 0.87 | 0.82 | 0.84 | 0.81 | n/a | n/a | 0.83 |
| SCimilarity | 0.87 | 0.81 | 0.84 | 0.80 | 0.99 | 0.98 | 0.83 |
| scANVI | 0.85 | 0.77 | 0.84 | 0.78 | 0.98 | 0.98 | 0.81 |
| PCA | 0.86 | 0.81 | 0.79 | 0.78 | 0.98 | 0.97 | 0.81 |
| CellPLM | 0.84 | 0.78 | 0.79 | 0.78 | 0.98 | 0.97 | 0.80 |
| scCello | 0.81 | 0.78 | 0.80 | 0.77 | 0.98 | 0.97 | 0.79 |
| Geneformer | 0.81 | 0.78 | 0.79 | 0.77 | 0.98 | 0.98 | 0.79 |
| UCE | 0.81 | 0.76 | 0.80 | 0.76 | 0.98 | 0.97 | 0.78 |
| scVI | 0.80 | 0.77 | 0.78 | 0.75 | 0.98 | 0.97 | 0.78 |
| scFoundation | 0.77 | 0.77 | 0.79 | 0.75 | 0.98 | 0.97 | 0.77 |
| LangCell | 0.77 | 0.74 | 0.77 | 0.73 | 0.98 | 0.94 | 0.75 |
| scGPT | 0.76 | 0.75 | 0.76 | 0.72 | 0.98 | 0.95 | 0.75 |
| scPRINT | 0.74 | 0.74 | 0.75 | 0.71 | 0.98 | 0.96 | 0.73 |
| CellFM | 0.79 | 0.75 | 0.71 | 0.67 | 0.98 | 0.96 | 0.73 |
| GenePT-w | 0.74 | 0.68 | 0.70 | 0.64 | 0.97 | 0.93 | 0.69 |
| CELLama | 0.61 | 0.57 | 0.56 | 0.54 | 0.93 | 0.83 | 0.57 |
| scBERT | 0.38 | 0.31 | 0.33 | 0.26 | 0.83 | 0.70 | 0.32 |

reference: v2, query: v1

| method | Annotation metrics | | | | | | Aggregated |
|---|---|---|---|---|---|---|---|
| | *Accuracy* | $Prec_{Macro}$ | $Recall_{Macro}$ | $F1_{Macro}$ | $AUC_{ROC}$ | $Acc_{top3}$ | *Annotation* |
| CellTypist | 0.88 | 0.85 | 0.82 | 0.81 | 0.98 | 0.99 | 0.84 |
| SCimilarity | 0.87 | 0.82 | 0.78 | 0.77 | 0.97 | 0.98 | 0.81 |
| Geneformer | 0.86 | 0.78 | 0.79 | 0.77 | 0.98 | 0.98 | 0.80 |
| SingleR | 0.83 | 0.81 | 0.79 | 0.76 | n/a | n/a | 0.80 |
| UCE | 0.82 | 0.80 | 0.79 | 0.77 | 0.98 | 0.98 | 0.80 |
| PCA | 0.85 | 0.80 | 0.76 | 0.75 | 0.98 | 0.99 | 0.79 |
| scFoundation | 0.88 | 0.76 | 0.77 | 0.74 | 0.98 | 0.98 | 0.79 |
| scGPT | 0.81 | 0.76 | 0.77 | 0.75 | 0.97 | 0.98 | 0.77 |
| scVI | 0.81 | 0.75 | 0.78 | 0.74 | 0.97 | 0.97 | 0.77 |
| scCello | 0.80 | 0.78 | 0.76 | 0.73 | 0.97 | 0.98 | 0.77 |
| CellFM | 0.74 | 0.77 | 0.75 | 0.73 | 0.97 | 0.97 | 0.75 |
| CellPLM | 0.80 | 0.76 | 0.72 | 0.71 | 0.97 | 0.97 | 0.75 |
| scPRINT | 0.78 | 0.69 | 0.76 | 0.69 | 0.98 | 0.96 | 0.73 |
| LangCell | 0.71 | 0.70 | 0.76 | 0.70 | 0.97 | 0.96 | 0.72 |
| scANVI | 0.67 | 0.64 | 0.79 | 0.66 | 0.96 | 0.93 | 0.69 |
| GenePT-w | 0.49 | 0.67 | 0.59 | 0.54 | 0.96 | 0.74 | 0.57 |
| CELLama | 0.43 | 0.50 | 0.59 | 0.50 | 0.91 | 0.78 | 0.51 |
| scBERT | 0.30 | 0.36 | 0.38 | 0.32 | 0.89 | 0.81 | 0.34 |

**Figure S176: Closed-set cross-dataset annotation for small intestine. a,** TSv1 to TSv2 (19,359 target cells; 31 shared cell types). **b,** TSv2 to TSv1 (9,104 target cells; 31 shared cell types). Label selection, methods and metrics are as in Fig. S161.

#### Spleen

reference: v1, query: v2

| method | Annotation metrics: Accuracy | $Prec_{Macro}$ | $Recall_{Macro}$ | $F1_{Macro}$ | $AUC_{ROC}$ | $Acc_{top3}$ | Aggregated: Annotation |
|---|---|---|---|---|---|---|---|
| scFoundation | 0.87 | 0.71 | 0.83 | 0.73 | 0.99 | 0.98 | 0.78 |
| Geneformer | 0.87 | 0.71 | 0.82 | 0.72 | 0.99 | 0.97 | 0.78 |
| SCimilarity | 0.86 | 0.70 | 0.83 | 0.71 | 0.99 | 0.98 | 0.77 |
| CellTypist | 0.86 | 0.71 | 0.81 | 0.72 | 0.98 | 0.97 | 0.77 |
| scANVI | 0.86 | 0.69 | 0.82 | 0.69 | 0.99 | 0.98 | 0.77 |
| SingleR | 0.86 | 0.69 | 0.81 | 0.69 | n/a | n/a | 0.76 |
| scGPT | 0.86 | 0.69 | 0.80 | 0.70 | 0.99 | 0.99 | 0.76 |
| scPRINT | 0.85 | 0.70 | 0.79 | 0.70 | 0.99 | 0.98 | 0.76 |
| CellPLM | 0.87 | 0.68 | 0.79 | 0.68 | 0.99 | 0.98 | 0.75 |
| UCE | 0.84 | 0.68 | 0.78 | 0.69 | 0.99 | 0.98 | 0.75 |
| LangCell | 0.85 | 0.67 | 0.80 | 0.68 | 0.98 | 0.97 | 0.75 |
| PCA | 0.85 | 0.66 | 0.79 | 0.68 | 0.99 | 0.99 | 0.75 |
| scVI | 0.84 | 0.67 | 0.78 | 0.67 | 0.99 | 0.98 | 0.74 |
| scCello | 0.84 | 0.65 | 0.78 | 0.66 | 0.99 | 0.98 | 0.73 |
| CellFM | 0.82 | 0.65 | 0.71 | 0.64 | 0.98 | 0.96 | 0.70 |
| GenePT-w | 0.78 | 0.58 | 0.67 | 0.55 | 0.98 | 0.94 | 0.65 |
| CELLama | 0.69 | 0.48 | 0.56 | 0.48 | 0.95 | 0.88 | 0.55 |
| scBERT | 0.64 | 0.43 | 0.47 | 0.41 | 0.93 | 0.85 | 0.49 |

reference: v2, query: v1

| method | Annotation metrics: Accuracy | $Prec_{Macro}$ | $Recall_{Macro}$ | $F1_{Macro}$ | $AUC_{ROC}$ | $Acc_{top3}$ | Aggregated: Annotation |
|---|---|---|---|---|---|---|---|
| scANVI | 0.86 | 0.70 | 0.82 | 0.73 | 0.98 | 0.98 | 0.78 |
| CellTypist | 0.85 | 0.70 | 0.78 | 0.73 | 0.98 | 0.97 | 0.77 |
| SCimilarity | 0.84 | 0.70 | 0.80 | 0.72 | 0.98 | 0.98 | 0.76 |
| UCE | 0.84 | 0.71 | 0.77 | 0.72 | 0.98 | 0.97 | 0.76 |
| scFoundation | 0.82 | 0.69 | 0.76 | 0.70 | 0.98 | 0.97 | 0.74 |
| scGPT | 0.82 | 0.69 | 0.74 | 0.69 | 0.97 | 0.97 | 0.74 |
| PCA | 0.83 | 0.67 | 0.76 | 0.68 | 0.98 | 0.97 | 0.74 |
| SingleR | 0.81 | 0.66 | 0.78 | 0.67 | n/a | n/a | 0.73 |
| LangCell | 0.80 | 0.69 | 0.74 | 0.69 | 0.97 | 0.95 | 0.73 |
| scVI | 0.81 | 0.65 | 0.77 | 0.67 | 0.98 | 0.97 | 0.73 |
| scCello | 0.81 | 0.66 | 0.76 | 0.68 | 0.98 | 0.97 | 0.73 |
| CellPLM | 0.79 | 0.65 | 0.78 | 0.67 | 0.96 | 0.95 | 0.72 |
| Geneformer | 0.79 | 0.64 | 0.75 | 0.64 | 0.97 | 0.94 | 0.71 |
| CellFM | 0.79 | 0.65 | 0.70 | 0.66 | 0.97 | 0.95 | 0.70 |
| scPRINT | 0.77 | 0.62 | 0.72 | 0.62 | 0.97 | 0.96 | 0.68 |
| GenePT-w | 0.66 | 0.52 | 0.53 | 0.46 | 0.96 | 0.88 | 0.54 |
| CELLama | 0.63 | 0.46 | 0.58 | 0.46 | 0.95 | 0.87 | 0.53 |
| scBERT | 0.53 | 0.36 | 0.42 | 0.35 | 0.89 | 0.79 | 0.41 |

**Figure S177: Closed-set cross-dataset annotation for spleen. a,** TSv1 to TSv2 (30,688 target cells; 23 shared cell types). **b,** TSv2 to TSv1 (32,711 target cells; 23 shared cell types). Label selection, methods and metrics are as in Fig. S161.

**Thymus**

reference: v1, query: v2

| method | Accuracy | $Prec_{Macro}$ | $Recall_{Macro}$ | $F1_{Macro}$ | $AUC_{ROC}$ | $Acc_{top3}$ | Aggregated Annotation |
|---|---|---|---|---|---|---|---|
| CellTypist | 0.86 | 0.85 | 0.86 | 0.84 | 0.99 | 0.97 | 0.85 |
| scANVI | 0.86 | 0.82 | 0.87 | 0.84 | 0.99 | 0.98 | 0.85 |
| scFoundation | 0.84 | 0.83 | 0.85 | 0.83 | 0.99 | 0.97 | 0.84 |
| Geneformer | 0.85 | 0.83 | 0.85 | 0.83 | 0.98 | 0.96 | 0.84 |
| SCimilarity | 0.83 | 0.82 | 0.84 | 0.82 | 0.99 | 0.96 | 0.83 |
| UCE | 0.84 | 0.82 | 0.84 | 0.82 | 0.98 | 0.96 | 0.83 |
| PCA | 0.83 | 0.78 | 0.82 | 0.78 | 0.99 | 0.97 | 0.80 |
| SingleR | 0.80 | 0.80 | 0.80 | 0.78 | n/a | n/a | 0.80 |
| scGPT | 0.81 | 0.78 | 0.80 | 0.78 | 0.99 | 0.96 | 0.79 |
| scVI | 0.80 | 0.78 | 0.79 | 0.78 | 0.98 | 0.96 | 0.79 |
| CellPLM | 0.77 | 0.78 | 0.80 | 0.75 | 0.99 | 0.95 | 0.77 |
| LangCell | 0.77 | 0.76 | 0.77 | 0.75 | 0.98 | 0.96 | 0.76 |
| scCello | 0.78 | 0.74 | 0.77 | 0.74 | 0.98 | 0.96 | 0.76 |
| CellFM | 0.73 | 0.74 | 0.76 | 0.73 | 0.98 | 0.94 | 0.74 |
| scPRINT | 0.74 | 0.74 | 0.74 | 0.72 | 0.98 | 0.95 | 0.74 |
| GenePT-w | 0.65 | 0.70 | 0.71 | 0.67 | 0.98 | 0.93 | 0.68 |
| CELLama | 0.60 | 0.52 | 0.59 | 0.53 | 0.95 | 0.86 | 0.56 |
| scBERT | 0.29 | 0.39 | 0.42 | 0.32 | 0.88 | 0.60 | 0.36 |

reference: v2, query: v1

| method | Accuracy | $Prec_{Macro}$ | $Recall_{Macro}$ | $F1_{Macro}$ | $AUC_{ROC}$ | $Acc_{top3}$ | Aggregated Annotation |
|---|---|---|---|---|---|---|---|
| CellTypist | 0.73 | 0.88 | 0.78 | 0.78 | 0.98 | 0.90 | 0.79 |
| scFoundation | 0.74 | 0.79 | 0.77 | 0.75 | 0.96 | 0.88 | 0.76 |
| SCimilarity | 0.72 | 0.79 | 0.75 | 0.74 | 0.95 | 0.91 | 0.75 |
| Geneformer | 0.73 | 0.77 | 0.76 | 0.73 | 0.95 | 0.88 | 0.75 |
| PCA | 0.71 | 0.79 | 0.75 | 0.73 | 0.96 | 0.88 | 0.74 |
| UCE | 0.71 | 0.77 | 0.76 | 0.74 | 0.96 | 0.88 | 0.74 |
| scANVI | 0.73 | 0.72 | 0.79 | 0.71 | 0.98 | 0.91 | 0.74 |
| scVI | 0.71 | 0.72 | 0.77 | 0.71 | 0.96 | 0.90 | 0.73 |
| scCello | 0.70 | 0.73 | 0.73 | 0.70 | 0.97 | 0.92 | 0.71 |
| CellPLM | 0.71 | 0.73 | 0.73 | 0.67 | 0.96 | 0.88 | 0.71 |
| SingleR | 0.60 | 0.82 | 0.70 | 0.70 | n/a | n/a | 0.71 |
| LangCell | 0.69 | 0.73 | 0.71 | 0.69 | 0.94 | 0.86 | 0.70 |
| CellFM | 0.66 | 0.73 | 0.70 | 0.68 | 0.96 | 0.87 | 0.69 |
| scPRINT | 0.67 | 0.71 | 0.71 | 0.67 | 0.95 | 0.87 | 0.69 |
| scGPT | 0.67 | 0.71 | 0.70 | 0.66 | 0.96 | 0.88 | 0.69 |
| GenePT-w | 0.56 | 0.68 | 0.59 | 0.48 | 0.96 | 0.77 | 0.58 |
| CELLama | 0.53 | 0.53 | 0.59 | 0.52 | 0.93 | 0.80 | 0.54 |
| scBERT | 0.34 | 0.32 | 0.34 | 0.26 | 0.88 | 0.62 | 0.31 |

**Figure S178: Closed-set cross-dataset annotation for thymus. a,** TSv1 to TSv2 (9,841 target cells; 26 shared cell types). **b,** TSv2 to TSv1 (28,845 target cells; 26 shared cell types). Label selection, methods and metrics are as in Fig. S161.

#### Tongue

reference: v1, query: v2

| method | Annotation metrics: Accuracy | $Prec_{Macro}$ | $Recall_{Macro}$ | $F1_{Macro}$ | $AUC_{ROC}$ | $Acc_{top3}$ | Aggregated: Annotation |
|---|---|---|---|---|---|---|---|
| CellTypist | 0.87 | 0.80 | 0.78 | 0.77 | 0.95 | 0.96 | 0.81 |
| SingleR | 0.85 | 0.75 | 0.82 | 0.76 | n/a | n/a | 0.79 |
| scFoundation | 0.84 | 0.77 | 0.79 | 0.72 | 0.98 | 0.94 | 0.78 |
| Geneformer | 0.84 | 0.70 | 0.82 | 0.73 | 0.99 | 0.96 | 0.77 |
| scVI | 0.85 | 0.72 | 0.79 | 0.73 | 0.98 | 0.96 | 0.77 |
| scPRINT | 0.84 | 0.75 | 0.77 | 0.73 | 0.95 | 0.96 | 0.77 |
| CellPLM | 0.82 | 0.72 | 0.81 | 0.73 | 0.98 | 0.96 | 0.77 |
| scCello | 0.83 | 0.72 | 0.79 | 0.72 | 0.96 | 0.96 | 0.76 |
| scGPT | 0.83 | 0.71 | 0.83 | 0.69 | 0.98 | 0.96 | 0.76 |
| CellFM | 0.81 | 0.68 | 0.75 | 0.69 | 0.99 | 0.96 | 0.73 |
| LangCell | 0.84 | 0.67 | 0.75 | 0.66 | 0.94 | 0.95 | 0.73 |
| SCimilarity | 0.82 | 0.65 | 0.79 | 0.65 | 0.94 | 0.96 | 0.73 |
| UCE | 0.82 | 0.68 | 0.74 | 0.68 | 0.97 | 0.95 | 0.73 |
| PCA | 0.81 | 0.69 | 0.73 | 0.66 | 0.99 | 0.96 | 0.72 |
| scANVI | 0.82 | 0.55 | 0.80 | 0.60 | 0.95 | 0.94 | 0.69 |
| GenePT-w | 0.72 | 0.53 | 0.68 | 0.51 | 0.95 | 0.93 | 0.61 |
| CELLama | 0.64 | 0.45 | 0.47 | 0.43 | 0.88 | 0.88 | 0.50 |
| scBERT | 0.63 | 0.32 | 0.34 | 0.28 | 0.79 | 0.84 | 0.39 |

reference: v2, query: v1

| method | Annotation metrics: Accuracy | $Prec_{Macro}$ | $Recall_{Macro}$ | $F1_{Macro}$ | $AUC_{ROC}$ | $Acc_{top3}$ | Aggregated: Annotation |
|---|---|---|---|---|---|---|---|
| CellTypist | 0.83 | 0.75 | 0.76 | 0.71 | 0.91 | 0.97 | 0.76 |
| SingleR | 0.81 | 0.77 | 0.74 | 0.71 | n/a | n/a | 0.76 |
| UCE | 0.80 | 0.69 | 0.72 | 0.68 | 0.94 | 0.95 | 0.72 |
| SCimilarity | 0.82 | 0.71 | 0.70 | 0.65 | 0.93 | 0.98 | 0.72 |
| CellFM | 0.78 | 0.68 | 0.69 | 0.68 | 0.93 | 0.97 | 0.71 |
| scCello | 0.75 | 0.67 | 0.72 | 0.67 | 0.94 | 0.96 | 0.70 |
| Geneformer | 0.78 | 0.66 | 0.70 | 0.66 | 0.94 | 0.94 | 0.70 |
| scFoundation | 0.79 | 0.69 | 0.68 | 0.64 | 0.92 | 0.96 | 0.70 |
| LangCell | 0.75 | 0.68 | 0.69 | 0.67 | 0.93 | 0.97 | 0.70 |
| scGPT | 0.77 | 0.63 | 0.72 | 0.65 | 0.94 | 0.94 | 0.69 |
| PCA | 0.80 | 0.65 | 0.68 | 0.64 | 0.91 | 0.97 | 0.69 |
| scPRINT | 0.74 | 0.71 | 0.62 | 0.59 | 0.92 | 0.95 | 0.66 |
| CellPLM | 0.69 | 0.67 | 0.62 | 0.58 | 0.90 | 0.93 | 0.64 |
| scVI | 0.70 | 0.58 | 0.56 | 0.51 | 0.92 | 0.92 | 0.59 |
| GenePT-w | 0.62 | 0.58 | 0.57 | 0.52 | 0.90 | 0.85 | 0.57 |
| scANVI | 0.63 | 0.42 | 0.68 | 0.45 | 0.90 | 0.87 | 0.55 |
| CELLama | 0.62 | 0.40 | 0.43 | 0.38 | 0.90 | 0.86 | 0.46 |
| scBERT | 0.47 | 0.34 | 0.29 | 0.23 | 0.79 | 0.70 | 0.33 |

**Figure S179: Closed-set cross-dataset annotation for tongue. a,** TSv1 to TSv2 (21,481 target cells; 21 shared cell types). **b,** TSv2 to TSv1 (11,789 target cells; 21 shared cell types). Label selection, methods and metrics are as in Fig. S161.

**Trachea**

reference: v1, query: v2

| method | Accuracy | $Prec_{Macro}$ | $Recall_{Macro}$ | $F1_{Macro}$ | $AUC_{ROC}$ | $Acc_{top3}$ | Aggregated Annotation |
|---|---|---|---|---|---|---|---|
| | Annotation metrics | | | | | | |
| CellTypist | 0.98 | 0.89 | 0.92 | 0.90 | 1.00 | 1.00 | 0.92 |
| Geneformer | 0.97 | 0.88 | 0.93 | 0.90 | 1.00 | 0.99 | 0.92 |
| SCimilarity | 0.96 | 0.87 | 0.94 | 0.90 | 1.00 | 0.99 | 0.92 |
| scPRINT | 0.97 | 0.88 | 0.92 | 0.90 | 1.00 | 1.00 | 0.92 |
| PCA | 0.96 | 0.88 | 0.89 | 0.87 | 1.00 | 1.00 | 0.90 |
| scFoundation | 0.96 | 0.85 | 0.91 | 0.87 | 1.00 | 0.99 | 0.90 |
| CellPLM | 0.96 | 0.87 | 0.88 | 0.86 | 1.00 | 0.99 | 0.89 |
| scVI | 0.93 | 0.85 | 0.91 | 0.87 | 1.00 | 1.00 | 0.89 |
| scGPT | 0.95 | 0.84 | 0.90 | 0.86 | 1.00 | 0.99 | 0.89 |
| CellFM | 0.96 | 0.87 | 0.86 | 0.83 | 1.00 | 1.00 | 0.88 |
| UCE | 0.95 | 0.80 | 0.90 | 0.83 | 1.00 | 0.99 | 0.87 |
| LangCell | 0.94 | 0.80 | 0.89 | 0.83 | 0.99 | 1.00 | 0.86 |
| scCello | 0.91 | 0.79 | 0.89 | 0.81 | 0.99 | 0.99 | 0.85 |
| SingleR | 0.93 | 0.79 | 0.87 | 0.77 | n/a | n/a | 0.84 |
| scANVI | 0.82 | 0.64 | 0.92 | 0.71 | 0.99 | 0.96 | 0.77 |
| CELLama | 0.84 | 0.57 | 0.66 | 0.60 | 0.97 | 0.96 | 0.67 |
| GenePT-w | 0.61 | 0.66 | 0.68 | 0.53 | 0.97 | 0.96 | 0.62 |
| scBERT | 0.85 | 0.51 | 0.58 | 0.51 | 0.96 | 0.96 | 0.61 |

reference: v2, query: v1

| method | Accuracy | $Prec_{Macro}$ | $Recall_{Macro}$ | $F1_{Macro}$ | $AUC_{ROC}$ | $Acc_{top3}$ | Aggregated Annotation |
|---|---|---|---|---|---|---|---|
| | Annotation metrics | | | | | | |
| CellTypist | 0.97 | 0.89 | 0.87 | 0.87 | 1.00 | 1.00 | 0.90 |
| scCello | 0.96 | 0.87 | 0.88 | 0.86 | 0.99 | 0.99 | 0.89 |
| SCimilarity | 0.96 | 0.86 | 0.88 | 0.86 | 1.00 | 0.99 | 0.89 |
| UCE | 0.96 | 0.86 | 0.86 | 0.85 | 0.99 | 0.99 | 0.88 |
| SingleR | 0.94 | 0.85 | 0.87 | 0.83 | n/a | n/a | 0.87 |
| LangCell | 0.96 | 0.84 | 0.84 | 0.83 | 1.00 | 0.99 | 0.87 |
| scFoundation | 0.95 | 0.84 | 0.84 | 0.83 | 0.99 | 0.99 | 0.87 |
| PCA | 0.96 | 0.84 | 0.84 | 0.83 | 1.00 | 0.99 | 0.86 |
| scGPT | 0.95 | 0.84 | 0.82 | 0.82 | 1.00 | 0.99 | 0.86 |
| Geneformer | 0.93 | 0.82 | 0.85 | 0.81 | 0.99 | 0.99 | 0.85 |
| scANVI | 0.90 | 0.77 | 0.87 | 0.80 | 0.97 | 0.98 | 0.83 |
| scVI | 0.94 | 0.80 | 0.80 | 0.78 | 0.99 | 0.98 | 0.83 |
| scPRINT | 0.92 | 0.78 | 0.82 | 0.78 | 0.99 | 0.99 | 0.83 |
| CellPLM | 0.93 | 0.75 | 0.78 | 0.75 | 0.95 | 0.98 | 0.80 |
| CellFM | 0.96 | 0.77 | 0.74 | 0.75 | 0.99 | 0.99 | 0.80 |
| CELLama | 0.84 | 0.63 | 0.68 | 0.64 | 0.97 | 0.95 | 0.70 |
| GenePT-w | 0.65 | 0.56 | 0.47 | 0.41 | 0.97 | 0.89 | 0.52 |
| scBERT | 0.69 | 0.39 | 0.44 | 0.38 | 0.90 | 0.90 | 0.47 |

**Figure S180: Closed-set cross-dataset annotation for trachea. a,** TSv1 to TSv2 (5,337 target cells; 22 shared cell types). **b,** TSv2 to TSv1 (9,212 target cells; 22 shared cell types). Label selection, methods and metrics are as in Fig. S161.

**Uterus**

reference: v1, query: v2

| method | Annotation metrics | | | | | | Aggregated |
|---|---|---|---|---|---|---|---|
| | *Accuracy* | *Prec*$_{Macro}$ | *Recall*$_{Macro}$ | *F1*$_{Macro}$ | *AUC*$_{ROC}$ | *Acc*$_{top3}$ | *Annotation* |
| CellTypist | 0.86 | 0.69 | 0.75 | 0.70 | 0.98 | 0.97 | 0.75 |
| SCimilarity | 0.80 | 0.71 | 0.75 | 0.69 | 0.98 | 0.99 | 0.74 |
| scFoundation | 0.83 | 0.70 | 0.74 | 0.67 | 0.97 | 0.98 | 0.73 |
| SingleR | 0.85 | 0.66 | 0.76 | 0.67 | n/a | n/a | 0.73 |
| scVI | 0.81 | 0.68 | 0.74 | 0.68 | 0.98 | 0.99 | 0.73 |
| CellPLM | 0.83 | 0.67 | 0.72 | 0.66 | 0.98 | 0.99 | 0.72 |
| UCE | 0.82 | 0.67 | 0.73 | 0.66 | 0.98 | 0.98 | 0.72 |
| LangCell | 0.82 | 0.69 | 0.72 | 0.66 | 0.98 | 0.97 | 0.72 |
| scGPT | 0.82 | 0.67 | 0.72 | 0.66 | 0.98 | 0.98 | 0.72 |
| Geneformer | 0.81 | 0.67 | 0.72 | 0.64 | 0.96 | 0.97 | 0.71 |
| scCello | 0.78 | 0.66 | 0.72 | 0.66 | 0.97 | 0.98 | 0.70 |
| PCA | 0.81 | 0.64 | 0.72 | 0.62 | 0.98 | 0.99 | 0.70 |
| CellFM | 0.81 | 0.64 | 0.66 | 0.62 | 0.97 | 0.98 | 0.69 |
| scANVI | 0.80 | 0.61 | 0.70 | 0.62 | 0.98 | 0.96 | 0.68 |
| scPRINT | 0.78 | 0.62 | 0.69 | 0.60 | 0.97 | 0.98 | 0.67 |
| GenePT-w | 0.75 | 0.60 | 0.66 | 0.58 | 0.97 | 0.96 | 0.65 |
| CELLama | 0.59 | 0.38 | 0.49 | 0.40 | 0.90 | 0.86 | 0.47 |
| scBERT | 0.46 | 0.35 | 0.36 | 0.32 | 0.82 | 0.74 | 0.37 |

reference: v2, query: v1

| method | Annotation metrics | | | | | | Aggregated |
|---|---|---|---|---|---|---|---|
| | *Accuracy* | *Prec*$_{Macro}$ | *Recall*$_{Macro}$ | *F1*$_{Macro}$ | *AUC*$_{ROC}$ | *Acc*$_{top3}$ | *Annotation* |
| CellTypist | 0.90 | 0.75 | 0.69 | 0.70 | 0.98 | 0.98 | 0.76 |
| SCimilarity | 0.84 | 0.72 | 0.71 | 0.69 | 0.96 | 0.98 | 0.74 |
| scGPT | 0.86 | 0.68 | 0.71 | 0.66 | 0.96 | 0.97 | 0.73 |
| UCE | 0.87 | 0.70 | 0.67 | 0.66 | 0.96 | 0.97 | 0.72 |
| scCello | 0.86 | 0.68 | 0.68 | 0.65 | 0.98 | 0.98 | 0.72 |
| scVI | 0.85 | 0.69 | 0.64 | 0.62 | 0.96 | 0.98 | 0.70 |
| CellFM | 0.86 | 0.64 | 0.62 | 0.58 | 0.95 | 0.96 | 0.67 |
| PCA | 0.84 | 0.62 | 0.62 | 0.59 | 0.95 | 0.96 | 0.67 |
| LangCell | 0.85 | 0.60 | 0.62 | 0.59 | 0.95 | 0.96 | 0.67 |
| SingleR | 0.80 | 0.63 | 0.66 | 0.55 | n/a | n/a | 0.66 |
| CellPLM | 0.82 | 0.59 | 0.65 | 0.58 | 0.95 | 0.97 | 0.66 |
| scFoundation | 0.82 | 0.63 | 0.60 | 0.58 | 0.93 | 0.97 | 0.66 |
| Geneformer | 0.83 | 0.57 | 0.60 | 0.56 | 0.95 | 0.94 | 0.64 |
| scPRINT | 0.78 | 0.58 | 0.56 | 0.53 | 0.87 | 0.93 | 0.61 |
| scANVI | 0.82 | 0.54 | 0.56 | 0.48 | 0.91 | 0.97 | 0.60 |
| GenePT-w | 0.64 | 0.54 | 0.48 | 0.43 | 0.90 | 0.87 | 0.52 |
| CELLama | 0.67 | 0.36 | 0.43 | 0.36 | 0.89 | 0.89 | 0.46 |
| scBERT | 0.59 | 0.32 | 0.39 | 0.32 | 0.75 | 0.84 | 0.40 |

**Figure S181: Closed-set cross-dataset annotation for uterus. a,** TSv1 to TSv2 (10,550 target cells; 17 shared cell types). **b,** TSv2 to TSv1 (5,853 target cells; 17 shared cell types). Label selection, methods and metrics are as in Fig. S161.

**Vasculature**

reference: v1, query: v2

| method | Annotation metrics | | | | | | Aggregated |
|---|---|---|---|---|---|---|---|
| | *Accuracy* | $Prec_{Macro}$ | $Recall_{Macro}$ | $F1_{Macro}$ | $AUC_{ROC}$ | $Acc_{top3}$ | *Annotation* |
| CellTypist | 0.85 | 0.83 | 0.85 | 0.83 | 0.99 | 0.99 | 0.84 |
| scGPT | 0.83 | 0.79 | 0.85 | 0.81 | 0.99 | 0.99 | 0.82 |
| SCimilarity | 0.83 | 0.78 | 0.86 | 0.80 | 0.99 | 0.98 | 0.82 |
| scCello | 0.83 | 0.79 | 0.85 | 0.80 | 0.99 | 0.97 | 0.82 |
| Geneformer | 0.83 | 0.78 | 0.85 | 0.80 | 0.99 | 0.96 | 0.82 |
| SingleR | 0.83 | 0.79 | 0.84 | 0.78 | n/a | n/a | 0.81 |
| UCE | 0.82 | 0.78 | 0.83 | 0.78 | 0.98 | 0.98 | 0.80 |
| scFoundation | 0.82 | 0.75 | 0.85 | 0.78 | 0.98 | 0.96 | 0.80 |
| scVI | 0.82 | 0.77 | 0.82 | 0.78 | 0.99 | 0.99 | 0.80 |
| scPRINT | 0.80 | 0.77 | 0.83 | 0.78 | 0.98 | 0.98 | 0.80 |
| PCA | 0.83 | 0.76 | 0.82 | 0.78 | 0.98 | 0.98 | 0.79 |
| scANVI | 0.82 | 0.74 | 0.85 | 0.77 | 0.98 | 0.96 | 0.79 |
| LangCell | 0.81 | 0.75 | 0.83 | 0.77 | 0.98 | 0.96 | 0.79 |
| CellPLM | 0.81 | 0.75 | 0.83 | 0.77 | 0.98 | 0.96 | 0.79 |
| CellFM | 0.82 | 0.77 | 0.76 | 0.74 | 0.99 | 0.99 | 0.77 |
| GenePT-w | 0.78 | 0.68 | 0.76 | 0.67 | 0.98 | 0.98 | 0.72 |
| CELLama | 0.73 | 0.56 | 0.66 | 0.57 | 0.97 | 0.93 | 0.63 |
| scBERT | 0.58 | 0.31 | 0.36 | 0.30 | 0.87 | 0.82 | 0.39 |

reference: v2, query: v1

| method | Annotation metrics | | | | | | Aggregated |
|---|---|---|---|---|---|---|---|
| | *Accuracy* | $Prec_{Macro}$ | $Recall_{Macro}$ | $F1_{Macro}$ | $AUC_{ROC}$ | $Acc_{top3}$ | *Annotation* |
| SingleR | 0.91 | 0.77 | 0.86 | 0.78 | n/a | n/a | 0.83 |
| CellTypist | 0.88 | 0.77 | 0.86 | 0.79 | 0.99 | 0.99 | 0.83 |
| SCimilarity | 0.87 | 0.73 | 0.87 | 0.77 | 0.99 | 0.99 | 0.81 |
| scCello | 0.85 | 0.75 | 0.84 | 0.77 | 0.99 | 0.98 | 0.80 |
| scFoundation | 0.81 | 0.75 | 0.83 | 0.75 | 0.99 | 0.96 | 0.78 |
| UCE | 0.83 | 0.71 | 0.82 | 0.73 | 0.99 | 0.98 | 0.77 |
| scVI | 0.83 | 0.70 | 0.84 | 0.71 | 0.99 | 0.98 | 0.77 |
| CellFM | 0.85 | 0.70 | 0.80 | 0.72 | 0.99 | 0.98 | 0.77 |
| PCA | 0.83 | 0.70 | 0.81 | 0.72 | 0.99 | 0.99 | 0.76 |
| Geneformer | 0.82 | 0.71 | 0.79 | 0.72 | 0.98 | 0.95 | 0.76 |
| scPRINT | 0.80 | 0.69 | 0.82 | 0.71 | 0.99 | 0.97 | 0.76 |
| scGPT | 0.80 | 0.71 | 0.79 | 0.71 | 0.98 | 0.97 | 0.75 |
| CellPLM | 0.80 | 0.71 | 0.78 | 0.70 | 0.98 | 0.97 | 0.75 |
| scANVI | 0.83 | 0.59 | 0.84 | 0.64 | 0.99 | 0.96 | 0.72 |
| LangCell | 0.77 | 0.67 | 0.78 | 0.67 | 0.99 | 0.97 | 0.72 |
| GenePT-w | 0.77 | 0.66 | 0.63 | 0.51 | 0.98 | 0.93 | 0.64 |
| CELLama | 0.73 | 0.48 | 0.61 | 0.47 | 0.96 | 0.92 | 0.57 |
| scBERT | 0.56 | 0.31 | 0.44 | 0.29 | 0.86 | 0.87 | 0.40 |

**Figure S182: Closed-set cross-dataset annotation for vasculature. a,** TSv1 to TSv2 (21,195 target cells; 19 shared cell types). **b,** TSv2 to TSv1 (15,555 target cells; 19 shared cell types). Label selection, methods and metrics are as in Fig. S161.

#### 2.7 Directional cross-dataset annotation and novel-type detection

The 44 directed TSv1–TSv2 tasks support two distinct views. The directional gallery below contrasts forced open-set annotation, which retains target-only labels, with closed-set evaluation over labels shared by source and target. Explicit novelty detection additionally requires both known and novel target cells; 37 of the 44 directions met that criterion.

Using a common Mahalanobis detector across embedding methods, mean per-task AUROC was 0.835 for scFoundation, 0.832 for LangCell, 0.825 for UCE, 0.824 for pretrained scVI and 0.730 for inductive PCA. Mahalanobis distance also provided strong AUPR and FPR95 across the common detector comparison.

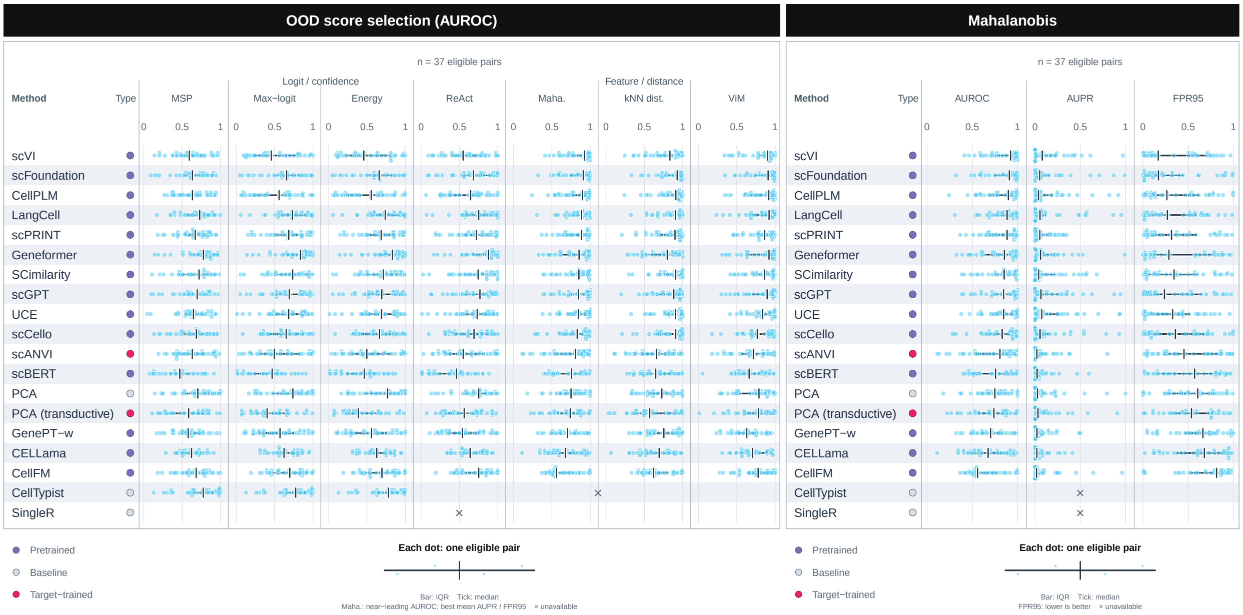

**Figure S183: Novel-type detection across eligible cross-dataset directions.** The first block compares four confidence or logit scores (maximum softmax probability, maximum logit, energy and ReAct energy) and three feature or distance scores (Mahalanobis, $k$NN distance and ViM) using per-task AUROC. The remaining blocks report the common Mahalanobis detector with AUROC, AUPR and FPR95. Each dot is one of 37 directed tasks containing both source-known and source-absent target types; seven of 44 tasks contain no novel target cell and are ineligible. AUROC and AUPR are higher-is-better; FPR95 is the known-cell false-positive rate at 95% novel-cell sensitivity and is lower-is-better. Horizontal bars span the interquartile range and vertical marks show medians. Crosses denote unavailable scores. SingleR's native reference similarity is not treated as softmax confidence and is excluded from the seven-score comparison.

Forced closed-set predictions for all 18,391 novel target cells often remained within a related biological category, although the relationship varied by method. UCE assigned 0.819 of novel cells to the same broad category, 0.936 to the same compartment and 0.411 to an annotated ancestor relation; the corresponding values were 0.816, 0.923 and 0.397 for scFoundation and 0.758, 0.906 and 0.331 for PCA. These values summarize the biological proximity of forced assignments before applying a novelty threshold.

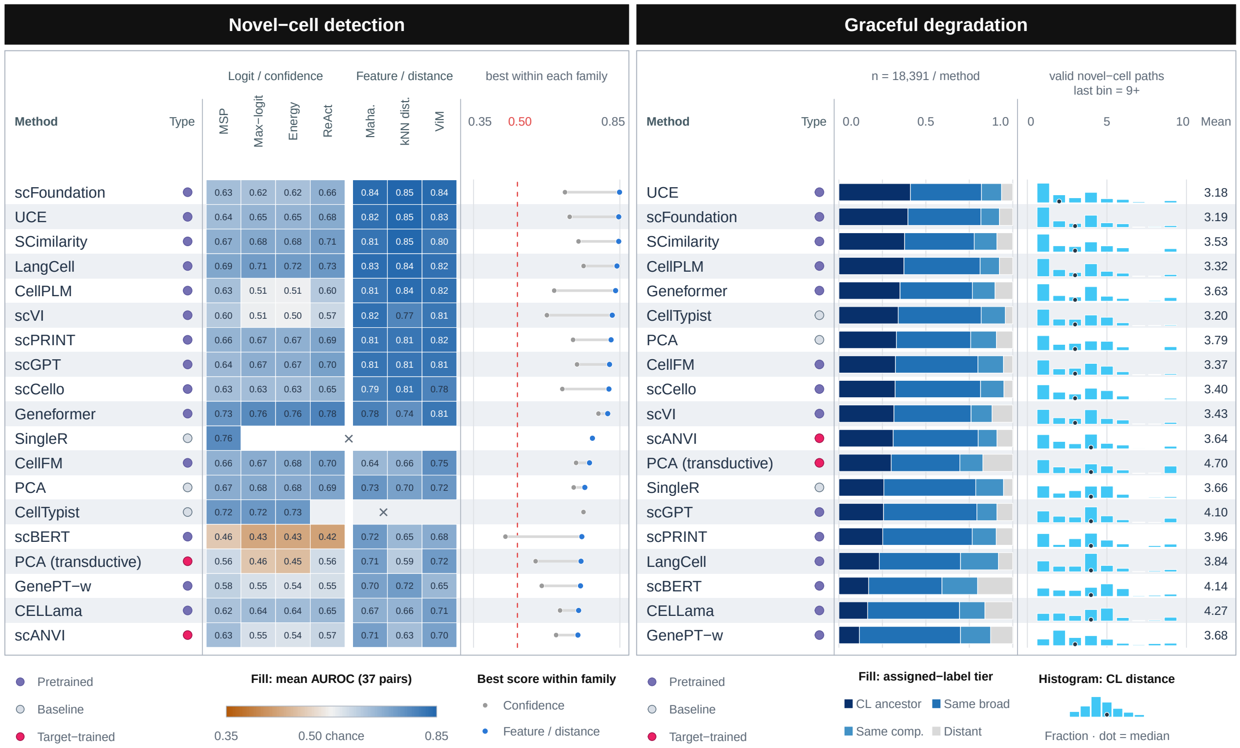

**Figure S184: Novel-type detection and semantic degradation of forced annotation.** The OOD-score block reports per-task AUROC averaged equally across the 37 eligible directed tasks for the available confidence/logit and feature/distance scores. The adjacent comparison contrasts the best available score in each detector family; the common-detector cross-method comparison is shown in Fig. S183. The ontology block assigns each forced source-label prediction for a novel target cell to mutually exclusive tiers in the order annotated ancestor, same broad category, same compartment and distant. Path-distance histograms pool cells with valid Cell Ontology paths within method; dots mark medians and numbers report uncapped means. The semantic analysis covers all 18,391 novel target cells before path-availability filtering and without an OOD-score threshold.

##### 2.7.1 *Directional open-set and shared-label annotation views*

The following gallery shows forced open-set predictions and shared-label results in both directions for each of the 22 matched tissues (Figs. S185–S228). Scalar novelty-detection results are summarized in Figs. S183 and S184.

**a** Bladder · reference: v1, query: v2 · open-world

| method | Accuracy | $Acc_{top3}$ | $Prec_{Macro}$ | $Recall_{Macro}$ | $F1_{Macro}$ | Aggregated Annotation |
|---|---|---|---|---|---|---|
| | Annotation metrics | | | | | |
| CellTypist | 0.95 | 0.99 | 0.72 | 0.83 | 0.76 | 0.82 |
| SingleR | 0.94 | n/a | 0.73 | 0.82 | 0.75 | 0.81 |
| PCA | 0.95 | 0.99 | 0.74 | 0.79 | 0.76 | 0.81 |
| scGPT | 0.94 | 0.99 | 0.74 | 0.79 | 0.75 | 0.81 |
| Geneformer | 0.94 | 0.99 | 0.71 | 0.81 | 0.73 | 0.80 |
| UCE | 0.93 | 0.98 | 0.73 | 0.79 | 0.74 | 0.80 |
| scFoundation | 0.94 | 0.98 | 0.71 | 0.80 | 0.73 | 0.80 |
| SCimilarity | 0.94 | 0.98 | 0.69 | 0.80 | 0.72 | 0.79 |
| CellFM | 0.92 | 0.99 | 0.72 | 0.77 | 0.73 | 0.78 |
| LangCell | 0.92 | 0.98 | 0.70 | 0.78 | 0.71 | 0.78 |
| scCello | 0.93 | 0.99 | 0.69 | 0.78 | 0.71 | 0.78 |
| CellPLM | 0.94 | 0.98 | 0.67 | 0.80 | 0.70 | 0.78 |
| scANVI | 0.92 | 0.98 | 0.66 | 0.79 | 0.70 | 0.77 |
| scPRINT | 0.93 | 0.98 | 0.67 | 0.78 | 0.69 | 0.76 |
| scVI | 0.93 | 0.99 | 0.66 | 0.77 | 0.69 | 0.76 |
| GenePT-w | 0.91 | 0.99 | 0.62 | 0.72 | 0.64 | 0.72 |
| CELLama | 0.74 | 0.92 | 0.35 | 0.54 | 0.39 | 0.50 |
| scBERT | 0.73 | 0.87 | 0.31 | 0.41 | 0.30 | 0.44 |

**b** Bladder · reference: v1, query: v2 · shared

| method | Accuracy | $Prec_{Macro}$ | $Recall_{Macro}$ | $F1_{Macro}$ | $AUC_{ROC}$ | $Acc_{top3}$ | Aggregated Annotation |
|---|---|---|---|---|---|---|---|
| | Annotation metrics | | | | | | |
| CellTypist | 0.95 | 0.77 | 0.83 | 0.79 | 0.98 | 0.99 | 0.84 |
| Geneformer | 0.95 | 0.75 | 0.81 | 0.76 | 0.99 | 0.99 | 0.82 |
| PCA | 0.95 | 0.75 | 0.79 | 0.76 | 0.99 | 1.00 | 0.81 |
| SingleR | 0.95 | 0.73 | 0.82 | 0.75 | n/a | n/a | 0.81 |
| SCimilarity | 0.94 | 0.74 | 0.80 | 0.75 | 1.00 | 0.99 | 0.81 |
| scGPT | 0.94 | 0.75 | 0.79 | 0.76 | 1.00 | 0.99 | 0.81 |
| UCE | 0.94 | 0.74 | 0.79 | 0.75 | 1.00 | 0.99 | 0.80 |
| scFoundation | 0.94 | 0.72 | 0.80 | 0.74 | 1.00 | 0.99 | 0.80 |
| CellPLM | 0.94 | 0.71 | 0.80 | 0.73 | 1.00 | 0.99 | 0.79 |
| CellFM | 0.93 | 0.73 | 0.77 | 0.73 | 0.99 | 0.99 | 0.79 |
| scCello | 0.94 | 0.70 | 0.78 | 0.72 | 0.99 | 0.99 | 0.78 |
| scVI | 0.94 | 0.70 | 0.77 | 0.72 | 1.00 | 0.99 | 0.78 |
| LangCell | 0.93 | 0.70 | 0.78 | 0.71 | 0.99 | 0.98 | 0.78 |
| scPRINT | 0.93 | 0.68 | 0.78 | 0.69 | 0.98 | 0.98 | 0.77 |
| scANVI | 0.92 | 0.66 | 0.79 | 0.70 | 0.99 | 0.98 | 0.77 |
| GenePT-w | 0.91 | 0.64 | 0.72 | 0.65 | 0.99 | 0.99 | 0.73 |
| CELLama | 0.74 | 0.35 | 0.54 | 0.39 | 0.94 | 0.92 | 0.51 |
| scBERT | 0.74 | 0.31 | 0.41 | 0.30 | 0.82 | 0.87 | 0.44 |

**Figure S185: Directional cross-dataset annotation for bladder, TSv1 to TSv2. a,** Evaluation of all 36,715 target cells, including 106 cells from labels absent from the source. **b,** Closed-set evaluation of 36,609 target cells from 20 labels shared by source and target. Embedding methods use source-trained logistic regression; CellTypist and SingleR use native predictions; scANVI is fitted transductively. Tables report accuracy, top-3 accuracy, macro precision, macro recall and macro-F1; panel **b** additionally reports macro one-versus-rest AUROC. Probability-based metrics are unavailable for SingleR.

**a** Bladder · reference: v2, query: v1 · open-world

| method | Annotation metrics | | | | | Aggregated |
|---|---|---|---|---|---|---|
| | *Accuracy* | $Acc_{top3}$ | $Prec_{Macro}$ | $Recall_{Macro}$ | $F1_{Macro}$ | *Annotation* |
| CellTypist | 0.90 | 0.99 | 0.79 | 0.75 | 0.75 | 0.80 |
| PCA | 0.89 | 0.99 | 0.77 | 0.76 | 0.75 | 0.79 |
| SCimilarity | 0.87 | 0.98 | 0.74 | 0.80 | 0.76 | 0.79 |
| UCE | 0.88 | 0.98 | 0.74 | 0.78 | 0.75 | 0.79 |
| scCello | 0.88 | 0.98 | 0.73 | 0.78 | 0.75 | 0.79 |
| scANVI | 0.87 | 0.97 | 0.73 | 0.79 | 0.75 | 0.79 |
| scGPT | 0.85 | 0.98 | 0.74 | 0.77 | 0.75 | 0.78 |
| scFoundation | 0.87 | 0.97 | 0.72 | 0.78 | 0.74 | 0.78 |
| Geneformer | 0.87 | 0.97 | 0.74 | 0.75 | 0.74 | 0.77 |
| SingleR | 0.84 | n/a | 0.74 | 0.77 | 0.73 | 0.77 |
| CellFM | 0.86 | 0.98 | 0.74 | 0.71 | 0.71 | 0.76 |
| CellPLM | 0.86 | 0.97 | 0.70 | 0.76 | 0.70 | 0.75 |
| scPRINT | 0.84 | 0.97 | 0.70 | 0.72 | 0.69 | 0.74 |
| LangCell | 0.83 | 0.98 | 0.68 | 0.73 | 0.69 | 0.73 |
| scVI | 0.85 | 0.98 | 0.65 | 0.76 | 0.68 | 0.73 |
| GenePT-w | 0.76 | 0.93 | 0.64 | 0.59 | 0.56 | 0.64 |
| CELLama | 0.73 | 0.90 | 0.48 | 0.56 | 0.49 | 0.57 |
| scBERT | 0.64 | 0.87 | 0.34 | 0.31 | 0.31 | 0.40 |

**b** Bladder · reference: v2, query: v1 · shared

| method | Annotation metrics | | | | | | Aggregated |
|---|---|---|---|---|---|---|---|
| | *Accuracy* | $Prec_{Macro}$ | $Recall_{Macro}$ | $F1_{Macro}$ | $AUC_{ROC}$ | $Acc_{top3}$ | *Annotation* |
| CellTypist | 0.90 | 0.79 | 0.75 | 0.75 | 0.98 | 0.99 | 0.80 |
| PCA | 0.89 | 0.77 | 0.76 | 0.75 | 0.98 | 0.99 | 0.79 |
| SCimilarity | 0.87 | 0.74 | 0.80 | 0.76 | 0.97 | 0.98 | 0.79 |
| UCE | 0.89 | 0.74 | 0.78 | 0.75 | 0.97 | 0.98 | 0.79 |
| scCello | 0.88 | 0.73 | 0.78 | 0.75 | 0.97 | 0.98 | 0.79 |
| scANVI | 0.87 | 0.73 | 0.79 | 0.75 | 0.96 | 0.98 | 0.79 |
| scGPT | 0.85 | 0.74 | 0.77 | 0.75 | 0.97 | 0.98 | 0.78 |
| scFoundation | 0.87 | 0.72 | 0.78 | 0.74 | 0.96 | 0.97 | 0.78 |
| Geneformer | 0.87 | 0.74 | 0.75 | 0.74 | 0.97 | 0.97 | 0.77 |
| SingleR | 0.84 | 0.74 | 0.77 | 0.73 | n/a | n/a | 0.77 |
| CellFM | 0.87 | 0.74 | 0.71 | 0.71 | 0.98 | 0.98 | 0.76 |
| CellPLM | 0.86 | 0.70 | 0.76 | 0.70 | 0.95 | 0.97 | 0.75 |
| scPRINT | 0.84 | 0.70 | 0.72 | 0.70 | 0.97 | 0.97 | 0.74 |
| LangCell | 0.84 | 0.68 | 0.73 | 0.69 | 0.97 | 0.98 | 0.73 |
| scVI | 0.85 | 0.65 | 0.76 | 0.68 | 0.95 | 0.98 | 0.73 |
| GenePT-w | 0.76 | 0.65 | 0.59 | 0.56 | 0.96 | 0.93 | 0.64 |
| CELLama | 0.73 | 0.48 | 0.56 | 0.49 | 0.95 | 0.90 | 0.57 |
| scBERT | 0.64 | 0.34 | 0.31 | 0.31 | 0.85 | 0.87 | 0.40 |

**Figure S186: Directional cross-dataset annotation for bladder, TSv2 to TSv1. a,** All 23,814 target cells, including 22 cells from source-absent labels. **b,** 23,792 target cells from 20 shared labels. Methods and metrics are as in Fig. S185.

**a** Blood · reference: v1, query: v2 · open-world

| method | Annotation metrics: *Accuracy* | *Acc*$_{top3}$ | *Prec*$_{Macro}$ | *Recall*$_{Macro}$ | *F1*$_{Macro}$ | Aggregated: *Annotation* |
|---|---|---|---|---|---|---|
| SCimilarity | 0.79 | 0.94 | 0.66 | 0.66 | 0.61 | 0.68 |
| CellPLM | 0.79 | 0.95 | 0.65 | 0.62 | 0.58 | 0.66 |
| scGPT | 0.77 | 0.95 | 0.63 | 0.58 | 0.58 | 0.64 |
| CellTypist | 0.77 | 0.94 | 0.63 | 0.58 | 0.57 | 0.64 |
| Geneformer | 0.77 | 0.95 | 0.63 | 0.58 | 0.57 | 0.64 |
| PCA | 0.77 | 0.94 | 0.61 | 0.57 | 0.55 | 0.63 |
| UCE | 0.76 | 0.93 | 0.60 | 0.57 | 0.55 | 0.62 |
| LangCell | 0.74 | 0.96 | 0.56 | 0.62 | 0.54 | 0.61 |
| SingleR | 0.73 | n/a | 0.61 | 0.59 | 0.53 | 0.61 |
| scFoundation | 0.73 | 0.95 | 0.57 | 0.57 | 0.54 | 0.60 |
| scVI | 0.76 | 0.95 | 0.56 | 0.56 | 0.52 | 0.60 |
| scANVI | 0.68 | 0.92 | 0.55 | 0.63 | 0.51 | 0.59 |
| scCello | 0.76 | 0.95 | 0.53 | 0.54 | 0.51 | 0.59 |
| scPRINT | 0.69 | 0.94 | 0.59 | 0.53 | 0.51 | 0.58 |
| CellFM | 0.74 | 0.93 | 0.52 | 0.55 | 0.51 | 0.58 |
| GenePT-w | 0.70 | 0.92 | 0.52 | 0.61 | 0.46 | 0.57 |
| CELLama | 0.64 | 0.90 | 0.45 | 0.56 | 0.46 | 0.53 |
| scBERT | 0.62 | 0.83 | 0.34 | 0.41 | 0.32 | 0.42 |

**b** Blood · reference: v1, query: v2 · shared

| method | Annotation metrics: *Accuracy* | *Prec*$_{Macro}$ | *Recall*$_{Macro}$ | *F1*$_{Macro}$ | *AUC*$_{ROC}$ | *Acc*$_{top3}$ | Aggregated: *Annotation* |
|---|---|---|---|---|---|---|---|
| SCimilarity | 0.79 | 0.67 | 0.66 | 0.61 | 0.97 | 0.95 | 0.68 |
| CellPLM | 0.80 | 0.66 | 0.62 | 0.58 | 0.97 | 0.96 | 0.66 |
| scGPT | 0.78 | 0.63 | 0.58 | 0.58 | 0.97 | 0.95 | 0.64 |
| CellTypist | 0.78 | 0.63 | 0.58 | 0.57 | 0.95 | 0.94 | 0.64 |
| Geneformer | 0.77 | 0.64 | 0.58 | 0.58 | 0.95 | 0.95 | 0.64 |
| PCA | 0.77 | 0.61 | 0.57 | 0.56 | 0.96 | 0.94 | 0.63 |
| UCE | 0.76 | 0.61 | 0.57 | 0.56 | 0.96 | 0.94 | 0.62 |
| LangCell | 0.74 | 0.56 | 0.62 | 0.54 | 0.97 | 0.96 | 0.62 |
| SingleR | 0.73 | 0.61 | 0.59 | 0.53 | n/a | n/a | 0.62 |
| scFoundation | 0.73 | 0.57 | 0.57 | 0.55 | 0.96 | 0.95 | 0.60 |
| scVI | 0.76 | 0.57 | 0.56 | 0.53 | 0.96 | 0.95 | 0.60 |
| scANVI | 0.68 | 0.55 | 0.63 | 0.51 | 0.95 | 0.92 | 0.59 |
| scCello | 0.76 | 0.54 | 0.54 | 0.51 | 0.97 | 0.95 | 0.59 |
| scPRINT | 0.69 | 0.60 | 0.53 | 0.51 | 0.96 | 0.94 | 0.58 |
| CellFM | 0.74 | 0.52 | 0.55 | 0.51 | 0.96 | 0.93 | 0.58 |
| GenePT-w | 0.71 | 0.52 | 0.61 | 0.46 | 0.95 | 0.92 | 0.58 |
| CELLama | 0.64 | 0.46 | 0.56 | 0.46 | 0.93 | 0.90 | 0.53 |
| scBERT | 0.62 | 0.34 | 0.41 | 0.32 | 0.90 | 0.83 | 0.42 |

**Figure S187: Directional cross-dataset annotation for blood, TSv1 to TSv2. a,** All 17,802 target cells, including 40 cells from source-absent labels. **b,** 17,762 target cells from 18 shared labels. Methods and metrics are as in Fig. S185.

**a** Blood · reference: v2, query: v1 · open-world

| method | Annotation metrics | | | | | Aggregated |
|---|---|---|---|---|---|---|
| | *Accuracy* | $Acc_{top3}$ | $Prec_{Macro}$ | $Recall_{Macro}$ | $F1_{Macro}$ | *Annotation* |
| scANVI | 0.68 | 0.93 | 0.52 | 0.65 | 0.54 | 0.60 |
| UCE | 0.66 | 0.91 | 0.54 | 0.63 | 0.54 | 0.59 |
| CellPLM | 0.71 | 0.92 | 0.49 | 0.65 | 0.50 | 0.59 |
| scVI | 0.69 | 0.92 | 0.52 | 0.61 | 0.52 | 0.58 |
| SCimilarity | 0.64 | 0.91 | 0.53 | 0.63 | 0.52 | 0.58 |
| scGPT | 0.66 | 0.91 | 0.51 | 0.63 | 0.51 | 0.58 |
| CellTypist | 0.65 | 0.92 | 0.54 | 0.58 | 0.51 | 0.57 |
| scFoundation | 0.66 | 0.89 | 0.49 | 0.64 | 0.49 | 0.57 |
| Geneformer | 0.64 | 0.89 | 0.50 | 0.60 | 0.50 | 0.56 |
| PCA | 0.64 | 0.91 | 0.52 | 0.56 | 0.51 | 0.56 |
| LangCell | 0.64 | 0.88 | 0.52 | 0.58 | 0.49 | 0.56 |
| SingleR | 0.58 | n/a | 0.50 | 0.61 | 0.48 | 0.55 |
| scPRINT | 0.62 | 0.88 | 0.48 | 0.55 | 0.49 | 0.53 |
| scCello | 0.60 | 0.89 | 0.47 | 0.58 | 0.47 | 0.53 |
| CellFM | 0.63 | 0.88 | 0.52 | 0.51 | 0.46 | 0.53 |
| CELLama | 0.59 | 0.88 | 0.40 | 0.46 | 0.40 | 0.46 |
| scBERT | 0.50 | 0.80 | 0.32 | 0.39 | 0.30 | 0.38 |
| GenePT-w | 0.46 | 0.56 | 0.21 | 0.26 | 0.18 | 0.28 |

**b** Blood · reference: v2, query: v1 · shared

| method | Annotation metrics | | | | | | Aggregated |
|---|---|---|---|---|---|---|---|
| | *Accuracy* | $Prec_{Macro}$ | $Recall_{Macro}$ | $F1_{Macro}$ | $AUC_{ROC}$ | $Acc_{top3}$ | *Annotation* |
| scANVI | 0.68 | 0.52 | 0.65 | 0.54 | 0.92 | 0.93 | 0.60 |
| UCE | 0.66 | 0.54 | 0.63 | 0.54 | 0.95 | 0.91 | 0.59 |
| CellPLM | 0.71 | 0.49 | 0.65 | 0.50 | 0.95 | 0.92 | 0.59 |
| scVI | 0.69 | 0.52 | 0.61 | 0.52 | 0.96 | 0.93 | 0.58 |
| SCimilarity | 0.64 | 0.53 | 0.63 | 0.52 | 0.94 | 0.91 | 0.58 |
| scGPT | 0.66 | 0.51 | 0.63 | 0.52 | 0.94 | 0.91 | 0.58 |
| CellTypist | 0.65 | 0.54 | 0.58 | 0.51 | 0.95 | 0.92 | 0.57 |
| scFoundation | 0.66 | 0.49 | 0.64 | 0.49 | 0.94 | 0.89 | 0.57 |
| Geneformer | 0.64 | 0.50 | 0.60 | 0.50 | 0.94 | 0.89 | 0.56 |
| PCA | 0.64 | 0.52 | 0.56 | 0.51 | 0.92 | 0.91 | 0.56 |
| LangCell | 0.64 | 0.52 | 0.58 | 0.49 | 0.94 | 0.88 | 0.56 |
| SingleR | 0.58 | 0.50 | 0.61 | 0.48 | n/a | n/a | 0.55 |
| scPRINT | 0.62 | 0.48 | 0.55 | 0.49 | 0.94 | 0.88 | 0.53 |
| scCello | 0.60 | 0.47 | 0.58 | 0.47 | 0.95 | 0.89 | 0.53 |
| CellFM | 0.63 | 0.52 | 0.51 | 0.46 | 0.95 | 0.88 | 0.53 |
| CELLama | 0.59 | 0.40 | 0.46 | 0.40 | 0.92 | 0.88 | 0.46 |
| scBERT | 0.50 | 0.32 | 0.39 | 0.30 | 0.87 | 0.80 | 0.38 |
| GenePT-w | 0.46 | 0.21 | 0.26 | 0.18 | 0.91 | 0.56 | 0.28 |

**Figure S188: Directional cross-dataset annotation for blood, TSv2 to TSv1. a,** All 46,423 target cells, including 12 cells from source-absent labels. **b,** 46,411 target cells from 18 shared labels. Methods and metrics are as in Fig. S185.

**a** Bone Marrow · reference: v1, query: v2 · open-world

| | Annotation metrics | | | | | Aggregated |
|---|---|---|---|---|---|---|
| method | *Accuracy* | $Acc_{top3}$ | $Prec_{Macro}$ | $Recall_{Macro}$ | $F1_{Macro}$ | *Annotation* |
| CellTypist | 0.78 | 0.89 | 0.62 | 0.62 | 0.58 | 0.65 |
| UCE | 0.75 | 0.88 | 0.54 | 0.63 | 0.53 | 0.61 |
| Geneformer | 0.76 | 0.88 | 0.53 | 0.59 | 0.51 | 0.60 |
| SCimilarity | 0.75 | 0.89 | 0.55 | 0.58 | 0.51 | 0.60 |
| scPRINT | 0.73 | 0.86 | 0.53 | 0.60 | 0.52 | 0.59 |
| CellPLM | 0.72 | 0.88 | 0.53 | 0.60 | 0.52 | 0.59 |
| SingleR | 0.73 | n/a | 0.52 | 0.63 | 0.49 | 0.59 |
| scANVI | 0.74 | 0.89 | 0.51 | 0.58 | 0.50 | 0.59 |
| scFoundation | 0.74 | 0.88 | 0.51 | 0.59 | 0.49 | 0.58 |
| scGPT | 0.76 | 0.89 | 0.52 | 0.54 | 0.50 | 0.58 |
| scCello | 0.73 | 0.88 | 0.48 | 0.58 | 0.49 | 0.57 |
| CellFM | 0.73 | 0.87 | 0.53 | 0.51 | 0.47 | 0.56 |
| scVI | 0.71 | 0.87 | 0.48 | 0.54 | 0.47 | 0.55 |
| LangCell | 0.71 | 0.89 | 0.49 | 0.52 | 0.47 | 0.55 |
| PCA | 0.74 | 0.88 | 0.50 | 0.48 | 0.45 | 0.54 |
| GenePT-w | 0.58 | 0.79 | 0.40 | 0.50 | 0.34 | 0.45 |
| CELLama | 0.57 | 0.81 | 0.33 | 0.36 | 0.31 | 0.39 |
| scBERT | 0.51 | 0.74 | 0.29 | 0.31 | 0.27 | 0.34 |

**b** Bone Marrow · reference: v1, query: v2 · shared

| | Annotation metrics | | | | | | Aggregated |
|---|---|---|---|---|---|---|---|
| method | *Accuracy* | $Prec_{Macro}$ | $Recall_{Macro}$ | $F1_{Macro}$ | $AUC_{ROC}$ | $Acc_{top3}$ | *Annotation* |
| CellTypist | 0.78 | 0.63 | 0.62 | 0.58 | 0.93 | 0.89 | 0.65 |
| UCE | 0.76 | 0.54 | 0.63 | 0.54 | 0.91 | 0.88 | 0.62 |
| Geneformer | 0.77 | 0.53 | 0.59 | 0.51 | 0.92 | 0.88 | 0.60 |
| SCimilarity | 0.75 | 0.55 | 0.58 | 0.51 | 0.94 | 0.89 | 0.60 |
| scPRINT | 0.73 | 0.53 | 0.60 | 0.52 | 0.94 | 0.87 | 0.60 |
| CellPLM | 0.72 | 0.53 | 0.60 | 0.52 | 0.92 | 0.88 | 0.59 |
| SingleR | 0.73 | 0.52 | 0.63 | 0.49 | n/a | n/a | 0.59 |
| scANVI | 0.75 | 0.51 | 0.58 | 0.50 | 0.94 | 0.89 | 0.59 |
| scFoundation | 0.75 | 0.51 | 0.59 | 0.49 | 0.91 | 0.88 | 0.59 |
| scGPT | 0.76 | 0.52 | 0.54 | 0.50 | 0.91 | 0.89 | 0.58 |
| scCello | 0.73 | 0.48 | 0.58 | 0.49 | 0.95 | 0.89 | 0.57 |
| CellFM | 0.73 | 0.53 | 0.51 | 0.47 | 0.91 | 0.87 | 0.56 |
| scVI | 0.72 | 0.48 | 0.54 | 0.47 | 0.95 | 0.87 | 0.55 |
| LangCell | 0.72 | 0.49 | 0.52 | 0.47 | 0.92 | 0.89 | 0.55 |
| PCA | 0.74 | 0.51 | 0.48 | 0.45 | 0.91 | 0.88 | 0.55 |
| GenePT-w | 0.58 | 0.40 | 0.50 | 0.35 | 0.88 | 0.79 | 0.46 |
| CELLama | 0.57 | 0.33 | 0.36 | 0.31 | 0.87 | 0.82 | 0.39 |
| scBERT | 0.51 | 0.29 | 0.31 | 0.27 | 0.84 | 0.74 | 0.34 |

**Figure S189: Directional cross-dataset annotation for bone marrow, TSv1 to TSv2. a,** All 8,045 target cells, including 22 cells from source-absent labels. **b,** 8,023 target cells from 22 shared labels. Methods and metrics are as in Fig. S185.

**a** Bone Marrow · reference: v2, query: v1 · open-world

| method | Annotation metrics: *Accuracy* | *$Acc_{top3}$* | *$Prec_{Macro}$* | *$Recall_{Macro}$* | *$F1_{Macro}$* | Aggregated: *Annotation* |
|---|---|---|---|---|---|---|
| SCimilarity | 0.78 | 0.94 | 0.54 | 0.65 | 0.54 | 0.63 |
| UCE | 0.76 | 0.93 | 0.56 | 0.62 | 0.54 | 0.62 |
| PCA | 0.82 | 0.96 | 0.58 | 0.56 | 0.53 | 0.62 |
| scFoundation | 0.79 | 0.94 | 0.51 | 0.62 | 0.52 | 0.61 |
| scVI | 0.78 | 0.96 | 0.55 | 0.60 | 0.52 | 0.61 |
| scCello | 0.76 | 0.95 | 0.55 | 0.60 | 0.52 | 0.61 |
| CellPLM | 0.72 | 0.90 | 0.58 | 0.59 | 0.53 | 0.60 |
| CellTypist | 0.83 | 0.96 | 0.54 | 0.52 | 0.49 | 0.59 |
| scGPT | 0.73 | 0.95 | 0.51 | 0.61 | 0.51 | 0.59 |
| SingleR | 0.73 | n/a | 0.51 | 0.59 | 0.49 | 0.58 |
| Geneformer | 0.72 | 0.88 | 0.50 | 0.54 | 0.49 | 0.56 |
| scANVI | 0.76 | 0.93 | 0.48 | 0.52 | 0.47 | 0.56 |
| scPRINT | 0.72 | 0.89 | 0.51 | 0.52 | 0.48 | 0.56 |
| LangCell | 0.74 | 0.92 | 0.52 | 0.49 | 0.47 | 0.55 |
| CellFM | 0.74 | 0.93 | 0.43 | 0.46 | 0.42 | 0.51 |
| CELLama | 0.61 | 0.84 | 0.37 | 0.39 | 0.35 | 0.43 |
| scBERT | 0.50 | 0.77 | 0.28 | 0.28 | 0.25 | 0.33 |
| GenePT-w | 0.44 | 0.66 | 0.32 | 0.31 | 0.21 | 0.32 |

**b** Bone Marrow · reference: v2, query: v1 · shared

| method | Annotation metrics: *Accuracy* | *$Prec_{Macro}$* | *$Recall_{Macro}$* | *$F1_{Macro}$* | *$AUC_{ROC}$* | *$Acc_{top3}$* | Aggregated: *Annotation* |
|---|---|---|---|---|---|---|---|
| SCimilarity | 0.78 | 0.54 | 0.65 | 0.54 | 0.97 | 0.94 | 0.63 |
| UCE | 0.76 | 0.56 | 0.62 | 0.54 | 0.95 | 0.93 | 0.62 |
| PCA | 0.82 | 0.58 | 0.56 | 0.53 | 0.96 | 0.96 | 0.62 |
| scFoundation | 0.79 | 0.51 | 0.62 | 0.52 | 0.96 | 0.94 | 0.61 |
| scVI | 0.78 | 0.55 | 0.60 | 0.52 | 0.96 | 0.96 | 0.61 |
| scCello | 0.76 | 0.55 | 0.60 | 0.52 | 0.96 | 0.95 | 0.61 |
| CellPLM | 0.72 | 0.58 | 0.59 | 0.53 | 0.91 | 0.90 | 0.60 |
| CellTypist | 0.83 | 0.54 | 0.52 | 0.49 | 0.97 | 0.96 | 0.59 |
| scGPT | 0.73 | 0.51 | 0.61 | 0.51 | 0.94 | 0.95 | 0.59 |
| SingleR | 0.73 | 0.51 | 0.59 | 0.49 | n/a | n/a | 0.58 |
| Geneformer | 0.72 | 0.50 | 0.54 | 0.49 | 0.96 | 0.88 | 0.56 |
| scANVI | 0.76 | 0.48 | 0.52 | 0.47 | 0.95 | 0.93 | 0.56 |
| scPRINT | 0.72 | 0.51 | 0.52 | 0.48 | 0.95 | 0.89 | 0.56 |
| LangCell | 0.74 | 0.52 | 0.49 | 0.47 | 0.95 | 0.92 | 0.55 |
| CellFM | 0.74 | 0.43 | 0.46 | 0.42 | 0.94 | 0.93 | 0.51 |
| CELLama | 0.61 | 0.37 | 0.39 | 0.35 | 0.92 | 0.84 | 0.43 |
| scBERT | 0.50 | 0.28 | 0.28 | 0.25 | 0.84 | 0.77 | 0.33 |
| GenePT-w | 0.44 | 0.32 | 0.31 | 0.21 | 0.94 | 0.66 | 0.32 |

**Figure S190: Directional cross-dataset annotation for bone marrow, TSv2 to TSv1. a,b,** Both panels contain 11,901 target cells from 22 shared labels because no target label is absent from the source in this direction. Methods and metrics are as in Fig. S185.

**a**

**Eye · reference: v1, query: v2 · open-world**

| method | Annotation metrics | | | | | Aggregated |
|---|---|---|---|---|---|---|
| | *Accuracy* | *Acc*$_{top3}$ | *Prec*$_{Macro}$ | *Recall*$_{Macro}$ | *F1*$_{Macro}$ | *Annotation* |
| **SCimilarity** | 0.85 | 0.95 | 0.60 | 0.69 | 0.60 | 0.68 |
| **scFoundation** | 0.85 | 0.94 | 0.61 | 0.67 | 0.60 | 0.68 |
| **CellTypist** | 0.86 | 0.95 | 0.63 | 0.64 | 0.60 | 0.68 |
| **scGPT** | 0.85 | 0.95 | 0.61 | 0.65 | 0.60 | 0.68 |
| **Geneformer** | 0.84 | 0.95 | 0.61 | 0.66 | 0.60 | 0.67 |
| **CellPLM** | 0.84 | 0.94 | 0.60 | 0.67 | 0.58 | 0.67 |
| **SingleR** | 0.83 | n/a | 0.58 | 0.69 | 0.57 | 0.67 |
| **PCA** | 0.83 | 0.95 | 0.62 | 0.64 | 0.59 | 0.67 |
| **UCE** | 0.82 | 0.93 | 0.60 | 0.64 | 0.58 | 0.66 |
| **LangCell** | 0.83 | 0.94 | 0.59 | 0.65 | 0.58 | 0.66 |
| **scANVI** | 0.84 | 0.95 | 0.57 | 0.67 | 0.57 | 0.66 |
| **scCello** | 0.82 | 0.94 | 0.59 | 0.66 | 0.58 | 0.66 |
| **scPRINT** | 0.84 | 0.95 | 0.58 | 0.63 | 0.57 | 0.65 |
| **CellFM** | 0.82 | 0.95 | 0.60 | 0.60 | 0.55 | 0.64 |
| **scVI** | 0.83 | 0.95 | 0.56 | 0.60 | 0.53 | 0.63 |
| **GenePT-w** | 0.76 | 0.92 | 0.57 | 0.61 | 0.49 | 0.61 |
| **CELLama** | 0.70 | 0.89 | 0.44 | 0.51 | 0.43 | 0.52 |
| **scBERT** | 0.36 | 0.67 | 0.25 | 0.25 | 0.19 | 0.26 |

**b**

**Eye · reference: v1, query: v2 · shared**

| method | Annotation metrics | | | | | | Aggregated |
|---|---|---|---|---|---|---|---|
| | *Accuracy* | *Prec*$_{Macro}$ | *Recall*$_{Macro}$ | *F1*$_{Macro}$ | *AUC*$_{ROC}$ | *Acc*$_{top3}$ | *Annotation* |
| **SCimilarity** | 0.85 | 0.60 | 0.69 | 0.60 | 0.98 | 0.95 | 0.69 |
| **scFoundation** | 0.85 | 0.61 | 0.67 | 0.60 | 0.97 | 0.94 | 0.69 |
| **CellTypist** | 0.86 | 0.63 | 0.64 | 0.60 | 0.98 | 0.95 | 0.68 |
| **scGPT** | 0.85 | 0.62 | 0.65 | 0.60 | 0.98 | 0.95 | 0.68 |
| **Geneformer** | 0.84 | 0.61 | 0.66 | 0.60 | 0.97 | 0.95 | 0.68 |
| **CellPLM** | 0.84 | 0.60 | 0.67 | 0.58 | 0.98 | 0.94 | 0.67 |
| **SingleR** | 0.83 | 0.58 | 0.69 | 0.57 | n/a | n/a | 0.67 |
| **PCA** | 0.83 | 0.62 | 0.64 | 0.59 | 0.98 | 0.95 | 0.67 |
| **UCE** | 0.83 | 0.61 | 0.64 | 0.58 | 0.97 | 0.93 | 0.67 |
| **scANVI** | 0.84 | 0.58 | 0.67 | 0.57 | 0.97 | 0.95 | 0.66 |
| **LangCell** | 0.83 | 0.60 | 0.65 | 0.58 | 0.97 | 0.94 | 0.66 |
| **scCello** | 0.82 | 0.60 | 0.66 | 0.59 | 0.97 | 0.94 | 0.66 |
| **scPRINT** | 0.84 | 0.58 | 0.63 | 0.57 | 0.98 | 0.95 | 0.66 |
| **CellFM** | 0.82 | 0.61 | 0.60 | 0.55 | 0.98 | 0.95 | 0.64 |
| **scVI** | 0.83 | 0.57 | 0.60 | 0.53 | 0.97 | 0.95 | 0.63 |
| **GenePT-w** | 0.76 | 0.57 | 0.61 | 0.49 | 0.98 | 0.92 | 0.61 |
| **CELLama** | 0.70 | 0.44 | 0.51 | 0.43 | 0.92 | 0.89 | 0.52 |
| **scBERT** | 0.36 | 0.25 | 0.25 | 0.19 | 0.77 | 0.67 | 0.26 |

**Figure S191: Directional cross-dataset annotation for eye, TSv1 to TSv2. a,** All 21,121 target cells, including 4 cells from source-absent labels. **b,** 21,117 target cells from 31 shared labels. Methods and metrics are as in Fig. S185.

**a** Eye · reference: v2, query: v1 · open-world

| method | Annotation metrics: *Accuracy* | $Acc_{top3}$ | $Prec_{Macro}$ | $Recall_{Macro}$ | $F1_{Macro}$ | Aggregated: *Annotation* |
|---|---|---|---|---|---|---|
| scFoundation | 0.88 | 0.97 | 0.69 | 0.69 | 0.66 | 0.73 |
| PCA | 0.89 | 0.98 | 0.68 | 0.69 | 0.64 | 0.72 |
| CellTypist | 0.89 | 0.98 | 0.66 | 0.67 | 0.62 | 0.71 |
| UCE | 0.86 | 0.96 | 0.70 | 0.65 | 0.63 | 0.71 |
| SCimilarity | 0.88 | 0.97 | 0.65 | 0.67 | 0.63 | 0.71 |
| Geneformer | 0.87 | 0.95 | 0.69 | 0.64 | 0.63 | 0.71 |
| scGPT | 0.86 | 0.97 | 0.67 | 0.66 | 0.63 | 0.71 |
| CellPLM | 0.88 | 0.97 | 0.63 | 0.68 | 0.62 | 0.70 |
| scPRINT | 0.85 | 0.96 | 0.69 | 0.62 | 0.63 | 0.70 |
| SingleR | 0.86 | n/a | 0.62 | 0.68 | 0.61 | 0.69 |
| scCello | 0.84 | 0.97 | 0.65 | 0.64 | 0.62 | 0.68 |
| LangCell | 0.85 | 0.96 | 0.63 | 0.63 | 0.59 | 0.68 |
| scVI | 0.82 | 0.96 | 0.60 | 0.64 | 0.58 | 0.66 |
| CellFM | 0.83 | 0.96 | 0.62 | 0.61 | 0.57 | 0.66 |
| scANVI | 0.84 | 0.95 | 0.55 | 0.64 | 0.56 | 0.65 |
| CELLama | 0.69 | 0.88 | 0.41 | 0.47 | 0.40 | 0.49 |
| GenePT-w | 0.60 | 0.82 | 0.57 | 0.41 | 0.35 | 0.48 |
| scBERT | 0.51 | 0.82 | 0.21 | 0.25 | 0.19 | 0.29 |

**b** Eye · reference: v2, query: v1 · shared

| method | Annotation metrics: *Accuracy* | $Prec_{Macro}$ | $Recall_{Macro}$ | $F1_{Macro}$ | $AUC_{ROC}$ | $Acc_{top3}$ | Aggregated: *Annotation* |
|---|---|---|---|---|---|---|---|
| scFoundation | 0.89 | 0.70 | 0.69 | 0.66 | 0.97 | 0.98 | 0.74 |
| PCA | 0.89 | 0.69 | 0.69 | 0.65 | 0.99 | 0.99 | 0.73 |
| CellTypist | 0.89 | 0.68 | 0.67 | 0.63 | 0.98 | 0.98 | 0.72 |
| SCimilarity | 0.89 | 0.66 | 0.67 | 0.64 | 0.98 | 0.98 | 0.72 |
| UCE | 0.87 | 0.71 | 0.65 | 0.63 | 0.98 | 0.97 | 0.72 |
| Geneformer | 0.88 | 0.70 | 0.64 | 0.64 | 0.96 | 0.96 | 0.71 |
| scGPT | 0.87 | 0.68 | 0.66 | 0.63 | 0.98 | 0.97 | 0.71 |
| CellPLM | 0.88 | 0.65 | 0.68 | 0.63 | 0.97 | 0.98 | 0.71 |
| scPRINT | 0.85 | 0.70 | 0.62 | 0.64 | 0.96 | 0.96 | 0.70 |
| SingleR | 0.87 | 0.64 | 0.68 | 0.62 | n/a | n/a | 0.70 |
| scCello | 0.84 | 0.66 | 0.64 | 0.62 | 0.98 | 0.98 | 0.69 |
| LangCell | 0.86 | 0.64 | 0.63 | 0.60 | 0.97 | 0.97 | 0.68 |
| scVI | 0.83 | 0.61 | 0.64 | 0.59 | 0.96 | 0.97 | 0.67 |
| CellFM | 0.84 | 0.63 | 0.61 | 0.58 | 0.98 | 0.97 | 0.66 |
| scANVI | 0.85 | 0.56 | 0.64 | 0.56 | 0.95 | 0.96 | 0.65 |
| CELLama | 0.70 | 0.42 | 0.47 | 0.41 | 0.94 | 0.88 | 0.50 |
| GenePT-w | 0.60 | 0.57 | 0.41 | 0.35 | 0.94 | 0.83 | 0.48 |
| scBERT | 0.52 | 0.21 | 0.25 | 0.19 | 0.84 | 0.83 | 0.29 |

**Figure S192: Directional cross-dataset annotation for eye, TSv2 to TSv1. a,** All 9,266 target cells, including 76 cells from source-absent labels. **b,** 9,190 target cells from 31 shared labels. Methods and metrics are as in Fig. S185.

**a** Fat · reference: v1, query: v2 · open-world

| method | Annotation metrics | | | | | Aggregated |
|---|---|---|---|---|---|---|
| | *Accuracy* | *Acc*$_{top3}$ | *Prec*$_{Macro}$ | *Recall*$_{Macro}$ | *F1*$_{Macro}$ | *Annotation* |
| **scGPT** | 0.83 | 0.95 | 0.58 | 0.68 | 0.58 | 0.67 |
| **scVI** | 0.83 | 0.95 | 0.57 | 0.63 | 0.56 | 0.65 |
| **scFoundation** | 0.76 | 0.95 | 0.56 | 0.69 | 0.57 | 0.65 |
| **UCE** | 0.76 | 0.94 | 0.57 | 0.67 | 0.57 | 0.64 |
| **scCello** | 0.79 | 0.96 | 0.54 | 0.67 | 0.55 | 0.64 |
| **LangCell** | 0.76 | 0.94 | 0.53 | 0.66 | 0.55 | 0.63 |
| **SCimilarity** | 0.71 | 0.94 | 0.54 | 0.64 | 0.53 | 0.61 |
| **CellTypist** | 0.73 | 0.93 | 0.51 | 0.63 | 0.51 | 0.60 |
| **CellPLM** | 0.65 | 0.82 | 0.53 | 0.64 | 0.51 | 0.58 |
| **scPRINT** | 0.70 | 0.92 | 0.51 | 0.60 | 0.50 | 0.58 |
| **Geneformer** | 0.62 | 0.86 | 0.50 | 0.63 | 0.51 | 0.57 |
| **scANVI** | 0.62 | 0.76 | 0.52 | 0.57 | 0.48 | 0.55 |
| **SingleR** | 0.56 | n/a | 0.50 | 0.64 | 0.45 | 0.54 |
| **PCA** | 0.66 | 0.87 | 0.47 | 0.58 | 0.44 | 0.54 |
| **CellFM** | 0.60 | 0.77 | 0.45 | 0.54 | 0.41 | 0.50 |
| **GenePT-w** | 0.51 | 0.74 | 0.42 | 0.50 | 0.37 | 0.45 |
| **CELLama** | 0.49 | 0.70 | 0.34 | 0.44 | 0.35 | 0.41 |
| **scBERT** | 0.37 | 0.63 | 0.29 | 0.30 | 0.23 | 0.30 |

**b** Fat · reference: v1, query: v2 · shared

| method | Annotation metrics | | | | | | Aggregated |
|---|---|---|---|---|---|---|---|
| | *Accuracy* | *Prec*$_{Macro}$ | *Recall*$_{Macro}$ | *F1*$_{Macro}$ | *AUC*$_{ROC}$ | *Acc*$_{top3}$ | *Annotation* |
| **scGPT** | 0.84 | 0.61 | 0.68 | 0.60 | 0.97 | 0.97 | 0.68 |
| **scVI** | 0.85 | 0.59 | 0.63 | 0.57 | 0.96 | 0.97 | 0.66 |
| **scFoundation** | 0.77 | 0.57 | 0.69 | 0.58 | 0.95 | 0.97 | 0.66 |
| **scCello** | 0.81 | 0.58 | 0.67 | 0.56 | 0.94 | 0.98 | 0.65 |
| **UCE** | 0.78 | 0.59 | 0.67 | 0.58 | 0.94 | 0.96 | 0.65 |
| **LangCell** | 0.78 | 0.58 | 0.66 | 0.56 | 0.97 | 0.96 | 0.64 |
| **SCimilarity** | 0.72 | 0.59 | 0.64 | 0.55 | 0.96 | 0.96 | 0.63 |
| **CellTypist** | 0.74 | 0.55 | 0.63 | 0.51 | 0.96 | 0.95 | 0.61 |
| **scPRINT** | 0.72 | 0.55 | 0.60 | 0.51 | 0.95 | 0.94 | 0.59 |
| **CellPLM** | 0.66 | 0.55 | 0.64 | 0.52 | 0.96 | 0.84 | 0.59 |
| **Geneformer** | 0.63 | 0.57 | 0.63 | 0.52 | 0.93 | 0.88 | 0.59 |
| **scANVI** | 0.63 | 0.55 | 0.57 | 0.49 | 0.94 | 0.78 | 0.56 |
| **SingleR** | 0.57 | 0.53 | 0.64 | 0.47 | n/a | n/a | 0.55 |
| **PCA** | 0.68 | 0.49 | 0.58 | 0.45 | 0.94 | 0.88 | 0.55 |
| **CellFM** | 0.62 | 0.50 | 0.54 | 0.41 | 0.93 | 0.78 | 0.52 |
| **GenePT-w** | 0.52 | 0.44 | 0.50 | 0.38 | 0.93 | 0.75 | 0.46 |
| **CELLama** | 0.50 | 0.35 | 0.44 | 0.36 | 0.85 | 0.71 | 0.41 |
| **scBERT** | 0.38 | 0.29 | 0.30 | 0.23 | 0.75 | 0.65 | 0.30 |

**Figure S193: Directional cross-dataset annotation for fat, TSv1 to TSv2. a,** All 71,021 target cells, including 1,426 cells from source-absent labels. **b,** 69,595 target cells from 18 shared labels. Methods and metrics are as in Fig. S185.

**a** Fat · reference: v2, query: v1 · open-world

| method | Annotation metrics: *Accuracy* | *Acc*$_{top3}$ | *Prec*$_{Macro}$ | *Recall*$_{Macro}$ | *F1*$_{Macro}$ | Aggregated: *Annotation* |
|---|---|---|---|---|---|---|
| SCimilarity | 0.83 | 0.98 | 0.61 | 0.68 | 0.56 | 0.67 |
| PCA | 0.82 | 0.98 | 0.60 | 0.66 | 0.56 | 0.66 |
| SingleR | 0.82 | n/a | 0.56 | 0.67 | 0.57 | 0.65 |
| scFoundation | 0.81 | 0.97 | 0.63 | 0.62 | 0.53 | 0.64 |
| CellTypist | 0.81 | 0.98 | 0.59 | 0.63 | 0.53 | 0.64 |
| scVI | 0.77 | 0.97 | 0.59 | 0.64 | 0.51 | 0.63 |
| scGPT | 0.75 | 0.97 | 0.60 | 0.64 | 0.51 | 0.63 |
| scPRINT | 0.80 | 0.96 | 0.62 | 0.60 | 0.48 | 0.62 |
| UCE | 0.76 | 0.98 | 0.58 | 0.63 | 0.50 | 0.62 |
| LangCell | 0.81 | 0.97 | 0.58 | 0.59 | 0.48 | 0.62 |
| scCello | 0.73 | 0.98 | 0.58 | 0.63 | 0.51 | 0.61 |
| CellPLM | 0.80 | 0.97 | 0.56 | 0.57 | 0.46 | 0.60 |
| Geneformer | 0.80 | 0.96 | 0.54 | 0.57 | 0.47 | 0.60 |
| scANVI | 0.75 | 0.95 | 0.48 | 0.64 | 0.47 | 0.59 |
| CellFM | 0.68 | 0.98 | 0.57 | 0.59 | 0.49 | 0.59 |
| GenePT-w | 0.47 | 0.93 | 0.50 | 0.56 | 0.37 | 0.48 |
| CELLama | 0.66 | 0.89 | 0.37 | 0.45 | 0.35 | 0.46 |
| scBERT | 0.67 | 0.86 | 0.28 | 0.36 | 0.26 | 0.39 |

**b** Fat · reference: v2, query: v1 · shared

| method | Annotation metrics: *Accuracy* | *Prec*$_{Macro}$ | *Recall*$_{Macro}$ | *F1*$_{Macro}$ | *AUC*$_{ROC}$ | *Acc*$_{top3}$ | Aggregated: *Annotation* |
|---|---|---|---|---|---|---|---|
| SCimilarity | 0.83 | 0.62 | 0.68 | 0.57 | 0.97 | 0.99 | 0.67 |
| PCA | 0.83 | 0.61 | 0.66 | 0.56 | 0.98 | 0.99 | 0.66 |
| SingleR | 0.83 | 0.58 | 0.67 | 0.58 | n/a | n/a | 0.66 |
| scFoundation | 0.81 | 0.63 | 0.62 | 0.53 | 0.97 | 0.98 | 0.65 |
| CellTypist | 0.81 | 0.59 | 0.63 | 0.53 | 0.98 | 0.99 | 0.64 |
| scVI | 0.78 | 0.61 | 0.64 | 0.52 | 0.96 | 0.98 | 0.64 |
| scGPT | 0.76 | 0.60 | 0.64 | 0.52 | 0.97 | 0.98 | 0.63 |
| scPRINT | 0.81 | 0.62 | 0.60 | 0.49 | 0.96 | 0.97 | 0.63 |
| UCE | 0.77 | 0.59 | 0.63 | 0.51 | 0.97 | 0.98 | 0.63 |
| LangCell | 0.82 | 0.59 | 0.59 | 0.48 | 0.98 | 0.98 | 0.62 |
| scCello | 0.74 | 0.59 | 0.63 | 0.52 | 0.96 | 0.98 | 0.62 |
| CellPLM | 0.81 | 0.56 | 0.57 | 0.46 | 0.89 | 0.98 | 0.60 |
| Geneformer | 0.80 | 0.55 | 0.57 | 0.47 | 0.98 | 0.97 | 0.60 |
| scANVI | 0.76 | 0.50 | 0.64 | 0.48 | 0.91 | 0.95 | 0.60 |
| CellFM | 0.69 | 0.59 | 0.59 | 0.50 | 0.96 | 0.98 | 0.59 |
| GenePT-w | 0.48 | 0.50 | 0.56 | 0.37 | 0.97 | 0.94 | 0.48 |
| CELLama | 0.67 | 0.37 | 0.45 | 0.35 | 0.88 | 0.89 | 0.46 |
| scBERT | 0.67 | 0.28 | 0.36 | 0.26 | 0.78 | 0.87 | 0.39 |

**Figure S194: Directional cross-dataset annotation for fat, TSv2 to TSv1. a,** All 19,666 target cells, including 131 cells from source-absent labels. **b,** 19,535 target cells from 18 shared labels. Methods and metrics are as in Fig. S185.

**a** Heart · reference: v1, query: v2 · open-world

| method | Accuracy | $Acc_{top3}$ | $Prec_{Macro}$ | $Recall_{Macro}$ | $F1_{Macro}$ | Annotation |
|---|---|---|---|---|---|---|
| | Annotation metrics | | | | | Aggregated |
| scGPT | 0.78 | 0.83 | 0.56 | 0.59 | 0.52 | 0.61 |
| scFoundation | 0.78 | 0.81 | 0.52 | 0.61 | 0.53 | 0.61 |
| Geneformer | 0.77 | 0.81 | 0.56 | 0.55 | 0.49 | 0.59 |
| scPRINT | 0.76 | 0.81 | 0.47 | 0.57 | 0.49 | 0.57 |
| UCE | 0.75 | 0.81 | 0.48 | 0.59 | 0.48 | 0.57 |
| scANVI | 0.76 | 0.82 | 0.46 | 0.54 | 0.46 | 0.55 |
| scCello | 0.75 | 0.81 | 0.52 | 0.48 | 0.45 | 0.55 |
| CellPLM | 0.76 | 0.80 | 0.46 | 0.53 | 0.45 | 0.55 |
| SingleR | 0.76 | n/a | 0.49 | 0.50 | 0.45 | 0.55 |
| LangCell | 0.77 | 0.81 | 0.51 | 0.49 | 0.44 | 0.55 |
| PCA | 0.78 | 0.82 | 0.40 | 0.49 | 0.43 | 0.52 |
| CellTypist | 0.77 | 0.81 | 0.41 | 0.48 | 0.42 | 0.52 |
| scVI | 0.76 | 0.81 | 0.44 | 0.46 | 0.41 | 0.52 |
| SCimilarity | 0.75 | 0.81 | 0.39 | 0.53 | 0.40 | 0.52 |
| GenePT-w | 0.60 | 0.79 | 0.38 | 0.48 | 0.36 | 0.46 |
| CellFM | 0.73 | 0.80 | 0.40 | 0.37 | 0.31 | 0.45 |
| CELLama | 0.53 | 0.71 | 0.32 | 0.37 | 0.27 | 0.37 |
| scBERT | 0.43 | 0.76 | 0.23 | 0.22 | 0.17 | 0.26 |

**b** Heart · reference: v1, query: v2 · shared

| method | Accuracy | $Prec_{Macro}$ | $Recall_{Macro}$ | $F1_{Macro}$ | $AUC_{ROC}$ | $Acc_{top3}$ | Annotation |
|---|---|---|---|---|---|---|---|
| | Annotation metrics | | | | | | Aggregated |
| scGPT | 0.94 | 0.63 | 0.59 | 0.57 | 0.97 | 0.99 | 0.68 |
| scFoundation | 0.94 | 0.58 | 0.61 | 0.58 | 0.97 | 0.97 | 0.68 |
| Geneformer | 0.92 | 0.63 | 0.55 | 0.54 | 0.98 | 0.97 | 0.66 |
| scPRINT | 0.91 | 0.53 | 0.57 | 0.53 | 0.95 | 0.97 | 0.63 |
| UCE | 0.89 | 0.52 | 0.59 | 0.51 | 0.91 | 0.97 | 0.63 |
| scANVI | 0.91 | 0.52 | 0.54 | 0.50 | 0.94 | 0.98 | 0.62 |
| CellPLM | 0.92 | 0.53 | 0.53 | 0.49 | 0.97 | 0.96 | 0.62 |
| scCello | 0.90 | 0.58 | 0.48 | 0.50 | 0.93 | 0.97 | 0.61 |
| SingleR | 0.91 | 0.55 | 0.50 | 0.49 | n/a | n/a | 0.61 |
| LangCell | 0.92 | 0.56 | 0.49 | 0.48 | 0.96 | 0.97 | 0.61 |
| PCA | 0.93 | 0.46 | 0.49 | 0.47 | 0.94 | 0.98 | 0.59 |
| scVI | 0.91 | 0.51 | 0.46 | 0.45 | 0.96 | 0.98 | 0.58 |
| CellTypist | 0.92 | 0.47 | 0.48 | 0.46 | 0.98 | 0.97 | 0.58 |
| SCimilarity | 0.89 | 0.44 | 0.53 | 0.44 | 0.98 | 0.97 | 0.58 |
| CellFM | 0.88 | 0.45 | 0.37 | 0.34 | 0.92 | 0.96 | 0.51 |
| GenePT-w | 0.72 | 0.44 | 0.48 | 0.39 | 0.95 | 0.95 | 0.51 |
| CELLama | 0.64 | 0.34 | 0.37 | 0.29 | 0.83 | 0.85 | 0.41 |
| scBERT | 0.52 | 0.26 | 0.22 | 0.19 | 0.77 | 0.91 | 0.30 |

**Figure S195: Directional cross-dataset annotation for heart, TSv1 to TSv2. a,** All 13,344 target cells, including 2,209 cells from source-absent labels. **b,** 11,135 target cells from 14 shared labels. Methods and metrics are as in Fig. S185.

**a** Heart · reference: v2, query: v1 · open-world

| method | Annotation metrics: *Accuracy* | *Acc*$_{top3}$ | *Prec*$_{Macro}$ | *Recall*$_{Macro}$ | *F1*$_{Macro}$ | Aggregated: *Annotation* |
|---|---|---|---|---|---|---|
| CellTypist | 0.73 | 0.95 | 0.55 | 0.60 | 0.55 | 0.61 |
| SingleR | 0.76 | n/a | 0.50 | 0.62 | 0.50 | 0.59 |
| scGPT | 0.57 | 0.89 | 0.51 | 0.69 | 0.53 | 0.58 |
| Geneformer | 0.66 | 0.83 | 0.46 | 0.70 | 0.45 | 0.57 |
| scVI | 0.73 | 0.98 | 0.41 | 0.62 | 0.43 | 0.55 |
| scCello | 0.57 | 0.89 | 0.43 | 0.62 | 0.46 | 0.52 |
| PCA | 0.59 | 0.90 | 0.44 | 0.56 | 0.47 | 0.52 |
| SCimilarity | 0.44 | 0.94 | 0.54 | 0.56 | 0.51 | 0.51 |
| scFoundation | 0.49 | 0.88 | 0.46 | 0.61 | 0.44 | 0.50 |
| LangCell | 0.57 | 0.94 | 0.41 | 0.62 | 0.40 | 0.50 |
| scPRINT | 0.39 | 0.81 | 0.43 | 0.65 | 0.39 | 0.47 |
| CellPLM | 0.44 | 0.95 | 0.45 | 0.54 | 0.40 | 0.46 |
| UCE | 0.32 | 0.82 | 0.45 | 0.51 | 0.42 | 0.42 |
| scANVI | 0.47 | 0.67 | 0.28 | 0.51 | 0.28 | 0.39 |
| CellFM | 0.42 | 0.70 | 0.31 | 0.48 | 0.31 | 0.38 |
| CELLama | 0.39 | 0.68 | 0.27 | 0.33 | 0.21 | 0.30 |
| GenePT-w | 0.35 | 0.59 | 0.31 | 0.21 | 0.19 | 0.27 |
| scBERT | 0.25 | 0.58 | 0.28 | 0.30 | 0.21 | 0.26 |

**b** Heart · reference: v2, query: v1 · shared

| method | Annotation metrics: *Accuracy* | *Prec*$_{Macro}$ | *Recall*$_{Macro}$ | *F1*$_{Macro}$ | *AUC*$_{ROC}$ | *Acc*$_{top3}$ | Aggregated: *Annotation* |
|---|---|---|---|---|---|---|---|
| CellTypist | 0.73 | 0.55 | 0.60 | 0.55 | 0.92 | 0.95 | 0.61 |
| SingleR | 0.76 | 0.50 | 0.62 | 0.50 | n/a | n/a | 0.59 |
| scGPT | 0.57 | 0.51 | 0.69 | 0.53 | 0.93 | 0.89 | 0.58 |
| Geneformer | 0.66 | 0.46 | 0.70 | 0.45 | 0.90 | 0.83 | 0.57 |
| scVI | 0.73 | 0.41 | 0.62 | 0.43 | 0.92 | 0.98 | 0.55 |
| scCello | 0.57 | 0.43 | 0.62 | 0.46 | 0.90 | 0.89 | 0.52 |
| PCA | 0.59 | 0.44 | 0.56 | 0.47 | 0.89 | 0.90 | 0.52 |
| SCimilarity | 0.44 | 0.54 | 0.56 | 0.51 | 0.88 | 0.94 | 0.51 |
| scFoundation | 0.49 | 0.46 | 0.61 | 0.44 | 0.85 | 0.88 | 0.50 |
| LangCell | 0.57 | 0.41 | 0.62 | 0.40 | 0.89 | 0.94 | 0.50 |
| scPRINT | 0.39 | 0.43 | 0.65 | 0.39 | 0.83 | 0.81 | 0.47 |
| CellPLM | 0.44 | 0.45 | 0.54 | 0.40 | 0.88 | 0.95 | 0.46 |
| UCE | 0.32 | 0.45 | 0.51 | 0.42 | 0.90 | 0.82 | 0.42 |
| scANVI | 0.47 | 0.28 | 0.51 | 0.28 | 0.86 | 0.67 | 0.39 |
| CellFM | 0.42 | 0.31 | 0.48 | 0.31 | 0.93 | 0.70 | 0.38 |
| CELLama | 0.39 | 0.27 | 0.33 | 0.21 | 0.80 | 0.68 | 0.30 |
| GenePT-w | 0.35 | 0.31 | 0.21 | 0.19 | 0.85 | 0.59 | 0.27 |
| scBERT | 0.25 | 0.28 | 0.30 | 0.21 | 0.81 | 0.58 | 0.26 |

**Figure S196: Directional cross-dataset annotation for heart, TSv2 to TSv1. a,b,** Both panels contain 10,581 target cells from 14 shared labels because no target label is absent from the source in this direction. Methods and metrics are as in Fig. S185.

**a** Large Intestine · reference: v1, query: v2 · open-world

| method | Annotation metrics: *Accuracy* | $Acc_{top3}$ | $Prec_{Macro}$ | $Recall_{Macro}$ | $F1_{Macro}$ | Aggregated: *Annotation* |
|---|---|---|---|---|---|---|
| SCimilarity | 0.71 | 0.87 | 0.60 | 0.68 | 0.60 | 0.65 |
| PCA | 0.72 | 0.86 | 0.57 | 0.65 | 0.57 | 0.63 |
| scANVI | 0.72 | 0.86 | 0.58 | 0.63 | 0.58 | 0.63 |
| CellTypist | 0.72 | 0.84 | 0.62 | 0.61 | 0.56 | 0.63 |
| UCE | 0.66 | 0.82 | 0.60 | 0.67 | 0.56 | 0.62 |
| scFoundation | 0.66 | 0.83 | 0.58 | 0.65 | 0.56 | 0.61 |
| scGPT | 0.69 | 0.85 | 0.56 | 0.65 | 0.55 | 0.61 |
| Geneformer | 0.66 | 0.82 | 0.59 | 0.63 | 0.56 | 0.61 |
| scCello | 0.69 | 0.82 | 0.54 | 0.64 | 0.55 | 0.60 |
| CellPLM | 0.63 | 0.84 | 0.60 | 0.61 | 0.52 | 0.59 |
| scPRINT | 0.66 | 0.85 | 0.57 | 0.55 | 0.50 | 0.57 |
| SingleR | 0.65 | n/a | 0.51 | 0.63 | 0.51 | 0.57 |
| LangCell | 0.66 | 0.85 | 0.54 | 0.57 | 0.52 | 0.57 |
| CellFM | 0.64 | 0.83 | 0.53 | 0.52 | 0.47 | 0.54 |
| scVI | 0.60 | 0.81 | 0.45 | 0.53 | 0.44 | 0.51 |
| GenePT-w | 0.55 | 0.69 | 0.39 | 0.46 | 0.34 | 0.44 |
| CELLama | 0.45 | 0.63 | 0.32 | 0.35 | 0.28 | 0.35 |
| scBERT | 0.23 | 0.42 | 0.12 | 0.16 | 0.10 | 0.15 |

**b** Large Intestine · reference: v1, query: v2 · shared

| method | Annotation metrics: *Accuracy* | $Prec_{Macro}$ | $Recall_{Macro}$ | $F1_{Macro}$ | $AUC_{ROC}$ | $Acc_{top3}$ | Aggregated: *Annotation* |
|---|---|---|---|---|---|---|---|
| SCimilarity | 0.77 | 0.62 | 0.68 | 0.61 | 0.98 | 0.94 | 0.67 |
| PCA | 0.78 | 0.59 | 0.65 | 0.59 | 0.98 | 0.92 | 0.65 |
| scANVI | 0.78 | 0.59 | 0.63 | 0.59 | 0.98 | 0.93 | 0.65 |
| UCE | 0.72 | 0.61 | 0.67 | 0.58 | 0.98 | 0.89 | 0.64 |
| CellTypist | 0.77 | 0.63 | 0.61 | 0.57 | 0.98 | 0.91 | 0.64 |
| scFoundation | 0.72 | 0.61 | 0.65 | 0.58 | 0.98 | 0.90 | 0.64 |
| scGPT | 0.74 | 0.58 | 0.65 | 0.57 | 0.98 | 0.92 | 0.63 |
| scCello | 0.75 | 0.57 | 0.64 | 0.57 | 0.98 | 0.88 | 0.63 |
| Geneformer | 0.71 | 0.60 | 0.63 | 0.57 | 0.97 | 0.88 | 0.63 |
| CellPLM | 0.68 | 0.61 | 0.61 | 0.53 | 0.97 | 0.91 | 0.61 |
| scPRINT | 0.72 | 0.58 | 0.55 | 0.51 | 0.97 | 0.92 | 0.59 |
| LangCell | 0.71 | 0.55 | 0.57 | 0.53 | 0.96 | 0.92 | 0.59 |
| SingleR | 0.70 | 0.51 | 0.63 | 0.51 | n/a | n/a | 0.59 |
| CellFM | 0.69 | 0.54 | 0.52 | 0.47 | 0.96 | 0.90 | 0.56 |
| scVI | 0.65 | 0.47 | 0.53 | 0.45 | 0.97 | 0.87 | 0.53 |
| GenePT-w | 0.59 | 0.44 | 0.46 | 0.37 | 0.92 | 0.74 | 0.47 |
| CELLama | 0.49 | 0.33 | 0.35 | 0.29 | 0.87 | 0.68 | 0.36 |
| scBERT | 0.25 | 0.12 | 0.16 | 0.11 | 0.67 | 0.46 | 0.16 |

**Figure S197: Directional cross-dataset annotation for large intestine, TSv1 to TSv2. a,** All 15,231 target cells, including 1,127 cells from source-absent labels. **b,** 14,104 target cells from 24 shared labels. Methods and metrics are as in Fig. S185.

**a** Large Intestine · reference: v2, query: v1 · open-world

| method | Annotation metrics: *Accuracy* | *Acc*$_{top3}$ | *Prec*$_{Macro}$ | *Recall*$_{Macro}$ | *F1*$_{Macro}$ | Aggregated: *Annotation* |
|---|---|---|---|---|---|---|
| UCE | 0.82 | 0.93 | 0.64 | 0.71 | 0.61 | 0.69 |
| PCA | 0.85 | 0.97 | 0.63 | 0.66 | 0.61 | 0.69 |
| SCimilarity | 0.84 | 0.95 | 0.61 | 0.68 | 0.59 | 0.68 |
| scFoundation | 0.83 | 0.95 | 0.62 | 0.63 | 0.59 | 0.67 |
| CellTypist | 0.86 | 0.97 | 0.61 | 0.62 | 0.58 | 0.67 |
| SingleR | 0.77 | n/a | 0.62 | 0.67 | 0.57 | 0.66 |
| Geneformer | 0.81 | 0.92 | 0.63 | 0.63 | 0.56 | 0.66 |
| scCello | 0.82 | 0.96 | 0.55 | 0.59 | 0.55 | 0.63 |
| scGPT | 0.76 | 0.93 | 0.58 | 0.59 | 0.54 | 0.62 |
| scPRINT | 0.76 | 0.94 | 0.57 | 0.53 | 0.51 | 0.59 |
| LangCell | 0.73 | 0.91 | 0.54 | 0.58 | 0.51 | 0.59 |
| scVI | 0.72 | 0.90 | 0.50 | 0.59 | 0.51 | 0.58 |
| scANVI | 0.75 | 0.90 | 0.47 | 0.60 | 0.49 | 0.58 |
| CellFM | 0.71 | 0.93 | 0.54 | 0.56 | 0.49 | 0.57 |
| CellPLM | 0.58 | 0.91 | 0.50 | 0.52 | 0.44 | 0.51 |
| GenePT-w | 0.57 | 0.76 | 0.46 | 0.37 | 0.34 | 0.44 |
| CELLama | 0.40 | 0.68 | 0.34 | 0.37 | 0.31 | 0.35 |
| scBERT | 0.23 | 0.46 | 0.25 | 0.25 | 0.18 | 0.23 |

**b** Large Intestine · reference: v2, query: v1 · shared

| method | Annotation metrics: *Accuracy* | *Prec*$_{Macro}$ | *Recall*$_{Macro}$ | *F1*$_{Macro}$ | *AUC*$_{ROC}$ | *Acc*$_{top3}$ | Aggregated: *Annotation* |
|---|---|---|---|---|---|---|---|
| UCE | 0.82 | 0.65 | 0.71 | 0.61 | 0.96 | 0.94 | 0.70 |
| PCA | 0.85 | 0.65 | 0.66 | 0.62 | 0.97 | 0.97 | 0.69 |
| SCimilarity | 0.84 | 0.62 | 0.68 | 0.60 | 0.97 | 0.96 | 0.68 |
| CellTypist | 0.86 | 0.63 | 0.62 | 0.59 | 0.97 | 0.97 | 0.67 |
| scFoundation | 0.83 | 0.63 | 0.63 | 0.60 | 0.95 | 0.95 | 0.67 |
| SingleR | 0.77 | 0.63 | 0.67 | 0.58 | n/a | n/a | 0.66 |
| Geneformer | 0.81 | 0.63 | 0.63 | 0.56 | 0.98 | 0.92 | 0.66 |
| scCello | 0.82 | 0.57 | 0.59 | 0.55 | 0.96 | 0.96 | 0.63 |
| scGPT | 0.76 | 0.59 | 0.59 | 0.55 | 0.96 | 0.93 | 0.62 |
| scPRINT | 0.76 | 0.57 | 0.53 | 0.51 | 0.93 | 0.94 | 0.59 |
| LangCell | 0.73 | 0.54 | 0.58 | 0.51 | 0.95 | 0.91 | 0.59 |
| scVI | 0.72 | 0.52 | 0.59 | 0.52 | 0.93 | 0.90 | 0.59 |
| scANVI | 0.75 | 0.47 | 0.60 | 0.49 | 0.92 | 0.90 | 0.58 |
| CellFM | 0.71 | 0.55 | 0.56 | 0.50 | 0.96 | 0.93 | 0.58 |
| CellPLM | 0.58 | 0.51 | 0.52 | 0.44 | 0.90 | 0.91 | 0.51 |
| GenePT-w | 0.57 | 0.46 | 0.37 | 0.34 | 0.90 | 0.76 | 0.44 |
| CELLama | 0.40 | 0.34 | 0.37 | 0.31 | 0.86 | 0.68 | 0.35 |
| scBERT | 0.23 | 0.25 | 0.25 | 0.18 | 0.79 | 0.46 | 0.23 |

**Figure S198: Directional cross-dataset annotation for large intestine, TSv2 to TSv1. a,** All 7,655 target cells, including 4 cells from source-absent labels. **b,** 7,651 target cells from 24 shared labels. Methods and metrics are as in Fig. S185.

**a** Liver · reference: v1, query: v2 · open-world

| method | Annotation metrics | | | | | Aggregated |
|---|---|---|---|---|---|---|
| | *Accuracy* | $Acc_{top3}$ | $Prec_{Macro}$ | $Recall_{Macro}$ | $F1_{Macro}$ | *Annotation* |
| UCE | 0.78 | 0.93 | 0.58 | 0.63 | 0.57 | 0.64 |
| SCimilarity | 0.79 | 0.94 | 0.59 | 0.62 | 0.56 | 0.64 |
| scCello | 0.74 | 0.91 | 0.58 | 0.61 | 0.52 | 0.61 |
| scANVI | 0.73 | 0.92 | 0.57 | 0.60 | 0.53 | 0.61 |
| CellTypist | 0.77 | 0.92 | 0.55 | 0.59 | 0.51 | 0.61 |
| scPRINT | 0.76 | 0.91 | 0.56 | 0.60 | 0.51 | 0.61 |
| scGPT | 0.76 | 0.91 | 0.55 | 0.58 | 0.52 | 0.60 |
| PCA | 0.77 | 0.93 | 0.59 | 0.55 | 0.51 | 0.60 |
| scVI | 0.77 | 0.94 | 0.55 | 0.58 | 0.50 | 0.60 |
| Geneformer | 0.76 | 0.91 | 0.55 | 0.58 | 0.50 | 0.60 |
| scFoundation | 0.75 | 0.90 | 0.53 | 0.58 | 0.51 | 0.60 |
| SingleR | 0.74 | n/a | 0.53 | 0.60 | 0.48 | 0.59 |
| CellPLM | 0.73 | 0.90 | 0.54 | 0.54 | 0.48 | 0.57 |
| LangCell | 0.77 | 0.91 | 0.50 | 0.53 | 0.47 | 0.57 |
| CellFM | 0.73 | 0.91 | 0.48 | 0.53 | 0.46 | 0.55 |
| GenePT-w | 0.62 | 0.84 | 0.45 | 0.43 | 0.34 | 0.46 |
| CELLama | 0.57 | 0.78 | 0.38 | 0.41 | 0.34 | 0.42 |
| scBERT | 0.41 | 0.67 | 0.27 | 0.22 | 0.21 | 0.28 |

**b** Liver · reference: v1, query: v2 · shared

| method | Annotation metrics | | | | | | Aggregated |
|---|---|---|---|---|---|---|---|
| | *Accuracy* | $Prec_{Macro}$ | $Recall_{Macro}$ | $F1_{Macro}$ | $AUC_{ROC}$ | $Acc_{top3}$ | *Annotation* |
| UCE | 0.80 | 0.61 | 0.63 | 0.60 | 0.97 | 0.95 | 0.66 |
| SCimilarity | 0.81 | 0.62 | 0.62 | 0.59 | 0.98 | 0.96 | 0.66 |
| scCello | 0.75 | 0.62 | 0.61 | 0.55 | 0.96 | 0.93 | 0.63 |
| scANVI | 0.75 | 0.60 | 0.60 | 0.56 | 0.96 | 0.94 | 0.63 |
| scPRINT | 0.77 | 0.59 | 0.60 | 0.55 | 0.97 | 0.93 | 0.63 |
| CellTypist | 0.79 | 0.59 | 0.59 | 0.54 | 0.94 | 0.94 | 0.63 |
| PCA | 0.79 | 0.62 | 0.55 | 0.54 | 0.95 | 0.95 | 0.63 |
| scGPT | 0.78 | 0.58 | 0.58 | 0.56 | 0.97 | 0.92 | 0.62 |
| scVI | 0.79 | 0.58 | 0.58 | 0.54 | 0.95 | 0.96 | 0.62 |
| Geneformer | 0.77 | 0.59 | 0.58 | 0.54 | 0.96 | 0.93 | 0.62 |
| scFoundation | 0.77 | 0.56 | 0.58 | 0.55 | 0.96 | 0.92 | 0.62 |
| SingleR | 0.75 | 0.55 | 0.60 | 0.50 | n/a | n/a | 0.60 |
| CellPLM | 0.74 | 0.57 | 0.54 | 0.52 | 0.96 | 0.91 | 0.59 |
| LangCell | 0.79 | 0.54 | 0.53 | 0.51 | 0.95 | 0.93 | 0.59 |
| CellFM | 0.75 | 0.50 | 0.53 | 0.49 | 0.95 | 0.93 | 0.57 |
| GenePT-w | 0.64 | 0.47 | 0.43 | 0.36 | 0.92 | 0.85 | 0.47 |
| CELLama | 0.58 | 0.39 | 0.41 | 0.35 | 0.90 | 0.80 | 0.43 |

**Figure S199: Directional cross-dataset annotation for liver, TSv1 to TSv2. a,** All 10,002 target cells, including 209 cells from source-absent labels. **b,** 9,793 target cells from 22 shared labels. Methods and metrics are as in Fig. S185.

**a** Liver · reference: v2, query: v1 · open-world

| method | Accuracy | $Acc_{top3}$ | $Prec_{Macro}$ | $Recall_{Macro}$ | $F1_{Macro}$ | Aggregated Annotation |
|---|---|---|---|---|---|---|
| | Annotation metrics | | | | | |
| SCimilarity | 0.74 | 0.94 | 0.61 | 0.64 | 0.57 | 0.64 |
| scGPT | 0.71 | 0.95 | 0.60 | 0.64 | 0.57 | 0.63 |
| UCE | 0.73 | 0.93 | 0.58 | 0.62 | 0.56 | 0.62 |
| scFoundation | 0.71 | 0.93 | 0.58 | 0.62 | 0.55 | 0.62 |
| CellTypist | 0.72 | 0.93 | 0.59 | 0.61 | 0.53 | 0.61 |
| scCello | 0.73 | 0.93 | 0.57 | 0.59 | 0.54 | 0.61 |
| SingleR | 0.69 | n/a | 0.61 | 0.59 | 0.54 | 0.61 |
| PCA | 0.69 | 0.93 | 0.60 | 0.59 | 0.54 | 0.60 |
| scVI | 0.70 | 0.92 | 0.57 | 0.61 | 0.53 | 0.60 |
| Geneformer | 0.70 | 0.93 | 0.57 | 0.59 | 0.53 | 0.60 |
| CellPLM | 0.73 | 0.93 | 0.53 | 0.59 | 0.51 | 0.59 |
| LangCell | 0.69 | 0.91 | 0.56 | 0.57 | 0.52 | 0.58 |
| scPRINT | 0.64 | 0.90 | 0.58 | 0.53 | 0.49 | 0.56 |
| CellFM | 0.66 | 0.89 | 0.48 | 0.50 | 0.47 | 0.53 |
| scANVI | 0.63 | 0.86 | 0.45 | 0.56 | 0.43 | 0.51 |
| GenePT-w | 0.65 | 0.83 | 0.38 | 0.47 | 0.33 | 0.46 |
| CELLama | 0.36 | 0.66 | 0.28 | 0.40 | 0.26 | 0.33 |
| scBERT | 0.37 | 0.68 | 0.25 | 0.30 | 0.22 | 0.28 |

**b** Liver · reference: v2, query: v1 · shared

| method | Accuracy | $Prec_{Macro}$ | $Recall_{Macro}$ | $F1_{Macro}$ | $AUC_{ROC}$ | $Acc_{top3}$ | Aggregated Annotation |
|---|---|---|---|---|---|---|---|
| | Annotation metrics | | | | | | |
| SCimilarity | 0.75 | 0.62 | 0.64 | 0.57 | 0.98 | 0.96 | 0.65 |
| scGPT | 0.73 | 0.61 | 0.64 | 0.57 | 0.98 | 0.97 | 0.63 |
| UCE | 0.74 | 0.58 | 0.62 | 0.56 | 0.94 | 0.94 | 0.62 |
| scFoundation | 0.72 | 0.58 | 0.62 | 0.56 | 0.96 | 0.95 | 0.62 |
| CellTypist | 0.73 | 0.59 | 0.61 | 0.53 | 0.94 | 0.95 | 0.62 |
| scCello | 0.74 | 0.58 | 0.59 | 0.54 | 0.96 | 0.95 | 0.61 |
| SingleR | 0.70 | 0.61 | 0.59 | 0.54 | n/a | n/a | 0.61 |
| scVI | 0.72 | 0.58 | 0.61 | 0.53 | 0.97 | 0.93 | 0.61 |
| PCA | 0.70 | 0.60 | 0.59 | 0.54 | 0.96 | 0.94 | 0.61 |
| CellPLM | 0.74 | 0.54 | 0.59 | 0.52 | 0.97 | 0.95 | 0.60 |
| Geneformer | 0.71 | 0.57 | 0.59 | 0.53 | 0.95 | 0.94 | 0.60 |
| LangCell | 0.70 | 0.56 | 0.57 | 0.52 | 0.94 | 0.93 | 0.59 |
| scPRINT | 0.65 | 0.58 | 0.53 | 0.49 | 0.95 | 0.92 | 0.56 |
| CellFM | 0.67 | 0.48 | 0.50 | 0.47 | 0.94 | 0.91 | 0.53 |
| scANVI | 0.64 | 0.46 | 0.56 | 0.43 | 0.92 | 0.87 | 0.52 |
| GenePT-w | 0.66 | 0.39 | 0.47 | 0.33 | 0.94 | 0.85 | 0.46 |
| CELLama | 0.36 | 0.29 | 0.40 | 0.27 | 0.86 | 0.67 | 0.33 |
| scBERT | 0.38 | 0.26 | 0.30 | 0.22 | 0.84 | 0.70 | 0.29 |

**Figure S200: Directional cross-dataset annotation for liver, TSv2 to TSv1. a,** All 4,665 target cells, including 75 cells from source-absent labels. **b,** 4,590 target cells from 22 shared labels. Methods and metrics are as in Fig. S185.

**a** **Lung · reference: v1, query: v2 · open-world**

| method | Annotation metrics | | | | | Aggregated |
|---|---|---|---|---|---|---|
| | *Accuracy* | *Acc*$_{top3}$ | *Prec*$_{Macro}$ | *Recall*$_{Macro}$ | *F1*$_{Macro}$ | *Annotation* |
| **SCimilarity** | 0.90 | 0.98 | 0.77 | 0.78 | 0.76 | 0.80 |
| **CellTypist** | 0.90 | 0.98 | 0.74 | 0.76 | 0.73 | 0.78 |
| **SingleR** | 0.88 | n/a | 0.74 | 0.77 | 0.73 | 0.78 |
| **scCello** | 0.88 | 0.98 | 0.72 | 0.76 | 0.73 | 0.77 |
| **scFoundation** | 0.84 | 0.97 | 0.72 | 0.76 | 0.72 | 0.76 |
| **Geneformer** | 0.87 | 0.98 | 0.71 | 0.73 | 0.71 | 0.76 |
| **PCA** | 0.86 | 0.97 | 0.71 | 0.75 | 0.71 | 0.76 |
| **scANVI** | 0.84 | 0.97 | 0.68 | 0.75 | 0.69 | 0.74 |
| **UCE** | 0.83 | 0.97 | 0.71 | 0.72 | 0.69 | 0.74 |
| **scVI** | 0.85 | 0.97 | 0.69 | 0.71 | 0.68 | 0.73 |
| **scGPT** | 0.84 | 0.97 | 0.69 | 0.72 | 0.69 | 0.73 |
| **scPRINT** | 0.85 | 0.97 | 0.68 | 0.69 | 0.68 | 0.73 |
| **LangCell** | 0.82 | 0.97 | 0.68 | 0.70 | 0.68 | 0.72 |
| **CellFM** | 0.84 | 0.98 | 0.69 | 0.68 | 0.67 | 0.72 |
| **CellPLM** | 0.85 | 0.97 | 0.67 | 0.70 | 0.65 | 0.72 |
| **GenePT-w** | 0.72 | 0.94 | 0.58 | 0.62 | 0.55 | 0.62 |
| **CELLama** | 0.65 | 0.87 | 0.45 | 0.50 | 0.45 | 0.52 |
| **scBERT** | 0.52 | 0.75 | 0.34 | 0.37 | 0.31 | 0.38 |

**b** **Lung · reference: v1, query: v2 · shared**

| method | Annotation metrics | | | | | | Aggregated |
|---|---|---|---|---|---|---|---|
| | *Accuracy* | *Prec*$_{Macro}$ | *Recall*$_{Macro}$ | *F1*$_{Macro}$ | *AUC*$_{ROC}$ | *Acc*$_{top3}$ | *Annotation* |
| **SCimilarity** | 0.90 | 0.77 | 0.78 | 0.76 | 0.99 | 0.98 | 0.80 |
| **CellTypist** | 0.90 | 0.74 | 0.76 | 0.73 | 0.99 | 0.98 | 0.78 |
| **SingleR** | 0.88 | 0.74 | 0.77 | 0.73 | n/a | n/a | 0.78 |
| **scCello** | 0.88 | 0.72 | 0.76 | 0.73 | 0.99 | 0.98 | 0.77 |
| **scFoundation** | 0.84 | 0.72 | 0.76 | 0.72 | 0.99 | 0.97 | 0.76 |
| **Geneformer** | 0.87 | 0.71 | 0.73 | 0.71 | 0.99 | 0.98 | 0.76 |
| **PCA** | 0.86 | 0.71 | 0.75 | 0.71 | 0.99 | 0.97 | 0.76 |
| **scANVI** | 0.84 | 0.68 | 0.75 | 0.69 | 0.99 | 0.97 | 0.74 |
| **UCE** | 0.83 | 0.71 | 0.72 | 0.69 | 0.99 | 0.97 | 0.74 |
| **scVI** | 0.85 | 0.69 | 0.71 | 0.68 | 0.99 | 0.97 | 0.73 |
| **scGPT** | 0.84 | 0.69 | 0.72 | 0.69 | 0.99 | 0.97 | 0.73 |
| **scPRINT** | 0.85 | 0.68 | 0.69 | 0.68 | 0.99 | 0.97 | 0.73 |
| **LangCell** | 0.82 | 0.68 | 0.70 | 0.68 | 0.98 | 0.97 | 0.72 |
| **CellFM** | 0.84 | 0.69 | 0.68 | 0.67 | 0.99 | 0.98 | 0.72 |
| **CellPLM** | 0.85 | 0.67 | 0.70 | 0.65 | 0.99 | 0.97 | 0.72 |
| **GenePT-w** | 0.72 | 0.58 | 0.62 | 0.55 | 0.98 | 0.94 | 0.62 |
| **CELLama** | 0.65 | 0.45 | 0.50 | 0.45 | 0.95 | 0.87 | 0.52 |
| **scBERT** | 0.52 | 0.34 | 0.37 | 0.31 | 0.86 | 0.75 | 0.38 |

**Figure S201: Directional cross-dataset annotation for lung, TSv1 to TSv2. a,b,** Both panels contain 11,716 target cells from 31 shared labels because no target label is absent from the source in this direction. Methods and metrics are as in Fig. S185.

**a** Lung · reference: v2, query: v1 · open-world

| method | Annotation metrics | | | | | Aggregated |
|---|---|---|---|---|---|---|
| | *Accuracy* | *Acc*$_{top3}$ | *Prec*$_{Macro}$ | *Recall*$_{Macro}$ | *F1*$_{Macro}$ | *Annotation* |
| CellTypist | 0.91 | 0.98 | 0.76 | 0.77 | 0.74 | 0.80 |
| scFoundation | 0.90 | 0.97 | 0.73 | 0.75 | 0.73 | 0.78 |
| PCA | 0.90 | 0.98 | 0.73 | 0.75 | 0.73 | 0.78 |
| SCimilarity | 0.89 | 0.96 | 0.72 | 0.76 | 0.73 | 0.78 |
| scCello | 0.89 | 0.97 | 0.73 | 0.75 | 0.73 | 0.77 |
| scGPT | 0.89 | 0.97 | 0.73 | 0.73 | 0.72 | 0.77 |
| SingleR | 0.89 | n/a | 0.73 | 0.73 | 0.69 | 0.76 |
| Geneformer | 0.89 | 0.96 | 0.74 | 0.71 | 0.70 | 0.76 |
| UCE | 0.87 | 0.97 | 0.69 | 0.74 | 0.70 | 0.75 |
| scANVI | 0.88 | 0.97 | 0.68 | 0.70 | 0.66 | 0.73 |
| scVI | 0.88 | 0.97 | 0.68 | 0.68 | 0.66 | 0.73 |
| CellPLM | 0.86 | 0.96 | 0.65 | 0.67 | 0.64 | 0.71 |
| LangCell | 0.86 | 0.95 | 0.66 | 0.65 | 0.63 | 0.70 |
| scPRINT | 0.86 | 0.95 | 0.66 | 0.63 | 0.61 | 0.69 |
| CellFM | 0.86 | 0.97 | 0.62 | 0.65 | 0.61 | 0.68 |
| CELLama | 0.81 | 0.93 | 0.58 | 0.55 | 0.52 | 0.62 |
| GenePT-w | 0.75 | 0.88 | 0.60 | 0.54 | 0.47 | 0.59 |
| scBERT | 0.63 | 0.88 | 0.32 | 0.36 | 0.31 | 0.40 |

**b** Lung · reference: v2, query: v1 · shared

| method | Annotation metrics | | | | | | Aggregated |
|---|---|---|---|---|---|---|---|
| | *Accuracy* | *Prec*$_{Macro}$ | *Recall*$_{Macro}$ | *F1*$_{Macro}$ | *AUC*$_{ROC}$ | *Acc*$_{top3}$ | *Annotation* |
| CellTypist | 0.92 | 0.76 | 0.77 | 0.75 | 0.98 | 0.98 | 0.80 |
| PCA | 0.90 | 0.74 | 0.75 | 0.73 | 0.99 | 0.98 | 0.78 |
| scFoundation | 0.90 | 0.73 | 0.75 | 0.73 | 0.99 | 0.97 | 0.78 |
| SCimilarity | 0.89 | 0.72 | 0.76 | 0.73 | 0.98 | 0.97 | 0.78 |
| scCello | 0.89 | 0.73 | 0.75 | 0.73 | 0.99 | 0.97 | 0.77 |
| scGPT | 0.89 | 0.74 | 0.73 | 0.72 | 0.99 | 0.97 | 0.77 |
| SingleR | 0.89 | 0.74 | 0.73 | 0.70 | n/a | n/a | 0.76 |
| Geneformer | 0.89 | 0.75 | 0.71 | 0.70 | 0.98 | 0.97 | 0.76 |
| UCE | 0.88 | 0.69 | 0.74 | 0.70 | 0.99 | 0.97 | 0.75 |
| scANVI | 0.88 | 0.68 | 0.70 | 0.67 | 0.98 | 0.97 | 0.73 |
| scVI | 0.88 | 0.69 | 0.68 | 0.66 | 0.99 | 0.97 | 0.73 |
| CellPLM | 0.87 | 0.66 | 0.67 | 0.64 | 0.99 | 0.97 | 0.71 |
| LangCell | 0.86 | 0.66 | 0.65 | 0.63 | 0.99 | 0.95 | 0.70 |
| scPRINT | 0.86 | 0.67 | 0.63 | 0.61 | 0.98 | 0.95 | 0.69 |
| CellFM | 0.86 | 0.62 | 0.65 | 0.61 | 0.99 | 0.97 | 0.69 |
| CELLama | 0.82 | 0.58 | 0.55 | 0.52 | 0.96 | 0.93 | 0.62 |
| GenePT-w | 0.75 | 0.60 | 0.54 | 0.47 | 0.98 | 0.89 | 0.59 |
| scBERT | 0.63 | 0.32 | 0.36 | 0.31 | 0.87 | 0.88 | 0.40 |

**Figure S202: Directional cross-dataset annotation for lung, TSv2 to TSv1.** **a,** All 34,400 target cells, including 58 cells from source-absent labels. **b,** 34,342 target cells from 31 shared labels. Methods and metrics are as in Fig. S185.

**a** Lymph Node · reference: v1, query: v2 · open-world

| | Annotation metrics | | | | | Aggregated |
|---|---|---|---|---|---|---|
| method | *Accuracy* | $Acc_{top3}$ | $Prec_{Macro}$ | $Recall_{Macro}$ | $F1_{Macro}$ | *Annotation* |
| SingleR | 0.76 | n/a | 0.57 | 0.69 | 0.55 | 0.64 |
| scFoundation | 0.82 | 0.98 | 0.52 | 0.68 | 0.53 | 0.64 |
| SCimilarity | 0.81 | 0.99 | 0.54 | 0.67 | 0.54 | 0.64 |
| CellTypist | 0.77 | 0.96 | 0.58 | 0.65 | 0.55 | 0.64 |
| scPRINT | 0.81 | 0.99 | 0.54 | 0.65 | 0.54 | 0.63 |
| scGPT | 0.79 | 0.98 | 0.54 | 0.66 | 0.52 | 0.63 |
| CellPLM | 0.82 | 0.98 | 0.53 | 0.64 | 0.52 | 0.63 |
| scANVI | 0.82 | 0.97 | 0.50 | 0.65 | 0.51 | 0.62 |
| scVI | 0.78 | 0.99 | 0.52 | 0.64 | 0.53 | 0.62 |
| Geneformer | 0.80 | 0.98 | 0.50 | 0.65 | 0.51 | 0.62 |
| PCA | 0.77 | 0.97 | 0.51 | 0.61 | 0.51 | 0.60 |
| UCE | 0.78 | 0.96 | 0.48 | 0.62 | 0.48 | 0.59 |
| scCello | 0.73 | 0.96 | 0.48 | 0.62 | 0.49 | 0.58 |
| LangCell | 0.72 | 0.97 | 0.47 | 0.58 | 0.48 | 0.56 |
| CellFM | 0.73 | 0.92 | 0.47 | 0.53 | 0.45 | 0.54 |
| GenePT-w | 0.66 | 0.93 | 0.46 | 0.58 | 0.44 | 0.54 |
| CELLama | 0.52 | 0.83 | 0.31 | 0.42 | 0.33 | 0.40 |
| scBERT | 0.48 | 0.73 | 0.29 | 0.40 | 0.29 | 0.36 |

**b** Lymph Node · reference: v1, query: v2 · shared

| | Annotation metrics | | | | | | Aggregated |
|---|---|---|---|---|---|---|---|
| method | *Accuracy* | $Prec_{Macro}$ | $Recall_{Macro}$ | $F1_{Macro}$ | $AUC_{ROC}$ | $Acc_{top3}$ | *Annotation* |
| SingleR | 0.76 | 0.57 | 0.69 | 0.55 | n/a | n/a | 0.64 |
| scFoundation | 0.82 | 0.52 | 0.68 | 0.53 | 0.99 | 0.98 | 0.64 |
| SCimilarity | 0.81 | 0.54 | 0.67 | 0.54 | 0.96 | 0.99 | 0.64 |
| CellTypist | 0.77 | 0.58 | 0.65 | 0.55 | 0.95 | 0.96 | 0.64 |
| scPRINT | 0.81 | 0.54 | 0.65 | 0.54 | 0.98 | 0.99 | 0.63 |
| scGPT | 0.79 | 0.54 | 0.66 | 0.52 | 0.98 | 0.98 | 0.63 |
| CellPLM | 0.82 | 0.53 | 0.64 | 0.52 | 0.96 | 0.98 | 0.63 |
| scANVI | 0.82 | 0.50 | 0.65 | 0.51 | 0.97 | 0.97 | 0.62 |
| scVI | 0.78 | 0.52 | 0.64 | 0.53 | 0.97 | 0.99 | 0.62 |
| Geneformer | 0.80 | 0.50 | 0.65 | 0.51 | 0.98 | 0.98 | 0.62 |
| PCA | 0.77 | 0.51 | 0.61 | 0.51 | 0.96 | 0.97 | 0.60 |
| UCE | 0.78 | 0.48 | 0.62 | 0.48 | 0.97 | 0.96 | 0.59 |
| scCello | 0.73 | 0.48 | 0.62 | 0.49 | 0.97 | 0.96 | 0.58 |
| LangCell | 0.72 | 0.47 | 0.58 | 0.48 | 0.95 | 0.97 | 0.56 |
| CellFM | 0.73 | 0.47 | 0.53 | 0.45 | 0.94 | 0.92 | 0.54 |
| GenePT-w | 0.66 | 0.46 | 0.58 | 0.44 | 0.95 | 0.93 | 0.54 |
| CELLama | 0.52 | 0.31 | 0.42 | 0.33 | 0.88 | 0.83 | 0.40 |
| scBERT | 0.48 | 0.29 | 0.40 | 0.29 | 0.85 | 0.73 | 0.36 |

**Figure S203: Directional cross-dataset annotation for lymph node, TSv1 to TSv2. a,b,** Both panels contain 72,290 target cells from 21 shared labels because no target label is absent from the source in this direction. Methods and metrics are as in Fig. S185.

**a** Lymph Node · reference: v2, query: v1 · open-world

| method | Accuracy | $Acc_{top3}$ | $Prec_{Macro}$ | $Recall_{Macro}$ | $F1_{Macro}$ | Aggregated Annotation |
|---|---|---|---|---|---|---|
| SCimilarity | 0.80 | 0.97 | 0.62 | 0.65 | 0.56 | 0.66 |
| SingleR | 0.83 | n/a | 0.59 | 0.63 | 0.57 | 0.65 |
| CellTypist | 0.79 | 0.96 | 0.57 | 0.60 | 0.53 | 0.62 |
| scGPT | 0.77 | 0.96 | 0.57 | 0.59 | 0.52 | 0.61 |
| scFoundation | 0.77 | 0.96 | 0.53 | 0.61 | 0.53 | 0.61 |
| UCE | 0.78 | 0.96 | 0.52 | 0.59 | 0.53 | 0.60 |
| CellPLM | 0.77 | 0.96 | 0.51 | 0.62 | 0.51 | 0.60 |
| PCA | 0.75 | 0.96 | 0.56 | 0.57 | 0.52 | 0.60 |
| Geneformer | 0.76 | 0.94 | 0.54 | 0.57 | 0.51 | 0.59 |
| scVI | 0.74 | 0.96 | 0.52 | 0.59 | 0.50 | 0.59 |
| scCello | 0.71 | 0.96 | 0.54 | 0.57 | 0.50 | 0.58 |
| LangCell | 0.71 | 0.95 | 0.51 | 0.54 | 0.48 | 0.56 |
| scPRINT | 0.70 | 0.94 | 0.49 | 0.55 | 0.49 | 0.56 |
| scANVI | 0.71 | 0.94 | 0.47 | 0.56 | 0.47 | 0.55 |
| CellFM | 0.69 | 0.90 | 0.50 | 0.49 | 0.44 | 0.53 |
| GenePT-w | 0.69 | 0.89 | 0.37 | 0.48 | 0.35 | 0.47 |
| CELLama | 0.49 | 0.78 | 0.34 | 0.45 | 0.35 | 0.40 |
| scBERT | 0.47 | 0.74 | 0.23 | 0.31 | 0.23 | 0.31 |

**b** Lymph Node · reference: v2, query: v1 · shared

| method | Accuracy | $Prec_{Macro}$ | $Recall_{Macro}$ | $F1_{Macro}$ | $AUC_{ROC}$ | $Acc_{top3}$ | Aggregated Annotation |
|---|---|---|---|---|---|---|---|
| SCimilarity | 0.80 | 0.63 | 0.65 | 0.57 | 0.97 | 0.98 | 0.66 |
| SingleR | 0.84 | 0.60 | 0.63 | 0.57 | n/a | n/a | 0.66 |
| CellTypist | 0.79 | 0.58 | 0.60 | 0.54 | 0.95 | 0.96 | 0.63 |
| scGPT | 0.77 | 0.61 | 0.59 | 0.53 | 0.94 | 0.97 | 0.62 |
| scFoundation | 0.78 | 0.55 | 0.61 | 0.55 | 0.95 | 0.97 | 0.62 |
| UCE | 0.79 | 0.53 | 0.59 | 0.53 | 0.94 | 0.97 | 0.61 |
| CellPLM | 0.77 | 0.52 | 0.62 | 0.51 | 0.92 | 0.97 | 0.61 |
| PCA | 0.76 | 0.56 | 0.57 | 0.53 | 0.95 | 0.97 | 0.60 |
| Geneformer | 0.76 | 0.55 | 0.57 | 0.51 | 0.93 | 0.94 | 0.60 |
| scVI | 0.74 | 0.52 | 0.59 | 0.51 | 0.94 | 0.96 | 0.59 |
| scCello | 0.71 | 0.55 | 0.57 | 0.51 | 0.94 | 0.96 | 0.58 |
| LangCell | 0.71 | 0.51 | 0.54 | 0.48 | 0.94 | 0.96 | 0.56 |
| scPRINT | 0.70 | 0.49 | 0.55 | 0.49 | 0.92 | 0.94 | 0.56 |
| scANVI | 0.71 | 0.47 | 0.56 | 0.47 | 0.93 | 0.94 | 0.55 |
| CellFM | 0.70 | 0.50 | 0.49 | 0.44 | 0.91 | 0.91 | 0.53 |
| GenePT-w | 0.69 | 0.37 | 0.48 | 0.35 | 0.93 | 0.90 | 0.47 |
| CELLama | 0.50 | 0.34 | 0.45 | 0.35 | 0.88 | 0.78 | 0.41 |
| scBERT | 0.47 | 0.23 | 0.31 | 0.23 | 0.79 | 0.75 | 0.31 |

**Figure S204: Directional cross-dataset annotation for lymph node, TSv2 to TSv1.** **a,** All 51,173 target cells, including 289 cells from source-absent labels. **b,** 50,884 target cells from 21 shared labels. Methods and metrics are as in Fig. S185.

**a** Mammary · reference: v1, query: v2 · open-world

| method | Accuracy | $Acc_{top3}$ | $Prec_{Macro}$ | $Recall_{Macro}$ | $F1_{Macro}$ | Annotation |
|---|---|---|---|---|---|---|
| | Annotation metrics | | | | | Aggregated |
| scCello | 0.89 | 0.92 | 0.81 | 0.84 | 0.77 | 0.83 |
| scVI | 0.89 | 0.91 | 0.77 | 0.85 | 0.80 | 0.83 |
| LangCell | 0.86 | 0.92 | 0.78 | 0.82 | 0.72 | 0.80 |
| PCA | 0.89 | 0.91 | 0.73 | 0.83 | 0.72 | 0.79 |
| UCE | 0.89 | 0.91 | 0.75 | 0.82 | 0.71 | 0.79 |
| scGPT | 0.86 | 0.95 | 0.73 | 0.82 | 0.71 | 0.78 |
| SingleR | 0.88 | n/a | 0.69 | 0.83 | 0.70 | 0.78 |
| SCimilarity | 0.88 | 0.91 | 0.68 | 0.83 | 0.71 | 0.78 |
| scFoundation | 0.87 | 0.91 | 0.68 | 0.83 | 0.70 | 0.77 |
| scPRINT | 0.83 | 0.91 | 0.74 | 0.79 | 0.70 | 0.77 |
| Geneformer | 0.86 | 0.93 | 0.67 | 0.83 | 0.63 | 0.75 |
| CellPLM | 0.83 | 0.91 | 0.64 | 0.84 | 0.68 | 0.75 |
| CellTypist | 0.90 | 0.91 | 0.67 | 0.70 | 0.64 | 0.73 |
| GenePT-w | 0.87 | 0.94 | 0.63 | 0.63 | 0.58 | 0.68 |
| CellFM | 0.86 | 0.91 | 0.66 | 0.63 | 0.57 | 0.68 |
| scANVI | 0.79 | 0.91 | 0.44 | 0.59 | 0.45 | 0.57 |
| CELLama | 0.65 | 0.83 | 0.34 | 0.49 | 0.37 | 0.46 |
| scBERT | 0.34 | 0.55 | 0.25 | 0.33 | 0.23 | 0.29 |

**b** Mammary · reference: v1, query: v2 · shared

| method | Accuracy | $Prec_{Macro}$ | $Recall_{Macro}$ | $F1_{Macro}$ | $AUC_{ROC}$ | $Acc_{top3}$ | Annotation |
|---|---|---|---|---|---|---|---|
| | Annotation metrics | | | | | | Aggregated |
| scVI | 0.89 | 0.81 | 0.85 | 0.82 | 1.00 | 0.91 | 0.84 |
| scCello | 0.89 | 0.83 | 0.84 | 0.78 | 0.97 | 0.92 | 0.84 |
| LangCell | 0.86 | 0.80 | 0.82 | 0.74 | 0.99 | 0.92 | 0.81 |
| PCA | 0.89 | 0.75 | 0.83 | 0.73 | 0.99 | 0.91 | 0.80 |
| UCE | 0.89 | 0.76 | 0.82 | 0.71 | 0.99 | 0.91 | 0.80 |
| SingleR | 0.88 | 0.73 | 0.83 | 0.72 | n/a | n/a | 0.79 |
| SCimilarity | 0.88 | 0.72 | 0.83 | 0.73 | 1.00 | 0.91 | 0.79 |
| scGPT | 0.86 | 0.75 | 0.82 | 0.72 | 0.99 | 0.95 | 0.79 |
| scFoundation | 0.87 | 0.72 | 0.83 | 0.73 | 0.98 | 0.91 | 0.78 |
| scPRINT | 0.83 | 0.79 | 0.79 | 0.72 | 0.99 | 0.92 | 0.78 |
| Geneformer | 0.86 | 0.69 | 0.83 | 0.65 | 0.99 | 0.93 | 0.76 |
| CellPLM | 0.83 | 0.64 | 0.84 | 0.68 | 1.00 | 0.91 | 0.75 |
| CellTypist | 0.90 | 0.69 | 0.70 | 0.66 | 0.99 | 0.91 | 0.74 |
| GenePT-w | 0.87 | 0.63 | 0.63 | 0.58 | 0.99 | 0.94 | 0.68 |
| CellFM | 0.86 | 0.66 | 0.63 | 0.57 | 0.99 | 0.91 | 0.68 |
| scANVI | 0.79 | 0.44 | 0.59 | 0.45 | 0.99 | 0.91 | 0.57 |
| CELLama | 0.65 | 0.34 | 0.49 | 0.37 | 0.93 | 0.83 | 0.46 |
| scBERT | 0.34 | 0.25 | 0.33 | 0.23 | 0.80 | 0.55 | 0.29 |

**Figure S205: Directional cross-dataset annotation for mammary, TSv1 to TSv2. a,** All 18,539 target cells, including 2 cells from source-absent labels. **b,** 18,537 target cells from 14 shared labels. Methods and metrics are as in Fig. S185.

**a** Mammary · reference: v2, query: v1 · open-world

| method | Accuracy | $Acc_{top3}$ | $Prec_{Macro}$ | $Recall_{Macro}$ | $F1_{Macro}$ | Aggregated Annotation |
|---|---|---|---|---|---|---|
| scFoundation | 0.93 | 0.99 | 0.77 | 0.81 | 0.77 | 0.82 |
| scCello | 0.94 | 0.99 | 0.74 | 0.75 | 0.73 | 0.79 |
| SingleR | 0.91 | n/a | 0.74 | 0.74 | 0.72 | 0.78 |
| LangCell | 0.92 | 0.99 | 0.74 | 0.69 | 0.70 | 0.76 |
| scGPT | 0.84 | 0.99 | 0.74 | 0.70 | 0.69 | 0.74 |
| SCimilarity | 0.91 | 0.99 | 0.74 | 0.68 | 0.64 | 0.74 |
| scPRINT | 0.88 | 0.97 | 0.69 | 0.70 | 0.65 | 0.73 |
| scVI | 0.90 | 0.98 | 0.64 | 0.73 | 0.64 | 0.73 |
| CellPLM | 0.86 | 0.97 | 0.63 | 0.72 | 0.64 | 0.71 |
| Geneformer | 0.86 | 0.98 | 0.73 | 0.64 | 0.62 | 0.71 |
| CellTypist | 0.93 | 0.99 | 0.67 | 0.63 | 0.60 | 0.71 |
| PCA | 0.91 | 0.99 | 0.66 | 0.62 | 0.57 | 0.69 |
| UCE | 0.88 | 0.99 | 0.64 | 0.62 | 0.60 | 0.68 |
| scANVI | 0.86 | 0.97 | 0.60 | 0.67 | 0.61 | 0.68 |
| CellFM | 0.89 | 0.98 | 0.58 | 0.63 | 0.55 | 0.66 |
| CELLama | 0.69 | 0.89 | 0.43 | 0.44 | 0.42 | 0.50 |
| scBERT | 0.46 | 0.80 | 0.27 | 0.32 | 0.24 | 0.32 |
| GenePT-w | 0.23 | 0.87 | 0.38 | 0.15 | 0.13 | 0.22 |

**b** Mammary · reference: v2, query: v1 · shared

| method | Accuracy | $Prec_{Macro}$ | $Recall_{Macro}$ | $F1_{Macro}$ | $AUC_{ROC}$ | $Acc_{top3}$ | Aggregated Annotation |
|---|---|---|---|---|---|---|---|
| scFoundation | 0.93 | 0.78 | 0.81 | 0.78 | 0.99 | 0.99 | 0.83 |
| scCello | 0.94 | 0.75 | 0.75 | 0.73 | 0.98 | 0.99 | 0.79 |
| SingleR | 0.91 | 0.74 | 0.74 | 0.72 | n/a | n/a | 0.78 |
| LangCell | 0.92 | 0.74 | 0.69 | 0.70 | 0.96 | 0.99 | 0.76 |
| scGPT | 0.84 | 0.74 | 0.70 | 0.69 | 0.98 | 0.99 | 0.75 |
| SCimilarity | 0.92 | 0.74 | 0.68 | 0.64 | 0.98 | 0.99 | 0.74 |
| scPRINT | 0.88 | 0.69 | 0.70 | 0.65 | 0.98 | 0.97 | 0.73 |
| scVI | 0.90 | 0.64 | 0.73 | 0.64 | 0.98 | 0.98 | 0.73 |
| CellPLM | 0.86 | 0.65 | 0.72 | 0.65 | 0.95 | 0.98 | 0.72 |
| Geneformer | 0.86 | 0.73 | 0.64 | 0.62 | 0.97 | 0.98 | 0.71 |
| CellTypist | 0.93 | 0.68 | 0.63 | 0.60 | 0.98 | 0.99 | 0.71 |
| PCA | 0.91 | 0.66 | 0.62 | 0.57 | 0.99 | 0.99 | 0.69 |
| UCE | 0.88 | 0.64 | 0.62 | 0.60 | 0.97 | 0.99 | 0.68 |
| scANVI | 0.86 | 0.60 | 0.67 | 0.61 | 0.95 | 0.97 | 0.68 |
| CellFM | 0.89 | 0.58 | 0.63 | 0.55 | 0.95 | 0.99 | 0.66 |
| CELLama | 0.70 | 0.43 | 0.44 | 0.42 | 0.88 | 0.89 | 0.50 |
| scBERT | 0.46 | 0.27 | 0.32 | 0.24 | 0.79 | 0.80 | 0.32 |
| GenePT-w | 0.23 | 0.38 | 0.15 | 0.13 | 0.82 | 0.88 | 0.22 |

**Figure S206: Directional cross-dataset annotation for mammary, TSv2 to TSv1. a,** All 10,710 target cells, including 13 cells from source-absent labels. **b,** 10,697 target cells from 14 shared labels. Methods and metrics are as in Fig. S185.

**a** Muscle · reference: v1, query: v2 · open-world

| method | Annotation metrics | | | | | Aggregated |
|---|---|---|---|---|---|---|
| | *Accuracy* | *Acc*$_{top3}$ | *Prec*$_{Macro}$ | *Recall*$_{Macro}$ | *F1*$_{Macro}$ | *Annotation* |
| CellTypist | 0.93 | 0.98 | 0.56 | 0.69 | 0.58 | 0.69 |
| SingleR | 0.92 | n/a | 0.63 | 0.62 | 0.55 | 0.68 |
| Geneformer | 0.92 | 0.97 | 0.57 | 0.60 | 0.55 | 0.66 |
| CellPLM | 0.90 | 0.96 | 0.55 | 0.62 | 0.53 | 0.65 |
| PCA | 0.89 | 0.97 | 0.54 | 0.62 | 0.53 | 0.64 |
| scVI | 0.91 | 0.97 | 0.57 | 0.58 | 0.52 | 0.64 |
| scANVI | 0.89 | 0.97 | 0.52 | 0.63 | 0.52 | 0.64 |
| scGPT | 0.88 | 0.96 | 0.53 | 0.61 | 0.51 | 0.64 |
| SCimilarity | 0.90 | 0.96 | 0.53 | 0.61 | 0.50 | 0.64 |
| LangCell | 0.89 | 0.97 | 0.54 | 0.59 | 0.50 | 0.63 |
| UCE | 0.90 | 0.95 | 0.48 | 0.60 | 0.49 | 0.62 |
| scFoundation | 0.88 | 0.96 | 0.50 | 0.60 | 0.49 | 0.62 |
| CellFM | 0.90 | 0.97 | 0.54 | 0.53 | 0.48 | 0.61 |
| scPRINT | 0.84 | 0.96 | 0.55 | 0.55 | 0.50 | 0.61 |
| scCello | 0.83 | 0.96 | 0.49 | 0.58 | 0.47 | 0.59 |
| GenePT-w | 0.85 | 0.95 | 0.43 | 0.47 | 0.38 | 0.53 |
| CELLama | 0.80 | 0.94 | 0.41 | 0.47 | 0.39 | 0.52 |
| scBERT | 0.32 | 0.83 | 0.26 | 0.23 | 0.19 | 0.25 |

**b** Muscle · reference: v1, query: v2 · shared

| method | Annotation metrics | | | | | | Aggregated |
|---|---|---|---|---|---|---|---|
| | *Accuracy* | *Prec*$_{Macro}$ | *Recall*$_{Macro}$ | *F1*$_{Macro}$ | *AUC*$_{ROC}$ | *Acc*$_{top3}$ | *Annotation* |
| CellTypist | 0.94 | 0.59 | 0.69 | 0.60 | 0.97 | 0.98 | 0.70 |
| SingleR | 0.92 | 0.65 | 0.62 | 0.56 | n/a | n/a | 0.69 |
| Geneformer | 0.93 | 0.60 | 0.60 | 0.56 | 0.98 | 0.97 | 0.67 |
| CellPLM | 0.91 | 0.58 | 0.62 | 0.54 | 0.97 | 0.96 | 0.66 |
| PCA | 0.90 | 0.56 | 0.62 | 0.54 | 0.96 | 0.98 | 0.65 |
| scVI | 0.91 | 0.60 | 0.58 | 0.52 | 0.99 | 0.98 | 0.65 |
| scANVI | 0.89 | 0.54 | 0.63 | 0.53 | 0.96 | 0.98 | 0.65 |
| scGPT | 0.89 | 0.56 | 0.61 | 0.53 | 0.98 | 0.97 | 0.65 |
| SCimilarity | 0.91 | 0.55 | 0.61 | 0.51 | 0.97 | 0.97 | 0.64 |
| LangCell | 0.90 | 0.55 | 0.59 | 0.51 | 0.98 | 0.98 | 0.64 |
| UCE | 0.90 | 0.50 | 0.60 | 0.51 | 0.96 | 0.96 | 0.63 |
| scFoundation | 0.89 | 0.52 | 0.60 | 0.50 | 0.95 | 0.97 | 0.63 |
| CellFM | 0.90 | 0.56 | 0.53 | 0.49 | 0.98 | 0.98 | 0.62 |
| scPRINT | 0.85 | 0.56 | 0.55 | 0.50 | 0.97 | 0.97 | 0.62 |
| scCello | 0.83 | 0.52 | 0.58 | 0.47 | 0.93 | 0.96 | 0.60 |
| GenePT-w | 0.86 | 0.45 | 0.47 | 0.40 | 0.97 | 0.95 | 0.54 |
| CELLama | 0.81 | 0.43 | 0.47 | 0.39 | 0.95 | 0.95 | 0.53 |
| scBERT | 0.32 | 0.28 | 0.23 | 0.19 | 0.84 | 0.83 | 0.26 |

**Figure S207: Directional cross-dataset annotation for muscle, TSv1 to TSv2. a,** All 13,035 target cells, including 94 cells from source-absent labels. **b,** 12,941 target cells from 25 shared labels. Methods and metrics are as in Fig. S185.

**Muscle · reference: v2, query: v1 · open-world**

| method | Annotation metrics: *Accuracy* | *Acc*top3 | *Prec*Macro | *Recall*Macro | *F1*Macro | Aggregated: *Annotation* |
|---|---|---|---|---|---|---|
| scCello | 0.86 | 0.94 | 0.54 | 0.63 | 0.54 | 0.64 |
| SingleR | 0.86 | n/a | 0.52 | 0.67 | 0.49 | 0.64 |
| PCA | 0.87 | 0.95 | 0.53 | 0.63 | 0.51 | 0.63 |
| SCimilarity | 0.80 | 0.93 | 0.53 | 0.64 | 0.50 | 0.62 |
| scGPT | 0.82 | 0.91 | 0.54 | 0.58 | 0.48 | 0.60 |
| scFoundation | 0.82 | 0.94 | 0.50 | 0.61 | 0.48 | 0.60 |
| CellTypist | 0.87 | 0.94 | 0.49 | 0.55 | 0.48 | 0.60 |
| UCE | 0.81 | 0.93 | 0.49 | 0.59 | 0.47 | 0.59 |
| scVI | 0.81 | 0.94 | 0.48 | 0.61 | 0.46 | 0.59 |
| CellPLM | 0.83 | 0.92 | 0.48 | 0.56 | 0.47 | 0.59 |
| LangCell | 0.80 | 0.92 | 0.53 | 0.53 | 0.46 | 0.58 |
| Geneformer | 0.80 | 0.89 | 0.51 | 0.55 | 0.45 | 0.58 |
| scPRINT | 0.80 | 0.91 | 0.48 | 0.57 | 0.44 | 0.57 |
| scANVI | 0.85 | 0.93 | 0.43 | 0.56 | 0.45 | 0.57 |
| CellFM | 0.85 | 0.95 | 0.42 | 0.49 | 0.41 | 0.54 |
| CELLama | 0.62 | 0.83 | 0.28 | 0.39 | 0.25 | 0.38 |
| scBERT | 0.45 | 0.72 | 0.21 | 0.24 | 0.18 | 0.27 |
| GenePT-w | 0.07 | 0.33 | 0.26 | 0.14 | 0.09 | 0.14 |

**b**

**Muscle · reference: v2, query: v1 · shared**

| method | Annotation metrics: *Accuracy* | *Prec*Macro | *Recall*Macro | *F1*Macro | *AUC*ROC | *Acc*top3 | Aggregated: *Annotation* |
|---|---|---|---|---|---|---|---|
| scCello | 0.88 | 0.56 | 0.63 | 0.54 | 0.96 | 0.97 | 0.65 |
| SingleR | 0.88 | 0.54 | 0.67 | 0.51 | n/a | n/a | 0.65 |
| PCA | 0.89 | 0.54 | 0.63 | 0.52 | 0.97 | 0.97 | 0.65 |
| SCimilarity | 0.82 | 0.54 | 0.64 | 0.50 | 0.93 | 0.95 | 0.63 |
| scFoundation | 0.84 | 0.52 | 0.61 | 0.49 | 0.96 | 0.97 | 0.61 |
| scGPT | 0.84 | 0.55 | 0.58 | 0.49 | 0.93 | 0.94 | 0.61 |
| CellTypist | 0.89 | 0.50 | 0.55 | 0.49 | 0.97 | 0.97 | 0.61 |
| UCE | 0.83 | 0.50 | 0.59 | 0.48 | 0.96 | 0.95 | 0.60 |
| scVI | 0.83 | 0.49 | 0.61 | 0.46 | 0.91 | 0.96 | 0.60 |
| CellPLM | 0.85 | 0.50 | 0.56 | 0.48 | 0.87 | 0.94 | 0.60 |
| LangCell | 0.83 | 0.54 | 0.53 | 0.47 | 0.96 | 0.94 | 0.59 |
| Geneformer | 0.82 | 0.52 | 0.55 | 0.45 | 0.95 | 0.91 | 0.58 |
| scPRINT | 0.82 | 0.49 | 0.57 | 0.45 | 0.95 | 0.93 | 0.58 |
| scANVI | 0.87 | 0.44 | 0.56 | 0.46 | 0.93 | 0.96 | 0.58 |
| CellFM | 0.87 | 0.44 | 0.49 | 0.41 | 0.96 | 0.97 | 0.55 |
| CELLama | 0.64 | 0.28 | 0.39 | 0.25 | 0.86 | 0.85 | 0.39 |
| scBERT | 0.47 | 0.21 | 0.24 | 0.18 | 0.71 | 0.74 | 0.27 |
| GenePT-w | 0.07 | 0.27 | 0.14 | 0.09 | 0.81 | 0.34 | 0.14 |

**Figure S208: Directional cross-dataset annotation for muscle, TSv2 to TSv1. a,** All 30,066 target cells, including 806 cells from source-absent labels. **b,** 29,260 target cells from 25 shared labels. Methods and metrics are as in Fig. S185.

**a** Prostate · reference: v1, query: v2 · open-world

| method | Accuracy | $Acc_{top3}$ | $Prec_{Macro}$ | $Recall_{Macro}$ | $F1_{Macro}$ | Aggregated Annotation |
|---|---|---|---|---|---|---|
| scFoundation | 0.94 | 1.00 | 0.84 | 0.86 | 0.85 | 0.88 |
| CellTypist | 0.93 | 1.00 | 0.83 | 0.88 | 0.85 | 0.87 |
| Geneformer | 0.91 | 1.00 | 0.83 | 0.86 | 0.84 | 0.86 |
| SCimilarity | 0.92 | 0.99 | 0.83 | 0.86 | 0.83 | 0.86 |
| scPRINT | 0.93 | 1.00 | 0.82 | 0.86 | 0.83 | 0.86 |
| PCA | 0.92 | 0.99 | 0.82 | 0.86 | 0.83 | 0.86 |
| scGPT | 0.91 | 0.99 | 0.81 | 0.85 | 0.82 | 0.85 |
| scANVI | 0.91 | 1.00 | 0.81 | 0.84 | 0.82 | 0.85 |
| CellPLM | 0.91 | 0.99 | 0.82 | 0.84 | 0.81 | 0.84 |
| LangCell | 0.90 | 0.99 | 0.80 | 0.85 | 0.81 | 0.84 |
| UCE | 0.89 | 0.99 | 0.80 | 0.84 | 0.81 | 0.84 |
| scCello | 0.89 | 0.99 | 0.80 | 0.84 | 0.81 | 0.83 |
| CellFM | 0.89 | 0.99 | 0.79 | 0.81 | 0.79 | 0.82 |
| scVI | 0.87 | 0.99 | 0.79 | 0.83 | 0.80 | 0.82 |
| SingleR | 0.83 | n/a | 0.80 | 0.85 | 0.79 | 0.82 |
| CELLama | 0.66 | 0.88 | 0.63 | 0.66 | 0.58 | 0.63 |
| GenePT-w | 0.59 | 0.84 | 0.59 | 0.58 | 0.50 | 0.57 |
| scBERT | 0.63 | 0.87 | 0.46 | 0.46 | 0.43 | 0.49 |

**b** Prostate · reference: v1, query: v2 · shared

| method | Accuracy | $Prec_{Macro}$ | $Recall_{Macro}$ | $F1_{Macro}$ | $AUC_{ROC}$ | $Acc_{top3}$ | Aggregated Annotation |
|---|---|---|---|---|---|---|---|
| scFoundation | 0.94 | 0.84 | 0.86 | 0.85 | 0.96 | 1.00 | 0.88 |
| CellTypist | 0.93 | 0.83 | 0.88 | 0.85 | 1.00 | 1.00 | 0.87 |
| Geneformer | 0.91 | 0.83 | 0.86 | 0.84 | 1.00 | 1.00 | 0.86 |
| SCimilarity | 0.92 | 0.83 | 0.86 | 0.83 | 0.99 | 0.99 | 0.86 |
| scPRINT | 0.93 | 0.82 | 0.86 | 0.83 | 0.97 | 1.00 | 0.86 |
| PCA | 0.92 | 0.82 | 0.86 | 0.83 | 0.97 | 0.99 | 0.86 |
| scGPT | 0.91 | 0.81 | 0.85 | 0.82 | 0.97 | 0.99 | 0.85 |
| scANVI | 0.91 | 0.81 | 0.84 | 0.82 | 0.98 | 1.00 | 0.85 |
| CellPLM | 0.91 | 0.82 | 0.84 | 0.81 | 0.94 | 0.99 | 0.84 |
| LangCell | 0.90 | 0.80 | 0.85 | 0.81 | 0.98 | 0.99 | 0.84 |
| UCE | 0.89 | 0.80 | 0.84 | 0.81 | 0.99 | 0.99 | 0.84 |
| scCello | 0.89 | 0.80 | 0.84 | 0.81 | 0.96 | 0.99 | 0.83 |
| CellFM | 0.89 | 0.79 | 0.81 | 0.79 | 0.92 | 0.99 | 0.82 |
| scVI | 0.87 | 0.79 | 0.83 | 0.80 | 0.93 | 0.99 | 0.82 |
| SingleR | 0.83 | 0.80 | 0.85 | 0.79 | n/a | n/a | 0.82 |
| CELLama | 0.66 | 0.63 | 0.66 | 0.58 | 0.95 | 0.88 | 0.63 |
| GenePT-w | 0.59 | 0.59 | 0.58 | 0.50 | 0.99 | 0.84 | 0.57 |
| scBERT | 0.63 | 0.46 | 0.46 | 0.43 | 0.91 | 0.87 | 0.49 |

**Figure S209: Directional cross-dataset annotation for prostate, TSv1 to TSv2. a,b,** Both panels contain 2,044 target cells from 14 shared labels because no target label is absent from the source in this direction. Methods and metrics are as in Fig. S185.

**a** Prostate · reference: v2, query: v1 · open-world

| method | Annotation metrics: $Accuracy$ | $Acc_{top3}$ | $Prec_{Macro}$ | $Recall_{Macro}$ | $F1_{Macro}$ | Aggregated: $Annotation$ |
|---|---|---|---|---|---|---|
| scFoundation | 0.93 | 0.98 | 0.79 | 0.72 | 0.71 | 0.79 |
| UCE | 0.89 | 0.98 | 0.80 | 0.72 | 0.73 | 0.78 |
| scGPT | 0.91 | 0.98 | 0.75 | 0.73 | 0.72 | 0.78 |
| CellTypist | 0.90 | 0.99 | 0.79 | 0.72 | 0.67 | 0.77 |
| PCA | 0.88 | 0.98 | 0.76 | 0.72 | 0.71 | 0.77 |
| LangCell | 0.90 | 0.98 | 0.75 | 0.69 | 0.69 | 0.76 |
| Geneformer | 0.87 | 0.98 | 0.74 | 0.72 | 0.68 | 0.75 |
| scPRINT | 0.86 | 0.97 | 0.73 | 0.73 | 0.68 | 0.75 |
| scVI | 0.84 | 0.99 | 0.73 | 0.68 | 0.66 | 0.73 |
| SingleR | 0.85 | n/a | 0.62 | 0.78 | 0.65 | 0.73 |
| CellFM | 0.83 | 0.98 | 0.68 | 0.70 | 0.64 | 0.71 |
| scCello | 0.72 | 0.96 | 0.70 | 0.69 | 0.66 | 0.69 |
| scANVI | 0.76 | 0.95 | 0.54 | 0.70 | 0.57 | 0.64 |
| SCimilarity | 0.54 | 0.83 | 0.61 | 0.71 | 0.57 | 0.61 |
| CellPLM | 0.49 | 0.71 | 0.67 | 0.62 | 0.55 | 0.58 |
| CELLama | 0.47 | 0.81 | 0.39 | 0.46 | 0.38 | 0.43 |
| scBERT | 0.49 | 0.82 | 0.32 | 0.44 | 0.32 | 0.39 |
| GenePT-w | 0.12 | 0.61 | 0.35 | 0.18 | 0.09 | 0.19 |

**b** Prostate · reference: v2, query: v1 · shared

| method | Annotation metrics: $Accuracy$ | $Prec_{Macro}$ | $Recall_{Macro}$ | $F1_{Macro}$ | $AUC_{ROC}$ | $Acc_{top3}$ | Aggregated: $Annotation$ |
|---|---|---|---|---|---|---|---|
| scFoundation | 0.93 | 0.79 | 0.72 | 0.71 | 0.92 | 0.98 | 0.79 |
| UCE | 0.89 | 0.80 | 0.72 | 0.73 | 0.90 | 0.98 | 0.78 |
| scGPT | 0.91 | 0.75 | 0.73 | 0.72 | 0.95 | 0.98 | 0.78 |
| CellTypist | 0.90 | 0.79 | 0.72 | 0.67 | 0.97 | 0.99 | 0.77 |
| PCA | 0.88 | 0.76 | 0.72 | 0.71 | 0.92 | 0.98 | 0.77 |
| LangCell | 0.90 | 0.75 | 0.69 | 0.69 | 0.95 | 0.98 | 0.76 |
| Geneformer | 0.87 | 0.74 | 0.72 | 0.68 | 0.93 | 0.98 | 0.75 |
| scPRINT | 0.86 | 0.73 | 0.73 | 0.68 | 0.94 | 0.97 | 0.75 |
| scVI | 0.84 | 0.73 | 0.68 | 0.66 | 0.92 | 0.99 | 0.73 |
| SingleR | 0.85 | 0.62 | 0.78 | 0.65 | n/a | n/a | 0.73 |
| CellFM | 0.83 | 0.68 | 0.70 | 0.64 | 0.87 | 0.98 | 0.71 |
| scCello | 0.72 | 0.70 | 0.69 | 0.66 | 0.87 | 0.96 | 0.69 |
| scANVI | 0.76 | 0.54 | 0.70 | 0.57 | 0.87 | 0.95 | 0.64 |
| SCimilarity | 0.54 | 0.61 | 0.71 | 0.57 | 0.93 | 0.83 | 0.61 |
| CellPLM | 0.49 | 0.67 | 0.62 | 0.55 | 0.92 | 0.71 | 0.58 |
| CELLama | 0.47 | 0.39 | 0.46 | 0.38 | 0.87 | 0.81 | 0.43 |
| scBERT | 0.49 | 0.32 | 0.44 | 0.32 | 0.82 | 0.82 | 0.39 |
| GenePT-w | 0.12 | 0.35 | 0.18 | 0.09 | 0.85 | 0.61 | 0.19 |

**Figure S210: Directional cross-dataset annotation for prostate, TSv2 to TSv1. a,b,** Both panels contain 15,798 target cells from 14 shared labels because no target label is absent from the source in this direction. Methods and metrics are as in Fig. S185.

**a** Salivary Gland · reference: v1, query: v2 · open-world

| method | Annotation metrics: *Accuracy* | *Acc*$_{top3}$ | *Prec*$_{Macro}$ | *Recall*$_{Macro}$ | *F1*$_{Macro}$ | Aggregated: *Annotation* |
|---|---|---|---|---|---|---|
| CellTypist | 0.63 | 0.96 | 0.64 | 0.71 | 0.62 | 0.65 |
| PCA | 0.63 | 0.97 | 0.64 | 0.70 | 0.60 | 0.64 |
| CellPLM | 0.73 | 0.96 | 0.54 | 0.65 | 0.55 | 0.62 |
| scFoundation | 0.59 | 0.96 | 0.58 | 0.70 | 0.56 | 0.61 |
| scANVI | 0.66 | 0.93 | 0.56 | 0.67 | 0.55 | 0.61 |
| SingleR | 0.58 | n/a | 0.61 | 0.67 | 0.53 | 0.60 |
| scPRINT | 0.66 | 0.96 | 0.57 | 0.62 | 0.54 | 0.60 |
| scCello | 0.57 | 0.95 | 0.57 | 0.64 | 0.55 | 0.59 |
| CellFM | 0.58 | 0.97 | 0.55 | 0.62 | 0.54 | 0.57 |
| Geneformer | 0.54 | 0.97 | 0.55 | 0.65 | 0.52 | 0.56 |
| scGPT | 0.58 | 0.95 | 0.53 | 0.63 | 0.50 | 0.56 |
| scVI | 0.56 | 0.96 | 0.54 | 0.63 | 0.49 | 0.55 |
| LangCell | 0.57 | 0.92 | 0.52 | 0.62 | 0.49 | 0.55 |
| UCE | 0.50 | 0.95 | 0.56 | 0.62 | 0.51 | 0.55 |
| SCimilarity | 0.51 | 0.95 | 0.53 | 0.63 | 0.49 | 0.54 |
| GenePT-w | 0.58 | 0.92 | 0.50 | 0.61 | 0.47 | 0.54 |
| CELLama | 0.48 | 0.86 | 0.44 | 0.51 | 0.40 | 0.46 |
| scBERT | 0.43 | 0.79 | 0.35 | 0.38 | 0.29 | 0.36 |

**b** Salivary Gland · reference: v1, query: v2 · shared

| method | Annotation metrics: *Accuracy* | *Prec*$_{Macro}$ | *Recall*$_{Macro}$ | *F1*$_{Macro}$ | *AUC*$_{ROC}$ | *Acc*$_{top3}$ | Aggregated: *Annotation* |
|---|---|---|---|---|---|---|---|
| CellTypist | 0.64 | 0.65 | 0.71 | 0.63 | 0.97 | 0.97 | 0.66 |
| PCA | 0.64 | 0.65 | 0.70 | 0.60 | 0.98 | 0.98 | 0.65 |
| CellPLM | 0.73 | 0.54 | 0.65 | 0.55 | 0.98 | 0.97 | 0.62 |
| scANVI | 0.67 | 0.58 | 0.67 | 0.55 | 0.98 | 0.94 | 0.62 |
| scFoundation | 0.60 | 0.59 | 0.70 | 0.57 | 0.98 | 0.97 | 0.61 |
| scPRINT | 0.67 | 0.59 | 0.62 | 0.54 | 0.98 | 0.97 | 0.60 |
| SingleR | 0.59 | 0.62 | 0.67 | 0.54 | n/a | n/a | 0.60 |
| scCello | 0.58 | 0.58 | 0.64 | 0.56 | 0.99 | 0.97 | 0.59 |
| CellFM | 0.58 | 0.57 | 0.62 | 0.54 | 0.98 | 0.98 | 0.58 |
| Geneformer | 0.54 | 0.57 | 0.65 | 0.52 | 0.99 | 0.98 | 0.57 |
| scGPT | 0.59 | 0.54 | 0.63 | 0.51 | 0.98 | 0.96 | 0.56 |
| scVI | 0.56 | 0.56 | 0.63 | 0.49 | 0.98 | 0.97 | 0.56 |
| LangCell | 0.58 | 0.55 | 0.62 | 0.49 | 0.98 | 0.93 | 0.56 |
| UCE | 0.50 | 0.58 | 0.62 | 0.51 | 0.97 | 0.96 | 0.55 |
| SCimilarity | 0.51 | 0.56 | 0.63 | 0.49 | 0.99 | 0.96 | 0.55 |
| GenePT-w | 0.58 | 0.51 | 0.61 | 0.47 | 0.97 | 0.93 | 0.54 |
| CELLama | 0.48 | 0.44 | 0.51 | 0.40 | 0.93 | 0.87 | 0.46 |
| scBERT | 0.44 | 0.35 | 0.38 | 0.30 | 0.85 | 0.80 | 0.37 |

**Figure S211: Directional cross-dataset annotation for salivary gland, TSv1 to TSv2. a,** All 11,561 target cells, including 141 cells from source-absent labels. **b,** 11,420 target cells from 20 shared labels. Methods and metrics are as in Fig. S185.

**a** Salivary Gland · reference: v2, query: v1 · open-world

| method | Annotation metrics | | | | | Aggregated |
|---|---|---|---|---|---|---|
| | *Accuracy* | $Acc_{top3}$ | $Prec_{Macro}$ | $Recall_{Macro}$ | $F1_{Macro}$ | *Annotation* |
| SCimilarity | 0.82 | 0.90 | 0.67 | 0.66 | 0.61 | 0.69 |
| scCello | 0.79 | 0.92 | 0.63 | 0.66 | 0.57 | 0.66 |
| Geneformer | 0.79 | 0.91 | 0.67 | 0.59 | 0.58 | 0.66 |
| scPRINT | 0.78 | 0.90 | 0.70 | 0.58 | 0.55 | 0.66 |
| scGPT | 0.79 | 0.91 | 0.64 | 0.63 | 0.55 | 0.66 |
| PCA | 0.82 | 0.91 | 0.65 | 0.59 | 0.57 | 0.66 |
| CellTypist | 0.83 | 0.93 | 0.63 | 0.58 | 0.57 | 0.65 |
| SingleR | 0.77 | n/a | 0.59 | 0.65 | 0.54 | 0.64 |
| scANVI | 0.77 | 0.90 | 0.59 | 0.63 | 0.55 | 0.63 |
| UCE | 0.76 | 0.91 | 0.64 | 0.56 | 0.55 | 0.63 |
| LangCell | 0.74 | 0.89 | 0.61 | 0.60 | 0.53 | 0.62 |
| scFoundation | 0.77 | 0.92 | 0.59 | 0.56 | 0.53 | 0.61 |
| CellFM | 0.76 | 0.92 | 0.56 | 0.50 | 0.50 | 0.58 |
| CellPLM | 0.69 | 0.87 | 0.51 | 0.45 | 0.41 | 0.51 |
| scVI | 0.58 | 0.83 | 0.54 | 0.41 | 0.38 | 0.48 |
| GenePT-w | 0.66 | 0.91 | 0.44 | 0.42 | 0.39 | 0.48 |
| CELLama | 0.46 | 0.71 | 0.34 | 0.31 | 0.28 | 0.35 |
| scBERT | 0.54 | 0.81 | 0.24 | 0.26 | 0.22 | 0.31 |

**b** Salivary Gland · reference: v2, query: v1 · shared

| method | Annotation metrics | | | | | | Aggregated |
|---|---|---|---|---|---|---|---|
| | *Accuracy* | $Prec_{Macro}$ | $Recall_{Macro}$ | $F1_{Macro}$ | $AUC_{ROC}$ | $Acc_{top3}$ | *Annotation* |
| SCimilarity | 0.86 | 0.70 | 0.66 | 0.63 | 0.97 | 0.94 | 0.71 |
| scCello | 0.83 | 0.66 | 0.66 | 0.59 | 0.98 | 0.96 | 0.68 |
| Geneformer | 0.83 | 0.70 | 0.59 | 0.60 | 0.96 | 0.95 | 0.68 |
| scPRINT | 0.82 | 0.73 | 0.58 | 0.57 | 0.95 | 0.94 | 0.68 |
| PCA | 0.86 | 0.67 | 0.59 | 0.58 | 0.95 | 0.96 | 0.68 |
| scGPT | 0.83 | 0.66 | 0.63 | 0.57 | 0.97 | 0.95 | 0.68 |
| CellTypist | 0.87 | 0.67 | 0.58 | 0.58 | 0.97 | 0.98 | 0.67 |
| SingleR | 0.81 | 0.61 | 0.65 | 0.55 | n/a | n/a | 0.66 |
| scANVI | 0.80 | 0.61 | 0.63 | 0.56 | 0.95 | 0.94 | 0.65 |
| UCE | 0.80 | 0.66 | 0.56 | 0.56 | 0.97 | 0.95 | 0.64 |
| LangCell | 0.78 | 0.63 | 0.60 | 0.55 | 0.96 | 0.94 | 0.64 |
| scFoundation | 0.81 | 0.62 | 0.56 | 0.54 | 0.96 | 0.97 | 0.63 |
| CellFM | 0.80 | 0.58 | 0.50 | 0.51 | 0.96 | 0.97 | 0.60 |
| CellPLM | 0.72 | 0.54 | 0.45 | 0.43 | 0.96 | 0.92 | 0.53 |
| scVI | 0.60 | 0.56 | 0.41 | 0.39 | 0.91 | 0.87 | 0.49 |
| GenePT-w | 0.69 | 0.46 | 0.42 | 0.39 | 0.96 | 0.95 | 0.49 |
| CELLama | 0.48 | 0.36 | 0.31 | 0.29 | 0.86 | 0.74 | 0.36 |
| scBERT | 0.57 | 0.25 | 0.26 | 0.23 | 0.86 | 0.85 | 0.33 |

**Figure S212: Directional cross-dataset annotation for salivary gland, TSv2 to TSv1. a,** All 22,577 target cells, including 1,060 cells from source-absent labels. **b,** 21,517 target cells from 20 shared labels. Methods and metrics are as in Fig. S185.

**a**

Skin · reference: v1, query: v2 · open-world

| method | Accuracy | $Acc_{top3}$ | $Prec_{Macro}$ | $Recall_{Macro}$ | $F1_{Macro}$ | Aggregated Annotation |
|---|---|---|---|---|---|---|
| Geneformer | 0.90 | 0.99 | 0.68 | 0.71 | 0.66 | 0.74 |
| LangCell | 0.89 | 0.98 | 0.68 | 0.70 | 0.65 | 0.73 |
| scFoundation | 0.90 | 0.99 | 0.69 | 0.68 | 0.66 | 0.73 |
| scGPT | 0.86 | 0.98 | 0.69 | 0.70 | 0.67 | 0.73 |
| CellTypist | 0.90 | 0.99 | 0.63 | 0.71 | 0.65 | 0.72 |
| SingleR | 0.89 | n/a | 0.61 | 0.74 | 0.64 | 0.72 |
| scCello | 0.88 | 0.98 | 0.62 | 0.69 | 0.63 | 0.71 |
| scPRINT | 0.89 | 0.98 | 0.63 | 0.66 | 0.63 | 0.70 |
| SCimilarity | 0.88 | 0.99 | 0.63 | 0.66 | 0.63 | 0.70 |
| scVI | 0.89 | 0.99 | 0.61 | 0.67 | 0.63 | 0.70 |
| PCA | 0.87 | 0.98 | 0.62 | 0.67 | 0.62 | 0.70 |
| scANVI | 0.85 | 0.98 | 0.64 | 0.65 | 0.61 | 0.69 |
| CellFM | 0.86 | 0.98 | 0.62 | 0.63 | 0.61 | 0.68 |
| UCE | 0.84 | 0.98 | 0.59 | 0.63 | 0.59 | 0.66 |
| CellPLM | 0.82 | 0.99 | 0.56 | 0.64 | 0.57 | 0.65 |
| GenePT-w | 0.83 | 0.97 | 0.54 | 0.62 | 0.56 | 0.64 |
| CELLama | 0.69 | 0.91 | 0.45 | 0.51 | 0.45 | 0.52 |
| scBERT | 0.58 | 0.82 | 0.28 | 0.32 | 0.27 | 0.37 |

**b**

Skin · reference: v1, query: v2 · shared

| method | Accuracy | $Prec_{Macro}$ | $Recall_{Macro}$ | $F1_{Macro}$ | $AUC_{ROC}$ | $Acc_{top3}$ | Aggregated Annotation |
|---|---|---|---|---|---|---|---|
| Geneformer | 0.90 | 0.69 | 0.71 | 0.67 | 0.99 | 0.99 | 0.74 |
| LangCell | 0.90 | 0.68 | 0.70 | 0.65 | 0.98 | 0.98 | 0.73 |
| scGPT | 0.87 | 0.70 | 0.70 | 0.67 | 0.98 | 0.99 | 0.73 |
| scFoundation | 0.91 | 0.69 | 0.68 | 0.66 | 0.98 | 0.99 | 0.73 |
| CellTypist | 0.90 | 0.64 | 0.71 | 0.65 | 0.99 | 0.99 | 0.73 |
| SingleR | 0.90 | 0.62 | 0.74 | 0.65 | n/a | n/a | 0.72 |
| scCello | 0.88 | 0.63 | 0.69 | 0.63 | 0.97 | 0.98 | 0.71 |
| SCimilarity | 0.89 | 0.65 | 0.66 | 0.64 | 0.98 | 0.99 | 0.71 |
| scPRINT | 0.89 | 0.64 | 0.66 | 0.63 | 0.98 | 0.99 | 0.71 |
| scVI | 0.89 | 0.61 | 0.67 | 0.63 | 0.97 | 0.99 | 0.70 |
| PCA | 0.87 | 0.62 | 0.67 | 0.62 | 0.98 | 0.99 | 0.70 |
| scANVI | 0.86 | 0.65 | 0.65 | 0.62 | 0.98 | 0.98 | 0.69 |
| CellFM | 0.87 | 0.63 | 0.63 | 0.62 | 0.98 | 0.98 | 0.69 |
| UCE | 0.85 | 0.60 | 0.63 | 0.60 | 0.97 | 0.98 | 0.67 |
| CellPLM | 0.82 | 0.57 | 0.64 | 0.58 | 0.97 | 0.99 | 0.65 |
| GenePT-w | 0.83 | 0.55 | 0.62 | 0.56 | 0.96 | 0.97 | 0.64 |
| CELLama | 0.69 | 0.45 | 0.51 | 0.45 | 0.93 | 0.91 | 0.52 |
| scBERT | 0.58 | 0.29 | 0.32 | 0.28 | 0.82 | 0.82 | 0.37 |

**Figure S213: Directional cross-dataset annotation for skin, TSv1 to TSv2. a,** All 6,486 target cells, including 20 cells from source-absent labels. **b,** 6,466 target cells from 17 shared labels. Methods and metrics are as in Fig. S185.

**a** Skin · reference: v2, query: v1 · open-world

| method | Accuracy | $Acc_{top3}$ | $Prec_{Macro}$ | $Recall_{Macro}$ | $F1_{Macro}$ | Aggregated Annotation |
|---|---|---|---|---|---|---|
| | Annotation metrics | | | | | |
| CellTypist | 0.88 | 0.97 | 0.65 | 0.72 | 0.67 | 0.73 |
| SingleR | 0.84 | n/a | 0.61 | 0.73 | 0.63 | 0.70 |
| scFoundation | 0.85 | 0.96 | 0.63 | 0.67 | 0.63 | 0.69 |
| PCA | 0.84 | 0.97 | 0.63 | 0.67 | 0.63 | 0.69 |
| scCello | 0.85 | 0.96 | 0.62 | 0.66 | 0.63 | 0.69 |
| UCE | 0.83 | 0.95 | 0.62 | 0.67 | 0.63 | 0.69 |
| SCimilarity | 0.83 | 0.97 | 0.60 | 0.66 | 0.62 | 0.68 |
| scANVI | 0.81 | 0.95 | 0.60 | 0.68 | 0.62 | 0.68 |
| CellFM | 0.82 | 0.96 | 0.60 | 0.66 | 0.62 | 0.68 |
| scVI | 0.83 | 0.96 | 0.60 | 0.65 | 0.61 | 0.67 |
| LangCell | 0.84 | 0.95 | 0.60 | 0.65 | 0.60 | 0.67 |
| CellPLM | 0.81 | 0.95 | 0.61 | 0.66 | 0.60 | 0.67 |
| Geneformer | 0.79 | 0.93 | 0.64 | 0.62 | 0.62 | 0.67 |
| scGPT | 0.80 | 0.95 | 0.61 | 0.65 | 0.60 | 0.67 |
| scPRINT | 0.78 | 0.94 | 0.65 | 0.59 | 0.59 | 0.65 |
| CELLama | 0.69 | 0.88 | 0.43 | 0.50 | 0.44 | 0.52 |
| GenePT-w | 0.61 | 0.85 | 0.49 | 0.35 | 0.30 | 0.44 |
| scBERT | 0.55 | 0.82 | 0.28 | 0.28 | 0.25 | 0.34 |

**b** Skin · reference: v2, query: v1 · shared

| method | Accuracy | $Prec_{Macro}$ | $Recall_{Macro}$ | $F1_{Macro}$ | $AUC_{ROC}$ | $Acc_{top3}$ | Aggregated Annotation |
|---|---|---|---|---|---|---|---|
| | Annotation metrics | | | | | | |
| CellTypist | 0.89 | 0.67 | 0.72 | 0.68 | 0.96 | 0.98 | 0.74 |
| SingleR | 0.85 | 0.62 | 0.73 | 0.64 | n/a | n/a | 0.71 |
| scFoundation | 0.86 | 0.64 | 0.67 | 0.64 | 0.94 | 0.97 | 0.70 |
| PCA | 0.85 | 0.63 | 0.67 | 0.64 | 0.93 | 0.98 | 0.70 |
| scCello | 0.86 | 0.63 | 0.66 | 0.64 | 0.96 | 0.97 | 0.70 |
| UCE | 0.84 | 0.63 | 0.67 | 0.64 | 0.94 | 0.96 | 0.69 |
| SCimilarity | 0.84 | 0.61 | 0.66 | 0.63 | 0.98 | 0.98 | 0.69 |
| scANVI | 0.82 | 0.62 | 0.68 | 0.63 | 0.89 | 0.96 | 0.69 |
| LangCell | 0.85 | 0.62 | 0.65 | 0.61 | 0.98 | 0.97 | 0.68 |
| CellFM | 0.83 | 0.61 | 0.66 | 0.62 | 0.94 | 0.97 | 0.68 |
| scVI | 0.84 | 0.61 | 0.65 | 0.62 | 0.94 | 0.97 | 0.68 |
| CellPLM | 0.82 | 0.62 | 0.66 | 0.61 | 0.92 | 0.96 | 0.68 |
| Geneformer | 0.80 | 0.65 | 0.62 | 0.63 | 0.95 | 0.94 | 0.68 |
| scGPT | 0.81 | 0.63 | 0.65 | 0.61 | 0.97 | 0.96 | 0.67 |
| scPRINT | 0.79 | 0.66 | 0.59 | 0.59 | 0.93 | 0.96 | 0.66 |
| CELLama | 0.70 | 0.43 | 0.50 | 0.45 | 0.90 | 0.89 | 0.52 |
| GenePT-w | 0.62 | 0.49 | 0.35 | 0.30 | 0.88 | 0.86 | 0.44 |
| scBERT | 0.56 | 0.28 | 0.28 | 0.25 | 0.72 | 0.83 | 0.34 |

**Figure S214: Directional cross-dataset annotation for skin, TSv2 to TSv1. a,** All 9,800 target cells, including 115 cells from source-absent labels. **b,** 9,685 target cells from 17 shared labels. Methods and metrics are as in Fig. S185.

**a** Small Intestine · reference: v1, query: v2 · open-world

| method | Annotation metrics | | | | | Aggregated |
|---|---|---|---|---|---|---|
| | *Accuracy* | *Acc*$_{top3}$ | *Prec*$_{Macro}$ | *Recall*$_{Macro}$ | *F1*$_{Macro}$ | *Annotation* |
| SCimilarity | 0.24 | 0.33 | 0.69 | 0.70 | 0.62 | 0.56 |
| scFoundation | 0.22 | 0.55 | 0.67 | 0.66 | 0.59 | 0.53 |
| UCE | 0.22 | 0.38 | 0.62 | 0.67 | 0.60 | 0.53 |
| SingleR | 0.24 | n/a | 0.62 | 0.67 | 0.57 | 0.52 |
| CellTypist | 0.24 | 0.60 | 0.63 | 0.65 | 0.57 | 0.52 |
| scVI | 0.22 | 0.29 | 0.61 | 0.66 | 0.55 | 0.51 |
| scCello | 0.22 | 0.30 | 0.59 | 0.65 | 0.56 | 0.51 |
| CellPLM | 0.22 | 0.28 | 0.59 | 0.65 | 0.56 | 0.51 |
| scGPT | 0.21 | 0.57 | 0.60 | 0.63 | 0.54 | 0.50 |
| PCA | 0.22 | 0.63 | 0.59 | 0.61 | 0.56 | 0.49 |
| Geneformer | 0.22 | 0.39 | 0.58 | 0.59 | 0.51 | 0.48 |
| LangCell | 0.21 | 0.40 | 0.54 | 0.60 | 0.48 | 0.46 |
| scANVI | 0.23 | 0.35 | 0.50 | 0.63 | 0.46 | 0.45 |
| GenePT-w | 0.39 | 0.62 | 0.46 | 0.52 | 0.41 | 0.44 |
| scPRINT | 0.19 | 0.29 | 0.51 | 0.55 | 0.48 | 0.43 |
| CellFM | 0.20 | 0.27 | 0.49 | 0.47 | 0.42 | 0.40 |
| CELLama | 0.21 | 0.47 | 0.34 | 0.38 | 0.30 | 0.31 |
| scBERT | 0.10 | 0.18 | 0.14 | 0.20 | 0.12 | 0.14 |

**b** Small Intestine · reference: v1, query: v2 · shared

| method | Annotation metrics | | | | | | Aggregated |
|---|---|---|---|---|---|---|---|
| | *Accuracy* | *Prec*$_{Macro}$ | *Recall*$_{Macro}$ | *F1*$_{Macro}$ | *AUC*$_{ROC}$ | *Acc*$_{top3}$ | *Annotation* |
| SCimilarity | 0.32 | 0.70 | 0.70 | 0.63 | 0.98 | 0.44 | 0.59 |
| scFoundation | 0.29 | 0.69 | 0.66 | 0.60 | 0.97 | 0.73 | 0.56 |
| UCE | 0.29 | 0.64 | 0.67 | 0.61 | 0.97 | 0.51 | 0.55 |
| SingleR | 0.31 | 0.65 | 0.67 | 0.58 | n/a | n/a | 0.55 |
| CellTypist | 0.32 | 0.65 | 0.65 | 0.58 | 0.99 | 0.81 | 0.55 |
| scVI | 0.29 | 0.63 | 0.66 | 0.56 | 0.97 | 0.38 | 0.53 |
| CellPLM | 0.30 | 0.61 | 0.65 | 0.57 | 0.98 | 0.38 | 0.53 |
| scCello | 0.30 | 0.60 | 0.65 | 0.57 | 0.97 | 0.40 | 0.53 |
| scGPT | 0.28 | 0.63 | 0.63 | 0.57 | 0.98 | 0.76 | 0.53 |
| PCA | 0.29 | 0.62 | 0.61 | 0.57 | 0.98 | 0.85 | 0.52 |
| Geneformer | 0.29 | 0.62 | 0.59 | 0.54 | 0.97 | 0.52 | 0.51 |
| scANVI | 0.31 | 0.58 | 0.63 | 0.52 | 0.97 | 0.47 | 0.51 |
| LangCell | 0.28 | 0.59 | 0.60 | 0.51 | 0.97 | 0.54 | 0.49 |
| GenePT-w | 0.52 | 0.50 | 0.52 | 0.43 | 0.96 | 0.83 | 0.49 |
| scPRINT | 0.26 | 0.53 | 0.55 | 0.50 | 0.96 | 0.39 | 0.46 |
| CellFM | 0.27 | 0.51 | 0.47 | 0.43 | 0.96 | 0.36 | 0.42 |
| CELLama | 0.28 | 0.38 | 0.38 | 0.32 | 0.90 | 0.63 | 0.34 |
| scBERT | 0.14 | 0.15 | 0.20 | 0.13 | 0.80 | 0.24 | 0.15 |

**Figure S215: Directional cross-dataset annotation for small intestine, TSv1 to TSv2. a,** All 25,834 target cells, including 6,475 cells from source-absent labels. **b,** 19,359 target cells from 31 shared labels. Methods and metrics are as in Fig. S185.

**a** Small Intestine · reference: v2, query: v1 · open-world

| method | Annotation metrics: *Accuracy* | *Acc*$_{top3}$ | *Prec*$_{Macro}$ | *Recall*$_{Macro}$ | *F1*$_{Macro}$ | Aggregated: *Annotation* |
|---|---|---|---|---|---|---|
| scFoundation | 0.72 | 0.87 | 0.73 | 0.74 | 0.70 | 0.72 |
| SCimilarity | 0.70 | 0.88 | 0.75 | 0.70 | 0.67 | 0.71 |
| CellTypist | 0.69 | 0.88 | 0.72 | 0.71 | 0.67 | 0.70 |
| Geneformer | 0.75 | 0.87 | 0.64 | 0.70 | 0.63 | 0.68 |
| scVI | 0.71 | 0.87 | 0.63 | 0.72 | 0.64 | 0.68 |
| UCE | 0.64 | 0.87 | 0.70 | 0.70 | 0.67 | 0.68 |
| PCA | 0.63 | 0.87 | 0.73 | 0.68 | 0.65 | 0.67 |
| CellPLM | 0.75 | 0.88 | 0.63 | 0.69 | 0.61 | 0.67 |
| scCello | 0.67 | 0.87 | 0.69 | 0.68 | 0.63 | 0.67 |
| SingleR | 0.59 | n/a | 0.65 | 0.72 | 0.62 | 0.65 |
| scGPT | 0.69 | 0.87 | 0.62 | 0.66 | 0.61 | 0.64 |
| scPRINT | 0.60 | 0.85 | 0.63 | 0.65 | 0.61 | 0.62 |
| LangCell | 0.60 | 0.84 | 0.59 | 0.69 | 0.58 | 0.62 |
| scANVI | 0.68 | 0.85 | 0.46 | 0.67 | 0.50 | 0.58 |
| CellFM | 0.57 | 0.83 | 0.60 | 0.60 | 0.54 | 0.58 |
| GenePT-w | 0.51 | 0.68 | 0.47 | 0.44 | 0.36 | 0.45 |
| CELLama | 0.32 | 0.62 | 0.33 | 0.51 | 0.34 | 0.37 |
| scBERT | 0.19 | 0.58 | 0.21 | 0.26 | 0.18 | 0.21 |

**b** Small Intestine · reference: v2, query: v1 · shared

| method | Annotation metrics: *Accuracy* | *Prec*$_{Macro}$ | *Recall*$_{Macro}$ | *F1*$_{Macro}$ | *AUC*$_{ROC}$ | *Acc*$_{top3}$ | Aggregated: *Annotation* |
|---|---|---|---|---|---|---|---|
| scFoundation | 0.79 | 0.76 | 0.74 | 0.71 | 0.97 | 0.96 | 0.75 |
| SCimilarity | 0.77 | 0.78 | 0.70 | 0.69 | 0.97 | 0.97 | 0.74 |
| Geneformer | 0.83 | 0.71 | 0.70 | 0.68 | 0.98 | 0.96 | 0.73 |
| CellTypist | 0.76 | 0.74 | 0.71 | 0.69 | 0.97 | 0.97 | 0.73 |
| CellPLM | 0.82 | 0.71 | 0.69 | 0.65 | 0.97 | 0.97 | 0.72 |
| scVI | 0.78 | 0.70 | 0.72 | 0.67 | 0.97 | 0.96 | 0.72 |
| UCE | 0.70 | 0.73 | 0.70 | 0.68 | 0.98 | 0.96 | 0.70 |
| scCello | 0.74 | 0.74 | 0.68 | 0.65 | 0.97 | 0.96 | 0.70 |
| PCA | 0.69 | 0.76 | 0.68 | 0.66 | 0.96 | 0.96 | 0.70 |
| SingleR | 0.65 | 0.74 | 0.72 | 0.67 | n/a | n/a | 0.70 |
| scGPT | 0.75 | 0.66 | 0.66 | 0.63 | 0.98 | 0.96 | 0.67 |
| LangCell | 0.66 | 0.65 | 0.69 | 0.63 | 0.97 | 0.93 | 0.66 |
| scPRINT | 0.66 | 0.66 | 0.65 | 0.62 | 0.96 | 0.93 | 0.65 |
| scANVI | 0.74 | 0.55 | 0.67 | 0.57 | 0.97 | 0.93 | 0.63 |
| CellFM | 0.62 | 0.66 | 0.60 | 0.57 | 0.96 | 0.91 | 0.61 |
| GenePT-w | 0.57 | 0.49 | 0.44 | 0.37 | 0.97 | 0.75 | 0.47 |
| CELLama | 0.35 | 0.39 | 0.51 | 0.37 | 0.91 | 0.68 | 0.41 |
| scBERT | 0.21 | 0.24 | 0.26 | 0.20 | 0.86 | 0.63 | 0.22 |

**Figure S216: Directional cross-dataset annotation for small intestine, TSv2 to TSv1. a,** All 10,014 target cells, including 910 cells from source-absent labels. **b,** 9,104 target cells from 31 shared labels. Methods and metrics are as in Fig. S185.

**a** Spleen · reference: v1, query: v2 · open-world

| method | Accuracy | $Acc_{top3}$ | $Prec_{Macro}$ | $Recall_{Macro}$ | $F1_{Macro}$ | Aggregated Annotation |
|---|---|---|---|---|---|---|
| | Annotation metrics | | | | | |
| SingleR | 0.86 | n/a | 0.61 | 0.79 | 0.63 | 0.72 |
| scPRINT | 0.86 | 0.98 | 0.63 | 0.74 | 0.64 | 0.72 |
| SCimilarity | 0.87 | 0.98 | 0.63 | 0.72 | 0.64 | 0.72 |
| scFoundation | 0.87 | 0.98 | 0.61 | 0.71 | 0.63 | 0.71 |
| CellPLM | 0.87 | 0.98 | 0.61 | 0.72 | 0.62 | 0.70 |
| CellTypist | 0.87 | 0.97 | 0.62 | 0.69 | 0.62 | 0.70 |
| Geneformer | 0.87 | 0.98 | 0.61 | 0.70 | 0.62 | 0.70 |
| scANVI | 0.85 | 0.97 | 0.62 | 0.70 | 0.61 | 0.70 |
| PCA | 0.86 | 0.99 | 0.61 | 0.69 | 0.62 | 0.69 |
| scGPT | 0.86 | 0.98 | 0.61 | 0.69 | 0.61 | 0.69 |
| scCello | 0.86 | 0.98 | 0.60 | 0.69 | 0.61 | 0.69 |
| scVI | 0.84 | 0.98 | 0.61 | 0.69 | 0.60 | 0.68 |
| UCE | 0.84 | 0.98 | 0.60 | 0.68 | 0.61 | 0.68 |
| LangCell | 0.86 | 0.97 | 0.58 | 0.69 | 0.59 | 0.68 |
| CellFM | 0.81 | 0.96 | 0.57 | 0.62 | 0.56 | 0.64 |
| GenePT-w | 0.78 | 0.94 | 0.53 | 0.59 | 0.49 | 0.60 |
| CELLama | 0.68 | 0.87 | 0.43 | 0.54 | 0.42 | 0.52 |
| scBERT | 0.65 | 0.86 | 0.37 | 0.40 | 0.36 | 0.44 |

**b** Spleen · reference: v1, query: v2 · shared

| method | Accuracy | $Prec_{Macro}$ | $Recall_{Macro}$ | $F1_{Macro}$ | $AUC_{ROC}$ | $Acc_{top3}$ | Aggregated Annotation |
|---|---|---|---|---|---|---|---|
| | Annotation metrics | | | | | | |
| SingleR | 0.86 | 0.61 | 0.79 | 0.63 | n/a | n/a | 0.72 |
| scPRINT | 0.86 | 0.63 | 0.74 | 0.64 | 0.99 | 0.98 | 0.72 |
| SCimilarity | 0.87 | 0.63 | 0.72 | 0.64 | 0.99 | 0.98 | 0.72 |
| scFoundation | 0.87 | 0.61 | 0.71 | 0.63 | 0.99 | 0.98 | 0.71 |
| CellPLM | 0.87 | 0.61 | 0.72 | 0.62 | 0.99 | 0.98 | 0.70 |
| CellTypist | 0.87 | 0.62 | 0.69 | 0.62 | 0.98 | 0.97 | 0.70 |
| Geneformer | 0.87 | 0.61 | 0.70 | 0.62 | 0.99 | 0.98 | 0.70 |
| scANVI | 0.85 | 0.62 | 0.70 | 0.61 | 0.98 | 0.97 | 0.70 |
| PCA | 0.86 | 0.61 | 0.69 | 0.62 | 0.99 | 0.99 | 0.69 |
| scGPT | 0.86 | 0.61 | 0.69 | 0.61 | 0.99 | 0.98 | 0.69 |
| scCello | 0.86 | 0.60 | 0.69 | 0.61 | 0.99 | 0.98 | 0.69 |
| scVI | 0.84 | 0.61 | 0.69 | 0.60 | 0.99 | 0.98 | 0.68 |
| UCE | 0.84 | 0.60 | 0.68 | 0.61 | 0.99 | 0.98 | 0.68 |
| LangCell | 0.86 | 0.58 | 0.69 | 0.60 | 0.99 | 0.97 | 0.68 |
| CellFM | 0.82 | 0.57 | 0.62 | 0.56 | 0.98 | 0.96 | 0.64 |
| GenePT-w | 0.78 | 0.53 | 0.59 | 0.49 | 0.98 | 0.94 | 0.60 |
| CELLama | 0.68 | 0.43 | 0.54 | 0.42 | 0.95 | 0.87 | 0.52 |
| scBERT | 0.65 | 0.37 | 0.40 | 0.36 | 0.90 | 0.86 | 0.44 |

**Figure S217: Directional cross-dataset annotation for spleen, TSv1 to TSv2. a,** All 30,690 target cells, including 2 cells from source-absent labels. **b,** 30,688 target cells from 23 shared labels. Methods and metrics are as in Fig. S185.

**a** Spleen · reference: v2, query: v1 · open-world

| method | Annotation metrics: Accuracy | Acc_top3 | Prec_Macro | Recall_Macro | F1_Macro | Aggregated: Annotation |
|---|---|---|---|---|---|---|
| scANVI | 0.84 | 0.97 | 0.61 | 0.75 | 0.64 | 0.71 |
| scFoundation | 0.83 | 0.97 | 0.63 | 0.72 | 0.64 | 0.70 |
| CellTypist | 0.85 | 0.97 | 0.61 | 0.68 | 0.63 | 0.69 |
| SingleR | 0.81 | n/a | 0.61 | 0.73 | 0.60 | 0.69 |
| PCA | 0.84 | 0.97 | 0.63 | 0.66 | 0.63 | 0.69 |
| UCE | 0.83 | 0.97 | 0.61 | 0.68 | 0.63 | 0.69 |
| scCello | 0.82 | 0.97 | 0.59 | 0.72 | 0.61 | 0.69 |
| scGPT | 0.81 | 0.97 | 0.62 | 0.70 | 0.62 | 0.69 |
| scVI | 0.81 | 0.97 | 0.62 | 0.71 | 0.60 | 0.68 |
| SCimilarity | 0.81 | 0.97 | 0.61 | 0.69 | 0.63 | 0.68 |
| Geneformer | 0.81 | 0.95 | 0.59 | 0.70 | 0.60 | 0.68 |
| CellPLM | 0.81 | 0.96 | 0.57 | 0.67 | 0.59 | 0.66 |
| LangCell | 0.81 | 0.95 | 0.58 | 0.65 | 0.60 | 0.66 |
| CellFM | 0.79 | 0.96 | 0.58 | 0.65 | 0.59 | 0.66 |
| scPRINT | 0.79 | 0.96 | 0.55 | 0.68 | 0.55 | 0.64 |
| GenePT-w | 0.65 | 0.83 | 0.50 | 0.47 | 0.37 | 0.50 |
| CELLama | 0.63 | 0.87 | 0.40 | 0.51 | 0.40 | 0.49 |
| scBERT | 0.53 | 0.77 | 0.33 | 0.37 | 0.32 | 0.39 |

**b** Spleen · reference: v2, query: v1 · shared

| method | Annotation metrics: Accuracy | Prec_Macro | Recall_Macro | F1_Macro | AUC_ROC | Acc_top3 | Aggregated: Annotatio |
|---|---|---|---|---|---|---|---|
| scANVI | 0.84 | 0.61 | 0.75 | 0.64 | 0.96 | 0.97 | 0.71 |
| scFoundation | 0.83 | 0.63 | 0.72 | 0.64 | 0.97 | 0.97 | 0.70 |
| CellTypist | 0.85 | 0.61 | 0.68 | 0.63 | 0.98 | 0.97 | 0.69 |
| SingleR | 0.81 | 0.61 | 0.73 | 0.60 | n/a | n/a | 0.69 |
| PCA | 0.84 | 0.63 | 0.66 | 0.63 | 0.99 | 0.97 | 0.69 |
| UCE | 0.83 | 0.61 | 0.68 | 0.63 | 0.98 | 0.97 | 0.69 |
| scCello | 0.82 | 0.59 | 0.72 | 0.61 | 0.98 | 0.97 | 0.69 |
| scGPT | 0.81 | 0.62 | 0.70 | 0.62 | 0.98 | 0.97 | 0.69 |
| scVI | 0.81 | 0.62 | 0.71 | 0.60 | 0.97 | 0.97 | 0.68 |
| SCimilarity | 0.81 | 0.61 | 0.69 | 0.63 | 0.98 | 0.97 | 0.68 |
| Geneformer | 0.81 | 0.59 | 0.70 | 0.60 | 0.96 | 0.95 | 0.68 |
| CellPLM | 0.81 | 0.57 | 0.67 | 0.59 | 0.97 | 0.96 | 0.66 |
| LangCell | 0.81 | 0.58 | 0.65 | 0.60 | 0.98 | 0.95 | 0.66 |
| CellFM | 0.79 | 0.58 | 0.65 | 0.59 | 0.97 | 0.96 | 0.66 |
| scPRINT | 0.79 | 0.55 | 0.68 | 0.55 | 0.97 | 0.96 | 0.64 |
| GenePT-w | 0.65 | 0.50 | 0.47 | 0.37 | 0.94 | 0.83 | 0.50 |
| CELLama | 0.63 | 0.40 | 0.51 | 0.40 | 0.94 | 0.87 | 0.49 |
| scBERT | 0.53 | 0.33 | 0.37 | 0.32 | 0.84 | 0.77 | 0.39 |

**Figure S218: Directional cross-dataset annotation for spleen, TSv2 to TSv1. a,b,** Both panels contain 32,711 target cells from 23 shared labels because no target label is absent from the source in this direction. Methods and metrics are as in Fig. S185.

**a** Thymus · reference: v1, query: v2 · open-world

| method | Annotation metrics: *Accuracy* | *Acc*$_{top3}$ | *Prec*$_{Macro}$ | *Recall*$_{Macro}$ | *F1*$_{Macro}$ | Aggregated: *Annotation* |
|---|---|---|---|---|---|---|
| CellTypist | 0.84 | 0.96 | 0.75 | 0.77 | 0.75 | 0.78 |
| PCA | 0.81 | 0.96 | 0.74 | 0.80 | 0.74 | 0.78 |
| scFoundation | 0.83 | 0.95 | 0.76 | 0.77 | 0.74 | 0.77 |
| SCimilarity | 0.81 | 0.94 | 0.75 | 0.80 | 0.72 | 0.77 |
| UCE | 0.81 | 0.95 | 0.76 | 0.76 | 0.74 | 0.77 |
| scANVI | 0.85 | 0.96 | 0.69 | 0.76 | 0.71 | 0.76 |
| CellPLM | 0.80 | 0.96 | 0.71 | 0.73 | 0.70 | 0.73 |
| Geneformer | 0.81 | 0.94 | 0.70 | 0.73 | 0.70 | 0.73 |
| SingleR | 0.76 | n/a | 0.69 | 0.75 | 0.69 | 0.72 |
| scGPT | 0.78 | 0.94 | 0.66 | 0.73 | 0.67 | 0.71 |
| scVI | 0.78 | 0.94 | 0.65 | 0.71 | 0.66 | 0.70 |
| scCello | 0.75 | 0.93 | 0.65 | 0.73 | 0.65 | 0.69 |
| LangCell | 0.76 | 0.93 | 0.65 | 0.66 | 0.65 | 0.68 |
| CellFM | 0.74 | 0.92 | 0.64 | 0.67 | 0.63 | 0.67 |
| scPRINT | 0.74 | 0.93 | 0.62 | 0.65 | 0.61 | 0.66 |
| GenePT-w | 0.70 | 0.92 | 0.58 | 0.66 | 0.58 | 0.63 |
| CELLama | 0.62 | 0.86 | 0.44 | 0.55 | 0.44 | 0.51 |
| scBERT | 0.26 | 0.58 | 0.27 | 0.34 | 0.23 | 0.28 |

**b** Thymus · reference: v1, query: v2 · shared

| method | Annotation metrics: *Accuracy* | *Prec*$_{Macro}$ | *Recall*$_{Macro}$ | *F1*$_{Macro}$ | *AUC*$_{ROC}$ | *Acc*$_{top3}$ | Aggregated: *Annotation* |
|---|---|---|---|---|---|---|---|
| SCimilarity | 0.82 | 0.78 | 0.80 | 0.74 | 0.98 | 0.95 | 0.79 |
| scFoundation | 0.84 | 0.78 | 0.77 | 0.76 | 0.96 | 0.96 | 0.79 |
| CellTypist | 0.85 | 0.76 | 0.77 | 0.76 | 0.98 | 0.97 | 0.78 |
| PCA | 0.82 | 0.75 | 0.80 | 0.75 | 0.98 | 0.97 | 0.78 |
| UCE | 0.82 | 0.77 | 0.76 | 0.74 | 0.98 | 0.96 | 0.77 |
| scANVI | 0.86 | 0.70 | 0.76 | 0.72 | 0.98 | 0.97 | 0.76 |
| CellPLM | 0.81 | 0.72 | 0.73 | 0.70 | 0.97 | 0.97 | 0.74 |
| Geneformer | 0.81 | 0.71 | 0.73 | 0.70 | 0.96 | 0.95 | 0.74 |
| SingleR | 0.77 | 0.71 | 0.75 | 0.69 | n/a | n/a | 0.73 |
| scGPT | 0.78 | 0.68 | 0.73 | 0.69 | 0.98 | 0.95 | 0.72 |
| scVI | 0.78 | 0.68 | 0.71 | 0.68 | 0.97 | 0.95 | 0.71 |
| scCello | 0.76 | 0.66 | 0.73 | 0.66 | 0.98 | 0.94 | 0.70 |
| LangCell | 0.77 | 0.67 | 0.66 | 0.66 | 0.97 | 0.94 | 0.69 |
| CellFM | 0.74 | 0.65 | 0.67 | 0.63 | 0.96 | 0.93 | 0.67 |
| scPRINT | 0.75 | 0.63 | 0.65 | 0.62 | 0.97 | 0.94 | 0.66 |
| GenePT-w | 0.71 | 0.59 | 0.66 | 0.59 | 0.97 | 0.92 | 0.64 |
| CELLama | 0.62 | 0.44 | 0.55 | 0.45 | 0.94 | 0.86 | 0.52 |
| scBERT | 0.27 | 0.28 | 0.34 | 0.23 | 0.82 | 0.58 | 0.28 |

**Figure S219: Directional cross-dataset annotation for thymus, TSv1 to TSv2. a,** All 9,933 target cells, including 92 cells from source-absent labels. **b,** 9,841 target cells from 26 shared labels. Methods and metrics are as in Fig. S185.

**a** Thymus · reference: v2, query: v1 · open-world

| method | Annotation metrics: *Accuracy* | *Acc*$_{top3}$ | *Prec*$_{Macro}$ | *Recall*$_{Macro}$ | *F1*$_{Macro}$ | Aggregated: *Annotation* |
|---|---|---|---|---|---|---|
| CellTypist | 0.72 | 0.89 | 0.77 | 0.73 | 0.70 | 0.73 |
| scFoundation | 0.72 | 0.87 | 0.70 | 0.74 | 0.68 | 0.71 |
| SCimilarity | 0.70 | 0.91 | 0.76 | 0.70 | 0.67 | 0.71 |
| UCE | 0.72 | 0.86 | 0.69 | 0.73 | 0.67 | 0.70 |
| PCA | 0.69 | 0.87 | 0.72 | 0.69 | 0.64 | 0.68 |
| scANVI | 0.70 | 0.87 | 0.63 | 0.71 | 0.61 | 0.67 |
| scVI | 0.69 | 0.89 | 0.64 | 0.71 | 0.62 | 0.66 |
| scCello | 0.68 | 0.90 | 0.68 | 0.67 | 0.61 | 0.66 |
| SingleR | 0.60 | n/a | 0.72 | 0.67 | 0.63 | 0.65 |
| Geneformer | 0.70 | 0.86 | 0.66 | 0.64 | 0.61 | 0.65 |
| scGPT | 0.67 | 0.87 | 0.65 | 0.67 | 0.60 | 0.65 |
| LangCell | 0.65 | 0.85 | 0.64 | 0.62 | 0.58 | 0.62 |
| scPRINT | 0.64 | 0.84 | 0.63 | 0.64 | 0.58 | 0.62 |
| CellPLM | 0.64 | 0.86 | 0.63 | 0.61 | 0.56 | 0.61 |
| CellFM | 0.65 | 0.85 | 0.62 | 0.60 | 0.55 | 0.61 |
| GenePT-w | 0.53 | 0.75 | 0.52 | 0.50 | 0.37 | 0.48 |
| CELLama | 0.52 | 0.80 | 0.47 | 0.48 | 0.42 | 0.47 |
| scBERT | 0.31 | 0.58 | 0.27 | 0.26 | 0.19 | 0.26 |

**b** Thymus · reference: v2, query: v1 · shared

| method | Annotation metrics: *Accuracy* | *Prec*$_{Macro}$ | *Recall*$_{Macro}$ | *F1*$_{Macro}$ | *AUC*$_{ROC}$ | *Acc*$_{top3}$ | Aggregated: *Annotation* |
|---|---|---|---|---|---|---|---|
| CellTypist | 0.72 | 0.78 | 0.73 | 0.70 | 0.98 | 0.90 | 0.73 |
| scFoundation | 0.73 | 0.73 | 0.74 | 0.69 | 0.96 | 0.87 | 0.72 |
| SCimilarity | 0.71 | 0.77 | 0.70 | 0.67 | 0.97 | 0.92 | 0.71 |
| UCE | 0.72 | 0.71 | 0.73 | 0.68 | 0.96 | 0.87 | 0.71 |
| PCA | 0.69 | 0.74 | 0.69 | 0.64 | 0.97 | 0.88 | 0.69 |
| scANVI | 0.70 | 0.65 | 0.71 | 0.63 | 0.93 | 0.87 | 0.67 |
| scCello | 0.68 | 0.71 | 0.67 | 0.63 | 0.96 | 0.91 | 0.67 |
| scVI | 0.69 | 0.65 | 0.71 | 0.63 | 0.92 | 0.90 | 0.67 |
| SingleR | 0.60 | 0.75 | 0.67 | 0.64 | n/a | n/a | 0.66 |
| Geneformer | 0.71 | 0.66 | 0.64 | 0.61 | 0.95 | 0.86 | 0.66 |
| scGPT | 0.68 | 0.66 | 0.67 | 0.61 | 0.96 | 0.88 | 0.65 |
| scPRINT | 0.65 | 0.64 | 0.64 | 0.58 | 0.95 | 0.85 | 0.63 |
| LangCell | 0.65 | 0.65 | 0.62 | 0.58 | 0.96 | 0.85 | 0.63 |
| CellPLM | 0.65 | 0.64 | 0.61 | 0.57 | 0.89 | 0.87 | 0.62 |
| CellFM | 0.66 | 0.64 | 0.60 | 0.56 | 0.95 | 0.86 | 0.61 |
| GenePT-w | 0.54 | 0.52 | 0.50 | 0.37 | 0.96 | 0.76 | 0.48 |
| CELLama | 0.52 | 0.47 | 0.48 | 0.42 | 0.93 | 0.80 | 0.47 |
| scBERT | 0.32 | 0.28 | 0.26 | 0.20 | 0.83 | 0.58 | 0.26 |

**Figure S220: Directional cross-dataset annotation for thymus, TSv2 to TSv1. a,** All 29,079 target cells, including 234 cells from source-absent labels. **b,** 28,845 target cells from 26 shared labels. Methods and metrics are as in Fig. S185.

**a** Tongue · reference: v1, query: v2 · open-world

| method | Annotation metrics: *Accuracy* | *Acc*$_{top3}$ | *Prec*$_{Macro}$ | *Recall*$_{Macro}$ | *F1*$_{Macro}$ | Aggregated: *Annotation* |
|---|---|---|---|---|---|---|
| CellPLM | 0.85 | 0.96 | 0.67 | 0.74 | 0.69 | 0.74 |
| SingleR | 0.83 | n/a | 0.65 | 0.78 | 0.66 | 0.73 |
| SCimilarity | 0.83 | 0.96 | 0.61 | 0.74 | 0.62 | 0.70 |
| Geneformer | 0.81 | 0.91 | 0.62 | 0.73 | 0.63 | 0.70 |
| scGPT | 0.81 | 0.95 | 0.64 | 0.70 | 0.61 | 0.69 |
| scPRINT | 0.83 | 0.95 | 0.64 | 0.68 | 0.62 | 0.69 |
| scFoundation | 0.80 | 0.93 | 0.62 | 0.72 | 0.62 | 0.69 |
| CellTypist | 0.85 | 0.95 | 0.63 | 0.63 | 0.61 | 0.68 |
| scVI | 0.82 | 0.95 | 0.60 | 0.67 | 0.60 | 0.67 |
| scCello | 0.81 | 0.94 | 0.61 | 0.66 | 0.59 | 0.67 |
| PCA | 0.84 | 0.95 | 0.61 | 0.59 | 0.56 | 0.65 |
| UCE | 0.78 | 0.93 | 0.63 | 0.61 | 0.57 | 0.65 |
| LangCell | 0.78 | 0.94 | 0.60 | 0.64 | 0.57 | 0.65 |
| scANVI | 0.86 | 0.95 | 0.57 | 0.62 | 0.54 | 0.65 |
| CellFM | 0.77 | 0.94 | 0.54 | 0.54 | 0.49 | 0.59 |
| GenePT-w | 0.59 | 0.86 | 0.54 | 0.54 | 0.46 | 0.53 |
| CELLama | 0.61 | 0.84 | 0.43 | 0.40 | 0.38 | 0.46 |
| scBERT | 0.62 | 0.81 | 0.26 | 0.25 | 0.20 | 0.33 |

**b** Tongue · reference: v1, query: v2 · shared

| method | Annotation metrics: *Accuracy* | *Prec*$_{Macro}$ | *Recall*$_{Macro}$ | *F1*$_{Macro}$ | *AUC*$_{ROC}$ | *Acc*$_{top3}$ | Aggregated: *Annotation* |
|---|---|---|---|---|---|---|---|
| CellPLM | 0.85 | 0.67 | 0.74 | 0.69 | 0.98 | 0.96 | 0.74 |
| SingleR | 0.83 | 0.65 | 0.78 | 0.66 | n/a | n/a | 0.73 |
| SCimilarity | 0.83 | 0.61 | 0.74 | 0.62 | 0.95 | 0.96 | 0.70 |
| Geneformer | 0.81 | 0.62 | 0.73 | 0.63 | 0.97 | 0.92 | 0.70 |
| scGPT | 0.81 | 0.64 | 0.70 | 0.61 | 0.96 | 0.95 | 0.69 |
| scPRINT | 0.83 | 0.64 | 0.68 | 0.62 | 0.96 | 0.95 | 0.69 |
| scFoundation | 0.80 | 0.62 | 0.72 | 0.62 | 0.96 | 0.93 | 0.69 |
| CellTypist | 0.85 | 0.63 | 0.63 | 0.61 | 0.95 | 0.95 | 0.68 |
| scVI | 0.82 | 0.60 | 0.67 | 0.60 | 0.95 | 0.95 | 0.67 |
| scCello | 0.81 | 0.62 | 0.66 | 0.59 | 0.96 | 0.94 | 0.67 |
| PCA | 0.84 | 0.61 | 0.59 | 0.56 | 0.98 | 0.95 | 0.65 |
| UCE | 0.78 | 0.63 | 0.61 | 0.57 | 0.95 | 0.93 | 0.65 |
| LangCell | 0.79 | 0.60 | 0.64 | 0.57 | 0.95 | 0.95 | 0.65 |
| scANVI | 0.86 | 0.57 | 0.62 | 0.54 | 0.96 | 0.95 | 0.65 |
| CellFM | 0.77 | 0.54 | 0.54 | 0.49 | 0.96 | 0.94 | 0.59 |
| GenePT-w | 0.59 | 0.55 | 0.54 | 0.46 | 0.94 | 0.86 | 0.53 |
| CELLama | 0.61 | 0.43 | 0.40 | 0.38 | 0.86 | 0.85 | 0.46 |
| scBERT | 0.62 | 0.27 | 0.25 | 0.20 | 0.74 | 0.82 | 0.33 |

**Figure S221: Directional cross-dataset annotation for tongue, TSv1 to TSv2. a,** All 21,499 target cells, including 18 cells from source-absent labels. **b,** 21,481 target cells from 21 shared labels. Methods and metrics are as in Fig. S185.

**a** Tongue · reference: v2, query: v1 · open-world

| method | Annotation metrics | | | | | Aggregated |
|---|---|---|---|---|---|---|
| | *Accuracy* | $Acc_{top3}$ | $Prec_{Macro}$ | $Recall_{Macro}$ | $F1_{Macro}$ | *Annotation* |
| CellTypist | 0.81 | 0.95 | 0.62 | 0.67 | 0.61 | 0.68 |
| PCA | 0.78 | 0.95 | 0.64 | 0.62 | 0.59 | 0.66 |
| scFoundation | 0.79 | 0.95 | 0.66 | 0.61 | 0.57 | 0.66 |
| scGPT | 0.77 | 0.93 | 0.63 | 0.62 | 0.60 | 0.65 |
| SCimilarity | 0.75 | 0.95 | 0.61 | 0.65 | 0.58 | 0.65 |
| SingleR | 0.75 | n/a | 0.62 | 0.65 | 0.58 | 0.65 |
| Geneformer | 0.76 | 0.89 | 0.59 | 0.61 | 0.58 | 0.64 |
| CellFM | 0.76 | 0.95 | 0.56 | 0.64 | 0.58 | 0.64 |
| UCE | 0.76 | 0.92 | 0.60 | 0.60 | 0.57 | 0.63 |
| scCello | 0.71 | 0.95 | 0.60 | 0.60 | 0.58 | 0.62 |
| LangCell | 0.71 | 0.88 | 0.56 | 0.58 | 0.52 | 0.59 |
| scPRINT | 0.70 | 0.90 | 0.58 | 0.51 | 0.50 | 0.57 |
| CellPLM | 0.69 | 0.94 | 0.46 | 0.58 | 0.48 | 0.55 |
| scANVI | 0.70 | 0.88 | 0.36 | 0.64 | 0.41 | 0.53 |
| scVI | 0.68 | 0.90 | 0.44 | 0.48 | 0.38 | 0.50 |
| GenePT-w | 0.63 | 0.91 | 0.52 | 0.40 | 0.34 | 0.47 |
| CELLama | 0.61 | 0.84 | 0.30 | 0.39 | 0.32 | 0.41 |
| scBERT | 0.44 | 0.63 | 0.24 | 0.25 | 0.16 | 0.27 |

**b** Tongue · reference: v2, query: v1 · shared

| method | Annotation metrics | | | | | | Aggregated |
|---|---|---|---|---|---|---|---|
| | *Accuracy* | $Prec_{Macro}$ | $Recall_{Macro}$ | $F1_{Macro}$ | $AUC_{ROC}$ | $Acc_{top3}$ | *Annotation* |
| CellTypist | 0.82 | 0.64 | 0.67 | 0.62 | 0.94 | 0.97 | 0.69 |
| PCA | 0.79 | 0.64 | 0.62 | 0.59 | 0.95 | 0.96 | 0.66 |
| scFoundation | 0.79 | 0.66 | 0.61 | 0.57 | 0.93 | 0.96 | 0.66 |
| scGPT | 0.78 | 0.63 | 0.62 | 0.60 | 0.95 | 0.95 | 0.66 |
| SCimilarity | 0.76 | 0.63 | 0.65 | 0.59 | 0.95 | 0.97 | 0.66 |
| SingleR | 0.76 | 0.62 | 0.65 | 0.58 | n/a | n/a | 0.65 |
| CellFM | 0.77 | 0.58 | 0.64 | 0.59 | 0.95 | 0.96 | 0.64 |
| Geneformer | 0.77 | 0.60 | 0.61 | 0.59 | 0.95 | 0.90 | 0.64 |
| UCE | 0.77 | 0.61 | 0.60 | 0.57 | 0.94 | 0.93 | 0.64 |
| scCello | 0.72 | 0.61 | 0.60 | 0.58 | 0.95 | 0.96 | 0.63 |
| LangCell | 0.72 | 0.57 | 0.58 | 0.53 | 0.94 | 0.89 | 0.60 |
| scPRINT | 0.71 | 0.59 | 0.51 | 0.50 | 0.92 | 0.91 | 0.58 |
| CellPLM | 0.70 | 0.48 | 0.58 | 0.49 | 0.94 | 0.95 | 0.56 |
| scANVI | 0.71 | 0.37 | 0.64 | 0.42 | 0.91 | 0.89 | 0.53 |
| scVI | 0.69 | 0.45 | 0.48 | 0.38 | 0.91 | 0.91 | 0.50 |
| GenePT-w | 0.64 | 0.55 | 0.40 | 0.34 | 0.93 | 0.92 | 0.48 |
| CELLama | 0.62 | 0.31 | 0.39 | 0.32 | 0.89 | 0.85 | 0.41 |
| scBERT | 0.44 | 0.24 | 0.25 | 0.16 | 0.79 | 0.64 | 0.27 |

**Figure S222: Directional cross-dataset annotation for tongue, TSv2 to TSv1. a,** All 11,928 target cells, including 139 cells from source-absent labels. **b,** 11,789 target cells from 21 shared labels. Methods and metrics are as in Fig. S185.

**a** Trachea · reference: v1, query: v2 · open-world

| method | Annotation metrics: *Accuracy* | $Acc_{top3}$ | $Prec_{Macro}$ | $Recall_{Macro}$ | $F1_{Macro}$ | Aggregated: *Annotation* |
|---|---|---|---|---|---|---|
| Geneformer | 0.94 | 0.99 | 0.67 | 0.78 | 0.70 | 0.77 |
| scPRINT | 0.91 | 0.99 | 0.66 | 0.77 | 0.67 | 0.75 |
| SCimilarity | 0.89 | 0.99 | 0.66 | 0.77 | 0.68 | 0.75 |
| scFoundation | 0.91 | 0.98 | 0.66 | 0.78 | 0.66 | 0.75 |
| scGPT | 0.92 | 0.99 | 0.65 | 0.75 | 0.66 | 0.75 |
| PCA | 0.92 | 0.98 | 0.67 | 0.73 | 0.64 | 0.74 |
| LangCell | 0.91 | 0.98 | 0.64 | 0.74 | 0.65 | 0.73 |
| scANVI | 0.90 | 0.98 | 0.62 | 0.74 | 0.64 | 0.73 |
| CellTypist | 0.93 | 0.99 | 0.62 | 0.70 | 0.62 | 0.72 |
| CellPLM | 0.90 | 0.98 | 0.63 | 0.70 | 0.61 | 0.71 |
| UCE | 0.86 | 0.98 | 0.62 | 0.73 | 0.63 | 0.71 |
| scCello | 0.80 | 0.97 | 0.62 | 0.77 | 0.62 | 0.70 |
| scVI | 0.82 | 0.97 | 0.64 | 0.71 | 0.62 | 0.70 |
| CellFM | 0.92 | 0.98 | 0.60 | 0.68 | 0.59 | 0.70 |
| SingleR | 0.76 | n/a | 0.64 | 0.75 | 0.60 | 0.69 |
| CELLama | 0.76 | 0.94 | 0.44 | 0.55 | 0.44 | 0.54 |
| scBERT | 0.84 | 0.95 | 0.38 | 0.44 | 0.37 | 0.51 |
| GenePT-w | 0.46 | 0.77 | 0.52 | 0.46 | 0.36 | 0.45 |

**b** Trachea · reference: v1, query: v2 · shared

| method | Annotation metrics: *Accuracy* | $Prec_{Macro}$ | $Recall_{Macro}$ | $F1_{Macro}$ | $AUC_{ROC}$ | $Acc_{top3}$ | Aggregated: *Annotation* |
|---|---|---|---|---|---|---|---|
| Geneformer | 0.94 | 0.68 | 0.78 | 0.70 | 0.99 | 0.99 | 0.77 |
| scPRINT | 0.91 | 0.66 | 0.77 | 0.67 | 0.99 | 0.99 | 0.75 |
| scFoundation | 0.91 | 0.66 | 0.78 | 0.66 | 0.99 | 0.99 | 0.75 |
| SCimilarity | 0.90 | 0.67 | 0.77 | 0.68 | 0.99 | 0.99 | 0.75 |
| scGPT | 0.93 | 0.66 | 0.75 | 0.66 | 0.99 | 0.99 | 0.75 |
| PCA | 0.92 | 0.68 | 0.73 | 0.64 | 0.99 | 0.98 | 0.74 |
| LangCell | 0.91 | 0.64 | 0.74 | 0.65 | 0.98 | 0.98 | 0.73 |
| scANVI | 0.90 | 0.63 | 0.74 | 0.65 | 0.98 | 0.99 | 0.73 |
| CellTypist | 0.93 | 0.62 | 0.70 | 0.62 | 0.99 | 0.99 | 0.72 |
| CellPLM | 0.90 | 0.63 | 0.70 | 0.61 | 0.99 | 0.99 | 0.71 |
| UCE | 0.86 | 0.63 | 0.73 | 0.63 | 0.99 | 0.98 | 0.71 |
| scCello | 0.80 | 0.62 | 0.77 | 0.63 | 0.98 | 0.97 | 0.70 |
| CellFM | 0.92 | 0.61 | 0.68 | 0.59 | 0.99 | 0.98 | 0.70 |
| scVI | 0.82 | 0.65 | 0.71 | 0.62 | 0.99 | 0.97 | 0.70 |
| SingleR | 0.76 | 0.65 | 0.75 | 0.60 | n/a | n/a | 0.69 |
| CELLama | 0.76 | 0.44 | 0.55 | 0.44 | 0.96 | 0.94 | 0.55 |
| scBERT | 0.84 | 0.38 | 0.44 | 0.37 | 0.93 | 0.95 | 0.51 |
| GenePT-w | 0.46 | 0.52 | 0.46 | 0.36 | 0.92 | 0.77 | 0.45 |

**Figure S223: Directional cross-dataset annotation for trachea, TSv1 to TSv2. a,** All 5,346 target cells, including 9 cells from source-absent labels. **b,** 5,337 target cells from 22 shared labels. Methods and metrics are as in Fig. S185.

**a** Trachea · reference: v2, query: v1 · open-world

| method | Accuracy | $Acc_{top3}$ | $Prec_{Macro}$ | $Recall_{Macro}$ | $F1_{Macro}$ | Aggregated Annotation |
|---|---|---|---|---|---|---|
| CellTypist | 0.94 | 0.99 | 0.73 | 0.68 | 0.67 | 0.75 |
| scCello | 0.93 | 0.99 | 0.69 | 0.71 | 0.69 | 0.75 |
| SingleR | 0.91 | n/a | 0.69 | 0.74 | 0.67 | 0.75 |
| SCimilarity | 0.92 | 0.98 | 0.68 | 0.72 | 0.69 | 0.75 |
| PCA | 0.93 | 0.99 | 0.69 | 0.68 | 0.67 | 0.75 |
| UCE | 0.92 | 0.98 | 0.67 | 0.71 | 0.66 | 0.74 |
| scFoundation | 0.92 | 0.98 | 0.69 | 0.68 | 0.65 | 0.74 |
| LangCell | 0.92 | 0.98 | 0.69 | 0.68 | 0.64 | 0.73 |
| Geneformer | 0.89 | 0.98 | 0.67 | 0.70 | 0.64 | 0.73 |
| scVI | 0.91 | 0.97 | 0.65 | 0.66 | 0.64 | 0.71 |
| scGPT | 0.89 | 0.98 | 0.64 | 0.68 | 0.64 | 0.71 |
| scPRINT | 0.88 | 0.98 | 0.62 | 0.66 | 0.61 | 0.69 |
| scANVI | 0.87 | 0.97 | 0.60 | 0.69 | 0.61 | 0.69 |
| CellPLM | 0.87 | 0.97 | 0.59 | 0.62 | 0.59 | 0.67 |
| CellFM | 0.93 | 0.99 | 0.63 | 0.56 | 0.55 | 0.67 |
| CELLama | 0.81 | 0.94 | 0.48 | 0.54 | 0.48 | 0.58 |
| GenePT-w | 0.62 | 0.84 | 0.53 | 0.33 | 0.28 | 0.44 |
| scBERT | 0.67 | 0.87 | 0.30 | 0.32 | 0.29 | 0.39 |

**b** Trachea · reference: v2, query: v1 · shared

| method | Accuracy | $Prec_{Macro}$ | $Recall_{Macro}$ | $F1_{Macro}$ | $AUC_{ROC}$ | $Acc_{top3}$ | Aggregated Annotation |
|---|---|---|---|---|---|---|---|
| scCello | 0.94 | 0.69 | 0.71 | 0.69 | 0.98 | 0.99 | 0.76 |
| CellTypist | 0.94 | 0.73 | 0.68 | 0.67 | 0.99 | 0.99 | 0.76 |
| SingleR | 0.91 | 0.69 | 0.74 | 0.67 | n/a | n/a | 0.75 |
| SCimilarity | 0.92 | 0.68 | 0.72 | 0.69 | 0.99 | 0.98 | 0.75 |
| PCA | 0.93 | 0.70 | 0.68 | 0.67 | 0.99 | 0.99 | 0.75 |
| UCE | 0.92 | 0.67 | 0.71 | 0.67 | 0.98 | 0.98 | 0.74 |
| scFoundation | 0.93 | 0.69 | 0.68 | 0.65 | 0.98 | 0.99 | 0.74 |
| LangCell | 0.92 | 0.69 | 0.68 | 0.64 | 0.98 | 0.99 | 0.73 |
| Geneformer | 0.90 | 0.67 | 0.70 | 0.65 | 0.97 | 0.98 | 0.73 |
| scVI | 0.91 | 0.65 | 0.66 | 0.64 | 0.97 | 0.97 | 0.71 |
| scGPT | 0.89 | 0.64 | 0.68 | 0.64 | 0.98 | 0.98 | 0.71 |
| scPRINT | 0.88 | 0.62 | 0.66 | 0.61 | 0.97 | 0.98 | 0.69 |
| scANVI | 0.87 | 0.60 | 0.69 | 0.61 | 0.94 | 0.97 | 0.69 |
| CellPLM | 0.87 | 0.59 | 0.62 | 0.59 | 0.96 | 0.97 | 0.67 |
| CellFM | 0.93 | 0.63 | 0.56 | 0.55 | 0.97 | 0.99 | 0.67 |
| CELLama | 0.82 | 0.48 | 0.54 | 0.48 | 0.96 | 0.94 | 0.58 |
| GenePT-w | 0.62 | 0.53 | 0.33 | 0.28 | 0.97 | 0.84 | 0.44 |
| scBERT | 0.67 | 0.30 | 0.32 | 0.29 | 0.88 | 0.88 | 0.39 |

**Figure S224: Directional cross-dataset annotation for trachea, TSv2 to TSv1.** **a,** All 9,227 target cells, including 15 cells from source-absent labels. **b,** 9,212 target cells from 22 shared labels. Methods and metrics are as in Fig. S185.

**a** Uterus · reference: v1, query: v2 · open-world

| method | Accuracy | $Acc_{top3}$ | $Prec_{Macro}$ | $Recall_{Macro}$ | $F1_{Macro}$ | Aggregated Annotation |
|---|---|---|---|---|---|---|
| | Annotation metrics | | | | | |
| scFoundation | 0.83 | 0.97 | 0.61 | 0.65 | 0.60 | 0.67 |
| scGPT | 0.83 | 0.98 | 0.66 | 0.63 | 0.57 | 0.67 |
| UCE | 0.81 | 0.96 | 0.63 | 0.63 | 0.60 | 0.67 |
| CellTypist | 0.86 | 0.97 | 0.62 | 0.62 | 0.57 | 0.67 |
| SingleR | 0.85 | n/a | 0.63 | 0.64 | 0.56 | 0.67 |
| Geneformer | 0.81 | 0.96 | 0.62 | 0.63 | 0.58 | 0.66 |
| LangCell | 0.82 | 0.96 | 0.59 | 0.61 | 0.57 | 0.65 |
| scVI | 0.82 | 0.99 | 0.55 | 0.63 | 0.56 | 0.64 |
| SCimilarity | 0.81 | 0.99 | 0.57 | 0.62 | 0.55 | 0.64 |
| CellPLM | 0.80 | 0.97 | 0.60 | 0.61 | 0.54 | 0.64 |
| scCello | 0.79 | 0.96 | 0.56 | 0.61 | 0.56 | 0.63 |
| scPRINT | 0.79 | 0.96 | 0.58 | 0.60 | 0.53 | 0.62 |
| PCA | 0.82 | 0.98 | 0.54 | 0.59 | 0.53 | 0.62 |
| scANVI | 0.82 | 0.95 | 0.55 | 0.56 | 0.51 | 0.61 |
| CellFM | 0.79 | 0.97 | 0.51 | 0.54 | 0.49 | 0.58 |
| GenePT-w | 0.74 | 0.96 | 0.50 | 0.58 | 0.47 | 0.57 |
| CELLama | 0.59 | 0.86 | 0.34 | 0.41 | 0.34 | 0.42 |
| scBERT | 0.46 | 0.71 | 0.28 | 0.29 | 0.26 | 0.32 |

**b** Uterus · reference: v1, query: v2 · shared

| method | Accuracy | $Prec_{Macro}$ | $Recall_{Macro}$ | $F1_{Macro}$ | $AUC_{ROC}$ | $Acc_{top3}$ | Aggregated Annotation |
|---|---|---|---|---|---|---|---|
| | Annotation metrics | | | | | | |
| scFoundation | 0.84 | 0.61 | 0.65 | 0.60 | 0.98 | 0.97 | 0.67 |
| scGPT | 0.83 | 0.66 | 0.63 | 0.57 | 0.98 | 0.98 | 0.67 |
| UCE | 0.81 | 0.63 | 0.63 | 0.60 | 0.98 | 0.96 | 0.67 |
| CellTypist | 0.86 | 0.62 | 0.62 | 0.58 | 0.98 | 0.97 | 0.67 |
| SingleR | 0.85 | 0.63 | 0.64 | 0.56 | n/a | n/a | 0.67 |
| Geneformer | 0.81 | 0.64 | 0.63 | 0.58 | 0.97 | 0.96 | 0.67 |
| scVI | 0.82 | 0.58 | 0.63 | 0.57 | 0.97 | 0.99 | 0.65 |
| LangCell | 0.82 | 0.59 | 0.61 | 0.57 | 0.98 | 0.96 | 0.65 |
| CellPLM | 0.80 | 0.60 | 0.61 | 0.54 | 0.99 | 0.97 | 0.64 |
| SCimilarity | 0.81 | 0.57 | 0.62 | 0.55 | 0.98 | 0.99 | 0.64 |
| scCello | 0.79 | 0.56 | 0.61 | 0.56 | 0.97 | 0.97 | 0.63 |
| scPRINT | 0.79 | 0.58 | 0.60 | 0.53 | 0.98 | 0.97 | 0.62 |
| PCA | 0.82 | 0.55 | 0.59 | 0.53 | 0.96 | 0.98 | 0.62 |
| scANVI | 0.82 | 0.55 | 0.56 | 0.51 | 0.97 | 0.96 | 0.61 |
| CellFM | 0.80 | 0.51 | 0.54 | 0.50 | 0.93 | 0.97 | 0.59 |
| GenePT-w | 0.75 | 0.50 | 0.58 | 0.47 | 0.98 | 0.96 | 0.57 |
| CELLama | 0.60 | 0.34 | 0.41 | 0.34 | 0.87 | 0.86 | 0.42 |
| scBERT | 0.46 | 0.28 | 0.29 | 0.26 | 0.77 | 0.71 | 0.32 |

**Figure S225: Directional cross-dataset annotation for uterus, TSv1 to TSv2. a,** All 10,580 target cells, including 30 cells from source-absent labels. **b,** 10,550 target cells from 17 shared labels. Methods and metrics are as in Fig. S185.

**a** Uterus · reference: v2, query: v1 · open-world

| method | Annotation metrics | | | | | Aggregated |
|---|---|---|---|---|---|---|
| | *Accuracy* | $Acc_{top3}$ | $Prec_{Macro}$ | $Recall_{Macro}$ | $F1_{Macro}$ | *Annotation* |
| CellTypist | 0.90 | 0.97 | 0.61 | 0.59 | 0.58 | 0.67 |
| PCA | 0.88 | 0.97 | 0.61 | 0.56 | 0.57 | 0.66 |
| UCE | 0.85 | 0.96 | 0.57 | 0.62 | 0.56 | 0.65 |
| SCimilarity | 0.86 | 0.97 | 0.59 | 0.58 | 0.57 | 0.65 |
| scCello | 0.85 | 0.98 | 0.57 | 0.56 | 0.54 | 0.63 |
| scGPT | 0.85 | 0.97 | 0.55 | 0.58 | 0.53 | 0.63 |
| scVI | 0.84 | 0.97 | 0.56 | 0.55 | 0.51 | 0.62 |
| scFoundation | 0.83 | 0.97 | 0.55 | 0.54 | 0.50 | 0.60 |
| CellFM | 0.85 | 0.96 | 0.52 | 0.51 | 0.49 | 0.59 |
| LangCell | 0.84 | 0.96 | 0.53 | 0.51 | 0.48 | 0.59 |
| scANVI | 0.80 | 0.95 | 0.47 | 0.60 | 0.48 | 0.59 |
| CellPLM | 0.74 | 0.94 | 0.51 | 0.59 | 0.49 | 0.58 |
| SingleR | 0.79 | n/a | 0.52 | 0.57 | 0.45 | 0.58 |
| Geneformer | 0.80 | 0.93 | 0.46 | 0.52 | 0.45 | 0.56 |
| scPRINT | 0.76 | 0.94 | 0.49 | 0.50 | 0.46 | 0.55 |
| GenePT-w | 0.55 | 0.87 | 0.50 | 0.39 | 0.32 | 0.44 |
| CELLama | 0.66 | 0.89 | 0.31 | 0.38 | 0.32 | 0.42 |
| scBERT | 0.57 | 0.81 | 0.29 | 0.30 | 0.26 | 0.35 |

**b** Uterus · reference: v2, query: v1 · shared

| method | Annotation metrics | | | | | | Aggregated |
|---|---|---|---|---|---|---|---|
| | *Accuracy* | $Prec_{Macro}$ | $Recall_{Macro}$ | $F1_{Macro}$ | $AUC_{ROC}$ | $Acc_{top3}$ | *Annotation* |
| CellTypist | 0.90 | 0.63 | 0.59 | 0.59 | 0.98 | 0.98 | 0.68 |
| PCA | 0.89 | 0.63 | 0.56 | 0.58 | 0.99 | 0.98 | 0.66 |
| UCE | 0.85 | 0.58 | 0.62 | 0.56 | 0.95 | 0.97 | 0.65 |
| SCimilarity | 0.86 | 0.60 | 0.58 | 0.57 | 0.97 | 0.98 | 0.65 |
| scCello | 0.85 | 0.58 | 0.56 | 0.55 | 0.97 | 0.98 | 0.64 |
| scGPT | 0.86 | 0.55 | 0.58 | 0.53 | 0.97 | 0.97 | 0.63 |
| scVI | 0.85 | 0.56 | 0.55 | 0.52 | 0.97 | 0.97 | 0.62 |
| scFoundation | 0.83 | 0.56 | 0.54 | 0.50 | 0.97 | 0.98 | 0.61 |
| CellFM | 0.85 | 0.52 | 0.51 | 0.49 | 0.93 | 0.97 | 0.59 |
| LangCell | 0.85 | 0.53 | 0.51 | 0.49 | 0.97 | 0.97 | 0.59 |
| scANVI | 0.80 | 0.47 | 0.60 | 0.48 | 0.93 | 0.95 | 0.59 |
| SingleR | 0.79 | 0.54 | 0.57 | 0.46 | n/a | n/a | 0.59 |
| CellPLM | 0.75 | 0.51 | 0.59 | 0.49 | 0.96 | 0.94 | 0.59 |
| Geneformer | 0.81 | 0.47 | 0.52 | 0.45 | 0.95 | 0.93 | 0.56 |
| scPRINT | 0.76 | 0.49 | 0.50 | 0.46 | 0.95 | 0.95 | 0.55 |
| GenePT-w | 0.56 | 0.50 | 0.39 | 0.32 | 0.91 | 0.87 | 0.44 |
| CELLama | 0.66 | 0.31 | 0.38 | 0.32 | 0.88 | 0.89 | 0.42 |
| scBERT | 0.57 | 0.29 | 0.30 | 0.26 | 0.70 | 0.82 | 0.35 |

**Figure S226: Directional cross-dataset annotation for uterus, TSv2 to TSv1. a,** All 5,882 target cells, including 29 cells from source-absent labels. **b,** 5,853 target cells from 17 shared labels. Methods and metrics are as in Fig. S185.

**a** Vasculature · reference: v1, query: v2 · open-world

| method | Annotation metrics: *Accuracy* | $Acc_{top3}$ | $Prec_{Macro}$ | $Recall_{Macro}$ | $F1_{Macro}$ | Aggregated: *Annotation* |
|---|---|---|---|---|---|---|
| SCimilarity | 0.75 | 0.88 | 0.67 | 0.79 | 0.66 | 0.72 |
| scVI | 0.75 | 0.89 | 0.67 | 0.76 | 0.66 | 0.71 |
| CellTypist | 0.76 | 0.89 | 0.62 | 0.77 | 0.66 | 0.70 |
| scFoundation | 0.73 | 0.87 | 0.63 | 0.78 | 0.65 | 0.70 |
| PCA | 0.75 | 0.88 | 0.62 | 0.76 | 0.65 | 0.69 |
| UCE | 0.74 | 0.86 | 0.61 | 0.78 | 0.64 | 0.69 |
| scANVI | 0.75 | 0.88 | 0.61 | 0.76 | 0.64 | 0.69 |
| scGPT | 0.75 | 0.89 | 0.60 | 0.76 | 0.64 | 0.69 |
| Geneformer | 0.74 | 0.86 | 0.60 | 0.77 | 0.64 | 0.69 |
| scCello | 0.75 | 0.88 | 0.60 | 0.76 | 0.64 | 0.69 |
| CellPLM | 0.74 | 0.87 | 0.60 | 0.77 | 0.64 | 0.69 |
| SingleR | 0.74 | n/a | 0.59 | 0.76 | 0.62 | 0.68 |
| LangCell | 0.73 | 0.87 | 0.58 | 0.76 | 0.62 | 0.67 |
| scPRINT | 0.73 | 0.87 | 0.57 | 0.75 | 0.61 | 0.67 |
| GenePT-w | 0.70 | 0.88 | 0.54 | 0.72 | 0.56 | 0.63 |
| CellFM | 0.73 | 0.89 | 0.57 | 0.65 | 0.55 | 0.63 |
| CELLama | 0.69 | 0.85 | 0.48 | 0.62 | 0.50 | 0.57 |
| scBERT | 0.51 | 0.73 | 0.28 | 0.36 | 0.28 | 0.36 |

**b** Vasculature · reference: v1, query: v2 · shared

| method | Annotation metrics: *Accuracy* | $Prec_{Macro}$ | $Recall_{Macro}$ | $F1_{Macro}$ | $AUC_{ROC}$ | $Acc_{top3}$ | Aggregated: *Annotation* |
|---|---|---|---|---|---|---|---|
| SCimilarity | 0.83 | 0.76 | 0.79 | 0.74 | 0.99 | 0.98 | 0.78 |
| scVI | 0.83 | 0.76 | 0.76 | 0.73 | 0.99 | 0.99 | 0.77 |
| CellTypist | 0.84 | 0.72 | 0.77 | 0.73 | 0.99 | 0.99 | 0.77 |
| PCA | 0.84 | 0.71 | 0.76 | 0.73 | 0.98 | 0.97 | 0.76 |
| scFoundation | 0.81 | 0.71 | 0.78 | 0.72 | 0.97 | 0.97 | 0.75 |
| UCE | 0.82 | 0.70 | 0.78 | 0.72 | 0.98 | 0.96 | 0.75 |
| Geneformer | 0.83 | 0.70 | 0.77 | 0.71 | 0.98 | 0.95 | 0.75 |
| scGPT | 0.83 | 0.69 | 0.76 | 0.72 | 0.99 | 0.99 | 0.75 |
| scCello | 0.83 | 0.69 | 0.76 | 0.71 | 0.98 | 0.98 | 0.75 |
| SingleR | 0.83 | 0.70 | 0.76 | 0.70 | n/a | n/a | 0.75 |
| CellPLM | 0.82 | 0.69 | 0.77 | 0.71 | 0.99 | 0.97 | 0.75 |
| scANVI | 0.84 | 0.66 | 0.76 | 0.68 | 0.98 | 0.98 | 0.73 |
| LangCell | 0.81 | 0.67 | 0.76 | 0.69 | 0.97 | 0.97 | 0.73 |
| scPRINT | 0.81 | 0.66 | 0.75 | 0.69 | 0.98 | 0.97 | 0.73 |
| CellFM | 0.82 | 0.66 | 0.65 | 0.63 | 0.98 | 0.99 | 0.69 |
| GenePT-w | 0.78 | 0.63 | 0.72 | 0.62 | 0.98 | 0.98 | 0.69 |
| CELLama | 0.76 | 0.52 | 0.62 | 0.54 | 0.97 | 0.95 | 0.61 |
| scBERT | 0.57 | 0.31 | 0.36 | 0.30 | 0.86 | 0.82 | 0.38 |

**Figure S227: Directional cross-dataset annotation for vasculature, TSv1 to TSv2. a,** All 23,569 target cells, including 2,374 cells from source-absent labels. **b,** 21,195 target cells from 19 shared labels. Methods and metrics are as in Fig. S185.

**a** Vasculature · reference: v2, query: v1 · open-world

| method | Accuracy | $Acc_{top3}$ | $Prec_{Macro}$ | $Recall_{Macro}$ | $F1_{Macro}$ | Aggregated Annotation |
|---|---|---|---|---|---|---|
| | Annotation metrics | | | | | |
| SingleR | 0.82 | n/a | 0.71 | 0.69 | 0.64 | 0.72 |
| PCA | 0.80 | 0.97 | 0.68 | 0.69 | 0.64 | 0.70 |
| scCello | 0.80 | 0.98 | 0.68 | 0.66 | 0.64 | 0.69 |
| CellTypist | 0.81 | 0.98 | 0.62 | 0.68 | 0.62 | 0.69 |
| scVI | 0.79 | 0.98 | 0.65 | 0.66 | 0.62 | 0.68 |
| SCimilarity | 0.77 | 0.98 | 0.60 | 0.66 | 0.59 | 0.65 |
| CellPLM | 0.76 | 0.98 | 0.62 | 0.65 | 0.59 | 0.65 |
| scPRINT | 0.72 | 0.97 | 0.64 | 0.66 | 0.58 | 0.65 |
| scANVI | 0.77 | 0.92 | 0.58 | 0.67 | 0.57 | 0.65 |
| CellFM | 0.79 | 0.97 | 0.61 | 0.61 | 0.57 | 0.65 |
| UCE | 0.75 | 0.98 | 0.64 | 0.62 | 0.57 | 0.65 |
| LangCell | 0.73 | 0.95 | 0.62 | 0.60 | 0.54 | 0.62 |
| Geneformer | 0.74 | 0.94 | 0.60 | 0.61 | 0.54 | 0.62 |
| scGPT | 0.74 | 0.97 | 0.58 | 0.61 | 0.55 | 0.62 |
| scFoundation | 0.70 | 0.93 | 0.60 | 0.61 | 0.55 | 0.62 |
| GenePT-w | 0.76 | 0.94 | 0.54 | 0.58 | 0.49 | 0.59 |
| CELLama | 0.70 | 0.91 | 0.45 | 0.53 | 0.41 | 0.52 |
| scBERT | 0.51 | 0.82 | 0.29 | 0.39 | 0.26 | 0.36 |

**b** Vasculature · reference: v2, query: v1 · shared

| method | Accuracy | $Prec_{Macro}$ | $Recall_{Macro}$ | $F1_{Macro}$ | $AUC_{ROC}$ | $Acc_{top3}$ | Aggregated Annotation |
|---|---|---|---|---|---|---|---|
| | Annotation metrics | | | | | | |
| SingleR | 0.82 | 0.71 | 0.69 | 0.64 | n/a | n/a | 0.72 |
| PCA | 0.80 | 0.68 | 0.69 | 0.64 | 0.98 | 0.97 | 0.70 |
| scCello | 0.80 | 0.68 | 0.66 | 0.64 | 0.99 | 0.98 | 0.69 |
| CellTypist | 0.81 | 0.62 | 0.68 | 0.63 | 0.99 | 0.98 | 0.69 |
| scVI | 0.80 | 0.65 | 0.66 | 0.62 | 0.99 | 0.98 | 0.68 |
| SCimilarity | 0.77 | 0.60 | 0.66 | 0.59 | 0.99 | 0.98 | 0.66 |
| CellPLM | 0.76 | 0.62 | 0.65 | 0.59 | 0.98 | 0.98 | 0.65 |
| scPRINT | 0.72 | 0.64 | 0.66 | 0.58 | 0.98 | 0.97 | 0.65 |
| scANVI | 0.77 | 0.58 | 0.67 | 0.57 | 0.96 | 0.92 | 0.65 |
| CellFM | 0.79 | 0.61 | 0.61 | 0.57 | 0.98 | 0.97 | 0.65 |
| UCE | 0.75 | 0.64 | 0.62 | 0.57 | 0.98 | 0.98 | 0.65 |
| LangCell | 0.73 | 0.62 | 0.60 | 0.54 | 0.98 | 0.95 | 0.62 |
| Geneformer | 0.74 | 0.60 | 0.61 | 0.54 | 0.97 | 0.94 | 0.62 |
| scGPT | 0.74 | 0.58 | 0.61 | 0.55 | 0.99 | 0.97 | 0.62 |
| scFoundation | 0.70 | 0.60 | 0.61 | 0.55 | 0.98 | 0.93 | 0.62 |
| GenePT-w | 0.76 | 0.54 | 0.58 | 0.49 | 0.98 | 0.94 | 0.59 |
| CELLama | 0.70 | 0.45 | 0.53 | 0.41 | 0.95 | 0.91 | 0.52 |
| scBERT | 0.51 | 0.29 | 0.39 | 0.26 | 0.86 | 0.82 | 0.36 |

**Figure S228: Directional cross-dataset annotation for vasculature, TSv2 to TSv1. a,** All 15,558 target cells, including 3 cells from source-absent labels. **b,** 15,555 target cells from 19 shared labels. Methods and metrics are as in Fig. S185.

#### 2.8 Gene-level capability and task-dependent biological signal

The capability inventory comprised 11 static gene representations, nine contextual gene representations and two completed model-native regulatory outputs (Fig. S229). These output classes were evaluated separately because a fixed gene representation, a gene state conditioned on expression data and a model-specific regulatory score address different biological questions.

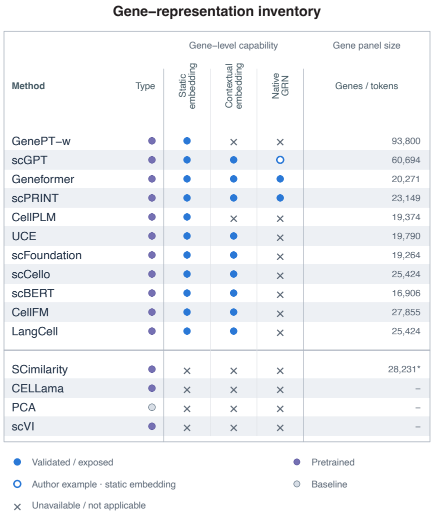

**Figure S229: Gene-level capability inventory.** The matrix distinguishes static gene representations, contextual gene representations and native GRN outputs across the 15 considered methods. Eleven methods contribute static representations, nine contribute contextual representations, and Geneformer and scPRINT contribute the two completed native regulatory outputs. Bars show native gene-vocabulary sizes before intersection with the common 16,508-gene universe.

##### *2.8.1 Functional coherence of static and contextual representations*

Gene-function performance varied substantially across representations (Fig. S230). GenePT-w gave the highest displayed value for each endpoint: STRING physical-interaction AUROC 0.827 and AUPRC 0.641, and MAP 0.156, 0.196 and 0.247 for GO Biological Process, Reactome and KEGG. The corresponding co-expression values were 0.585, 0.286, 0.070, 0.072 and 0.075. Contextual computation did not have a uniform direction: scFoundation increased from 0.508 to 0.639 in PPI AUROC, whereas Geneformer decreased from 0.675 to 0.629. These results show model-specific functional signal rather than a general advantage of transcriptomic pretraining or contextualization.

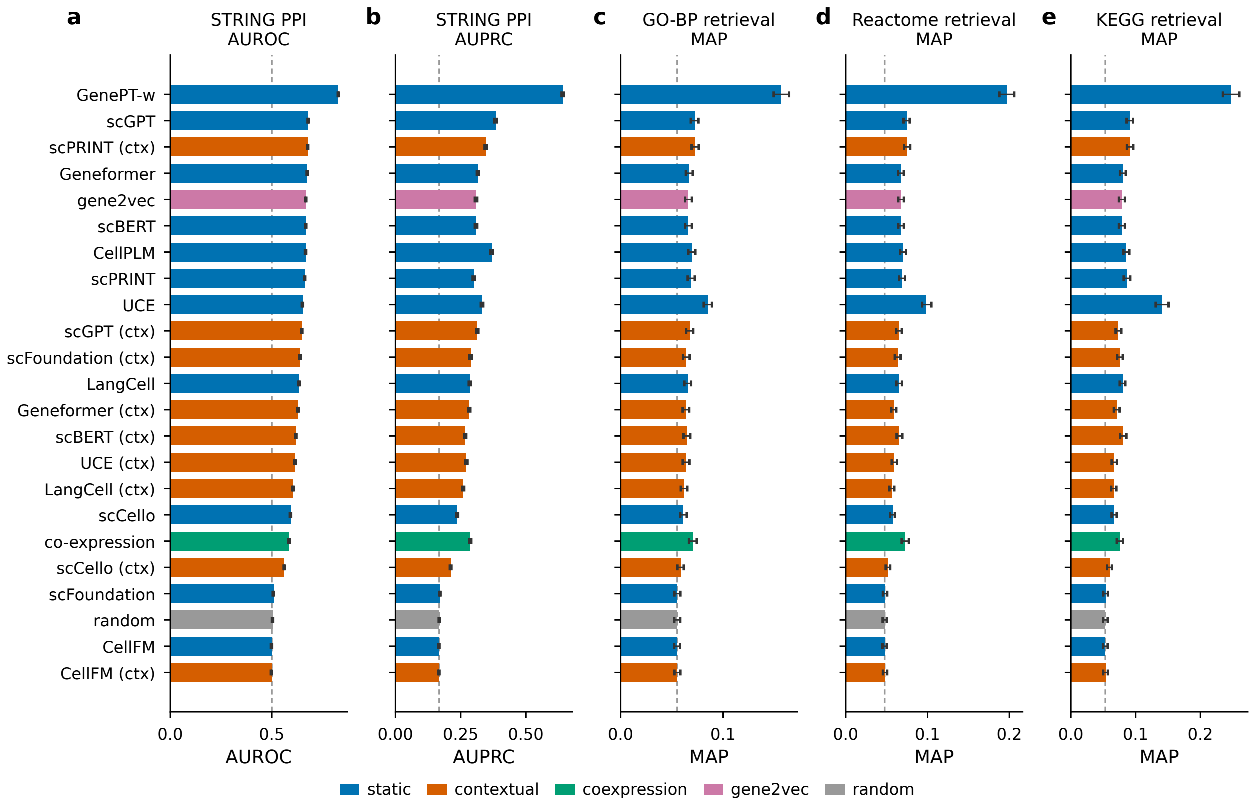

**Figure S230: Functional coherence of static and contextual gene representations.** Eleven static and nine contextual representations were evaluated with co-expression, gene2vec and random references. Contextual states were obtained from the deterministic 112-cell reference spanning seven compartments, 26 tissues and 55 cell types. The evaluation universe was fixed as the 16,508-gene identity intersection before task filters; undefined contextual similarities received neutral score zero and did not remove genes. **a,b,** STRING v12 physical-interaction link prediction with 40,000 positive and 200,000 degree-matched negative pairs, evaluated by AUROC and AUPRC. **c–e,** GO Biological Process, Reactome and KEGG retrieval, evaluated by MAP. Error bars are 95% percentile-bootstrap intervals from 1,000 pair or query-gene resamples. Dashed lines denote chance AUROC or the empirical random-provider value.

GenePT-w also retained the highest precision over the leading retrieved genes, and UCE was the next strongest representation across GO Biological Process, Reactome and KEGG (Fig. S231). This agreement between MAP and precision at multiple retrieval depths shows that the functional signal was not driven only by the tail of the ranked lists.

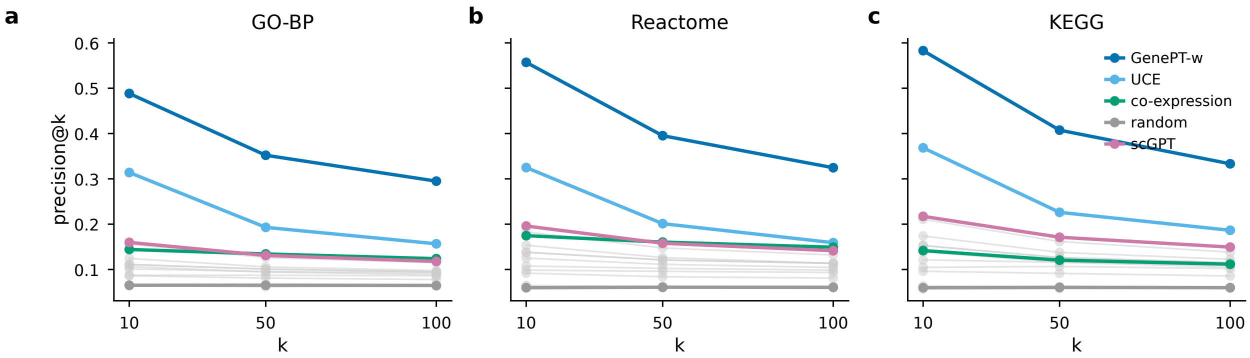

**Figure S231: Precision of pathway and ontology retrieval.** Precision at 10, 50 and 100 retrieved genes for GO Biological Process (**a**), Reactome (**b**) and KEGG (**c**). GenePT-w, UCE, scGPT, co-expression and random references are highlighted; the remaining representations are shown in grey.

##### *2.8.2 Regulatory-edge prioritization in BEELINE*

Broad functional organization did not translate directly into context-specific regulatory-edge recovery (Fig. S232). GENIE3 was strongest among the established GRN methods, with a study-equal aggregate AUPRC ratio of 1.27. GenePT-w led the static scFM representations (1.12), and UCE led the contextual representations (1.15). Contextualization increased the combined regulatory score for six of the nine eligible models, but the magnitude and direction varied across models. Neither completed native output showed consistently strong recovery across the evaluated datasets.

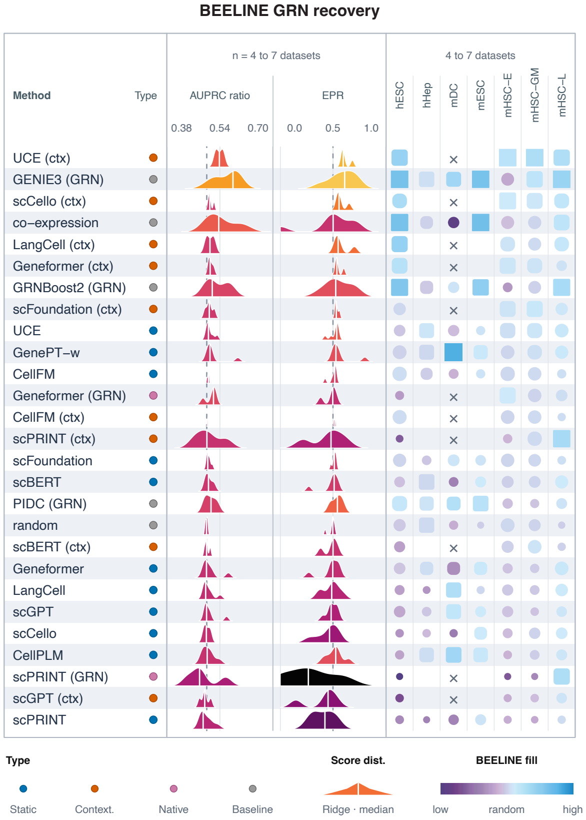

**Figure S232: BEELINE regulatory-edge recovery.** Static representations and the established GRN methods GENIE3, GRNBoost2 and PIDC were evaluated on seven datasets from five studies. Contextual representations and the completed Geneformer and scPRINT native outputs were evaluated on four count-traceable datasets from two studies. Panels show method class, distributions of AUPRC ratio and early-precision ratio, and aggregate performance and ranking. The three mHSC datasets were first averaged within GSE81682 before studies received equal weight. Ratios are relative to random expectation 1.

##### *2.8.3 Perturbation-response and curated-regulon retrieval*

Perturbation-response retrieval produced a different ordering from broad gene function (Fig. S233). Co-expression achieved a study-equal AUROC of 0.599 and AUPRC lift of 1.678. Contextual Geneformer, LangCell and scPRINT achieved AUPRC lifts of 2.454, 2.429 and 2.118, respectively, whereas GenePT-w was close to random expectation (AUROC 0.512; AUPRC lift 1.134). Contextual representations therefore improved retrieval for selected models, but the effect was not shared across the panel.

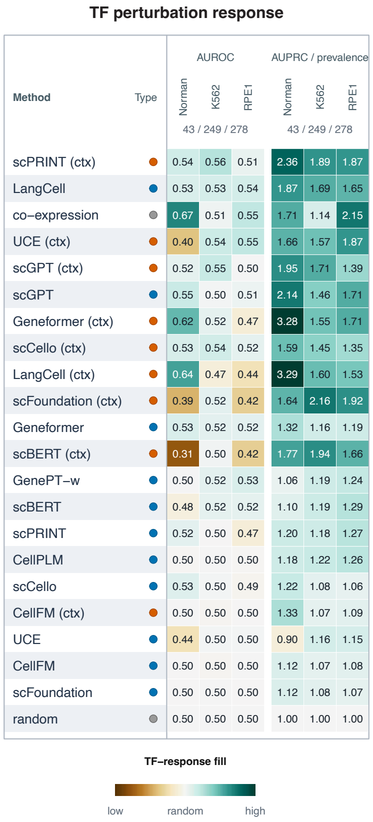

**Figure S233: Retrieval of transcription-factor perturbation-response genes.** For 570 eligible single-TF perturbations (Norman CRISPRa, 43; Replogle K562 CRISPRi, 249; Replogle RPE1 CRISPRi, 278), positives were the 100 genes with the largest absolute normalized mean-expression response relative to control, excluding the perturbed TF. Contextual states were computed from 112 control cells per dataset. Heatmap cells show per-dataset mean AUROC (left) and AUPRC divided by positive prevalence (right).

Curated-regulon recovery likewise differed by representation and resource (Fig. S234). GenePT-w had the highest displayed median for TRRUST (0.689), and contextual scPRINT led DoRothEA ABC (0.685). ChEA recovery was generally weaker, although co-expression (0.560) and contextual scPRINT (0.551) were modestly above chance. Contextualization did not uniformly improve static vectors.

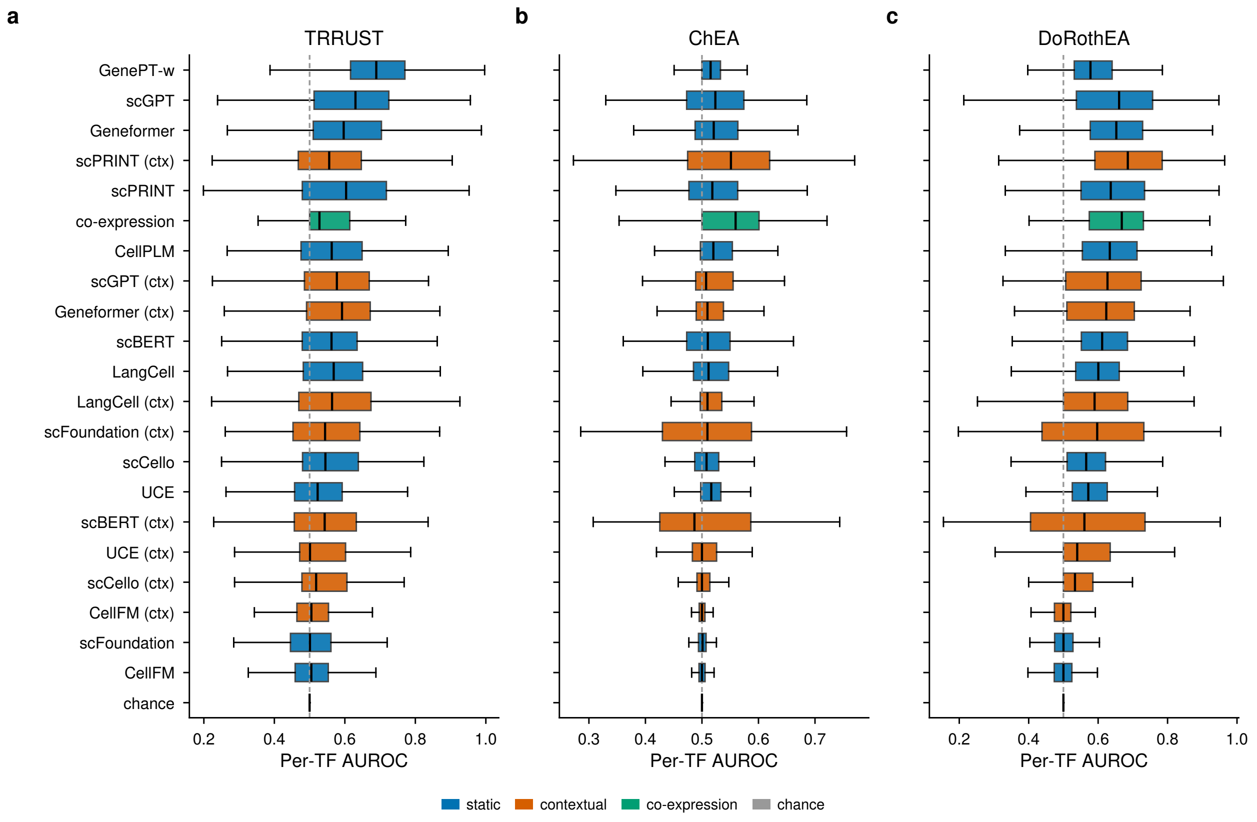

**Figure S234: Recovery of curated transcription-factor regulons.** All other genes in the fixed 16,508-gene universe were ranked for each eligible TF. Boxplots show per-TF AUROC for TRRUST (319 TFs), ChEA (272 TFs) and DoRothEA ABC (295 TFs), with at least five recorded targets per TF. Eleven static and nine contextual representations are shown with co-expression and chance. Contextual states use the 112-cell pan-tissue reference.

The changes in ordering across physical interaction, pathways, regulatory edges, curated regulons and perturbation-associated responses separate broad functional relatedness from context-specific regulation. A representation that led one gene-level endpoint did not generally lead the others.

#### 2.9 Cell-level double-perturbation response prediction

The additive reference remained the strongest predictor after standardizing the decoder comparison across 13 scFMs. It had the lowest normalized L2 error over the 65 held-out double perturbations in the Norman dataset (mean 0.567), followed by GenePT-w, the strongest scFM-based decoder (0.692; Fig. S235). For the CEBPA+CEBPE example, the additive prediction closely followed the observed response ($R^2 = 0.93$) and exceeded every scFM-based decoder. Several scFM representations approached the additive reference, but none surpassed it. Training-only standardization, a common 30-dimensional projection and a tuned ridge decoder therefore strengthened the conclusion that the additive model remains the most accurate approach in this setting.

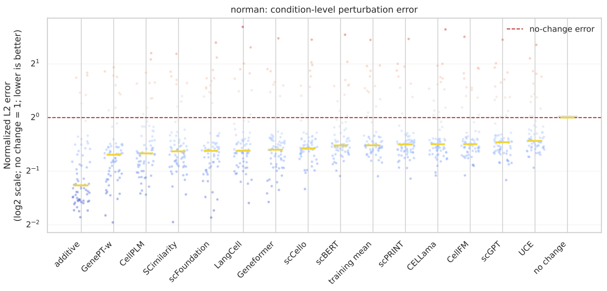

(a)

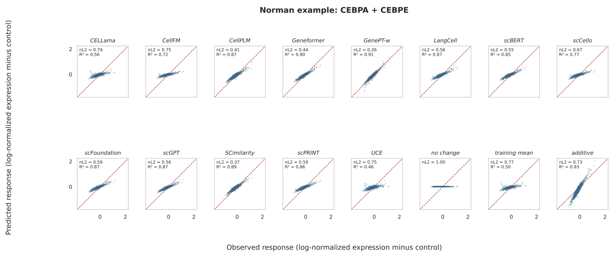

(b)

**Figure S235: Cell-level double-perturbation response prediction in the Norman dataset. a,** Normalized L2 error across 65 held-out double perturbations after training-only standardization, a common 30-dimensional projection and tuned ridge decoding; lower values indicate better prediction. **b,** Predicted and observed gene-wise expression changes for CEBPA+CEBPE. The additive reference had $R^2 = 0.93$.

#### 2.10 Evaluation on spatial transcriptomics

Across 17 methods and 25 sections, the coordinate-aware spatial model NOVAE had the lowest overall mean rank (2.873) and appeared among the five best methods in 22 sections. It was followed by scFoundation (5.979), CellPLM (6.007), scVI trained de novo (6.318) and scGPT (6.391); pretrained scVI and PCA had mean ranks of 7.558 and 7.613, respectively. NOVAE had a lower paired section-level aggregate rank than each of the other 16 methods (all Holm-adjusted $P = 1.96 \times 10^{-4}$).

**Table S10: Spatial-domain aggregate across 25 sections.** Mean ranks and 95% section-bootstrap intervals were calculated from the equal-weight average of the four metric-family ranks. Lower ranks indicate better performance; top-five counts are the number of sections in which a method ranked among the five best methods.

| Method | Role | Mean rank | 95% interval | Top five |
|---|---|---|---|---|
| NOVAE | Spatial-native | 2.873 | 2.290–3.547 | 22/25 |
| scFoundation | Frozen scFM | 5.979 | 5.243–6.789 | 16/25 |
| CellPLM | Frozen scFM | 6.007 | 5.201–6.893 | 20/25 |
| scVI (de novo) | Target-trained | 6.318 | 5.695–7.006 | 16/25 |
| scGPT | Frozen scFM | 6.391 | 5.672–7.130 | 15/25 |
| scVI (pretrained) | Pretrained reference | 7.558 | 6.843–8.350 | 7/25 |
| HVGs+PCA | Target-fitted reference | 7.613 | 6.474–8.780 | 11/25 |
| SCimilarity | Frozen scFM | 8.315 | 7.594–9.072 | 3/25 |
| Geneformer | Frozen scFM | 8.503 | 7.746–9.236 | 2/25 |
| LangCell | Frozen scFM | 8.681 | 7.684–9.723 | 7/25 |
| UCE | Frozen scFM | 9.494 | 8.815–10.182 | 2/25 |
| scPRINT | Frozen scFM | 9.777 | 8.890–10.708 | 2/25 |
| CELLama | Frozen scFM | 11.536 | 10.531–12.509 | 1/25 |
| CellFM | Frozen scFM | 12.070 | 10.817–13.312 | 1/25 |
| scCello | Frozen scFM | 12.884 | 12.290–13.452 | 0/25 |
| GenePT-w | Frozen scFM | 13.806 | 13.242–14.346 | 0/25 |
| scBERT | Frozen scFM | 15.199 | 14.736–15.656 | 0/25 |

CellPLM was among the strongest frozen scFMs and showed its clearest advantage on SpatialLIBD, where it had the highest mean ARI among the frozen scFMs (0.243), exceeding PCA (0.177) and approaching scVI trained de novo (0.258), although NOVAE remained higher (0.309). Absolute spatial-domain ARIs were low for all methods on HER2ST and MERFISH.

The ordering changed in the separate MERFISH cell-type analysis. CELLama had the highest mean ARI (0.358), followed by scVI trained de novo (0.350), PCA (0.317) and CellPLM (0.244). CellPLM was therefore the second-ranked frozen scFM and fourth overall for this endpoint. NOVAE, which led spatial-domain recovery, had a mean cell-type ARI of 0.041 and ranked 16th of 17. Spatial domains and cell identity therefore produced distinct method orderings.

The complete cross-section composite is reported in the corresponding main-text figure. The 30 panels below provide section-level maps and metric tables; their 15- or 13-method displays are selected visual subsets of the complete 17-method analysis.

###### *2.10.1 SpatialLIBD spatial-domain clustering*

Section-level maps and metric tables for all 12 SpatialLIBD sections are shown in Figs. S236–S247.

**a**

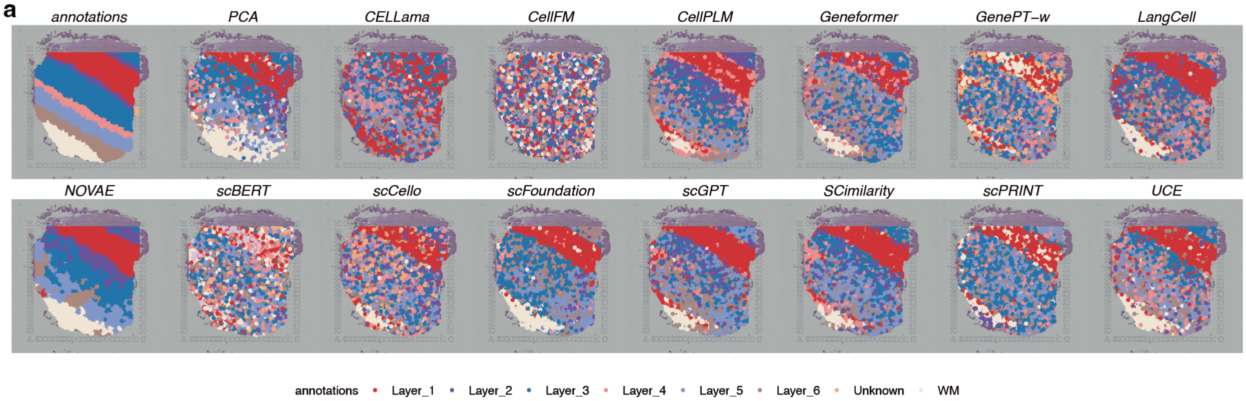

**b**

| | Cluster concordance | | | | | Local Structure | | Spatial coherence | | Spatial marker | | Aggregated scores | | | |
|---|---|---|---|---|---|---|---|---|---|---|---|---|---|---|---|
| method | ARI | NMI | HOM | COM | FMI | Acc@kNN | GC | 1-CHAOS | 1-PAS | Moran's I | 1-Geary's C | Cluster | Local | Coherence | Marker |
| NOVAE | 0.43 | 0.55 | 0.53 | 0.57 | 0.54 | 0.86 | 0.99 | 0.94 | 0.94 | 0.25 | 0.25 | 0.52 | 0.92 | 0.94 | 0.25 |
| scFoundation | 0.26 | 0.32 | 0.31 | 0.33 | 0.40 | 0.62 | 0.72 | 0.93 | 0.54 | 0.24 | 0.24 | 0.33 | 0.67 | 0.74 | 0.24 |
| CellPLM | 0.26 | 0.35 | 0.35 | 0.35 | 0.39 | 0.62 | 0.79 | 0.93 | 0.46 | 0.22 | 0.22 | 0.34 | 0.70 | 0.69 | 0.22 |
| scGPT | 0.24 | 0.29 | 0.29 | 0.29 | 0.37 | 0.57 | 0.72 | 0.93 | 0.39 | 0.28 | 0.28 | 0.30 | 0.65 | 0.66 | 0.28 |
| SCimilarity | 0.26 | 0.30 | 0.30 | 0.30 | 0.39 | 0.53 | 0.67 | 0.93 | 0.41 | 0.25 | 0.24 | 0.31 | 0.60 | 0.67 | 0.25 |
| Geneformer | 0.14 | 0.22 | 0.23 | 0.22 | 0.29 | 0.63 | 0.61 | 0.93 | 0.30 | 0.26 | 0.26 | 0.22 | 0.62 | 0.62 | 0.26 |
| scPRINT | 0.17 | 0.24 | 0.23 | 0.25 | 0.32 | 0.54 | 0.64 | 0.93 | 0.35 | 0.24 | 0.24 | 0.24 | 0.59 | 0.64 | 0.24 |
| LangCell | 0.22 | 0.25 | 0.25 | 0.25 | 0.35 | 0.52 | 0.55 | 0.93 | 0.33 | 0.26 | 0.25 | 0.27 | 0.53 | 0.63 | 0.25 |
| PCA | 0.24 | 0.27 | 0.27 | 0.28 | 0.38 | 0.49 | 0.32 | 0.93 | 0.54 | 0.26 | 0.25 | 0.29 | 0.40 | 0.73 | 0.25 |
| UCE | 0.23 | 0.28 | 0.28 | 0.27 | 0.36 | 0.54 | 0.31 | 0.93 | 0.37 | 0.22 | 0.22 | 0.28 | 0.42 | 0.65 | 0.22 |
| scCello | 0.13 | 0.17 | 0.18 | 0.17 | 0.27 | 0.47 | 0.48 | 0.92 | 0.18 | 0.21 | 0.21 | 0.18 | 0.47 | 0.55 | 0.21 |
| CELLama | 0.05 | 0.08 | 0.08 | 0.08 | 0.22 | 0.46 | 0.59 | 0.93 | 0.16 | 0.21 | 0.21 | 0.10 | 0.53 | 0.54 | 0.21 |
| GenePT-w | 0.08 | 0.14 | 0.15 | 0.14 | 0.23 | 0.43 | 0.29 | 0.93 | 0.17 | 0.22 | 0.22 | 0.15 | 0.36 | 0.55 | 0.22 |
| scBERT | 0.06 | 0.12 | 0.15 | 0.10 | 0.18 | 0.37 | 0.11 | 0.91 | 0.05 | 0.17 | 0.17 | 0.12 | 0.24 | 0.48 | 0.17 |
| CellFM | 0.00 | 0.01 | 0.01 | 0.01 | 0.14 | 0.27 | 0.06 | 0.91 | 0.00 | 0.03 | 0.03 | 0.03 | 0.16 | 0.46 | 0.03 |

**Figure S236: Spatial-domain clustering in SpatialLIBD section 151507. a,** Predicted domains from 15 displayed methods and the reference annotation on the Visium section. **b,** Scores for cluster concordance (ARI, NMI, homogeneity, completeness and FMI), local neighbourhood structure (Acc@kNN and graph connectivity), spatial coherence (1−CHAOS and 1−PAS), and spatial-marker autocorrelation (Moran's I and 1−Geary's C). The four family means receive equal weight; rows are ordered by their aggregate score.

**a**

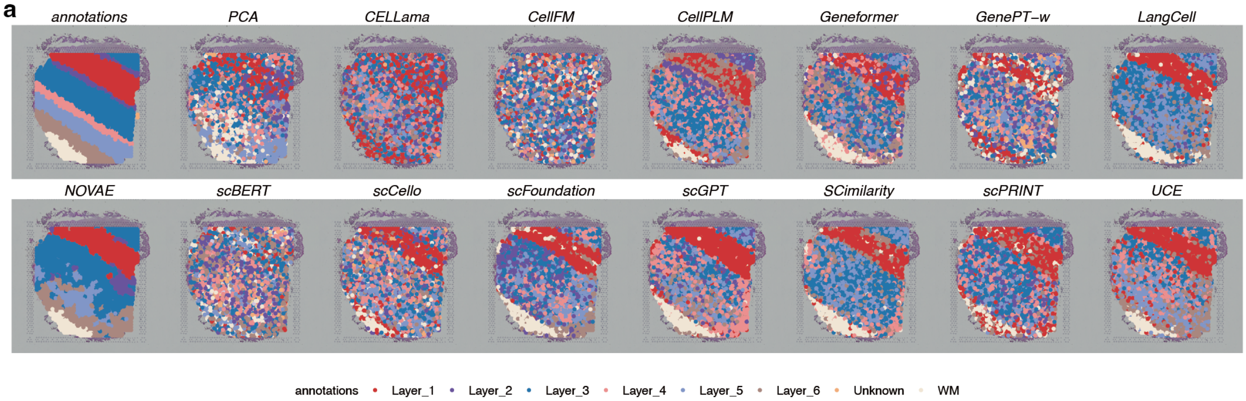

**b**

| | Cluster concordance | | | | | Local Structure | | Spatial coherence | | Spatial marker | | Aggregated scores | | | |
|---|---|---|---|---|---|---|---|---|---|---|---|---|---|---|---|
| method | ARI | NMI | HOM | COM | FMI | Acc@kNN | GC | 1-CHAOS | 1-PAS | Moran's I | 1-Geary's C | Cluster | Local | Coherence | Marker |
| NOVAE | 0.42 | 0.49 | 0.46 | 0.52 | 0.55 | 0.83 | 0.97 | 0.94 | 0.94 | 0.23 | 0.23 | 0.49 | 0.90 | 0.94 | 0.23 |
| scGPT | 0.26 | 0.31 | 0.32 | 0.30 | 0.39 | 0.60 | 0.74 | 0.93 | 0.41 | 0.23 | 0.23 | 0.32 | 0.67 | 0.67 | 0.23 |
| scFoundation | 0.21 | 0.28 | 0.29 | 0.27 | 0.35 | 0.62 | 0.72 | 0.94 | 0.45 | 0.21 | 0.21 | 0.28 | 0.67 | 0.69 | 0.21 |
| CellPLM | 0.16 | 0.27 | 0.27 | 0.26 | 0.32 | 0.60 | 0.74 | 0.93 | 0.43 | 0.21 | 0.21 | 0.26 | 0.67 | 0.68 | 0.21 |
| SCimilarity | 0.20 | 0.29 | 0.28 | 0.29 | 0.36 | 0.54 | 0.65 | 0.93 | 0.47 | 0.23 | 0.23 | 0.28 | 0.59 | 0.70 | 0.23 |
| LangCell | 0.21 | 0.26 | 0.25 | 0.27 | 0.37 | 0.53 | 0.46 | 0.94 | 0.47 | 0.20 | 0.20 | 0.27 | 0.50 | 0.70 | 0.20 |
| scPRINT | 0.16 | 0.20 | 0.20 | 0.19 | 0.32 | 0.54 | 0.60 | 0.93 | 0.30 | 0.23 | 0.23 | 0.21 | 0.57 | 0.62 | 0.23 |
| Geneformer | 0.12 | 0.21 | 0.21 | 0.20 | 0.28 | 0.61 | 0.58 | 0.93 | 0.28 | 0.20 | 0.20 | 0.20 | 0.60 | 0.61 | 0.20 |
| UCE | 0.22 | 0.27 | 0.26 | 0.27 | 0.37 | 0.53 | 0.22 | 0.93 | 0.46 | 0.20 | 0.20 | 0.28 | 0.37 | 0.70 | 0.20 |
| scCello | 0.13 | 0.15 | 0.16 | 0.14 | 0.28 | 0.49 | 0.46 | 0.92 | 0.19 | 0.18 | 0.18 | 0.17 | 0.47 | 0.56 | 0.18 |
| PCA | 0.13 | 0.17 | 0.18 | 0.17 | 0.28 | 0.46 | 0.18 | 0.93 | 0.42 | 0.19 | 0.19 | 0.19 | 0.32 | 0.67 | 0.19 |
| GenePT-w | 0.09 | 0.16 | 0.18 | 0.15 | 0.24 | 0.46 | 0.27 | 0.93 | 0.15 | 0.19 | 0.19 | 0.16 | 0.37 | 0.54 | 0.19 |
| CELLama | 0.04 | 0.07 | 0.07 | 0.07 | 0.22 | 0.46 | 0.48 | 0.93 | 0.12 | 0.17 | 0.16 | 0.09 | 0.47 | 0.52 | 0.16 |
| scBERT | 0.06 | 0.14 | 0.18 | 0.12 | 0.19 | 0.39 | 0.11 | 0.91 | 0.04 | 0.15 | 0.15 | 0.14 | 0.25 | 0.48 | 0.15 |
| CellFM | 0.00 | 0.01 | 0.01 | 0.01 | 0.16 | 0.25 | 0.03 | 0.91 | 0.01 | 0.02 | 0.02 | 0.04 | 0.14 | 0.46 | 0.02 |

**Figure S237: Spatial-domain clustering in SpatialLIBD section 151508. a,** Predicted domains and reference annotation. **b,** Scores for 15 displayed methods. Display and scoring conventions are as in Fig. S236.

**a**

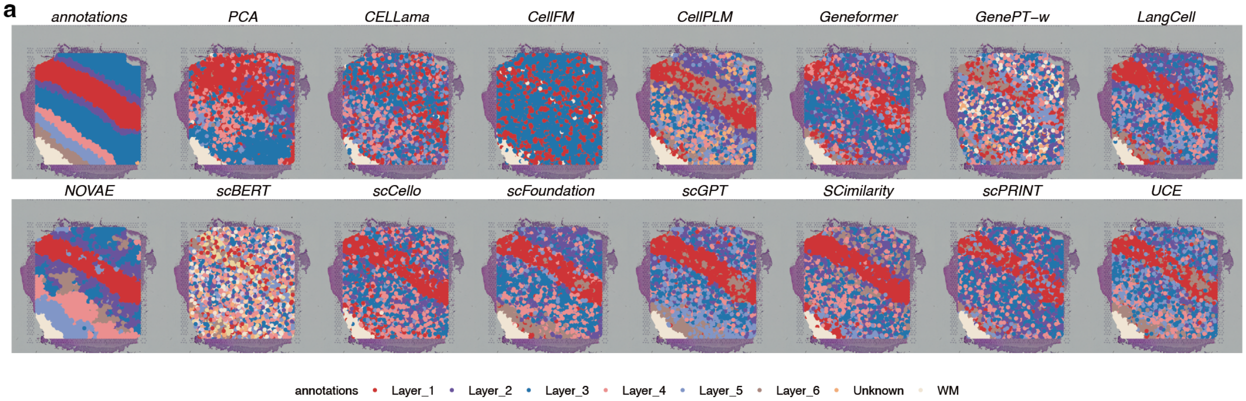

**b**

| | Cluster concordance | | | | | Local Structure | | Spatial coherence | | Spatial marker | | Aggregated scores | | | |
|---|---|---|---|---|---|---|---|---|---|---|---|---|---|---|---|
| method | ARI | NMI | HOM | COM | FMI | Acc@kNN | GC | 1-CHAOS | 1-PAS | Moran's I | 1-Geary's C | Cluster | Local | Coherence | Marker |
| NOVAE | 0.36 | 0.51 | 0.54 | 0.48 | 0.49 | 0.86 | 0.98 | 0.95 | 0.91 | 0.26 | 0.26 | 0.48 | 0.92 | 0.93 | 0.26 |
| scFoundation | 0.30 | 0.37 | 0.39 | 0.35 | 0.45 | 0.70 | 0.83 | 0.94 | 0.56 | 0.28 | 0.28 | 0.37 | 0.76 | 0.75 | 0.28 |
| scGPT | 0.28 | 0.37 | 0.39 | 0.35 | 0.43 | 0.68 | 0.74 | 0.94 | 0.48 | 0.26 | 0.26 | 0.36 | 0.71 | 0.71 | 0.26 |
| CellPLM | 0.22 | 0.35 | 0.39 | 0.31 | 0.37 | 0.70 | 0.74 | 0.93 | 0.40 | 0.25 | 0.25 | 0.33 | 0.72 | 0.67 | 0.25 |
| SCimilarity | 0.29 | 0.35 | 0.37 | 0.33 | 0.44 | 0.63 | 0.67 | 0.94 | 0.40 | 0.27 | 0.27 | 0.36 | 0.65 | 0.67 | 0.27 |
| Geneformer | 0.22 | 0.27 | 0.28 | 0.27 | 0.40 | 0.70 | 0.69 | 0.93 | 0.41 | 0.26 | 0.26 | 0.29 | 0.69 | 0.67 | 0.26 |
| scPRINT | 0.25 | 0.32 | 0.33 | 0.31 | 0.42 | 0.65 | 0.61 | 0.94 | 0.42 | 0.26 | 0.26 | 0.32 | 0.63 | 0.68 | 0.26 |
| LangCell | 0.23 | 0.30 | 0.32 | 0.28 | 0.39 | 0.63 | 0.52 | 0.94 | 0.39 | 0.27 | 0.27 | 0.30 | 0.58 | 0.66 | 0.27 |
| scCello | 0.20 | 0.23 | 0.23 | 0.23 | 0.39 | 0.60 | 0.50 | 0.93 | 0.37 | 0.23 | 0.23 | 0.25 | 0.55 | 0.65 | 0.23 |
| PCA | 0.10 | 0.19 | 0.19 | 0.19 | 0.32 | 0.55 | 0.30 | 0.94 | 0.61 | 0.26 | 0.26 | 0.20 | 0.43 | 0.77 | 0.26 |
| UCE | 0.21 | 0.29 | 0.31 | 0.28 | 0.37 | 0.60 | 0.29 | 0.93 | 0.35 | 0.27 | 0.27 | 0.29 | 0.45 | 0.64 | 0.27 |
| CELLama | 0.05 | 0.12 | 0.12 | 0.12 | 0.27 | 0.58 | 0.53 | 0.93 | 0.23 | 0.27 | 0.27 | 0.14 | 0.55 | 0.58 | 0.27 |
| CellFM | 0.05 | 0.09 | 0.06 | 0.14 | 0.41 | 0.38 | 0.06 | 0.94 | 0.71 | 0.28 | 0.28 | 0.15 | 0.22 | 0.83 | 0.28 |
| GenePT-w | 0.12 | 0.20 | 0.22 | 0.18 | 0.29 | 0.58 | 0.32 | 0.93 | 0.19 | 0.24 | 0.24 | 0.20 | 0.45 | 0.56 | 0.24 |
| scBERT | 0.08 | 0.17 | 0.23 | 0.14 | 0.22 | 0.53 | 0.14 | 0.91 | 0.04 | 0.16 | 0.16 | 0.17 | 0.34 | 0.48 | 0.16 |

**Figure S238: Spatial-domain clustering in SpatialLIBD section 151509. a,** Predicted domains and reference annotation. **b,** Scores for 15 displayed methods. Display and scoring conventions are as in Fig. S236.

**a**

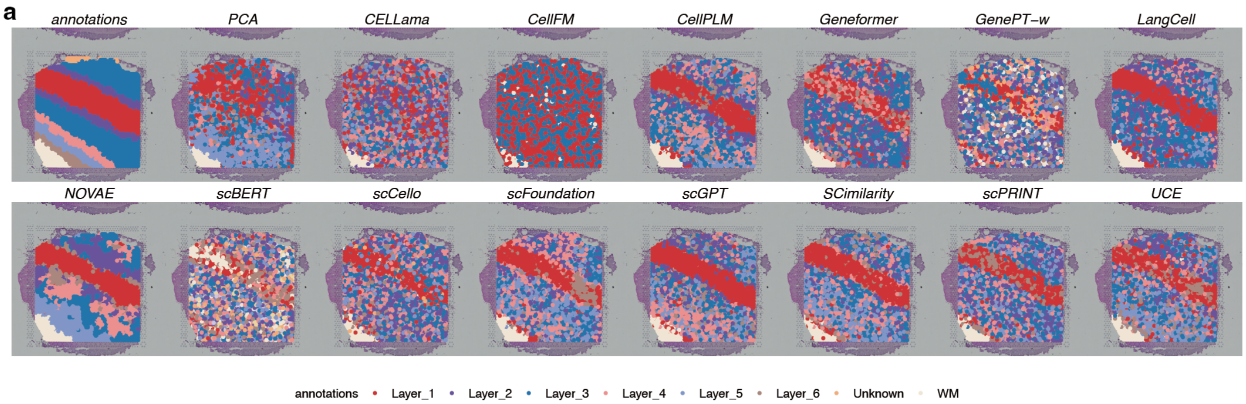

**b**

| | Cluster concordance | | | | | Local Structure | | Spatial coherence | | Spatial marker | | Aggregated scores | | | |
|---|---|---|---|---|---|---|---|---|---|---|---|---|---|---|---|
| method | ARI | NMI | HOM | COM | FMI | Acc@kNN | GC | 1-CHAOS | 1-PAS | Moran's I | 1-Geary's C | Cluster | Local | Coherence | Marker |
| NOVAE | 0.40 | 0.52 | 0.56 | 0.49 | 0.53 | 0.83 | 0.98 | 0.94 | 0.90 | 0.22 | 0.22 | 0.50 | 0.90 | 0.92 | 0.22 |
| CellPLM | 0.26 | 0.34 | 0.36 | 0.32 | 0.42 | 0.69 | 0.67 | 0.93 | 0.43 | 0.22 | 0.22 | 0.34 | 0.68 | 0.68 | 0.22 |
| scFoundation | 0.20 | 0.30 | 0.32 | 0.28 | 0.37 | 0.66 | 0.69 | 0.94 | 0.41 | 0.26 | 0.26 | 0.30 | 0.68 | 0.67 | 0.26 |
| scGPT | 0.28 | 0.32 | 0.35 | 0.30 | 0.43 | 0.64 | 0.62 | 0.93 | 0.39 | 0.26 | 0.26 | 0.34 | 0.63 | 0.66 | 0.26 |
| LangCell | 0.25 | 0.29 | 0.28 | 0.29 | 0.42 | 0.59 | 0.45 | 0.94 | 0.47 | 0.23 | 0.24 | 0.31 | 0.52 | 0.70 | 0.24 |
| SCimilarity | 0.26 | 0.30 | 0.32 | 0.28 | 0.42 | 0.60 | 0.53 | 0.93 | 0.36 | 0.24 | 0.24 | 0.31 | 0.57 | 0.65 | 0.24 |
| Geneformer | 0.15 | 0.20 | 0.21 | 0.20 | 0.34 | 0.67 | 0.59 | 0.93 | 0.33 | 0.24 | 0.24 | 0.22 | 0.63 | 0.63 | 0.24 |
| scPRINT | 0.20 | 0.27 | 0.29 | 0.26 | 0.36 | 0.59 | 0.51 | 0.93 | 0.26 | 0.25 | 0.25 | 0.28 | 0.55 | 0.60 | 0.25 |
| UCE | 0.17 | 0.24 | 0.26 | 0.22 | 0.34 | 0.57 | 0.26 | 0.93 | 0.33 | 0.24 | 0.24 | 0.25 | 0.42 | 0.63 | 0.24 |
| PCA | 0.08 | 0.16 | 0.17 | 0.16 | 0.30 | 0.50 | 0.29 | 0.93 | 0.48 | 0.22 | 0.22 | 0.17 | 0.39 | 0.71 | 0.22 |
| scCello | 0.16 | 0.19 | 0.20 | 0.18 | 0.33 | 0.56 | 0.46 | 0.93 | 0.26 | 0.16 | 0.16 | 0.21 | 0.51 | 0.59 | 0.16 |
| CELLama | 0.05 | 0.10 | 0.10 | 0.09 | 0.24 | 0.54 | 0.49 | 0.93 | 0.13 | 0.20 | 0.21 | 0.12 | 0.52 | 0.53 | 0.21 |
| GenePT-w | 0.08 | 0.16 | 0.19 | 0.14 | 0.25 | 0.56 | 0.32 | 0.93 | 0.11 | 0.22 | 0.22 | 0.16 | 0.44 | 0.52 | 0.22 |
| CellFM | 0.01 | 0.03 | 0.02 | 0.05 | 0.35 | 0.37 | 0.04 | 0.94 | 0.59 | 0.15 | 0.15 | 0.09 | 0.21 | 0.77 | 0.15 |
| scBERT | 0.06 | 0.14 | 0.19 | 0.12 | 0.21 | 0.50 | 0.12 | 0.91 | 0.06 | 0.13 | 0.13 | 0.14 | 0.31 | 0.48 | 0.13 |

**Figure S239: Spatial-domain clustering in SpatialLIBD section 151510. a,** Predicted domains and reference annotation. **b,** Scores for 15 displayed methods. Display and scoring conventions are as in Fig. S236.

**a**

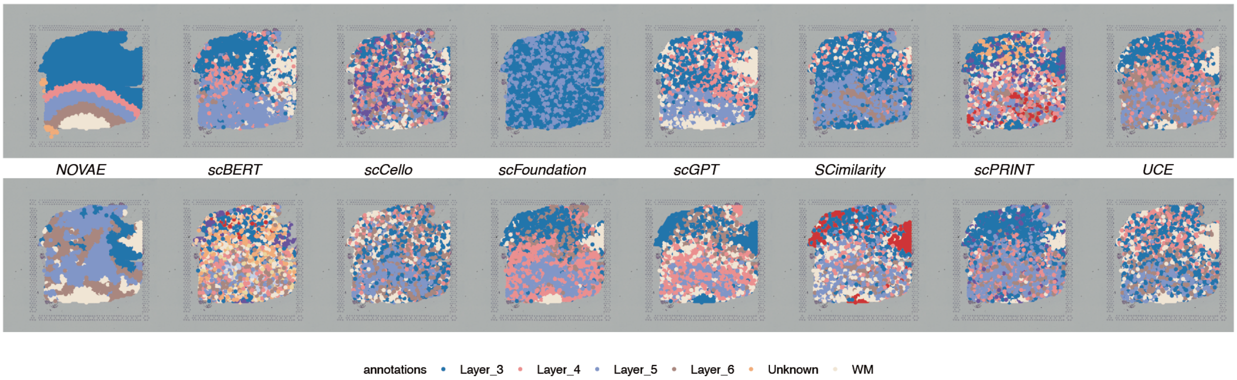

**b**

| method | Cluster concordance: ARI | NMI | HOM | COM | FMI | Local Structure: Acc@kNN | GC | Spatial coherence: 1-CHAOS | 1-PAS | Spatial marker: Moran's I | 1-Geary's C | Aggregated scores: Cluster | Local | Coherence | Marker |
|---|---|---|---|---|---|---|---|---|---|---|---|---|---|---|---|
| NOVAE | 0.04 | 0.17 | 0.17 | 0.16 | 0.37 | 0.86 | 0.98 | 0.94 | 0.90 | 0.19 | 0.18 | 0.18 | 0.92 | 0.92 | 0.18 |
| scFoundation | 0.10 | 0.22 | 0.24 | 0.20 | 0.38 | 0.71 | 0.77 | 0.93 | 0.62 | 0.22 | 0.22 | 0.22 | 0.74 | 0.77 | 0.22 |
| CellPLM | 0.16 | 0.22 | 0.23 | 0.21 | 0.43 | 0.75 | 0.85 | 0.93 | 0.51 | 0.19 | 0.18 | 0.25 | 0.80 | 0.72 | 0.18 |
| Geneformer | 0.15 | 0.18 | 0.19 | 0.17 | 0.45 | 0.75 | 0.73 | 0.93 | 0.58 | 0.24 | 0.24 | 0.23 | 0.74 | 0.75 | 0.24 |
| PCA | 0.17 | 0.24 | 0.25 | 0.23 | 0.45 | 0.68 | 0.35 | 0.93 | 0.70 | 0.21 | 0.21 | 0.27 | 0.52 | 0.82 | 0.21 |
| scGPT | 0.11 | 0.20 | 0.22 | 0.17 | 0.38 | 0.69 | 0.73 | 0.93 | 0.44 | 0.19 | 0.18 | 0.22 | 0.71 | 0.69 | 0.18 |
| LangCell | 0.12 | 0.18 | 0.21 | 0.17 | 0.38 | 0.67 | 0.59 | 0.93 | 0.51 | 0.22 | 0.22 | 0.21 | 0.63 | 0.72 | 0.22 |
| scPRINT | 0.11 | 0.13 | 0.15 | 0.12 | 0.39 | 0.65 | 0.67 | 0.92 | 0.43 | 0.18 | 0.17 | 0.18 | 0.66 | 0.68 | 0.18 |
| SCimilarity | 0.05 | 0.13 | 0.16 | 0.12 | 0.32 | 0.65 | 0.68 | 0.92 | 0.32 | 0.19 | 0.18 | 0.16 | 0.67 | 0.62 | 0.19 |
| UCE | 0.08 | 0.11 | 0.13 | 0.10 | 0.35 | 0.65 | 0.42 | 0.92 | 0.25 | 0.18 | 0.18 | 0.15 | 0.54 | 0.59 | 0.18 |
| scCello | 0.03 | 0.07 | 0.08 | 0.07 | 0.31 | 0.61 | 0.50 | 0.93 | 0.25 | 0.16 | 0.16 | 0.11 | 0.55 | 0.59 | 0.16 |
| CELLama | 0.02 | 0.05 | 0.06 | 0.04 | 0.28 | 0.58 | 0.63 | 0.92 | 0.10 | 0.19 | 0.19 | 0.09 | 0.60 | 0.51 | 0.19 |
| GenePT-w | 0.04 | 0.15 | 0.21 | 0.12 | 0.26 | 0.62 | 0.29 | 0.92 | 0.15 | 0.19 | 0.19 | 0.16 | 0.46 | 0.53 | 0.19 |
| CellFM | -0.01 | 0.00 | 0.00 | 0.00 | 0.45 | 0.35 | 0.17 | 0.94 | 0.66 | 0.02 | 0.02 | 0.09 | 0.26 | 0.80 | 0.02 |
| scBERT | 0.05 | 0.14 | 0.23 | 0.10 | 0.23 | 0.60 | 0.12 | 0.90 | 0.05 | 0.10 | 0.10 | 0.15 | 0.36 | 0.48 | 0.10 |

**Figure S240: Spatial-domain clustering in SpatialLIBD section 151669. a,** Predicted domains and reference annotation. **b,** Scores for 15 displayed methods. Display and scoring conventions are as in Fig. S236.

**a**

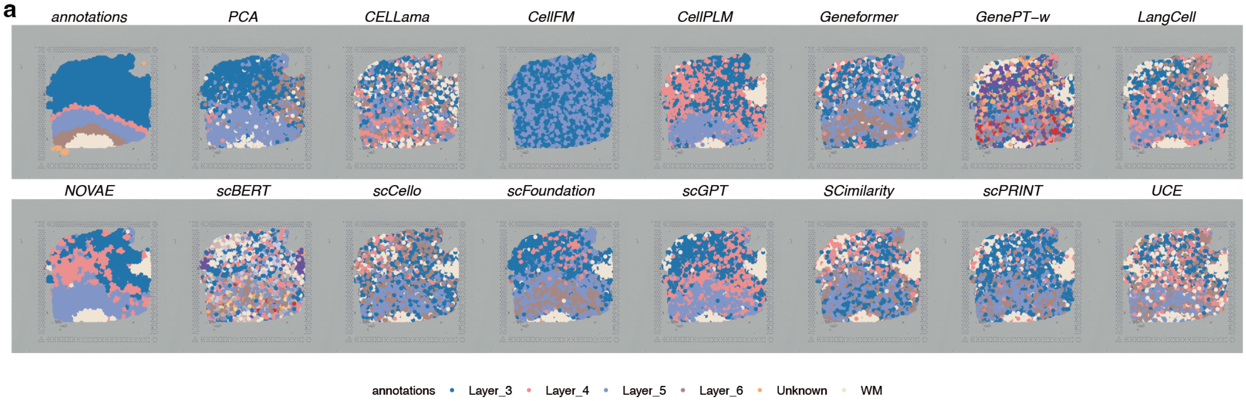

**b**

| | Cluster concordance | | | | | Local Structure | | Spatial coherence | | Spatial marker | | Aggregated scores | | | |
|---|---|---|---|---|---|---|---|---|---|---|---|---|---|---|---|
| method | ARI | NMI | HOM | COM | FMI | Acc@kNN | GC | 1-CHAOS | 1-PAS | Moran's I | 1-Geary's C | Cluster | Local | Coherence | Marker |
| NOVAE | 0.23 | 0.35 | 0.37 | 0.33 | 0.51 | 0.89 | 0.97 | 0.94 | 0.92 | 0.19 | 0.20 | 0.36 | 0.93 | 0.93 | 0.19 |
| CellPLM | 0.24 | 0.28 | 0.29 | 0.26 | 0.52 | 0.79 | 0.82 | 0.93 | 0.68 | 0.21 | 0.21 | 0.32 | 0.81 | 0.81 | 0.21 |
| scGPT | 0.18 | 0.22 | 0.23 | 0.20 | 0.47 | 0.74 | 0.70 | 0.93 | 0.65 | 0.21 | 0.21 | 0.26 | 0.72 | 0.79 | 0.21 |
| scFoundation | 0.13 | 0.21 | 0.23 | 0.19 | 0.43 | 0.75 | 0.75 | 0.93 | 0.61 | 0.20 | 0.20 | 0.24 | 0.75 | 0.77 | 0.20 |
| PCA | 0.22 | 0.24 | 0.25 | 0.23 | 0.52 | 0.71 | 0.35 | 0.93 | 0.70 | 0.21 | 0.21 | 0.29 | 0.53 | 0.82 | 0.21 |
| Geneformer | 0.09 | 0.16 | 0.18 | 0.14 | 0.40 | 0.78 | 0.67 | 0.93 | 0.47 | 0.18 | 0.19 | 0.19 | 0.73 | 0.70 | 0.18 |
| scPRINT | 0.11 | 0.12 | 0.13 | 0.10 | 0.41 | 0.68 | 0.62 | 0.93 | 0.45 | 0.19 | 0.19 | 0.18 | 0.65 | 0.69 | 0.19 |
| LangCell | 0.07 | 0.16 | 0.18 | 0.13 | 0.35 | 0.70 | 0.53 | 0.93 | 0.42 | 0.20 | 0.20 | 0.18 | 0.62 | 0.67 | 0.20 |
| SCimilarity | 0.01 | 0.10 | 0.12 | 0.09 | 0.31 | 0.69 | 0.60 | 0.93 | 0.33 | 0.18 | 0.19 | 0.13 | 0.64 | 0.63 | 0.18 |
| UCE | 0.09 | 0.15 | 0.18 | 0.13 | 0.37 | 0.71 | 0.38 | 0.92 | 0.29 | 0.18 | 0.18 | 0.18 | 0.55 | 0.61 | 0.18 |
| scCello | 0.04 | 0.08 | 0.09 | 0.07 | 0.35 | 0.64 | 0.45 | 0.92 | 0.29 | 0.18 | 0.18 | 0.13 | 0.55 | 0.61 | 0.18 |
| CELLama | 0.05 | 0.06 | 0.07 | 0.05 | 0.34 | 0.64 | 0.55 | 0.92 | 0.18 | 0.18 | 0.18 | 0.11 | 0.59 | 0.55 | 0.18 |
| GenePT-w | 0.06 | 0.14 | 0.20 | 0.10 | 0.29 | 0.65 | 0.32 | 0.92 | 0.16 | 0.17 | 0.17 | 0.16 | 0.49 | 0.54 | 0.17 |
| CellFM | -0.00 | 0.00 | 0.00 | 0.00 | 0.47 | 0.40 | 0.16 | 0.94 | 0.62 | 0.01 | 0.01 | 0.09 | 0.28 | 0.78 | 0.01 |
| scBERT | 0.05 | 0.13 | 0.22 | 0.09 | 0.24 | 0.65 | 0.14 | 0.90 | 0.06 | 0.09 | 0.09 | 0.15 | 0.40 | 0.48 | 0.09 |

**Figure S241: Spatial-domain clustering in SpatialLIBD section 151670. a,** Predicted domains and reference annotation. **b,** Scores for 15 displayed methods. Display and scoring conventions are as in Fig. S236.

**a**

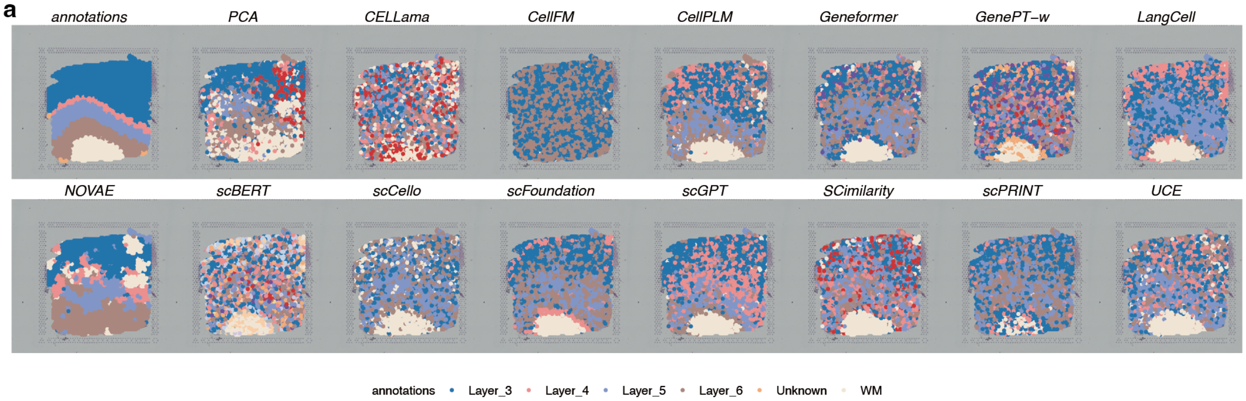

**b**

| | Cluster concordance | | | | | Local Structure | | Spatial coherence | | Spatial marker | | Aggregated scores | | | |
|---|---|---|---|---|---|---|---|---|---|---|---|---|---|---|---|
| method | ARI | NMI | HOM | COM | FMI | Acc@kNN | GC | 1-CHAOS | 1-PAS | Moran's I | 1-Geary's C | Cluster | Local | Coherence | Marker |
| NOVAE | 0.31 | 0.39 | 0.41 | 0.37 | 0.49 | 0.87 | 0.98 | 0.94 | 0.91 | 0.27 | 0.27 | 0.39 | 0.93 | 0.93 | 0.27 |
| scFoundation | 0.21 | 0.29 | 0.30 | 0.28 | 0.43 | 0.71 | 0.80 | 0.94 | 0.61 | 0.30 | 0.30 | 0.30 | 0.76 | 0.77 | 0.30 |
| CellPLM | 0.23 | 0.31 | 0.32 | 0.29 | 0.43 | 0.76 | 0.80 | 0.93 | 0.51 | 0.26 | 0.26 | 0.32 | 0.78 | 0.72 | 0.26 |
| scGPT | 0.26 | 0.31 | 0.33 | 0.30 | 0.46 | 0.72 | 0.74 | 0.93 | 0.54 | 0.25 | 0.25 | 0.33 | 0.73 | 0.74 | 0.25 |
| Geneformer | 0.15 | 0.23 | 0.25 | 0.21 | 0.37 | 0.75 | 0.73 | 0.93 | 0.46 | 0.33 | 0.33 | 0.24 | 0.74 | 0.70 | 0.33 |
| LangCell | 0.15 | 0.25 | 0.26 | 0.25 | 0.39 | 0.63 | 0.62 | 0.93 | 0.57 | 0.34 | 0.33 | 0.26 | 0.63 | 0.75 | 0.33 |
| SCimilarity | 0.10 | 0.19 | 0.21 | 0.17 | 0.32 | 0.62 | 0.76 | 0.93 | 0.24 | 0.29 | 0.28 | 0.20 | 0.69 | 0.59 | 0.28 |
| scPRINT | 0.08 | 0.14 | 0.14 | 0.14 | 0.35 | 0.60 | 0.71 | 0.93 | 0.39 | 0.26 | 0.26 | 0.17 | 0.66 | 0.66 | 0.26 |
| scCello | 0.07 | 0.13 | 0.13 | 0.13 | 0.33 | 0.57 | 0.61 | 0.93 | 0.35 | 0.28 | 0.28 | 0.16 | 0.59 | 0.64 | 0.28 |
| UCE | 0.13 | 0.19 | 0.20 | 0.17 | 0.35 | 0.65 | 0.36 | 0.93 | 0.32 | 0.32 | 0.32 | 0.21 | 0.50 | 0.63 | 0.32 |
| PCA | 0.15 | 0.16 | 0.17 | 0.15 | 0.37 | 0.57 | 0.25 | 0.93 | 0.55 | 0.30 | 0.30 | 0.20 | 0.41 | 0.74 | 0.30 |
| GenePT-w | 0.08 | 0.19 | 0.24 | 0.15 | 0.26 | 0.57 | 0.46 | 0.92 | 0.14 | 0.23 | 0.23 | 0.18 | 0.52 | 0.53 | 0.23 |
| CELLama | 0.02 | 0.06 | 0.07 | 0.05 | 0.25 | 0.54 | 0.58 | 0.93 | 0.10 | 0.20 | 0.20 | 0.09 | 0.56 | 0.51 | 0.20 |
| scBERT | 0.07 | 0.17 | 0.25 | 0.13 | 0.23 | 0.55 | 0.19 | 0.90 | 0.09 | 0.18 | 0.18 | 0.17 | 0.37 | 0.50 | 0.18 |
| CellFM | 0.00 | 0.00 | 0.00 | 0.00 | 0.39 | 0.40 | 0.14 | 0.94 | 0.62 | 0.02 | 0.02 | 0.08 | 0.27 | 0.78 | 0.02 |

**Figure S242: Spatial-domain clustering in SpatialLIBD section 151671. a,** Predicted domains and reference annotation. **b,** Scores for 15 displayed methods. Display and scoring conventions are as in Fig. S236.

**a**

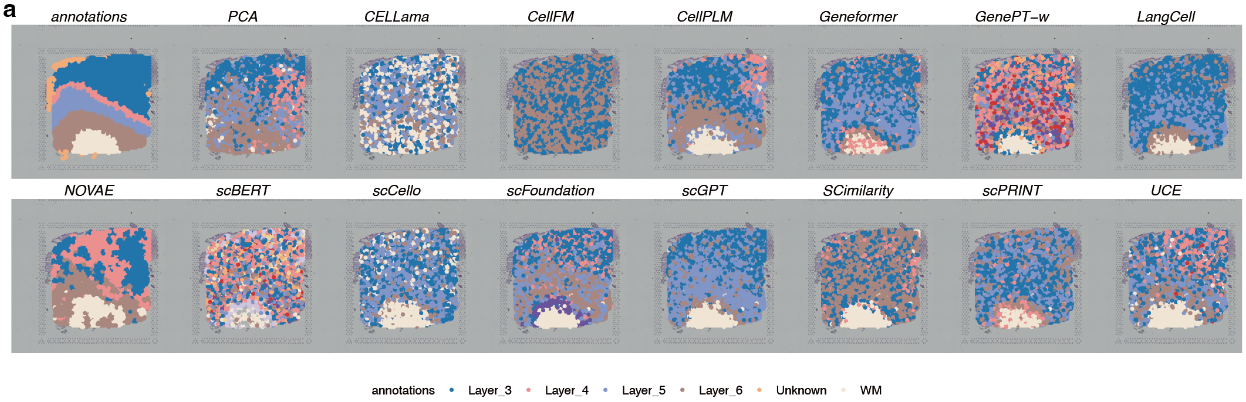

**b**

| | Cluster concordance | | | | | Local Structure | | Spatial coherence | | Spatial marker | | Aggregated scores | | | |
|---|---|---|---|---|---|---|---|---|---|---|---|---|---|---|---|
| method | ARI | NMI | HOM | COM | FMI | Acc@kNN | GC | 1-CHAOS | 1-PAS | Moran's I | 1-Geary's C | Cluster | Local | Coherence | Marker |
| NOVAE | 0.29 | 0.36 | 0.35 | 0.37 | 0.48 | 0.86 | 0.98 | 0.94 | 0.92 | 0.28 | 0.28 | 0.37 | 0.92 | 0.93 | 0.28 |
| CellPLM | 0.21 | 0.30 | 0.30 | 0.30 | 0.42 | 0.73 | 0.84 | 0.93 | 0.59 | 0.24 | 0.24 | 0.30 | 0.79 | 0.76 | 0.24 |
| scGPT | 0.16 | 0.26 | 0.24 | 0.28 | 0.41 | 0.67 | 0.75 | 0.93 | 0.62 | 0.25 | 0.25 | 0.27 | 0.71 | 0.78 | 0.25 |
| scFoundation | 0.16 | 0.27 | 0.29 | 0.25 | 0.36 | 0.67 | 0.81 | 0.93 | 0.47 | 0.25 | 0.25 | 0.27 | 0.74 | 0.70 | 0.25 |
| LangCell | 0.14 | 0.22 | 0.20 | 0.25 | 0.42 | 0.59 | 0.62 | 0.94 | 0.69 | 0.27 | 0.27 | 0.25 | 0.61 | 0.81 | 0.27 |
| SCimilarity | 0.09 | 0.20 | 0.18 | 0.22 | 0.37 | 0.59 | 0.78 | 0.93 | 0.57 | 0.28 | 0.28 | 0.21 | 0.68 | 0.75 | 0.28 |
| Geneformer | 0.11 | 0.19 | 0.18 | 0.20 | 0.38 | 0.70 | 0.74 | 0.93 | 0.58 | 0.24 | 0.24 | 0.21 | 0.72 | 0.76 | 0.24 |
| scPRINT | 0.07 | 0.15 | 0.15 | 0.15 | 0.32 | 0.55 | 0.71 | 0.93 | 0.36 | 0.22 | 0.22 | 0.17 | 0.63 | 0.65 | 0.22 |
| scCello | 0.06 | 0.10 | 0.10 | 0.11 | 0.31 | 0.51 | 0.65 | 0.93 | 0.38 | 0.25 | 0.25 | 0.13 | 0.58 | 0.66 | 0.25 |
| UCE | 0.15 | 0.23 | 0.24 | 0.22 | 0.35 | 0.61 | 0.36 | 0.93 | 0.37 | 0.22 | 0.22 | 0.24 | 0.48 | 0.65 | 0.22 |
| PCA | 0.15 | 0.16 | 0.16 | 0.16 | 0.37 | 0.52 | 0.23 | 0.93 | 0.52 | 0.21 | 0.21 | 0.20 | 0.38 | 0.73 | 0.21 |
| CELLama | 0.03 | 0.04 | 0.04 | 0.04 | 0.29 | 0.49 | 0.57 | 0.93 | 0.25 | 0.22 | 0.22 | 0.09 | 0.53 | 0.59 | 0.22 |
| GenePT-w | 0.08 | 0.17 | 0.22 | 0.14 | 0.24 | 0.51 | 0.44 | 0.92 | 0.09 | 0.21 | 0.21 | 0.17 | 0.48 | 0.50 | 0.21 |
| CellFM | 0.00 | 0.00 | 0.00 | 0.00 | 0.37 | 0.36 | 0.12 | 0.94 | 0.64 | 0.05 | 0.05 | 0.07 | 0.24 | 0.79 | 0.05 |
| scBERT | 0.06 | 0.16 | 0.22 | 0.13 | 0.22 | 0.49 | 0.17 | 0.90 | 0.08 | 0.16 | 0.16 | 0.16 | 0.33 | 0.49 | 0.16 |

**Figure S243: Spatial-domain clustering in SpatialLIBD section 151672. a,** Predicted domains and reference annotation. **b,** Scores for 15 displayed methods. Display and scoring conventions are as in Fig. S236.

a

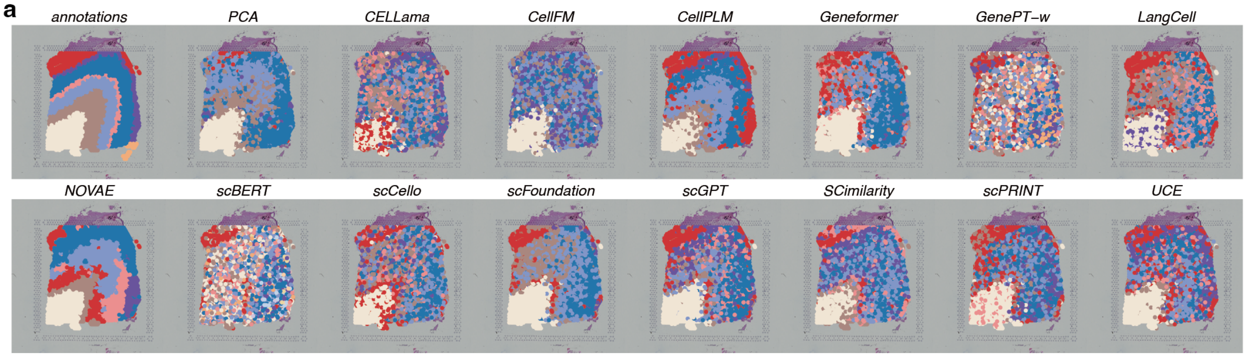

b

| | Cluster concordance | | | | | Local Structure | | Spatial coherence | | Spatial marker | | Aggregated scores | | | |
|---|---|---|---|---|---|---|---|---|---|---|---|---|---|---|---|
| method | ARI | NMI | HOM | COM | FMI | Acc@kNN | GC | 1-CHAOS | 1-PAS | Moran's I | 1-Geary's C | Cluster | Local | Coherence | Marker |
| NOVAE | 0.32 | 0.50 | 0.51 | 0.49 | 0.44 | 0.85 | 0.95 | 0.94 | 0.94 | 0.38 | 0.38 | 0.45 | 0.90 | 0.94 | 0.38 |
| CellPLM | 0.37 | 0.47 | 0.45 | 0.49 | 0.49 | 0.71 | 0.81 | 0.93 | 0.74 | 0.39 | 0.39 | 0.45 | 0.76 | 0.84 | 0.39 |
| scFoundation | 0.21 | 0.32 | 0.30 | 0.34 | 0.36 | 0.63 | 0.68 | 0.93 | 0.66 | 0.39 | 0.39 | 0.31 | 0.66 | 0.79 | 0.39 |
| Geneformer | 0.19 | 0.26 | 0.25 | 0.26 | 0.35 | 0.68 | 0.69 | 0.93 | 0.55 | 0.39 | 0.39 | 0.26 | 0.68 | 0.74 | 0.39 |
| scGPT | 0.23 | 0.31 | 0.32 | 0.31 | 0.36 | 0.61 | 0.72 | 0.93 | 0.47 | 0.38 | 0.38 | 0.31 | 0.66 | 0.70 | 0.38 |
| PCA | 0.24 | 0.33 | 0.30 | 0.36 | 0.40 | 0.55 | 0.33 | 0.93 | 0.72 | 0.38 | 0.38 | 0.33 | 0.44 | 0.82 | 0.38 |
| SCimilarity | 0.16 | 0.28 | 0.28 | 0.28 | 0.31 | 0.53 | 0.68 | 0.93 | 0.40 | 0.40 | 0.40 | 0.26 | 0.61 | 0.66 | 0.40 |
| LangCell | 0.16 | 0.28 | 0.28 | 0.27 | 0.30 | 0.54 | 0.61 | 0.93 | 0.42 | 0.41 | 0.41 | 0.26 | 0.57 | 0.67 | 0.41 |
| scPRINT | 0.15 | 0.26 | 0.27 | 0.26 | 0.30 | 0.53 | 0.63 | 0.93 | 0.36 | 0.40 | 0.40 | 0.25 | 0.58 | 0.64 | 0.40 |
| UCE | 0.20 | 0.29 | 0.30 | 0.29 | 0.34 | 0.59 | 0.42 | 0.92 | 0.38 | 0.40 | 0.40 | 0.28 | 0.50 | 0.65 | 0.40 |
| CELLama | 0.09 | 0.20 | 0.20 | 0.20 | 0.25 | 0.51 | 0.61 | 0.92 | 0.23 | 0.43 | 0.43 | 0.19 | 0.56 | 0.58 | 0.43 |
| scCello | 0.14 | 0.21 | 0.21 | 0.21 | 0.28 | 0.48 | 0.53 | 0.92 | 0.31 | 0.40 | 0.40 | 0.21 | 0.50 | 0.62 | 0.40 |
| CellFM | 0.12 | 0.18 | 0.17 | 0.20 | 0.29 | 0.38 | 0.17 | 0.93 | 0.23 | 0.48 | 0.48 | 0.19 | 0.28 | 0.58 | 0.48 |
| GenePT-w | 0.03 | 0.08 | 0.09 | 0.08 | 0.18 | 0.42 | 0.33 | 0.92 | 0.11 | 0.25 | 0.25 | 0.09 | 0.38 | 0.51 | 0.25 |
| scBERT | 0.04 | 0.09 | 0.11 | 0.08 | 0.16 | 0.32 | 0.18 | 0.89 | 0.08 | 0.14 | 0.14 | 0.10 | 0.25 | 0.49 | 0.14 |

**Figure S244: Spatial-domain clustering in SpatialLIBD section 151673. a,** Predicted domains and reference annotation. **b,** Scores for 15 displayed methods. Display and scoring conventions are as in Fig. S236.

**a**

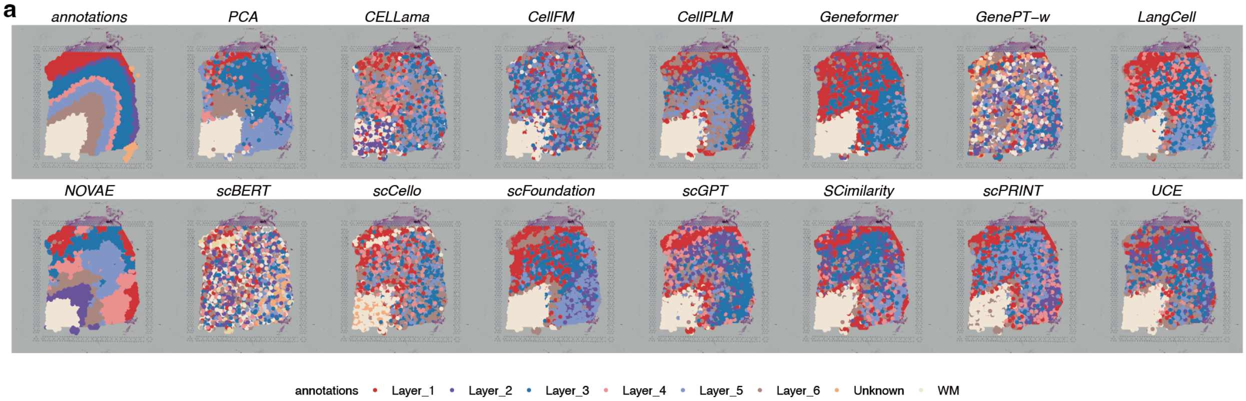

**b**

| | Cluster concordance | | | | | Local Structure | | Spatial coherence | | Spatial marker | | Aggregated scores | | | |
|---|---|---|---|---|---|---|---|---|---|---|---|---|---|---|---|
| method | ARI | NMI | HOM | COM | FMI | Acc@kNN | GC | 1-CHAOS | 1-PAS | Moran's I | 1-Geary's C | Cluster | Local | Coherence | Marker |
| NOVAE | 0.29 | 0.46 | 0.46 | 0.45 | 0.41 | 0.83 | 0.97 | 0.94 | 0.93 | 0.42 | 0.42 | 0.41 | 0.90 | 0.94 | 0.42 |
| CellPLM | 0.30 | 0.40 | 0.39 | 0.41 | 0.42 | 0.66 | 0.80 | 0.93 | 0.60 | 0.34 | 0.35 | 0.38 | 0.73 | 0.77 | 0.35 |
| scFoundation | 0.22 | 0.30 | 0.30 | 0.31 | 0.36 | 0.61 | 0.70 | 0.93 | 0.67 | 0.42 | 0.42 | 0.30 | 0.65 | 0.80 | 0.42 |
| Geneformer | 0.21 | 0.30 | 0.26 | 0.33 | 0.39 | 0.66 | 0.65 | 0.93 | 0.71 | 0.34 | 0.34 | 0.30 | 0.66 | 0.82 | 0.34 |
| scGPT | 0.23 | 0.30 | 0.30 | 0.30 | 0.36 | 0.58 | 0.73 | 0.93 | 0.48 | 0.36 | 0.36 | 0.30 | 0.65 | 0.70 | 0.36 |
| LangCell | 0.23 | 0.29 | 0.28 | 0.30 | 0.37 | 0.55 | 0.54 | 0.93 | 0.53 | 0.43 | 0.44 | 0.29 | 0.54 | 0.73 | 0.43 |
| SCimilarity | 0.22 | 0.27 | 0.27 | 0.26 | 0.35 | 0.54 | 0.68 | 0.93 | 0.44 | 0.42 | 0.42 | 0.27 | 0.61 | 0.68 | 0.42 |
| PCA | 0.23 | 0.30 | 0.29 | 0.31 | 0.37 | 0.54 | 0.34 | 0.93 | 0.75 | 0.39 | 0.39 | 0.30 | 0.44 | 0.84 | 0.39 |
| scPRINT | 0.20 | 0.27 | 0.27 | 0.27 | 0.34 | 0.54 | 0.64 | 0.93 | 0.43 | 0.42 | 0.42 | 0.27 | 0.59 | 0.68 | 0.42 |
| UCE | 0.23 | 0.27 | 0.27 | 0.27 | 0.36 | 0.53 | 0.45 | 0.92 | 0.35 | 0.35 | 0.35 | 0.28 | 0.49 | 0.64 | 0.35 |
| scCello | 0.12 | 0.20 | 0.20 | 0.20 | 0.27 | 0.47 | 0.53 | 0.92 | 0.27 | 0.45 | 0.45 | 0.20 | 0.50 | 0.60 | 0.45 |
| CELLama | 0.10 | 0.18 | 0.18 | 0.18 | 0.25 | 0.50 | 0.62 | 0.92 | 0.20 | 0.36 | 0.36 | 0.18 | 0.56 | 0.56 | 0.36 |
| CellFM | 0.14 | 0.14 | 0.15 | 0.14 | 0.28 | 0.36 | 0.19 | 0.92 | 0.18 | 0.38 | 0.38 | 0.17 | 0.28 | 0.55 | 0.38 |
| GenePT-w | 0.03 | 0.08 | 0.09 | 0.08 | 0.17 | 0.43 | 0.25 | 0.92 | 0.11 | 0.28 | 0.28 | 0.09 | 0.34 | 0.51 | 0.28 |
| scBERT | 0.02 | 0.09 | 0.11 | 0.08 | 0.14 | 0.32 | 0.11 | 0.89 | 0.02 | 0.21 | 0.21 | 0.09 | 0.22 | 0.46 | 0.21 |

**Figure S245: Spatial-domain clustering in SpatialLIBD section 151674. a,** Predicted domains and reference annotation. **b,** Scores for 15 displayed methods. Display and scoring conventions are as in Fig. S236.

**a**

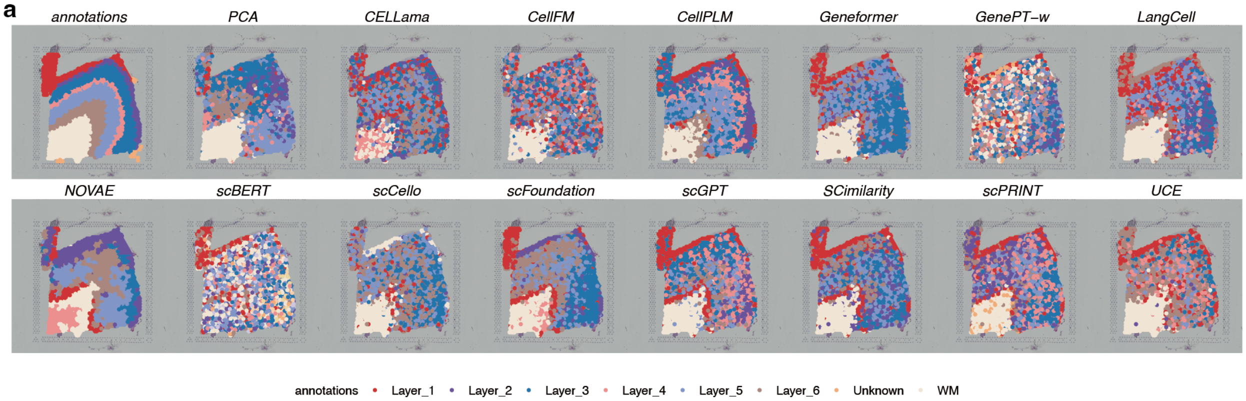

**b**

| | Cluster concordance | | | | | Local Structure | | Spatial coherence | | Spatial marker | | Aggregated scores | | | |
|---|---|---|---|---|---|---|---|---|---|---|---|---|---|---|---|
| **method** | ARI | NMI | HOM | COM | FMI | Acc@kNN | GC | 1-CHAOS | 1-PAS | Moran's I | 1-Geary's C | Cluster | Local | Coherence | Marker |
| **NOVAE** | 0.26 | 0.43 | 0.43 | 0.44 | 0.38 | 0.80 | 0.96 | 0.94 | 0.92 | 0.35 | 0.35 | 0.39 | 0.88 | 0.93 | 0.35 |
| **CellPLM** | 0.28 | 0.40 | 0.40 | 0.41 | 0.40 | 0.65 | 0.83 | 0.93 | 0.56 | 0.33 | 0.33 | 0.38 | 0.74 | 0.74 | 0.33 |
| **scFoundation** | 0.23 | 0.34 | 0.33 | 0.34 | 0.36 | 0.60 | 0.78 | 0.93 | 0.61 | 0.36 | 0.36 | 0.32 | 0.69 | 0.77 | 0.36 |
| **scGPT** | 0.25 | 0.33 | 0.33 | 0.32 | 0.37 | 0.61 | 0.79 | 0.93 | 0.45 | 0.33 | 0.33 | 0.32 | 0.70 | 0.69 | 0.33 |
| **Geneformer** | 0.22 | 0.29 | 0.27 | 0.31 | 0.37 | 0.63 | 0.66 | 0.93 | 0.58 | 0.32 | 0.32 | 0.29 | 0.65 | 0.75 | 0.32 |
| **LangCell** | 0.25 | 0.31 | 0.31 | 0.31 | 0.37 | 0.52 | 0.58 | 0.93 | 0.48 | 0.36 | 0.36 | 0.31 | 0.55 | 0.70 | 0.36 |
| **SCimilarity** | 0.22 | 0.28 | 0.27 | 0.29 | 0.35 | 0.51 | 0.69 | 0.93 | 0.40 | 0.38 | 0.38 | 0.28 | 0.60 | 0.66 | 0.38 |
| **scPRINT** | 0.20 | 0.29 | 0.30 | 0.28 | 0.33 | 0.52 | 0.65 | 0.93 | 0.36 | 0.31 | 0.31 | 0.28 | 0.59 | 0.64 | 0.31 |
| **PCA** | 0.22 | 0.27 | 0.26 | 0.28 | 0.36 | 0.48 | 0.26 | 0.93 | 0.67 | 0.31 | 0.31 | 0.28 | 0.37 | 0.80 | 0.31 |
| **scCello** | 0.16 | 0.20 | 0.18 | 0.21 | 0.32 | 0.45 | 0.45 | 0.93 | 0.39 | 0.37 | 0.37 | 0.21 | 0.45 | 0.66 | 0.37 |
| **UCE** | 0.21 | 0.27 | 0.26 | 0.28 | 0.35 | 0.52 | 0.39 | 0.93 | 0.39 | 0.30 | 0.30 | 0.27 | 0.45 | 0.66 | 0.30 |
| **CELLama** | 0.09 | 0.18 | 0.18 | 0.18 | 0.24 | 0.43 | 0.54 | 0.92 | 0.19 | 0.31 | 0.31 | 0.17 | 0.48 | 0.56 | 0.31 |
| **CellFM** | 0.12 | 0.13 | 0.13 | 0.13 | 0.26 | 0.32 | 0.12 | 0.92 | 0.15 | 0.39 | 0.39 | 0.15 | 0.22 | 0.53 | 0.39 |
| **GenePT-w** | 0.04 | 0.10 | 0.11 | 0.09 | 0.17 | 0.40 | 0.30 | 0.91 | 0.07 | 0.24 | 0.24 | 0.10 | 0.35 | 0.49 | 0.24 |
| **scBERT** | 0.04 | 0.11 | 0.13 | 0.09 | 0.15 | 0.30 | 0.15 | 0.90 | 0.05 | 0.18 | 0.18 | 0.10 | 0.23 | 0.48 | 0.18 |

**Figure S246: Spatial-domain clustering in SpatialLIBD section 151675. a,** Predicted domains and reference annotation. **b,** Scores for 15 displayed methods. Display and scoring conventions are as in Fig. S236.

**a**

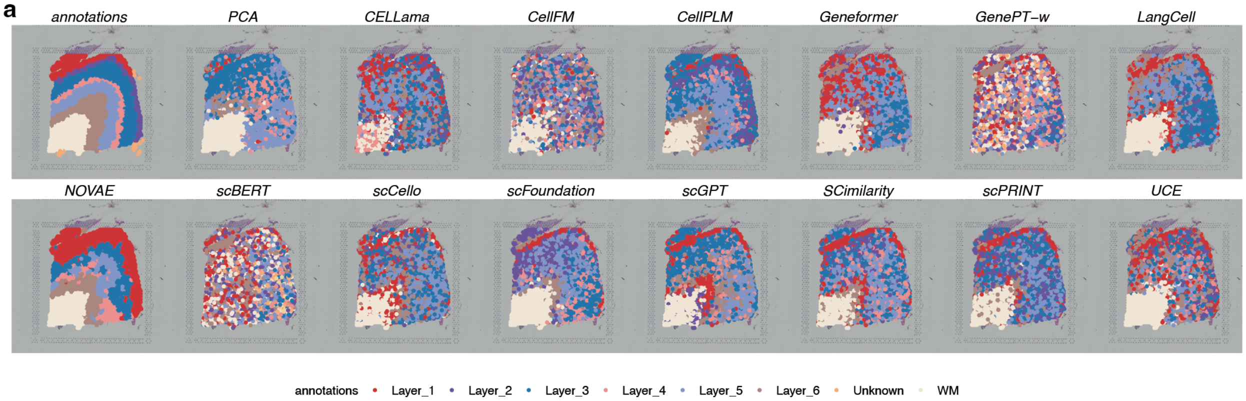

**b**

| | Cluster concordance | | | | | Local Structure | | Spatial coherence | | Spatial marker | | Aggregated scores | | | |
|---|---|---|---|---|---|---|---|---|---|---|---|---|---|---|---|
| method | ARI | NMI | HOM | COM | FMI | Acc@kNN | GC | 1-CHAOS | 1-PAS | Moran's I | 1-Geary's C | Cluster | Local | Coherence | Marker |
| NOVAE | 0.37 | 0.53 | 0.51 | 0.55 | 0.48 | 0.82 | 0.95 | 0.94 | 0.93 | 0.35 | 0.34 | 0.49 | 0.89 | 0.93 | 0.35 |
| CellPLM | 0.23 | 0.38 | 0.37 | 0.39 | 0.37 | 0.62 | 0.79 | 0.93 | 0.58 | 0.28 | 0.29 | 0.35 | 0.70 | 0.75 | 0.29 |
| scFoundation | 0.22 | 0.31 | 0.31 | 0.32 | 0.35 | 0.61 | 0.68 | 0.93 | 0.57 | 0.36 | 0.36 | 0.30 | 0.65 | 0.75 | 0.36 |
| scGPT | 0.19 | 0.31 | 0.31 | 0.31 | 0.33 | 0.59 | 0.76 | 0.93 | 0.53 | 0.33 | 0.33 | 0.29 | 0.67 | 0.73 | 0.33 |
| Geneformer | 0.18 | 0.25 | 0.24 | 0.26 | 0.33 | 0.65 | 0.71 | 0.92 | 0.51 | 0.32 | 0.32 | 0.25 | 0.68 | 0.72 | 0.32 |
| LangCell | 0.19 | 0.27 | 0.26 | 0.28 | 0.34 | 0.51 | 0.57 | 0.93 | 0.52 | 0.35 | 0.35 | 0.27 | 0.54 | 0.72 | 0.35 |
| scPRINT | 0.16 | 0.24 | 0.23 | 0.25 | 0.31 | 0.54 | 0.62 | 0.93 | 0.35 | 0.39 | 0.39 | 0.24 | 0.58 | 0.64 | 0.39 |
| SCimilarity | 0.16 | 0.26 | 0.26 | 0.26 | 0.30 | 0.51 | 0.65 | 0.92 | 0.38 | 0.36 | 0.36 | 0.25 | 0.58 | 0.65 | 0.36 |
| PCA | 0.19 | 0.28 | 0.26 | 0.30 | 0.35 | 0.49 | 0.27 | 0.93 | 0.69 | 0.27 | 0.27 | 0.28 | 0.38 | 0.81 | 0.27 |
| UCE | 0.19 | 0.25 | 0.25 | 0.25 | 0.32 | 0.54 | 0.39 | 0.92 | 0.29 | 0.36 | 0.36 | 0.25 | 0.46 | 0.60 | 0.36 |
| scCello | 0.14 | 0.19 | 0.18 | 0.20 | 0.29 | 0.43 | 0.47 | 0.92 | 0.33 | 0.33 | 0.33 | 0.20 | 0.45 | 0.62 | 0.33 |
| CELLama | 0.08 | 0.16 | 0.16 | 0.17 | 0.24 | 0.44 | 0.57 | 0.92 | 0.23 | 0.30 | 0.30 | 0.16 | 0.51 | 0.58 | 0.30 |
| CellFM | 0.09 | 0.10 | 0.10 | 0.10 | 0.24 | 0.29 | 0.12 | 0.91 | 0.13 | 0.26 | 0.26 | 0.13 | 0.21 | 0.52 | 0.26 |
| GenePT-w | 0.02 | 0.07 | 0.07 | 0.06 | 0.17 | 0.41 | 0.20 | 0.91 | 0.10 | 0.22 | 0.22 | 0.08 | 0.30 | 0.51 | 0.22 |
| scBERT | 0.03 | 0.08 | 0.10 | 0.07 | 0.17 | 0.29 | 0.14 | 0.90 | 0.11 | 0.16 | 0.16 | 0.09 | 0.21 | 0.51 | 0.16 |

**Figure S247: Spatial-domain clustering in SpatialLIBD section 151676. a,** Predicted domains and reference annotation. **b,** Scores for 15 displayed methods. Display and scoring conventions are as in Fig. S236.

#### 2.10.2 *HER2ST spatial-domain clustering*

The eight HER2ST section-level displays are shown in Figs. S248–S255.

**a**

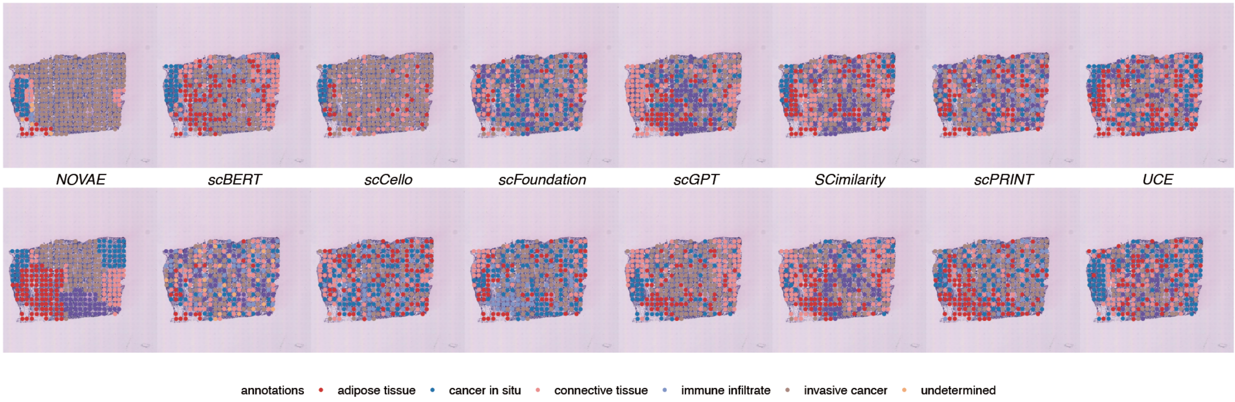

**b**

| | Cluster concordance | | | | | Local Structure | | Spatial coherence | | Spatial marker | | Aggregated scores | | | |
|---|---|---|---|---|---|---|---|---|---|---|---|---|---|---|---|
| method | ARI | NMI | HOM | COM | FMI | Acc@kNN | GC | 1-CHAOS | 1-PAS | Moran's I | 1-Geary's C | Cluster | Local | Coherence | Marker |
| NOVAE | 0.03 | 0.15 | 0.25 | 0.11 | 0.42 | 0.88 | 0.83 | 0.85 | 0.91 | 0.22 | 0.22 | 0.19 | 0.86 | 0.88 | 0.22 |
| scGPT | 0.13 | 0.18 | 0.27 | 0.14 | 0.53 | 0.85 | 0.78 | 0.79 | 0.58 | 0.20 | 0.20 | 0.25 | 0.82 | 0.68 | 0.20 |
| CELLama | 0.26 | 0.14 | 0.16 | 0.13 | 0.72 | 0.84 | 0.54 | 0.81 | 0.73 | 0.11 | 0.12 | 0.28 | 0.69 | 0.77 | 0.12 |
| CellPLM | 0.03 | 0.19 | 0.32 | 0.14 | 0.42 | 0.86 | 0.80 | 0.79 | 0.39 | 0.19 | 0.20 | 0.22 | 0.83 | 0.59 | 0.20 |
| PCA | 0.11 | 0.16 | 0.25 | 0.12 | 0.51 | 0.88 | 0.37 | 0.80 | 0.63 | 0.20 | 0.20 | 0.23 | 0.63 | 0.72 | 0.20 |
| scFoundation | 0.07 | 0.13 | 0.21 | 0.09 | 0.45 | 0.87 | 0.73 | 0.79 | 0.36 | 0.16 | 0.17 | 0.19 | 0.80 | 0.58 | 0.16 |
| SCimilarity | -0.01 | 0.13 | 0.22 | 0.10 | 0.40 | 0.86 | 0.63 | 0.78 | 0.42 | 0.19 | 0.19 | 0.17 | 0.75 | 0.60 | 0.19 |
| Geneformer | 0.04 | 0.14 | 0.24 | 0.10 | 0.43 | 0.88 | 0.63 | 0.79 | 0.29 | 0.18 | 0.18 | 0.19 | 0.76 | 0.54 | 0.18 |
| UCE | -0.01 | 0.14 | 0.23 | 0.10 | 0.39 | 0.84 | 0.49 | 0.78 | 0.23 | 0.16 | 0.17 | 0.17 | 0.66 | 0.50 | 0.16 |
| LangCell | -0.01 | 0.08 | 0.13 | 0.06 | 0.38 | 0.85 | 0.56 | 0.78 | 0.20 | 0.16 | 0.16 | 0.13 | 0.71 | 0.49 | 0.16 |
| scPRINT | 0.04 | 0.08 | 0.12 | 0.06 | 0.46 | 0.86 | 0.45 | 0.77 | 0.35 | 0.10 | 0.10 | 0.15 | 0.66 | 0.56 | 0.10 |
| scCello | 0.03 | 0.11 | 0.19 | 0.08 | 0.41 | 0.85 | 0.49 | 0.75 | 0.11 | 0.13 | 0.13 | 0.16 | 0.67 | 0.43 | 0.13 |
| CellFM | 0.05 | 0.09 | 0.15 | 0.07 | 0.44 | 0.79 | 0.40 | 0.76 | 0.16 | 0.11 | 0.11 | 0.16 | 0.59 | 0.46 | 0.11 |
| GenePT-w | 0.02 | 0.07 | 0.12 | 0.05 | 0.38 | 0.82 | 0.51 | 0.74 | 0.11 | 0.09 | 0.09 | 0.13 | 0.67 | 0.43 | 0.09 |
| scBERT | 0.03 | 0.09 | 0.18 | 0.06 | 0.37 | 0.80 | 0.38 | 0.72 | 0.06 | 0.04 | 0.05 | 0.14 | 0.59 | 0.39 | 0.04 |

**Figure S248: Spatial-domain clustering in HER2ST section A1. a,** Predicted domains from 15 displayed methods and the reference annotation. **b,** Scores for cluster concordance, local neighbourhood structure, spatial coherence and spatial-marker autocorrelation. The four family means receive equal weight; rows are ordered by their aggregate score.

**a**

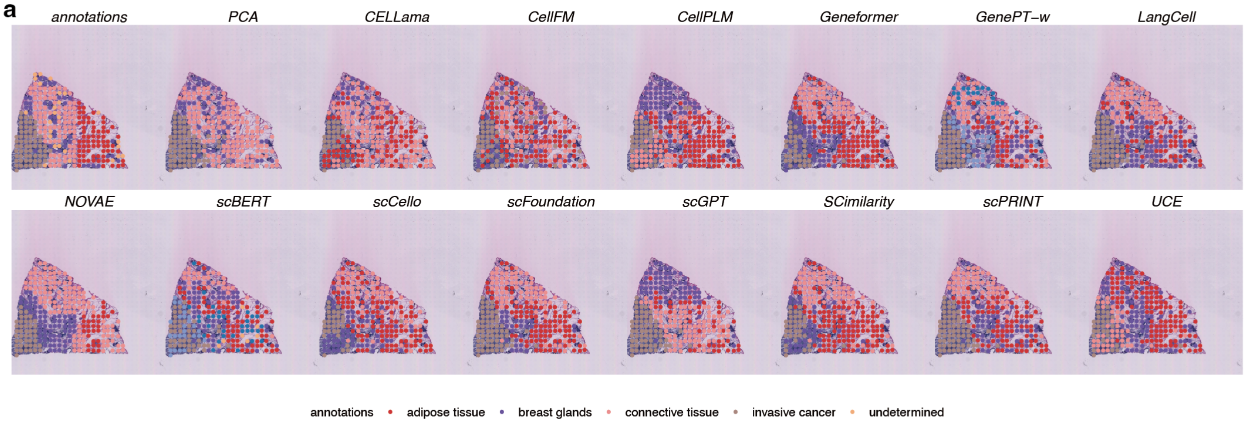

**b**

| | Cluster concordance | | | | | Local Structure | | Spatial coherence | | Spatial marker | | Aggregated scores | | | |
|---|---|---|---|---|---|---|---|---|---|---|---|---|---|---|---|
| method | ARI | NMI | HOM | COM | FMI | Acc@kNN | GC | 1-CHAOS | 1-PAS | Moran's I | 1-Geary's C | Cluster | Local | Coherence | Marker |
| NOVAE | 0.26 | 0.42 | 0.40 | 0.44 | 0.48 | 0.77 | 0.82 | 0.83 | 0.97 | 0.26 | 0.27 | 0.40 | 0.79 | 0.90 | 0.27 |
| CellPLM | 0.25 | 0.33 | 0.32 | 0.33 | 0.46 | 0.65 | 0.83 | 0.80 | 0.73 | 0.27 | 0.27 | 0.34 | 0.74 | 0.77 | 0.27 |
| scPRINT | 0.30 | 0.35 | 0.35 | 0.34 | 0.48 | 0.61 | 0.87 | 0.80 | 0.70 | 0.25 | 0.27 | 0.37 | 0.74 | 0.75 | 0.26 |
| scGPT | 0.26 | 0.32 | 0.32 | 0.32 | 0.46 | 0.63 | 0.80 | 0.80 | 0.79 | 0.22 | 0.24 | 0.34 | 0.72 | 0.80 | 0.23 |
| scFoundation | 0.29 | 0.35 | 0.34 | 0.36 | 0.49 | 0.67 | 0.67 | 0.80 | 0.80 | 0.24 | 0.24 | 0.37 | 0.67 | 0.80 | 0.24 |
| Geneformer | 0.21 | 0.31 | 0.30 | 0.31 | 0.44 | 0.64 | 0.69 | 0.81 | 0.79 | 0.28 | 0.27 | 0.31 | 0.67 | 0.80 | 0.27 |
| SCimilarity | 0.19 | 0.29 | 0.28 | 0.30 | 0.43 | 0.65 | 0.70 | 0.79 | 0.76 | 0.27 | 0.30 | 0.29 | 0.67 | 0.78 | 0.29 |
| PCA | 0.32 | 0.38 | 0.33 | 0.45 | 0.56 | 0.56 | 0.42 | 0.81 | 0.81 | 0.30 | 0.31 | 0.41 | 0.49 | 0.81 | 0.31 |
| UCE | 0.29 | 0.40 | 0.38 | 0.41 | 0.50 | 0.61 | 0.44 | 0.79 | 0.71 | 0.29 | 0.32 | 0.40 | 0.52 | 0.75 | 0.31 |
| LangCell | 0.31 | 0.35 | 0.36 | 0.35 | 0.49 | 0.62 | 0.67 | 0.79 | 0.68 | 0.19 | 0.19 | 0.37 | 0.65 | 0.73 | 0.19 |
| CELLama | 0.11 | 0.20 | 0.19 | 0.21 | 0.39 | 0.57 | 0.58 | 0.79 | 0.65 | 0.25 | 0.27 | 0.22 | 0.57 | 0.72 | 0.26 |
| GenePT-w | 0.22 | 0.29 | 0.33 | 0.26 | 0.40 | 0.58 | 0.55 | 0.77 | 0.47 | 0.21 | 0.22 | 0.30 | 0.56 | 0.62 | 0.21 |
| scCello | 0.14 | 0.21 | 0.21 | 0.21 | 0.38 | 0.52 | 0.48 | 0.78 | 0.61 | 0.19 | 0.19 | 0.23 | 0.50 | 0.69 | 0.19 |
| scBERT | 0.18 | 0.31 | 0.38 | 0.26 | 0.36 | 0.58 | 0.46 | 0.76 | 0.46 | 0.15 | 0.14 | 0.30 | 0.52 | 0.61 | 0.14 |
| CellFM | 0.04 | 0.06 | 0.06 | 0.06 | 0.30 | 0.36 | 0.22 | 0.76 | 0.30 | 0.07 | 0.08 | 0.11 | 0.29 | 0.53 | 0.08 |

**Figure S249: Spatial-domain clustering in HER2ST section B1. a,** Predicted domains and reference annotation. **b,** Scores for 15 displayed methods. Display and scoring conventions are as in Fig. S248.

**a**

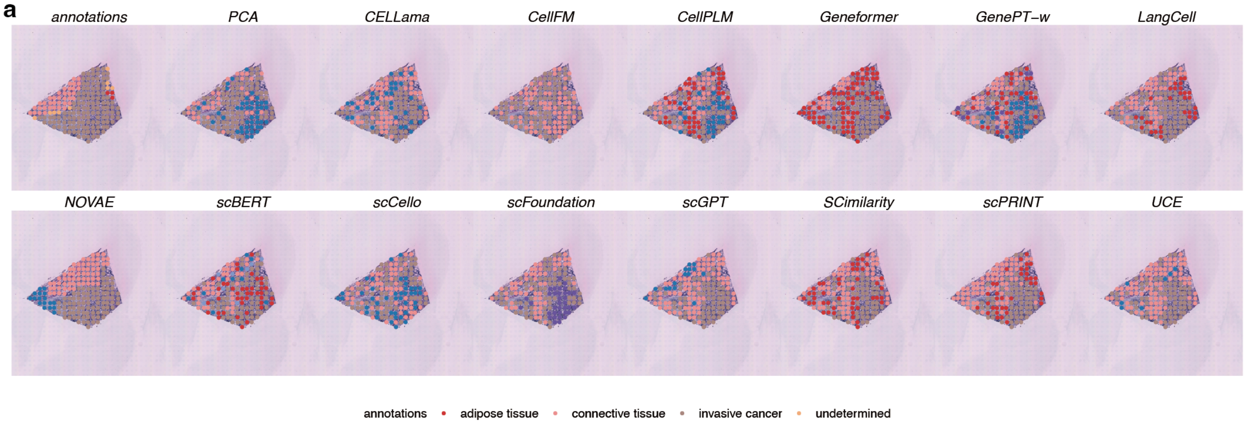

**b**

| | Cluster concordance | | | | | Local Structure | | Spatial coherence | | Spatial marker | | Aggregated scores | | | |
|---|---|---|---|---|---|---|---|---|---|---|---|---|---|---|---|
| method | ARI | NMI | HOM | COM | FMI | Acc@kNN | GC | 1-CHAOS | 1-PAS | Moran's I | 1-Geary's C | Cluster | Local | Coherence | Marker |
| NOVAE | 0.25 | 0.27 | 0.34 | 0.22 | 0.64 | 0.88 | 0.98 | 0.78 | 0.99 | 0.18 | 0.20 | 0.35 | 0.93 | 0.88 | 0.19 |
| scGPT | 0.12 | 0.16 | 0.20 | 0.13 | 0.57 | 0.78 | 0.84 | 0.74 | 0.70 | 0.22 | 0.22 | 0.24 | 0.81 | 0.72 | 0.22 |
| scPRINT | 0.11 | 0.21 | 0.29 | 0.17 | 0.53 | 0.84 | 0.69 | 0.74 | 0.65 | 0.26 | 0.25 | 0.26 | 0.77 | 0.69 | 0.26 |
| Geneformer | 0.16 | 0.18 | 0.24 | 0.15 | 0.58 | 0.83 | 0.60 | 0.74 | 0.72 | 0.22 | 0.22 | 0.26 | 0.72 | 0.73 | 0.22 |
| scFoundation | 0.04 | 0.13 | 0.17 | 0.10 | 0.50 | 0.81 | 0.78 | 0.75 | 0.64 | 0.25 | 0.24 | 0.19 | 0.79 | 0.69 | 0.25 |
| LangCell | 0.09 | 0.13 | 0.18 | 0.11 | 0.54 | 0.81 | 0.60 | 0.75 | 0.62 | 0.26 | 0.27 | 0.21 | 0.71 | 0.68 | 0.27 |
| UCE | 0.01 | 0.06 | 0.06 | 0.05 | 0.54 | 0.81 | 0.73 | 0.74 | 0.66 | 0.24 | 0.23 | 0.14 | 0.77 | 0.70 | 0.24 |
| SCimilarity | 0.02 | 0.05 | 0.07 | 0.04 | 0.47 | 0.81 | 0.76 | 0.73 | 0.51 | 0.24 | 0.24 | 0.13 | 0.79 | 0.62 | 0.24 |
| PCA | 0.03 | 0.06 | 0.08 | 0.05 | 0.52 | 0.76 | 0.32 | 0.73 | 0.69 | 0.22 | 0.23 | 0.15 | 0.54 | 0.71 | 0.23 |
| CellFM | -0.01 | 0.00 | 0.00 | 0.00 | 0.56 | 0.72 | 0.46 | 0.74 | 0.66 | 0.21 | 0.21 | 0.11 | 0.59 | 0.70 | 0.21 |
| CELLama | 0.03 | 0.04 | 0.06 | 0.03 | 0.49 | 0.77 | 0.52 | 0.73 | 0.51 | 0.19 | 0.19 | 0.13 | 0.64 | 0.62 | 0.19 |
| CellPLM | -0.00 | 0.05 | 0.08 | 0.04 | 0.40 | 0.81 | 0.68 | 0.71 | 0.30 | 0.22 | 0.20 | 0.11 | 0.75 | 0.50 | 0.21 |
| scBERT | 0.13 | 0.14 | 0.22 | 0.10 | 0.51 | 0.79 | 0.49 | 0.66 | 0.43 | 0.16 | 0.16 | 0.22 | 0.64 | 0.54 | 0.16 |
| GenePT-w | 0.05 | 0.12 | 0.21 | 0.09 | 0.42 | 0.78 | 0.66 | 0.70 | 0.26 | 0.16 | 0.14 | 0.18 | 0.72 | 0.48 | 0.15 |
| scCello | -0.01 | 0.01 | 0.02 | 0.01 | 0.46 | 0.75 | 0.48 | 0.71 | 0.40 | 0.20 | 0.18 | 0.10 | 0.62 | 0.55 | 0.19 |

**Figure S250: Spatial-domain clustering in HER2ST section C1. a,** Predicted domains and reference annotation. **b,** Scores for 15 displayed methods. Display and scoring conventions are as in Fig. S248.

**a**

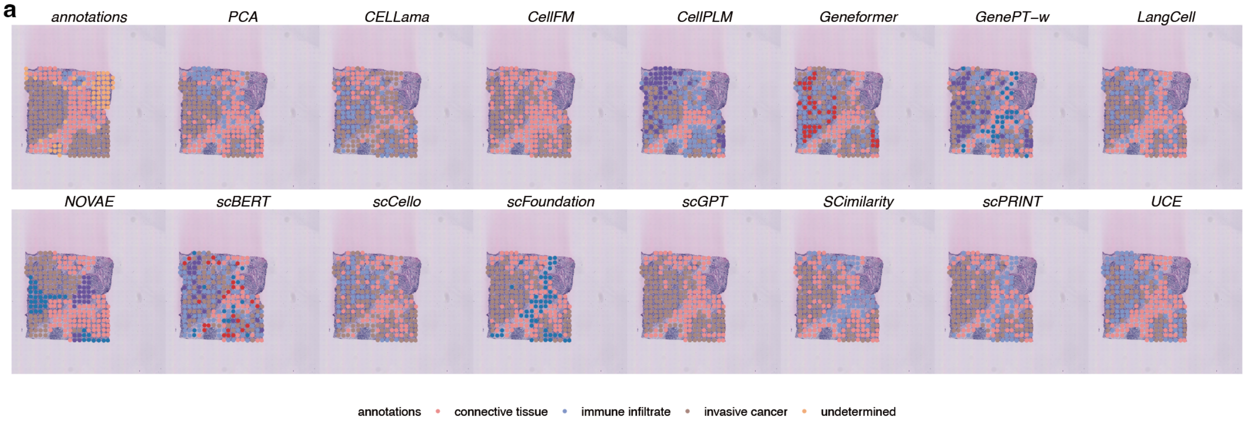

**b**

| | Cluster concordance | | | | | Local Structure | | Spatial coherence | | Spatial marker | | Aggregated scores | | | |
|---|---|---|---|---|---|---|---|---|---|---|---|---|---|---|---|
| method | ARI | NMI | HOM | COM | FMI | Acc@kNN | GC | 1-CHAOS | 1-PAS | Moran's I | 1-Geary's C | Cluster | Local | Coherence | Marker |
| scGPT | 0.35 | 0.26 | 0.25 | 0.28 | 0.67 | 0.78 | 0.72 | 0.81 | 0.85 | 0.40 | 0.40 | 0.36 | 0.75 | 0.83 | 0.40 |
| NOVAE | 0.04 | 0.15 | 0.19 | 0.12 | 0.42 | 0.84 | 0.90 | 0.81 | 0.88 | 0.34 | 0.35 | 0.18 | 0.87 | 0.85 | 0.35 |
| SCimilarity | 0.20 | 0.23 | 0.27 | 0.20 | 0.53 | 0.80 | 0.86 | 0.80 | 0.67 | 0.33 | 0.34 | 0.29 | 0.83 | 0.73 | 0.33 |
| scFoundation | 0.25 | 0.22 | 0.26 | 0.20 | 0.56 | 0.79 | 0.82 | 0.79 | 0.67 | 0.33 | 0.34 | 0.30 | 0.81 | 0.73 | 0.33 |
| scPRINT | 0.29 | 0.22 | 0.26 | 0.20 | 0.58 | 0.76 | 0.80 | 0.78 | 0.60 | 0.37 | 0.37 | 0.31 | 0.78 | 0.69 | 0.37 |
| CellPLM | 0.11 | 0.16 | 0.21 | 0.12 | 0.43 | 0.76 | 0.93 | 0.79 | 0.56 | 0.33 | 0.33 | 0.21 | 0.85 | 0.67 | 0.33 |
| LangCell | 0.20 | 0.19 | 0.22 | 0.16 | 0.53 | 0.73 | 0.80 | 0.78 | 0.56 | 0.35 | 0.35 | 0.26 | 0.77 | 0.67 | 0.35 |
| UCE | 0.16 | 0.17 | 0.20 | 0.15 | 0.50 | 0.73 | 0.65 | 0.79 | 0.64 | 0.33 | 0.33 | 0.24 | 0.69 | 0.71 | 0.33 |
| CellFM | 0.15 | 0.17 | 0.15 | 0.18 | 0.58 | 0.62 | 0.47 | 0.79 | 0.81 | 0.37 | 0.38 | 0.24 | 0.54 | 0.80 | 0.37 |
| PCA | 0.19 | 0.22 | 0.25 | 0.19 | 0.54 | 0.65 | 0.40 | 0.79 | 0.74 | 0.38 | 0.38 | 0.28 | 0.52 | 0.77 | 0.38 |
| Geneformer | 0.13 | 0.19 | 0.25 | 0.15 | 0.45 | 0.76 | 0.77 | 0.78 | 0.47 | 0.26 | 0.26 | 0.23 | 0.76 | 0.62 | 0.26 |
| CELLama | 0.05 | 0.10 | 0.12 | 0.09 | 0.46 | 0.74 | 0.74 | 0.78 | 0.55 | 0.29 | 0.30 | 0.16 | 0.74 | 0.66 | 0.30 |
| scCello | 0.06 | 0.06 | 0.07 | 0.06 | 0.46 | 0.66 | 0.75 | 0.78 | 0.51 | 0.30 | 0.31 | 0.14 | 0.70 | 0.65 | 0.31 |
| GenePT-w | 0.13 | 0.16 | 0.23 | 0.12 | 0.42 | 0.71 | 0.58 | 0.75 | 0.27 | 0.27 | 0.26 | 0.21 | 0.64 | 0.51 | 0.27 |
| scBERT | 0.07 | 0.10 | 0.16 | 0.08 | 0.38 | 0.69 | 0.46 | 0.74 | 0.23 | 0.17 | 0.18 | 0.16 | 0.58 | 0.48 | 0.17 |

**Figure S251: Spatial-domain clustering in HER2ST section D1. a,** Predicted domains and reference annotation. **b,** Scores for 15 displayed methods. Display and scoring conventions are as in Fig. S248.

**a**

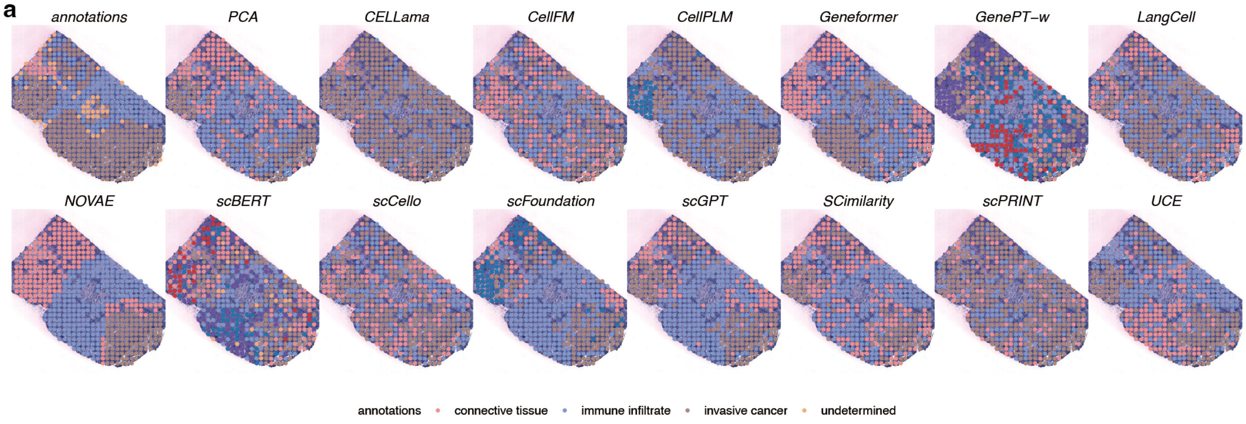

**b**

| | Cluster concordance | | | | | Local Structure | | Spatial coherence | | Spatial marker | | Aggregated scores | | | |
|---|---|---|---|---|---|---|---|---|---|---|---|---|---|---|---|
| **method** | ARI | NMI | HOM | COM | FMI | Acc@kNN | GC | 1-CHAOS | 1-PAS | Moran's I | 1-Geary's C | Cluster | Local | Coherence | Marker |
| **NOVAE** | 0.04 | 0.16 | 0.19 | 0.14 | 0.45 | 0.86 | 0.95 | 0.87 | 0.96 | 0.34 | 0.33 | 0.19 | 0.90 | 0.92 | 0.34 |
| **CellPLM** | 0.09 | 0.08 | 0.09 | 0.08 | 0.54 | 0.76 | 0.82 | 0.86 | 0.76 | 0.30 | 0.31 | 0.17 | 0.79 | 0.81 | 0.31 |
| **scFoundation** | 0.06 | 0.10 | 0.12 | 0.08 | 0.46 | 0.78 | 0.85 | 0.86 | 0.72 | 0.30 | 0.31 | 0.16 | 0.82 | 0.79 | 0.31 |
| **scGPT** | 0.12 | 0.16 | 0.19 | 0.14 | 0.51 | 0.75 | 0.71 | 0.86 | 0.75 | 0.30 | 0.30 | 0.22 | 0.73 | 0.81 | 0.30 |
| **SCimilarity** | 0.07 | 0.13 | 0.16 | 0.11 | 0.47 | 0.74 | 0.78 | 0.86 | 0.60 | 0.27 | 0.27 | 0.19 | 0.76 | 0.73 | 0.27 |
| **Geneformer** | 0.01 | 0.03 | 0.04 | 0.03 | 0.42 | 0.74 | 0.80 | 0.86 | 0.74 | 0.24 | 0.25 | 0.11 | 0.77 | 0.80 | 0.25 |
| **UCE** | 0.16 | 0.19 | 0.23 | 0.16 | 0.52 | 0.72 | 0.35 | 0.86 | 0.70 | 0.30 | 0.30 | 0.25 | 0.54 | 0.78 | 0.30 |
| **LangCell** | 0.02 | 0.04 | 0.05 | 0.04 | 0.45 | 0.71 | 0.81 | 0.86 | 0.50 | 0.25 | 0.25 | 0.12 | 0.76 | 0.68 | 0.25 |
| **PCA** | 0.07 | 0.12 | 0.15 | 0.11 | 0.46 | 0.64 | 0.37 | 0.86 | 0.64 | 0.30 | 0.29 | 0.18 | 0.50 | 0.75 | 0.30 |
| **CELLama** | 0.02 | 0.04 | 0.04 | 0.05 | 0.60 | 0.69 | 0.62 | 0.86 | 0.82 | 0.04 | 0.04 | 0.15 | 0.65 | 0.84 | 0.04 |
| **scCello** | 0.09 | 0.06 | 0.07 | 0.05 | 0.48 | 0.66 | 0.68 | 0.85 | 0.44 | 0.13 | 0.13 | 0.15 | 0.67 | 0.65 | 0.13 |
| **scPRINT** | -0.00 | 0.00 | 0.00 | 0.00 | 0.51 | 0.77 | 0.64 | 0.85 | 0.64 | 0.01 | 0.02 | 0.10 | 0.70 | 0.74 | 0.02 |
| **GenePT-w** | 0.01 | 0.04 | 0.06 | 0.03 | 0.32 | 0.61 | 0.36 | 0.84 | 0.34 | 0.22 | 0.23 | 0.09 | 0.49 | 0.59 | 0.22 |
| **CellFM** | 0.02 | 0.02 | 0.03 | 0.02 | 0.42 | 0.60 | 0.08 | 0.85 | 0.37 | 0.22 | 0.22 | 0.10 | 0.34 | 0.61 | 0.22 |
| **scBERT** | -0.00 | 0.04 | 0.07 | 0.03 | 0.28 | 0.64 | 0.31 | 0.82 | 0.27 | 0.15 | 0.15 | 0.08 | 0.47 | 0.54 | 0.15 |

**Figure S252: Spatial-domain clustering in HER2ST section E1. a,** Predicted domains and reference annotation. **b,** Scores for 15 displayed methods. Display and scoring conventions are as in Fig. S248.

**a**

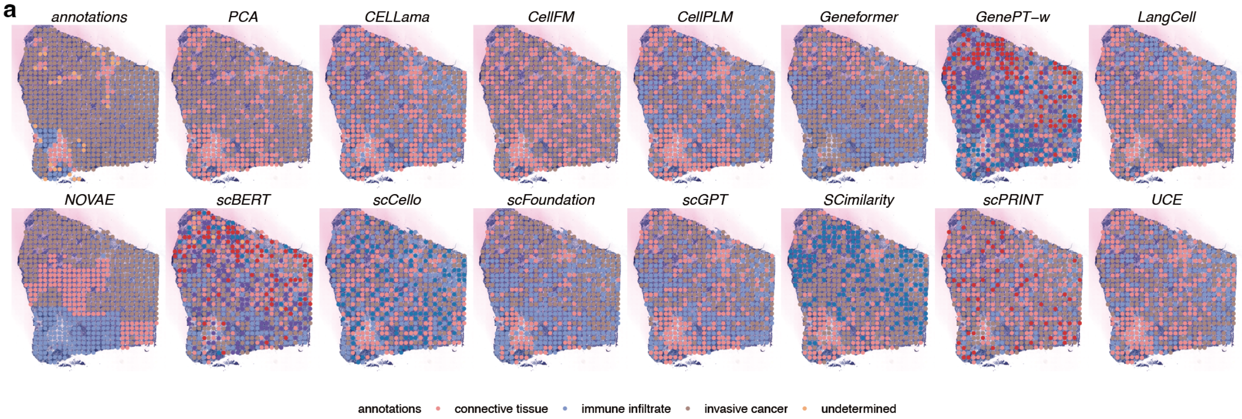

**b**

| | Cluster concordance | | | | | Local Structure | | Spatial coherence | | Spatial marker | | Aggregated scores | | | |
|---|---|---|---|---|---|---|---|---|---|---|---|---|---|---|---|
| method | ARI | NMI | HOM | COM | FMI | Acc@kNN | GC | 1-CHAOS | 1-PAS | Moran's I | 1-Geary's C | Cluster | Local | Coherence | Marker |
| NOVAE | 0.08 | 0.11 | 0.17 | 0.08 | 0.59 | 0.91 | 0.74 | 0.87 | 0.96 | 0.12 | 0.12 | 0.20 | 0.82 | 0.91 | 0.12 |
| PCA | 0.22 | 0.23 | 0.28 | 0.19 | 0.73 | 0.85 | 0.42 | 0.87 | 0.82 | 0.16 | 0.16 | 0.33 | 0.64 | 0.84 | 0.16 |
| scFoundation | 0.12 | 0.13 | 0.21 | 0.09 | 0.60 | 0.90 | 0.78 | 0.86 | 0.67 | 0.07 | 0.08 | 0.23 | 0.84 | 0.76 | 0.07 |
| SCimilarity | 0.08 | 0.13 | 0.22 | 0.09 | 0.58 | 0.87 | 0.70 | 0.85 | 0.63 | 0.09 | 0.09 | 0.22 | 0.79 | 0.74 | 0.09 |
| scGPT | 0.04 | 0.13 | 0.22 | 0.09 | 0.53 | 0.88 | 0.72 | 0.86 | 0.58 | 0.10 | 0.11 | 0.20 | 0.80 | 0.72 | 0.11 |
| CellPLM | 0.02 | 0.16 | 0.28 | 0.11 | 0.52 | 0.91 | 0.71 | 0.86 | 0.48 | 0.10 | 0.11 | 0.22 | 0.81 | 0.67 | 0.11 |
| UCE | 0.02 | 0.13 | 0.22 | 0.09 | 0.53 | 0.89 | 0.55 | 0.86 | 0.44 | 0.10 | 0.11 | 0.20 | 0.72 | 0.65 | 0.10 |
| Geneformer | -0.03 | 0.02 | 0.03 | 0.01 | 0.57 | 0.89 | 0.59 | 0.86 | 0.64 | 0.04 | 0.04 | 0.12 | 0.74 | 0.75 | 0.04 |
| CELLama | -0.01 | 0.11 | 0.18 | 0.08 | 0.51 | 0.88 | 0.58 | 0.85 | 0.39 | 0.08 | 0.08 | 0.17 | 0.73 | 0.62 | 0.08 |
| CellFM | 0.05 | 0.07 | 0.09 | 0.06 | 0.64 | 0.82 | 0.26 | 0.87 | 0.61 | 0.09 | 0.08 | 0.18 | 0.54 | 0.74 | 0.08 |
| LangCell | -0.02 | 0.04 | 0.06 | 0.03 | 0.52 | 0.87 | 0.57 | 0.85 | 0.45 | 0.05 | 0.05 | 0.13 | 0.72 | 0.65 | 0.05 |
| scPRINT | -0.00 | 0.01 | 0.01 | 0.00 | 0.53 | 0.87 | 0.65 | 0.84 | 0.36 | 0.02 | 0.02 | 0.11 | 0.76 | 0.60 | 0.02 |
| scCello | 0.00 | 0.04 | 0.08 | 0.03 | 0.44 | 0.85 | 0.52 | 0.84 | 0.16 | 0.03 | 0.04 | 0.12 | 0.69 | 0.50 | 0.04 |
| GenePT-w | 0.00 | 0.02 | 0.06 | 0.01 | 0.37 | 0.86 | 0.44 | 0.82 | 0.11 | 0.02 | 0.02 | 0.09 | 0.65 | 0.46 | 0.02 |
| scBERT | -0.00 | 0.01 | 0.02 | 0.00 | 0.39 | 0.90 | 0.20 | 0.82 | 0.11 | 0.01 | 0.01 | 0.08 | 0.55 | 0.46 | 0.01 |

**Figure S253: Spatial-domain clustering in HER2ST section F1. a,** Predicted domains and reference annotation. **b,** Scores for 15 displayed methods. Display and scoring conventions are as in Fig. S248.

**a**

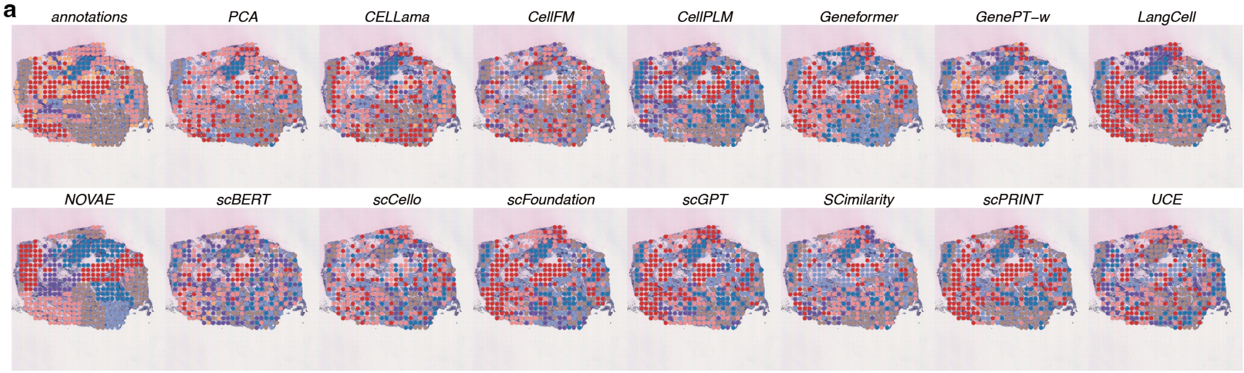

**b**

| method | Cluster concordance: ARI | NMI | HOM | COM | FMI | Local Structure: Acc@kNN | GC | Spatial coherence: 1-CHAOS | 1-PAS | Spatial marker: Moran's I | 1-Geary's C | Aggregated scores: Cluster | Local | Coherence | Marker |
|---|---|---|---|---|---|---|---|---|---|---|---|---|---|---|---|
| NOVAE | 0.09 | 0.20 | 0.21 | 0.19 | 0.29 | 0.69 | 0.79 | 0.86 | 0.88 | 0.20 | 0.20 | 0.20 | 0.74 | 0.87 | 0.20 |
| scFoundation | 0.17 | 0.26 | 0.28 | 0.25 | 0.36 | 0.66 | 0.73 | 0.81 | 0.37 | 0.24 | 0.24 | 0.27 | 0.70 | 0.59 | 0.24 |
| scGPT | 0.17 | 0.26 | 0.27 | 0.24 | 0.36 | 0.65 | 0.74 | 0.81 | 0.38 | 0.23 | 0.23 | 0.26 | 0.69 | 0.60 | 0.23 |
| LangCell | 0.16 | 0.24 | 0.24 | 0.24 | 0.38 | 0.60 | 0.59 | 0.81 | 0.49 | 0.24 | 0.24 | 0.25 | 0.59 | 0.65 | 0.24 |
| scPRINT | 0.19 | 0.23 | 0.23 | 0.23 | 0.40 | 0.60 | 0.67 | 0.81 | 0.43 | 0.22 | 0.22 | 0.26 | 0.64 | 0.62 | 0.22 |
| Geneformer | 0.17 | 0.24 | 0.25 | 0.23 | 0.36 | 0.66 | 0.66 | 0.82 | 0.35 | 0.20 | 0.20 | 0.25 | 0.66 | 0.58 | 0.20 |
| CellPLM | 0.14 | 0.22 | 0.24 | 0.20 | 0.33 | 0.60 | 0.77 | 0.80 | 0.29 | 0.23 | 0.23 | 0.22 | 0.69 | 0.55 | 0.23 |
| SCimilarity | 0.11 | 0.23 | 0.25 | 0.21 | 0.31 | 0.63 | 0.80 | 0.80 | 0.32 | 0.18 | 0.18 | 0.22 | 0.71 | 0.56 | 0.18 |
| PCA | 0.14 | 0.24 | 0.24 | 0.24 | 0.36 | 0.60 | 0.39 | 0.82 | 0.41 | 0.28 | 0.29 | 0.25 | 0.50 | 0.61 | 0.28 |
| CELLama | 0.15 | 0.23 | 0.25 | 0.21 | 0.34 | 0.56 | 0.50 | 0.80 | 0.37 | 0.23 | 0.23 | 0.23 | 0.53 | 0.58 | 0.23 |
| UCE | 0.12 | 0.18 | 0.20 | 0.17 | 0.31 | 0.56 | 0.45 | 0.80 | 0.24 | 0.23 | 0.23 | 0.19 | 0.51 | 0.52 | 0.23 |
| GenePT-w | 0.12 | 0.19 | 0.22 | 0.17 | 0.29 | 0.46 | 0.42 | 0.78 | 0.17 | 0.19 | 0.19 | 0.20 | 0.44 | 0.48 | 0.19 |
| scCello | 0.04 | 0.12 | 0.13 | 0.11 | 0.26 | 0.49 | 0.40 | 0.79 | 0.12 | 0.20 | 0.20 | 0.13 | 0.44 | 0.46 | 0.20 |
| scBERT | 0.09 | 0.20 | 0.25 | 0.17 | 0.26 | 0.48 | 0.26 | 0.74 | 0.08 | 0.15 | 0.15 | 0.19 | 0.37 | 0.41 | 0.15 |
| CellFM | 0.03 | 0.06 | 0.06 | 0.06 | 0.27 | 0.39 | 0.19 | 0.78 | 0.15 | 0.15 | 0.14 | 0.10 | 0.29 | 0.47 | 0.14 |

**Figure S254: Spatial-domain clustering in HER2ST section G2. a,** Predicted domains and reference annotation. **b,** Scores for 15 displayed methods. Display and scoring conventions are as in Fig. S248.

**a**

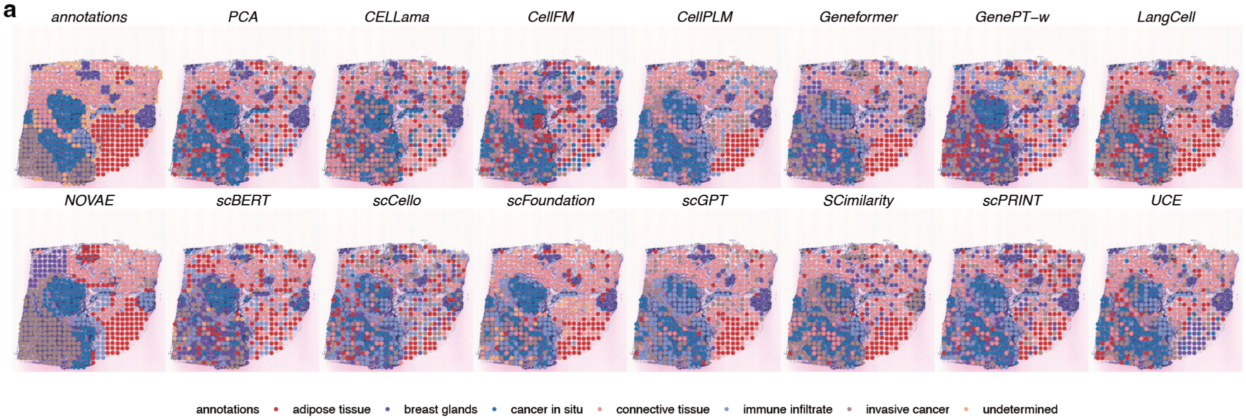

**b**

| | Cluster concordance | | | | | Local Structure | | Spatial coherence | | Spatial marker | | Aggregated scores | | | |
|---|---|---|---|---|---|---|---|---|---|---|---|---|---|---|---|
| method | ARI | NMI | HOM | COM | FMI | Acc@kNN | GC | 1-CHAOS | 1-PAS | Moran's I | 1-Geary's C | Cluster | Local | Coherence | Marker |
| NOVAE | 0.32 | 0.44 | 0.45 | 0.44 | 0.46 | 0.80 | 0.84 | 0.87 | 0.89 | 0.21 | 0.22 | 0.42 | 0.82 | 0.88 | 0.22 |
| CellPLM | 0.27 | 0.36 | 0.37 | 0.36 | 0.42 | 0.69 | 0.86 | 0.83 | 0.49 | 0.24 | 0.23 | 0.35 | 0.77 | 0.66 | 0.24 |
| scFoundation | 0.34 | 0.39 | 0.40 | 0.38 | 0.48 | 0.63 | 0.81 | 0.83 | 0.56 | 0.21 | 0.21 | 0.40 | 0.72 | 0.70 | 0.21 |
| scGPT | 0.27 | 0.36 | 0.36 | 0.35 | 0.43 | 0.57 | 0.81 | 0.83 | 0.54 | 0.24 | 0.23 | 0.35 | 0.69 | 0.68 | 0.23 |
| Geneformer | 0.24 | 0.31 | 0.32 | 0.30 | 0.40 | 0.63 | 0.83 | 0.84 | 0.49 | 0.18 | 0.19 | 0.31 | 0.73 | 0.66 | 0.19 |
| LangCell | 0.22 | 0.30 | 0.28 | 0.32 | 0.41 | 0.59 | 0.68 | 0.84 | 0.56 | 0.23 | 0.24 | 0.30 | 0.63 | 0.70 | 0.23 |
| SCimilarity | 0.20 | 0.29 | 0.28 | 0.29 | 0.38 | 0.62 | 0.75 | 0.83 | 0.51 | 0.20 | 0.21 | 0.29 | 0.68 | 0.67 | 0.20 |
| scPRINT | 0.15 | 0.24 | 0.24 | 0.23 | 0.31 | 0.57 | 0.83 | 0.83 | 0.29 | 0.21 | 0.22 | 0.23 | 0.70 | 0.56 | 0.22 |
| UCE | 0.27 | 0.30 | 0.30 | 0.29 | 0.43 | 0.53 | 0.42 | 0.83 | 0.47 | 0.19 | 0.19 | 0.32 | 0.47 | 0.65 | 0.19 |
| PCA | 0.22 | 0.27 | 0.28 | 0.27 | 0.38 | 0.51 | 0.26 | 0.81 | 0.43 | 0.23 | 0.23 | 0.28 | 0.38 | 0.62 | 0.23 |
| CELLama | 0.08 | 0.15 | 0.15 | 0.15 | 0.29 | 0.42 | 0.57 | 0.82 | 0.30 | 0.15 | 0.16 | 0.16 | 0.50 | 0.56 | 0.16 |
| scCello | 0.06 | 0.13 | 0.13 | 0.12 | 0.25 | 0.46 | 0.62 | 0.81 | 0.13 | 0.17 | 0.17 | 0.14 | 0.54 | 0.47 | 0.17 |
| GenePT-w | 0.11 | 0.22 | 0.25 | 0.20 | 0.27 | 0.46 | 0.38 | 0.80 | 0.15 | 0.15 | 0.15 | 0.21 | 0.42 | 0.47 | 0.15 |
| scBERT | 0.15 | 0.20 | 0.22 | 0.19 | 0.31 | 0.45 | 0.28 | 0.81 | 0.25 | 0.13 | 0.12 | 0.22 | 0.36 | 0.53 | 0.12 |
| CellFM | 0.09 | 0.12 | 0.11 | 0.12 | 0.29 | 0.21 | 0.10 | 0.82 | 0.26 | 0.19 | 0.19 | 0.15 | 0.16 | 0.54 | 0.19 |

**Figure S255: Spatial-domain clustering in HER2ST section H1. a,** Predicted domains and reference annotation. **b,** Scores for 15 displayed methods. Display and scoring conventions are as in Fig. S248.

##### *2.10.3 MERFISH spatial-domain clustering*

The five MERFISH spatial-domain displays are shown in Figs. S256–S260.

**a**

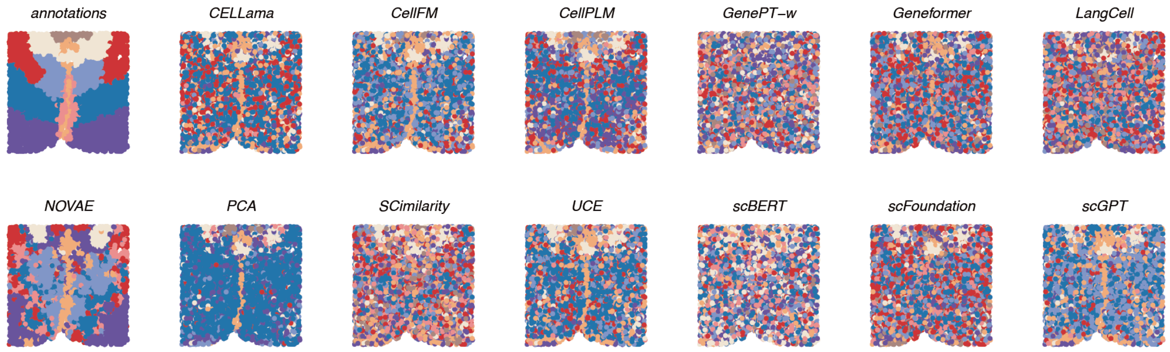

**b**

| method | Cluster concordance | | | | | Local Structure | | Spatial coherence | | Spatial marker | | Aggregated scores | | | |
|---|---|---|---|---|---|---|---|---|---|---|---|---|---|---|---|
| | ARI | NMI | HOM | COM | FMI | Acc@kNN | GC | 1-CHAOS | 1-PAS | Moran's I | 1-Geary's C | Cluster | Local | Coherence | Marker |
| NOVAE | 0.16 | 0.27 | 0.27 | 0.27 | 0.31 | 0.76 | 0.96 | 0.97 | 0.80 | 0.19 | 0.19 | 0.25 | 0.86 | 0.88 | 0.19 |
| PCA | 0.08 | 0.19 | 0.16 | 0.24 | 0.34 | 0.54 | 0.62 | 0.95 | 0.64 | 0.15 | 0.16 | 0.20 | 0.58 | 0.80 | 0.15 |
| CELLama | 0.04 | 0.09 | 0.08 | 0.09 | 0.23 | 0.46 | 0.55 | 0.94 | 0.28 | 0.14 | 0.14 | 0.11 | 0.51 | 0.61 | 0.14 |
| CellPLM | 0.07 | 0.13 | 0.13 | 0.13 | 0.23 | 0.42 | 0.52 | 0.94 | 0.22 | 0.15 | 0.15 | 0.14 | 0.47 | 0.58 | 0.15 |
| UCE | 0.05 | 0.11 | 0.10 | 0.11 | 0.23 | 0.42 | 0.53 | 0.94 | 0.25 | 0.15 | 0.15 | 0.12 | 0.47 | 0.60 | 0.15 |
| scGPT | 0.04 | 0.09 | 0.08 | 0.09 | 0.23 | 0.39 | 0.50 | 0.94 | 0.26 | 0.15 | 0.15 | 0.11 | 0.45 | 0.60 | 0.15 |
| scFoundation | 0.04 | 0.08 | 0.08 | 0.07 | 0.19 | 0.36 | 0.51 | 0.94 | 0.10 | 0.18 | 0.18 | 0.09 | 0.44 | 0.52 | 0.18 |
| Geneformer | 0.03 | 0.07 | 0.07 | 0.07 | 0.19 | 0.37 | 0.48 | 0.93 | 0.09 | 0.16 | 0.16 | 0.08 | 0.42 | 0.51 | 0.16 |
| SCimilarity | 0.03 | 0.07 | 0.08 | 0.07 | 0.17 | 0.29 | 0.53 | 0.93 | 0.07 | 0.15 | 0.14 | 0.08 | 0.41 | 0.50 | 0.14 |
| CellFM | 0.05 | 0.10 | 0.10 | 0.11 | 0.23 | 0.40 | 0.14 | 0.94 | 0.22 | 0.16 | 0.16 | 0.12 | 0.27 | 0.58 | 0.16 |
| GenePT-w | 0.01 | 0.03 | 0.03 | 0.03 | 0.15 | 0.38 | 0.40 | 0.93 | 0.01 | 0.18 | 0.18 | 0.05 | 0.39 | 0.47 | 0.18 |
| LangCell | 0.02 | 0.05 | 0.06 | 0.05 | 0.17 | 0.29 | 0.27 | 0.93 | 0.05 | 0.17 | 0.16 | 0.07 | 0.28 | 0.49 | 0.16 |
| scBERT | 0.01 | 0.03 | 0.03 | 0.03 | 0.17 | 0.24 | 0.20 | 0.93 | 0.04 | 0.18 | 0.18 | 0.06 | 0.22 | 0.49 | 0.18 |

**Figure S256: Spatial-domain clustering in MERFISH slice 0.04. a,** Predicted domains from 13 displayed methods and the reference annotation. **b,** Cluster-concordance, local-neighbourhood, spatial-coherence and spatial-marker scores. The four family means receive equal weight; rows are ordered by their aggregate score.

**a**

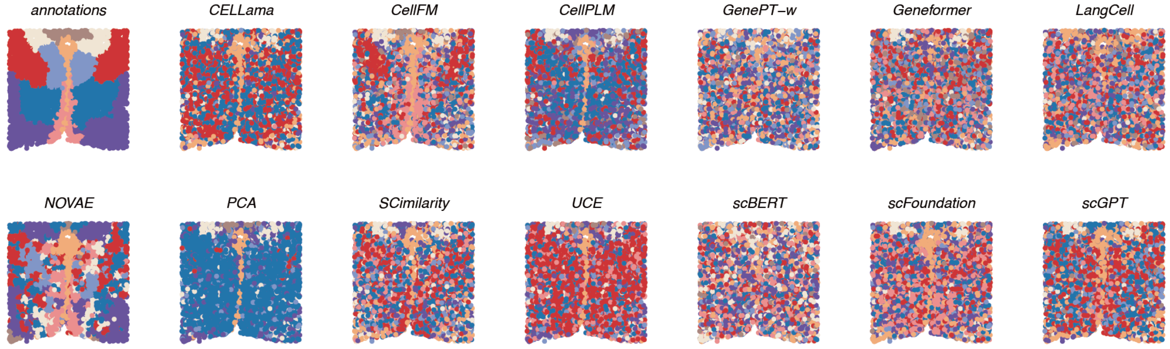

**b**

| method | Cluster concordance | | | | | Local Structure | | Spatial coherence | | Spatial marker | | Aggregated scores | | | |
|---|---|---|---|---|---|---|---|---|---|---|---|---|---|---|---|
| | ARI | NMI | HOM | COM | FMI | Acc@kNN | GC | 1-CHAOS | 1-PAS | Moran's I | 1-Geary's C | Cluster | Local | Coherence | Marker |
| NOVAE | 0.16 | 0.29 | 0.31 | 0.28 | 0.29 | 0.77 | 0.95 | 0.96 | 0.79 | 0.19 | 0.19 | 0.26 | 0.86 | 0.88 | 0.19 |
| PCA | 0.09 | 0.21 | 0.17 | 0.27 | 0.36 | 0.60 | 0.63 | 0.96 | 0.68 | 0.18 | 0.19 | 0.22 | 0.61 | 0.82 | 0.19 |
| CellPLM | 0.09 | 0.16 | 0.16 | 0.16 | 0.25 | 0.47 | 0.56 | 0.94 | 0.27 | 0.15 | 0.14 | 0.16 | 0.51 | 0.60 | 0.14 |
| CELLama | 0.05 | 0.10 | 0.09 | 0.11 | 0.26 | 0.50 | 0.53 | 0.95 | 0.32 | 0.14 | 0.13 | 0.12 | 0.52 | 0.63 | 0.14 |
| UCE | 0.06 | 0.14 | 0.14 | 0.15 | 0.25 | 0.46 | 0.50 | 0.94 | 0.25 | 0.17 | 0.16 | 0.15 | 0.48 | 0.60 | 0.16 |
| scGPT | 0.04 | 0.09 | 0.09 | 0.09 | 0.22 | 0.43 | 0.50 | 0.94 | 0.18 | 0.15 | 0.15 | 0.11 | 0.47 | 0.56 | 0.15 |
| scFoundation | 0.04 | 0.09 | 0.09 | 0.09 | 0.20 | 0.40 | 0.52 | 0.94 | 0.13 | 0.15 | 0.15 | 0.10 | 0.46 | 0.53 | 0.15 |
| CellFM | 0.11 | 0.15 | 0.16 | 0.15 | 0.25 | 0.45 | 0.15 | 0.94 | 0.25 | 0.17 | 0.16 | 0.16 | 0.30 | 0.59 | 0.17 |
| SCimilarity | 0.04 | 0.08 | 0.08 | 0.08 | 0.21 | 0.31 | 0.54 | 0.94 | 0.16 | 0.15 | 0.14 | 0.10 | 0.42 | 0.55 | 0.14 |
| Geneformer | 0.03 | 0.08 | 0.08 | 0.07 | 0.18 | 0.41 | 0.48 | 0.94 | 0.08 | 0.17 | 0.17 | 0.09 | 0.45 | 0.51 | 0.17 |
| GenePT-w | 0.01 | 0.03 | 0.03 | 0.03 | 0.17 | 0.43 | 0.43 | 0.93 | 0.03 | 0.19 | 0.19 | 0.06 | 0.43 | 0.48 | 0.19 |
| LangCell | 0.02 | 0.04 | 0.04 | 0.04 | 0.17 | 0.32 | 0.28 | 0.93 | 0.06 | 0.14 | 0.14 | 0.06 | 0.30 | 0.49 | 0.14 |
| scBERT | 0.01 | 0.02 | 0.03 | 0.02 | 0.16 | 0.25 | 0.17 | 0.93 | 0.03 | 0.17 | 0.17 | 0.05 | 0.21 | 0.48 | 0.17 |

**Figure S257: Spatial-domain clustering in MERFISH slice 0.09. a,** Predicted domains and reference annotation. **b,** Scores for 13 displayed methods. Display and scoring conventions are as in Fig. S256.

**a**

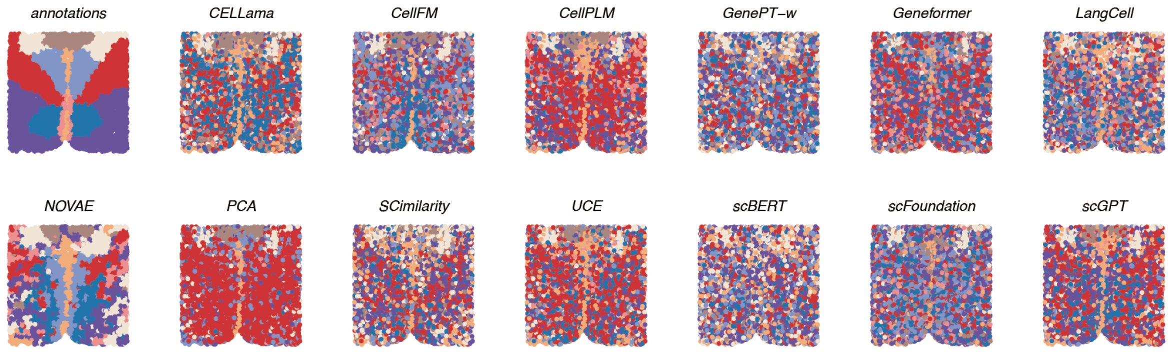

**b**

| method | Cluster concordance: ARI | NMI | HOM | COM | FMI | Local Structure: Acc@kNN | GC | Spatial coherence: 1-CHAOS | 1-PAS | Spatial marker: Moran's I | 1-Geary's C | Aggregated scores: Cluster | Local | Coherence | Marker |
|---|---|---|---|---|---|---|---|---|---|---|---|---|---|---|---|
| NOVAE | 0.18 | 0.32 | 0.35 | 0.30 | 0.31 | 0.79 | 0.95 | 0.97 | 0.79 | 0.19 | 0.19 | 0.29 | 0.87 | 0.88 | 0.19 |
| PCA | 0.11 | 0.22 | 0.19 | 0.28 | 0.38 | 0.57 | 0.61 | 0.95 | 0.69 | 0.17 | 0.17 | 0.23 | 0.59 | 0.82 | 0.17 |
| CellPLM | 0.08 | 0.16 | 0.16 | 0.16 | 0.26 | 0.46 | 0.53 | 0.94 | 0.37 | 0.17 | 0.17 | 0.17 | 0.50 | 0.65 | 0.17 |
| CELLama | 0.06 | 0.13 | 0.13 | 0.13 | 0.24 | 0.50 | 0.54 | 0.94 | 0.26 | 0.15 | 0.15 | 0.14 | 0.52 | 0.60 | 0.15 |
| scGPT | 0.07 | 0.13 | 0.13 | 0.13 | 0.25 | 0.45 | 0.53 | 0.94 | 0.25 | 0.18 | 0.18 | 0.14 | 0.49 | 0.59 | 0.18 |
| UCE | 0.06 | 0.12 | 0.12 | 0.12 | 0.24 | 0.46 | 0.50 | 0.94 | 0.24 | 0.17 | 0.17 | 0.13 | 0.48 | 0.59 | 0.17 |
| SCimilarity | 0.05 | 0.08 | 0.08 | 0.08 | 0.22 | 0.34 | 0.53 | 0.94 | 0.15 | 0.17 | 0.17 | 0.10 | 0.43 | 0.55 | 0.17 |
| Geneformer | 0.04 | 0.08 | 0.08 | 0.08 | 0.21 | 0.42 | 0.47 | 0.94 | 0.11 | 0.19 | 0.18 | 0.10 | 0.44 | 0.52 | 0.18 |
| scFoundation | 0.03 | 0.08 | 0.08 | 0.07 | 0.18 | 0.40 | 0.53 | 0.93 | 0.05 | 0.19 | 0.19 | 0.09 | 0.47 | 0.49 | 0.19 |
| GenePT-w | 0.02 | 0.04 | 0.04 | 0.04 | 0.18 | 0.42 | 0.50 | 0.94 | 0.04 | 0.19 | 0.19 | 0.06 | 0.46 | 0.49 | 0.19 |
| CellFM | 0.07 | 0.16 | 0.17 | 0.15 | 0.22 | 0.43 | 0.13 | 0.94 | 0.19 | 0.19 | 0.19 | 0.15 | 0.28 | 0.56 | 0.19 |
| LangCell | 0.02 | 0.05 | 0.05 | 0.05 | 0.18 | 0.33 | 0.26 | 0.94 | 0.06 | 0.20 | 0.19 | 0.07 | 0.30 | 0.50 | 0.20 |
| scBERT | 0.01 | 0.02 | 0.02 | 0.02 | 0.18 | 0.29 | 0.25 | 0.94 | 0.05 | 0.18 | 0.17 | 0.05 | 0.27 | 0.50 | 0.17 |

**Figure S258: Spatial-domain clustering in MERFISH slice 0.14. a,** Predicted domains and reference annotation. **b,** Scores for 13 displayed methods. Display and scoring conventions are as in Fig. S256.

**a**

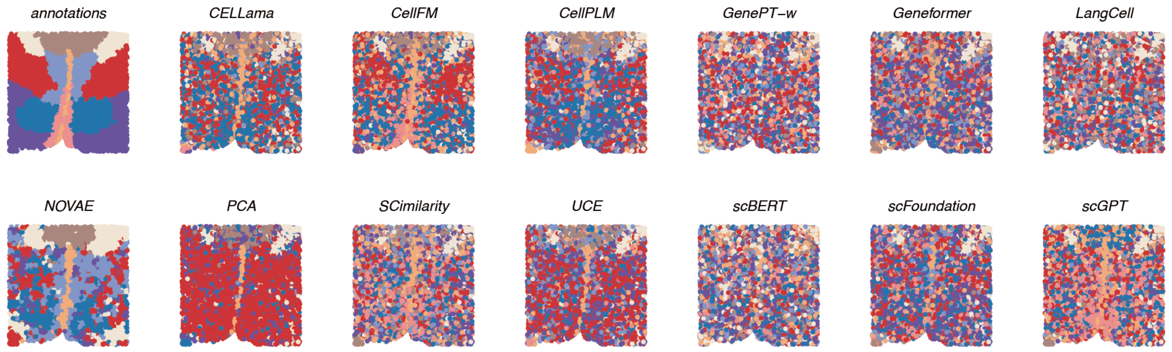

**b**

| | Cluster concordance | | | | | Local Structure | | Spatial coherence | | Spatial marker | | Aggregated scores | | | |
|---|---|---|---|---|---|---|---|---|---|---|---|---|---|---|---|
| method | ARI | NMI | HOM | COM | FMI | Acc@kNN | GC | 1-CHAOS | 1-PAS | Moran's I | 1-Geary's C | Cluster | Local | Coherence | Marker |
| NOVAE | 0.18 | 0.33 | 0.32 | 0.34 | 0.32 | 0.77 | 0.96 | 0.97 | 0.82 | 0.19 | 0.19 | 0.30 | 0.86 | 0.89 | 0.19 |
| PCA | 0.10 | 0.24 | 0.19 | 0.30 | 0.35 | 0.57 | 0.66 | 0.96 | 0.68 | 0.18 | 0.18 | 0.24 | 0.62 | 0.82 | 0.18 |
| CellPLM | 0.10 | 0.18 | 0.18 | 0.18 | 0.25 | 0.47 | 0.58 | 0.94 | 0.27 | 0.18 | 0.18 | 0.18 | 0.52 | 0.61 | 0.18 |
| CELLama | 0.08 | 0.14 | 0.14 | 0.15 | 0.25 | 0.49 | 0.53 | 0.94 | 0.29 | 0.16 | 0.16 | 0.15 | 0.51 | 0.62 | 0.16 |
| UCE | 0.07 | 0.13 | 0.13 | 0.13 | 0.22 | 0.45 | 0.56 | 0.94 | 0.19 | 0.16 | 0.16 | 0.13 | 0.50 | 0.56 | 0.16 |
| CellFM | 0.14 | 0.19 | 0.18 | 0.19 | 0.29 | 0.43 | 0.17 | 0.95 | 0.34 | 0.18 | 0.18 | 0.20 | 0.30 | 0.64 | 0.18 |
| scGPT | 0.04 | 0.10 | 0.10 | 0.10 | 0.20 | 0.45 | 0.53 | 0.94 | 0.18 | 0.16 | 0.16 | 0.11 | 0.49 | 0.56 | 0.16 |
| scFoundation | 0.04 | 0.08 | 0.08 | 0.08 | 0.20 | 0.40 | 0.60 | 0.94 | 0.10 | 0.18 | 0.18 | 0.10 | 0.50 | 0.52 | 0.18 |
| SCimilarity | 0.04 | 0.09 | 0.09 | 0.09 | 0.18 | 0.32 | 0.58 | 0.94 | 0.11 | 0.18 | 0.18 | 0.10 | 0.45 | 0.52 | 0.18 |
| Geneformer | 0.03 | 0.08 | 0.08 | 0.08 | 0.18 | 0.40 | 0.48 | 0.94 | 0.10 | 0.18 | 0.18 | 0.09 | 0.44 | 0.52 | 0.18 |
| GenePT-w | 0.01 | 0.03 | 0.03 | 0.03 | 0.17 | 0.41 | 0.55 | 0.94 | 0.03 | 0.17 | 0.17 | 0.05 | 0.48 | 0.48 | 0.17 |
| LangCell | 0.02 | 0.04 | 0.04 | 0.04 | 0.17 | 0.31 | 0.32 | 0.94 | 0.05 | 0.18 | 0.18 | 0.06 | 0.31 | 0.49 | 0.18 |
| scBERT | 0.01 | 0.03 | 0.03 | 0.03 | 0.15 | 0.24 | 0.19 | 0.93 | 0.02 | 0.18 | 0.18 | 0.05 | 0.21 | 0.48 | 0.18 |

**Figure S259: Spatial-domain clustering in MERFISH slice 0.19. a,** Predicted domains and reference annotation. **b,** Scores for 13 displayed methods. Display and scoring conventions are as in Fig. S256.

**a**

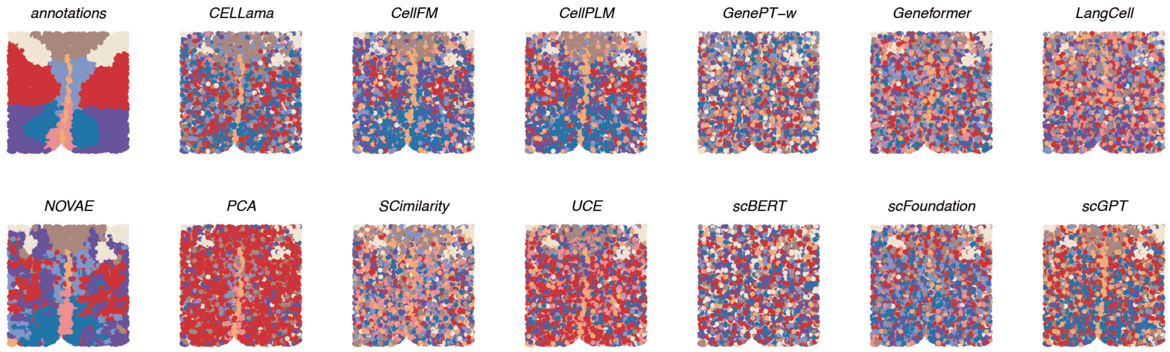

**b**

| | Cluster concordance | | | | | Local Structure | | Spatial coherence | | Spatial marker | | Aggregated scores | | | |
|---|---|---|---|---|---|---|---|---|---|---|---|---|---|---|---|
| **method** | ARI | NMI | HOM | COM | FMI | Acc@kNN | GC | 1-CHAOS | 1-PAS | Moran's I | 1-Geary's C | Cluster | Local | Coherence | Marker |
| **NOVAE** | 0.24 | 0.39 | 0.41 | 0.38 | 0.36 | 0.76 | 0.96 | 0.97 | 0.80 | 0.18 | 0.17 | 0.36 | 0.86 | 0.88 | 0.18 |
| **PCA** | 0.08 | 0.18 | 0.16 | 0.21 | 0.31 | 0.56 | 0.56 | 0.95 | 0.55 | 0.15 | 0.14 | 0.19 | 0.56 | 0.75 | 0.14 |
| **CellPLM** | 0.11 | 0.19 | 0.19 | 0.19 | 0.25 | 0.49 | 0.57 | 0.94 | 0.28 | 0.16 | 0.16 | 0.19 | 0.53 | 0.61 | 0.16 |
| **CELLama** | 0.08 | 0.17 | 0.17 | 0.17 | 0.24 | 0.51 | 0.51 | 0.94 | 0.22 | 0.16 | 0.14 | 0.17 | 0.51 | 0.58 | 0.15 |
| **UCE** | 0.06 | 0.12 | 0.12 | 0.12 | 0.23 | 0.46 | 0.55 | 0.94 | 0.19 | 0.15 | 0.14 | 0.13 | 0.51 | 0.57 | 0.15 |
| **scGPT** | 0.04 | 0.11 | 0.11 | 0.11 | 0.21 | 0.45 | 0.53 | 0.94 | 0.20 | 0.13 | 0.13 | 0.12 | 0.49 | 0.57 | 0.13 |
| **scFoundation** | 0.03 | 0.08 | 0.08 | 0.08 | 0.18 | 0.40 | 0.56 | 0.94 | 0.07 | 0.16 | 0.15 | 0.09 | 0.48 | 0.50 | 0.15 |
| **CellFM** | 0.08 | 0.16 | 0.16 | 0.16 | 0.23 | 0.43 | 0.17 | 0.94 | 0.26 | 0.15 | 0.14 | 0.16 | 0.30 | 0.60 | 0.15 |
| **SCimilarity** | 0.03 | 0.08 | 0.08 | 0.08 | 0.17 | 0.33 | 0.56 | 0.94 | 0.09 | 0.16 | 0.16 | 0.09 | 0.45 | 0.51 | 0.16 |
| **Geneformer** | 0.03 | 0.08 | 0.08 | 0.07 | 0.17 | 0.42 | 0.49 | 0.93 | 0.07 | 0.14 | 0.14 | 0.09 | 0.45 | 0.50 | 0.14 |
| **GenePT-w** | 0.01 | 0.03 | 0.03 | 0.03 | 0.16 | 0.43 | 0.51 | 0.93 | 0.03 | 0.16 | 0.15 | 0.05 | 0.47 | 0.48 | 0.15 |
| **LangCell** | 0.02 | 0.05 | 0.05 | 0.05 | 0.17 | 0.34 | 0.30 | 0.93 | 0.05 | 0.17 | 0.17 | 0.07 | 0.32 | 0.49 | 0.17 |
| **scBERT** | 0.01 | 0.02 | 0.02 | 0.02 | 0.17 | 0.26 | 0.19 | 0.93 | 0.04 | 0.15 | 0.14 | 0.05 | 0.22 | 0.48 | 0.15 |

**Figure S260: Spatial-domain clustering in MERFISH slice 0.24. a,** Predicted domains and reference annotation. **b,** Scores for 13 displayed methods. Display and scoring conventions are as in Fig. S256.

##### *2.10.4 MERFISH cell-type clustering*

The separate MERFISH cell-type endpoint is shown for all five z-sections in Figs. S261–S265.

**a**

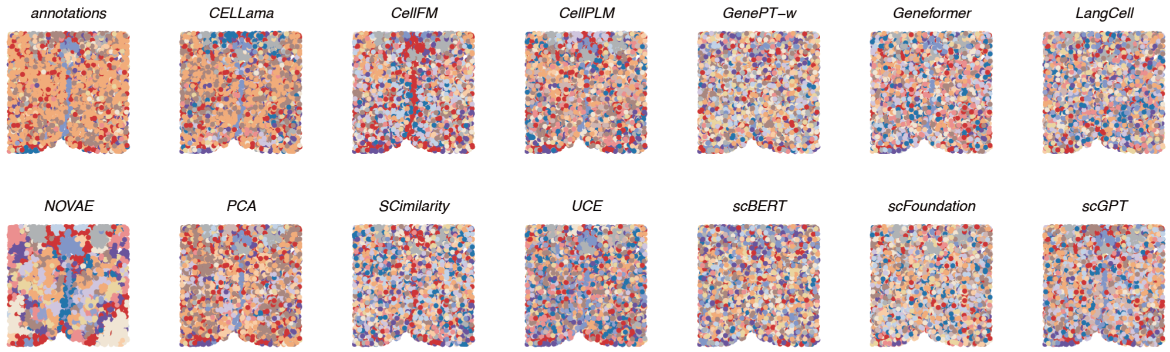

**b**

| method | Cluster concordance | | | | | Local Structure | | Spatial coherence | | Spatial marker | | Aggregated scores | | | |
|---|---|---|---|---|---|---|---|---|---|---|---|---|---|---|---|
| | ARI | NMI | HOM | COM | FMI | Acc@kNN | GC | 1-CHAOS | 1-PAS | Moran's I | 1-Geary's C | Cluster | Local | Coherence | Marker |
| CELLama | 0.40 | 0.60 | 0.65 | 0.55 | 0.52 | 0.85 | 0.70 | 0.93 | 0.20 | 0.15 | 0.15 | 0.54 | 0.77 | 0.57 | 0.15 |
| PCA | 0.31 | 0.59 | 0.66 | 0.53 | 0.45 | 0.86 | 0.71 | 0.93 | 0.20 | 0.13 | 0.13 | 0.51 | 0.79 | 0.57 | 0.13 |
| CellPLM | 0.25 | 0.48 | 0.57 | 0.42 | 0.39 | 0.80 | 0.65 | 0.92 | 0.11 | 0.15 | 0.15 | 0.42 | 0.72 | 0.52 | 0.15 |
| scGPT | 0.17 | 0.43 | 0.51 | 0.37 | 0.32 | 0.77 | 0.64 | 0.92 | 0.09 | 0.15 | 0.15 | 0.36 | 0.70 | 0.50 | 0.15 |
| UCE | 0.18 | 0.43 | 0.51 | 0.37 | 0.32 | 0.78 | 0.63 | 0.92 | 0.07 | 0.13 | 0.13 | 0.36 | 0.71 | 0.50 | 0.13 |
| Geneformer | 0.17 | 0.38 | 0.46 | 0.33 | 0.31 | 0.73 | 0.55 | 0.92 | 0.04 | 0.15 | 0.15 | 0.33 | 0.64 | 0.48 | 0.15 |
| scFoundation | 0.13 | 0.31 | 0.37 | 0.26 | 0.27 | 0.69 | 0.56 | 0.92 | 0.04 | 0.15 | 0.16 | 0.27 | 0.62 | 0.48 | 0.16 |
| SCimilarity | 0.14 | 0.31 | 0.37 | 0.27 | 0.28 | 0.64 | 0.58 | 0.92 | 0.04 | 0.15 | 0.14 | 0.27 | 0.61 | 0.48 | 0.14 |
| NOVAE | 0.05 | 0.12 | 0.14 | 0.10 | 0.19 | 0.42 | 0.33 | 0.96 | 0.70 | 0.16 | 0.16 | 0.12 | 0.37 | 0.83 | 0.16 |
| GenePT-w | 0.06 | 0.15 | 0.18 | 0.13 | 0.20 | 0.72 | 0.50 | 0.91 | 0.01 | 0.18 | 0.18 | 0.15 | 0.61 | 0.46 | 0.18 |
| CellFM | 0.15 | 0.36 | 0.42 | 0.32 | 0.30 | 0.64 | 0.12 | 0.92 | 0.12 | 0.14 | 0.14 | 0.31 | 0.38 | 0.52 | 0.14 |
| LangCell | 0.08 | 0.21 | 0.26 | 0.18 | 0.22 | 0.58 | 0.37 | 0.91 | 0.02 | 0.16 | 0.16 | 0.19 | 0.48 | 0.47 | 0.16 |
| scBERT | 0.04 | 0.10 | 0.12 | 0.09 | 0.18 | 0.39 | 0.20 | 0.91 | 0.01 | 0.16 | 0.16 | 0.11 | 0.29 | 0.46 | 0.16 |

**Figure S261: Cell-type clustering in MERFISH slice 0.04. a,** Predicted cell-type clusters from 13 displayed methods and the reference annotation. **b,** Cluster-concordance, local-neighbourhood, spatial-coherence and spatial-marker scores. The four family means receive equal weight; rows are ordered by their aggregate score.

**a**

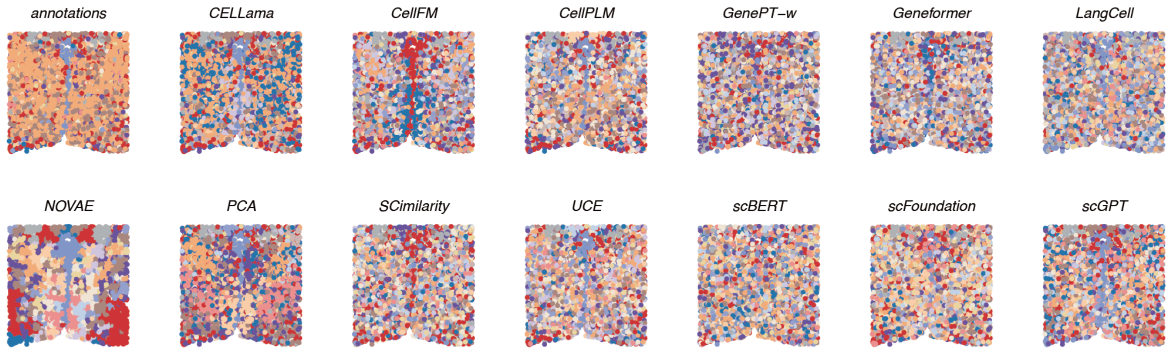

**b**

| | Cluster concordance | | | | | Local Structure | | Spatial coherence | | Spatial marker | | Aggregated scores | | | |
|---|---|---|---|---|---|---|---|---|---|---|---|---|---|---|---|
| method | ARI | NMI | HOM | COM | FMI | Acc@kNN | GC | 1-CHAOS | 1-PAS | Moran's I | 1-Geary's C | Cluster | Local | Coherence | Marker |
| PCA | 0.26 | 0.56 | 0.68 | 0.47 | 0.41 | 0.89 | 0.66 | 0.93 | 0.29 | 0.13 | 0.13 | 0.48 | 0.77 | 0.61 | 0.13 |
| CELLama | 0.35 | 0.58 | 0.67 | 0.52 | 0.49 | 0.87 | 0.70 | 0.93 | 0.17 | 0.13 | 0.12 | 0.52 | 0.78 | 0.55 | 0.12 |
| CellPLM | 0.24 | 0.48 | 0.60 | 0.40 | 0.40 | 0.83 | 0.64 | 0.92 | 0.12 | 0.14 | 0.14 | 0.42 | 0.74 | 0.52 | 0.14 |
| scGPT | 0.17 | 0.43 | 0.53 | 0.37 | 0.32 | 0.79 | 0.59 | 0.92 | 0.09 | 0.16 | 0.16 | 0.36 | 0.69 | 0.51 | 0.16 |
| UCE | 0.16 | 0.41 | 0.51 | 0.35 | 0.31 | 0.79 | 0.58 | 0.92 | 0.07 | 0.14 | 0.13 | 0.35 | 0.68 | 0.49 | 0.13 |
| NOVAE | 0.03 | 0.12 | 0.15 | 0.10 | 0.19 | 0.47 | 0.30 | 0.96 | 0.72 | 0.17 | 0.17 | 0.12 | 0.38 | 0.84 | 0.17 |
| Geneformer | 0.12 | 0.34 | 0.43 | 0.28 | 0.26 | 0.73 | 0.51 | 0.91 | 0.02 | 0.14 | 0.14 | 0.29 | 0.62 | 0.47 | 0.14 |
| scFoundation | 0.11 | 0.31 | 0.39 | 0.26 | 0.26 | 0.71 | 0.50 | 0.91 | 0.03 | 0.14 | 0.14 | 0.27 | 0.60 | 0.47 | 0.14 |
| SCimilarity | 0.09 | 0.27 | 0.35 | 0.23 | 0.24 | 0.66 | 0.54 | 0.91 | 0.04 | 0.14 | 0.14 | 0.24 | 0.60 | 0.48 | 0.14 |
| GenePT-w | 0.08 | 0.21 | 0.27 | 0.18 | 0.22 | 0.75 | 0.51 | 0.91 | 0.01 | 0.15 | 0.14 | 0.19 | 0.63 | 0.46 | 0.14 |
| CellFM | 0.12 | 0.31 | 0.39 | 0.26 | 0.27 | 0.66 | 0.14 | 0.91 | 0.14 | 0.14 | 0.13 | 0.27 | 0.40 | 0.53 | 0.14 |
| LangCell | 0.07 | 0.18 | 0.23 | 0.15 | 0.21 | 0.60 | 0.35 | 0.91 | 0.03 | 0.12 | 0.12 | 0.17 | 0.48 | 0.47 | 0.12 |
| scBERT | 0.04 | 0.11 | 0.14 | 0.09 | 0.19 | 0.42 | 0.20 | 0.91 | 0.00 | 0.16 | 0.15 | 0.11 | 0.31 | 0.46 | 0.15 |

**Figure S262: Cell-type clustering in MERFISH slice 0.09. a,** Predicted cell-type clusters and reference annotation. **b,** Scores for 13 displayed methods. Display and scoring conventions are as in Fig. S261.

**a**

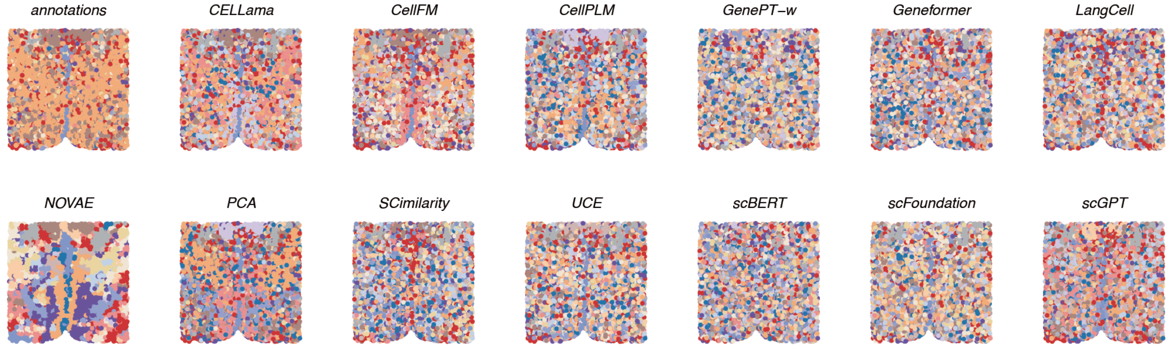

**b**

| method | Cluster concordance | | | | | Local Structure | | Spatial coherence | | Spatial marker | | Aggregated scores | | | |
|---|---|---|---|---|---|---|---|---|---|---|---|---|---|---|---|
| | ARI | NMI | HOM | COM | FMI | Acc@kNN | GC | 1-CHAOS | 1-PAS | Moran's I | 1-Geary's C | Cluster | Local | Coherence | Marker |
| PCA | 0.31 | 0.58 | 0.69 | 0.50 | 0.46 | 0.88 | 0.65 | 0.93 | 0.28 | 0.16 | 0.16 | 0.51 | 0.77 | 0.61 | 0.16 |
| CELLama | 0.29 | 0.56 | 0.69 | 0.48 | 0.45 | 0.87 | 0.69 | 0.93 | 0.13 | 0.16 | 0.16 | 0.49 | 0.78 | 0.53 | 0.16 |
| CellPLM | 0.22 | 0.47 | 0.59 | 0.38 | 0.38 | 0.83 | 0.64 | 0.92 | 0.12 | 0.18 | 0.17 | 0.41 | 0.73 | 0.52 | 0.17 |
| scGPT | 0.17 | 0.43 | 0.54 | 0.36 | 0.32 | 0.79 | 0.66 | 0.92 | 0.10 | 0.18 | 0.18 | 0.36 | 0.72 | 0.51 | 0.18 |
| UCE | 0.16 | 0.42 | 0.53 | 0.34 | 0.32 | 0.80 | 0.61 | 0.92 | 0.08 | 0.17 | 0.17 | 0.35 | 0.70 | 0.50 | 0.17 |
| Geneformer | 0.12 | 0.33 | 0.42 | 0.28 | 0.27 | 0.74 | 0.56 | 0.92 | 0.04 | 0.18 | 0.18 | 0.28 | 0.65 | 0.48 | 0.18 |
| NOVAE | 0.04 | 0.12 | 0.15 | 0.09 | 0.19 | 0.49 | 0.32 | 0.96 | 0.71 | 0.18 | 0.18 | 0.12 | 0.40 | 0.83 | 0.18 |
| scFoundation | 0.11 | 0.30 | 0.38 | 0.25 | 0.26 | 0.70 | 0.53 | 0.92 | 0.04 | 0.18 | 0.18 | 0.26 | 0.62 | 0.48 | 0.18 |
| GenePT-w | 0.06 | 0.18 | 0.23 | 0.15 | 0.21 | 0.75 | 0.55 | 0.91 | 0.01 | 0.18 | 0.18 | 0.16 | 0.65 | 0.46 | 0.18 |
| SCimilarity | 0.08 | 0.24 | 0.31 | 0.20 | 0.23 | 0.66 | 0.58 | 0.91 | 0.03 | 0.15 | 0.15 | 0.21 | 0.62 | 0.47 | 0.15 |
| CellFM | 0.16 | 0.36 | 0.43 | 0.30 | 0.31 | 0.66 | 0.13 | 0.93 | 0.14 | 0.16 | 0.16 | 0.31 | 0.40 | 0.54 | 0.16 |
| LangCell | 0.05 | 0.17 | 0.21 | 0.14 | 0.20 | 0.61 | 0.33 | 0.91 | 0.01 | 0.18 | 0.17 | 0.16 | 0.47 | 0.46 | 0.17 |
| scBERT | 0.03 | 0.10 | 0.13 | 0.09 | 0.17 | 0.43 | 0.23 | 0.91 | 0.00 | 0.19 | 0.18 | 0.10 | 0.33 | 0.46 | 0.18 |

**Figure S263: Cell-type clustering in MERFISH slice 0.14. a,** Predicted cell-type clusters and reference annotation. **b,** Scores for 13 displayed methods. Display and scoring conventions are as in Fig. S261.

**a**

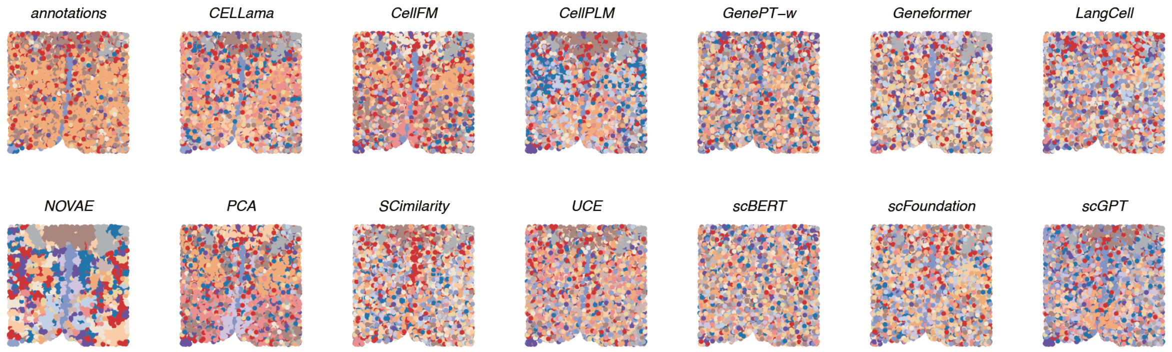

**b**

| method | Cluster concordance | | | | | Local Structure | | Spatial coherence | | Spatial marker | | Aggregated scores | | | |
|---|---|---|---|---|---|---|---|---|---|---|---|---|---|---|---|
| | ARI | NMI | HOM | COM | FMI | Acc@kNN | GC | 1-CHAOS | 1-PAS | Moran's I | 1-Geary's C | Cluster | Local | Coherence | Marker |
| PCA | 0.34 | 0.60 | 0.71 | 0.52 | 0.48 | 0.89 | 0.67 | 0.94 | 0.31 | 0.16 | 0.16 | 0.53 | 0.78 | 0.62 | 0.16 |
| CELLama | 0.34 | 0.60 | 0.72 | 0.52 | 0.49 | 0.88 | 0.71 | 0.93 | 0.18 | 0.16 | 0.16 | 0.53 | 0.80 | 0.55 | 0.16 |
| CellPLM | 0.23 | 0.47 | 0.59 | 0.39 | 0.39 | 0.83 | 0.64 | 0.92 | 0.15 | 0.16 | 0.16 | 0.42 | 0.73 | 0.54 | 0.16 |
| scGPT | 0.15 | 0.43 | 0.53 | 0.36 | 0.30 | 0.79 | 0.60 | 0.92 | 0.12 | 0.17 | 0.17 | 0.36 | 0.70 | 0.52 | 0.17 |
| UCE | 0.18 | 0.43 | 0.52 | 0.36 | 0.33 | 0.81 | 0.58 | 0.92 | 0.06 | 0.16 | 0.16 | 0.36 | 0.70 | 0.49 | 0.16 |
| Geneformer | 0.12 | 0.34 | 0.43 | 0.28 | 0.26 | 0.74 | 0.52 | 0.92 | 0.05 | 0.18 | 0.18 | 0.29 | 0.63 | 0.48 | 0.18 |
| SCimilarity | 0.12 | 0.28 | 0.35 | 0.24 | 0.27 | 0.67 | 0.56 | 0.92 | 0.06 | 0.18 | 0.18 | 0.25 | 0.62 | 0.49 | 0.18 |
| NOVAE | 0.04 | 0.13 | 0.17 | 0.11 | 0.18 | 0.47 | 0.33 | 0.96 | 0.68 | 0.18 | 0.18 | 0.12 | 0.40 | 0.82 | 0.18 |
| scFoundation | 0.12 | 0.29 | 0.36 | 0.24 | 0.27 | 0.71 | 0.51 | 0.92 | 0.04 | 0.16 | 0.16 | 0.26 | 0.61 | 0.48 | 0.16 |
| CellFM | 0.20 | 0.38 | 0.45 | 0.33 | 0.36 | 0.67 | 0.13 | 0.93 | 0.19 | 0.17 | 0.17 | 0.35 | 0.40 | 0.56 | 0.17 |
| GenePT-w | 0.08 | 0.20 | 0.25 | 0.17 | 0.23 | 0.75 | 0.49 | 0.91 | 0.01 | 0.16 | 0.16 | 0.18 | 0.62 | 0.46 | 0.16 |
| LangCell | 0.06 | 0.17 | 0.21 | 0.14 | 0.22 | 0.60 | 0.33 | 0.92 | 0.02 | 0.16 | 0.16 | 0.16 | 0.47 | 0.47 | 0.16 |
| scBERT | 0.04 | 0.11 | 0.14 | 0.09 | 0.19 | 0.42 | 0.18 | 0.91 | 0.00 | 0.18 | 0.18 | 0.11 | 0.30 | 0.46 | 0.18 |

**Figure S264: Cell-type clustering in MERFISH slice 0.19. a,** Predicted cell-type clusters and reference annotation. **b,** Scores for 13 displayed methods. Display and scoring conventions are as in Fig. S261.

**a**

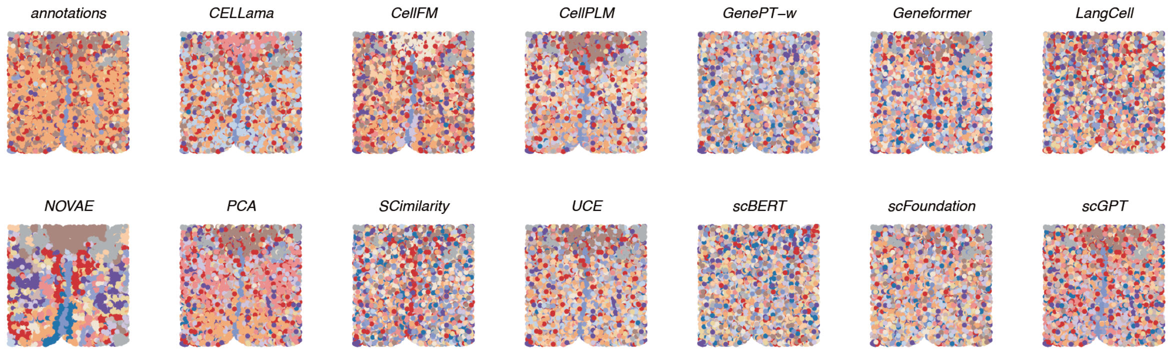

**b**

| method | ARI | NMI | HOM | COM | FMI | Acc@kNN | GC | 1-CHAOS | 1-PAS | Moran's I | 1-Geary's C | Cluster | Local | Coherence | Marker |
|---|---|---|---|---|---|---|---|---|---|---|---|---|---|---|---|
| | Cluster concordance | | | | | Local Structure | | Spatial coherence | | Spatial marker | | Aggregated scores | | | |
| PCA | 0.37 | 0.61 | 0.71 | 0.54 | 0.50 | 0.89 | 0.67 | 0.93 | 0.28 | 0.14 | 0.13 | 0.55 | 0.78 | 0.61 | 0.14 |
| CELLama | 0.41 | 0.63 | 0.72 | 0.56 | 0.53 | 0.88 | 0.70 | 0.93 | 0.17 | 0.14 | 0.13 | 0.57 | 0.79 | 0.55 | 0.13 |
| CellPLM | 0.28 | 0.51 | 0.62 | 0.43 | 0.42 | 0.83 | 0.66 | 0.92 | 0.13 | 0.16 | 0.16 | 0.45 | 0.75 | 0.53 | 0.16 |
| scGPT | 0.22 | 0.49 | 0.59 | 0.42 | 0.36 | 0.80 | 0.63 | 0.92 | 0.10 | 0.16 | 0.15 | 0.42 | 0.71 | 0.51 | 0.15 |
| UCE | 0.23 | 0.48 | 0.58 | 0.41 | 0.37 | 0.81 | 0.62 | 0.92 | 0.09 | 0.15 | 0.15 | 0.41 | 0.72 | 0.50 | 0.15 |
| Geneformer | 0.15 | 0.39 | 0.47 | 0.33 | 0.29 | 0.75 | 0.54 | 0.91 | 0.04 | 0.16 | 0.16 | 0.33 | 0.64 | 0.48 | 0.16 |
| scFoundation | 0.12 | 0.33 | 0.41 | 0.28 | 0.26 | 0.72 | 0.54 | 0.91 | 0.04 | 0.16 | 0.16 | 0.28 | 0.63 | 0.48 | 0.16 |
| SCimilarity | 0.14 | 0.33 | 0.40 | 0.28 | 0.28 | 0.68 | 0.57 | 0.92 | 0.06 | 0.15 | 0.15 | 0.29 | 0.63 | 0.49 | 0.15 |
| CellFM | 0.24 | 0.43 | 0.49 | 0.38 | 0.38 | 0.67 | 0.15 | 0.93 | 0.18 | 0.15 | 0.14 | 0.38 | 0.41 | 0.55 | 0.15 |
| NOVAE | 0.04 | 0.13 | 0.15 | 0.11 | 0.18 | 0.42 | 0.30 | 0.96 | 0.67 | 0.17 | 0.16 | 0.12 | 0.36 | 0.82 | 0.16 |
| GenePT-w | 0.09 | 0.21 | 0.26 | 0.18 | 0.23 | 0.77 | 0.50 | 0.91 | 0.01 | 0.16 | 0.16 | 0.19 | 0.64 | 0.46 | 0.16 |
| LangCell | 0.08 | 0.20 | 0.24 | 0.17 | 0.22 | 0.61 | 0.35 | 0.92 | 0.02 | 0.17 | 0.16 | 0.18 | 0.48 | 0.47 | 0.16 |
| scBERT | 0.04 | 0.11 | 0.13 | 0.09 | 0.17 | 0.41 | 0.21 | 0.91 | 0.01 | 0.17 | 0.17 | 0.11 | 0.31 | 0.46 | 0.17 |

**Figure S265: Cell-type clustering in MERFISH slice 0.24. a,** Predicted cell-type clusters and reference annotation. **b,** Scores for 13 displayed methods. Display and scoring conventions are as in Fig. S261.

##### 2.11 Paired evaluation of DRIFT spatial adaptation

The effect of the common DRIFT operator varied across datasets and methods (Table S11). Averaged over the common 12-method panel, mean ΔARI was +0.031 for SpatialLIBD, −0.009 for HER2ST and +0.053 for the MERFISH spatial-domain endpoint. Seven of 12 methods had a positive mean change on SpatialLIBD, five of 12 on HER2ST and 11 of 12 on MERFISH. scGPT improved in 0/12 SpatialLIBD sections and 2/8 HER2ST sections, but in all five MERFISH sections (mean ΔARI +0.068).

The response was also method dependent. CellPLM increased more than PCA on SpatialLIBD (+0.169 versus +0.119), and CellFM increased more than PCA on MERFISH (+0.151 versus +0.117). Conversely, PCA itself improved on SpatialLIBD and MERFISH. Coordinate-based diffusion therefore benefited both selected scFM representations and PCA, with effects that differed across datasets and methods.

#### Supplementary Tables

**Table S11:** Effect of DRIFT spatial adaptation on spatial-domain ARI. The common panel contains 11 eligible scFMs and HVGs+PCA; NOVAE, pretrained and de novo scVI, scCello and scPRINT are outside this paired estimand. DRIFT used $k_{\text{graph}} = 3$ and $t = 5$. Clustering used k-means with $K_{\text{clusters}}$ fixed to the number of annotated domains. $\Delta$ is adapted minus unadapted ARI. For each method, the final column reports sections with positive $\Delta$ARI; in each dataset mean row it instead reports methods with positive mean $\Delta$ARI.

| Method | Unadapted | Adapted | ΔARI | Positive units |
|---|---|---|---|---|
| *spatialLIBD* (spot-level) | | | | |
| scGPT | 0.187 | 0.123 | −0.064 | 0/12 |
| scFoundation | 0.167 | 0.150 | −0.016 | 2/12 |
| Geneformer | 0.140 | 0.113 | −0.028 | 4/12 |
| CellPLM | 0.179 | 0.348 | +0.169 | 12/12 |
| CellFM | 0.150 | 0.181 | +0.031 | 9/12 |
| CELLama | 0.088 | 0.151 | +0.063 | 11/12 |
| SCimilarity | 0.124 | 0.169 | +0.045 | 11/12 |
| LangCell | 0.153 | 0.207 | +0.054 | 9/12 |
| UCE | 0.160 | 0.238 | +0.078 | 11/12 |
| GenePT-w | 0.083 | 0.080 | −0.003 | 4/12 |
| scBERT | 0.107 | 0.033 | −0.074 | 1/12 |
| PCA | 0.199 | 0.318 | +0.119 | 10/12 |
| *mean* | | | +0.031 | 7/12 methods |
| *HER2ST* (spot-level) | | | | |
| scGPT | 0.130 | 0.115 | −0.015 | 2/8 |
| scFoundation | 0.136 | 0.113 | −0.023 | 2/8 |
| Geneformer | 0.152 | 0.091 | −0.061 | 2/8 |
| CellPLM | 0.131 | 0.173 | +0.042 | 4/8 |
| CellFM | 0.072 | 0.111 | +0.038 | 5/8 |
| CELLama | 0.114 | 0.132 | +0.018 | 3/8 |
| SCimilarity | 0.102 | 0.144 | +0.041 | 6/8 |
| LangCell | 0.118 | 0.135 | +0.017 | 4/8 |
| UCE | 0.159 | 0.152 | −0.008 | 3/8 |
| GenePT-w | 0.124 | 0.083 | −0.041 | 2/8 |
| scBERT | 0.129 | 0.019 | −0.110 | 0/8 |
| PCA | 0.199 | 0.198 | −0.001 | 4/8 |
| *mean* | | | −0.009 | 5/12 methods |
| *MERFISH* (single-cell) | | | | |
| scGPT | 0.041 | 0.109 | +0.068 | 5/5 |
| scFoundation | 0.032 | 0.070 | +0.038 | 5/5 |
| Geneformer | 0.026 | 0.058 | +0.032 | 5/5 |
| CellPLM | 0.094 | 0.153 | +0.059 | 5/5 |
| CellFM | 0.067 | 0.218 | +0.151 | 5/5 |
| CELLama | 0.054 | 0.135 | +0.082 | 5/5 |
| SCimilarity | 0.037 | 0.051 | +0.014 | 4/5 |
| LangCell | 0.024 | 0.017 | −0.007 | 1/5 |
| UCE | 0.056 | 0.091 | +0.035 | 5/5 |
| GenePT-w | 0.011 | 0.032 | +0.021 | 5/5 |
| scBERT | 0.005 | 0.029 | +0.024 | 5/5 |
| PCA | 0.134 | 0.251 | +0.117 | 5/5 |
| *mean* | | | +0.053 | 11/12 methods |

**Table S12: Datasets analysed in this study.** The 94 transcriptomic datasets comprise tissue-specific atlas datasets, developmental systems, spatial sections, regulatory-network datasets and perturbation datasets. Source–target directions, transcription factors and repeated label draws are analysis units derived from these datasets rather than additional datasets.

| Collection | Dataset units | Analysed scale | Principal analyses |
|---|---|---|---|
| Tabula Sapiens v1 | 24 tissue datasets | 444,116 cells | Cross-release robustness and cross-dataset transfer |
| Tabula Sapiens v2 | 26 tissue datasets | 548,977 cells | Cell-representation benchmarking, representation geometry and transfer |
| Developmental trajectories | 9 datasets | 4,751 cells; three linear, three bifurcating and three tree-structured systems | Trajectory recovery with Slingshot, PAGA, DPT and Monocle 3 |
| Spatial transcriptomics | 25 sections | 79,479 observations: 12 SpatialLIBD, 8 HER2ST and 5 MERFISH sections | Spatial-domain recovery, MERFISH cell-type clustering and DRIFT adaptation |
| BEELINE | 7 datasets from 5 studies | Seven static and four count-traceable contextual evaluations | Regulatory-edge prioritization |
| Perturbation data | 3 datasets | Norman CRISPRa and Replogle K562 and RPE1 CRISPRi | Perturbation-response-gene retrieval and cell-level double-perturbation prediction |

**Table S13: Metric families for cell-representation and spatial-domain evaluation.** All reported metrics were oriented so that higher values indicate better performance.

| Family | Metrics | Biological or technical quantity |
|---|---|---|
| Cluster concordance | ARI, NMI, homogeneity, completeness and FMI | Agreement between unsupervised clusters and annotated biological groups |
| Continuous separation | Cell-type ASW | Separation of annotated cell types relative to within-type dispersion |
| Local neighbourhood structure | Acc@kNN, cLISI and graph connectivity | Local recoverability and purity of biological labels, and connectivity of cells sharing a label |
| Batch mixing | BRAS, kBET and CiLISI | Mixing of batches within biologically matched populations |
| Spatial coherence | $1-$CHAOS and $1-$PAS | Compactness and local spatial continuity of predicted domains |
| Spatial-marker autocorrelation | Moran's I and $1-$Geary's C | Spatial autocorrelation of marker-gene expression within predicted domains |

**Table S14: Trajectory-inference metric families.** The 12 retained `dyneval` effect-size metrics were oriented so that higher values indicate greater agreement with the reference trajectory.

| Family | Retained metrics | Trajectory property |
|---|---|---|
| Cell-position distance | Geodesic-distance correlation | Agreement of global cell–cell positions along the inferred and reference trajectories |
| Reference-position recoverability | Linear-model and random-forest normalized MSE and $R^2$ | Recovery of reference progression from inferred cell positions |
| Topology | Edge-flip similarity, Hamming–Ipsen–Mikhailov similarity and graph isomorphism | Agreement of milestone-graph structure |
| Feature relevance | Correlation and weighted correlation of trajectory-associated feature importance | Recovery of genes varying along the reference trajectory |
| Branch assignment | Branch F1 and milestone F1 | Agreement of cell allocation to branches and milestones |

DPT yields a directed linear trajectory; topology metrics and branch F1 were therefore unavailable for this backend.

**Table S15: Primary endpoints for supervised transfer and gene-level evaluation.**

| Analysis | Primary endpoint | Scientific quantity | Complementary endpoints |
|---|---|---|---|
| Closed-set transfer | Macro-F1 | Accuracy across direction-specific shared cell types without weighting abundant types more strongly | Accuracy, balanced accuracy, macro precision, macro recall, AUROC and top-3 accuracy |
| Label efficiency | Smallest tested source-label budget reaching 90% of the method's $N$ = 500 macro-F1 | Amount of labelled reference data required to recover the method's data-rich transfer performance | Macro-F1 curves and reference-scarce-type recall |
| Open-set analysis | Novel-type AUROC | Separation of source-absent from source-represented query types | AUPR, FPR95 and forced-annotation performance |
| Supervised adaptation | Change in macro-F1 relative to the frozen representation with logistic regression | Effect of each model's implemented source-supervised training procedure | Per-method adapted and frozen macro-F1 |
| Physical interactions | AUROC and AUPRC | Ranking of STRING physical-interaction partners within the fixed gene universe | Degree-matched negative sampling |
| Functional sets | Mean average precision | Retrieval of genes sharing GO Biological Process, Reactome or KEGG annotations | Precision at 10, 50 and 100 |
| BEELINE regulatory edges | AUPRC divided by positive prevalence | Enrichment of known regulatory edges over the dataset-specific chance level | Early precision ratio |
| Perturbation-response genes | AUROC and AUPRC divided by positive prevalence | Retrieval of the 100 genes with the largest absolute response to each TF perturbation | Dataset-level means over 570 TF perturbations |
| Curated regulons | AUROC | Recovery of TF targets in TRRUST, ChEA and DoRothEA ABC | Per-TF distributions |
| Double perturbations | Normalized L2 error | Error in the predicted control-relative expression response of held-out double perturbations | Gene-wise $R^2$ for the CEBPA+CEBPE illustration |

**Table S16: Reproducibility of publicly released scFM execution routes.** Author-matched comparisons retain the released data, split and metric where available. Cross-study comparisons reproduce the stated external frozen- representation evaluation protocol using the scFoundry execution route.

| Evidence | Method | Data or protocol | Metric | Published or external | scFoundry | Source |
|---|---|---|---|---|---|---|
| **Author-matched supervised annotation** | | | | | | |
| Author-matched | scGPT | MS, 7,844 reference → 13,468 query cells | Accuracy / macro-F1 | 0.856 / 0.703 | 0.864 / 0.710 | Cui et al., Supplementary Table 2 [20] |
| Author-matched | scGPT | hPancreas, 10,600 reference → 4,218 query cells | Accuracy / macro-F1 | 0.968 / 0.718 | 0.978 / 0.672; shared-class macro-F1 0.873 | Cui et al., Supplementary Table 2 [20] |
| Author-matched | scGPT | Myeloid, 9,748 reference → 3,430 query cells | Accuracy / macro-F1 | 0.642 / 0.346 | 0.642 / 0.341 | Cui et al., Supplementary Table 2 [20] |
| Author-matched | CellPLM | MS, authors' split | Macro-F1 / macro-precision | 0.766 ± 0.007 / 0.803 ± 0.008 | 0.741 / 0.787 | Wen et al., Table 4 [21] |
| Author-matched | CellPLM | hPancreas, authors' split | Macro-F1 / macro-precision | 0.749 ± 0.010 / 0.753 ± 0.010 | 0.756 / 0.745 | Wen et al., Table 4 [21] |
| Author-matched | CellFM-80M | hPancreas, authors' split | Accuracy | 0.976 | 0.984 | Zeng et al., Supplementary Fig. S17 [22] |
| Author-matched | GenePT-w | scGPT MS query set; frozen weighted embeddings; 10-NN cosine | Accuracy / macro-F1 | 0.38 / 0.24 | 0.341 / 0.201 | Chen and Zou, Table C4 [23] |
| Author-matched | GenePT-w | scGPT hPancreas query set; frozen weighted embeddings; 10-NN cosine | Accuracy / macro-F1 | 0.95 / 0.66 | 0.947 / 0.631 | Chen and Zou, Table C4 [23] |
| Author-matched | GenePT-w | scGPT Myeloid query set; frozen weighted embeddings; 10-NN cosine | Accuracy / macro-F1 | 0.50 / 0.31 | 0.502 / 0.294 | Chen and Zou, Table C4 [23] |
| **Cross-study reproduction of published evaluation protocols** | | | | | | |
| External protocol | scBERT | Segerstolpe, scFoundation authors' split | Macro-F1 | 0.670 | 0.699 | Hao et al., Supplementary Table 4 [24] |
| External protocol | scBERT | Zheng68K, scFoundation authors' split | Accuracy / macro-F1 | — / 0.670 | 0.771 / 0.663 | Hao et al., Supplementary Table 4 [24] |
| External protocol | scGPT | scGPT MS query set; frozen embeddings; 10-NN cosine | Accuracy / macro-F1 | 0.76 / 0.61 | 0.735 / 0.583 | Chen and Zou, Table C4 [23] |
| External protocol | scGPT | scGPT hPancreas query set; frozen embeddings; 10-NN cosine | Accuracy / macro-F1 | 0.77 / 0.55 | 0.784 / 0.566 | Chen and Zou, Table C4 [23] |

*continued on next page*

*Table S16 continued*

| Evidence | Method | Data or protocol | Metric | Published or external | scFoundry | Source |
|---|---|---|---|---|---|---|
| External protocol | scGPT | scGPT Myeloid query set; frozen embeddings; 10-NN cosine | Accuracy / macro-F1 | 0.53 / 0.30 | 0.551 / 0.329 | Chen and Zou, Table C4 [23] |
| External protocol | scGPT | scIB pancreas 16k; `scib opt_louvain`; NMI-optimal resolution | AvgBIO | 0.44 | 0.446 | Kedzierska et al., Table S6 [25] |
| External protocol | UCE | Tabula Sapiens v2; published UCE evaluation framework | Bio / Batch / Total | 0.623 / 0.498 / 0.573 | 0.605 / 0.450 / 0.543 | Rosen et al., Supplementary Table 1 [26] |
| External protocol | Geneformer | Tabula Sapiens v2; published UCE evaluation framework | Bio / Batch / Total | 0.536 / 0.452 / 0.503 | 0.565 / 0.417 / 0.506 | Rosen et al., Supplementary Table 1 [26] |
| External protocol | scGPT | Tabula Sapiens v2; published UCE evaluation framework | Bio / Batch / Total | 0.436 / 0.374 / 0.411 | 0.588 / 0.428 / 0.524 | Rosen et al., Supplementary Table 1 [26] |
| External protocol | scPRINT-medium | scIB pancreas; scPRINT Benchmarker, `min_max_scale=False` | Total | 0.44 | 0.458 | Kalfon et al., Source Data Fig. 4C [27] |
| External protocol | scPRINT-medium | scIB lung; same protocol | Total | 0.51 | 0.569 | Kalfon et al., Source Data Fig. 4C [27] |
| External protocol | scGPT | scIB pancreas; scPRINT protocol | Total | 0.37 | 0.464 | Kalfon et al., Source Data Fig. 4C [27] |
| External protocol | scGPT | scIB lung; scPRINT protocol | Total | 0.36 | 0.573 | Kalfon et al., Source Data Fig. 4C [27] |

[2] CZI Single-Cell Biology Program, Shibla Abdulla, Brian Aevermann, Pedro Assis, Seve Badajoz, Sidney M. Bell, Emanuele Bezzi, Batuhan Cakir, Jim Chaffer, Signe Chambers, J. Michael Cherry, Tiffany Chi, Jennifer Chien, Leah Dorman, Pablo Garcia-Nieto, Nayib Gloria, Mim Hastie, Daniel Hegeman, Jason Hilton, Timmy Huang, Amanda Infeld, Ana-Maria Istrate, Ivana Jelic, Kuni Katsuya, Yang Joon Kim, Karen Liang, Mike Lin, Maximilian Lombardo, Bailey Marshall, Bruce Martin, Fran McDade, Colin Megill, Nikhil Patel, Alexander Predeus, Brian Raymor, Behnam Robatmili, Dave Rogers, Erica Rutherford, Dana Sadgat, Andrew Shin, Corinn Small, Trent Smith, Prathap Sridharan, Alexander Tarashansky, Norbert Tavares, Harley Thomas, Andrew Tolopko, Meghan Urisko, Joyce Yan, Garabet Yeretssian, Jennifer Zamanian, Arathi Mani, Jonah Cool, and Ambrose Carr. CZ CELL×GENE Discover: A single-cell data platform for scalable exploration, analysis and modeling of aggregated data, November 2023.
